# Epilipidomics platform for holistic profiling of oxidized complex lipids in blood plasma of obese individuals

**DOI:** 10.1101/2021.12.23.473968

**Authors:** Angela Criscuolo, Palina Nepachalovich, Diego Fernando Garcia-del Rio, Mike Lange, Zhixu Ni, Matthias Blüher, Maria Fedorova

**Author notes:** **Corresponding author:** Dr. Maria Fedorova, Lipid metabolism: analysis and integration, Center of Membrane Biochemistry and Lipid Metabolism, University Hospital and Faculty of Medicine Carl Gustav Carus of TU Dresden, Germany;. Lipid metabolism: analysis and integration, Center for Membrane Biochemistry and Lipid Metabolism, University Hospital and Faculty of Medicine Carl Gustav Carus of TU Dresden, Germany. Authors equally contributed to the work (Angela Criscuolo, Palina Nepachalovich).

## Abstract

Lipids are a structurally diverse class of biomolecules which can undergo a variety of chemical modifications. Among them, lipid (per)oxidation attracts most of the attention due to its significance in regulation of inflammation, cell proliferation and death programs. Despite their apparent regulatory significance, the molecular repertoire of oxidized lipids remains largely elusive as accurate annotation of lipid modifications is challenged by their low abundance and largely unknown, biological context-dependent structural diversity. Here we provide a holistic workflow based on the combination of bioinformatics and LC-MS/MS technologies to support identification and relative quantification of oxidized complex lipids in a modification type- and position-specific manner. The developed methodology was used to identify epilipidomics signatures of lean and obese individuals with and without type II diabetes. Characteristic signature of lipid modifications in lean individuals, dominated by the presence of modified octadecanoid acyl chains in phospho- and neutral lipids, was drastically shifted towards lipid peroxidation-driven accumulation of oxidized eicosanoids, suggesting significant alteration of endocrine signalling by oxidized lipids in metabolic disorders.

## Introduction

Lipids are extremely diverse class of small biomolecules, for which the full natural variety is yet not fully defined. So far more than 46 000 of lipid molecular species are annotated (LIPID MAPS; November 2021), and over hundreds of thousands being computationally predicted. In addition to the high diversity of original species derived from classical biosynthesis routes, lipids can undergo various modifications by the introduction of small chemical groups via enzymatic and non-enzymatic reactions, forming higher levels of structural complexity, termed “epilipidome”^1^. Modifications of lipids including oxidation, nitration, and halogenation are described for numerous physiological and pathological conditions and are generally linked to the regulation of various signalling events^2^. Thus, epilipidome, in a manner similar to epigenetic and post-translational protein modifications, increases the regulatory capacity of biological systems by supporting prompt responses to different stimuli and stressors.

Among different modifications, lipid (per)oxidation was so far studied the most. Oxylipins, oxygenated derivatives of polyunsaturated fatty acids (PUFA), are commonly analysed as markers of systemic and tissue specific inflammation and its resolution^3^. Although usually measured in the form of free fatty acids, the majority of oxylipins *in vivo* are believed to be esterified to complex lipids, including phospholipids (PLs), cholesteryl esters (CEs) and triglycerides (TGs)^4^. Positive correlation between endogenous levels of oxidized complex lipids and the development and progression of numerous human diseases including cardiovascular (CVD), pulmonary and neurological disorders as well as non-alcoholic steatohepatitis (NASH) was demonstrated^5–8^. Oxidized lipids have been shown to accumulate in apoptotic cells as well as microsomes released by activated or dying cells, and are generally associated with inflammatory responses, mediated mainly via the innate immune system^2^. Inflammation modulating potential of oxidized lipids is attributed to their recognition by and thus activation of a wide range of pattern recognition receptors including macrophage scavenger receptors, Toll-like receptors, CD36, and C reactive protein^9^. Moreover, oxidized lipids and their protein adducts can be recognized by natural IgM antibodies and thus sequestered from the circulation^5^. Furthermore, recently formulated concept of ferroptosis placed oxidized membrane phospholipids (PL) in the center of the cell death execution program via necrotic rupture of cellular plasma membranes^10^.

Despite overwhelming amount of data on their biological significance, the pathways leading to the intracellular and extracellular generation of complex oxidized lipids and thus their *in vivo* structural diversity remain largely elusive. Both enzymatic (via action of lipoxygenase, and cytochrome P450 systems^11,12^) and non-enzymatic (via free radical driven lipid peroxidation^13,14^) systems are proposed to contribute to the accumulation of oxidized lipids. Overall, an increasing body of evidence allows to propose that action of oxidized complex lipids is context dependent and their bioactivities might be manifested either in adaptive or pathological manner^9,11^. Understanding the underlying regularities in epilipidomics patterns would require holistic mapping of oxidized lipids in different conditions in a variety of biological matrices. Liquid chromatography coupled on-line to mass spectrometry (LC-MS) was actively applied for the identification and quantification of oxidized complex lipids with a main focus on oxidized phosphatidylcholines (oxPC) and oxCE. With just a few reports addressing the high diversity of oxidized PLs^15–17^, the vast majority of the available LC-MS-derived data report only a handful of oxidized species analysed by targeted LC-MS workflows.

Indeed, due to their low natural abundances, analysis of oxidized lipids in complex biological matrices mostly relies on the targeted LC-MS utilizing a limited list of previously identified oxidized species independent of the biological context preventing the discovery of sample-specific epilipidome.

Here we describe the LC-MS/MS workflow developed for the holistic, biological context-specific epilipidome profiling and relative quantification. The method combines biological intelligence-driven *in silico* prediction of a sample-specific epilipidome followed by its semi-targeted detection using a set of optimized LC, MS and MS/MS parameters aiming to increase the accuracy of the annotation. The developed workflow, validated using blood plasma samples of lean, non-diabetic and type II diabetic obese individuals, allowed the identification and relative quantification of oxidized PC, CE and TG lipids, showing both physiological (lean) and pathological (obese) patterns, thus supporting endocrine signalling role of the oxidized epilipidome.

## Results and Discussion

### Fragmentation rules for annotation of oxidized complex lipids

Mass spectrometry-based annotation of oxidized lipids requires accurate detection of the precursor ion *m/z* (MS1) from which lipid elemental composition can be deduced, as well as an informative fragment mass spectrum (MS2) allowing the assignment of lipid class, molecular species, modification type, and modification position. For instance, oxPLs ionized in negative ion mode upon collision-induced dissociation (CID) produce intense fragment ions specific to the head groups, and fatty acyl chains, including the one carrying the modification. Studies on oxygenated free fatty acids (oxylipins) demonstrated that the anion of oxygenated fatty acid upon CID undergoes charge-remote fragmentation resulting in a set of fragments characteristic to the modification type and position along the hydrocarbon chain^18–21^. Recently, to induce similar fragmentation patterns for oxidized complex PLs, multistage fragmentation techniques (MS3) on tribrid MS instruments were applied for the annotation of oxidized PC and PE lipids in complex biological samples^22^ (Figure 1A). However, the multistage fragmentation reduces sensitivity (when MS3 spectra are recorded in the orbitrap) and/or resolution (when the spectra are recorded in the ion trap). Here, using a set of *in vitro* oxidized PC lipid standards, we demonstrate that MS2 spectra obtained using elevated energy HCD display similar set of fragment ions without the need of multistage ion activation (Figure 1B). Thus, HCD of oxPC formate adduct ion at *m/z* 818.5545 obtained using stepped normalized collision energy of 20-30-40 units, provided lipid class (head group-specific ions at *m/z* 758.5352, 168.0431 and 224.0695), molecular species (anions of fatty acyl chains at *m/z* 255.2332 and 295.2281), modification type (water loss characteristic to the presence of hydroxyl group at *m/z* 277.2175), and modification position (*m/z* 195.1392, the product of cleavage of the adjacent to the carbinol C-C bond) specific fragment ions allowing to annotate this oxidized lipid as PC(16:0_18:2<OH{13}>). Similarly, each type of modification and its positional isomers in oxPLs can be characterized by a set of specific fragment ions obtained either in MS3 or elevated energy HCD MS2 experiments.

**Figure 1.**
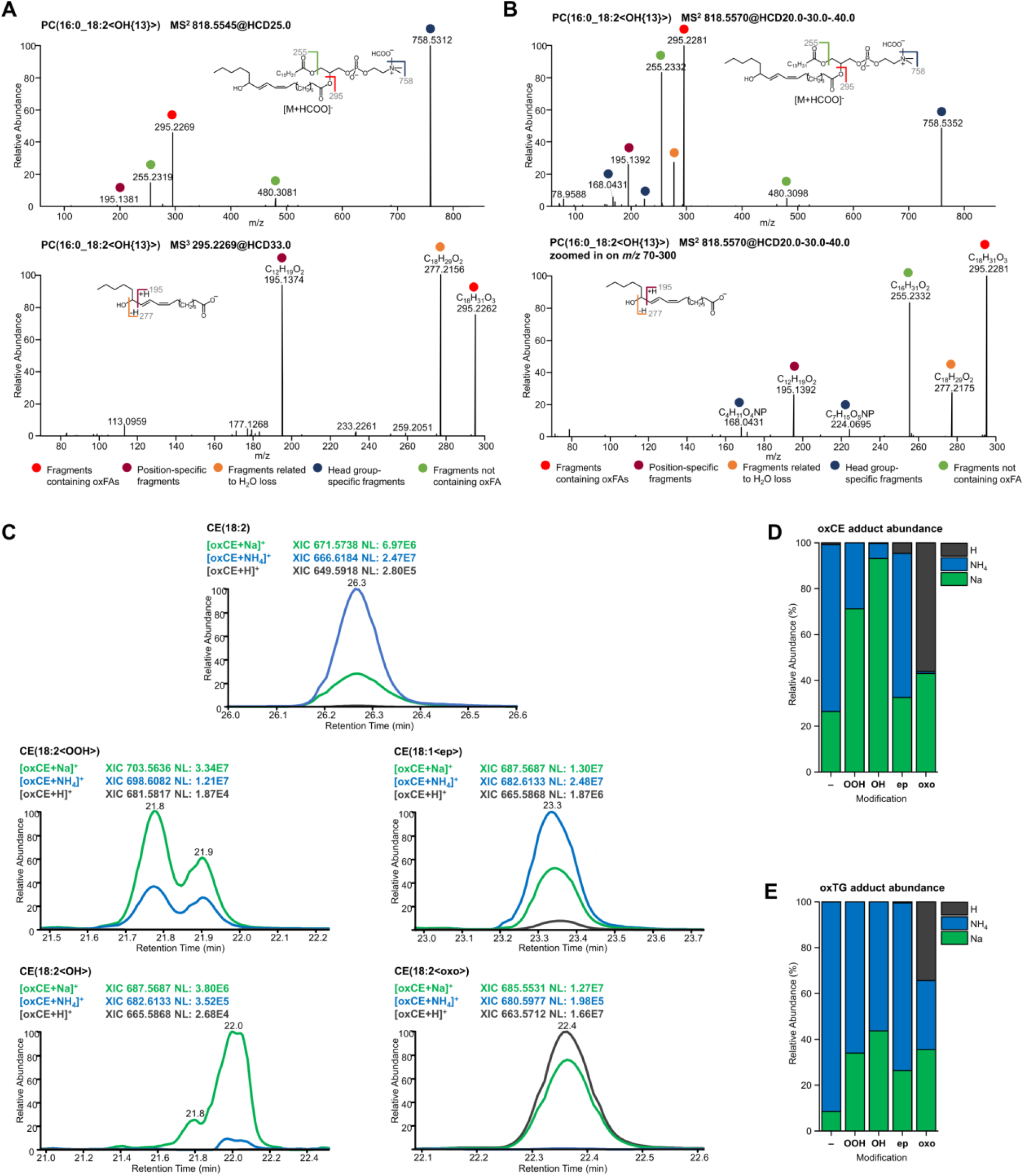
Optimization of MS and MS2 parameters for the structural elucidation of oxidized complex lipids structures. Tandem mass spectrometry analysis of oxidized PC lipid using multistage MS3 (**A**) or optimized elevated energy stepped HCD MS2 experiments (**B**) both allowed structural assignment of PC(16:0_18:2<OH{13}>). Preferential positive ion mode ionization adducts were evaluated for *in vitro* oxidized neutral lipids and showed modification type specific preferences. Qualitative (**C**; illustrated using extracted ion chromatograms for oxCE(18:2) species) and quantitative (**D, E**) distribution of modification specific preferences to form protonated (dark grey), ammoniated (blue) or sodiated (green) adducts for unmodified (-), hydroperoxy-(OOH), hydroxy (OH), epoxy (ep), or keto (oxo) derivatives of CE and TG. Relative abundances of differential adducts for oxCE (D) and oxTG (E) represent average values calculated for 10 species.

Unfortunately, chemically defined standards of complex oxidized lipids necessary to create a fragment spectra library, are limited to just a few, commercially available molecular species and do not reflect the diversity of possible endogenous analytes. To facilitate further high-throughput annotation of oxPLs, we compiled our results obtained by MS3 and elevated energy HCD MS2 fragmentation of *in vitro* oxidized PC lipids and selected oxylipin standards (Tables S1.1 and S1.2), literature data on the fragmentation of oxidized free fatty acids and complex lipids^17,19,23–31^, as well as available MS2 spectra from METLIN, LIPID MAPS, and MS DIAL.msp library in a form of fragmentation rules exemplified here for different modification types and positions on oxidized oleoyl (18:1), linoleoyl (18:2), and arachidonoyl (20:4) chains in PC lipids (File S1).

To extend these rules beyond oxPL lipids, we performed *in vitro* oxidation of neutral lipids, namely CE and TG. Neutral lipids do not ionize in negative ion mode, and are usually monitored as positively charged ammonium adducts. However, when *in vitro* prepared standards were analysed by LC-MS/MS, we noticed that upon oxidation both CE and TG have higher tendency to form sodiated, and in some case protonated, adducts (Figure 1C). Moreover, adduct preferences were modification type specific (Figure 1D and E). Thus, unmodified CEs were preferentially ionized as NH_4_ adducts (73% of total abundance) with minor contribution of sodiated forms (26%), whereas CE hydroperoxides showed clear preference towards Na adduct formation (71%). The preference for Na adduct was even more evident for hydroxylated CE derivatives with 93% detected in the form of sodiated ions. Majority of epoxyCE, however, remained in ammoniated forms (63%), and oxoCE derivatives showed preferences towards protonated (56%) adducts (Figure 1D). Oxidized TG displayed overall similar modification type specific adducts distribution, however ammoniated adducts remained dominant for almost all modification types (Figure 1E).

Considering the trend of oxCE and oxTG to form differential adducts upon ESI, we further compared the MS2 spectra of *in vitro* oxidized standards obtained from different precursor ions in terms of their utility towards accurate oxidized lipids annotations. Protonated species provided non-informative MS2 (data not shown), whereas both Na and NH_4_ adducts showed some structure related fragments (Figure 2A). Thus, the HCD spectrum of CE(18:2<OOH>) ammonium adduct displayed single meaningful fragment ion corresponding to the protonated form of cholestene, whereas the sodiated precursor resulted in much more informative MS2 spectrum with modification type- and position-specific neutral loss and fragment ions. Indeed, alkali metal adducts of oxidized lipids and oxylipins were previously reported to follow the similar charge-remote fragmentation mechanism described above for oxidized fatty acyl chain anions^18–20^. Here, we further optimized HCD collision energy to obtain informative MS2 patterns allowing to identify both modification type and position specific fragment ions of sodiated oxCE and oxTG. Overall, Na adducts were more stable than their ammoniated counterparts and required higher collision energies for their efficient fragmentation (Figure 2B). Stepped elevated energy HCD of 30-40-50 normalized units was chosen for further experiments and allowed to define fragmentation rules for hydroperoxy derivatives of CE and TG lipids (Figure 2C and D) as well as other types of modifications (Table S1.3 and Table S1.4). Similar to oxPC, we compiled MS2 data on sodiated precursors from *in vitro* oxidized CE/TG as well as available literature data in a form of modification and position specific fragmentation rules to be used for high-throughput annotation of modified neutral lipids (File S1).

**Figure 2.**
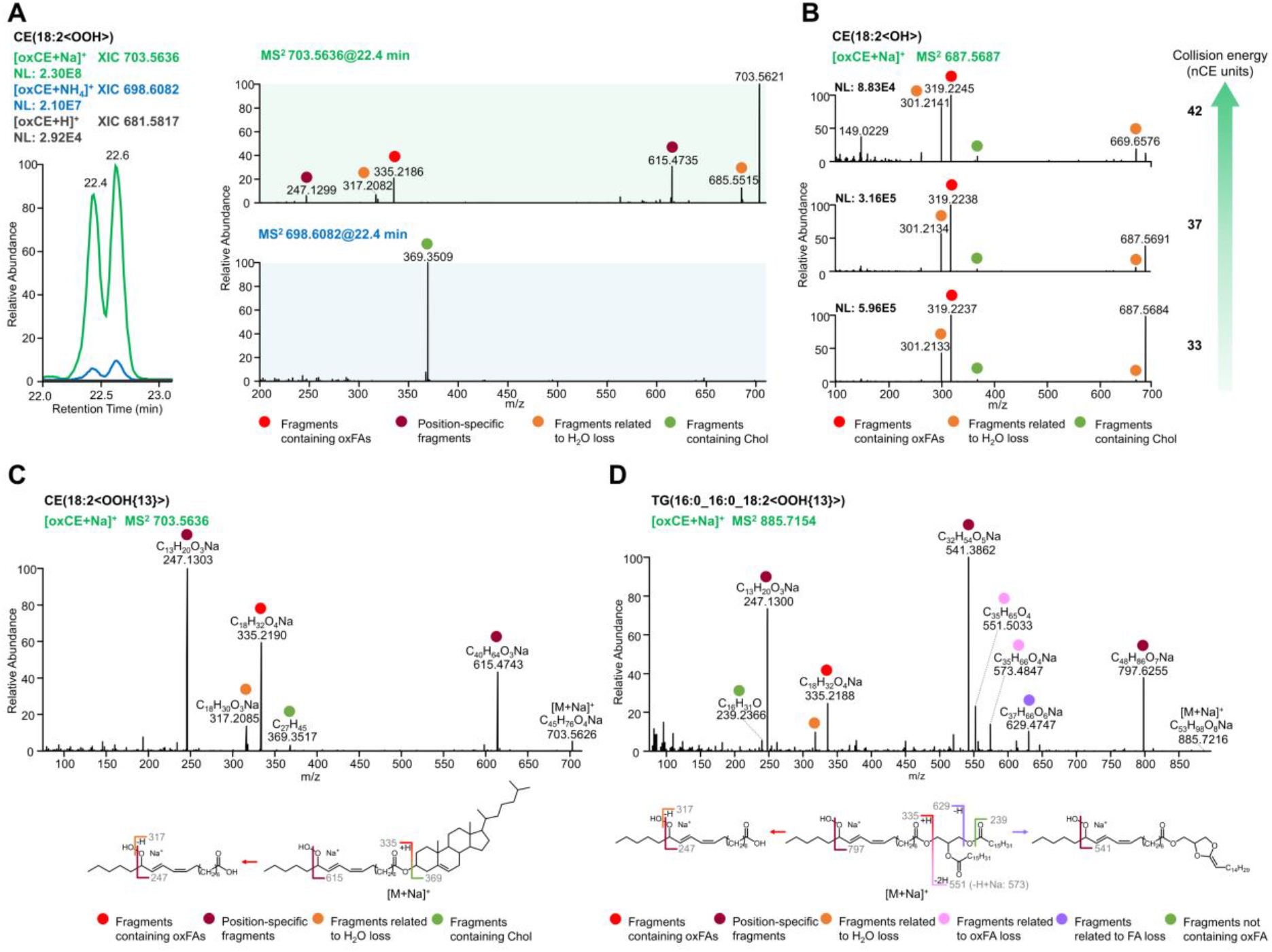
Optimization of elevated energy HCD of sodiated precursors to support structural elucidation of oxidized neutral lipids. Utility of tandem mass spectra to provide structure-relevant fragment ions was compared for ammoniated (blue) and sodiated (green) oxCE precursor ions (**A**). Considering larger number of structure-relevant fragment ions obtained by HCD of sodiated oxCE precursors, the collision energy was optimized to enhance fragment ions formation (**B**). Elevated energy stepped HCD MS2 experiments using sodiated precursors were shown to provide highly informative spectra, exemplified here for CE(18:2<OH{13}) (**C**) and TG(16:0_16:0_18:2<OOH{13}>) (**D**).

To validate ionization preferences and fragmentation rules in complex biological matrices, we performed *in vitro* oxidation of commercially available human blood plasma and were able to confirm the presence of the described adduct types and to annotate selected oxidized species using a set of fragmentation rules provided in File S1 (Table S2). Taken together, using *in vitro* oxidized lipid standards and human blood plasma, we defined the preferential ionization adducts to be utilized for MS2 experiments aiming for the identification of oxidized complex lipids. The optimized elevated energy HCD was shown to be efficient in providing informative MS2 spectra of oxidized lipids. Charge-remote fragmentation of acyl anions or their sodiated analogues in oxPC and oxCE/oxTG respectively, allowed efficient assignment of hydroxy-, epoxy-, keto-, and hydroperoxy-modified lipids as well as oxidatively truncated forms in a modification type and position specific manner. The fragmentation rules defined for each modification type and position can be further used to support high-throughput annotation of lipid modifications.

### Increasing annotation accuracy by retention time mapping of isomeric oxidized lipids

Lipid oxidation might lead to the formation of a large number of isomeric species including isomers carrying different types of modifications as well as positional isomers. Although unresolvable by MS, isomeric oxidized lipids can be separated by reversed-phase chromatography (RPC)^32^. Here, to characterize *in vitro* oxidized standards and blood plasma we applied C18 RPC coupled on-line to MS. LC-MS coupling allowed to separate multiple isomeric lipids providing clean MS2 spectra for the annotation. Moreover, we were able to define the elution order for mono- and dioxygenated isomeric species (Figure 3A). The elution order was consistent within different lipid classes (PC, CE, TG, and oxylipin standards) and was identical to the one described for isomeric oxylipins in the literature^33^. Thus, for RPC-separated monooxygenated lipids the elution order was *OH < oxo < epoxy*, and for dioxygenated it was *2OH < OH, O < ep, O < OOH*. Dioxygenated lipids eluted earlier than their monooxygenated counterparts. For regioisomers the general trend was defined as well with earlier elution times for the species carrying the modification closer to the ω-end of the acyl chains (e.g. *C13-OOH < C9-OOH*). Furthermore, previously introduced retention time (RT) mapping of oxidized lipids using Kendrick mass defect (KMD) plots^34^ was employed here for visualization and manual inspection of the defined elution order to increase accuracy of the annotations in high-throughput epilipidome profiling experiments (Figure 3B).

**Figure 3.**
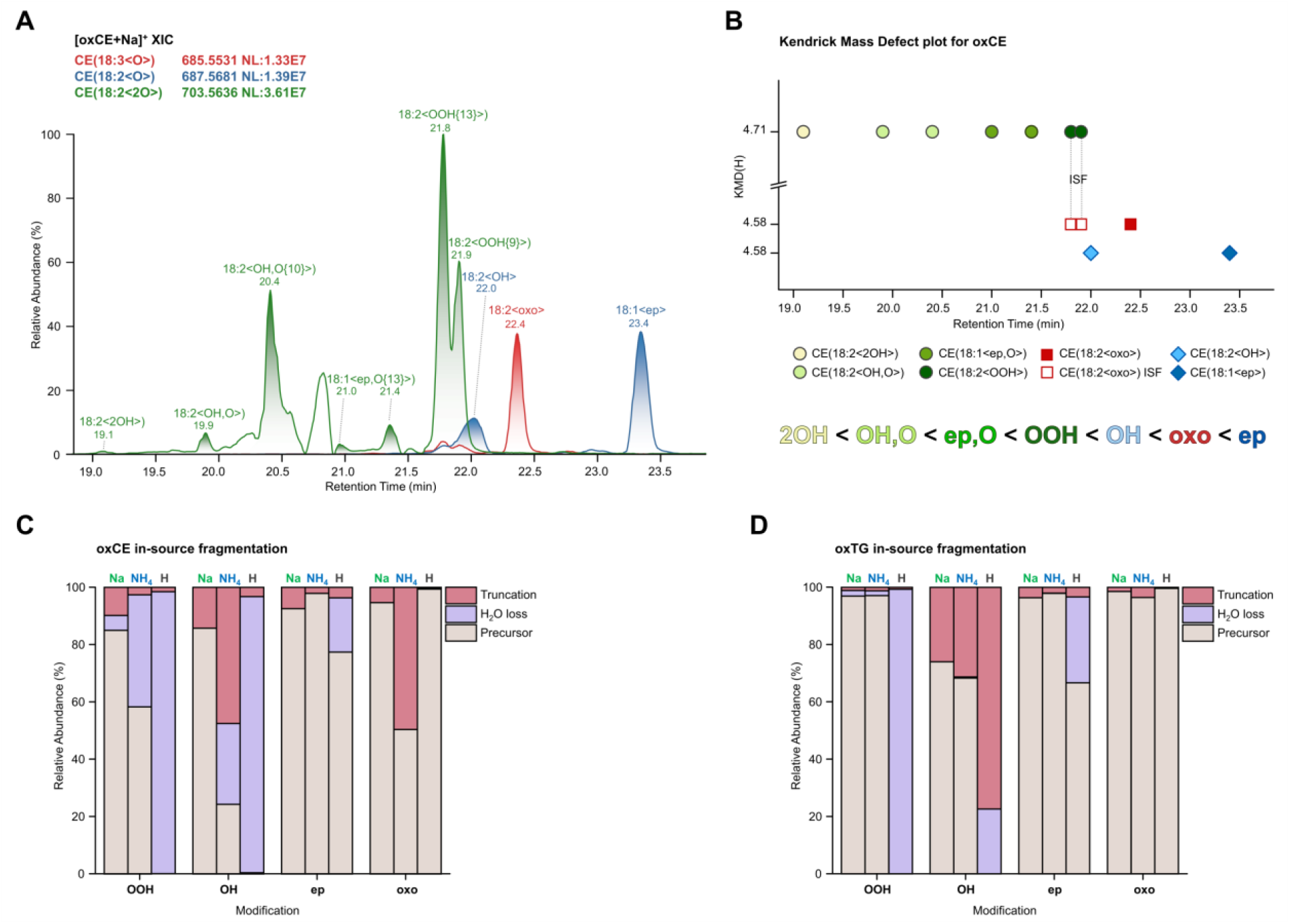
LC-MS/MS coupling and retention time mapping facilitate resolution of isomeric oxidized lipids and allow to correct for in-source fragmentation (ISF). Isomeric oxidized lipids can be successfully resolved by RPC, as illustrated here for mono-(CE(18:3<O>) in red and CE(18:2<O>) in blue) and dioxygenated (CE(18:2<2O>) oxCE lipids (A). Retention time mapping using KMD(H) plots allows visual inspection of isomer-specific elution order as well as identification of false positive annotation due to the ISF (B). The extent of ISF for oxCE (C) and oxTG (D) lipids was shown to be ionization and modification type specific. ISF was calculated based on the parent and product peak areas using the formula 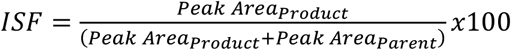. Relative abundances of ISF derived species (truncation and water loss) for oxCE (C) and oxTG (D) represent average values calculated for 10 species.

Importantly, using KMD vs RT plots we were able to spot previously unconsidered in-source fragmentation (ISF) of oxidized lipids. Despite the application of soft ionization techniques such as electrospray ionization, lipid ions remain quite labile and can undergo undesired fragmentation via collision with neutral gas molecules in the intermediate region of MS instruments between the atmospheric pressure ion source and deep vacuum environment of the mass analyzer. Those fragment ions often are identical to the endogenous lipids of other subclasses and thus ISF is a known source of false positive identifications in conventional lipidomics^35,36^. Similarly, we demonstrated that oxidized lipids can undergo substantial ISF with the formation of ions related to other types of modifications. For instance, CE hydroperoxides undergo ISF accompanied by the loss of water with the formation of the ion with *m/z* equal to the endogenous oxoCE derivative (Figure 3A, red trace). As ISF occurs after LC separation, it can be recognized and thus corrected by RT mapping in a form of KMD vs RT plots (Figure 3B, open red squares).

Using this approach, we systematically evaluated the extent of ISF for different type of adducts and modifications for oxPC, oxCE, and oxTG. At the settings for the ion transfer optics used in this experiments, oxPC did not display any significant ISF, whereas both oxCE and oxTG showed adduct- and modification type-specific ISF. Among the most significant fragmentation events driven by the presence of the modification were the loss of water and gas-phase acyl chain cleavage (truncation) at the site adjacent to the modification. Protonated precursors underwent the most significant ISF for all studied modifications except oxo-derivatives (Figure 3C and D), which probably explains the observed adduct abundance described above for this modification type (Figure 1C and D). Similarly, in line with the observation that sodiated adducts required higher collision energies to obtain fragment-rich MS2, Na adducts of both oxCE and oxTG were more resistant to ISF (Figure 3C and D), whereas ammoniated adducts showed ISF up to 75%. Overall sodiated adducts showed highest analytical robustness and thus selectivity of oxidized lipid profiling, although they are not necessarily represented the most intense species. Among the modification types, hydroxy- and hydroperoxy-modified CE and TG were the most labile whereas epoxy- and keto-derivatives generally showed high stability. Thus, we demonstrated the efficiency of RT mapping to increase the accuracy of oxidized lipid annotations both in terms of isomer resolution and the correction for ISF. Taken together, we provided multi-level annotation guidelines for oxPC, oxCE, and oxTG lipids based on the selection of the preferential ionization adducts showing minimized ISF (*MS1 level*) and the most informative MS2 patterns, modification type- and position-specific fragmentation rules (*MS2 level*), as well as RT mapping (*LC level*) to be used for the accurate and high-throughput identification of oxidized complex lipids from LC-MS/MS datasets.

### Semi-targeted LC-MS/MS for context dependent high-throughput epilipidome profiling

Multi-level (LC-MS1-MS2) annotation guidelines developed above could support identification of oxidized complex lipids in biological matrices but only when high quality MS2 spectra are made available. Application of targeted MS methods limits epilipidome coverage to a predefined subset of oxidized species which not necessarily reflect the biological context of the sample in question. On the other hand, application of untargeted data-dependent (DDA) method showed to be generally inefficient for detection of oxidized lipids as their low endogenous abundance prevents them to be selected for MS2 events in the presence of highly abundant unmodified lipids. To find a compromise between targeted solution limiting epilipidome coverage and “undersampling” of low abundant oxidized lipids by untargeted DDA, we developed a semi-targeted LC-MS/MS method. The method relies on the combination of biological context dependent *in silico* prediction of a sample specific epilipidome which is used as an inclusion list for semi-targeted DDA (stDDA) considering MS1 (preferential adduct selection) and MS2 (lipid class specific elevated energy HCD) settings optimized as described above. For the method development and validation we used lipid extracts from blood plasma of lean non-diabetic (LND), obese non-diabetic (OND) and obese with type II diabetes (OT2D) individuals for whom endogenous oxidized complex lipids with largely unknown diversity are expected.

Considering that native lipidome serves as the substrate for lipid (per)oxidation, epilipidome of a particular tissue can be predicted from its native lipidome. In blood, lipids are transported mainly in the form of lipoproteins with PC, CE, and TG representing the main lipid classes. We and others previously reported the lipid composition of human blood plasma^37,38^, from which here we selected 45 most abundant PUFA-containing PC, CE and TG lipid molecular species to be used as sample-specific endogenous substrates for *in silico* oxidation. LPPtiger software was employed to predict the sample-specific epilipidome using 17 PC, 17 TG, and 11 CE lipid molecular species as the substrates. Rather than performing simple enumeration with oxygen atoms, the *in silico* oxidation algorithm within LPPtiger relies on molecular networks manually reconstructed for 10 PUFAs based on the literature data available on their enzymatic and non-enzymatic oxidation products^39^. Moreover, a user can specify the level of *in silico* oxidation which was here constrained to the maximum of two oxidation sites with up to 2 hydroxy, 2 epoxy, 1 oxo, or 1 hydroperoxy modifications in long-chain and truncated oxidized lipids, thus limiting the predicted chemical space to the most probable lipid modifications. Using 45 native lipids as *in silico* oxidation substrates, LPPtiger predicted 956 oxidized species corresponding to 559 unique elemental compositions. Those were used to compose inclusion lists for stDDA LC-MS/MS analysis considering their preferential ionization mode and adducts defined above for each lipids class. Thus, oxPCs were analysed in negative ion mode as formate adducts, and oxCEs/oxTGs as sodium adducts in positive ion mode using elevated energy HCD levels specific for each lipid class. Considering efficient RPC separation of oxidized lipid classes, we further used polarity switching with the 1^st^ half of the LC gradient measured in negative (oxPC) and the 2^nd^ in positive (oxCE and oxTG) ion modes to facilitate higher throughput of the analysis. Using this setup 559 unique *m/z* for predicted oxidized lipids were targeted within two LC-MS/MS DDA analysis per group specific plasma pool (e.g. LND, OND and OT2D) using a sample amount equivalent to as low as 1.3 μL of blood plasma per sample.

The developed semi-targeted method (Figure 4A) provided several important advantages including large coverage of the epilipidome predicted in a sample-specific manner and thus addressing the biological context-driven profiling of modified lipid species (Figure 4B). The use of stDDA instead of multiple reaction monitoring (MRM) or parallel reaction monitoring (PRM) approach allowed to profile a larger number of analytes without repetitive sample injections. Indeed, within the large inclusion list of possible targets, only those above certain intensity threshold and abundance rank (here top 6 mode was used) were selected for MS2 ensuring sufficient quality of MS2 spectra, without wasting instrument cycle time on extremely low intense or even absent analytes, fragmentation of which would not result in senseful MS2 spectra anyway. The use of relatively low number of precursors for MS2 selection (top 6) within each DDA cycle further allowed to increase injection time (200 ms) to accumulate larger number of ions for the fragmentation without compromising the resolution provided by RPC. Elevated energy HCD provided informative MS2 spectra without the need of multistage ion activation (MS3) thus increasing the sensitivity and resolution necessary for the accurate annotation of isobaric modification type- and position-specific fragments. Moreover, RT-scheduled polarity switching allowed to record MS2 spectra for both oxPC and oxCE/oxTG within the same LC-MS/MS analysis. Taken together, the developed stDDA method allowed profiling of oxidized complex lipids in a holistic, biological context-dependent, and sensitive way.

**Figure 4.**
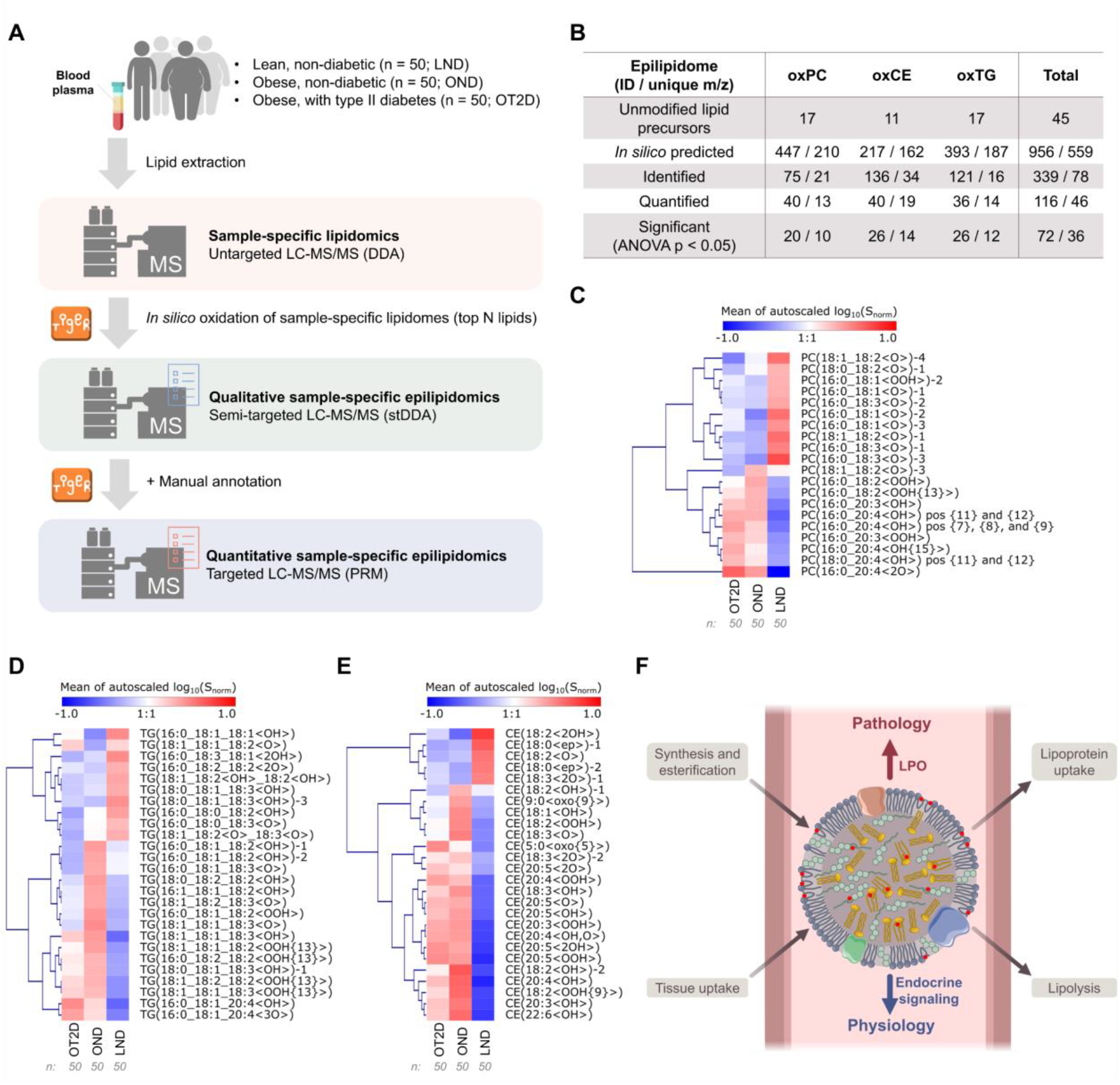
Semi-targeted identification and targeted quantification of oxidized complex lipids in blood plasma lipid extracts of lean non-diabetic (LND), obese non-diabetic (OND) and obese with type II diabetes (OT2D) individuals. Dedicated workflow (**A**) based on *in silico* prediction of sample specific epilipidome, its identification using stDDA followed by PRM based targeted quantification allowed to annotate and quantify oxidized lipids significantly different between their phenotypes (**B**). Heatmaps illustrate lean (LND), obese (OND) and T2D obese (OT2D) specific epilipidomics signatures for oxPC (**C**), oxTG (**D**) and oxCE (**E**) lipids with significantly different levels (ANOVA < 0.05). Based on the identified phenotype-specific epilipidomes, the endocrine functionality was proposed for complex lipids esterified octadecanoids, which was altered upon obesity and type II diabetes via enrichment of lipid peroxidation (LPO) derived eicosanoids (**F**).

### Annotation and relative quantification of oxidized complex lipids in blood plasma of lean and obese individuals

Using the lipidome-specific prediction and stDDA method described above, we analysed group-specific pooled samples (OND vs LND vs OT2D) in which 339 oxidized complex lipids corresponding to 78 unique *m/z* were identified (Figure 4A and B; Table S3, Files S2-4). The initial identification of modified lipids was performed by LPPtiger, followed by the manual annotation based on the fragmentation rules to define modification type- and position-specific species (File S1). Large number of isomeric oxidized lipids were annotated and confirmed using elution rules and RT mapping via KMD(H) plots (an example for oxCE is presented in Figure S1).

Having identified oxidized complex lipids in group-specific pooled samples, we further performed their relative quantification in individual blood plasma lipid extracts of LND (n=50), OND (n=50) and OT2D (n=50) individuals. To this end, the targeted PRM method was employed utilizing retention time scheduling and polarity switching. Out of 339 identified oxidized complex lipids, 116 species (46 unique *m/z*) were quantified (Figure 4A and B; Table S4).

Out of 116 quantified oxidized lipids, 20 oxPC, 26 oxTG and 26 oxCE species showed significant differences between three groups of samples (ANOVA p < 0.05) (Figure 4 C-E, Table S5). Interestingly, out of 72 oxidized lipids, 25 showed higher levels in LND relative to obese (OND and OT2D) individuals. Importantly, those lipids were specifically represented by octadecanoids, a family of oxidized lipids bearing 18 carbon long fatty acyl chains. Thus, oxidized complex lipids enriched in the blood plasma of LND individuals exclusively carried 18:1, 18:2 and 18:3 modified acyl chains. Out of 10 oxPC species enriched in LND vs obese individuals, 9 corresponded to the monooxygenated octadecanoids (Figure 4C). Unfortunately, due to the extremely low abundance of those lipids, the confident structural assignment of the modification type and position was not possible. However, within oxTGs with higher abundance in LND vs obese individuals, the majority was represented by hydroxy-modified octadecanoids (Figure 4D), indicating that lipid class independent octadecanoid accumulation is a hallmark of healthy human plasma. It was previously established that de-esterified octadecanoid alcohols are enriched in triglyceride rich lipoproteins, especially VLDL^4^, and it is worthwhile to speculate that their TG esterified forms enriched in LND samples represent a potential storage and/or transport mechanisms. On the other hand, oxCE elevated in LND individuals were represented by epoxy- and diol-modified octadecanoids, indicating enrichment of CYP450- and soluble epoxide hydrolase pathway-derived metabolites (Figure 4E). Indeed, the role of octadecanoids as physiologically relevant signalling molecules emerged in several recent reports. Thus, elevated levels of linoleic- and α-linolenic acid-derived oxylipins were shown to be negatively associated with the development of type 1 diabetes^40^, cardiovascular events^41^, and acutely decompensated cirrhosis^42^. Esterified diols of linoleic acid in triglyceride-rich lipoproteins formed a characteristic signature promoting anti-atherogenic endothelial phenotype^43^. However, in all those studies only oxylipins (free and released from complex lipids by basic hydrolysis) were measured. Here we demonstrated that those oxidized acyl chains are present within complex lipids of different classes with a certain specificity of modification patterns (e.g. alcohols for oxTG and epoxides/diols for oxCE) and enriched in blood plasma of LND vs obese individuals. This finding provides direct evidence for the previously proposed endocrine signalling role of oxylipins usually referred only as paracrine and autocrine molecules^4^. Indeed, complex oxidized lipids incorporated in blood lipoproteins can be delivered to or collected from different organs and tissue to support their controlled release or clearance via the coordinated action of lipoprotein lipases and acyltransferases, respectively.

The pattern of oxidized complex lipids was remarkably altered in obese individuals, with overall enrichment of non-enzymatic LPO products evident for all analysed lipid classes. Thus, in the blood plasma of obese individuals we quantified elevated levels of hydroperoxides as well as a variety of hydroxy-derivates. In this case, the majority of oxPCs contained modified eicosatetraenoic and eicosatrienoic acyl chains. Multiple positional isomers carrying hydroxyeicosatetraenoic (HETE) acyl chains including 7-, 8-, 9-, 11-, 12-, and 15-OH were enriched in the samples from obese individuals, indicating the free-radical origin of these oxPCs (Figure 4C). Overall, the levels of LPO-derived oxPCs were higher in OT2D individuals with the prevalence of C20:x modified acyl chains, whereas in OND samples, C18:x oxidized oxPCs were detected as well. This pattern discriminating between OND vs OT2D individuals was even more evident for oxTG (Figure 4D), with C18:x hydroxy lipids enriched in OND, and the majority of LPO-derived hydroperoxides (both C18:x and C20:x origin) elevated in both OND and OT2D samples. The difference between OND and OT2D individuals in terms of oxCE followed the same pattern, with C18:x-derived oxCEs enriched specifically in OND, and the general LPO products derived mostly from C20:x acyl chains characteristic for both obese phenotypes (Figure 4E). Taken together, we could demonstrate that complex lipids within human blood carry a large variety of oxidized acyl chains supporting their endocrine functionality (Figure 4F). Moreover, LND individuals have a distinct epilipidomic signature enriched in oxygenated octadecanoid acyl chains with modification type specificity between different lipid classes. This physiological signature is disrupted by the obesity-induced metabolic distress with the formation of various products, mostly of non-enzymatic origin. Both OND and OT2D individuals were characterized by this LPO-driven epilipidomics shift, however in OND a number of C18:x modified species, different from the ones present in lean individuals, was enriched indicating possible attempts towards systemic adaptation.

### Conclusion

Here we presented a holistic bioinformatics- and LC-MS/MS-based workflow for the accurate identification and relative quantification of oxidized complex lipids. Using *in vitro* generated standards we optimized MS (selection of preferential ionization adducts) and MS2 (lipid class-specific elevated energy HCD) methodology for the accurate annotation of oxidized lipids complimented with the RPC-based retention time mapping for the resolution of isomeric species and correction for in-source fragmentation. To address sample-specific epilipidome composition, blood plasma PUFA-containing lipids served as substrates for *in silico* prediction of oxidized species, which in turn were used as an inclusion list for semi-targeted DDA analysis. Using optimized LC-MS/MS workflow and defined fragmentation rules, oxidized complex lipids can be identified in modification type- and position-specific manner.

The new workflow was used for holistic profiling and relative quantification of oxPC, oxCE and oxTG lipids in blood plasma from lean, non-diabetic and diabetic obese individuals, with over 300 modified species identified of which 116 were relatively quantified. We could illustrate significant remodelling of the blood plasma epilipidome upon the development of obesity and its complications. Epilipidomic signature specific to lean individuals was dominated by lipid class-specific patterns of modified octadecanoid acyl chains with an endocrine regulatory potential. Obesity and especially obesity-associated type II diabetes induced significant shift in the epilipidome characterized by the depletion of regulatory octadecanoid signature and the elevation of LPO-derived eicosanoids. This new methodology opens up new perspectives in addressing the complexity of epilipidomic (dys)regulation beyond oxylipin profiling towards holistic understanding of complex lipid modifications and their role as endocrine signals in pathophysiology of metabolic disorders.

## Acknowledgments

Financial support from the German Federal Ministry of Education and Research (BMBF) within the framework of the e:Med research and funding concept for SysMedOS project (to MF) are gratefully acknowledged. We thank Prof. Ralf Hoffmann (Institute of Bioanalytical Chemistry, University of Leipzig) for providing access to his laboratory.

## Author Contributions

MF conceived the project, guided the research, assisted with the experiments and data interpretation, and wrote the manuscript. AC designed and performed most of the experiments, analyzed and interpreted data. PN performed oxidized lipid identification and most of the data analysis and validation. ML prepared blood plasma lipid extracts. DG performed stDDA, PRM analysis and oxidized lipid identification. ZN performed *in silico* epilipidome predictions. MB provided human blood plasma samples. All authors edited and approved the manuscript.

## Supplementary information figures and table legends

**Figure S1.**
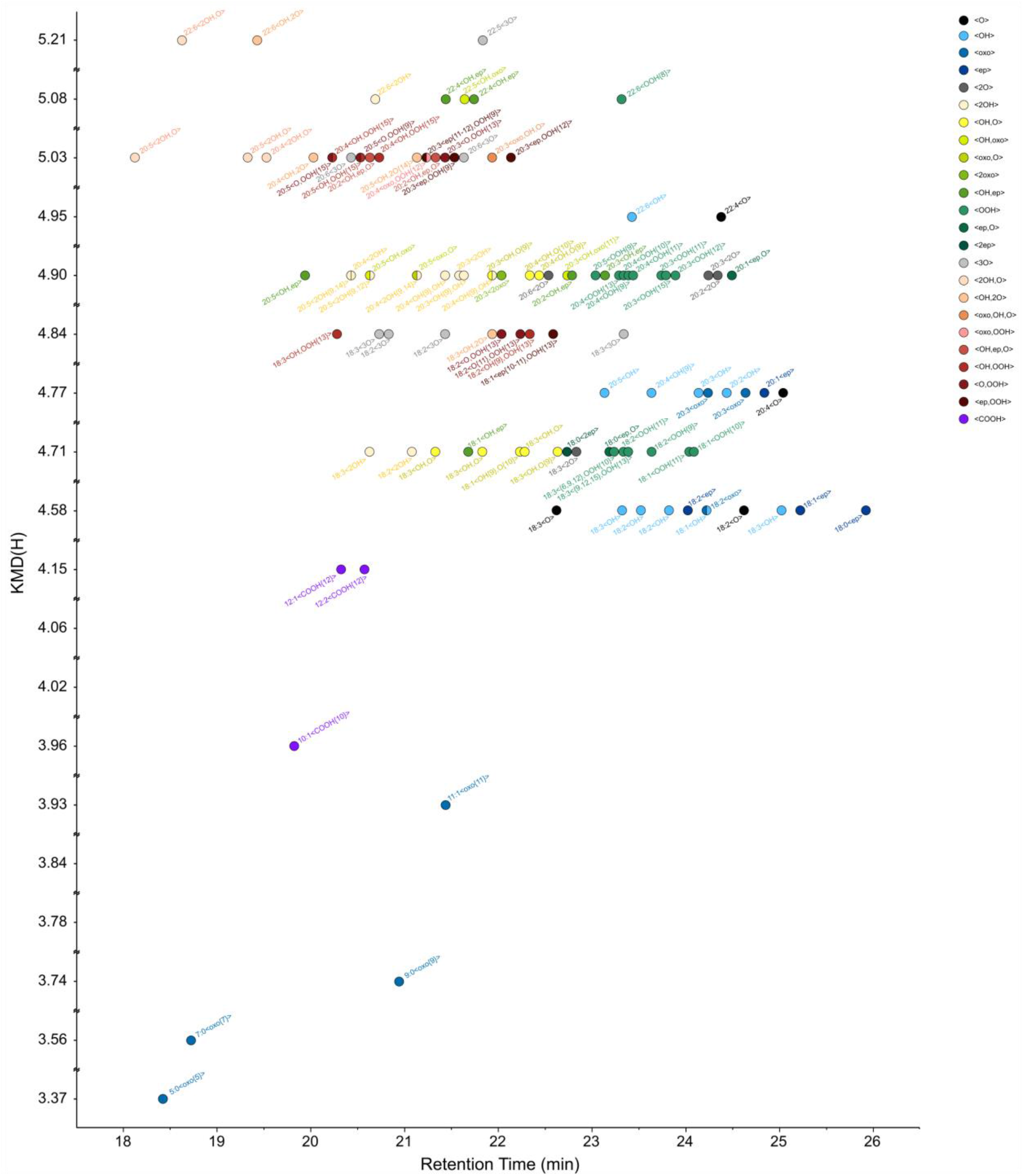
KMD(H) vs retention time plot for annotation of oxCE lipids identified in blood plasma lipid extracts of obese non-diabetic and obese with type II diabetes individuals.

**File S1**. MS2 fragmentation patterns for oxidatively truncated and full-length oxygenated lipids.

**File S2**. MS2 spectra acquired using stDDA and used for the annotation of oxidized PC (oxPC) species in group-pooled blood plasma samples of obese non-diabetic (OND) and obese with type 2 diabetes (OT2D) individuals. Structure-related fragmentation ions are colour-coded according to the legend provided. Annotated lipids for each group pool are sorted by their precursor *m/z*.

**File S3**. MS2 spectra acquired using stDDA and used for the annotation of oxidized CE (oxCE) species in group-pooled blood plasma samples of obese non-diabetic (OND) and obese with type 2 diabetes (OT2D) individuals. Structure-related fragmentation ions are colour-coded according to the legend provided. Annotated lipids for each group pool are sorted by their precursor *m/z*.

**File S4**. MS2 spectra acquired using stDDA and used for the annotation of oxidized TG (oxTG) species in group-pooled blood plasma samples of obese non-diabetic (OND) and obese with type 2 diabetes (OT2D) individuals. Structure-related fragmentation ions are colour-coded according to the legend provided. Annotated lipids for each group pool are sorted by their precursor *m/z*.

**Table S1**. List of annotated oxylipins standards (S1.1), *in vitro* oxPC (S1.2), oxCE (S1.3) and oxTG (S1.4) with corresponding IDs at fatty acyl, modification type and modification position levels. Neutral/ion formula, theoretical and measured mass, detected adducts and their *m/z* as well as all detected structure specific fragments are listed.

**Table S2**. List of annotated oxPC (S2.1), oxCE (S2.2) and oxTG (S2.3) in *in vitro* oxidized human blood plasma with corresponding IDs at fatty acyl, modification type and modification position levels. Neutral/ion formula, theoretical and measured mass, detected adducts and their *m/z* as well as all detected structure specific fragments are listed.

**Table S3**. List of annotated oxPC (S3.1 for OND and S3.2 for OT2D pools), oxCE (S3.4 for OND and S3.4 for OT2D pools) and oxTG (S3.5 for OND and S3.6 for OT2D pools) in human blood plasma of obese non-diabetic (OND) and obese diabetic (OT2D) individulas with corresponding IDs at fatty acyl, modification type and modification position levels. Neutral/ion formula, theoretical and measured mass, detected adducts and their *m/z* as well as all detected structure specific fragments are listed.

**Table S4**. Normalized intensities for PRM quantified oxidized complex lipids in individual blood plasma lipid extracts of lean non-diabetic (LND), obese non-diabetic (OND), and obese with type II diabetes (OT2D) individuals.

**Table S5**. List of oxPC (S5.1), oxCE (S5.2) and oxTG (S5.3) species showed significant differences (ANOVA < 0.05) between the groups upon targeted PRM based quantification.

**Table S6**. List of human blood plasma PUFA-containing PC, CE and TG molecular species used for *in silico* oxidation.

**Table S7**. Annotation of the raw MS data files uploaded to MassIVE MSV000088608.

## Material and Methods

### Chemicals

Acetonitrile (MeCN), isopropanol (*i*-PrOH), methanol (MeOH), and formic acid (all ULC/MS-CC/SFC grade) were purchased from Biosolve (Valkenswaard, Netherlands). Chloroform (CHCl_3_; Emsure®), methyl-*tert*-butyl-ether (MTBE; ≥99%), 3,5-di-*tert*-4-butylhydroxytoluol (BHT), Na-L-ascorbate, CuSO_4_·5H_2_O and the NIST^®^ SRM^®^ 1950 Metabolites in Frozen Plasma were purchased from Sigma-Aldrich (Taufkirchen, Germany). Ammonium formate (NH_4_HCO_2_; MS grade) was purchased from Fluka Analytical (München, Germany). Water (ddH_2_O) was ultrapurified by an ELGA PURELAB Ultra Analytic (Berlin, Germany) instrument delivering water quality of a resistivity ≥ 18.2 MΩ·cm.

### Lipid standards

1-Palmitoyl-2-linoleoyl-*sn*-glycero-3-phosphocholine (PC 16:0/18:2), 1-palmitoyl-2-oleoyl-*sn*-glycero-3-phosphocholine (PC 16:0/18:1), and SPLASH® LIPIDOMIX® were purchased from Avanti Polar Lipids (Avanti Polar Lipids, Inc., Alabama, USA). 1,2-Dipalmitate-3-linoleate-glycerol (TG 16:0/16:0/18:2) and cholesteryl linoleate (CE 18:2) were purchased from Larodan (Solna, Sweden). Oxylipin standards (15(S)-hydroperoxy eicosatetraenoic acid, 15(S)-hydroxy eicosatetraenoic acid, 14(15)-epoxy eicosatetraenoic acid, 15-oxo eicosatetraenoic acid, 8(S),15(S)-dihydroxy eicosatetraenoic acid) were from Cayman Chemical (Ann Arbor, USA).

### *In vitro* oxidation of lipid standards and human plasma

Lipid standards (PC 16:0/18:2, TG 16:0/16:0/18:2, and CE 18:2) and human plasma were oxidized in the presence of copper sulfate and ascorbic acid. Liposomes (PC 16:0/18:2) or micelles (TG 16:0/16:0/18:2 or CE 18:2 mixed with PC 16:0/18:1 in a molar ratio 3:1) were dried in amber vials under the stream of nitrogen and reconstituted in NH_4_HCO_2_ (10 mM, pH 7.5) at the final concentrations of 1.5 mM. CuSO_4_ (final concentration 0.075 mM) and ascorbic acid (final concentration 0.15 mM) were added to the liposomes or micelles (final concentration 1.125 mM) and incubated at 37°C, 1300 rpm for 24 h. Lipids were diluted with *i*-PrOH to the final concentration of 0.15 mM prior to the analysis. Lipids from *in vitro* oxidized blood plasma were extracted as described below.

### Human blood plasma lipid extraction

Plasma samples (n=150) were selected from the Leipzig Obesity BioBank that has been approved by the Ethics committee of the University of Leipzig (approval number: 159-12-21052012). All subjects gave written informed consent before contributing samples and data to the biobank. After collection, samples were frozen and stored at -80°C until the analysis. Donors of plasma samples were selected to compare groups patients with obesity (n=100body mass index (BMI) 30-50kg/m^2^) without (obesity, no diabetes (OND); n=50) or with type 2 diabetes (obesity, type 2 diabetes (OT2D); n=50), and a group of healthy lean controls with a BMI < 25kg/m^2^ (lean, no diabetes (LND); n=50). Data on BMI, gender, and age are provided for these three groups in Table S4.

### Lipid extraction from human blood plasma

For lipid extraction samples were thawed by incubating tubes containing 10 µL of plasma on ice for 1 h. SPLASH® LIPIDOMIX® was added (5 μL) and incubated on ice for 15 min. Lipids were extracted by adding ice cold MeOH (375 µL) and MTBE (1250 µL) with subsequent vortexing^44^. Samples were incubated for 1 h at 4°C (Orbital shaker, 32 rpm). Phase separation was induced by addition of H_2_O (375 µL). Samples were vortexed, incubated for 10 min at 4°C (Orbital shaker, 32 rpm), centrifuged (10 min, 4°C, 1000 x g), the organic phase was collected into a new tube, and the solvent was removed *in vacuo* (Eppendorf concentrator 5301, 1 mbar). A Quality Control (QC) sample was prepared by mixing obtained lipid extracts in an equivolumetric manner. All extraction solvents contained 0.01% (w/v) BHT and were cooled on ice before use.

### *In silico* oxidation

Lipid standards or previously reported^37^ most abundant PUFA-containing human blood plasma PCs (17 molecular species), TGs (17 molecular species) and CEs (11 molecular species)(Table S6) were used for *in silico* oxidation by LPPtiger software source code version^39^ (https://github.com/SysMedOs/lpptiger). The list of modifications included hydroperoxy, hydroxy, epoxy, and keto groups as well as truncated products. The oxidation was performed at the level 1 considering maximum of 2 sites and a maximum one <oxo> and one <OOH>group. Elemental composition of predicted oxidized lipids was used to compose inclusion lists considering preferential ionization adducts used for MS analysis (fomate adduct anions for oxPC, sodiated adduct cations for oxCE and oxTG).

### Chromatography

Ultra-high-performance LC (RP-UHPLC) was carried out on a Vanquish Horizon (Thermo Fisher Scientific, Germering, Germany) equipped with an Accucore C18 column (150 × 2.1 mm: 2.6 µm, 150 Å, Thermo Fisher Scientific, Sunnyvale, CA, USA). Lipids were separated by gradient elution with solvent A (MeCN/H_2_O, 1:1, v/v) and B (IPA/MeCN/H_2_O, 85:10:5, v/v/v) both containing 5 mM HCOONH_4_ and 0.1% (v/v) formic acid. The separation was performed at 50°C with a flow rate of 0.3 mL/min using the following gradient: 0-20 min – 10 to 86 % B, 20-22 min – 86 to 95 % B, 22-26 min –95 % B (isocratic), 26-26.1 min – 95 to 10 % B, 26.1-34.0 min – 10 % B (isocratic, column re-equilibration).

### Mass spectrometry

Method development and optimization were performed using RP-UHPLC coupled on-line either to a Thermo Scientific Q Exactive Plus Quadrupole-Orbitrap (Thermo Fisher Scientific, Bremen, Germany) or Orbitrap Fusion Lumos Tribrid (Thermo Fisher Scientific, San Jose, USA) mass spectrometers.

Q Exactive Plus Quadrupole-Orbitrap was equipped with a heated electrospray (HESI) source and operated in both positive and negative ion modes with the following parameters: sheath gas – 40 arbitrary units, auxiliary gas – 10 arbitrary units, sweep gas – 1 arbitrary units, spray voltage – +3.5 kV and -2.5 kV, capillary temperature – 300 °C, S-lens RF level – 35%, and aux gas heater temperature – 370 °C.

Orbitrap Fusion Lumos Tribrid mass spectrometer using a HESI source was operated in both positive and negative ion modes with the following parameters: spray voltage – +3.5 kV and - 2.8 kV, ion transfer tube temperature – 250°C, sheath gas – 25 arbitrary units, aux gas – 10 arbitrary units, vaporizer temperature – 200°C, RF level – 25%.

### Multistage ion activation (MS3) for analysis of *in vitro* oxidized standards and human plasma

Multistage ion activation experiments were performed on Orbitrap Fusion Lumos Tribrid mass spectrometer using inclusion list DDA mode. Full MS was acquired at the resolution 120 000 at *m/z* 200, scan range 450 – 1000 *m/z*, AGC target 4e^5^ counts, maximum injection time (max IT) 100 ms. MS/MS events were triggered for precursor ions from the inclusion lists. MS2 spectra were acquired at the resolution 15 000 at *m/z* 200, AGC target 8e^4^, max IT 100 ms, isolation window 1 *m/z*, normalized collision energy (nCE) 25% (negative mode) and 37% (positive mode). The filters used were MIPS (small molecule), charge state (1), dynamic exclusion (exclusion duration 6 s, mass tolerance ± 10 ppm), target exclusion (polarity-specific) and an inclusion list (class-specific).

MS3 analysis was conducted using the orbitrap at the resolution setting of 15 000 for *m/z* 200, AGC target 1e^5^, max IT 250 ms, injection ions for all available parallelizable time, isolation window 2 m/z, nCE 33%). The following filters were used: precursor selection range (100-450 *m/z*); targeted exclusion mass list (including non-oxidized fatty acid list, mass tolerance ± 50 ppm); product ion trigger (including non-oxidized fatty acid list, mass tolerance ± 25 ppm).

### Semi-targeted data-dependent acquisition (stDDA) for oxidized lipids identification

stDDA analysis was performed on Q Exactive Plus Quadrupole-Orbitrap mass spectrometer. Full MS spectra were acquired at the resolution 140 000 at *m/z* 200, scan range 500 – 900 *m/z* (negative ion mode, 0-17 min) and 600 – 1000 *m/z* (positive ion mode, 17-34 min), AGC target 1e^6^ counts, max IT 100 ms. MS/MS events (top 6) were triggered for precursor ions from the inclusion lists (*in-silico* oxidized PC in negative, and oxCE and oxTG in positive mode). MS/MS spectra were acquired at the resolution 17 500 at *m/z* 200, AGC target 1e^5^, max IT 200 ms, loop count 6, isolation window 1.2 *m/z*, fixed first mass 100.0 *m/z*, stepped nCE 20-30-40 (negative mode) and 30-40-50 (positive mode). The filters used were minimum AGC target (1e^1^), charge state (1), isotope exclusion (on), apex trigger (up to 6 s), and dynamic exclusion (3 s). All spectra were acquired in profile mode.

### Targeted parallel reaction monitoring (PRM) for oxidized lipids relative quantification

For PRM, inclusion lists were used in retention time-scheduled negative (0-17 min) and positive modes (17-34 min). MS/MS spectra were acquired at the resolution 17 500 at *m/z* 200, AGC target 2e^5^ counts, max IT 200 ms, isolation window 1.2 *m/z*, stepped nCE 20-30-40 (negative mode) and 30-40-50 (positive mode).

### Identification and quantification of oxidized lipids

Oxidized lipids were identified by LPPtiger^39^ and further annotated by manual control for oxidized fatty acyl (oxFA)-, modification type- and position-specific fragment ion signals in MS/MS spectra generated in stDDA epilipidomics experiments. Lists of *in-silico* oxidized lipids generated by LPPtiger with possible “precursor-oxFA fragment” pairs as well as fragmentation rules defined for the *in vitro* oxidized standards (File S1) were used as a support. All MS/MS spectra were screened in Qual Browser, Thermo Xcalibur v. 4.2.47 (Thermo Fisher Scientific Inc.).

Quantification of oxidized lipids was performed using PRM-derived data in Skyline v.21.1.0.146 (MacCoss Lab)^45^. For each targeted precursor, the oxFA fragment ion was selected for the peak integration. The peak boundaries were delimited, manually corrected and verified. The obtained peak areas were normalized by appropriate lipid species from SPLASH® Lipidomix Mass Spec Standard (Avanti): LPC(18:1)(d7) for truncated oxPC, PC(15:0/18:1)(d7) for full-length oxPC, CE(18:1)(d7) and TG(15:0/18:1(d7)/15:0 for oxCE and oxTG, correspondingly.

## Data analysis and visualization

For RT mapping, hydrogen-based Kendrick mass defects were calculated for each identified lipid and plotted against RT using R-scripts available at https://github.com/SysMedOs/AdipoAtlasScripts/tree/main/DataVisualization.

Quantified normalized peak areas were median-centred, log-transformed, and autoscaled in MetaboAnalyst v.5 (https://www.metaboanalyst.ca, Xia Lab)^46^. Data processed by MetaboAnalyst were exported as CSV files and used in other software for visualisation.

Oxidized lipids with statistically significant difference were selected for heatmaps. The latter were created in Genesis v.1.8.1 (Bioinformatics TU-Graz) ^47^. Features (oxidized lipids) were clustered by average linkage WPGMA (weighted pair group method with arithmetic mean) agglomeration rule. Other graphs were created in OriginPro 2019 v. 9.6.0.172 (Academic) (OriginLab Corporation).

## Data availability

All raw LC-MS/MS files are available at MassIVE MSV000088608. File names correspond to the following LC-MS/MS datasets (Table S7).

