## Supplementary material for "Epilipidomics platform for holistic profiling of oxidized complex lipids in blood plasma of obese individuals": FileS1

#### File S1. MS2 fragmentation patterns for oxidatively truncated and full-length oxygenated lipids.

**General notes:**

1. All *m/z* values given in the table were calculated considering the charge.
2. Representative $MS^2$ spectra do not always contain signals corresponding to the position-specific fragments (the latter were acquired in $MS^3$ experiments).
3. For simplicity, only *E*-conformers of PUFAs are depicted. Fragmentation patterns do not allow to elucidate the configuration of a double bond, but keep in mind the natural *Z*-configuration for some double bonds in (ox)PUFAs.
4. Modification- and position-specific fragments: the most intense ones are highlighted in **bold**, the intermediate intensity ones are regular, and the weak ones are grayed out.

### PHOSPHATIDYLCHOLINES

#### Oxidatively truncated: <COOH>

[M-H]$^-$ adducts (precursors)

| Mod. position | Precursor | Chemical Formula | *m/z* | MS$^2$ Fragmentation | Mod./pos.-specific fragment[1] | Chemical Formula | *m/z* |
|---|---|---|---|---|---|---|---|
| **4** | PC(16:0/4:0<COOH>) | $C_{28}H_{53}O_{10}NP$ | 594.3413 | 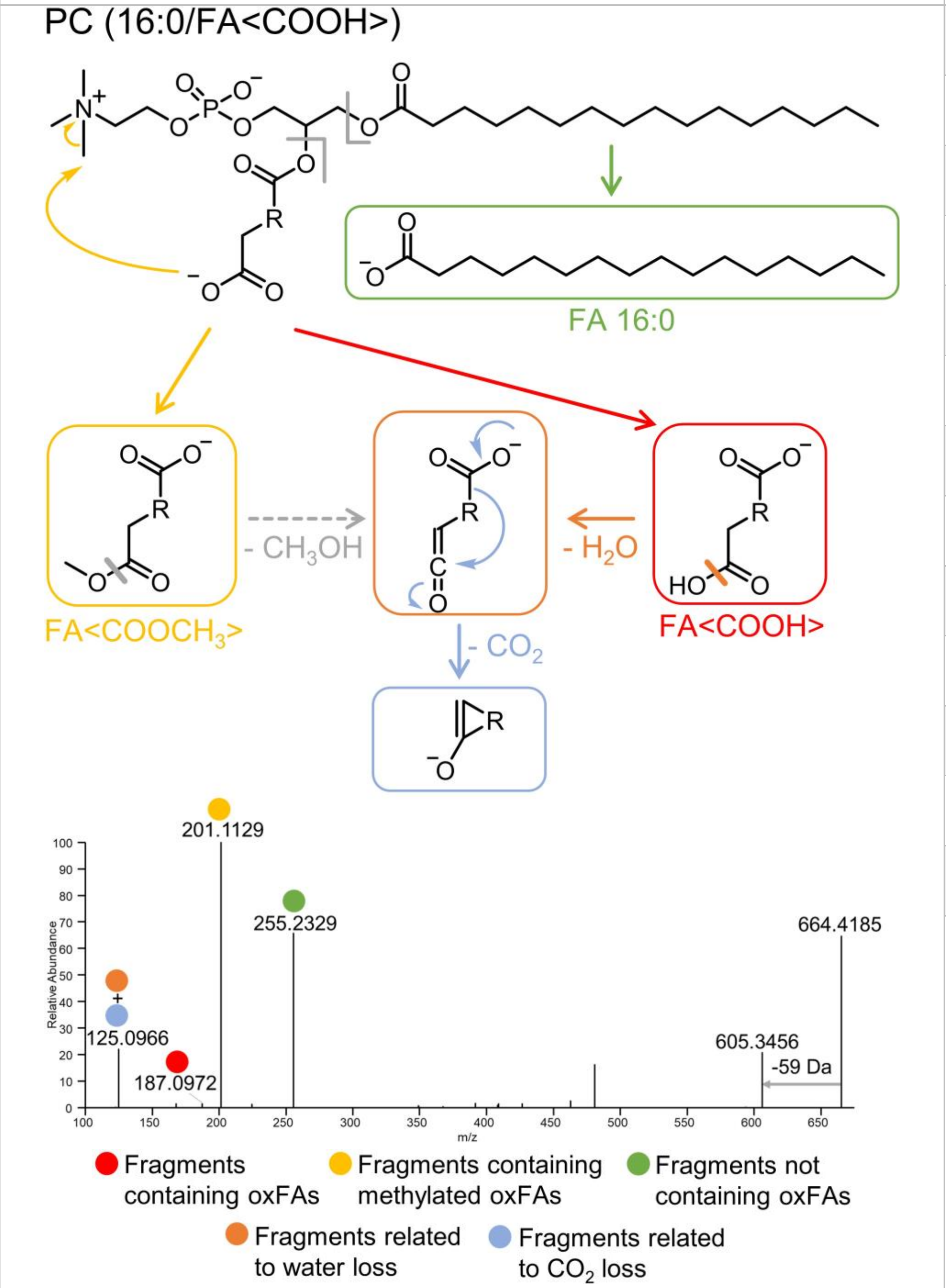 | FA(4:0<COOCH$_3$>) | $C_5H_7O_4$ | 131.0350 |
| **5** | PC(16:0/5:0<COOH>) | $C_{29}H_{55}O_{10}NP$ | 608.3569 | | FA(5:0<COOCH$_3$>) | $C_6H_9O_4$ | 145.0506 |
| **7** | PC(16:0/7:0<COOH>) | $C_{31}H_{59}O_{10}NP$ | 636.3882 | | FA(7:0<COOCH$_3$>) | $C_8H_{13}O_4$ | 173.0819 |
| **8** | PC(16:0/8:0<COOH>) | $C_{32}H_{61}O_{10}NP$ | 650.4039 | | FA(8:0<COOCH$_3$>) | $C_9H_{15}O_4$ | 187.0976 |
| **9** | PC(16:0/9:0<COOH>) | $C_{33}H_{63}O_{10}NP$ | 664.4195 | | FA(9:0<COOCH3>) | $C_{10}H_{17}O_4$ | 201.1132 |
| **10** | PC(16:0/10:0<COOH>) | $C_{34}H_{65}O_{10}NP$ | 678.4352 | | FA(10:0<COOCH$_3$>) | $C_{11}H_{19}O_4$ | 215.1289 |
| | PC(16:0/10:1<COOH>) | $C_{34}H_{63}O_{10}NP$ | 676.4195 | | FA(10:1<COOCH$_3$>) | $C_{11}H_{17}O_4$ | 213.1132 |
| **11** | PC(16:0/11:1<COOH>) | $C_{35}H_{65}O_{10}NP$ | 690.4352 | | FA(11:1<COOCH$_3$>) | $C_{12}H_{19}O_4$ | 227.1289 |
| | PC(16:0/11:2<COOH>) | $C_{35}H_{63}O_{10}NP$ | 688.4195 | | FA(11:2<COOCH$_3$>) | $C_{12}H_{17}O_4$ | 225.1132 |
| **12** | PC(16:0/12:1<COOH>) | $C_{36}H_{67}O_{10}NP$ | 704.4508 | | FA(12:1<COOCH$_3$>) | $C_{13}H_{21}O_4$ | 241.1445 |
| | PC(16:0/12:2<COOH>) | $C_{36}H_{65}O_{10}NP$ | 702.4352 | | FA(12:2<COOCH$_3$>) | $C_{13}H_{19}O_4$ | 239.1289 |
| **13** | PC(16:0/13:2<COOH>) | $C_{37}H_{67}O_{10}NP$ | 716.4508 | | FA(13:2<COOCH$_3$>) | $C_{14}H_{21}O_4$ | 253.1445 |
| | PC(16:0/13:3<COOH>) | $C_{37}H_{65}O_{10}NP$ | 714.4352 | | FA(13:3<COOCH$_3$>) | $C_{14}H_{19}O_4$ | 251.1289 |
| **14** | PC(16:0/14:3<COOH>) | $C_{38}H_{67}O_{10}NP$ | 728.4508 | | FA(14:3<COOCH$_3$>) | $C_{15}H_{21}O_4$ | 265.1445 |

[1] Although other characteristic fragments can be formed occasionally, "Mod./pos.-specific fragment" column contains the most reproducible fragment ion *m/z*. Such fragments as FA<COOH>, FA<COOH>-$CO_2$, FA<COOH>-$H_2O$ (or FA<COOCH$_3$>-$CH_3OH$) were occasionally observed, and might be also used as supporting the annotation.

#### Oxidatively truncated: <oxo>

[M+HCOO]⁻ adducts (precursors)

| Mod. position | Precursor | Chemical Formula | *m/z* | MS² Fragmentation | Mod./pos.-specific fragment[2] | Chemical Formula | *m/z* |
|---|---|---|---|---|---|---|---|
| **4** | PC(16:0/4:0<oxo>) | $C_{29}H_{55}O_{11}NP$ | 624.3518 | 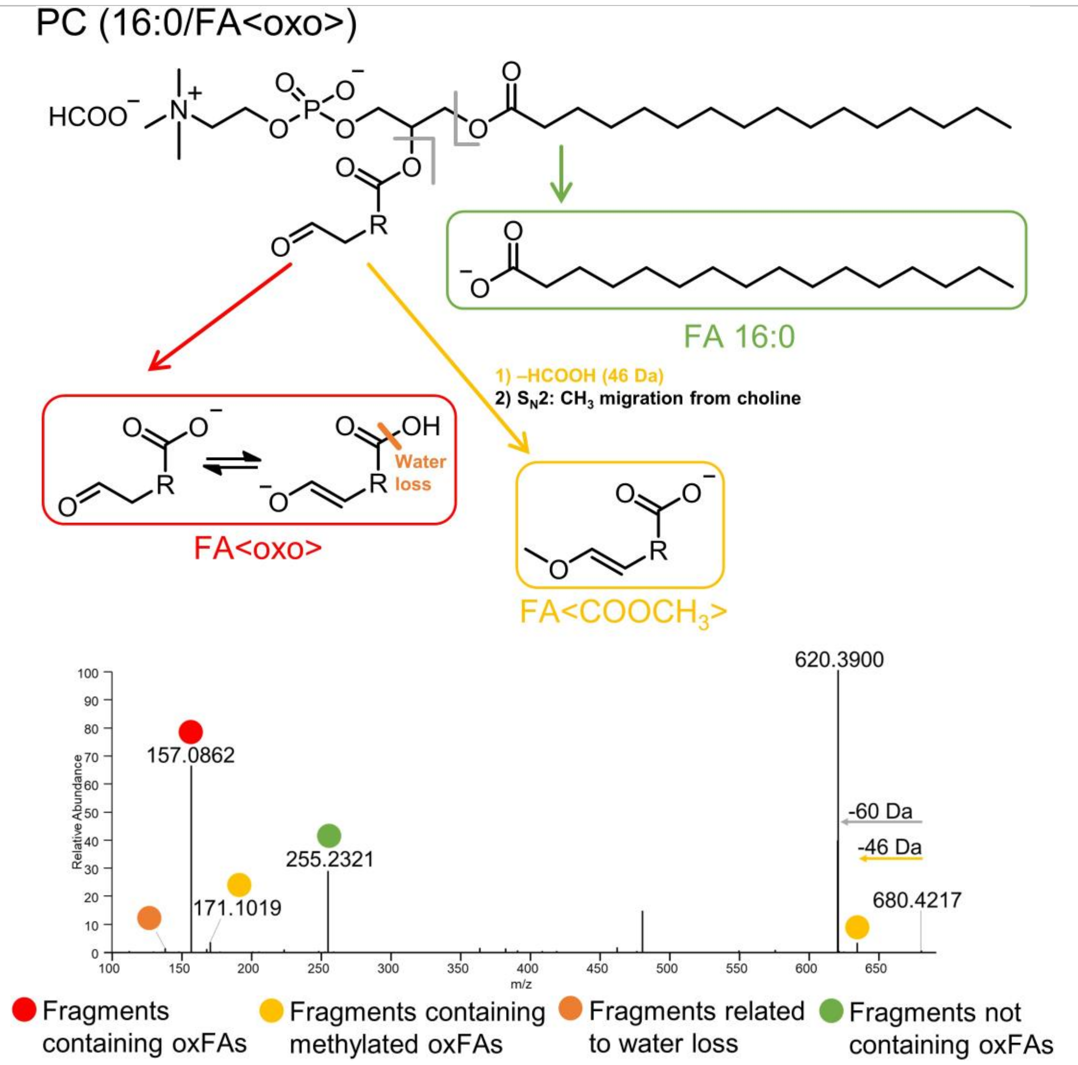 | FA(4:0<oxo>) | $C_4H_5O_3$ | 101.0244 |
| **5** | PC(16:0/5:0<oxo>) | $C_{30}H_{57}O_{11}NP$ | 638.3675 | | FA(5:0<oxo>) | $C_5H_7O_3$ | 115.0401 |
| **7** | PC(16:0/7:0<oxo>) | $C_{32}H_{61}O_{11}NP$ | 666.3988 | | FA(7:0<oxo>) | $C_7H_{11}O_3$ | 143.0714 |
| **8** | PC(16:0/8:0<oxo>) | $C_{33}H_{63}O_{11}NP$ | 680.4144 | | FA(8:0<oxo>) | $C_8H_{13}O_3$ | 157.0870 |
| **9** | PC(16:0/9:0<oxo>) | $C_{34}H_{65}O_{11}NP$ | 694.4301 | | FA(9:0<oxo>) | $C_9H_{15}O_3$ | 171.1027 |
| **10** | PC(16:0/10:0<oxo>) | $C_{35}H_{67}O_{11}NP$ | 708.4457 | | FA(10:0<oxo>) | $C_{10}H_{17}O_3$ | 185.1183 |
| | PC(16:0/10:1<oxo>) | $C_{35}H_{65}O_{11}NP$ | 706.4301 | | FA(10:1<oxo>) | $C_{10}H_{15}O_3$ | 183.1027 |
| **11** | PC(16:0/11:1<oxo>) | $C_{36}H_{67}O_{11}NP$ | 720.4457 | | FA(11:1<oxo>) | $C_{11}H_{17}O_3$ | 197.1183 |
| | PC(16:0/11:2<oxo>) | $C_{36}H_{65}O_{11}NP$ | 718.4301 | | FA(11:2<oxo>) | $C_{11}H_{15}O_3$ | 195.1027 |
| **12** | PC(16:0/12:1<oxo>) | $C_{37}H_{69}O_{11}NP$ | 734.4614 | | FA(12:1<oxo>) | $C_{12}H_{19}O_3$ | 211.1340 |
| | PC(16:0/12:2<oxo>) | $C_{37}H_{67}O_{11}NP$ | 732.4457 | | FA(12:2<oxo>) | $C_{12}H_{17}O_3$ | 209.1183 |
| **13** | PC(16:0/13:2<oxo>) | $C_{38}H_{69}O_{11}NP$ | 746.4614 | | FA(13:2<oxo>) | $C_{13}H_{19}O_3$ | 223.1340 |
| | PC(16:0/13:3<oxo>) | $C_{38}H_{67}O_{11}NP$ | 744.4457 | | FA(13:3<oxo>) | $C_{13}H_{17}O_3$ | 221.1183 |
| **14** | PC(16:0/14:3<oxo>) | $C_{39}H_{69}O_{11}NP$ | 758.4614 | | FA(14:3<oxo>) | $C_{14}H_{19}O_3$ | 235.1340 |

[2] + additional weak signals in MS² spectra: FA<COOH> -$H_2O$ (or FA<COOCH₃>-$CH_3OH$), FA<COOH>-$CO_2$.

#### Full-length oxygenated: <oxo>

[M+HCOO]$^{-}$ adducts (precursors)

| Precursor | Chemical Formula & *m/z* | MS$^2$ Fragmentation | Mod.-specific fragment | Chemical Formula & *m/z* | Pos.-specific fragmentation | Chemical Formula | *m/z* |
|---|---|---|---|---|---|---|---|
| **PC(16:0/18:1<oxo>)** | $C_{43}H_{81}O_{11}NP$ 818.5553 | PC (16:0/FA<oxo>) 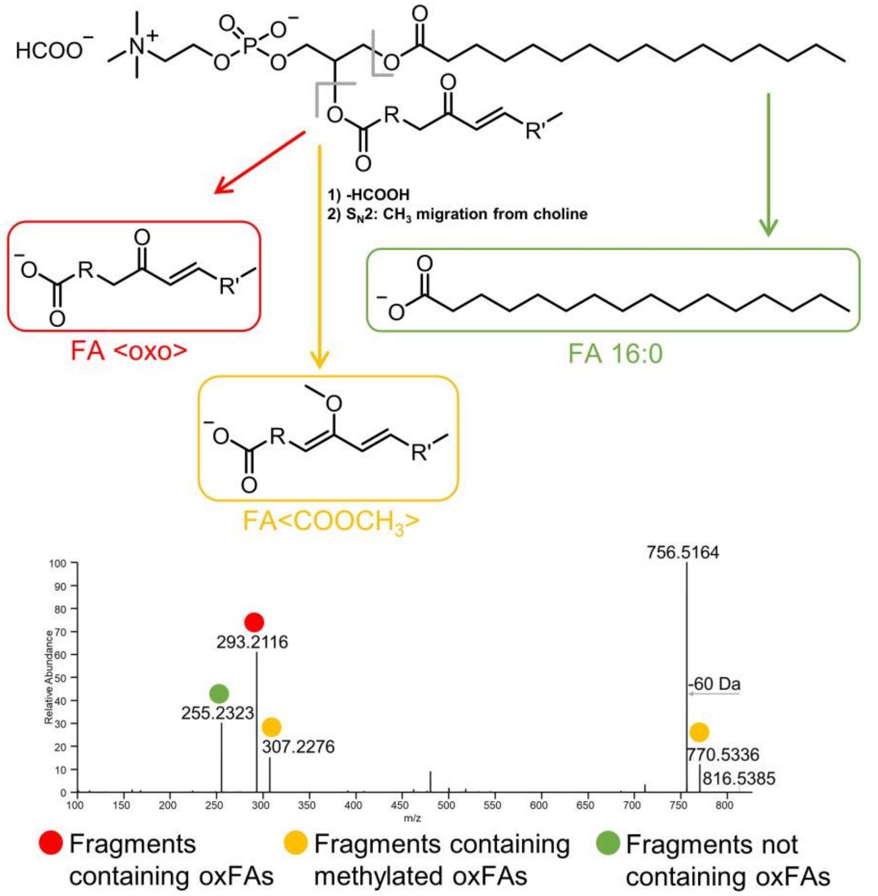  * Signal of precursor ion might not always be present in MS$^2$ spectra due to the intensive fragmentation **FA fragmentation pattern:** 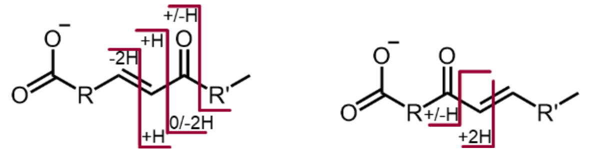 | **FA(18:1<oxo>)** Methylation CO2 loss | **$C_{18}H_{31}O_3$ 295.2279** $C_{19}H_{33}O_3$ 309.2435 $C_{17}H_{31}O$ 251.2380 | 9 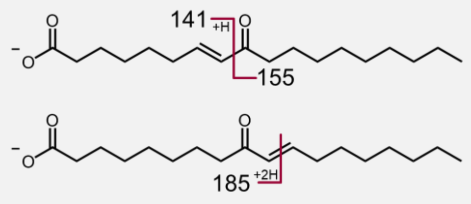 | $C_8H_{13}O_2$ $C_{10}H_{19}O$ $C_{10}H_{17}O_3$ | 141.0921 155.1441 185.1183 |
| **PC(16:0/18:2<oxo>)** | $C_{43}H_{79}O_{11}NP$ 816.5396 | | **FA(18:2<oxo>)** Methylation CO2 loss | **$C_{18}H_{29}O_3$ 293.2122** $C_{19}H_{31}O_3$ 307.2279 $C_{17}H_{29}O$ 249.2224 | 8 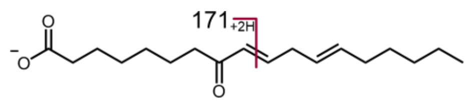 | **$C_9H_{15}O_3$** | **171.1027** |
| | | | | | 9 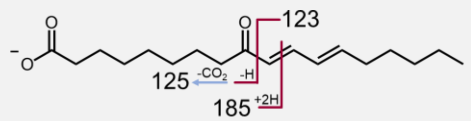 | $C_9H_{15}$ $C_8H_{13}O$ **$C_{10}H_{17}O_3$** | 123.1179 125.0972 **185.1183** |
| | | | | | 10 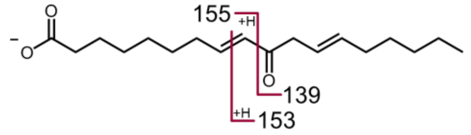 | **$C_9H_{15}O$** $C_{10}H_{17}O$ $C_9H_{15}O_2$ | **139.1128** 153.1285 155.1078 |
| | | | | | 11 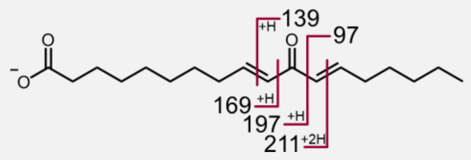 | $C_7H_{13}$ $C_9H_{15}O$ $C_{10}H_{17}O_2$ $C_{11}H_{17}O_3$ $C_{12}H_{19}O_3$ | 97.1023 139.1128 169.1234 197.1183 211.1340 |
| | | | | | 12 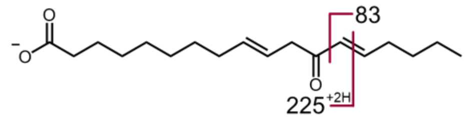 | $C_6H_{11}$ $C_{13}H_{21}O_3$ | 83.0866 225.1496 |
| | | | | | 13 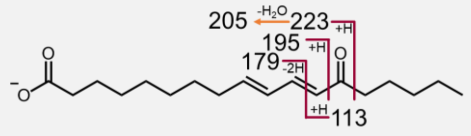 | **$C_7H_{13}O$** $C_{11}H_{15}O_2$ $C_{12}H_{19}O_2$ $C_{13}H_{17}O_2$ $C_{13}H_{19}O_3$ | **113.0972** 179.1078 195.1391 205.1234[3] 223.1340[4] |
| | | | | | 14 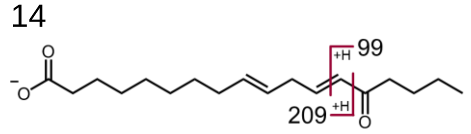 | $C_6H_{11}O$ $C_{13}H_{21}O_2$ | 99.0815 209.1547 |

[3] The signal is seen in the case of FA(18:2<OOH{13}>).

[4] The signal is seen in the case of FA(18:2<OOH{13}>).

#### Full-length oxygenated: <oxo> (continued)

[M+HCOO]⁻ adducts (precursors)

| Precursor | Chemical Formula & *m/z* | MS² Fragmentation | Mod.-specific fragment | Chemical Formula & *m/z* | Pos.-specific fragmentation | Chemical Formula | *m/z* |
|---|---|---|---|---|---|---|---|
| **PC(16:0/20:4<oxo>)** | $C_{45}H_{79}O_{11}NP$<br>840.5396 | 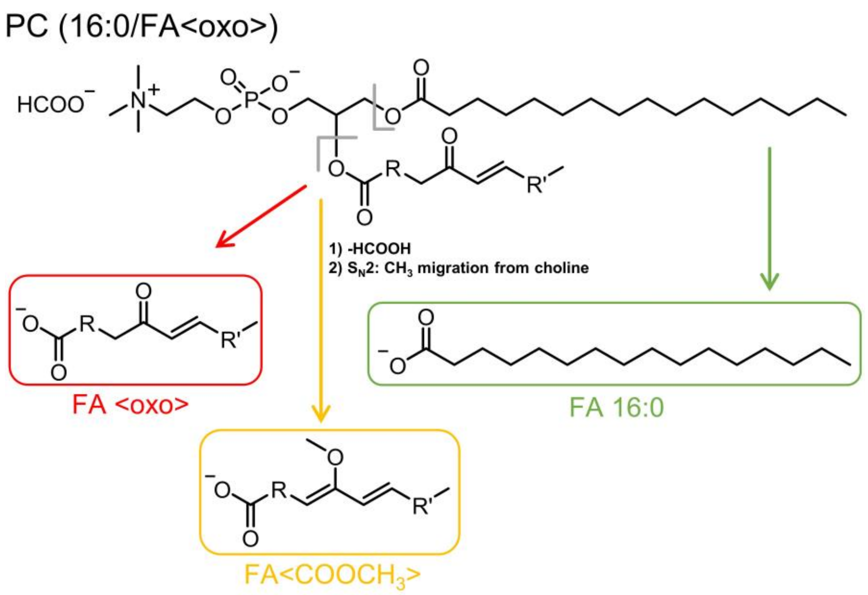<br>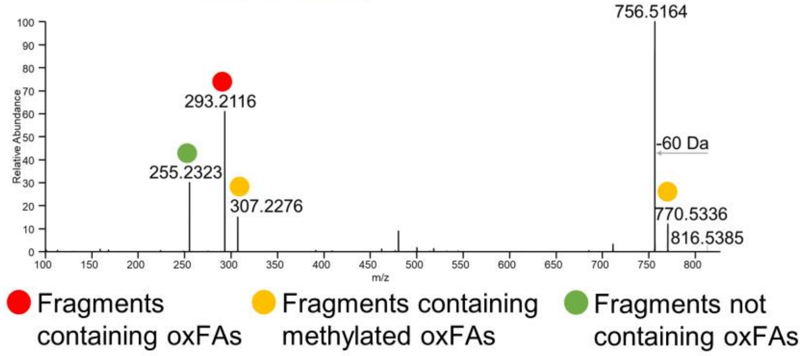<br>* The precursor ion m/z may not always present in MS² spectra due to the intensive fragmentation<br>**FA fragmentation pattern:**<br>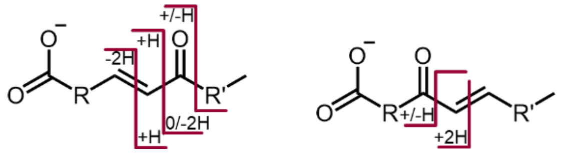 | **FA(20:4<oxo>)**<br>Methylation<br>CO2 loss | **$C_{20}H_{29}O_3$**<br>**317.2122**<br>$C_{21}H_{31}O_3$<br>331.2279<br>$C_{19}H_{29}O$<br>273.2224 | 5<br>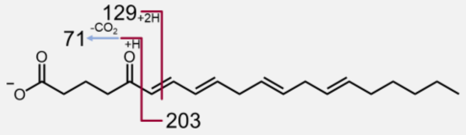 | $C_4H_7O$<br>$C_6H_9O_3$<br>**$C_{15}H_{23}$** | 71.0502<br>129.0557<br>**203.1805** |
| | | | | | 6<br>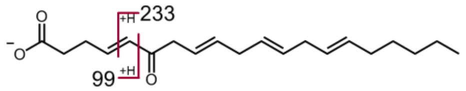 | $C_5H_7O_2$<br>$C_{16}H_{25}O$ | 99.0452<br>233.1911 |
| | | | | | 7<br>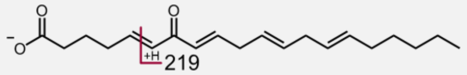 | $C_{15}H_{23}O$ | 219.1754 |
| | | | | | 8<br>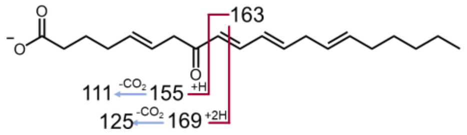 | $C_7H_{11}O$<br>$C_8H_{13}O$<br>$C_8H_{11}O_3$<br>$C_{12}H_{19}$<br>$C_9H_{13}O_3$ | 111.0815<br>125.0972<br>155.0714<br>163.1492<br>169.0870 |
| | | | | | 9<br>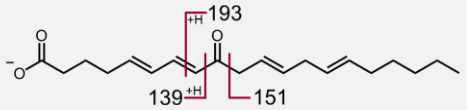 | $C_8H_{11}O_2$<br>$C_{11}H_{19}$<br>$C_{13}H_{21}O$ | 139.0765<br>151.1492<br>193.1598 |
| | | | | | 11<br>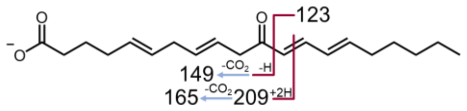 | $C_9H_{15}$<br>$C_{10}H_{13}O$<br>$C_{11}H_{17}O$<br>$C_{12}H_{17}O_3$ | 123.1128<br>149.0972<br>165.1285<br>209.1183 |
| | | | | | 12<br>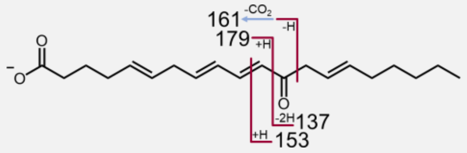 | $C_9H_{13}O$<br>**$C_{10}H_{17}O$**<br>$C_{11}H_{13}O$<br>$C_{11}H_{15}O_2$ | 137.0972<br>**153.1285**<br>161.0971<br>179.1078 |
| | | | | | 13<br>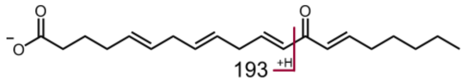 | $C_{12}H_{17}O_2$ | 193.1234 |
| | | | | | 14<br>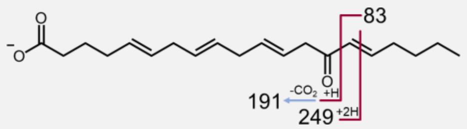 | $C_6H_{11}$<br>**$C_{13}H_{19}O$**<br>$C_{15}H_{21}O_3$ | 83.0866<br>**191.1441**<br>249.1496 |
| | | | | | 15<br>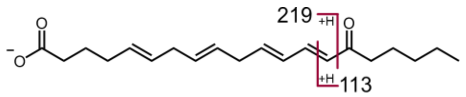 | $C_7H_{13}O$<br>$C_9H_{15}O$<br>$C_{14}H_{19}O_2$ | **113.0972**<br>139.1128<br>219.1391 |

#### Full-length oxygenated: <OH>

[M+HCOO]$^-$ adducts (precursors)

| Precursor | Chemical Formula & *m/z* | MS$^2$ Fragmentation | Mod.-specific fragment | Chemical Formula & *m/z* | Pos.-specific fragmentation | Chemical Formula | *m/z* |
|---|---|---|---|---|---|---|---|
| **PC(16:0/18:1<OH>)** | $C_{43}H_{83}O_{11}NP$ 820.5709 | 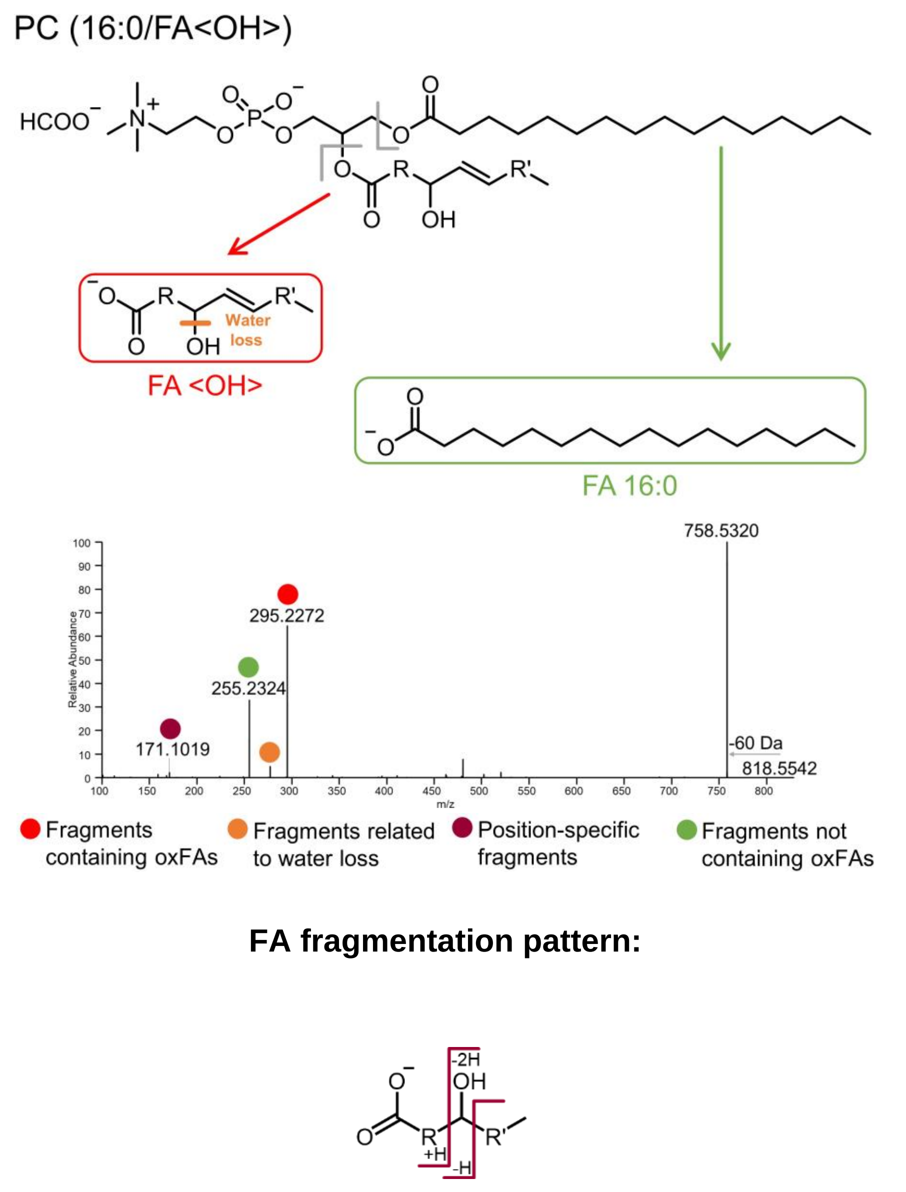 | **FA(18:1<OH>)**<br>$H_2O$ loss<br>$CO_2$ loss<br>$H_2O$&$CO_2$ loss | **$C_{18}H_{33}O_3$ 297.2435**<br>$C_{18}H_{31}O_2$ 279.2330<br>$C_{17}H_{33}O$ 253.2536<br>$C_{17}H_{31}$ 235.2431 | 9 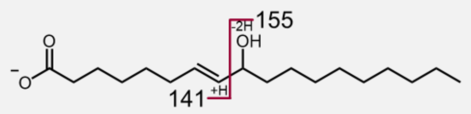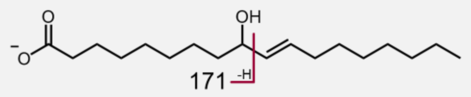 | $C_8H_{13}O_2$<br>$C_{10}H_{19}O$<br>**$C_9H_{15}O_3$** | 141.0921<br>155.1441<br>**171.1027** |
| **PC(16:0/18:2<OH>)** | $C_{43}H_{81}O_{11}NP$ 818.5553 | | **FA(18:2<OH>)**<br>$H_2O$ loss<br>$CO_2$ loss<br>$H_2O$&$CO_2$ loss | **$C_{18}H_{31}O_3$ 295.2279**<br>$C_{18}H_{29}O_2$ 277.2173<br>$C_{17}H_{31}O$ 251.2380<br>$C_{17}H_{29}$ 233.2275 | 9 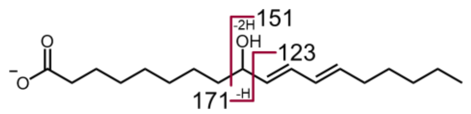 | $C_9H_{15}$<br>$C_{10}H_{15}O$<br>**$C_9H_{15}O_3$** | 123.1179<br>151.1128<br>**171.1027** |
| | | | | | 10 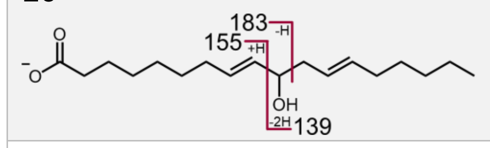 | $C_9H_{15}O$<br>$C_9H_{15}O_2$<br>$C_{10}H_{15}O_3$ | 139.1128<br>155.1078<br>183.1027 |
| | | | | | 11 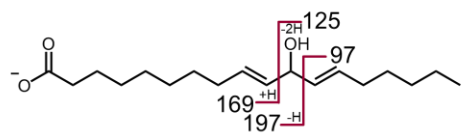 | $C_7H_{13}$<br>$C_8H_{13}O$<br>$C_{10}H_{17}O_2$<br>$C_{11}H_{17}O_3$ | 97.1023<br>125.0972<br>169.1234<br>197.1183 |
| | | | | | 12 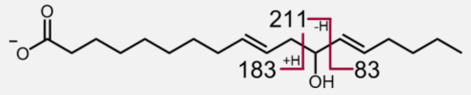 | $C_6H_{11}$<br>**$C_{11}H_{19}O_2$**<br>$C_{12}H_{19}O_3$ | 83.0866<br>**183.1391**<br>211.1340 |
| | | | | | 13 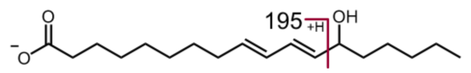 | **$C_{12}H_{19}O_2$** | **195.1391** |
| | | | | | 14 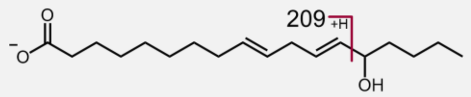 | $C_{13}H_{21}O_2$ | 209.1547 |

#### Full-length oxygenated: <OH> (continued)

[M+HCOO]⁻ adducts

| Precursor | Chemical Formula & *m/z* | MS² Fragmentation | Mod.-specific fragment | Chemical Formula & *m/z* | Pos.-specific fragmentation | Chemical Formula | *m/z* |
|---|---|---|---|---|---|---|---|
| **PC(16:0/20:4<OH>)** | $C_{45}H_{81}O_{11}NP$<br>842.5553 | 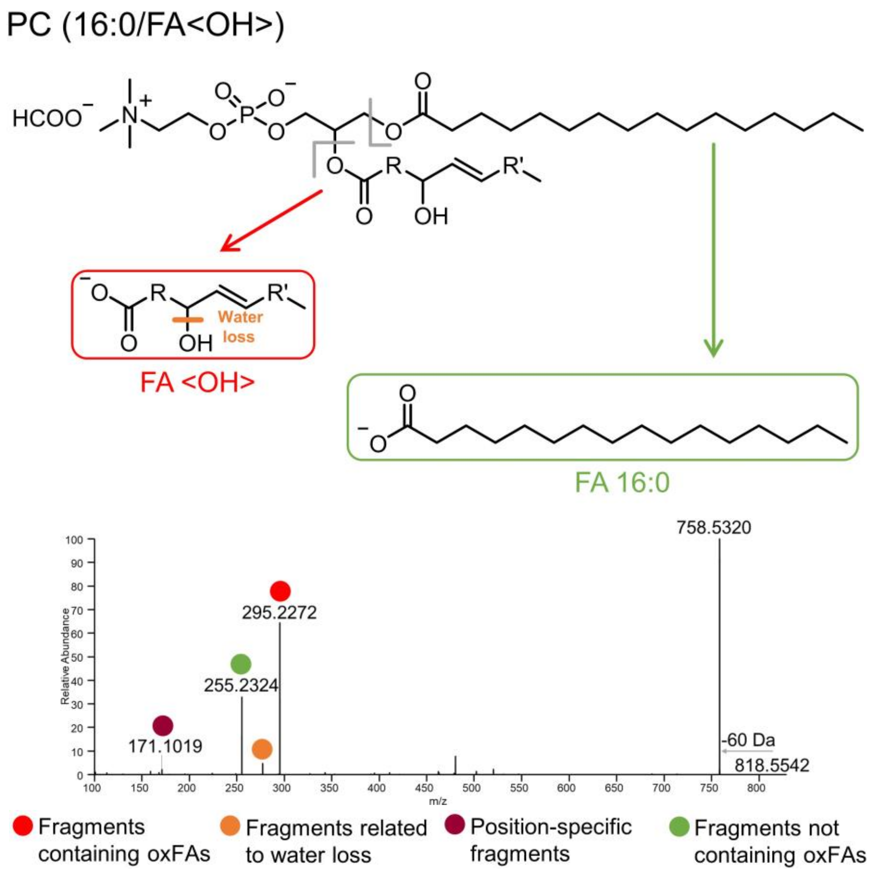<br>**FA fragmentation pattern:**<br>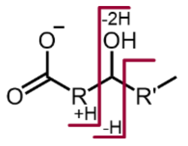 | **FA(20:4<OH>)**<br>$H_2O$ loss<br>$CO_2$ loss<br>$H_2O$&$CO_2$ loss | **$C_{20}H_{31}O_3$**<br>**319.2279**<br>$C_{20}H_{29}O_2$<br>301.2173<br>$C_{19}H_{31}O$<br>275.2380<br>$C_{19}H_{29}$<br>257.2275 | 5<br>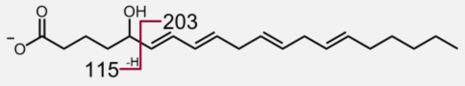 | **$C_5H_7O_3$**<br>$C_{15}H_{23}$ | **115.0401**<br>203.1805[5] |
| | | | | | 6<br>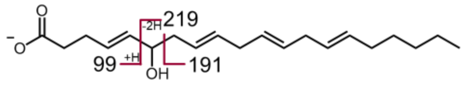 | $C_5H_7O_2$<br>$C_{14}H_{23}$<br>$C_{15}H_{23}O$ | 99.0452<br>191.1805<br>219.1754 |
| | | | | | 7<br>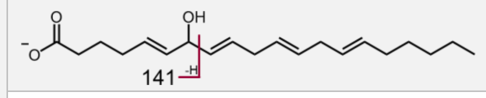 | $C_7H_9O_3$ | 141.0557 |
| | | | | | 8<br>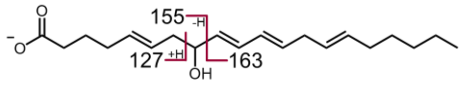 | $C_7H_{11}O_2$<br>**$C_8H_{11}O_3$**<br>**$C_{12}H_{19}$** | 127.0765<br>**155.0714**<br>**163.1492** |
| | | | | | 9<br>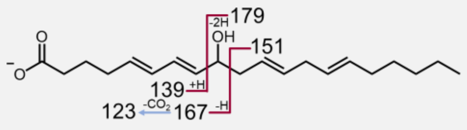 | **$C_8H_{11}O$**<br>$C_8H_{11}O_2$<br>$C_{11}H_{19}$<br>$C_9H_{11}O_3$<br>**$C_{12}H_{19}O$** | **123.0815**<br>139.0765<br>151.1492<br>167.0714<br>**179.1441** |
| | | | | | 11<br>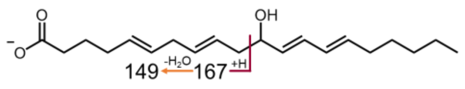 | $C_{10}H_{13}O$<br>**$C_{10}H_{15}O_2$** | 149.0972<br>**167.1078** |
| | | | | | 12<br>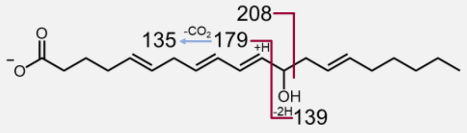 | $C_{10}H_{15}$<br>$C_9H_{15}O$<br>**$C_{11}H_{15}O_2$**<br>$C_{12}H_{16}O_3$ | 135.1179<br>139.1128<br>**179.1078**<br>208.1105[6] |
| | | | | | 13<br>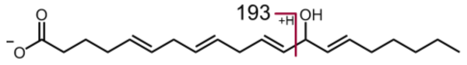 | $C_{12}H_{17}O_2$ | 193.1234 |
| | | | | | 14<br>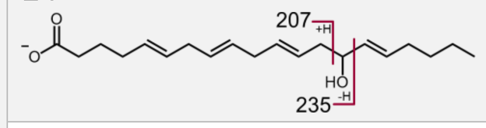 | **$C_{13}H_{19}O_2$**<br>$C_{14}H_{19}O_3$ | **207.1391**<br>235.1340 |
| | | | | | 15<br>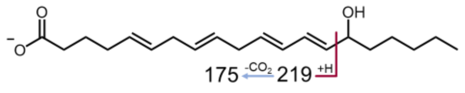 | **$C_{13}H_{19}$**<br>**$C_{14}H_{19}O_2$** | **175.1492**<br>**219.1391** |

[5] The signal is not OH{5}-specific (can be found in the spectra of other isomers, but less intense)
[6] The signal should correspond to an anion-radical that is the unique case for position 12.

#### Full-length oxygenated: <ep>

$[M+HCOO]^-$ adducts

| Precursor | Chemical Formula & *m/z* | MS$^2$ Fragmentation | Mod.-specific fragment | Chemical Formula & *m/z* | Pos.-specific fragmentation | Chemical Formula | *m/z* |
|---|---|---|---|---|---|---|---|
| **PC(16:0/18:0<ep>)** | $C_{43}H_{83}O_{11}NP$<br>820.5709 | 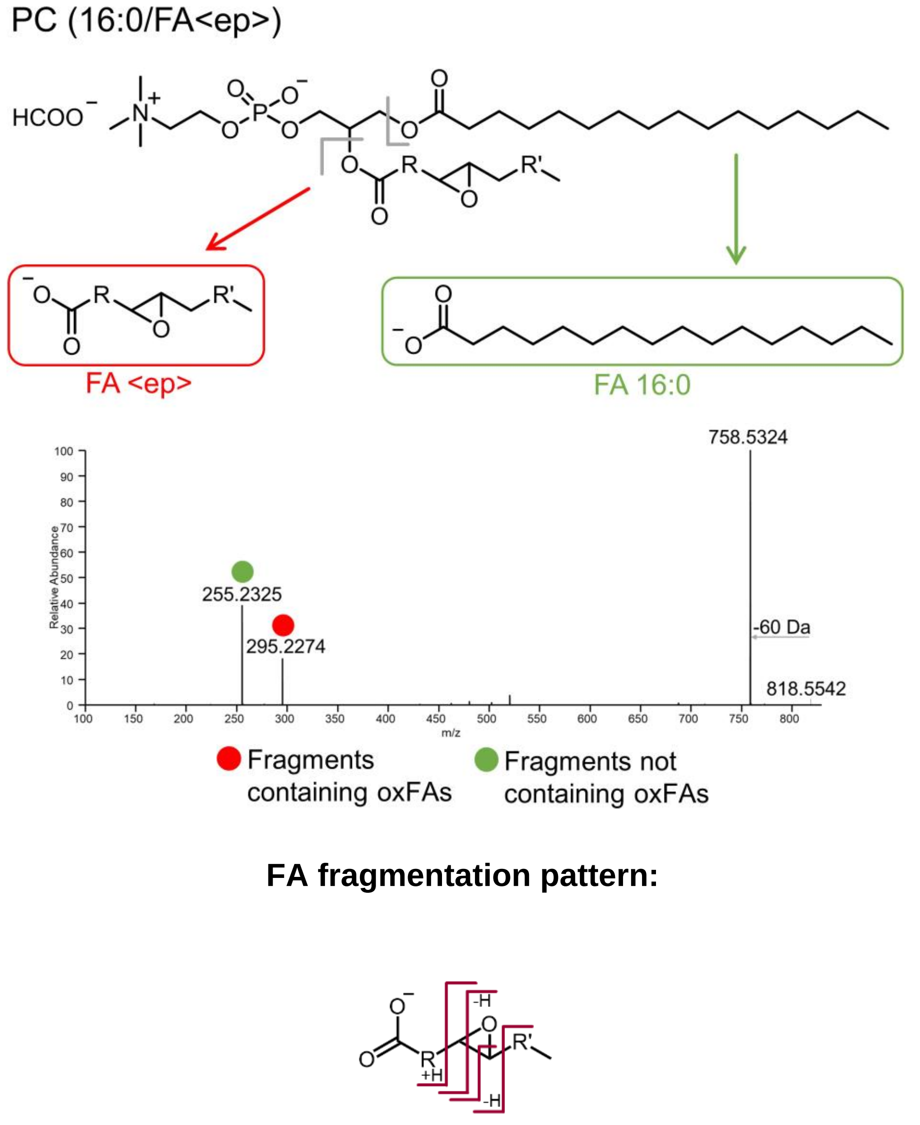 | **FA(18:0<ep>)**<br>$H_2O$ loss | **$C_{18}H_{33}O_3$**<br>**297.2435**<br>$C_{18}H_{31}O_2$<br>279.2330 | 9-10<br>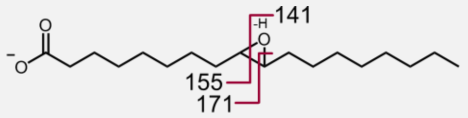 | $C_9H_{17}O$<br>$C_9H_{15}O_2$<br>**$C_9H_{15}O_3$** | 141.1285<br>155.1078<br>**171.1027** |
| **PC(16:0/18:1<ep>)** | $C_{43}H_{81}O_{11}NP$<br>818.5553 | | **FA(18:1<ep>)**<br>$H_2O$ loss | **$C_{18}H_{31}O_3$**<br>**295.2279**<br>$C_{18}H_{29}O_2$<br>277.2173 | 9-10<br>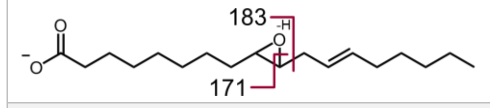 | **$C_9H_{15}O_3$**<br>$C_{10}H_{15}O_3$ | **171.1027**<br>183.1027 |
| | | | | | 12-13<br>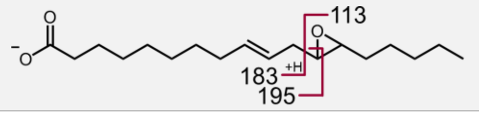 | $C_7H_{13}O$<br>**$C_{11}H_{19}O_2$**<br>**$C_{12}H_{19}O_2$** | 113.0972<br>**183.1391**<br>**195.1391** |
| **PC(16:0/20:3<ep>)** | $C_{45}H_{81}O_{11}NP$<br>842.5553 | | **FA(20:3<ep>)**<br>$H_2O$ loss<br>$H_2O$&$CO_2$ loss | **$C_{20}H_{31}O_3$**<br>**319.2279**<br>$C_{20}H_{29}O_2$<br>301.2173<br>$C_{19}H_{29}$<br>257.2275 | 5-6<br>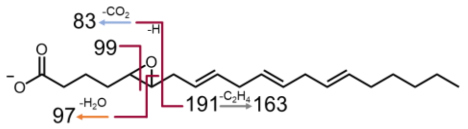 | $C_5H_7O$<br>$C_5H_5O_2$<br>$C_5H_7O_2$<br>**$C_{12}H_{19}$**<br>**$C_{14}H_{23}$** | 83.0502<br>97.0295<br>99.0452<br>**163.1492**<br>**191.1805** |
| | | | | | 8-9<br>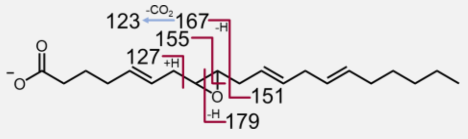 | **$C_7H_{11}O_2$**<br>**$C_8H_{11}O$**<br>**$C_{11}H_{19}$**<br>**$C_8H_{11}O_3$**<br>**$C_9H_{11}O_3$**<br>**$C_{12}H_{19}O$** | **127.0765**[7]<br>**123.0815**<br>**151.1492**<br>**155.0714**<br>**167.0714**<br>**179.1441** |
| | | | | | 11-12<br>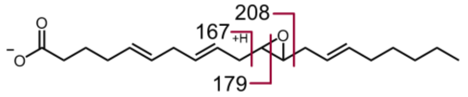 | **$C_{10}H_{15}O_2$**<br>**$C_{11}H_{15}O_2$**<br>$C_{12}H_{15}O_3$ | **167.1078**<br>**179.1078**<br>208.1104[8] |
| | | | | | 14-15<br>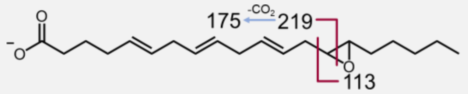 | $C_7H_{13}O$<br>**$C_{13}H_{19}$**<br>**$C_{14}H_{19}O_2$** | 113.0972<br>**175.1492**<br>**219.1391** |

[7] The abundances of fragments do not differ much, so it is difficult to choose the major one.
[8] The signal should correspond to an anion-radical that is the unique case for position 12.

#### Full-length oxygenated: <OOH>

$[M+HCOO]^-$ adducts

| Precursor | Chemical Formula & *m/z* | MS$^2$ Fragmentation | Mod.-specific fragment | Chemical Formula & *m/z* | Pos.-specific fragmentation |
|---|---|---|---|---|---|
| **PC(16:0/18:1<OOH>)** | $C_{43}H_{83}O_{12}NP$<br>836.5658 | 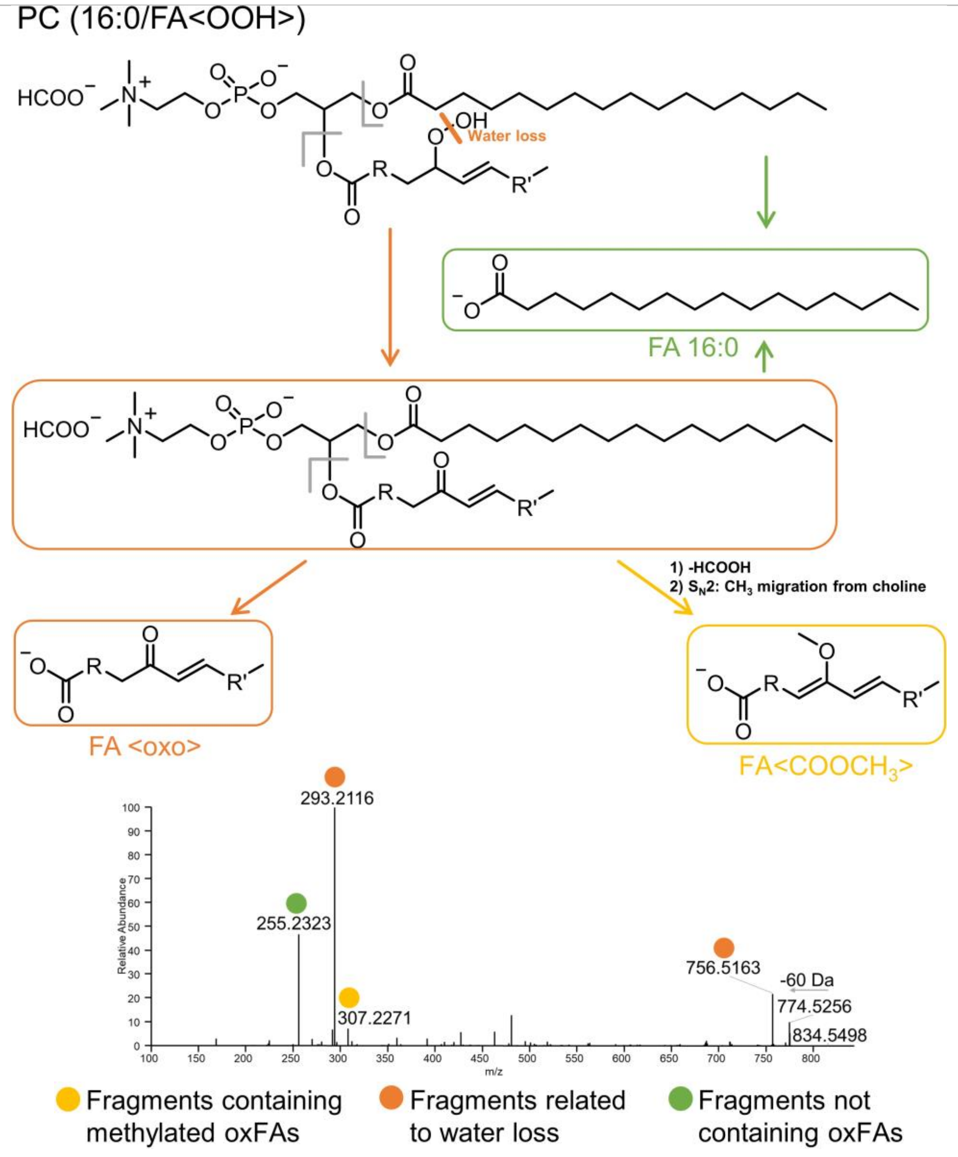 | FA(18:1<OOH>)<br>**$H_2O$ loss**<br>Methylation<br>$H_2O$&$CO_2$ loss<br>$2H_2O$ loss<br>$H_2O_2$ loss | $C_{18}H_{33}O_4$<br>313.2384<br>**$C_{18}H_{31}O_3$**<br>**295.2279**<br>$C_{19}H_{33}O_3$<br>309.2435<br>$C_{17}H_{31}O$<br>251.2380<br>$C_{18}H_{29}O_2$<br>277.2173<br>$C_{18}H_{27}O_2$<br>275.2016 | |
| **PC(16:0/18:2<OOH>)** | $C_{43}H_{81}O_{12}NP$<br>834.5502 | | FA(18:2<OOH>)<br>**$H_2O$ loss**<br>Methylation<br>$H_2O$&$CO_2$ loss<br>$2H_2O$ loss<br>$H_2O_2$ loss | $C_{18}H_{31}O_4$<br>311.2228<br>**$C_{18}H_{29}O_3$**<br>**293.2122**<br>$C_{19}H_{31}O_3$<br>307.2279<br>$C_{17}H_{29}O$<br>249.2224<br>$C_{18}H_{27}O_2$<br>275.2016<br>$C_{18}H_{29}O_2$<br>277.2173 | See above (Full-length oxygenated: <oxo>) |

#### Full-length oxygenated: <OOH> (continued)

$[M+HCOO]^-$ adducts

| Precursor | Chemical Formula & *m/z* | MS² Fragmentation | Mod.-specific fragment | Chemical Formula & *m/z* | Pos.-specific fragmentation |
|---|---|---|---|---|---|
| **PC(16:0/20:4<OOH>)** | $C_{45}H_{81}O_{12}NP$<br>858.5502 | 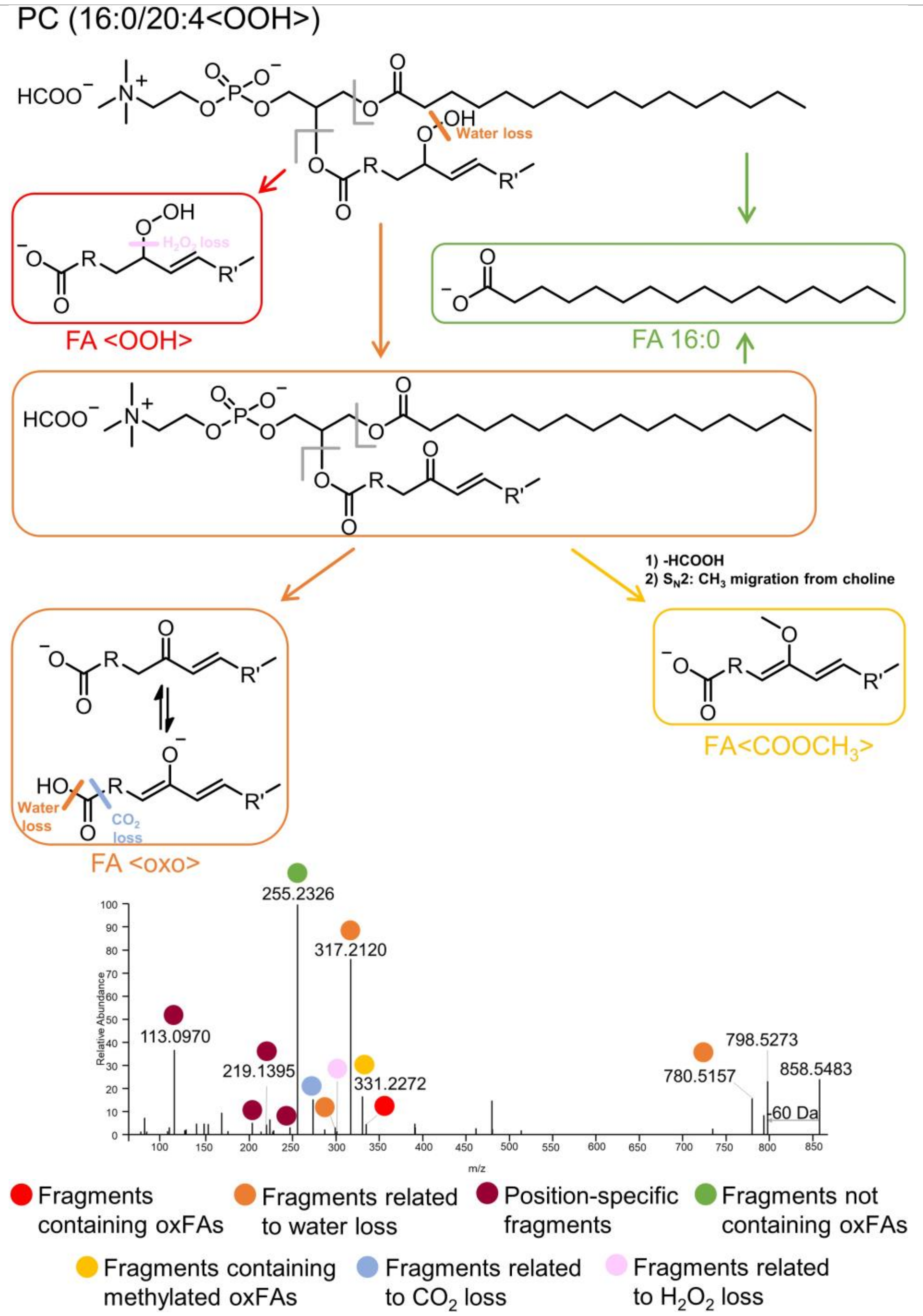 | FA(20:4<OOH>)<br><br>**$H_2O$ loss**<br><br>Methylation<br><br>$H_2O$&$CO_2$ loss<br><br>$2H_2O$ loss<br><br>$H_2O_2$ loss | $C_{20}H_{31}O_4$<br>335.2228<br>**$C_{20}H_{29}O_3$**<br>**317.2122**<br>$C_{21}H_{31}O_3$<br>331.2279<br>$C_{19}H_{29}O$<br>273.2224<br>$C_{20}H_{27}O_2$<br>299.2017<br>$C_{20}H_{29}O_2$<br>301.2173 | See above (Full-length oxygenated: <oxo>) |

### CHOLESTERYL ESTERS

#### Oxidatively truncated: <COOH>

$[M+Na]^+$ adducts

| Mod. position | Precursor | Chemical Formula | *m/z* | MS² Fragmentation | Mod./pos.-specific fragment | Chemical Formula | *m/z* |
|---|---|---|---|---|---|---|---|
| **4** | CE(4:0<COOH>) | $C_{31}H_{50}O_4Na$ | 509.3601 | | FA(4:0<COOH>) | $C_4H_6O_4Na$ | 141.0158 |
| **5** | CE(5:0<COOH>) | $C_{32}H_{52}O_4Na$ | 523.3758 | | FA(5:0<COOH>) | $C_5H_8O_4Na$ | 155.0315 |
| **7** | CE(7:0<COOH>) | $C_{34}H_{56}O_4Na$ | 551.4071 | | FA(7:0<COOH>) | $C_7H_{12}O_4Na$ | 183.0628 |
| **8** | CE(8:0<COOH>) | $C_{35}H_{58}O_4Na$ | 565.4227 | | FA(8:0<COOH>) | $C_8H_{14}O_4Na$ | 197.0784 |
| **9** | CE(9:0<COOH>) | $C_{36}H_{60}O_4Na$ | 579.4384 | | FA(9:0<COOH>) | $C_9H_{16}O_4Na$ | 211.0941 |
| **10** | CE(10:0<COOH>) | $C_{37}H_{62}O_4Na$ | 593.4540 | | FA(10:0<COOH>) | $C_{10}H_{18}O_4Na$ | 225.1097 |
| | CE(10:1<COOH>) | $C_{37}H_{60}O_4Na$ | 591.4384 | | FA(10:1<COOH>) | $C_{10}H_{16}O_4Na$ | 223.0941 |
| **11** | CE(11:1<COOH>) | $C_{38}H_{62}O_4Na$ | 605.4540 | | FA(11:1<COOH>) | $C_{11}H_{18}O_4Na$ | 237.1097 |
| | CE(11:2<COOH>) | $C_{38}H_{60}O_4Na$ | 603.4384 | | FA(11:2<COOH>) | $C_{11}H_{16}O_4Na$ | 235.0941 |
| **12** | CE(12:1<COOH>) | $C_{39}H_{64}O_4Na$ | 619.4697 | | FA(12:1<COOH>) | $C_{12}H_{20}O_4Na$ | 251.1254 |
| | CE(12:2<COOH>) | $C_{39}H_{62}O_4Na$ | 617.4540 | | FA(12:2<COOH>) | $C_{12}H_{18}O_4Na$ | 249.1097 |
| **13** | CE(13:2<COOH>) | $C_{40}H_{64}O_4Na$ | 631.4697 | | FA(13:2<COOH>) | $C_{13}H_{20}O_4Na$ | 263.1254 |
| | CE(13:3<COOH>) | $C_{40}H_{62}O_4Na$ | 629.4540 | | FA(13:3<COOH>) | $C_{13}H_{18}O_4Na$ | 261.1097 |
| **14** | CE(14:3<COOH>) | $C_{41}H_{64}O_4Na$ | 643.4697 | | FA(14:3<COOH>) | $C_{14}H_{20}O_4Na$ | 275.1254 |

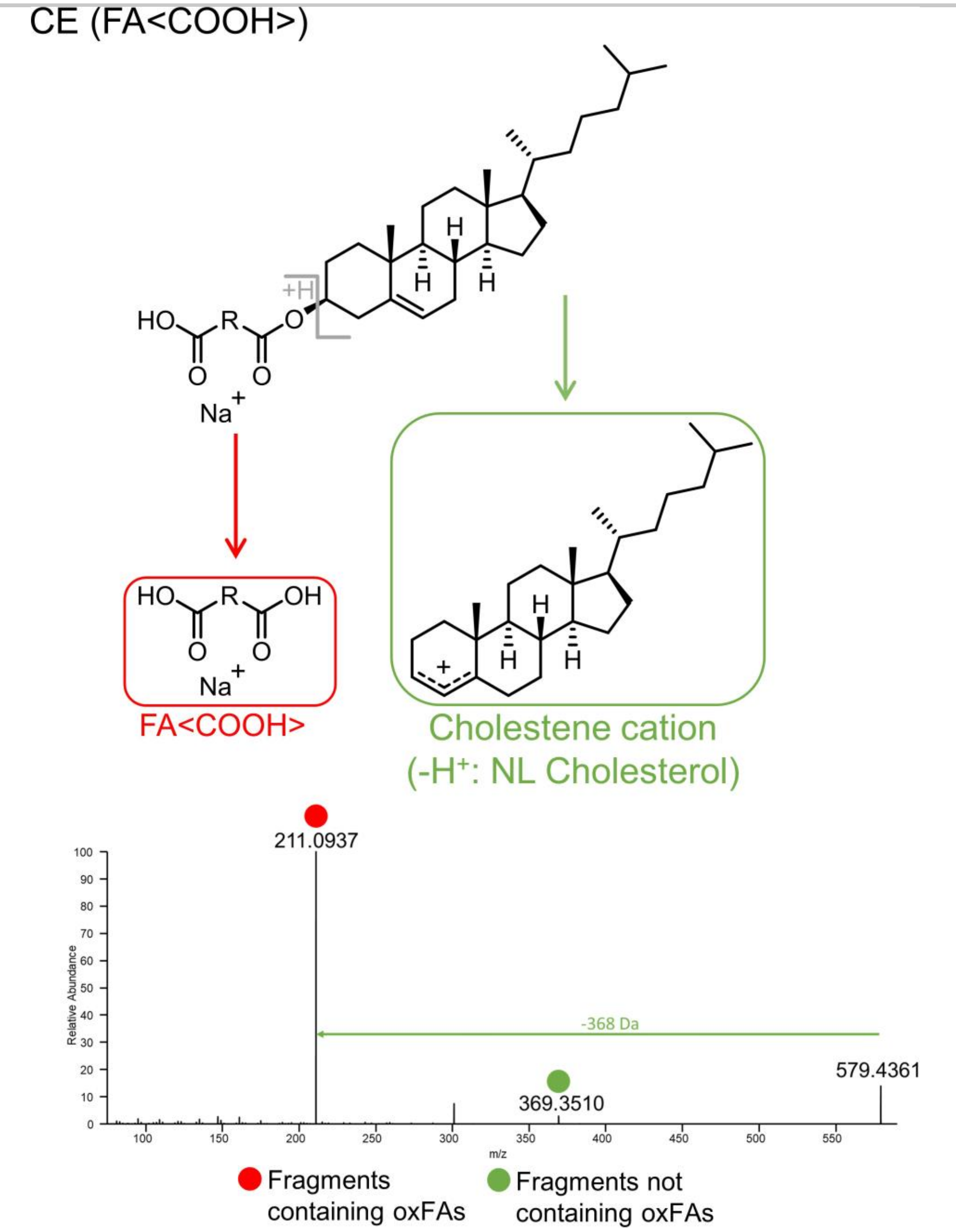

#### Oxidatively truncated: <oxo>

$[M+Na]^+$ adducts

| Mod. position | Precursor | Chemical Formula | *m/z* | MS² Fragmentation | Mod./pos.-specific fragment | Chemical Formula | *m/z* |
|---|---|---|---|---|---|---|---|
| **4** | CE(4:0<oxo>) | $C_{31}H_{50}O_3Na$ | 493.3652 | | FA(4:0<oxo>) | $C_4H_6O_3Na$ | 125.0209 |
| **5** | CE(5:0<oxo>) | $C_{32}H_{52}O_3Na$ | 507.3809 | | FA(5:0<oxo>) | $C_5H_8O_3Na$ | 139.0366 |
| **7** | CE(7:0<oxo>) | $C_{34}H_{56}O_3Na$ | 535.4122 | | FA(7:0<oxo>) | $C_7H_{12}O_3Na$ | 167.0679 |
| **8** | CE(8:0<oxo>) | $C_{35}H_{58}O_3Na$ | 549.4278 | | FA(8:0<oxo>) | $C_8H_{14}O_3Na$ | 181.0835 |
| **9** | CE(9:0<oxo>) | $C_{36}H_{60}O_3Na$ | 563.4435 | | FA(9:0<oxo>) | $C_9H_{16}O_3Na$ | 195.0992 |
| **10** | CE(10:0<oxo>) | $C_{37}H_{62}O_3Na$ | 577.4591 | | FA(10:0<oxo>) | $C_{10}H_{18}O_3Na$ | 209.1148 |
| | CE(10:1<oxo>) | $C_{37}H_{60}O_3Na$ | 575.4435 | | FA(10:1<oxo>) | $C_{10}H_{16}O_3Na$ | 207.0992 |
| **11** | CE(11:1<oxo>) | $C_{38}H_{62}O_3Na$ | 589.4591 | | FA(11:1<oxo>) | $C_{11}H_{18}O_3Na$ | 221.1148 |
| | CE(11:2<oxo>) | $C_{38}H_{60}O_3Na$ | 587.4435 | | FA(11:2<oxo>) | $C_{11}H_{16}O_3Na$ | 219.0992 |
| **12** | CE(12:1<oxo>) | $C_{39}H_{64}O_3Na$ | 603.4748 | | FA(12:1<oxo>) | $C_{12}H_{20}O_3Na$ | 235.1305 |
| | CE(12:2<oxo>) | $C_{39}H_{62}O_3Na$ | 601.4591 | | FA(12:2<oxo>) | $C_{12}H_{18}O_3Na$ | 233.1148 |
| **13** | CE(13:2<oxo>) | $C_{40}H_{64}O_3Na$ | 615.4748 | | FA(13:2<oxo>) | $C_{13}H_{20}O_3Na$ | 247.1305 |
| | CE(13:3<oxo>) | $C_{40}H_{62}O_3Na$ | 613.4591 | | FA(13:3<oxo>) | $C_{13}H_{18}O_3Na$ | 245.1148 |
| **14** | CE(14:3<oxo>) | $C_{41}H_{64}O_3Na$ | 627.4748 | | FA(14:3<oxo>) | $C_{14}H_{20}O_3Na$ | 259.1305 |

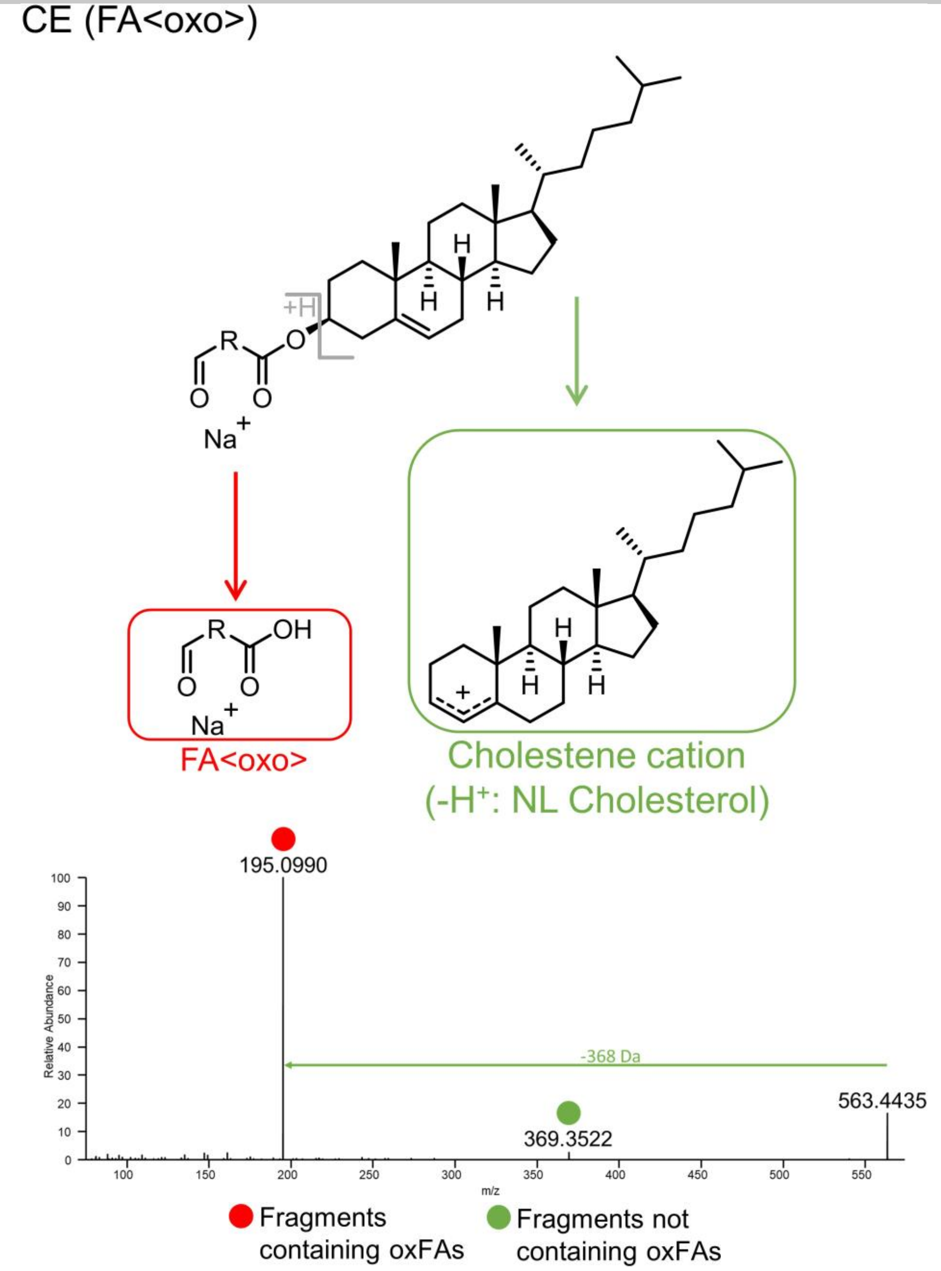

**Full-length oxygenated: <oxo>**

$[M+Na]^+$ adducts

| Precursor | Chemical Formula | *m/z* | MS$^2$ Fragmentation | Mod.-specific fragment | Chemical Formula | *m/z* |
|---|---|---|---|---|---|---|
| **CE(18:1)<oxo>** | $C_{45}H_{76}O_3Na$ | 687.5687 | 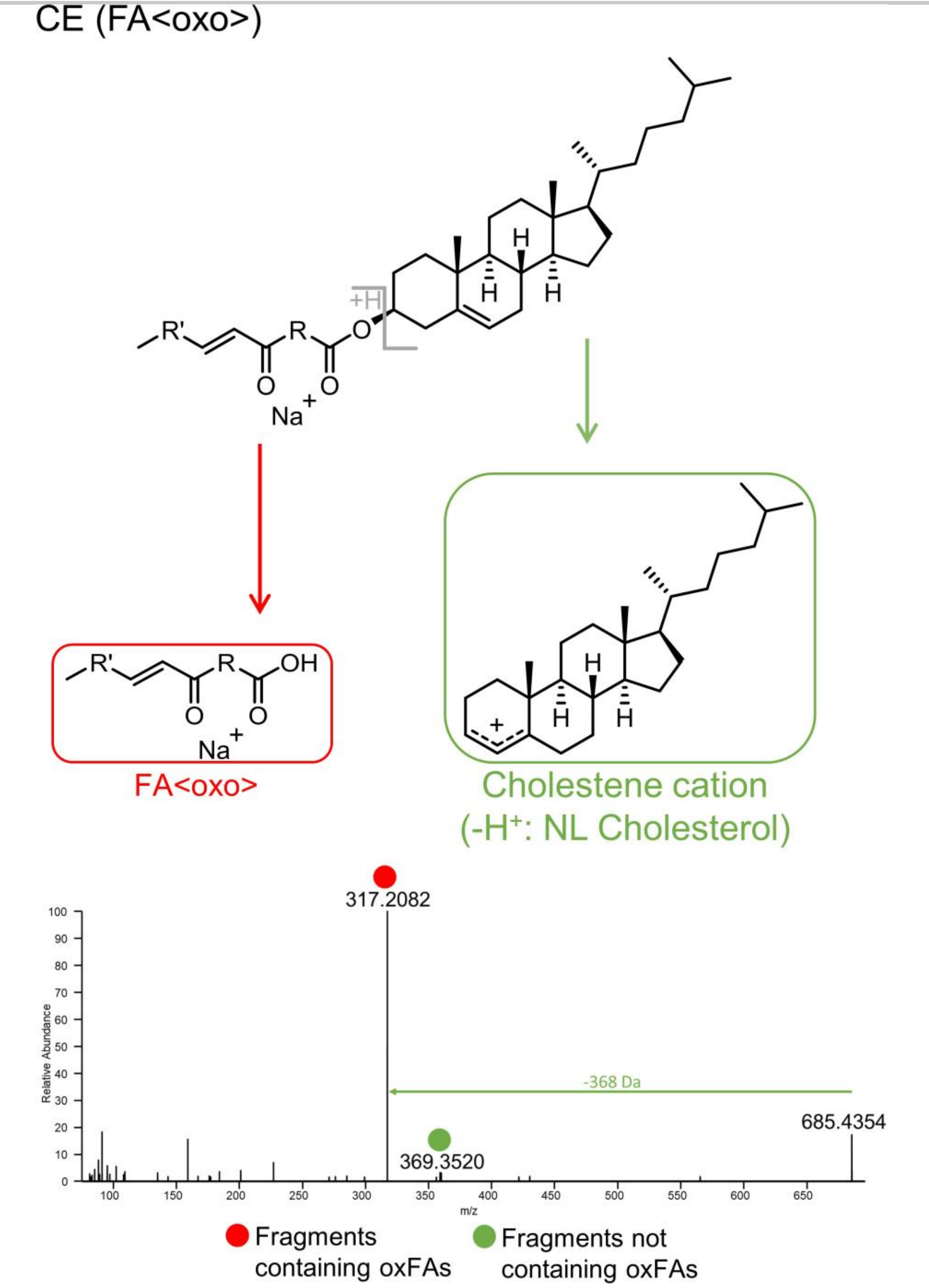  | FA(18:1)<oxo> | $C_{18}H_{32}O_3Na$ | 319.2244 |
| **CE(18:2)<oxo>** | $C_{45}H_{74}O_3Na$ | 685.5530 | | FA(18:2)<oxo> | $C_{18}H_{30}O_3Na$ | 317.2087 |
| **CE(20:4)<oxo>** | $C_{47}H_{74}O_3Na$ | 709.5530 | | FA(20:4)<oxo> | $C_{20}H_{30}O_3Na$ | 341.2087 |

**Full-length oxygenated: <OH>**

$[M+Na]^+$ adducts

| Precursor | Chemical Formula | *m/z* | $MS^2$ Fragmentation | Mod.-specific fragment | Chemical Formula | *m/z* |
|---|---|---|---|---|---|---|
| **CE(18:1<OH>)** | $C_{45}H_{78}O_3Na$ | 689.5843 | 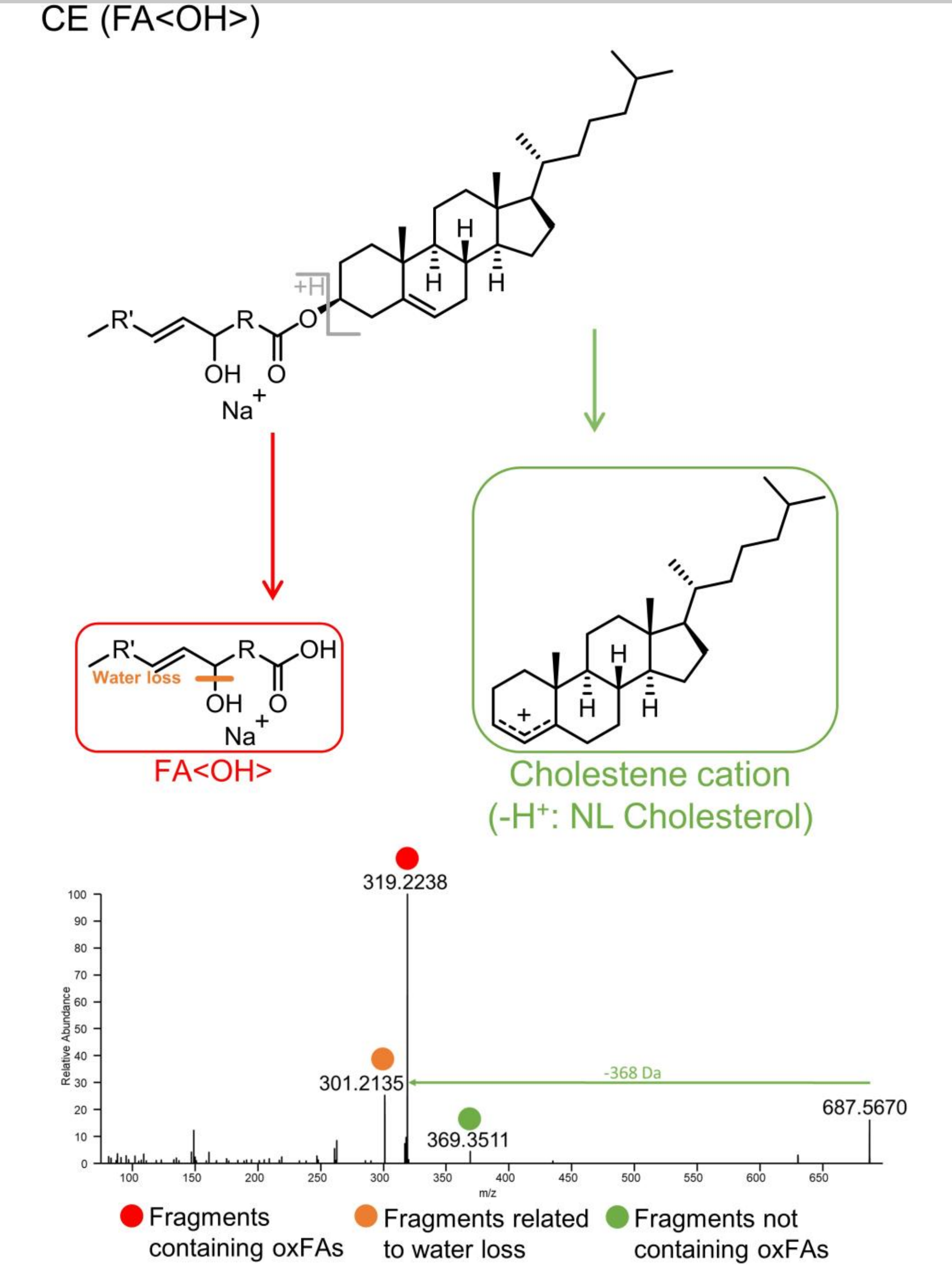 | **FA(18:1<OH>)**<br>H2O loss | $\mathbf{C_{18}H_{34}O_3Na}$<br>$C_{18}H_{32}O_2Na$ | **321.2400**<br>303.2295 |
| **CE(18:2<OH>)** | $C_{45}H_{76}O_3Na$ | 687.5687 | | **FA(18:2<OH>)**<br>H2O loss | $\mathbf{C_{18}H_{32}O_3Na}$<br>$C_{18}H_{30}O_2Na$ | **319.2244**<br>301.2138 |
| **CE(20:4<OH>)** | $C_{47}H_{76}O_3Na$ | 711.5687 | | **FA(20:4<OH>)**<br>H2O loss | $\mathbf{C_{20}H_{32}O_3Na}$<br>$C_{20}H_{30}O_2Na$ | **343.2244**<br>325.2138 |

**Full-length oxygenated: <ep>**

$[M+Na]^+$ adducts

| Precursor | Chemical Formula | *m/z* | $MS^2$ Fragmentation | Mod.-specific fragment | Chemical Formula | *m/z* |
|---|---|---|---|---|---|---|
| **CE(18:0<ep>)** | $C_{45}H_{78}O_3Na$ | 689.5843 | 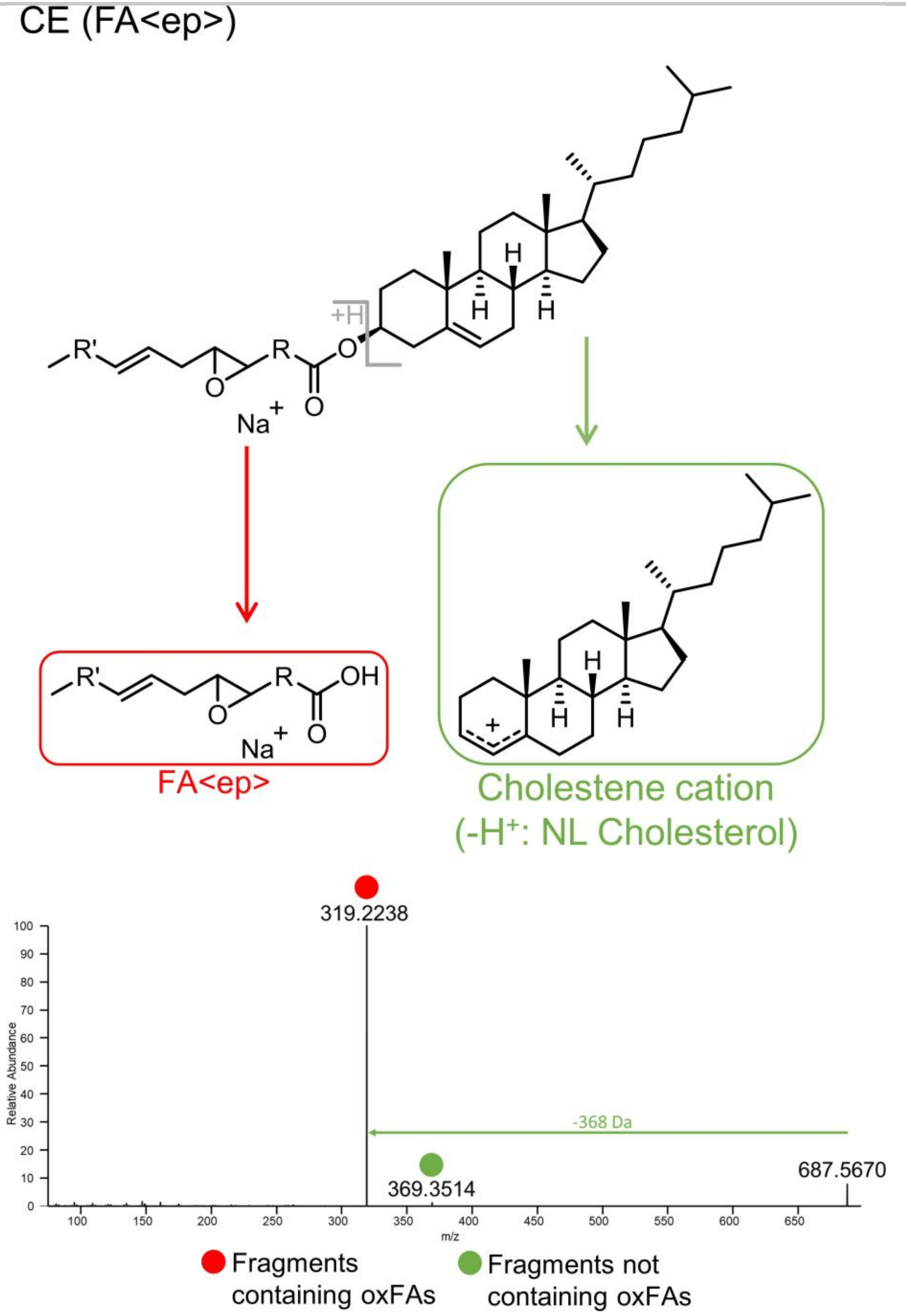 | FA(18:0<ep>) | $C_{18}H_{34}O_3Na$ | 321.2400 |
| **CE(18:1<ep>)** | $C_{45}H_{76}O_3Na$ | 687.5687 | | FA(18:1<ep>) | $C_{18}H_{32}O_3Na$ | 319.2244 |
| **CE(20:3<ep>)** | $C_{47}H_{76}O_3Na$ | 711.5687 | | FA(20:3<ep>) | $C_{20}H_{32}O_3Na$ | 343.2244 |

**Full-length oxygenated: <OOH>**

$[M+Na]^+$ adducts

| Precursor | Chemical Formula & *m/z* | MS² Fragmentation | Mod.-specific fragment | Chemical Formula & *m/z* | Pos.-specific fragment | Chemical Formula | *m/z* |
|---|---|---|---|---|---|---|---|
| **CE(18:1<OOH>)** | $C_{45}H_{78}O_4Na$ 705.5792 | 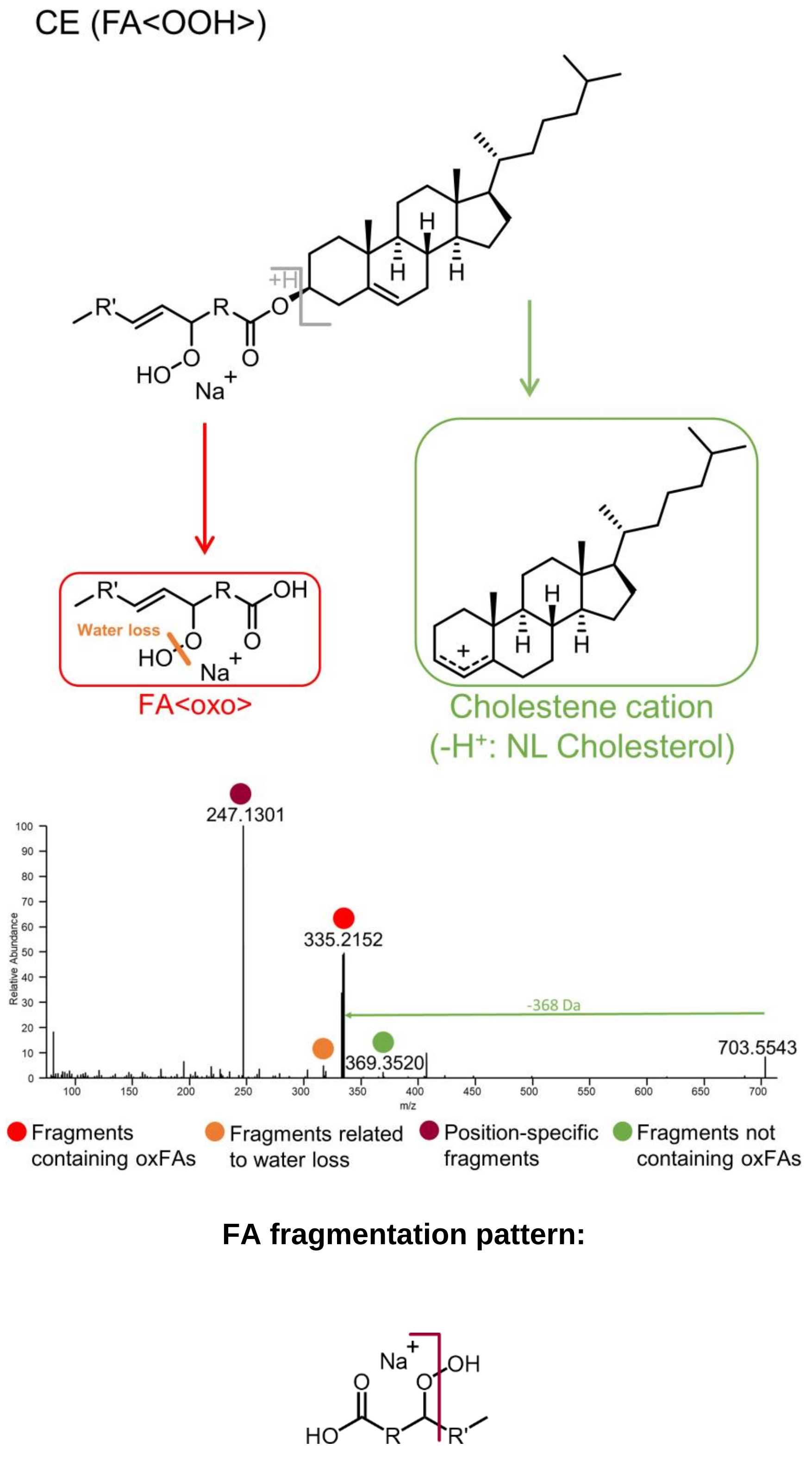 | **FA(18:1<OOH>)** H2O loss | **$C_{18}H_{34}O_4Na$ 337.2349** $C_{18}H_{32}O_3Na$ 319.2244 | 9 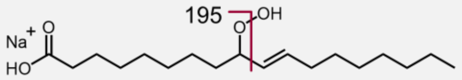 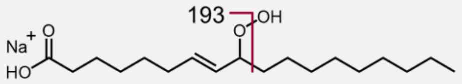 | $C_9H_{16}O_3Na$<br>$C_9H_{14}O_3Na$ | 195.0992<br>193.0835 |
| **CE(18:2<OOH>)** | $C_{45}H_{76}O_4Na$ 703.5636 | | **FA(18:2<OOH>)** H2O loss | **$C_{18}H_{32}O_4Na$ 335.2193** $C_{18}H_{30}O_3Na$ 317.2087 | 9 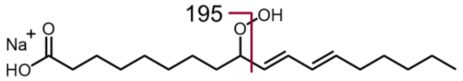 | $C_9H_{16}O_3Na$ | 195.0992 |
| | | | | | 10 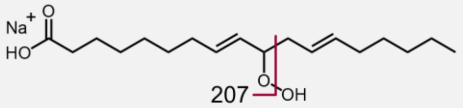 | $C_{10}H_{16}O_3Na$ | 207.0992 |
| | | | | | 11 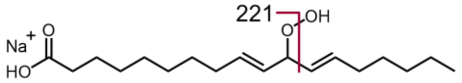 | $C_{11}H_{18}O_3Na$ | 221.1148 |
| | | | | | 12 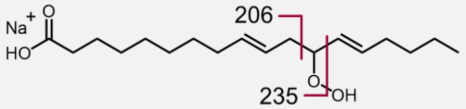 | $C_{11}H_{19}O_2Na$<br>$C_{12}H_{20}O_3Na$ | 206.1277[10]<br>235.1305 |
| | | | | | 13 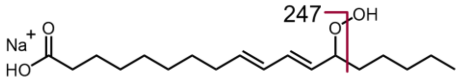 | $C_{13}H_{20}O_3Na$ | 247.1305 |
| **CE(20:4<OOH>)** | $C_{47}H_{76}O_4Na$ 727.5636 | | **FA(20:4<OOH>)** H2O loss | **$C_{20}H_{32}O_4Na$ 359.2193** $C_{20}H_{30}O_3Na$ 341.2087 | 5 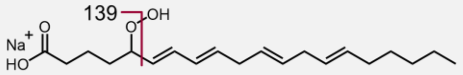 | $C_5H_8O_3Na$ | 139.0366 |
| | | | | | 6 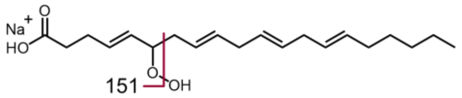 | $C_6H_8O_3Na$ | 151.0366 |
| | | | | | 8 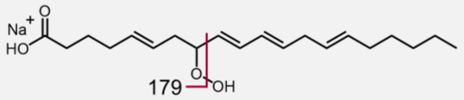 | $C_8H_{12}O_3Na$ | 179.0679 |
| | | | | | 9 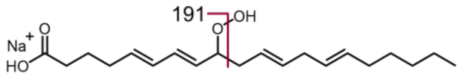 | $C_9H_{12}O_3Na$ | 191.0679 |

#### Full-length oxygenated: <OOH> (continued)

$[M+Na]^+$ adducts

| Precursor | Chemical Formula & *m/z* | MS² Fragmentation | Mod.-specific fragment | Chemical Formula & *m/z* | Pos.-specific fragment | Chemical Formula | *m/z* |
|---|---|---|---|---|---|---|---|
| **CE(20:4<OOH>)** | $C_{47}H_{76}O_4Na$ 727.5636 | | **FA(20:4<OOH>)** <br> $H_2O$ loss | **$C_{20}H_{32}O_4Na$ 359.2193** <br> $C_{20}H_{30}O_3Na$ 341.2087 | 11 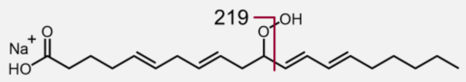  | $C_{11}H_{16}O_3Na$ | 219.0992 |
| | | | | | 12 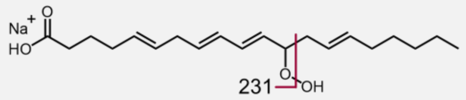  | $C_{12}H_{16}O_3Na$ | 231.0992 |
| | | | | | 14 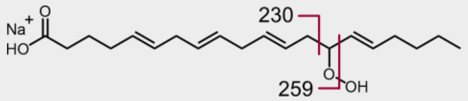  | $C_{13}H_{19}O_2Na$ <br> $C_{14}H_{20}O_3Na$ | 230.1277[9] <br> 259.1305 |
| | | | | | 15 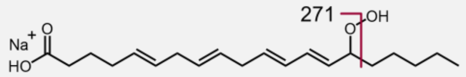  | $C_{15}H_{20}O_3Na$ | 271.1305 |

[9] See *Ito, J., Mizuochi, S., Nakagawa, K., Kato, S., Miyazawa, T., 2015. Tandem Mass Spectrometry Analysis of Linoleic and Arachidonic Acid Hydroperoxides via Promotion of Alkali Metal Adduct Formation. Anal. Chem. 87, 4980–4987.*

### TRIACYLGLYCEROLS

#### Oxidatively truncated: <COOH>

[M+Na]$^+$ adducts

| Mod. position | Precursor | Chemical Formula | *m/z* | MS$^2$ Fragmentation | Mod./pos.-specific fragment | Chemical Formula | *m/z* |
|---|---|---|---|---|---|---|---|
| **4** | TG(16:0/16:0/4:0<COOH>) | $C_{39}H_{72}O_8Na$ | 691.5119 | | FA(4:0<COOH>) | $C_4H_6O_4Na$ | 141.0158 |
| **5** | TG(16:0/16:0/5:0<COOH>) | $C_{40}H_{74}O_8Na$ | 705.5276 | | FA(5:0<COOH>) | $C_5H_8O_4Na$ | 155.0315 |
| **7** | TG(16:0/16:0/7:0<COOH>) | $C_{42}H_{78}O_8Na$ | 733.5589 | | FA(7:0<COOH>) | $C_7H_{12}O_4Na$ | 183.0628 |
| **8** | TG(16:0/16:0/8:0<COOH>) | $C_{43}H_{80}O_8Na$ | 747.5745 | | FA(8:0<COOH>) | $C_8H_{14}O_4Na$ | 197.0784 |
| **9** | TG(16:0/16:0/9:0<COOH>) | $C_{44}H_{82}O_8Na$ | 761.5902 | | FA(9:0<COOH>) | $C_9H_{16}O_4Na$ | 211.0941 |
| **10** | TG(16:0/16:0/10:0<COOH>) | $C_{45}H_{84}O_8Na$ | 775.6058 | | FA(10:0<COOH>) | $C_{10}H_{18}O_4Na$ | 225.1097 |
| | TG(16:0/16:0/10:1<COOH>) | $C_{45}H_{82}O_8Na$ | 773.5902 | | FA(10:1<COOH>) | $C_{10}H_{16}O_4Na$ | 223.0941 |
| **11** | TG(16:0/16:0/11:1<COOH>) | $C_{46}H_{84}O_8Na$ | 787.6058 | | FA(11:1<COOH>) | $C_{11}H_{18}O_4Na$ | 237.1097 |
| | TG(16:0/16:0/11:2<COOH>) | $C_{46}H_{82}O_8Na$ | 785.5902 | | FA(11:2<COOH>) | $C_{11}H_{16}O_4Na$ | 235.0941 |
| **12** | TG(16:0/16:0/12:1<COOH>) | $C_{47}H_{86}O_8Na$ | 801.6215 | | FA(12:1<COOH>) | $C_{12}H_{20}O_4Na$ | 251.1254 |
| | TG(16:0/16:0/12:2<COOH>) | $C_{47}H_{84}O_8Na$ | 799.6058 | | FA(12:2<COOH>) | $C_{12}H_{18}O_4Na$ | 249.1097 |
| **13** | TG(16:0/16:0/13:2<COOH>) | $C_{48}H_{86}O_8Na$ | 813.6215 | | FA(13:2<COOH>) | $C_{13}H_{20}O_4Na$ | 263.1254 |
| | TG(16:0/16:0/13:3<COOH>) | $C_{48}H_{84}O_8Na$ | 811.6058 | | FA(13:3<COOH>) | $C_{13}H_{18}O_4Na$ | 261.1097 |
| **14** | TG(16:0/16:0/14:3<COOH>) | $C_{49}H_{86}O_8Na$ | 825.6215 | | FA(14:3<COOH>) | $C_{14}H_{20}O_4Na$ | 275.1254 |

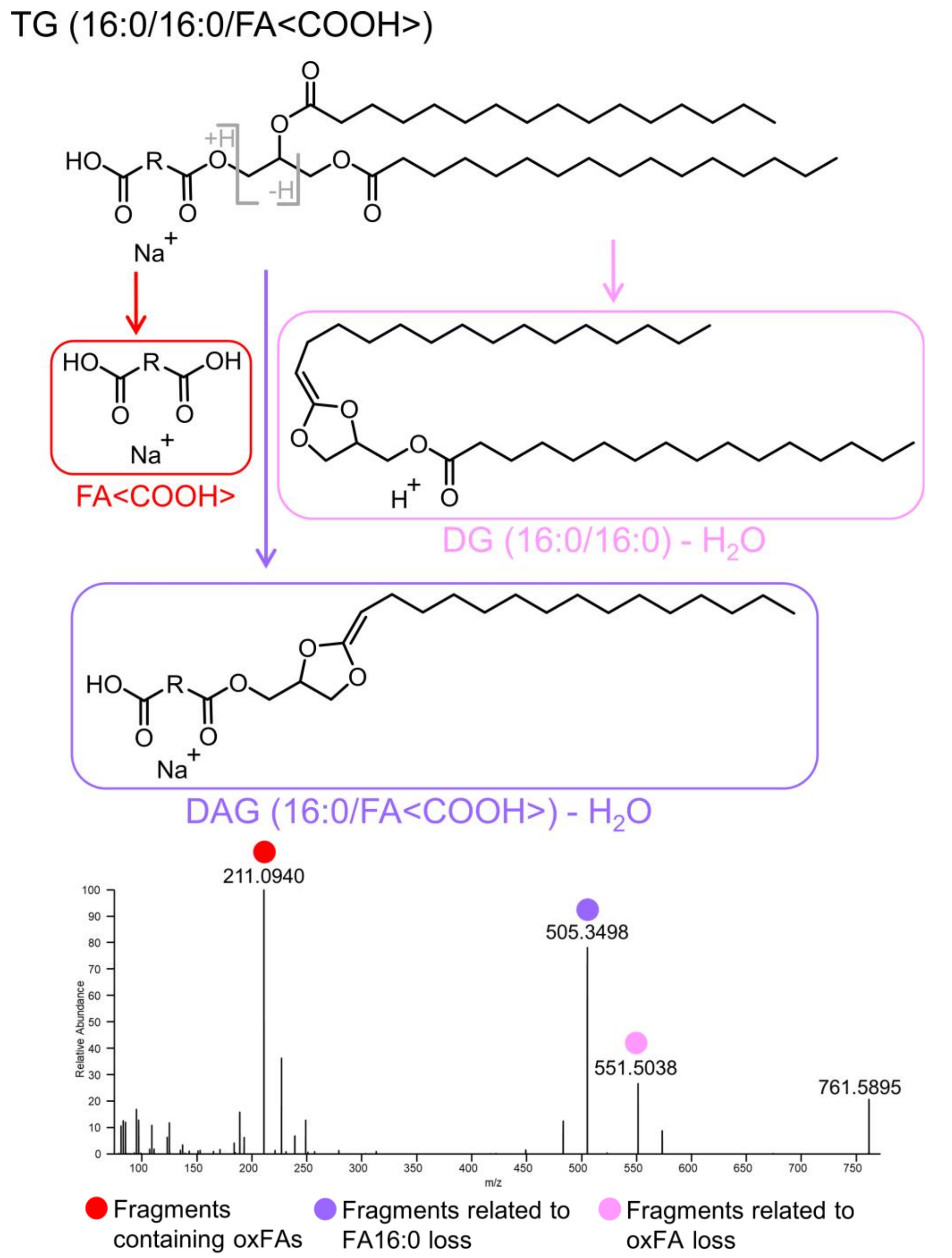

**Oxidatively truncated: <oxo>**

$[M+Na]^+$ adducts

| Mod. position | Precursor | Chemical Formula | *m/z* | MS² Fragmentation | Mod./pos.-specific fragment | Chemical Formula | *m/z* |
|---|---|---|---|---|---|---|---|
| **4** | TG(16:0/16:0/4:0<oxo>) | $C_{39}H_{72}O_7Na$ | 675.5170 | 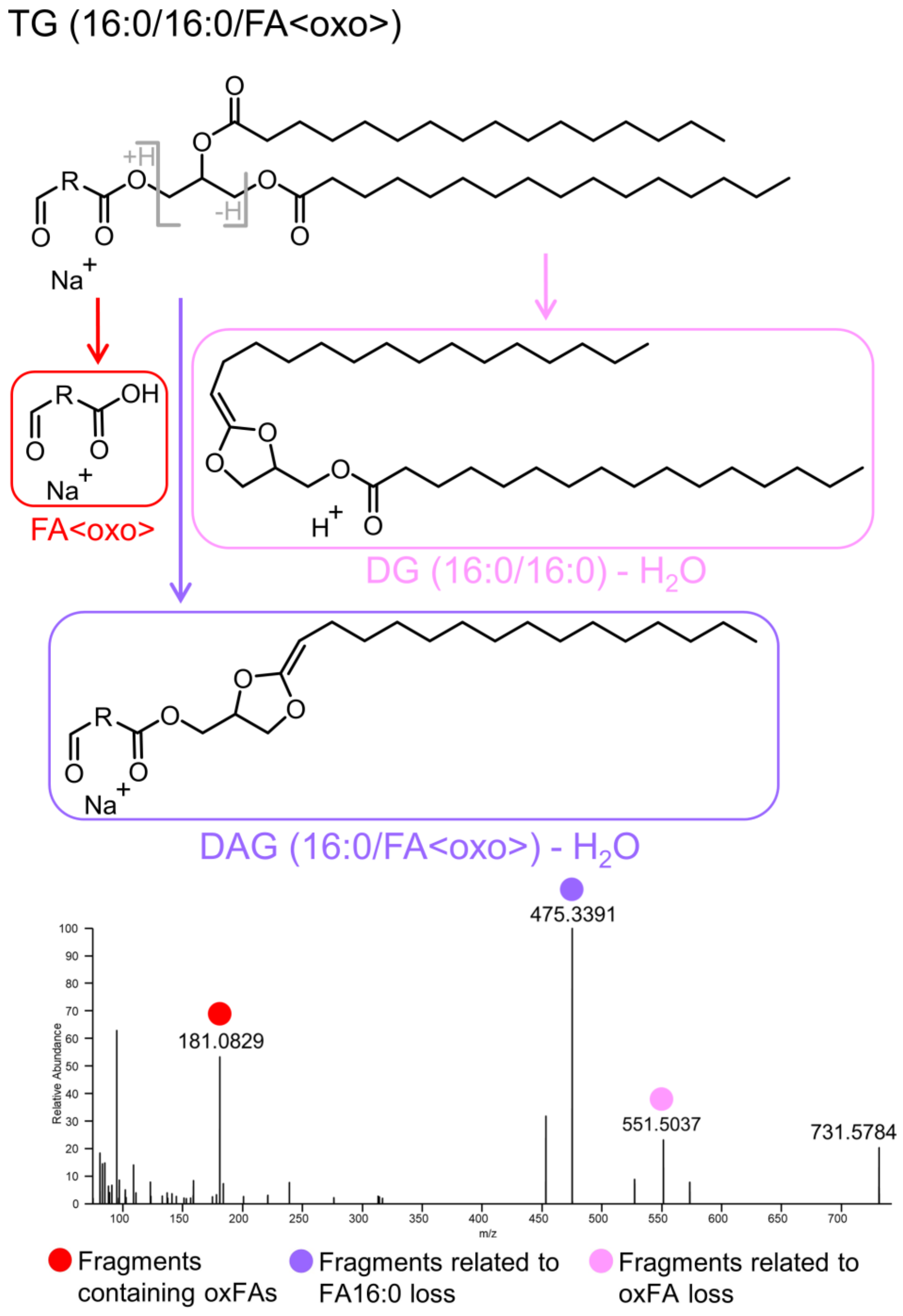  | FA(4:0<oxo>) | $C_4H_6O_3Na$ | 125.0209 |
| **5** | TG(16:0/16:0/5:0<oxo>) | $C_{40}H_{74}O_7Na$ | 689.5327 | | FA(5:0<oxo>) | $C_5H_8O_3Na$ | 139.0366 |
| **7** | TG(16:0/16:0/7:0<oxo>) | $C_{42}H_{78}O_7Na$ | 717.5640 | | FA(7:0<oxo>) | $C_7H_{12}O_3Na$ | 167.0679 |
| **8** | TG(16:0/16:0/8:0<oxo>) | $C_{43}H_{80}O_7Na$ | 731.5796 | | FA(8:0<oxo>) | $C_8H_{14}O_3Na$ | 181.0835 |
| **9** | TG(16:0/16:0/9:0<oxo>) | $C_{44}H_{82}O_7Na$ | 745.5953 | | FA(9:0<oxo>) | $C_9H_{16}O_3Na$ | 195.0992 |
| **10** | TG(16:0/16:0/10:0<oxo>) | $C_{45}H_{84}O_7Na$ | 759.6109 | | FA(10:0<oxo>) | $C_{10}H_{18}O_3Na$ | 209.1148 |
| | TG(16:0/16:0/10:1<oxo>) | $C_{45}H_{82}O_7Na$ | 757.5953 | | FA(10:1<oxo>) | $C_{10}H_{16}O_3Na$ | 207.0992 |
| **11** | TG(16:0/16:0/11:1<oxo>) | $C_{46}H_{84}O_7Na$ | 771.6109 | | FA(11:1<oxo>) | $C_{11}H_{18}O_3Na$ | 221.1148 |
| | TG(16:0/16:0/11:2<oxo>) | $C_{46}H_{82}O_7Na$ | 769.5953 | | FA(11:2<oxo>) | $C_{11}H_{16}O_3Na$ | 219.0992 |
| **12** | TG(16:0/16:0/12:1<oxo>) | $C_{47}H_{86}O_7Na$ | 785.6266 | | FA(12:1<oxo>) | $C_{12}H_{20}O_3Na$ | 235.1305 |
| | TG(16:0/16:0/12:2<oxo>) | $C_{47}H_{84}O_7Na$ | 783.6109 | | FA(12:2<oxo>) | $C_{12}H_{18}O_3Na$ | 233.1148 |
| **13** | TG(16:0/16:0/13:2<oxo>) | $C_{48}H_{86}O_7Na$ | 797.6266 | | FA(13:2<oxo>) | $C_{13}H_{20}O_3Na$ | 247.1305 |
| | TG(16:0/16:0/13:3<oxo>) | $C_{48}H_{84}O_7Na$ | 795.6109 | | FA(13:3<oxo>) | $C_{13}H_{18}O_3Na$ | 245.1148 |
| **14** | TG(16:0/16:0/14:3<oxo>) | $C_{49}H_{86}O_7Na$ | 809.6266 | | FA(14:3<oxo>) | $C_{14}H_{20}O_3Na$ | 259.1305 |

**Full-length oxygenated: <oxo>**

$[M+Na]^+$ adducts

| Precursor | Chemical Formula | *m/z* | MS² Fragmentation | Mod.-specific fragment | Chemical Formula | *m/z* |
|---|---|---|---|---|---|---|
| **TG(16:0/16:0/18:1<oxo>)** | $C_{53}H_{98}O_7Na$ | 869.7205 | (see figure below) | FA(18:1<oxo>) | $C_{18}H_{32}O_3Na$ | 319.2244 |
| **TG(16:0/16:0/18:2<oxo>)** | $C_{53}H_{96}O_7Na$ | 867.7048 | | FA(18:2<oxo>) | $C_{18}H_{30}O_3Na$ | 317.2087 |
| **TG(16:0/16:0/20:4<oxo>)** | $C_{55}H_{96}O_7Na$ | 891.7048 | | FA(20:4<oxo>) | $C_{20}H_{30}O_3Na$ | 341.2087 |

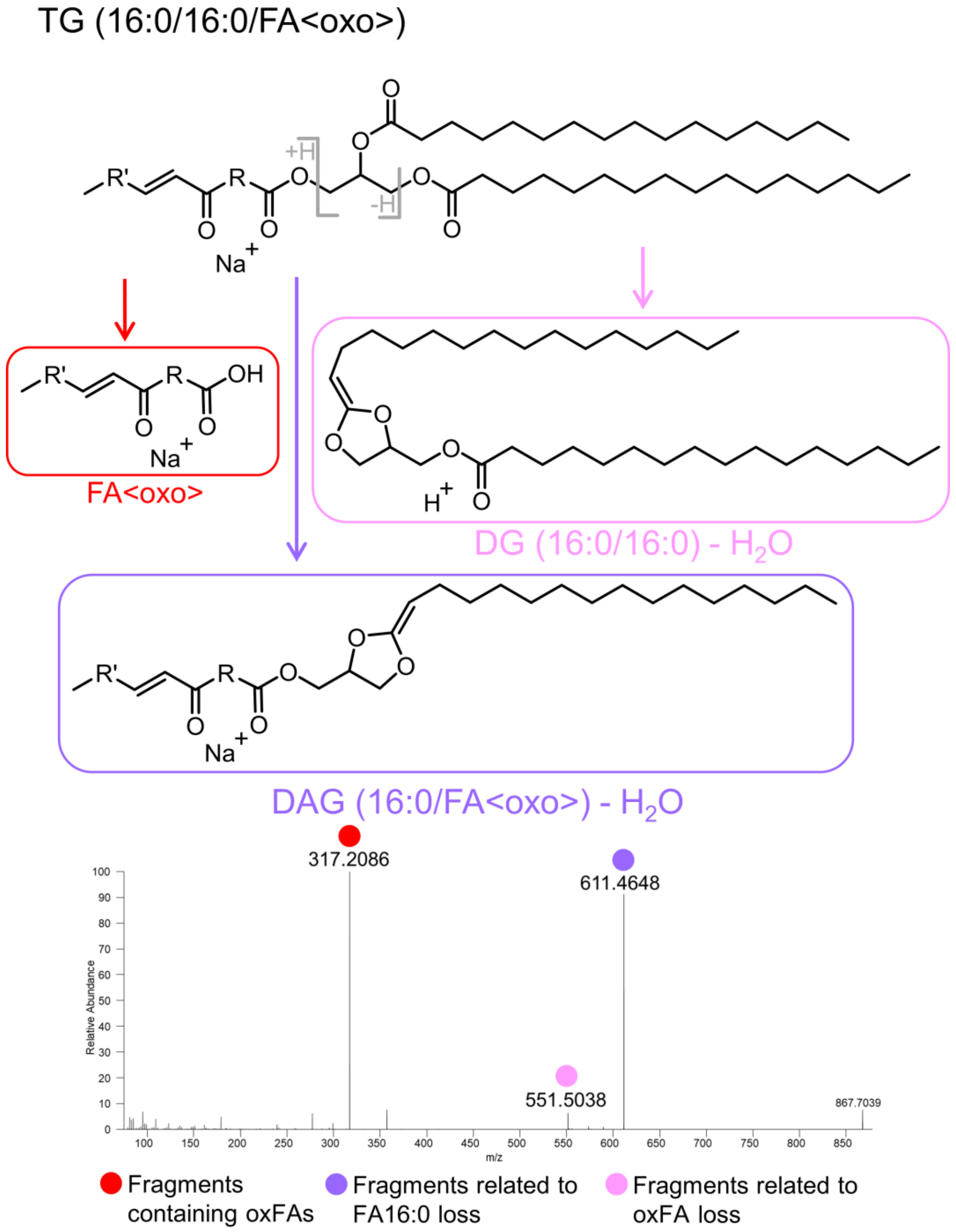

#### Full-length oxygenated: <OH>

[M+Na]+ adducts

| Precursor | Chemical Formula | *m/z* | MS² Fragmentation | Mod.-specific fragment | Chemical Formula | *m/z* |
|---|---|---|---|---|---|---|
| **TG(16:0/16:0/18:1<OH>)** | $C_{53}H_{100}O_7Na$ | 871.7361 | 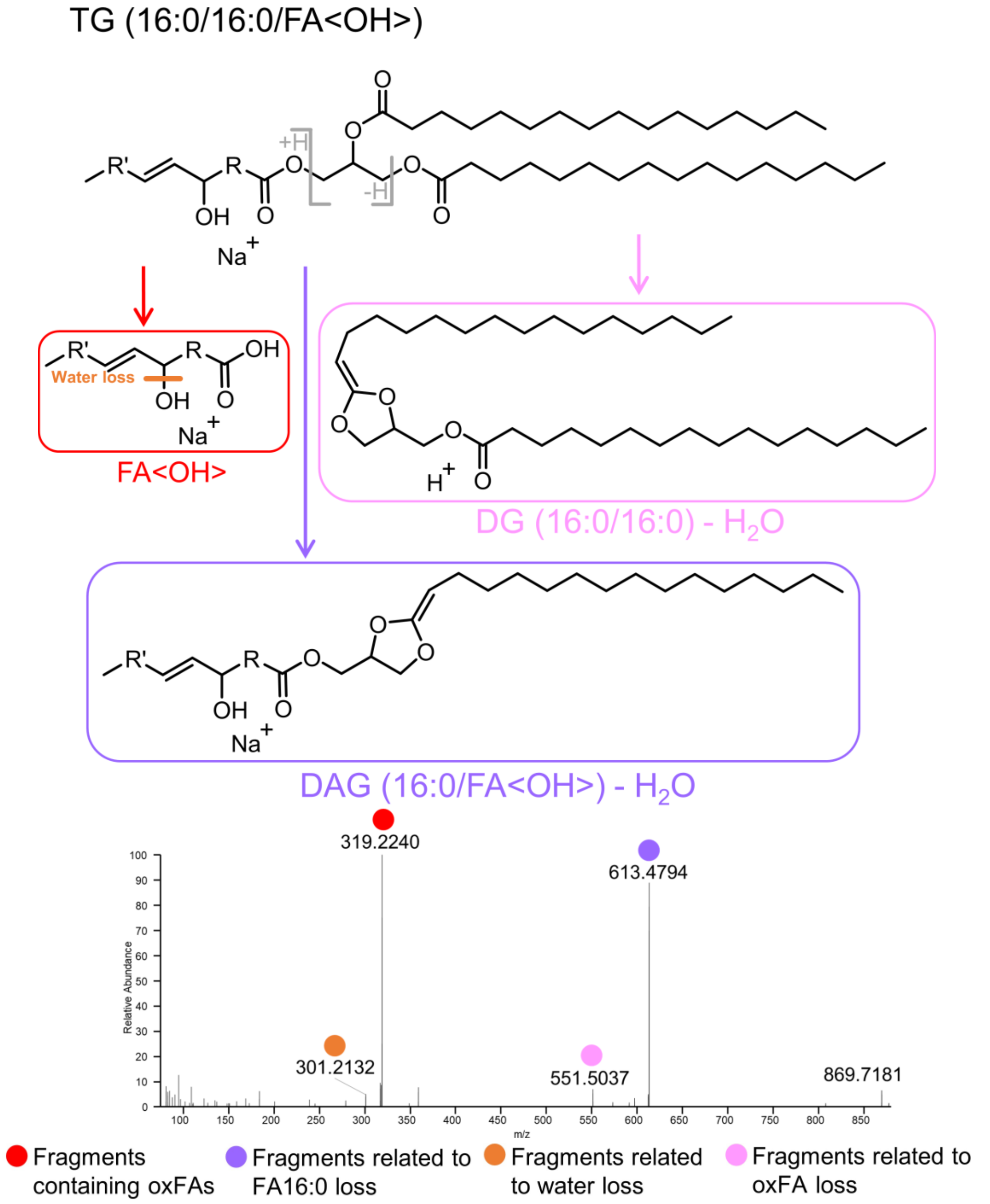 | **FA(18:1<OH>)**<br>$H_2O$ loss | **$C_{18}H_{34}O_3Na$**<br>$C_{18}H_{32}O_2Na$ | **321.2400**<br>303.2295 |
| **TG(16:0/16:0/18:2<OH>)** | $C_{53}H_{98}O_7Na$ | 869.7205 | | **FA(18:2<OH>)**<br>$H_2O$ loss | **$C_{18}H_{32}O_3Na$**<br>$C_{18}H_{30}O_2Na$ | **319.2244**<br>301.2138 |
| **TG(16:0/16:0/20:4<OH>)** | $C_{55}H_{98}O_7Na$ | 893.7205 | | **FA(20:4<OH>)**<br>$H_2O$ loss | **$C_{20}H_{32}O_3Na$**<br>$C_{20}H_{30}O_2Na$ | **343.2244**<br>325.2138 |

**Full-length oxygenated: <ep>**

$[M+Na]^+$ adducts

| Precursor | Chemical Formula | *m/z* | MS² Fragmentation | Mod.-specific fragment | Chemical Formula | *m/z* |
|---|---|---|---|---|---|---|
| **TG(16:0/16:0/18:0<ep>)** | $C_{53}H_{100}O_7Na$ | 871.7361 | 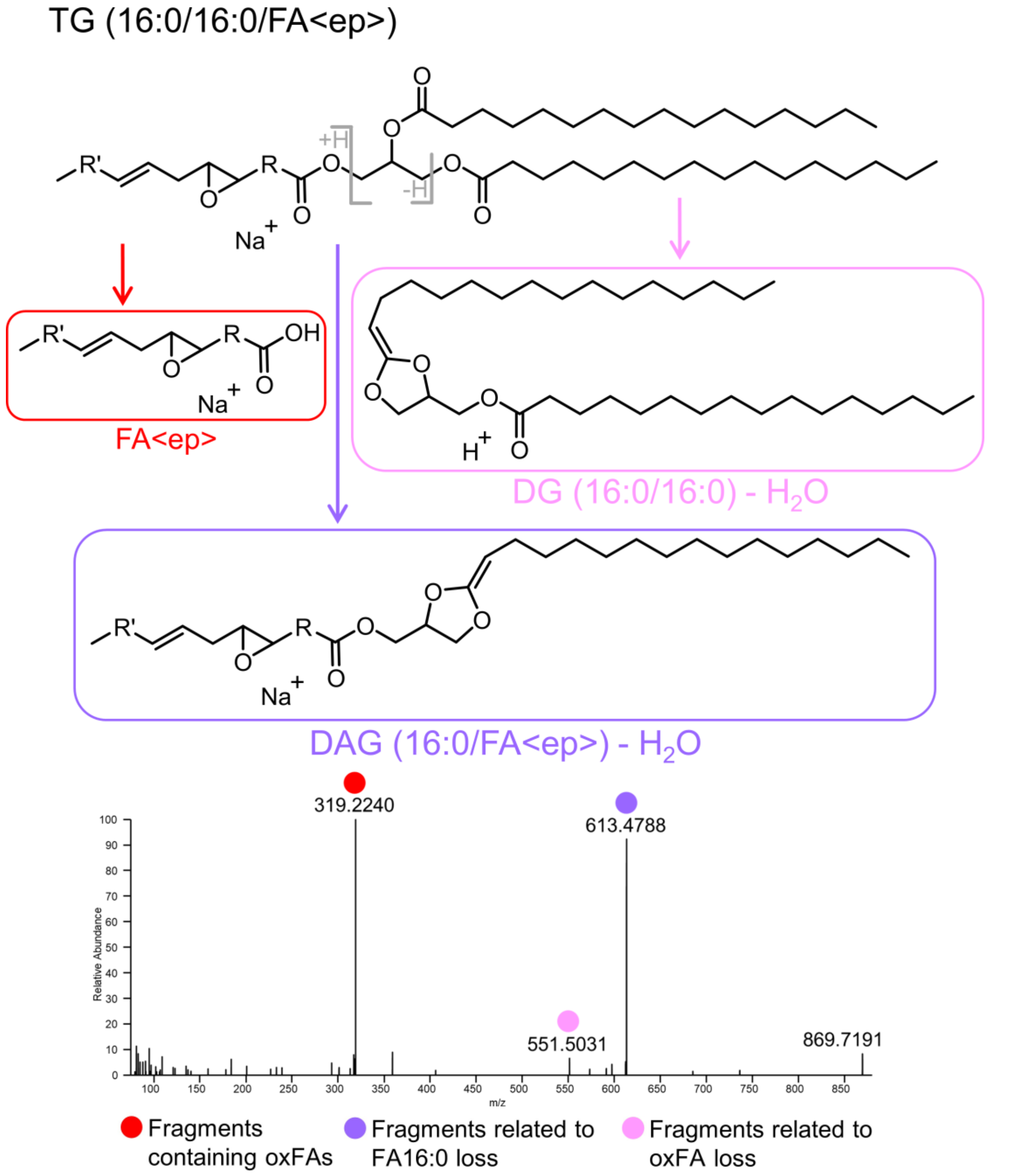 | FA(18:0<ep>) | $C_{18}H_{34}O_3Na$ | 321.2400 |
| **TG(16:0/16:0/18:1<ep>)** | $C_{53}H_{98}O_7Na$ | 869.7205 | | FA(18:1<ep>) | $C_{18}H_{32}O_3Na$ | 319.2244 |
| **TG(16:0/16:0/20:3<ep>)** | $C_{55}H_{98}O_7Na$ | 893.7205 | | FA(20:3<ep>) | $C_{20}H_{32}O_3Na$ | 343.2244 |

**Full-length oxygenated: <OOH>**

$[M+Na]^+$ adducts

| Precursor | Chemical Formula & *m/z* | MS$^2$ Fragmentation | Mod.-specific fragment | Chemical Formula & *m/z* | Pos.-specific fragment | Chemical Formula | *m/z* |
|---|---|---|---|---|---|---|---|
| **TG(16:0/16:0/18:1<OOH>)** | $C_{53}H_{100}O_8Na$ 887.7310 | 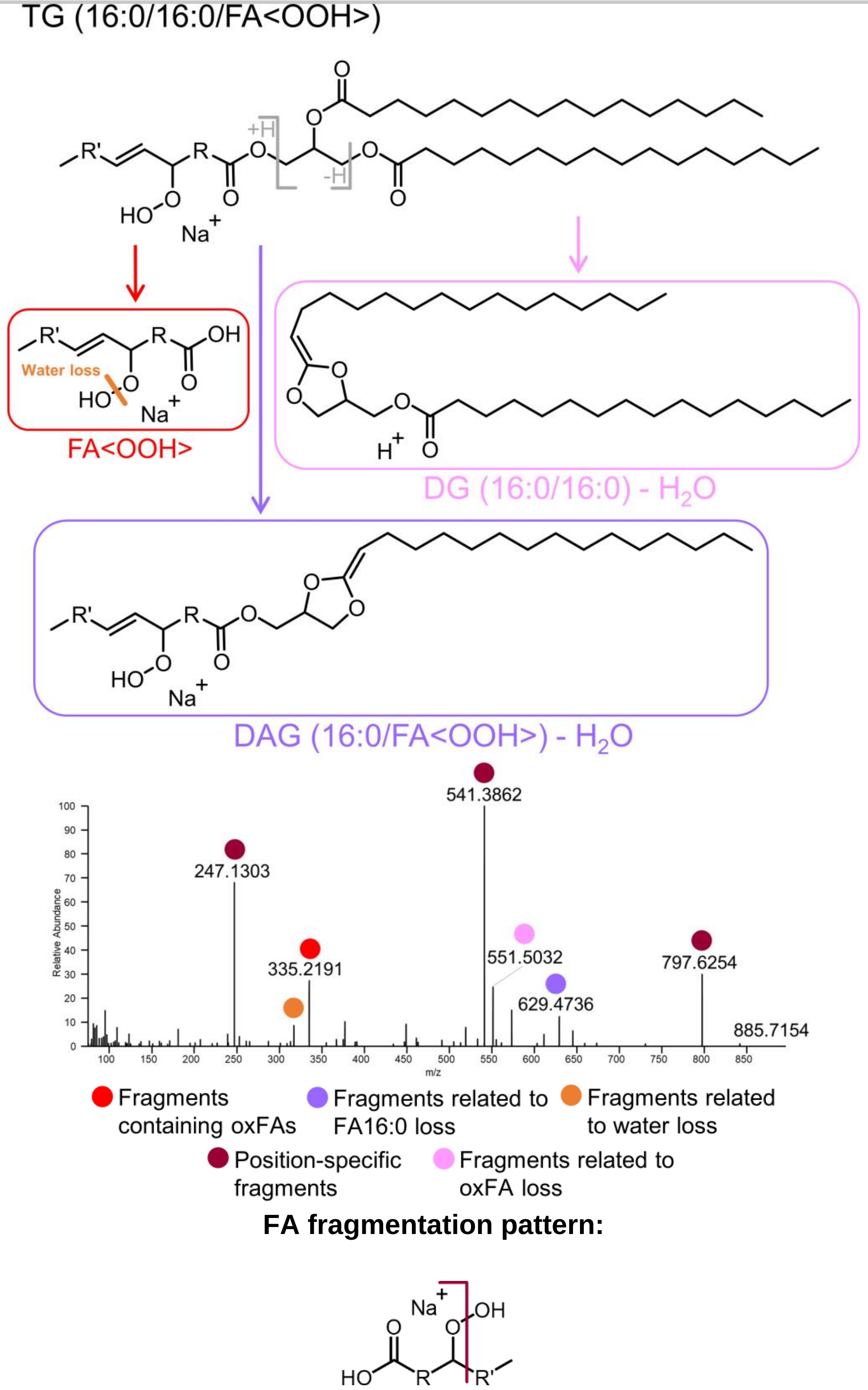 | **FA(18:1<OOH>)**<br>$H_2O$ loss | **$C_{18}H_{34}O_4Na$ 337.2349**<br>$C_{18}H_{32}O_3Na$ 319.2244 | 9 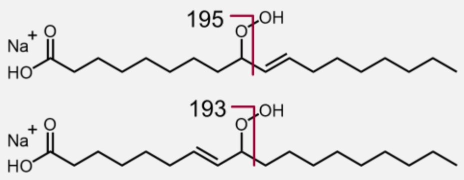 | $C_9H_{16}O_3Na$<br>$C_9H_{14}O_3Na$ | 195.0992<br>193.0835 |
| **TG(16:0/16:0/18:2<OOH>)** | $C_{53}H_{98}O_8Na$ 885.7154 | | **FA(18:2<OOH>)**<br>$H_2O$ loss | **$C_{18}H_{32}O_4Na$ 335.2193**<br>$C_{18}H_{30}O_3Na$ 317.2087 | 9 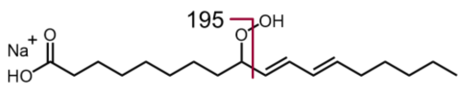 | $C_9H_{16}O_3Na$ | 195.0992 |
| | | | | | 10 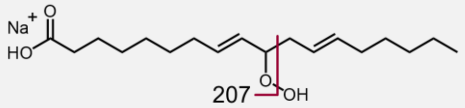 | $C_{10}H_{16}O_3Na$ | 207.0992 |
| | | | | | 11 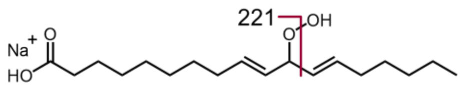 | $C_{11}H_{18}O_3Na$ | 221.1148 |
| | | | | | 12 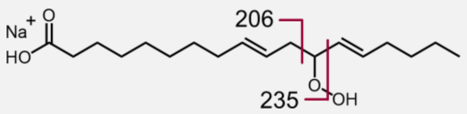 | $C_{11}H_{19}O_2Na$<br>$C_{12}H_{20}O_3Na$ | 206.1277[11]<br>235.1305 |
| | | | | | 13 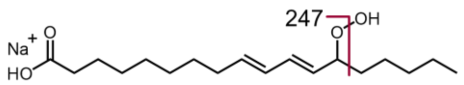 | $C_{13}H_{20}O_3Na$ | 247.1305 |
| **TG(16:0/16:0/20:4<OOH>)** | $C_{55}H_{98}O_8Na$ 909.7154 | | **FA(20:4<OOH>)**<br>$H_2O$ loss | **$C_{20}H_{32}O_4Na$ 359.2193**<br>$C_{20}H_{30}O_3Na$ 341.2087 | 5 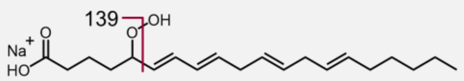 | $C_5H_8O_3Na$ | 139.0366 |
| | | | | | 6 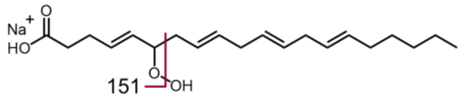 | $C_6H_8O_3Na$ | 151.0366 |
| | | | | | 8 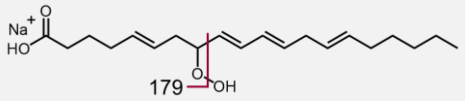 | $C_8H_{12}O_3Na$ | 179.0679 |

#### Full-length oxygenated: <OOH> (continued)

[M+Na]+ adducts

| Precursor | Chemical Formula & *m/z* | MS² Fragmentation | Mod.-specific fragment | Chemical Formula & *m/z* | Pos.-specific fragment | Chemical Formula | *m/z* |
|---|---|---|---|---|---|---|---|
| **TG(16:0/16:0/20:4<OOH>)** | $C_{55}H_{98}O_8Na$ 909.7154 | | **FA(20:4<OOH>)**<br><br>$H_2O$ loss | **$C_{20}H_{30}O_3Na$ 359.2193**<br><br>$C_{20}H_{32}O_4Na$ 341.2087 | 9 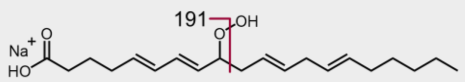  | $C_9H_{12}O_3Na$ | 191.0679 |
| | | | | | 11 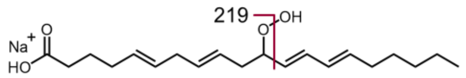  | $C_{11}H_{16}O_3Na$ | 219.0992 |
| | | | | | 12 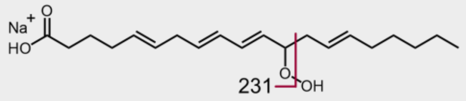  | $C_{12}H_{16}O_3Na$ | 231.0992 |
| | | | | | 14 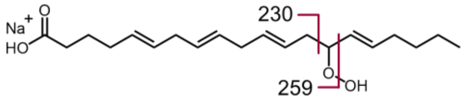  | $C_{13}H_{19}O_2Na$<br>$C_{14}H_{20}O_3Na$ | 230.1277[10]<br>259.1305 |
| | | | | | 15 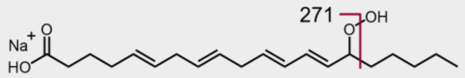  | $C_{15}H_{20}O_3Na$ | 271.1305 |

[10] See *Ito, J., Mizuochi, S., Nakagawa, K., Kato, S., Miyazawa, T., 2015. Tandem Mass Spectrometry Analysis of Linoleic and Arachidonic Acid Hydroperoxides via Promotion of Alkali Metal Adduct Formation. Anal. Chem. 87, 4980–4987.*

**LIPIDMAPS MS/MS spectra**

(negative ionization mode, CE 30V)
https://www.lipidmaps.org/resources/standards/index.php?lipid_category=FA

Name / LIPIDMAPS ID
FA(18:1<ep{9-10}>) / LMFA02000037
FA(18:1<ep{12-13}>) / LMFA02000038
FA(18:2<oxo{13}>) / LMFA02000016
FA(18:2<OH{9}>) / LMFA02000036, LMFA02000188
FA(18:2<OH{13}>) / LMFA02000035, LMFA02000228
FA(18:2<OOH{9}>) / LMFA02000012
FA(18:2<OOH{13}>) / LMFA02000034
FA(20:3<ep{5-6}>) / LMFA03080002
FA(20:3<ep{8-9}>) / LMFA03080003
FA(20:3<ep{11-12}>) / LMFA03080004
FA(20:3<ep{14-15}>) / LMFA03080005
FA(20:4<oxo{5}>) / LMFA03060011
FA(20:4<oxo{12}>) / LMFA03060019
FA(20:4<OH{5}>) / LMFA03060002
FA(20:4<OH{8}>) / LMFA03060006
FA(20:4<OH{9}>) / LMFA03060089
FA(20:4<OH{11}>) / LMFA03060003
FA(20:4<OH{12}>) / LMFA03060007, LMFA03060008
FA(20:4<OH{15}>) / LMFA03060001
FA(20:4<OOH{5}>) / LMFA03060012
FA(20:4<OOH{12}>) / LMFA03060013
FA(20:4<OOH{15}>) / LMFA03060014

**METLIN MS/MS spectra**

(negative ion mode, CE 10, 20V)
https://metlin.scripps.edu/landing_page.php?pgcontent=mainPage

Name / METLIN ID

FA(18:0<ep{9-10}>) / 36008
FA(18:1<ep{9-10}>) / 43441
FA(18:1<ep{12-13}>) / 43442
FA(18:2<OH{9}>) / 35487, 45660, 45662
FA(18:2<OH{13}>) / 35490, 45665, 45667
FA(18:2<oxo{9}>) / 35860
FA(18:2<oxo{13}>) / 36023
FA(18:2<OOH{9}>) / 36019, 64785
FA(18:2<OOH{13}>) / 36036, 64784
FA(20:4<OH{5}>) / 36336, 45646, 45684
FA(20:4<OH{8}>) / 36286, 3840, 45730
FA(20:4<OH{9}>) / 36290, 45649, 45650
FA(20:4<OH{11}>) / 36337, 3838, 45056
FA(20:4<OH{12}>) / 3841, 45054, 45653
FA(20:4<OH{15}>) / 3836, 45651
FA(20:4<oxo{5}>) / 3844
FA(20:4<oxo{12}>) / 36285
FA(20:4<OOH{5}>) / 36281
FA(20:4<OOH{12}>) / 3845
FA(20:4<OOH{15}>) / 3846
