## Supplementary material for "Epilipidomics platform for holistic profiling of oxidized complex lipids in blood plasma of obese individuals": FileS2

**File S2.** MS2 spectra acquired using stDDA and used for the annotation of **oxidized PC** (oxPC) species in group pooled blood plasma samples of obese non-diabetic (OND) and obese with type 2 diabetes (OT2D) individuals. Structure-related fragmentation ions are colour-codes according to the legend provided. Annotated lipids for each group pool are sorted by their precursor *m/z*.

### OND_pool

# PC(16:1_18:2<O>)
# RT 12.0

**[oxPC+HCOO]-    XIC 816.5396**

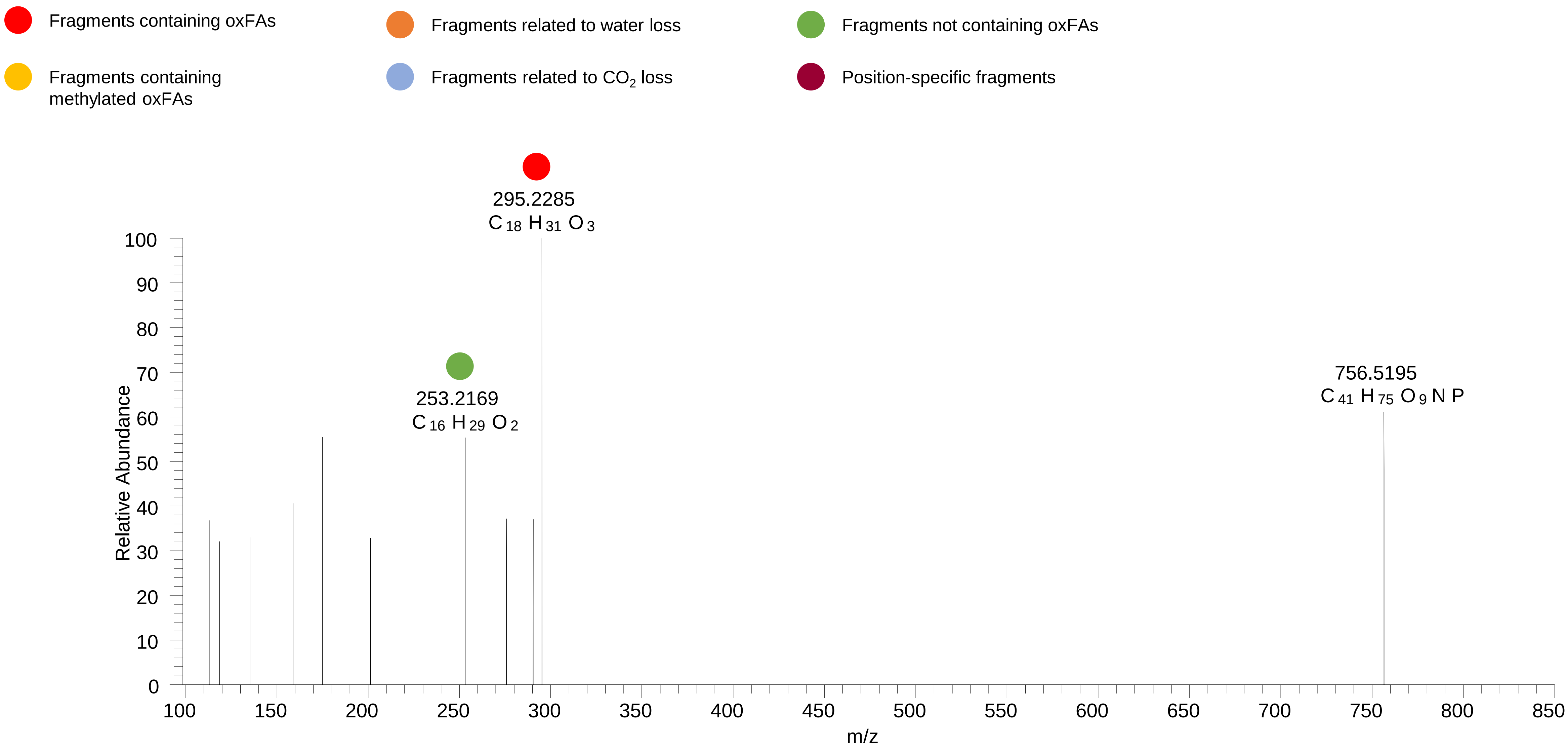

# PC(16:0_18:3<O>)
# RT 12.4

**[oxPC+HCOO]- XIC 816.5396**

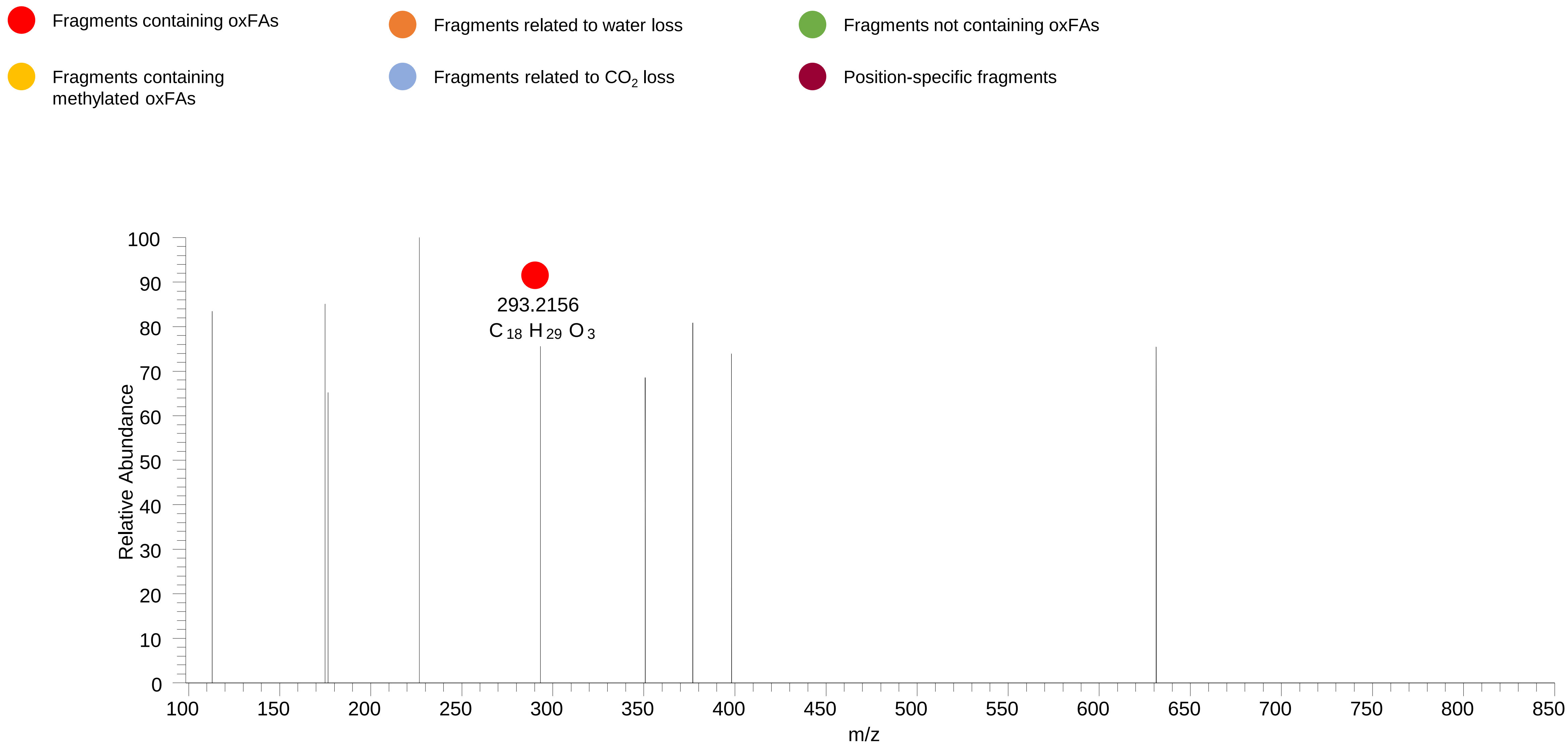

# PC(16:0_18:2<OH>)
# RT 12.6

**[oxPC+HCOO]- XIC 818.5553**

- Fragments containing oxFAs
- Fragments related to water loss
- Fragments not containing oxFAs
- Fragments containing methylated oxFAs
- Fragments related to $CO_2$ loss
- Position-specific fragments

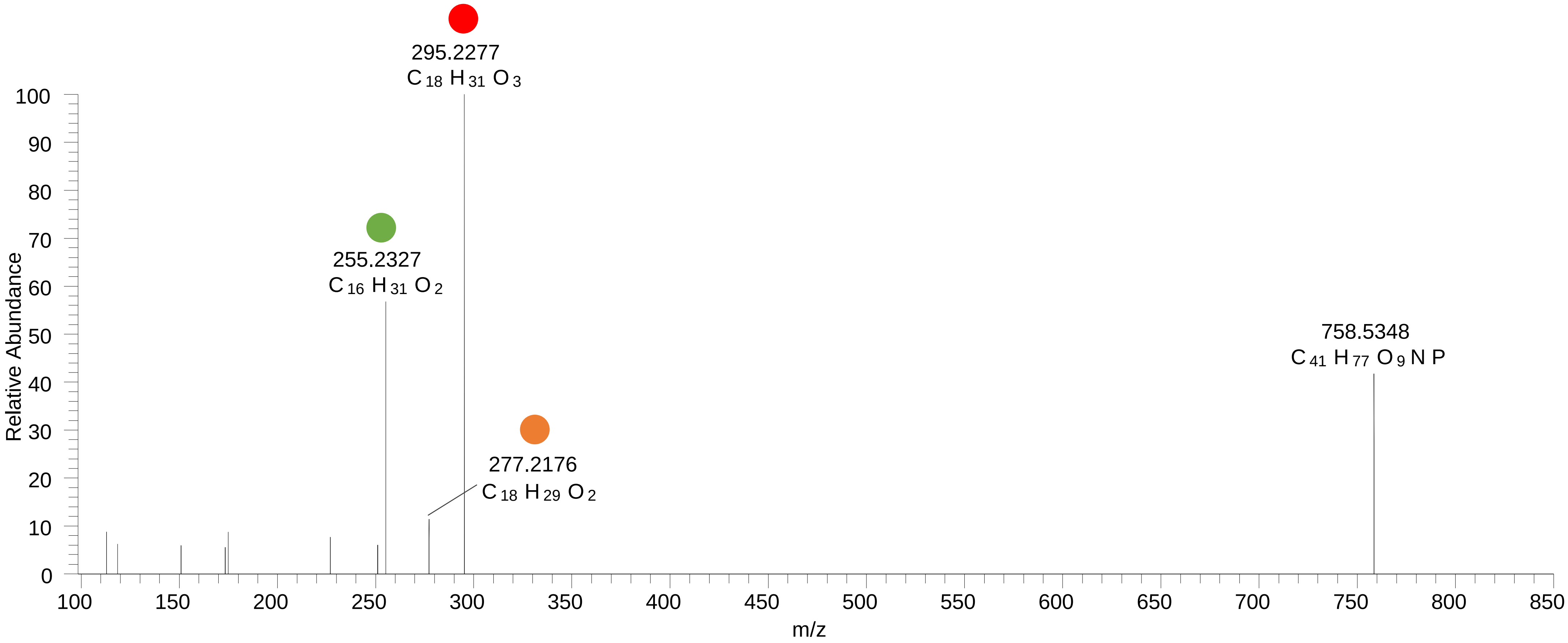

# PC(16:0_18:2<OH{13}>)
# RT 13.4

**[oxPC+HCOO]- XIC 818.5553**

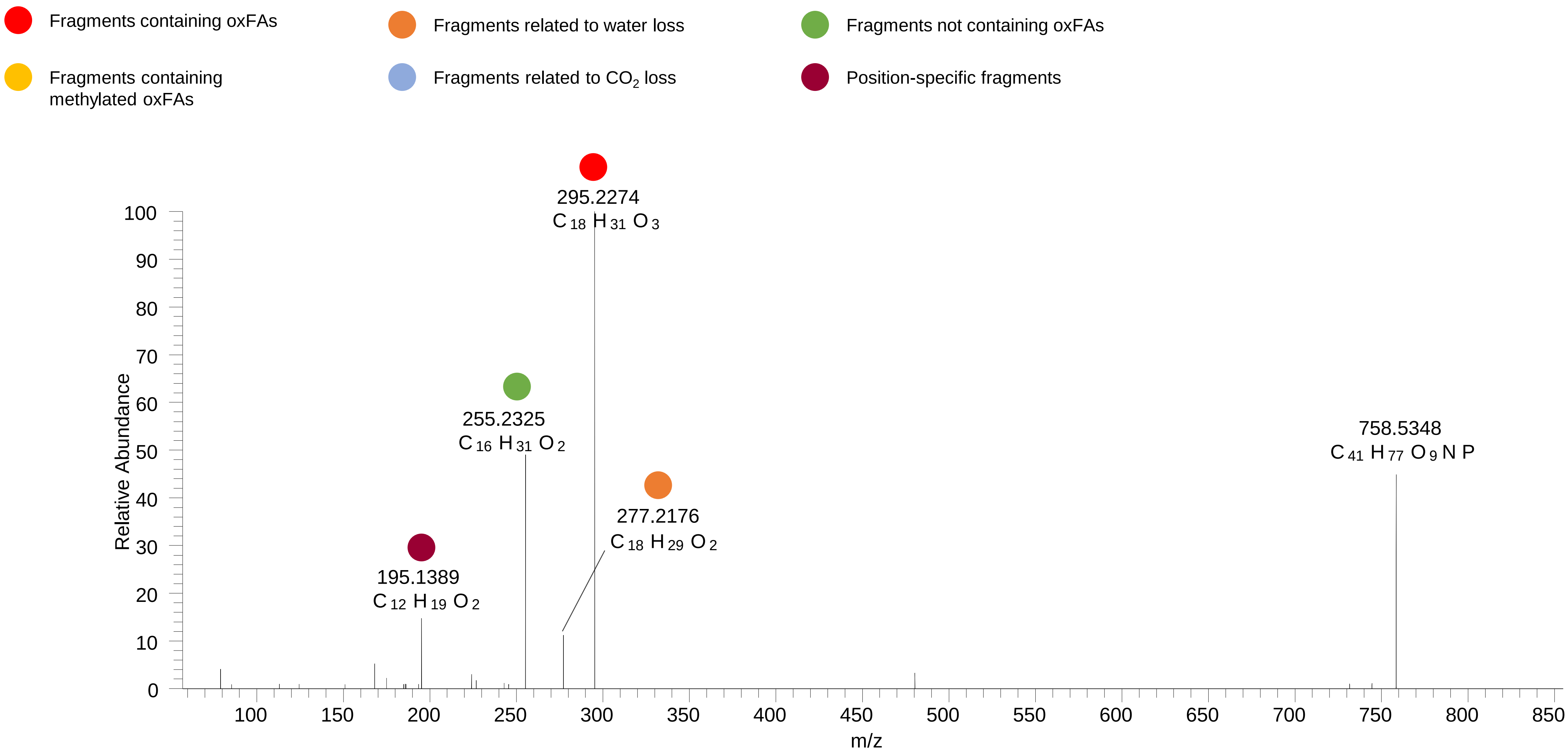

# PC(16:0_18:2<OH{9}>)
# RT 13.6

**[oxPC+HCOO]- XIC 818.5553**

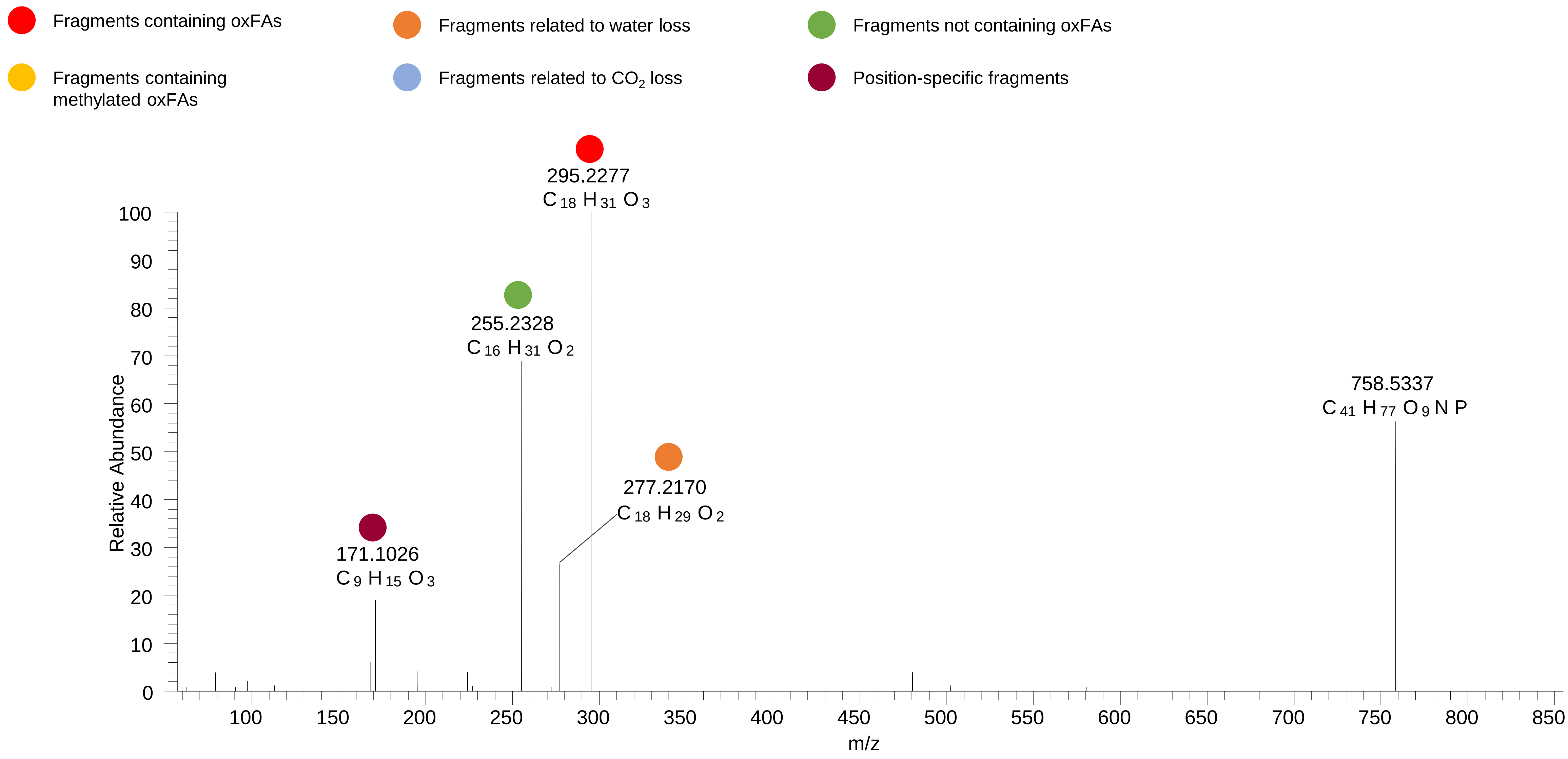

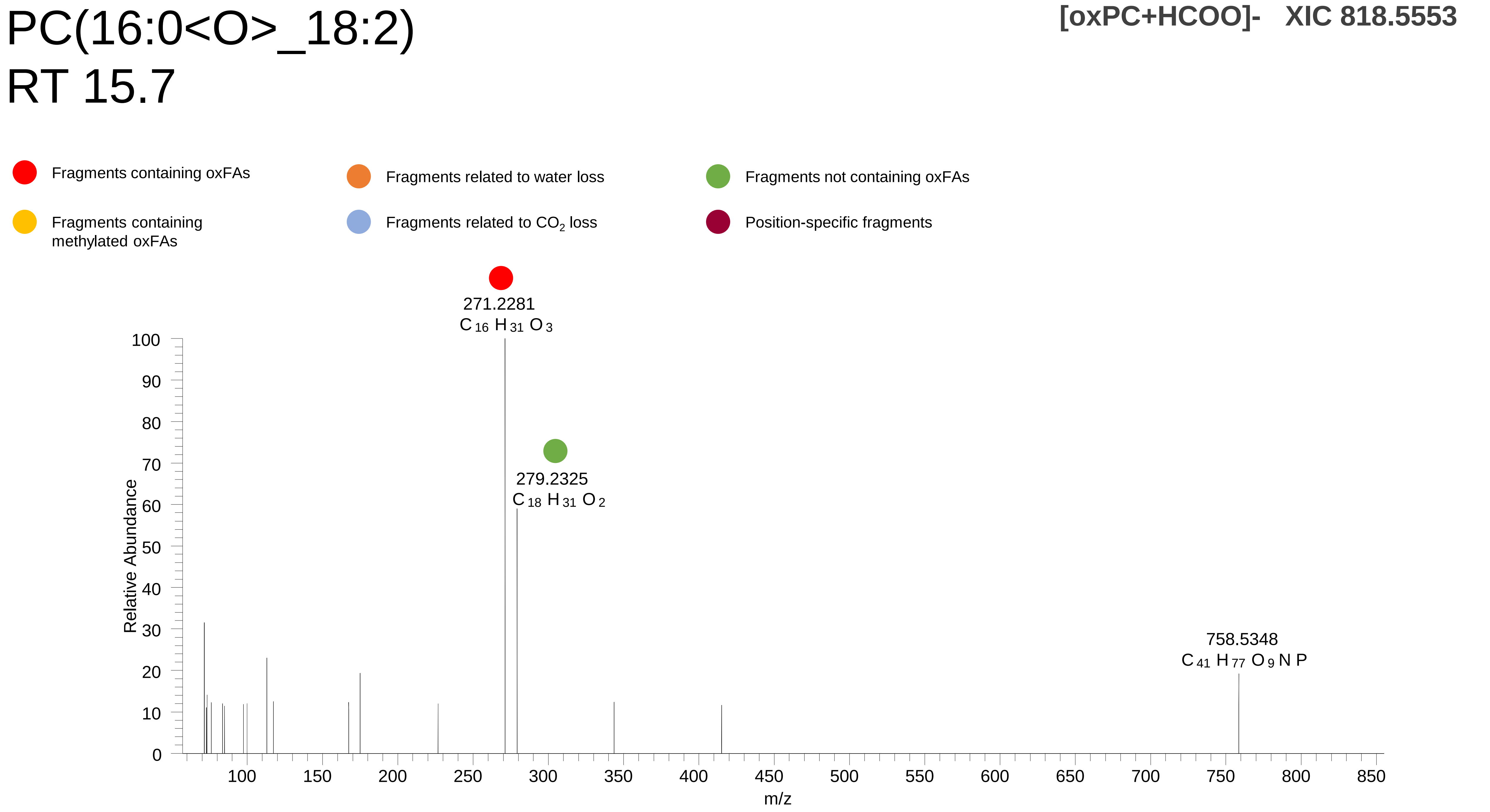

PC(16:0<O>_18:2)
RT 15.7
[oxPC+HCOO]- XIC 818.5553
Fragments containing oxFAs
Fragments related to water loss
Fragments not containing oxFAs
Fragments containing methylated oxFAs
Fragments related to $CO_2$ loss
Position-specific fragments
271.2281
$C_{16}H_{31}O_3$
279.2325
$C_{18}H_{31}O_2$
758.5348
$C_{41}H_{77}O_9NP$
Relative Abundance
m/z

# PC(16:0_18:1<O>)
# RT 14.2

**[oxPC+HCOO]- XIC 820.5709**

- Fragments containing oxFAs
- Fragments related to water loss
- Fragments not containing oxFAs
- Fragments containing methylated oxFAs
- Fragments related to $CO_2$ loss
- Position-specific fragments

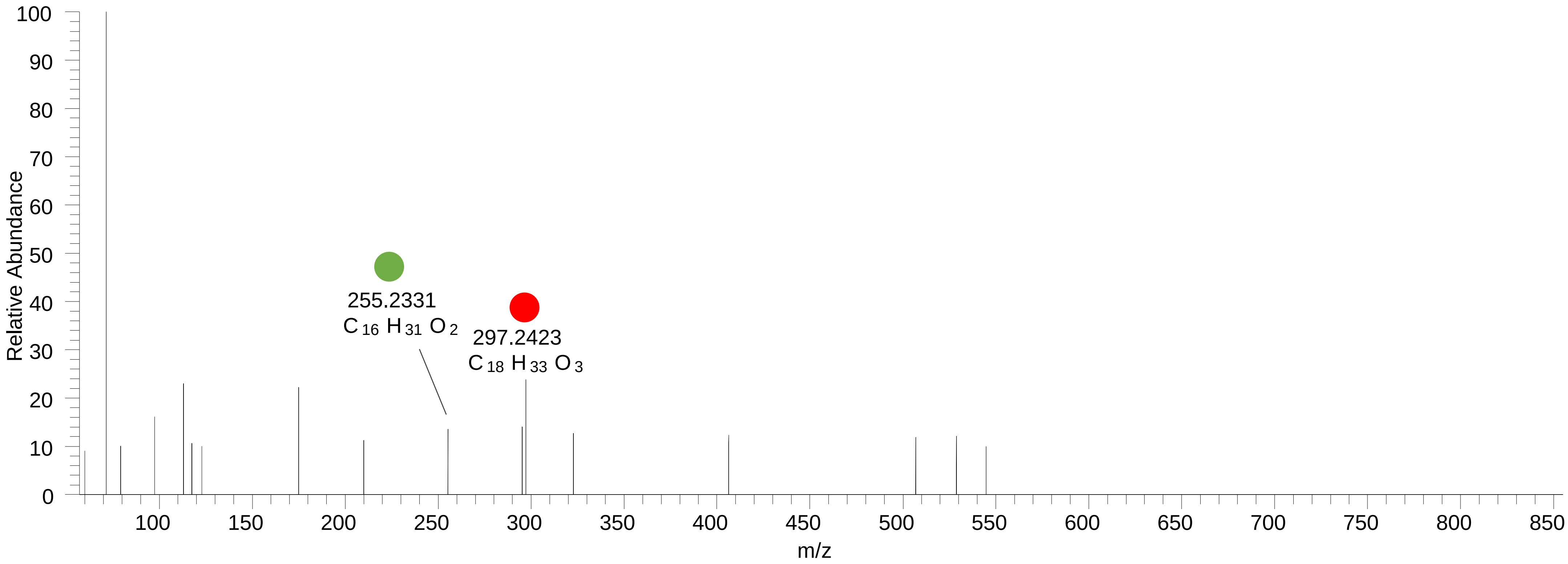

# PC(16:0<O>_18:1)
# RT 16.8-17.1

**[oxPC+HCOO]- XIC 820.5709**

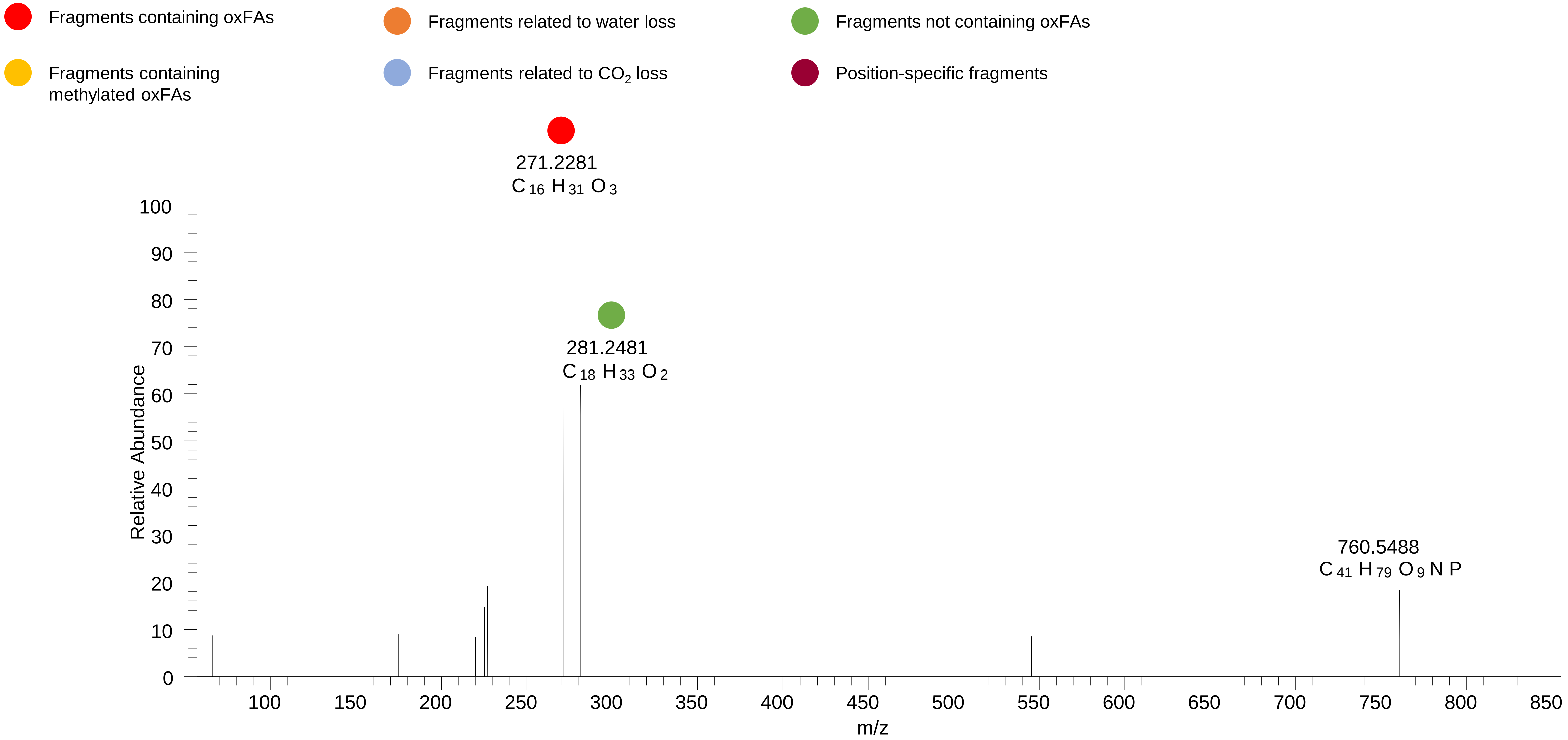

# PC(16:0_18:2<OH,O{9}>)
# RT 11.6

**[oxPC+HCOO]- XIC 834.5502**

- Fragments containing oxFAs
- Fragments related to water loss
- Fragments not containing oxFAs
- Fragments containing methylated oxFAs
- Fragments related to $CO_2$ loss
- Position-specific fragments

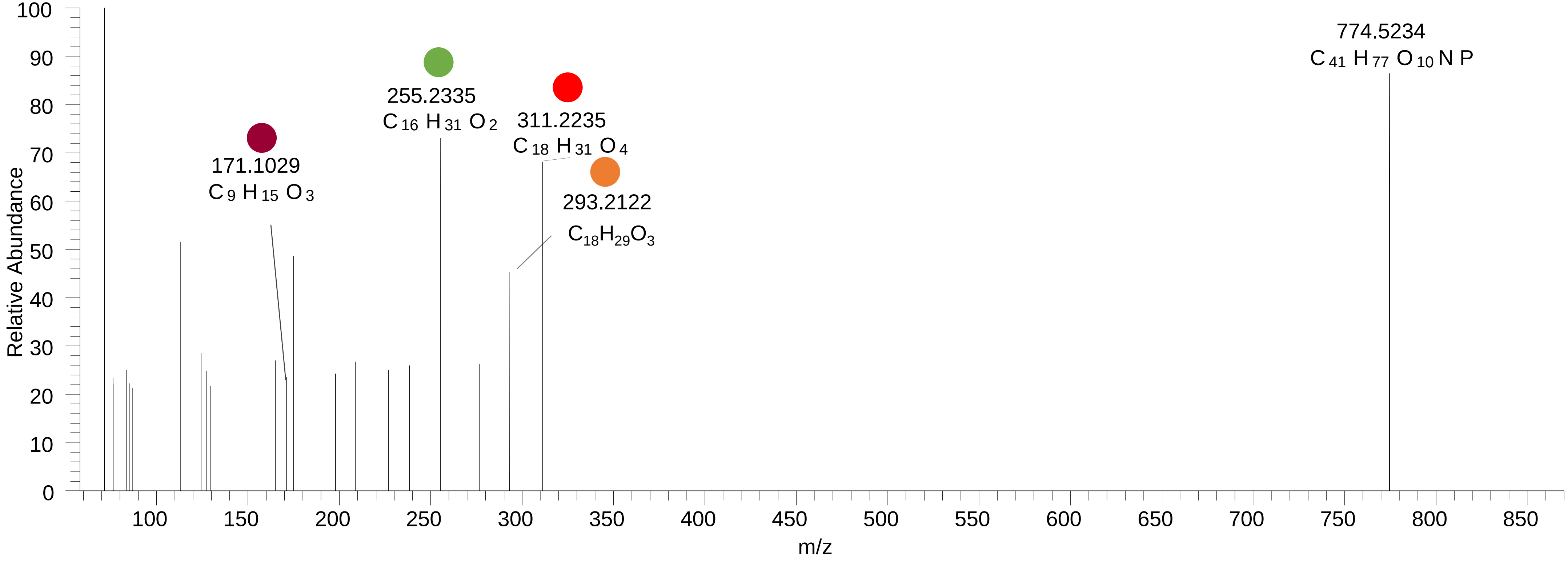

### PC(16:0_18:2<OOH>)
# RT 12.8

**[oxPC+HCOO]- XIC 834.5502**

- Fragments containing oxFAs
- Fragments related to water loss
- Fragments not containing oxFAs
- Fragments containing methylated oxFAs
- Fragments related to $CO_2$ loss
- Position-specific fragments

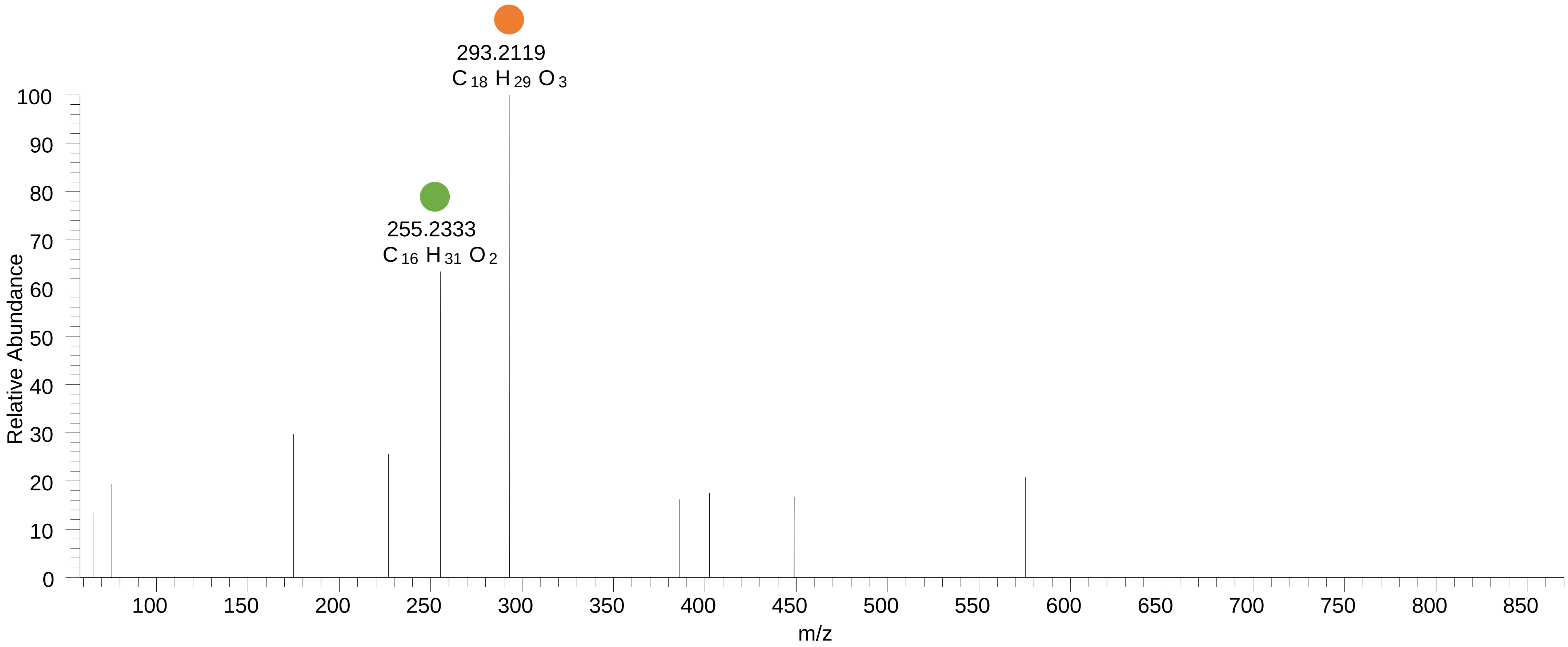

### PC(16:0_18:2<OOH{13}>)
# RT 13.1-13.4

**[oxPC+HCOO]- XIC 834.5502**

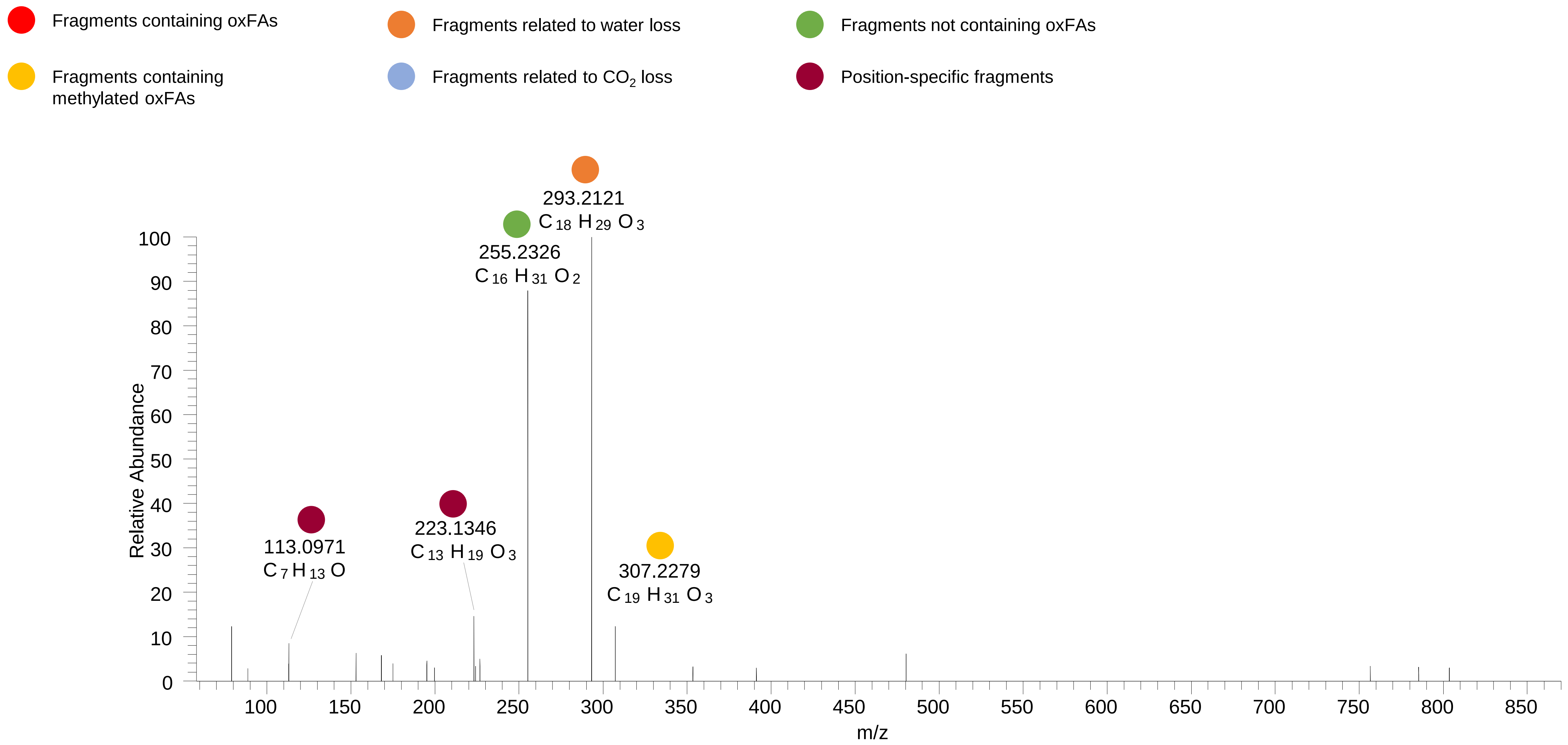

### PC(16:0_18:2<OOH{9}>)
# RT 13.6

**[oxPC+HCOO]- XIC 834.5502**

- Fragments containing oxFAs
- Fragments related to water loss
- Fragments not containing oxFAs
- Fragments containing methylated oxFAs
- Fragments related to $CO_2$ loss
- Position-specific fragments

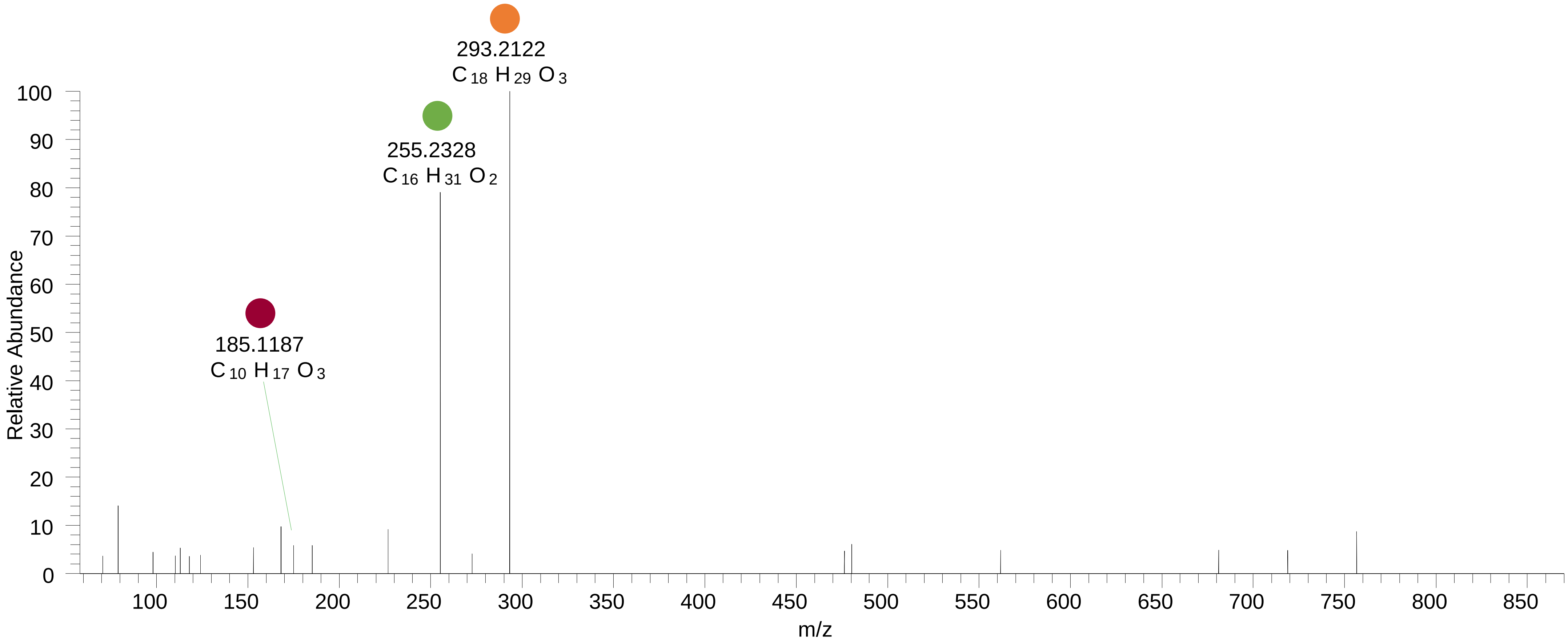

### PC(16:0_18:1<OOH>)
# RT 14.3

**[oxPC+HCOO]- XIC 836.5658**

- Fragments containing oxFAs
- Fragments related to water loss
- Fragments not containing oxFAs
- Fragments containing methylated oxFAs
- Fragments related to $CO_2$ loss
- Position-specific fragments

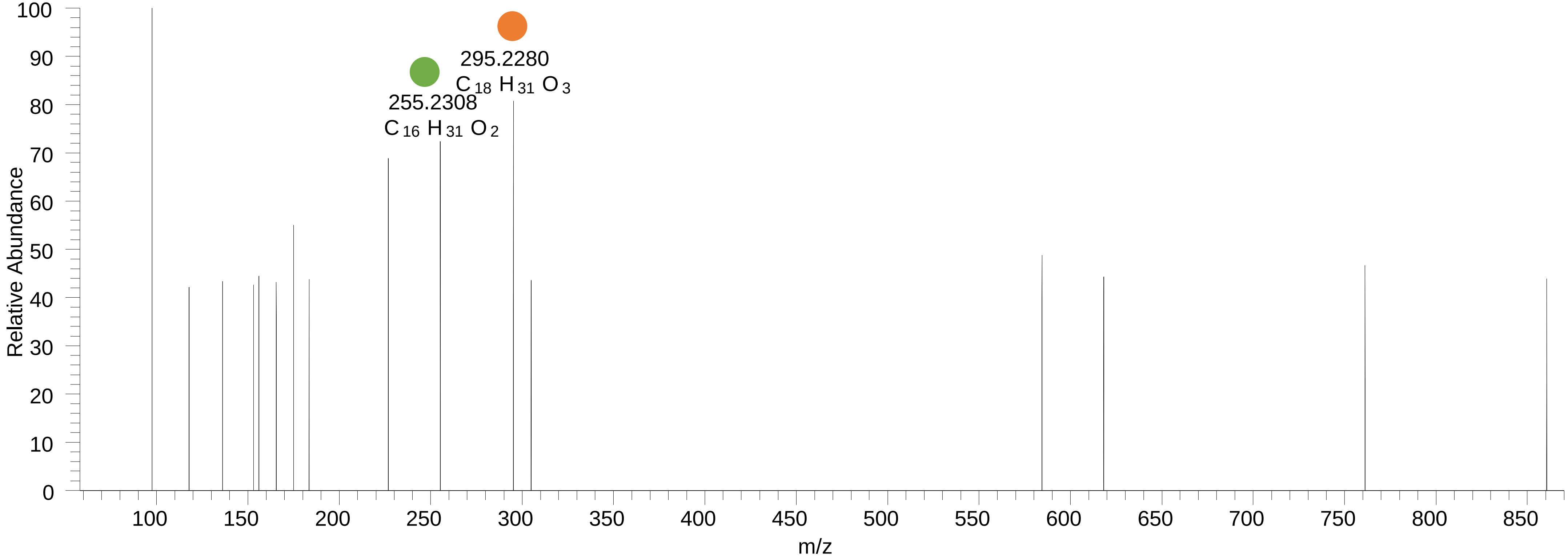

# PC(16:0_20:5<O>)
# RT 12.6-13.0

**[oxPC+HCOO]- XIC 840.5396**

- Fragments containing oxFAs
- Fragments related to water loss
- Fragments not containing oxFAs
- Fragments containing methylated oxFAs
- Fragments related to $CO_2$ loss
- Position-specific fragments

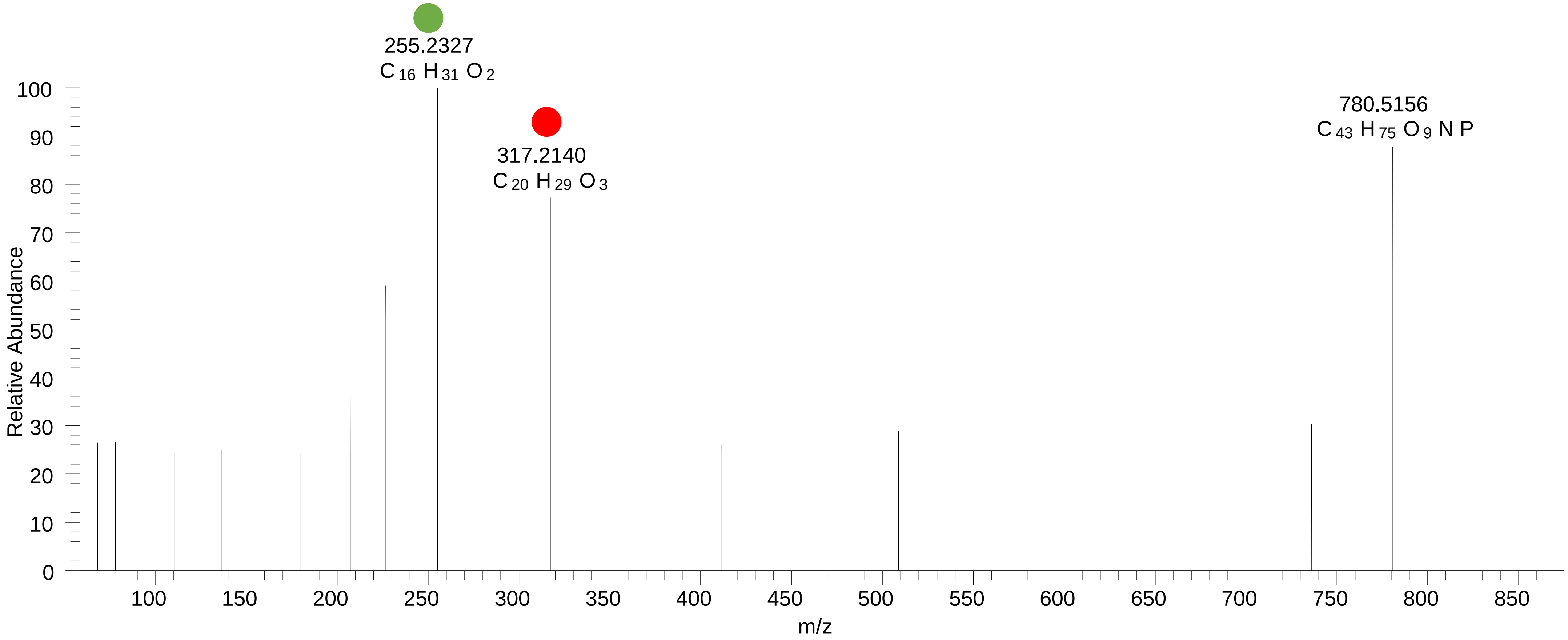

# PC(18:2_18:2<OH>)

# RT 12.5

**[oxPC+HCOO]- XIC 842.5553**

- Fragments containing oxFAs
- Fragments related to water loss
- Fragments not containing oxFAs
- Fragments containing methylated oxFAs
- Fragments related to $CO_2$ loss
- Position-specific fragments

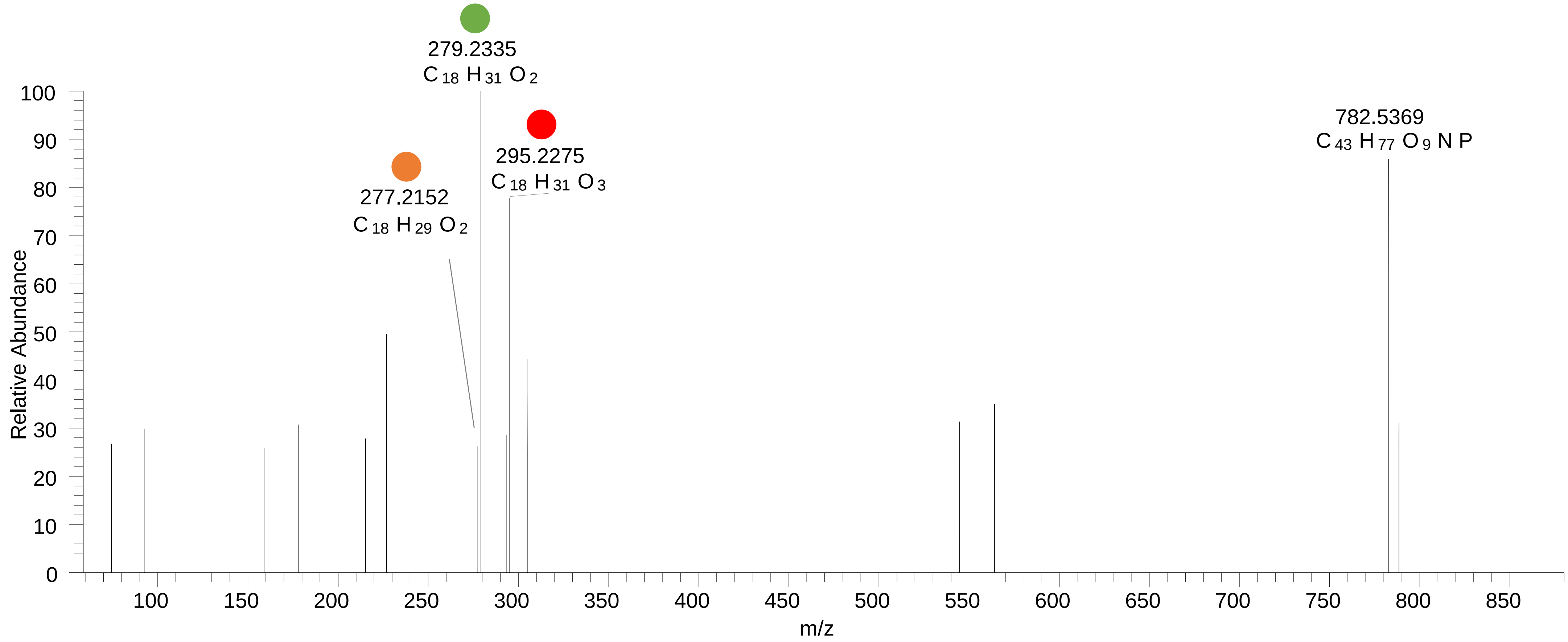

# PC(16:0_20:4<OH{13}>)
# RT 13.2

**[oxPC+HCOO]-   XIC 842.5553**

- Fragments containing oxFAs
- Fragments related to water loss
- Fragments not containing oxFAs
- Fragments containing methylated oxFAs
- Fragments related to $CO_2$ loss
- Position-specific fragments

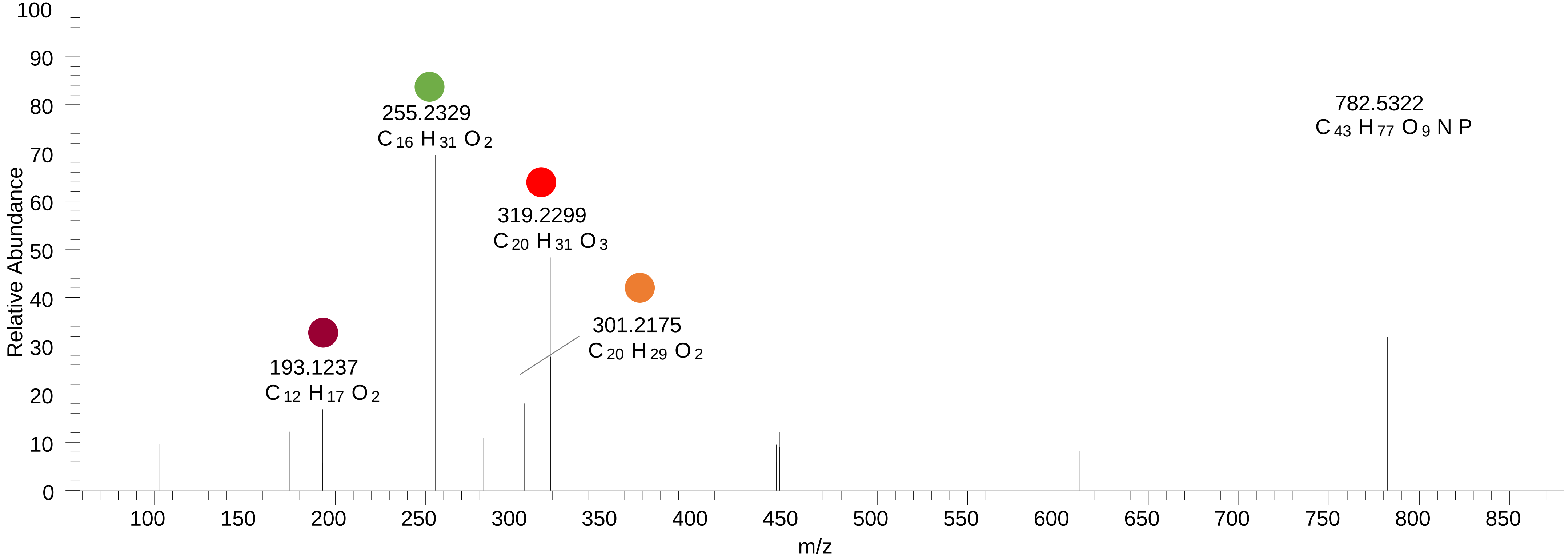

# PC(16:0_20:4<OH{15}>)
# RT 13.6

[oxPC+HCOO]- XIC 842.5553

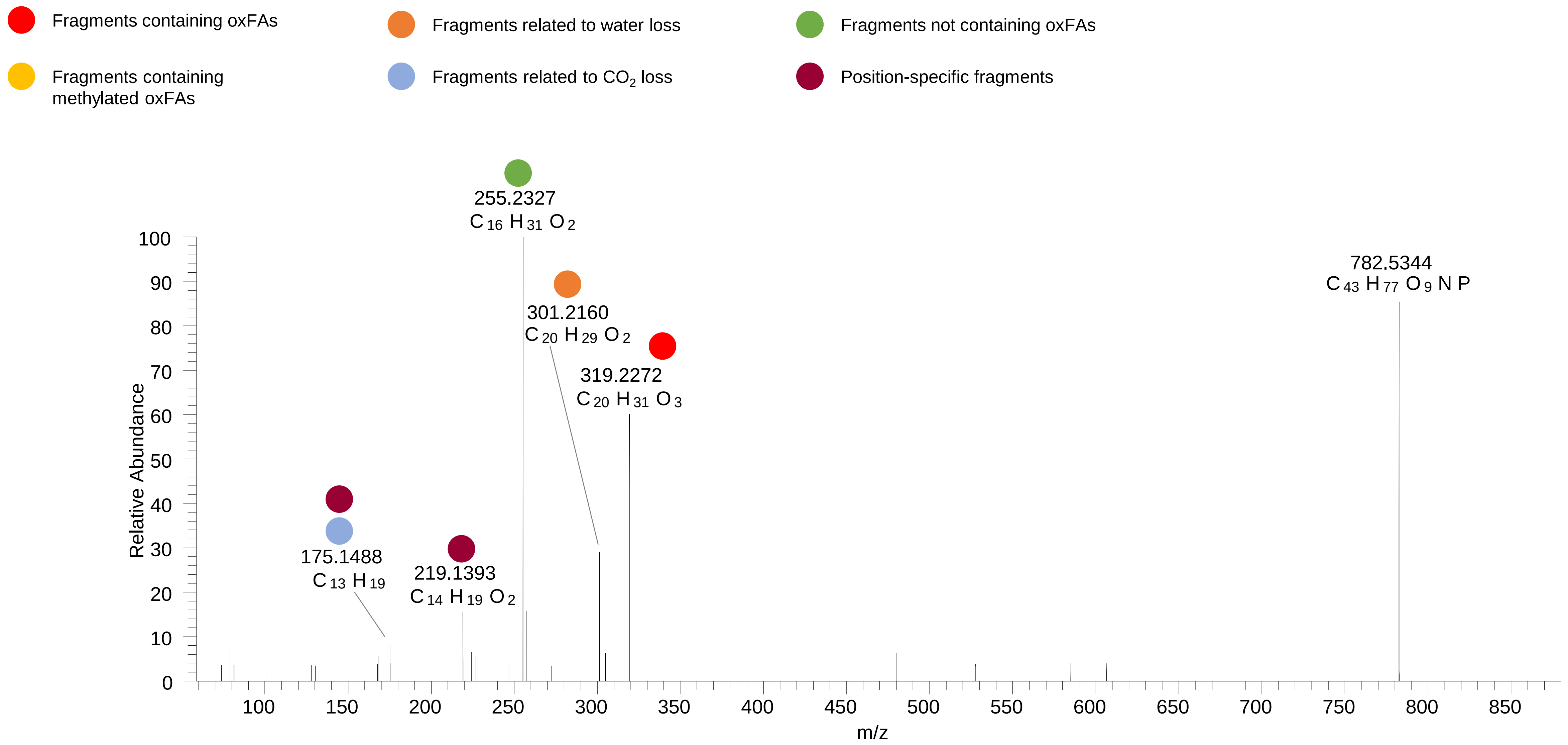

# PC(16:0_20:4<OH{11}>)
# PC(16:0_20:4<OH{12}>)
# RT 14.0

**[oxPC+HCOO]- XIC 842.5553**

- Fragments containing oxFAs
- Fragments related to water loss
- Fragments not containing oxFAs
- Fragments containing methylated oxFAs
- Fragments related to $CO_2$ loss
- Position-specific fragments

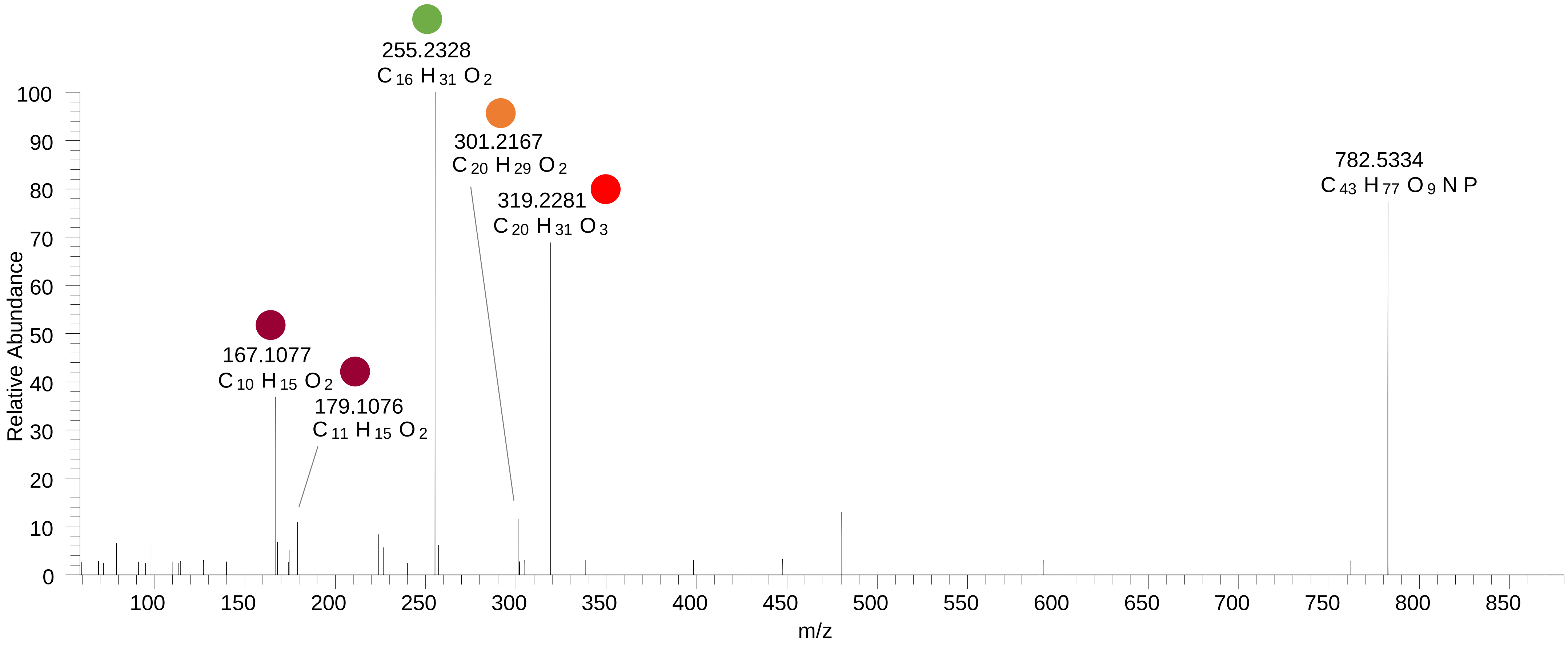

# PC(16:0_20:4<OH{8}>)
# PC(16:0_20:4<OH{9}>)
# RT 14.5

**[oxPC+HCOO]- XIC 842.5553**

- Fragments containing oxFAs
- Fragments related to water loss
- Fragments not containing oxFAs
- Fragments containing methylated oxFAs
- Fragments related to $CO_2$ loss
- Position-specific fragments

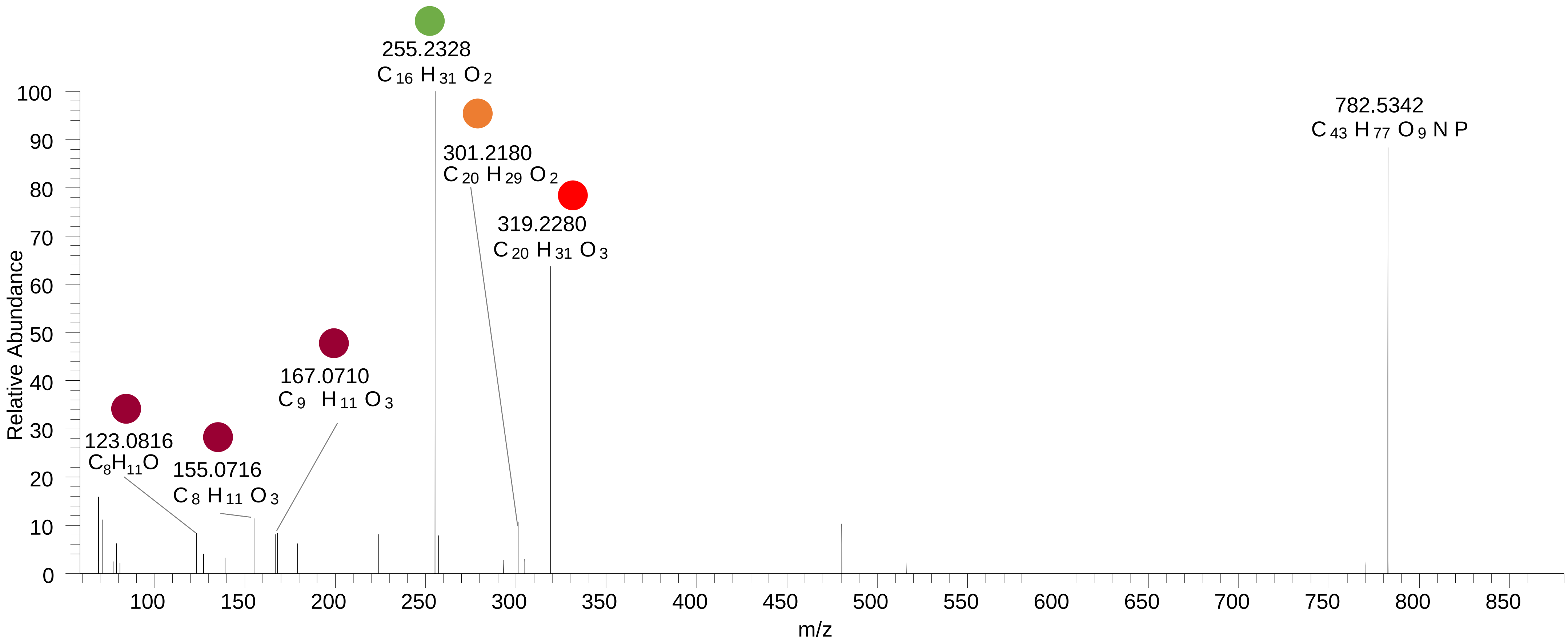

# PC(16:0_20:4<OH{5}>)
# RT 15.1

**[oxPC+HCOO]- XIC 842.5553**

- Fragments containing oxFAs
- Fragments related to water loss
- Fragments not containing oxFAs
- Fragments containing methylated oxFAs
- Fragments related to $CO_2$ loss
- Position-specific fragments

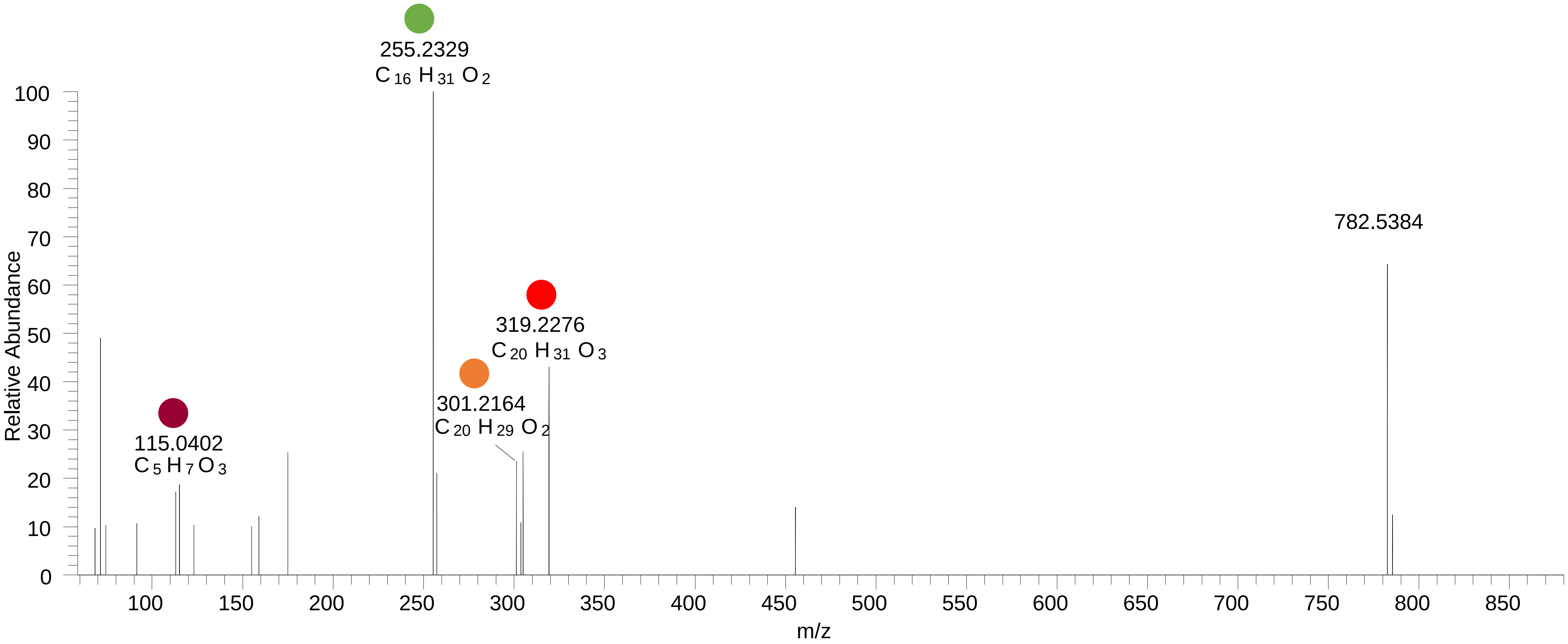

# PC(18:1_18:2<O>)
# RT 12.8

**[oxPC+HCOO]- XIC 844.5709**

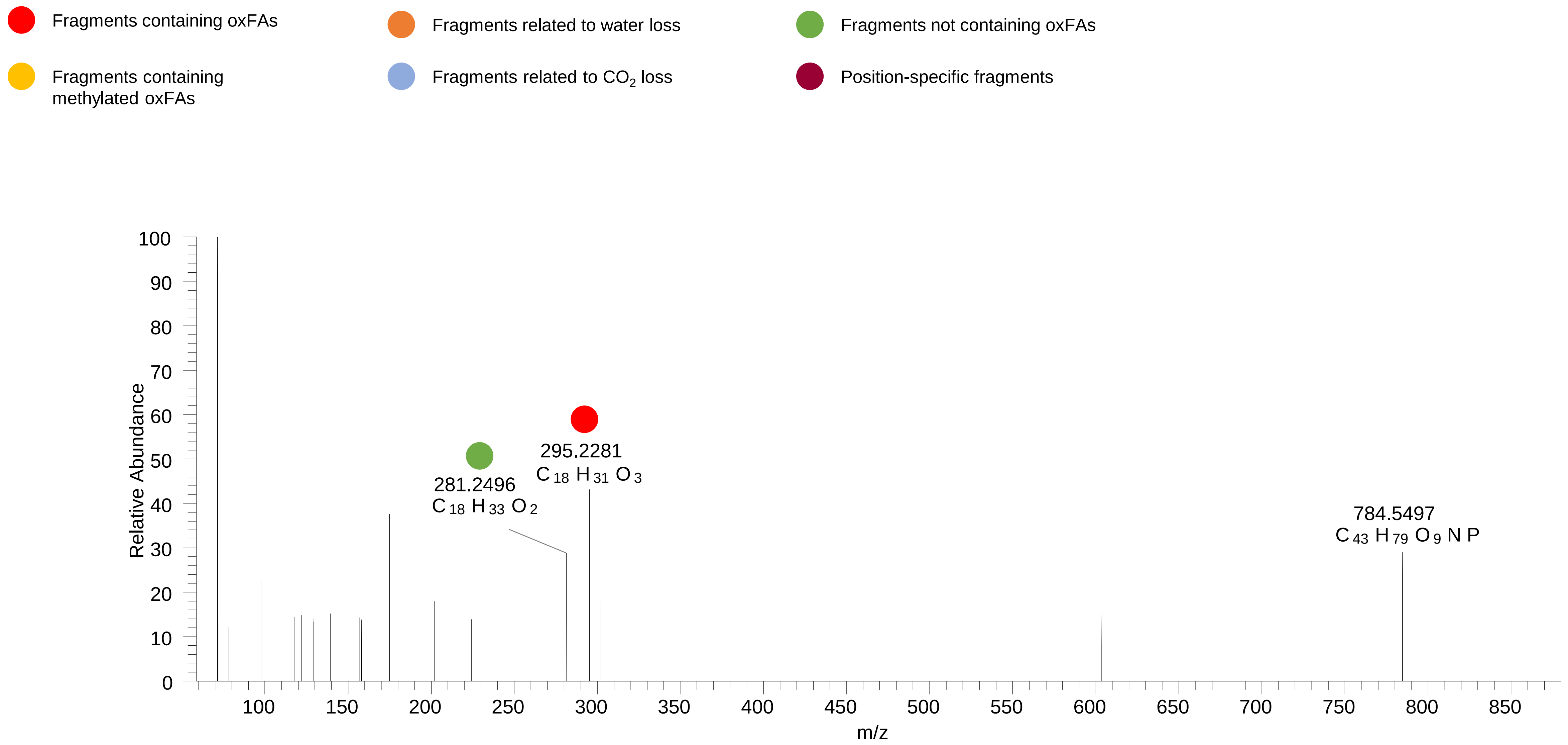

# PC(16:0_20:3<OH{12}>)
# RT 14.3

**[oxPC+HCOO]- XIC 844.5709**

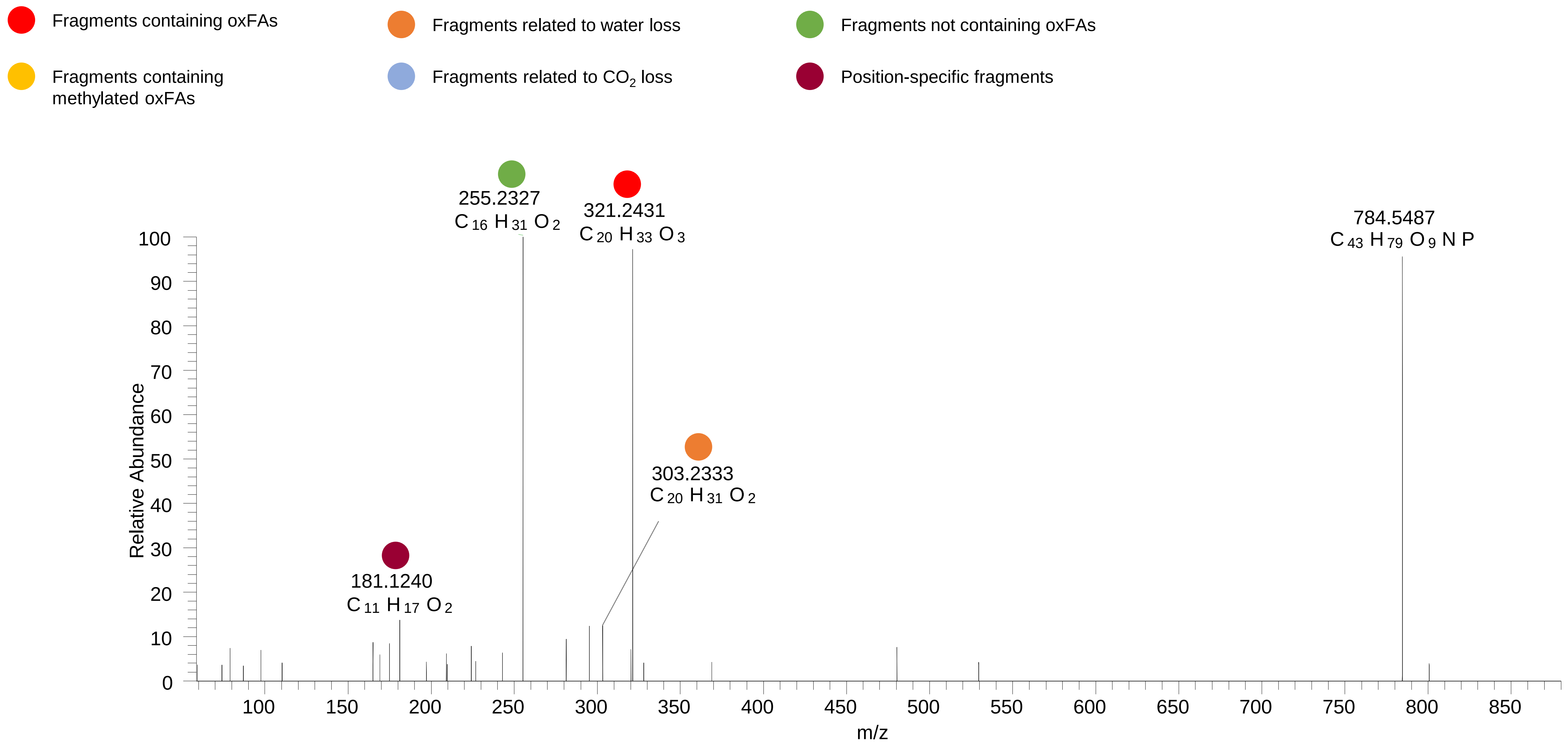

# PC(18:0_18:2<O>)
# PC(16:0_20:2<OH>)
# RT 14.6

**[oxPC+HCOO]- XIC 846.5866**

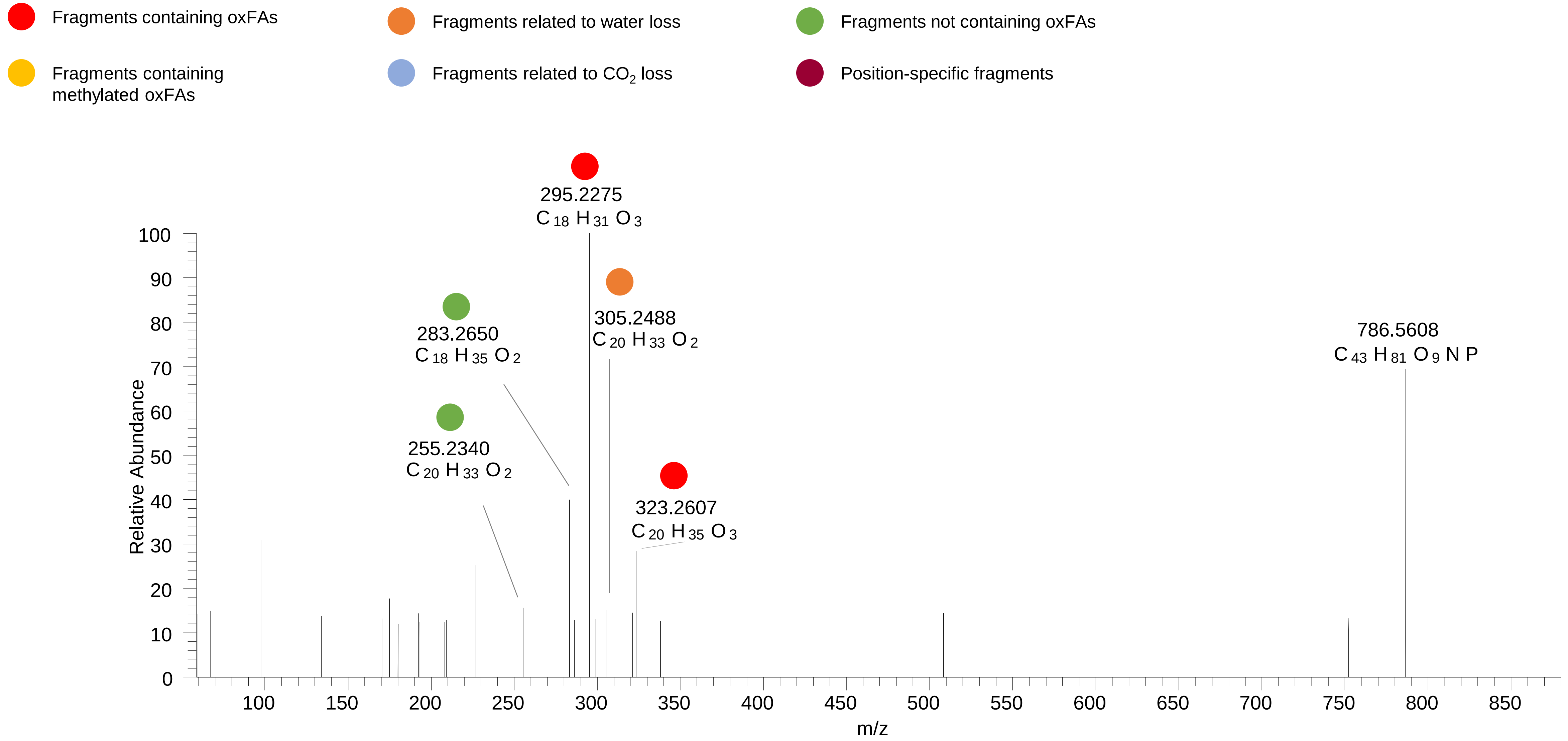

# PC(18:0_18:2<O>)
# RT 14.8

**[oxPC+HCOO]-   XIC 846.5866**

- Fragments containing oxFAs
- Fragments related to water loss
- Fragments not containing oxFAs
- Fragments containing methylated oxFAs
- Fragments related to $CO_2$ loss
- Position-specific fragments

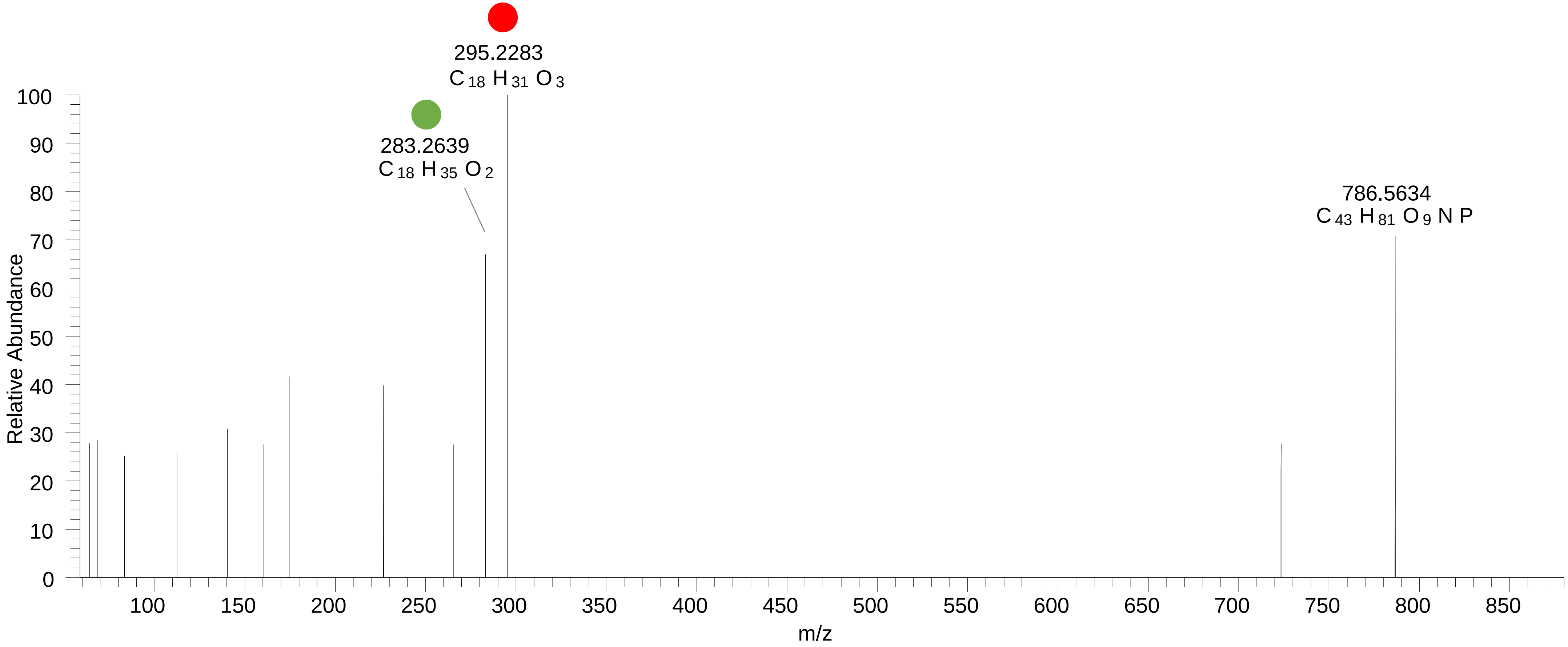

# PC(18:0_18:2<OH{13}>)
# RT 15.0

**[oxPC+HCOO]- XIC 846.5866**

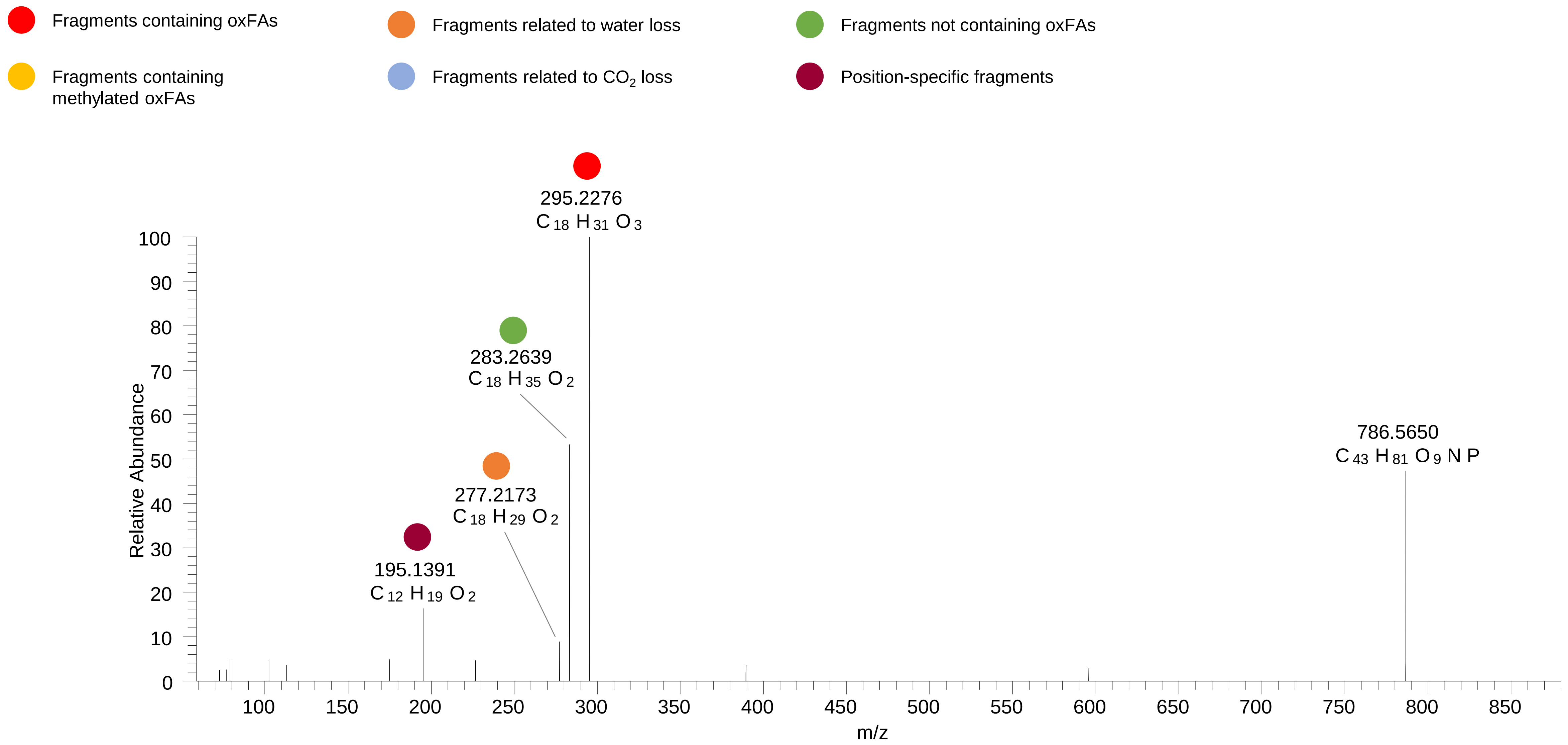

# PC(18:0_18:2<OH{9}>)
# RT 15.3

**[oxPC+HCOO]- XIC 846.5866**

- Fragments containing oxFAs
- Fragments related to water loss
- Fragments not containing oxFAs
- Fragments containing methylated oxFAs
- Fragments related to $CO_2$ loss
- Position-specific fragments

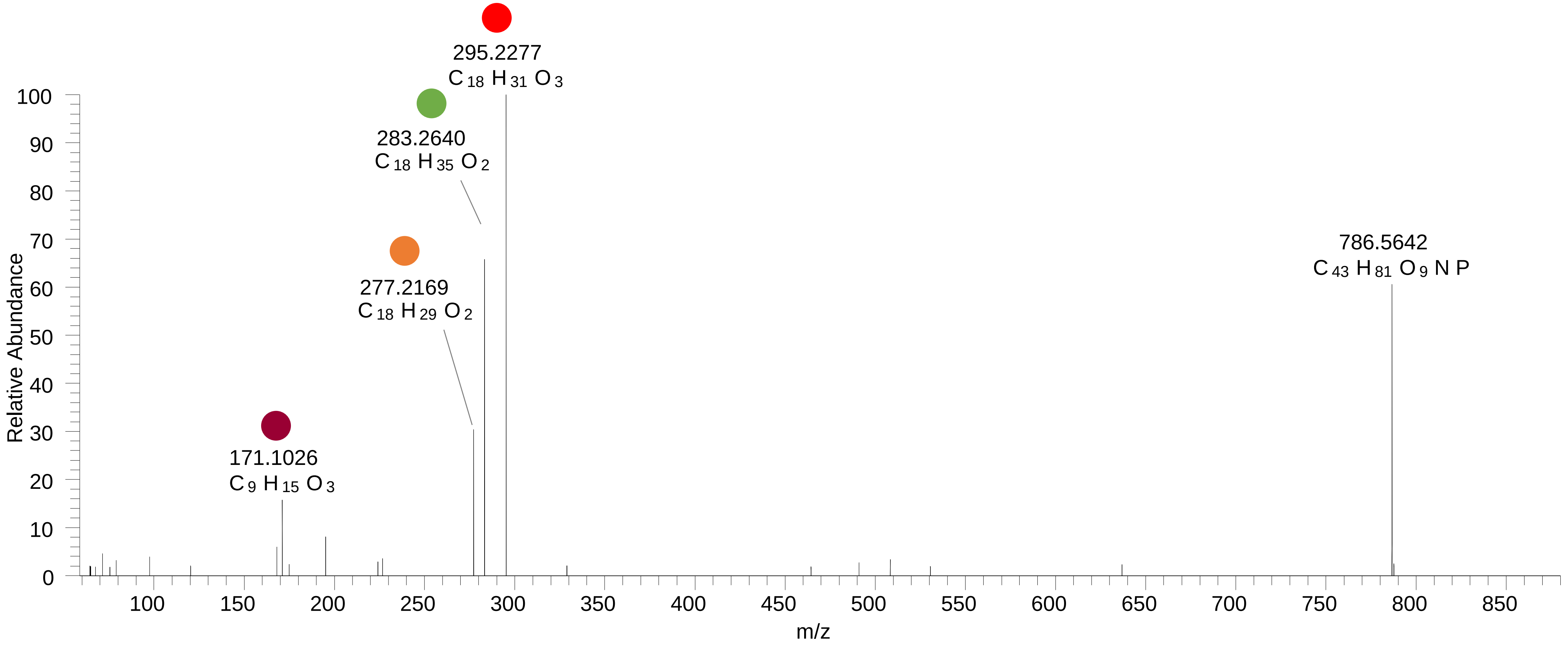

### PC(16:0_18:2<OOH,OH>)
# RT 11.7

**[oxPC+HCOO]-   XIC 850.5451**

- Fragments containing oxFAs
- Fragments related to water loss
- Fragments not containing oxFAs
- Fragments containing methylated oxFAs
- Fragments related to $CO_2$ loss
- Position-specific fragments

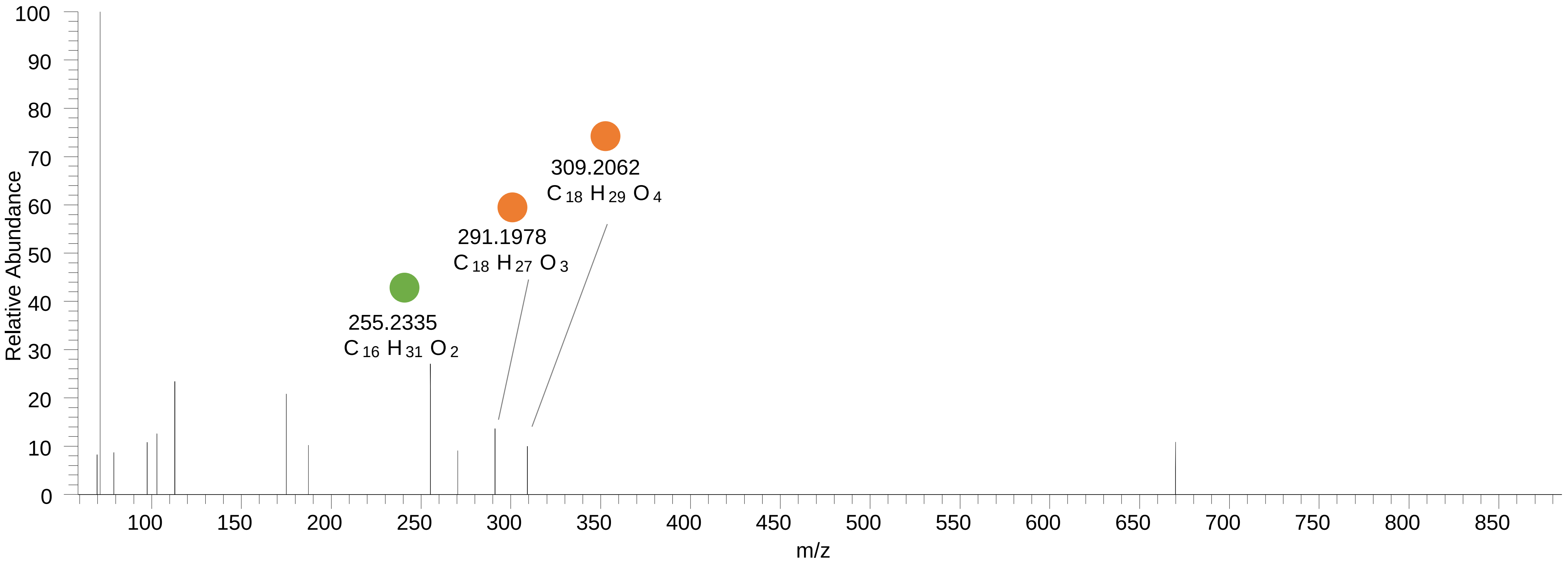

# PC(16:0_20:4<OH,O>)
# RT 11.1

**[oxPC+HCOO]- XIC 858.5502**

- Fragments containing oxFAs
- Fragments related to water loss
- Fragments not containing oxFAs
- Fragments containing methylated oxFAs
- Fragments related to $CO_2$ loss
- Position-specific fragments

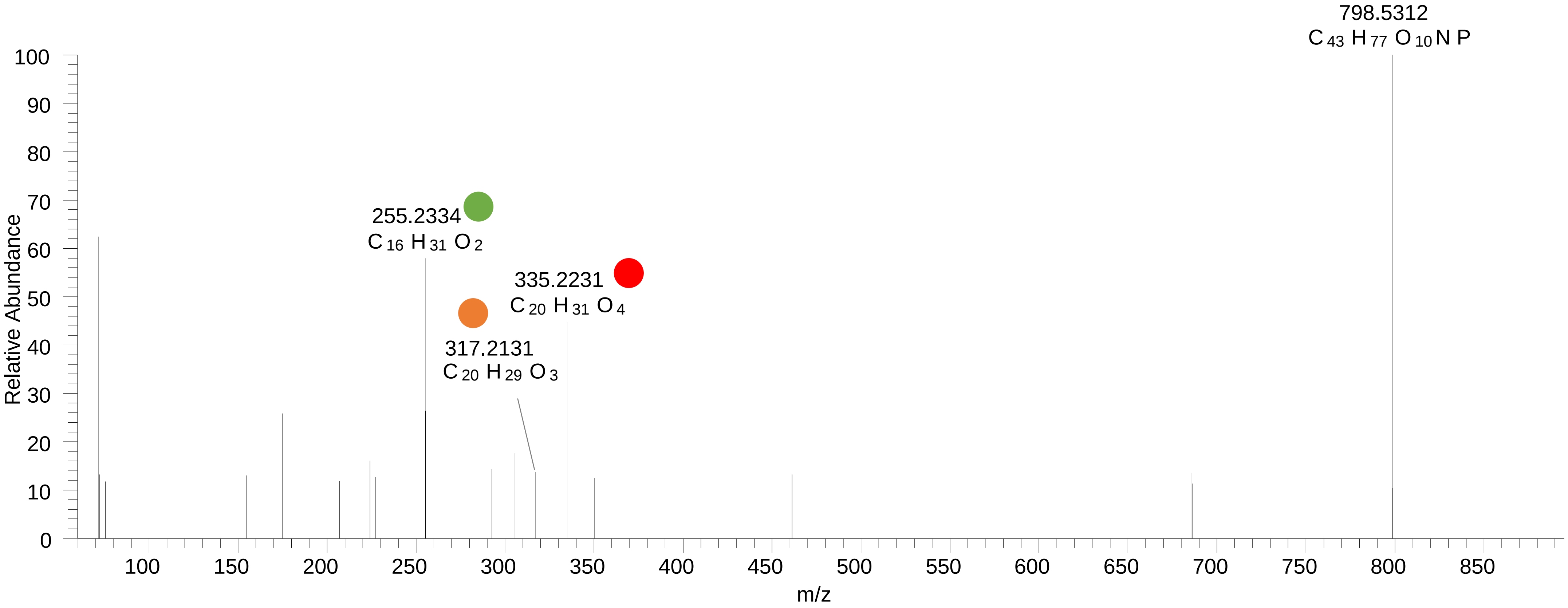

# PC(16:0_20:4<2O>)
# RT 12.0

**[oxPC+HCOO]- XIC 858.5502**

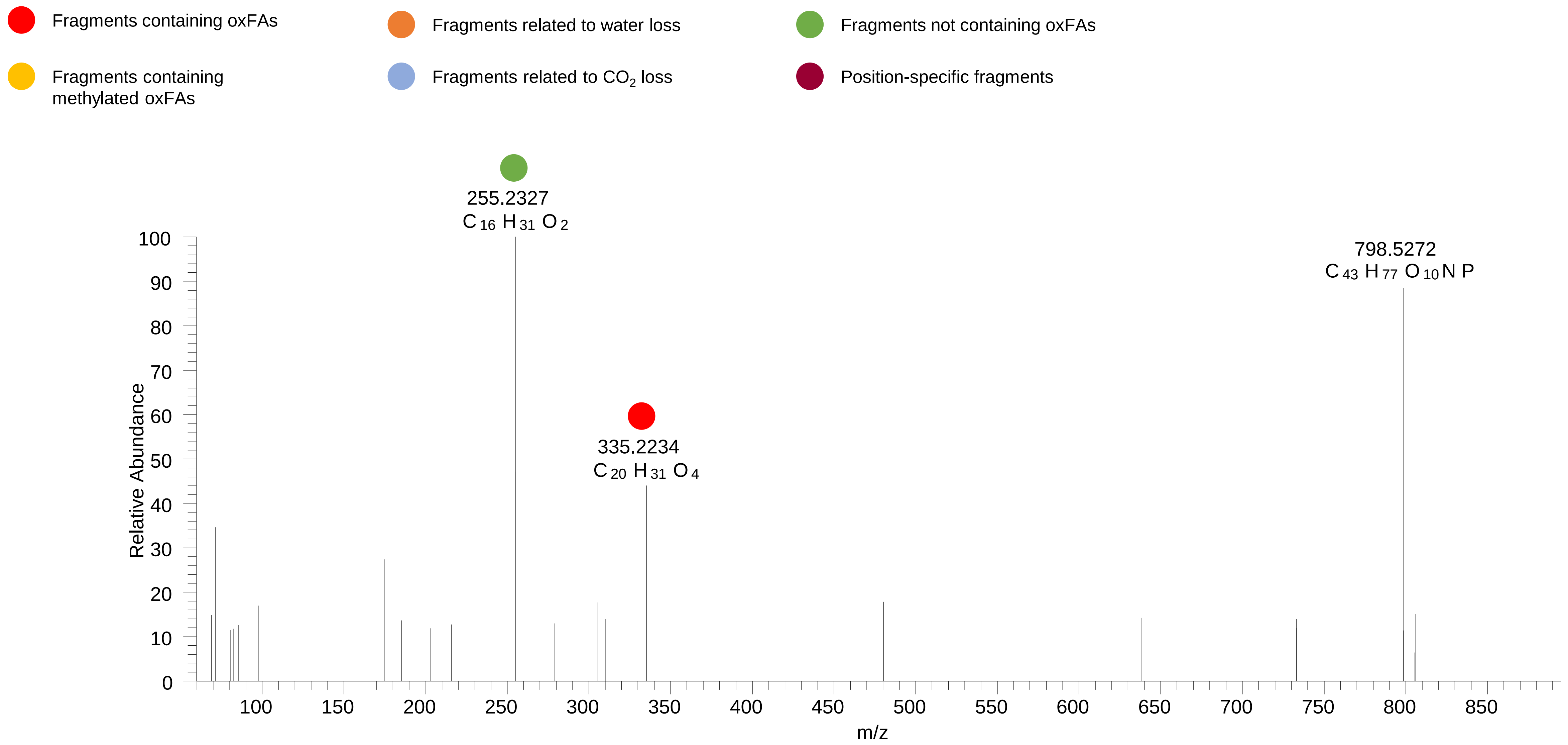

### PC(16:0_20:4<OOH>)
# RT 13.3

**[oxPC+HCOO]- XIC 858.5502**

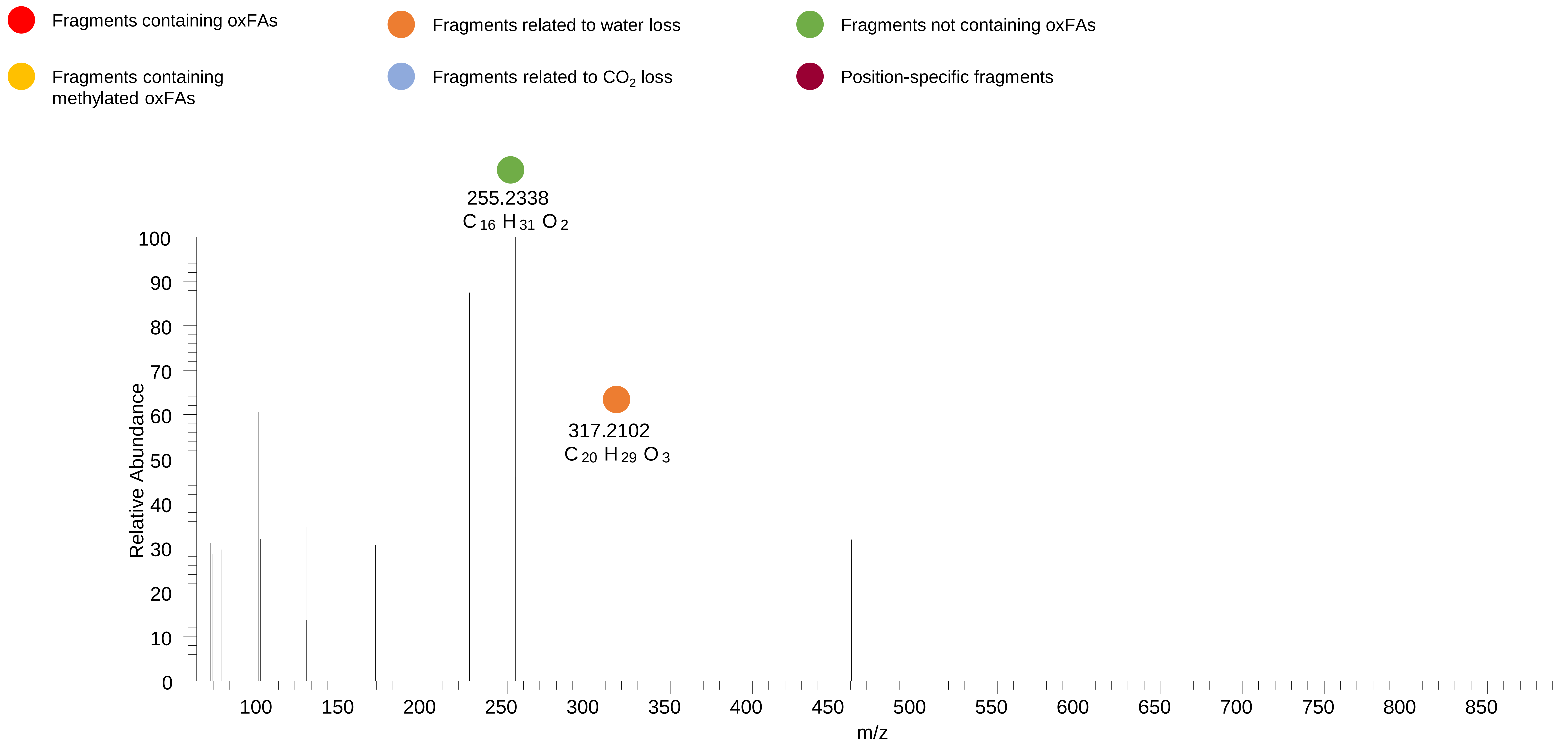

### PC(16:0_20:4<OOH{15}>)
# RT 13.5

**[oxPC+HCOO]- XIC 858.5502**

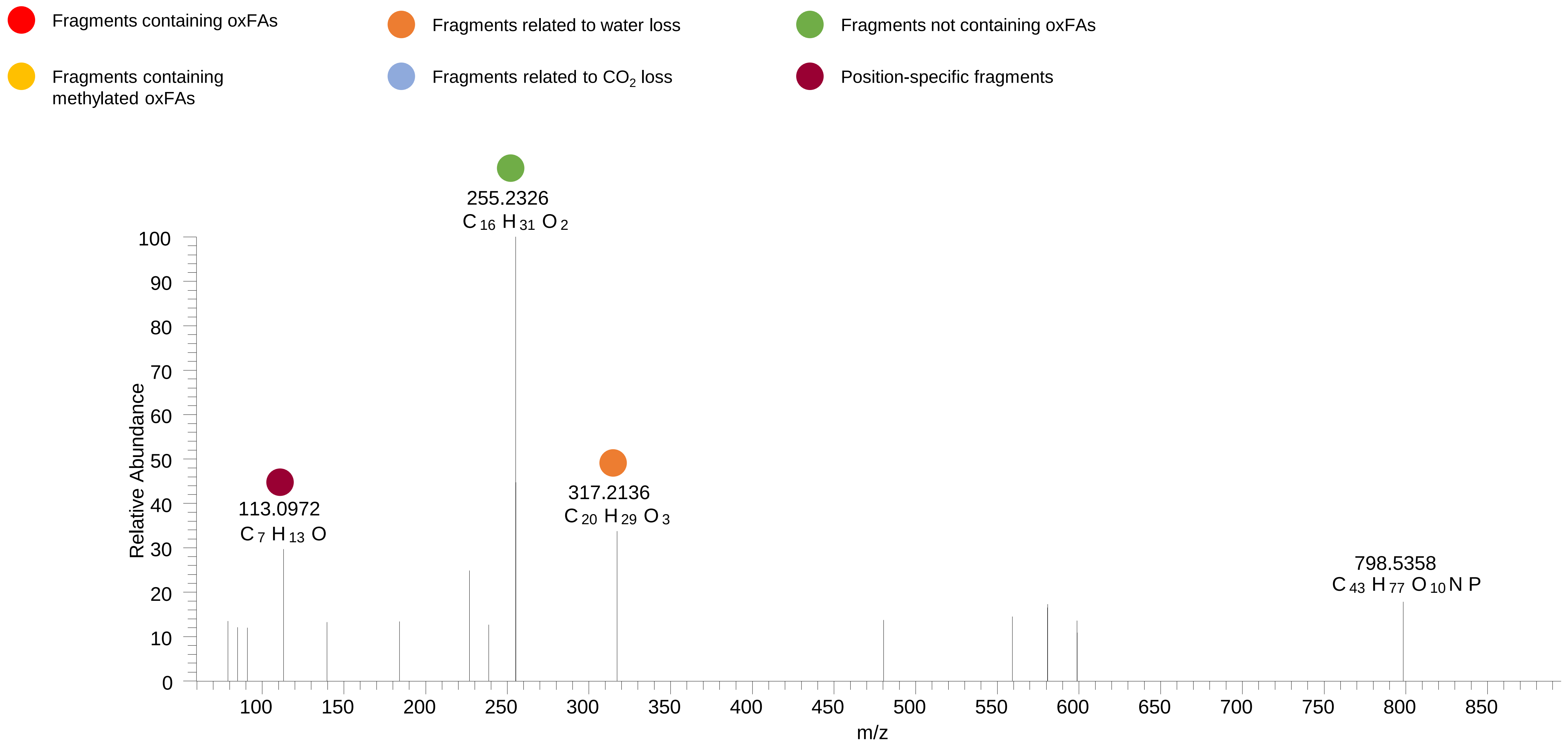

### PC(16:0_20:4<OOH{12}>)
# RT 13.9

**[oxPC+HCOO]- XIC 858.5502**

- Fragments containing oxFAs
- Fragments related to water loss
- Fragments not containing oxFAs
- Fragments containing methylated oxFAs
- Fragments related to $CO_2$ loss
- Position-specific fragments

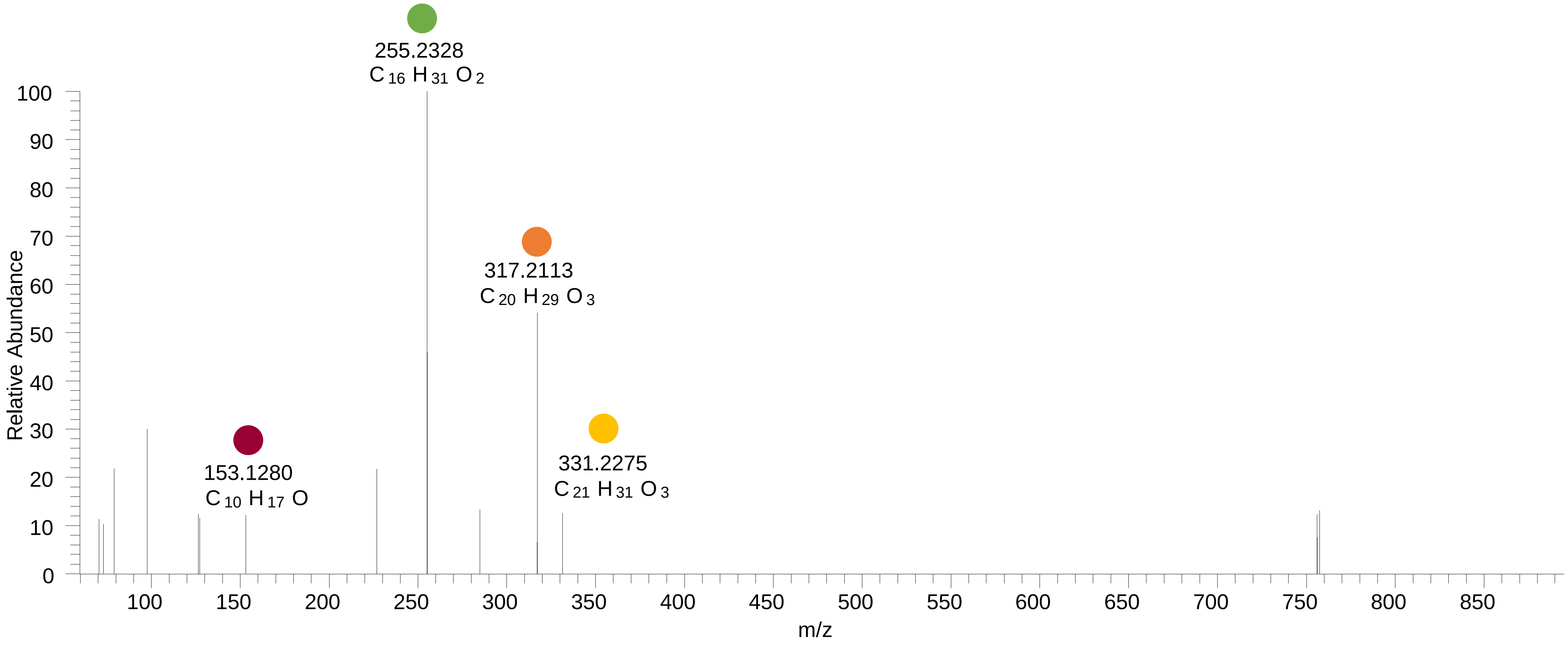

### PC(16:0_20:4<OOH{7}>)

# RT 14.4

**[oxPC+HCOO]-   XIC 858.5502**

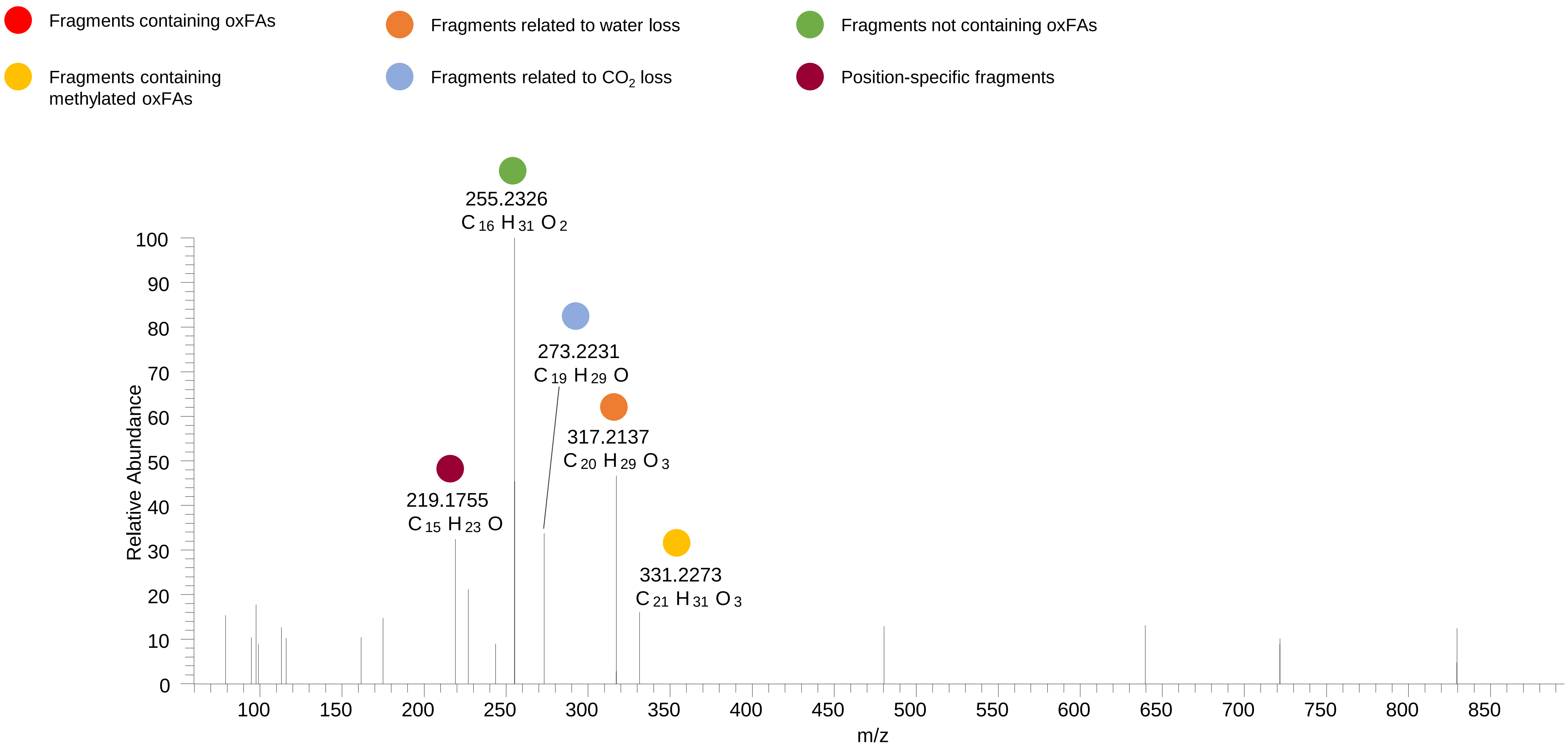

### PC(18:1_18:2<OOH>)
### PC(16:0_20:3<OOH>)
# RT 13.8

**[oxPC+HCOO]- XIC 860.5658**

- Fragments containing oxFAs
- Fragments related to water loss
- Fragments not containing oxFAs
- Fragments containing methylated oxFAs
- Fragments related to $CO_2$ loss
- Position-specific fragments

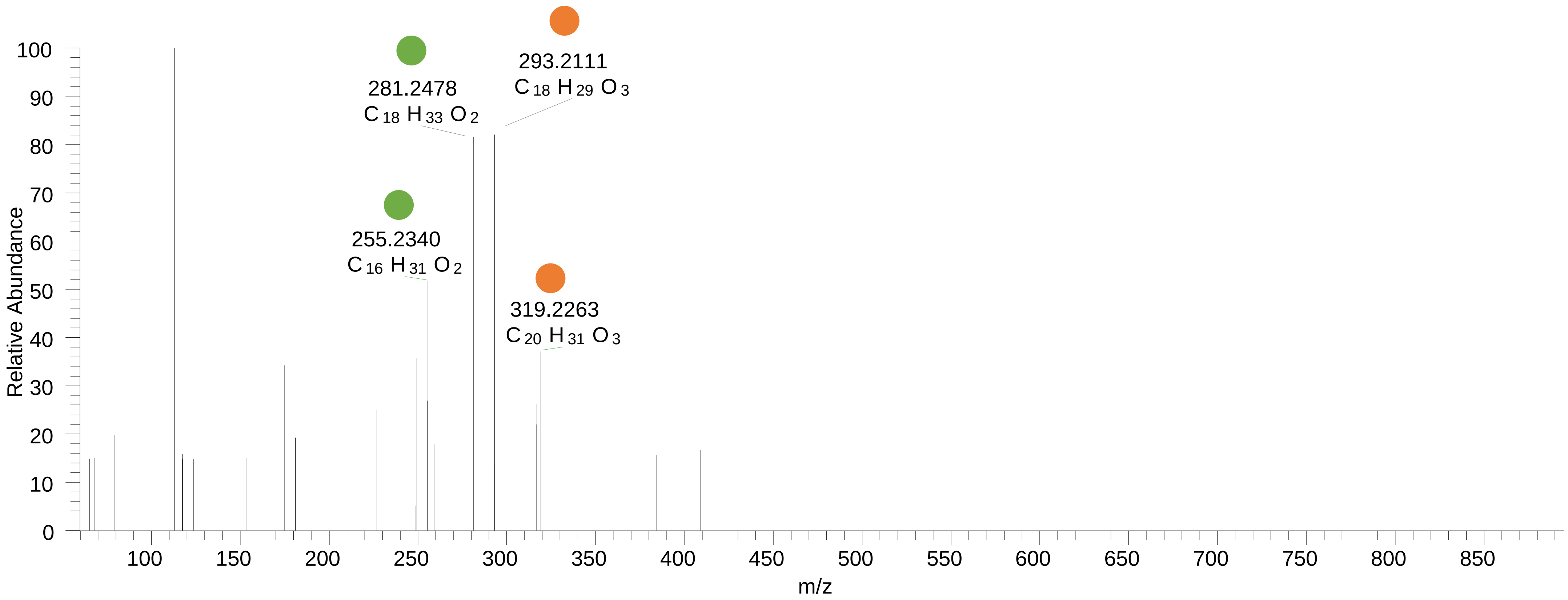

### PC(16:0_20:3<OOH>)
# RT 14.2

**[oxPC+HCOO]- XIC 860.5658**

Fragments containing oxFAs
Fragments related to water loss
Fragments not containing oxFAs
Fragments containing methylated oxFAs
Fragments related to $CO_2$ loss
Position-specific fragments

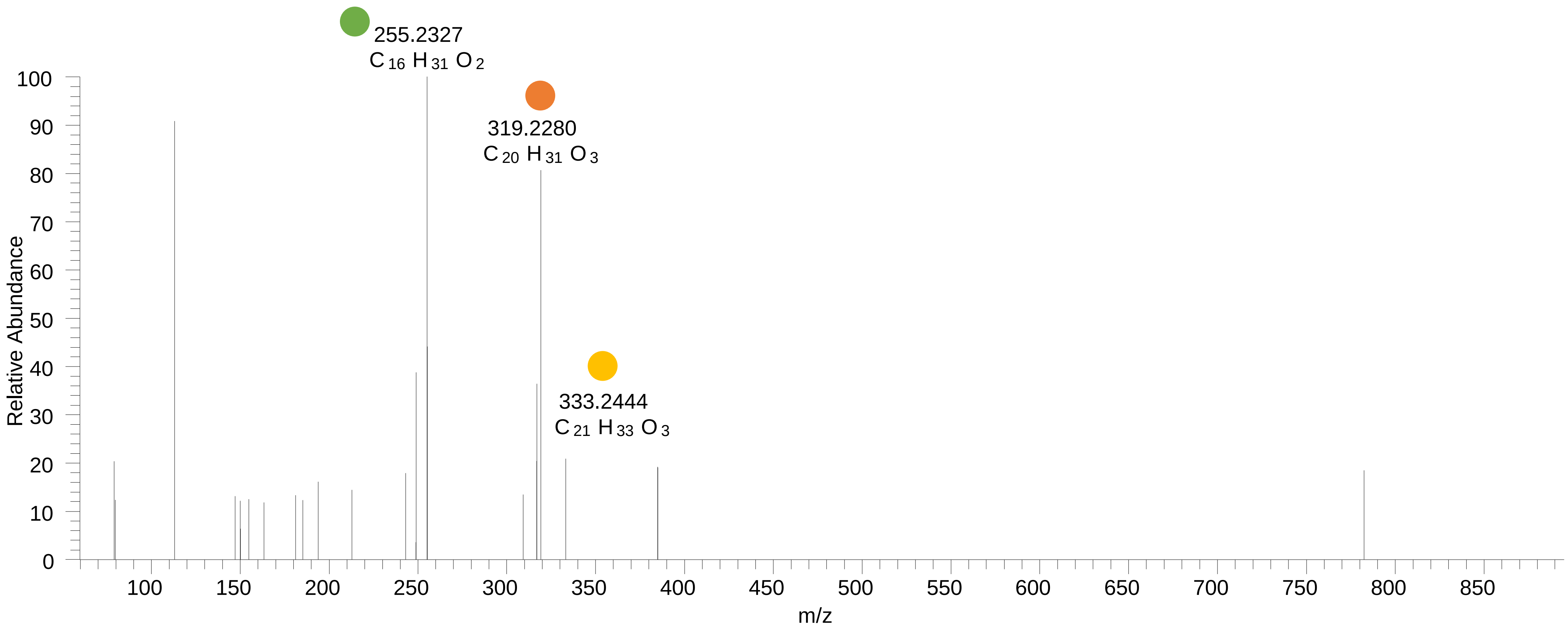

### PC(18:0_18:2<OOH>)
# RT 14.6

**[oxPC+HCOO]- XIC 862.5815**

- Fragments containing oxFAs
- Fragments related to water loss
- Fragments not containing oxFAs
- Fragments containing methylated oxFAs
- Fragments related to $CO_2$ loss
- Position-specific fragments

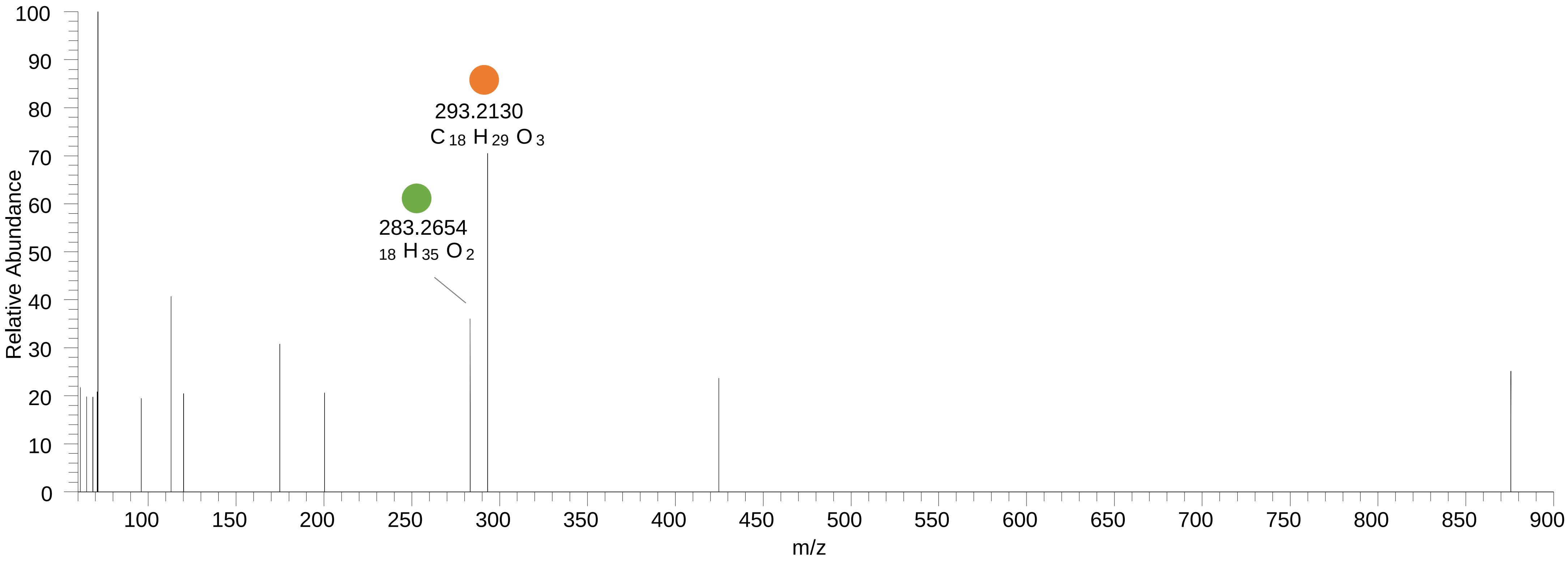

### PC(18:0_18:2<OOH{13}>)
# RT 15.1

**[oxPC+HCOO]- XIC 862.5815**

- Fragments containing oxFAs
- Fragments related to water loss
- Fragments not containing oxFAs
- Fragments containing methylated oxFAs
- Fragments related to $CO_2$ loss
- Position-specific fragments

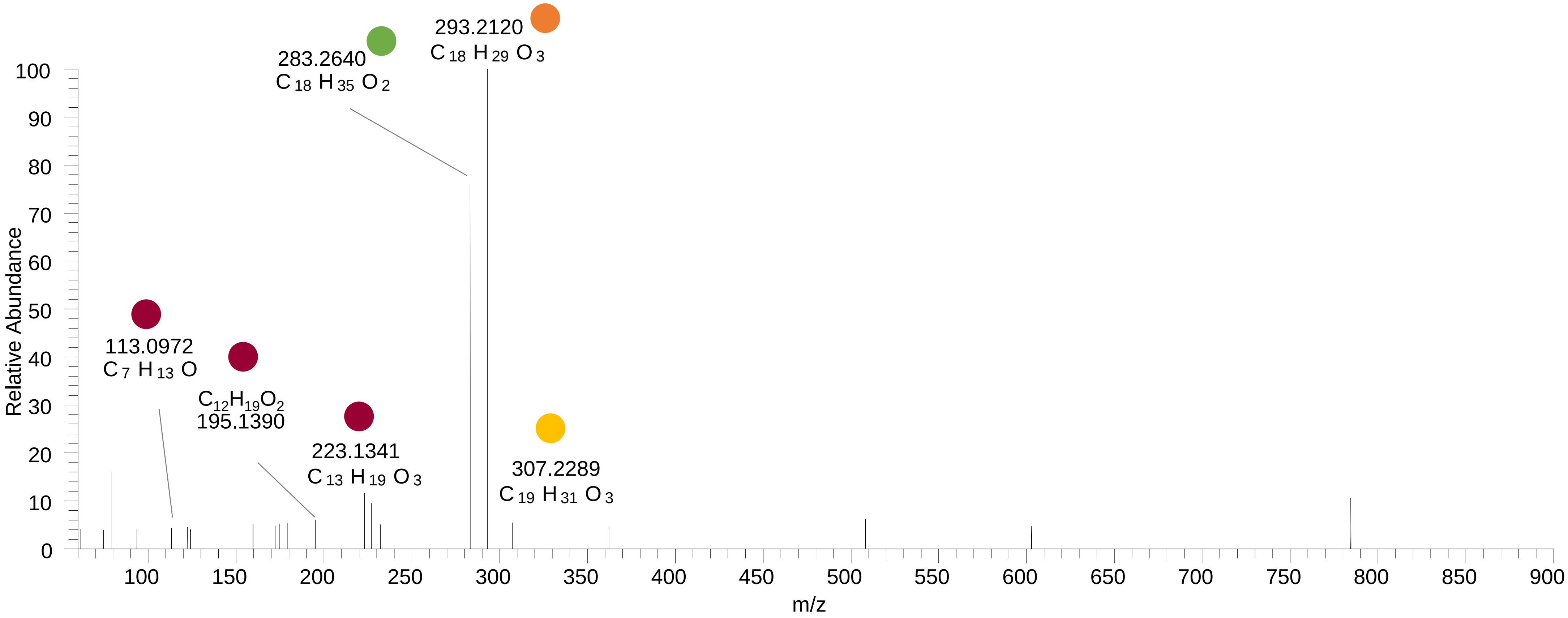

# PC(16:0_22:6<O{16}>)
# RT 13.5

**[oxPC+HCOO]- XIC 866.5553**

- Fragments containing oxFAs
- Fragments related to water loss
- Fragments not containing oxFAs
- Fragments containing methylated oxFAs
- Fragments related to $CO_2$ loss
- Position-specific fragments

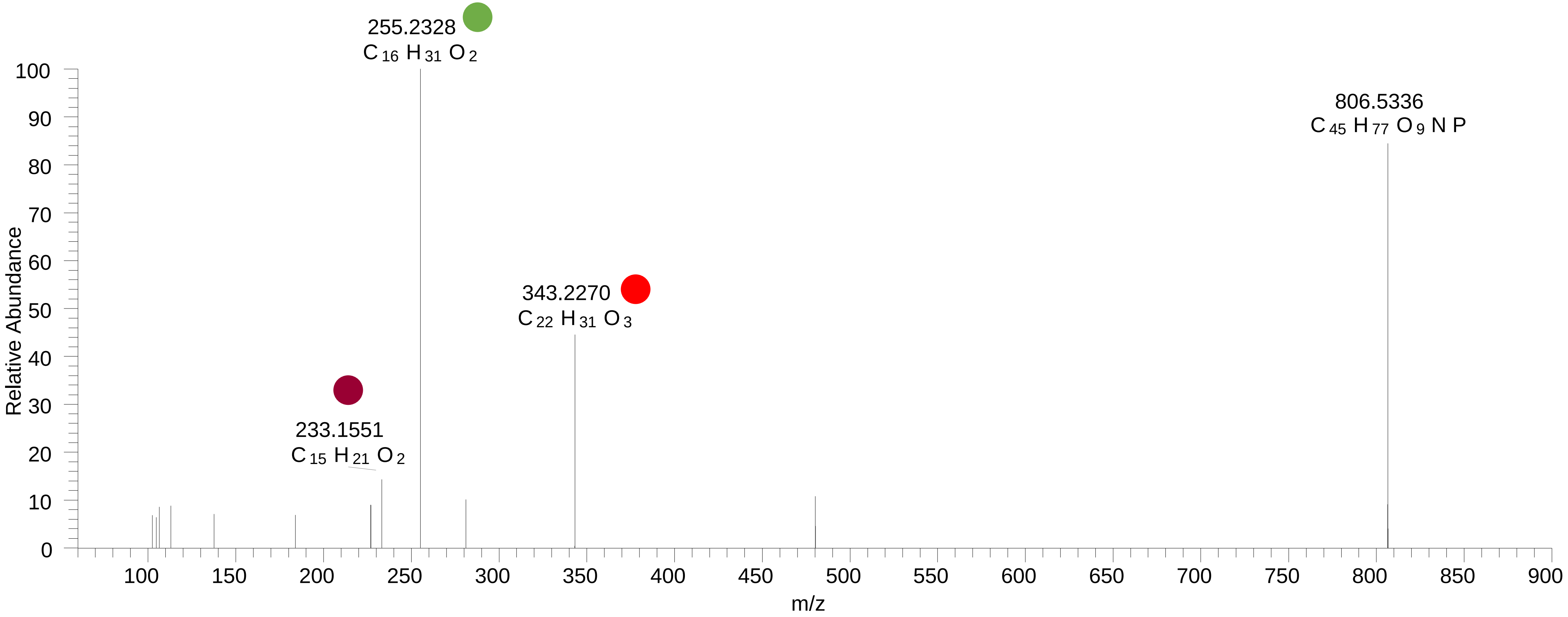

# PC(16:0_22:5<ep{7-8}>)
# RT 14.5

**[oxPC+HCOO]- XIC 866.5553**

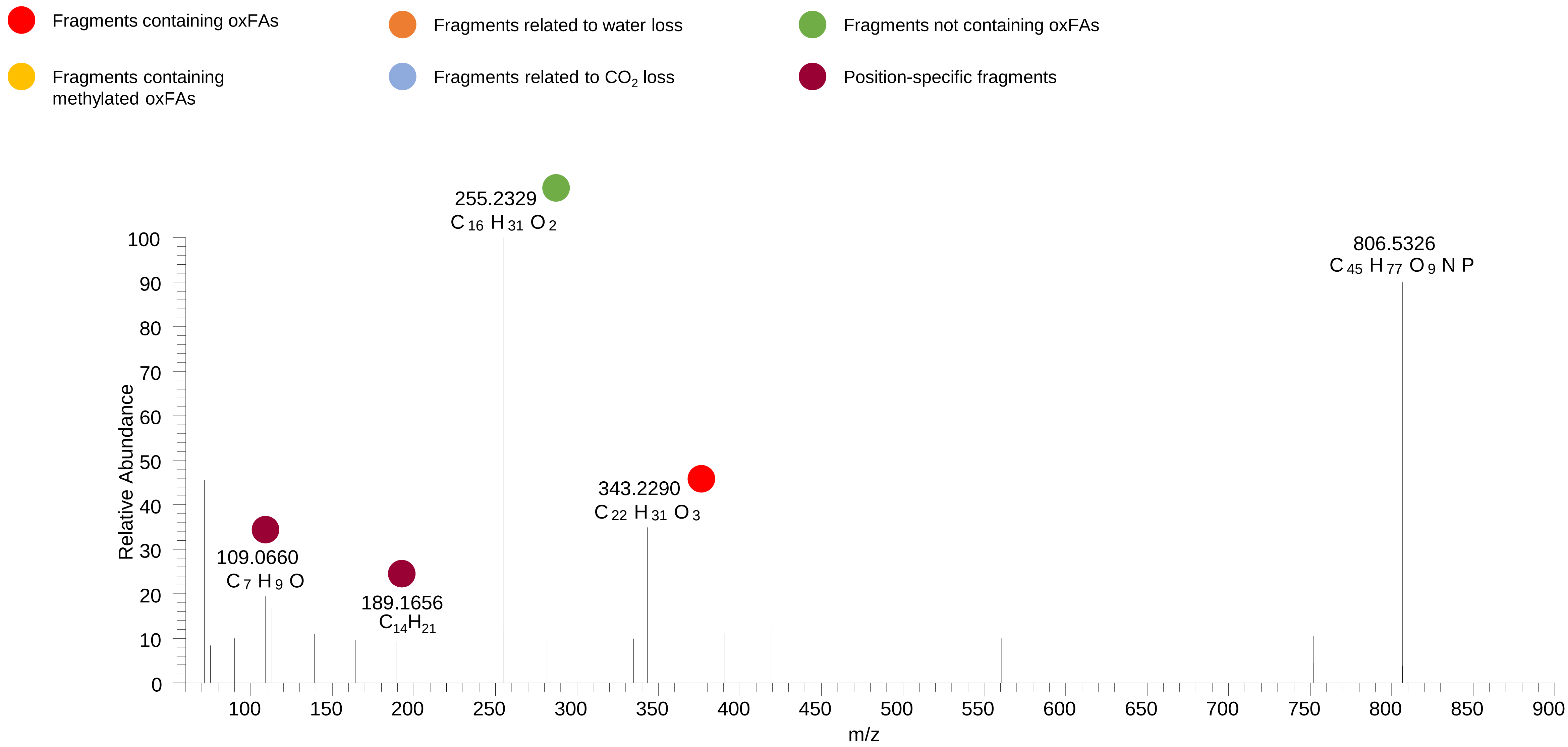

# PC(16:0_22:5<OH>)
# RT 13.7

**[oxPC+HCOO]- XIC 868.5709**

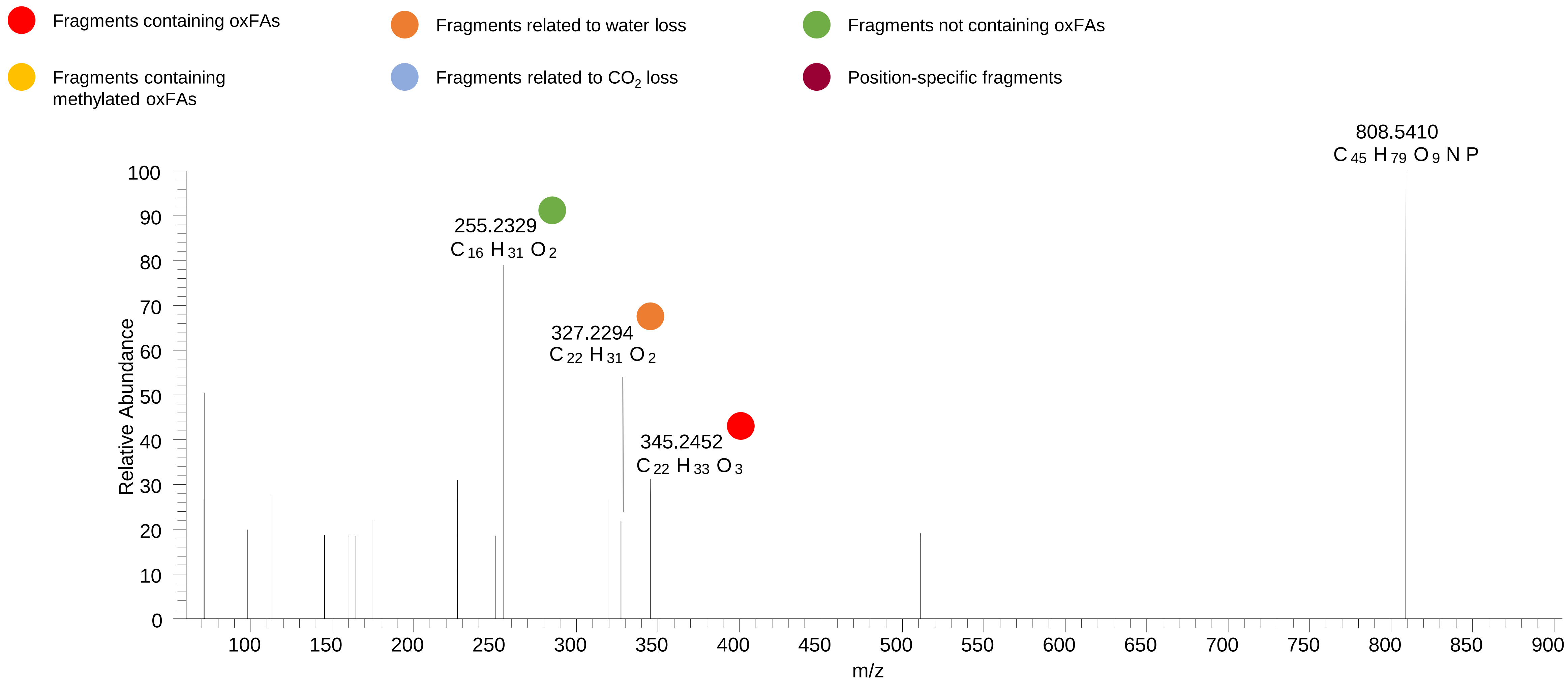

# PC(16:0_22:5<OH>)
# PC(18:1_20:4<O>)
# RT 13.8-14.5

**[oxPC+HCOO]- XIC 868.5709**

- Fragments containing oxFAs
- Fragments related to water loss
- Fragments not containing oxFAs
- Fragments containing methylated oxFAs
- Fragments related to $CO_2$ loss
- Position-specific fragments

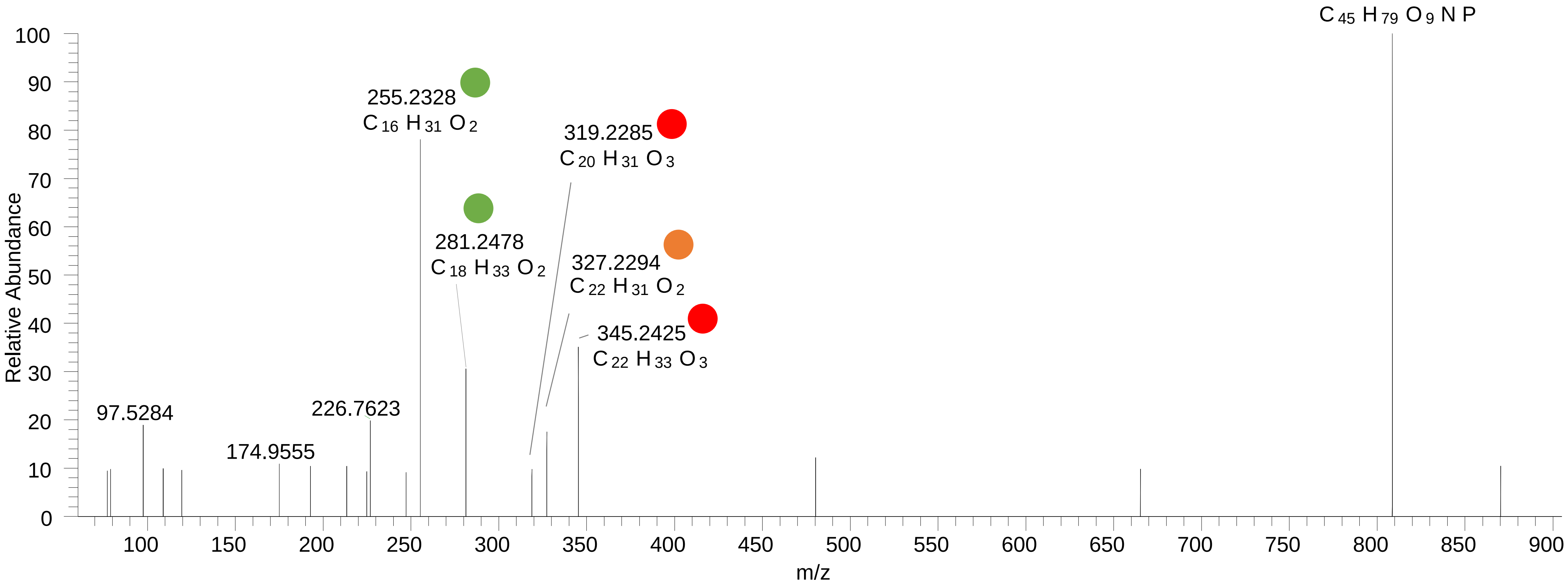

# PC(18:0_20:5<O>)
# PC(18:1_20:4<O>)
# RT 14.7-15.0

**[oxPC+HCOO]- XIC 868.5709**

- Fragments containing oxFAs
- Fragments related to water loss
- Fragments not containing oxFAs
- Fragments containing methylated oxFAs
- Fragments related to $CO_2$ loss
- Position-specific fragments

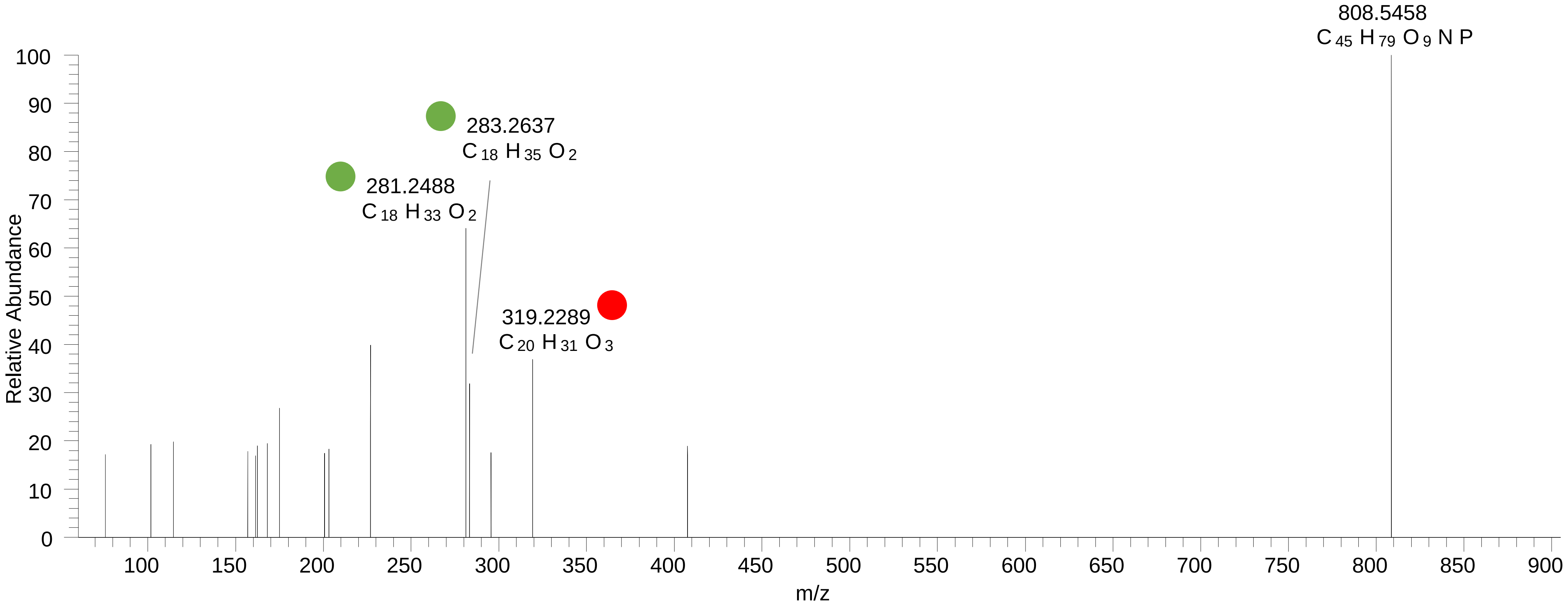

# PC(18:1_20:3<O>)
# RT 14.4

**[oxPC+HCOO]- XIC 870.5866**

Fragments containing oxFAs

Fragments related to water loss

Fragments not containing oxFAs

Fragments containing methylated oxFAs

Fragments related to $CO_2$ loss

Position-specific fragments

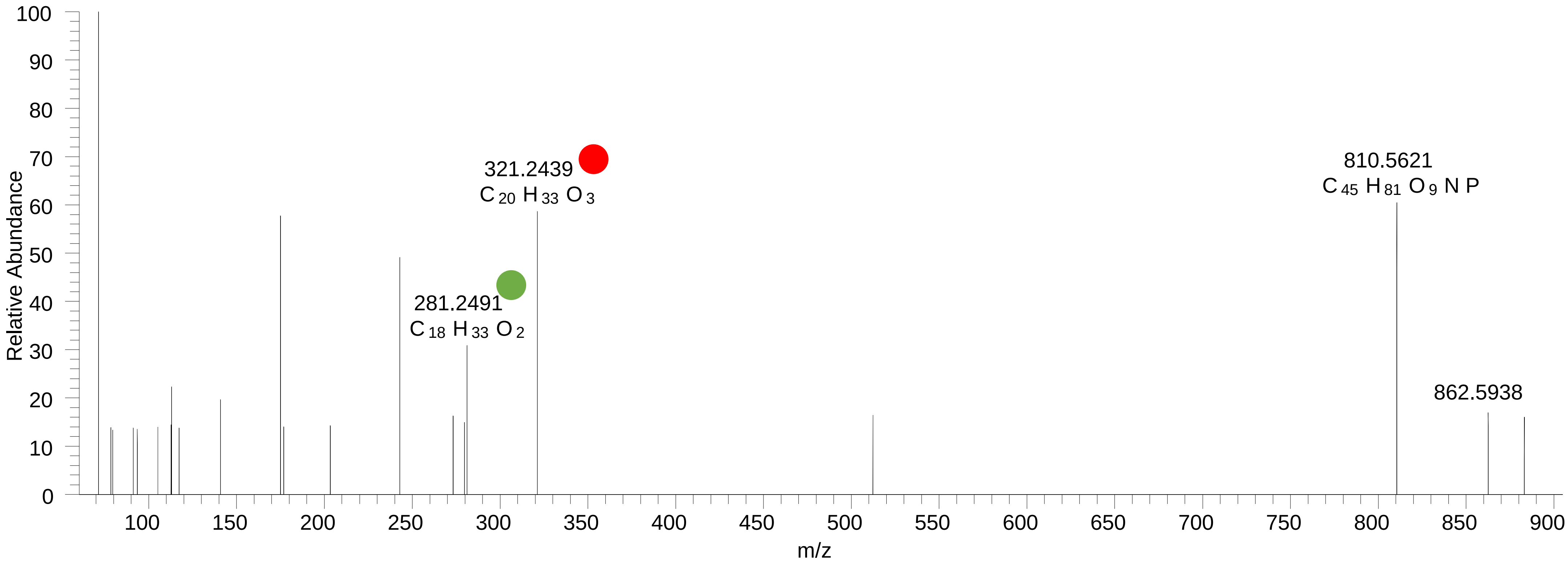

# PC(18:0_20:4<OH>)
# RT 15.3

**[oxPC+HCOO]- XIC 870.5866**

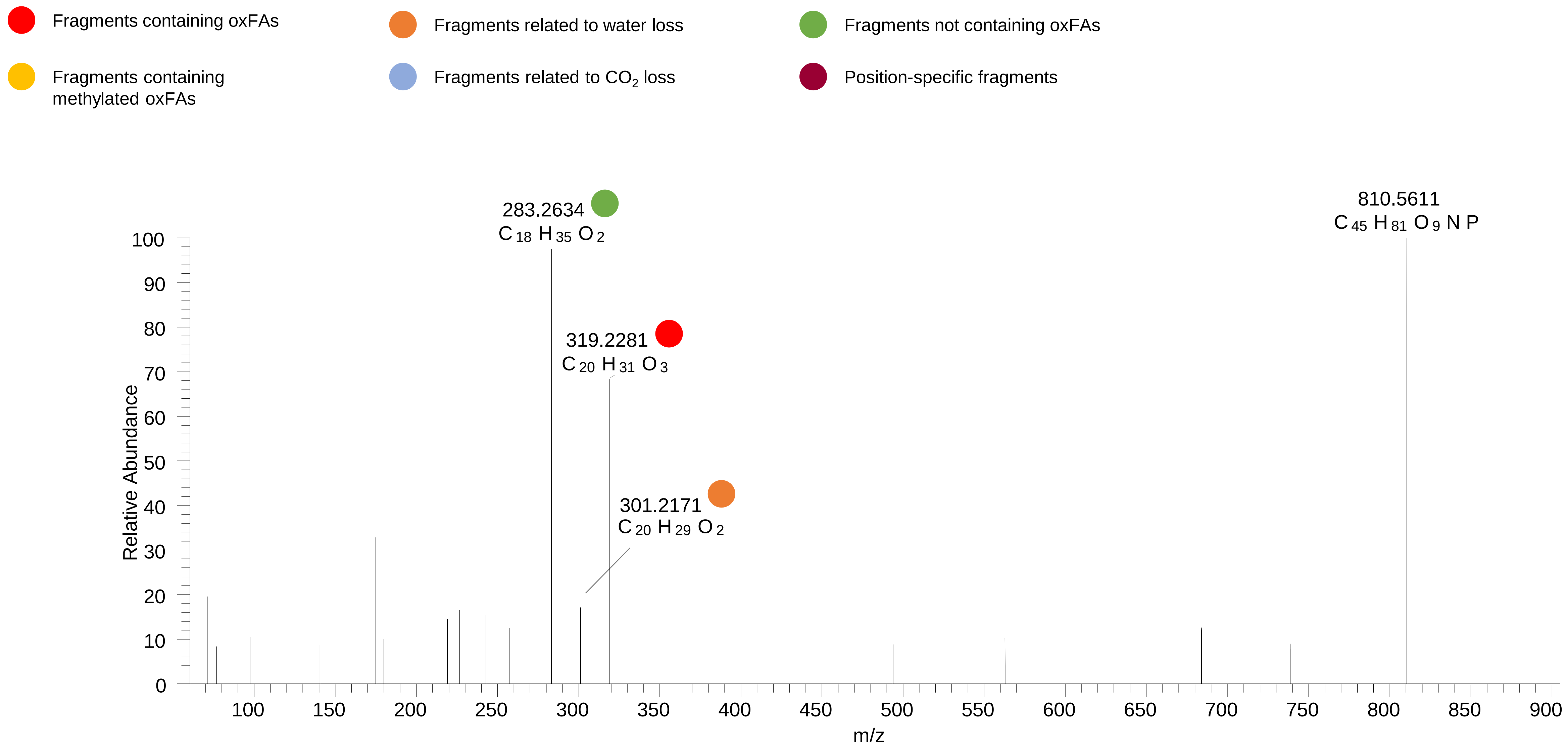

# PC(18:0_20:4<OH{11}>)
# RT 15.7

**[oxPC+HCOO]- XIC 870.5866**

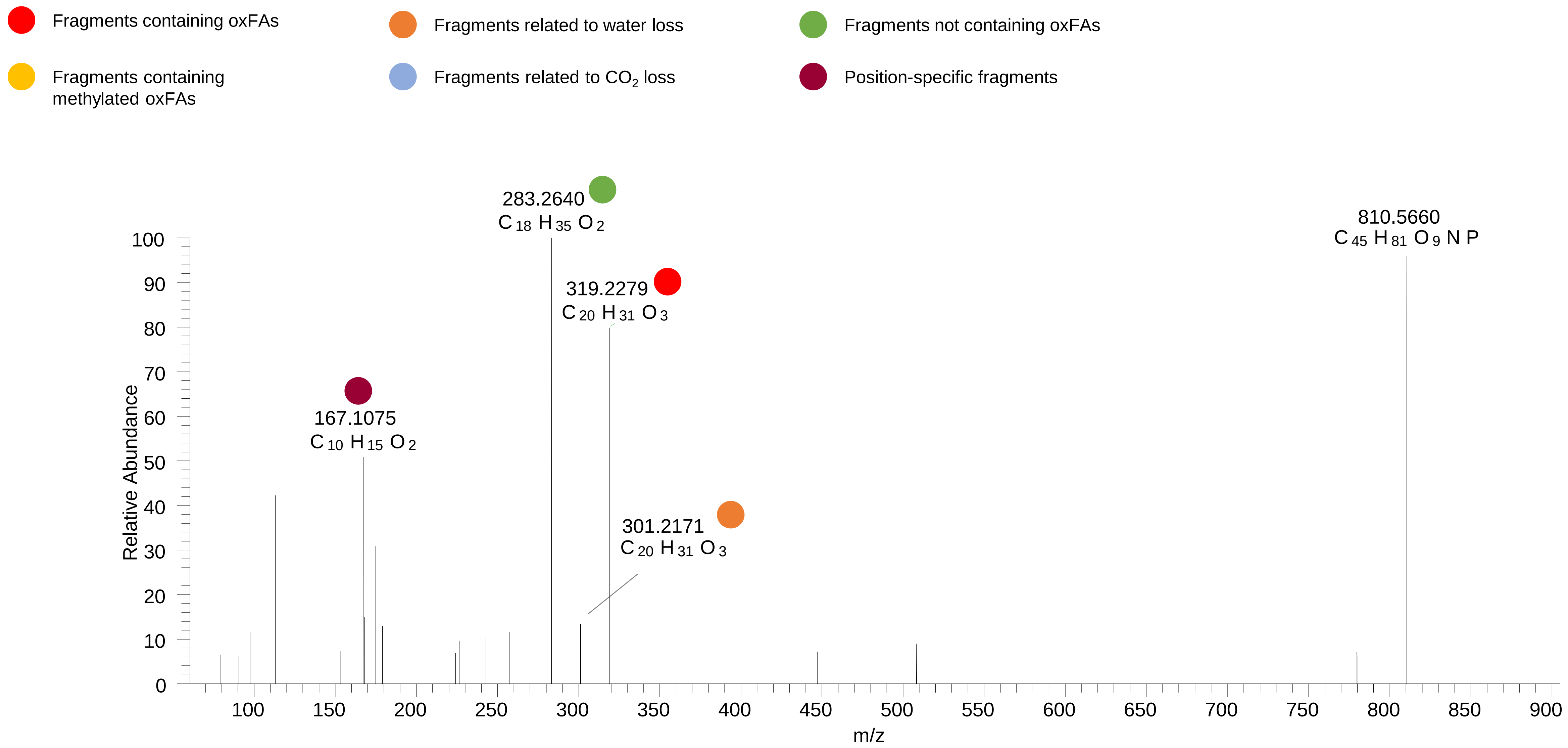

# PC(18:0_20:4<OH{8}>)
# PC(18:0_20:4<OH{9}>)
# RT 16.2

**[oxPC+HCOO]- XIC 870.5866**

- Fragments containing oxFAs
- Fragments related to water loss
- Fragments not containing oxFAs
- Fragments containing methylated oxFAs
- Fragments related to $CO_2$ loss
- Position-specific fragments

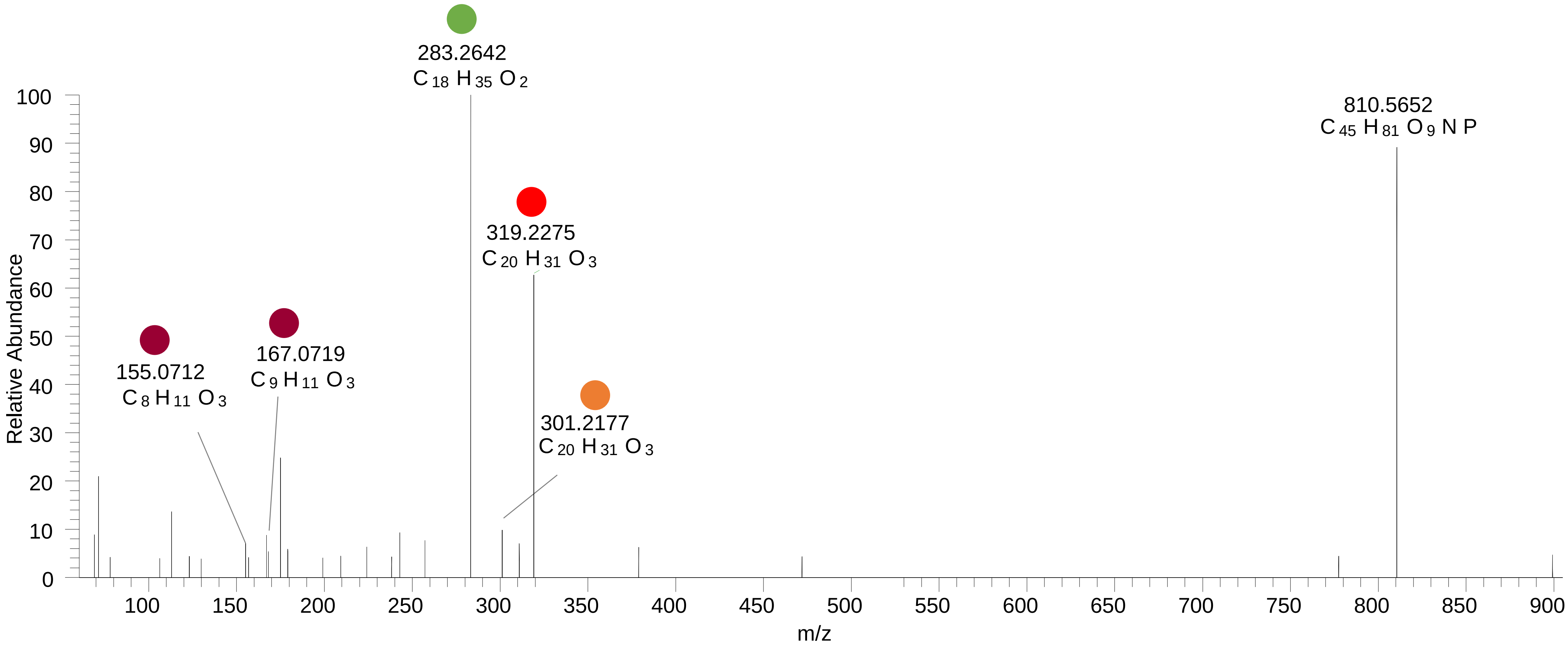

# PC(16:0_22:6<2O>)
# RT 10.8-14.4

**[oxPC+HCOO]- XIC 882.5502**

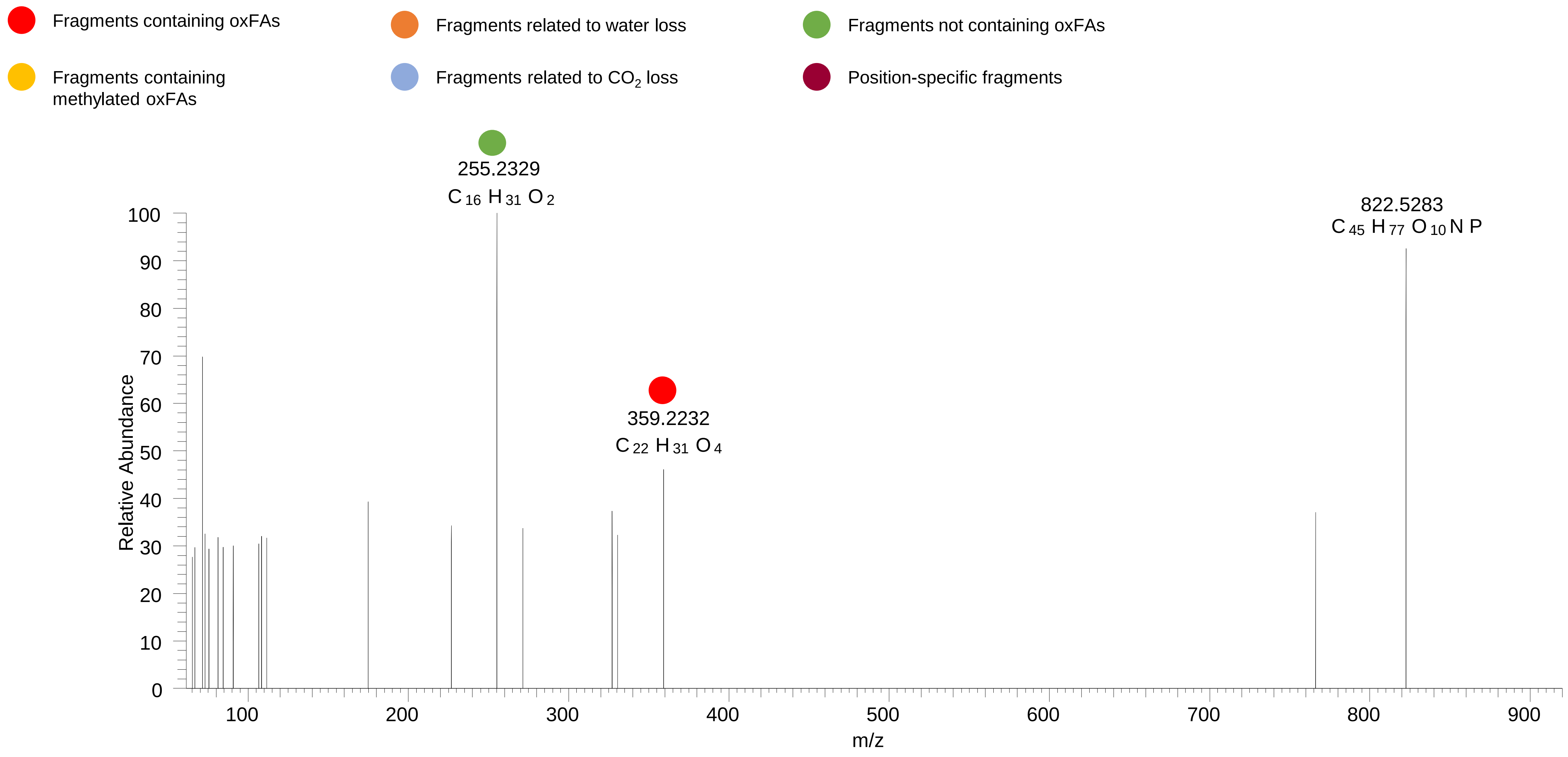

# PC(18:0_20:4<2O>)
# RT 13.1

**[oxPC+HCOO]- XIC 886.5815**

- Fragments containing oxFAs
- Fragments related to water loss
- Fragments not containing oxFAs
- Fragments containing methylated oxFAs
- Fragments related to $CO_2$ loss
- Position-specific fragments

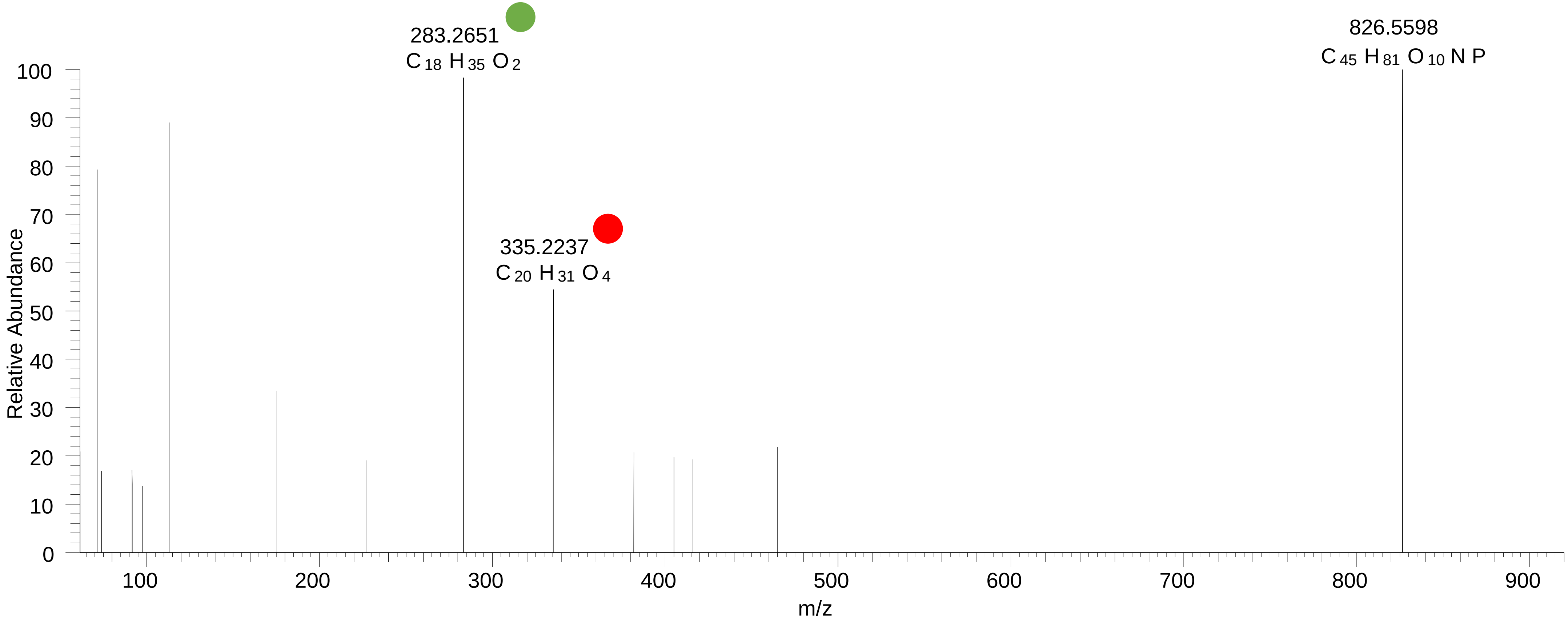

### PC(18:0_20:4<OOH{15}>)
# RT 15.3

**[oxPC+HCOO]-   XIC 886.5815**

- Fragments containing oxFAs
- Fragments related to water loss
- Fragments not containing oxFAs
- Fragments containing methylated oxFAs
- Fragments related to $CO_2$ loss
- Position-specific fragments

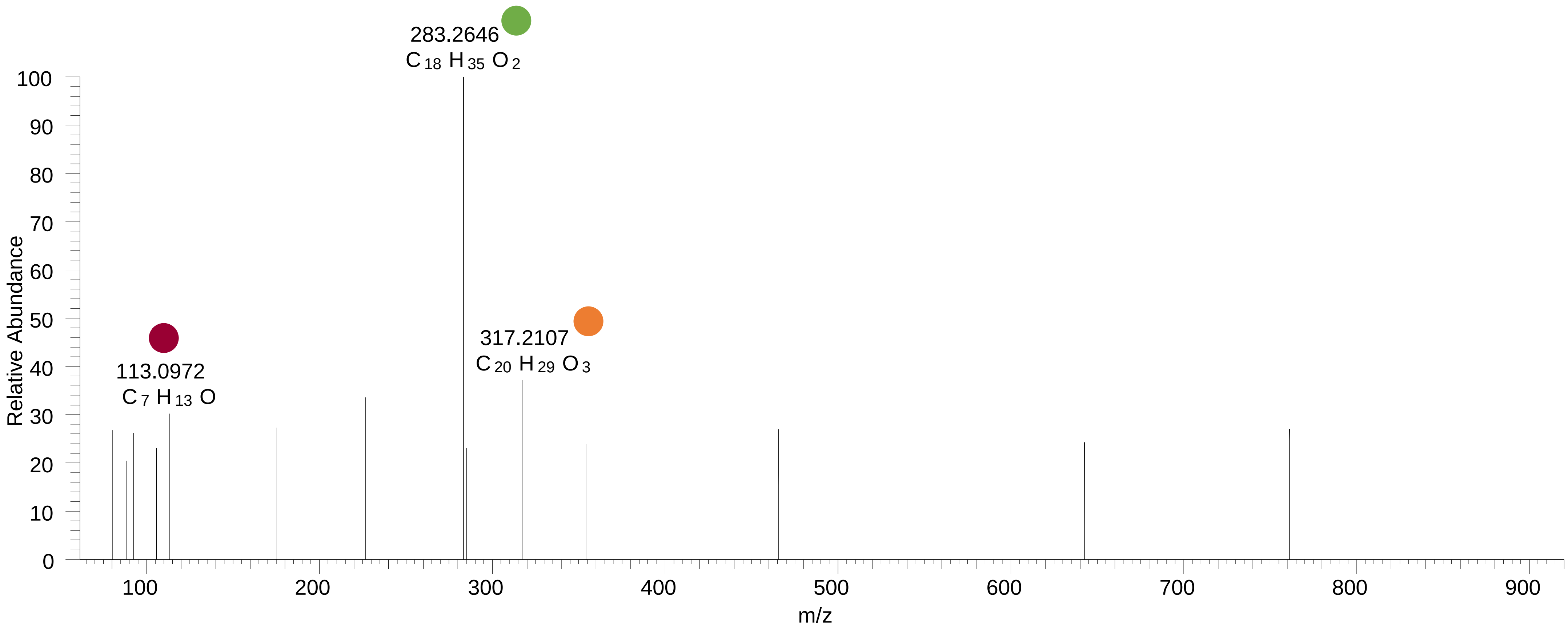

### PC(18:0_20:4<OOH>)
# RT 15.6

**[oxPC+HCOO]- XIC 886.5815**

Fragments containing oxFAs

Fragments related to water loss

Fragments not containing oxFAs

Fragments containing methylated oxFAs

Fragments related to $CO_2$ loss

Position-specific fragments

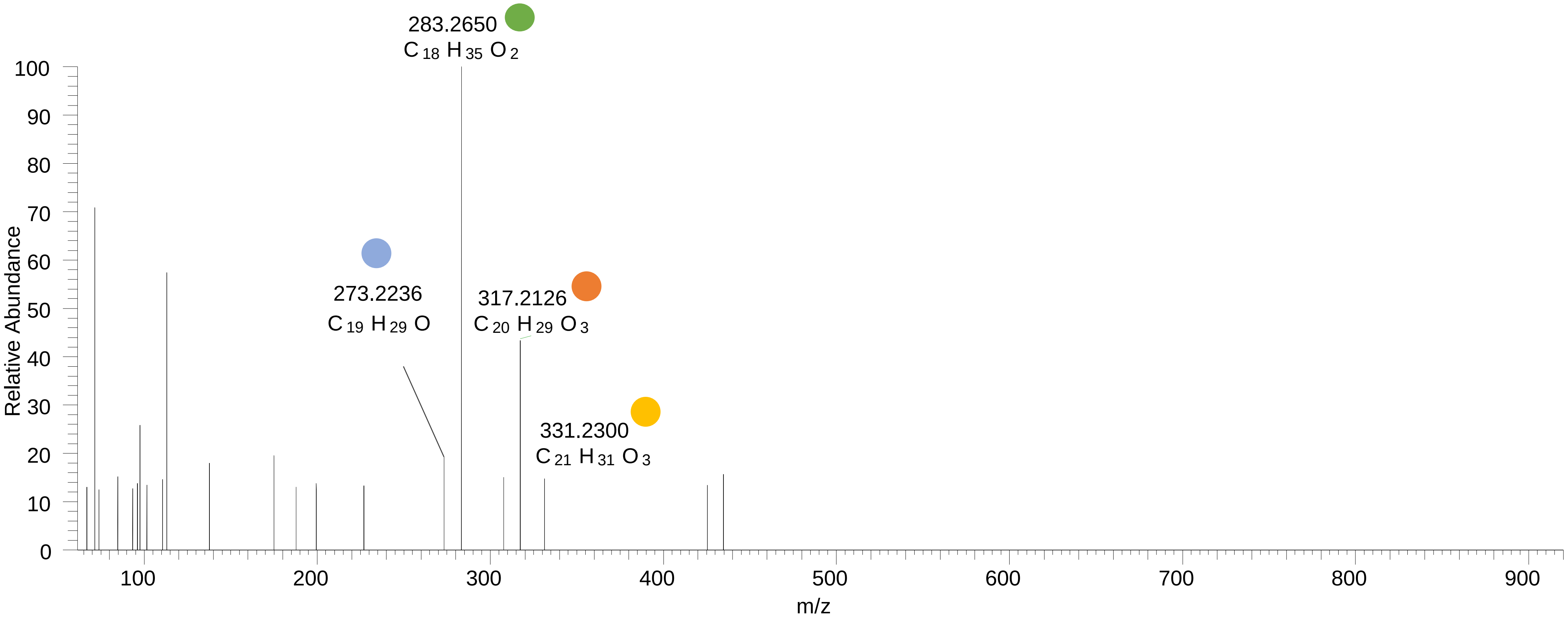

### PC(18:0_20:4<OOH{7}>)
# RT 16.0

**[oxPC+HCOO]- XIC 886.5815**

- Fragments containing oxFAs
- Fragments related to water loss
- Fragments not containing oxFAs
- Fragments containing methylated oxFAs
- Fragments related to $CO_2$ loss
- Position-specific fragments

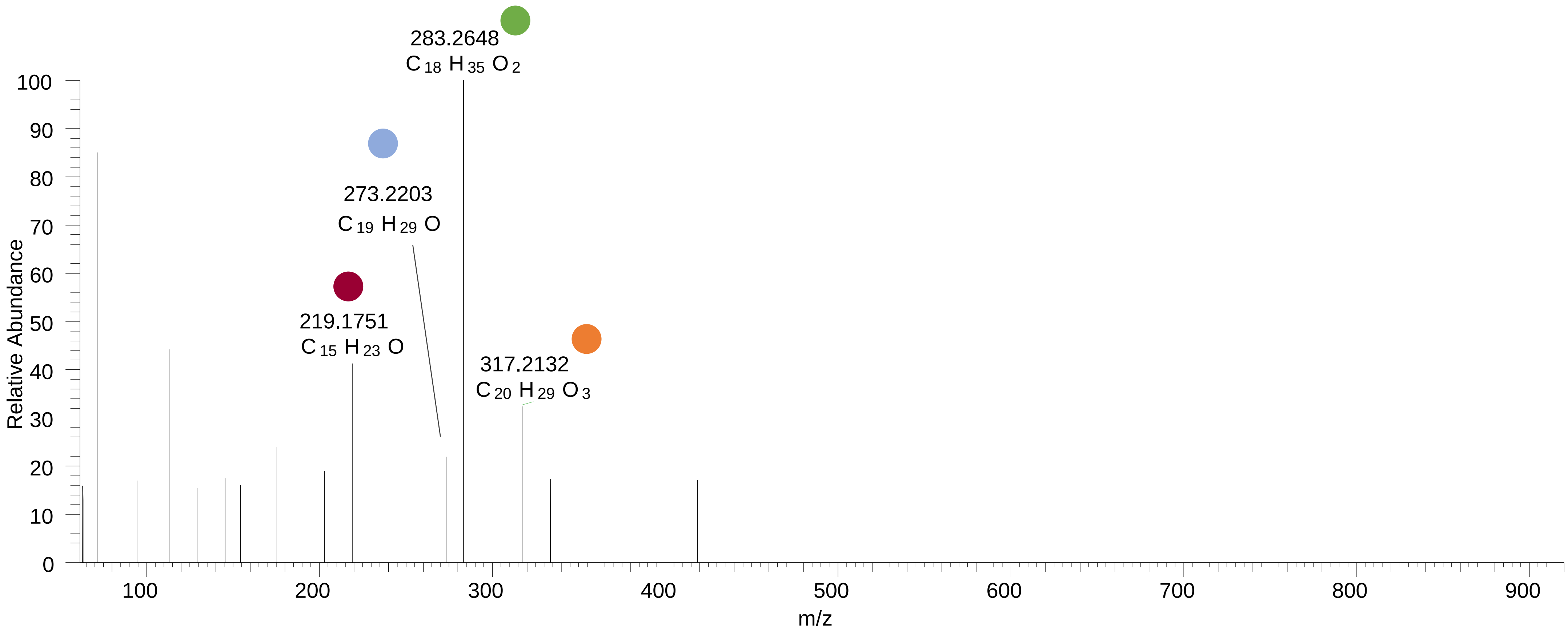

# PC(18:0_22:6<O>)
# RT 15.2-16.2

**[oxPC+HCOO]-   XIC 894.5866**

- Fragments containing oxFAs
- Fragments related to water loss
- Fragments not containing oxFAs
- Fragments containing methylated oxFAs
- Fragments related to $CO_2$ loss
- Position-specific fragments

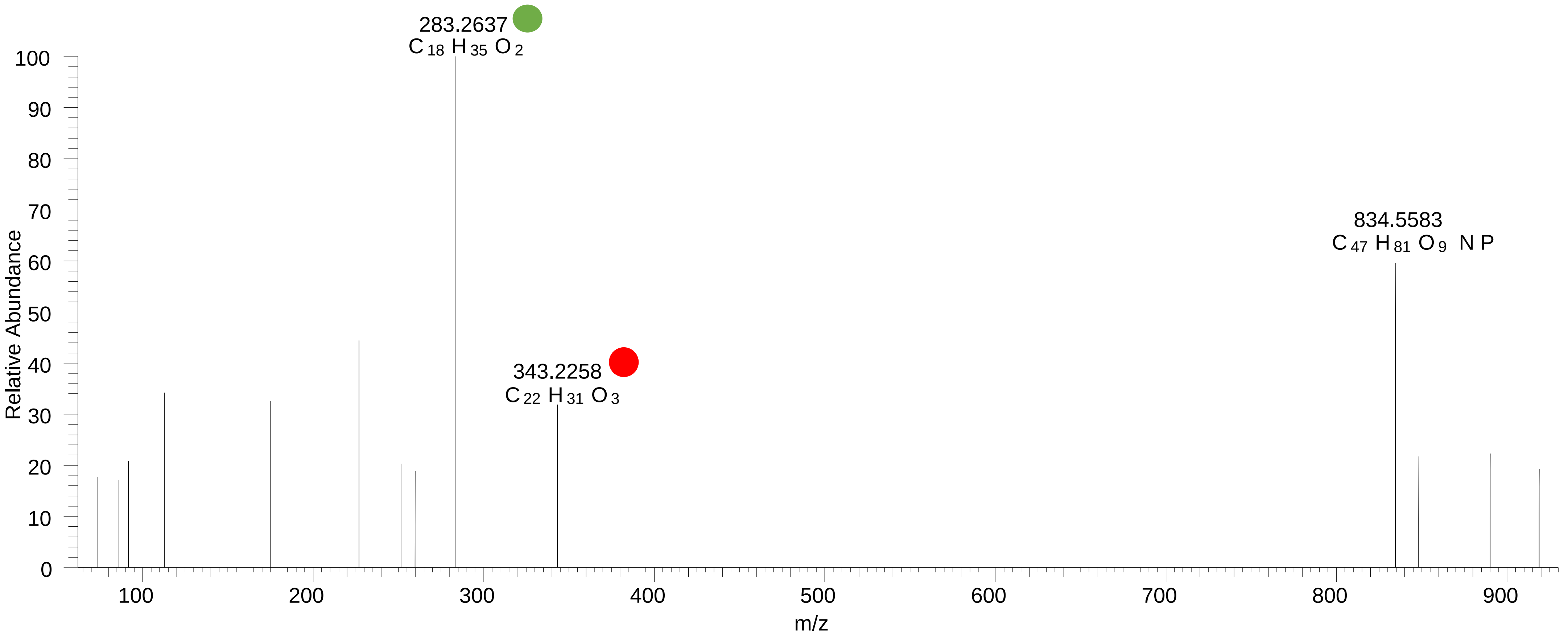

### OT2D_pool

# PC(16:1_18:2<O>)
# RT 11.9-12.0

**[oxPC+HCOO]- XIC 816.5396**

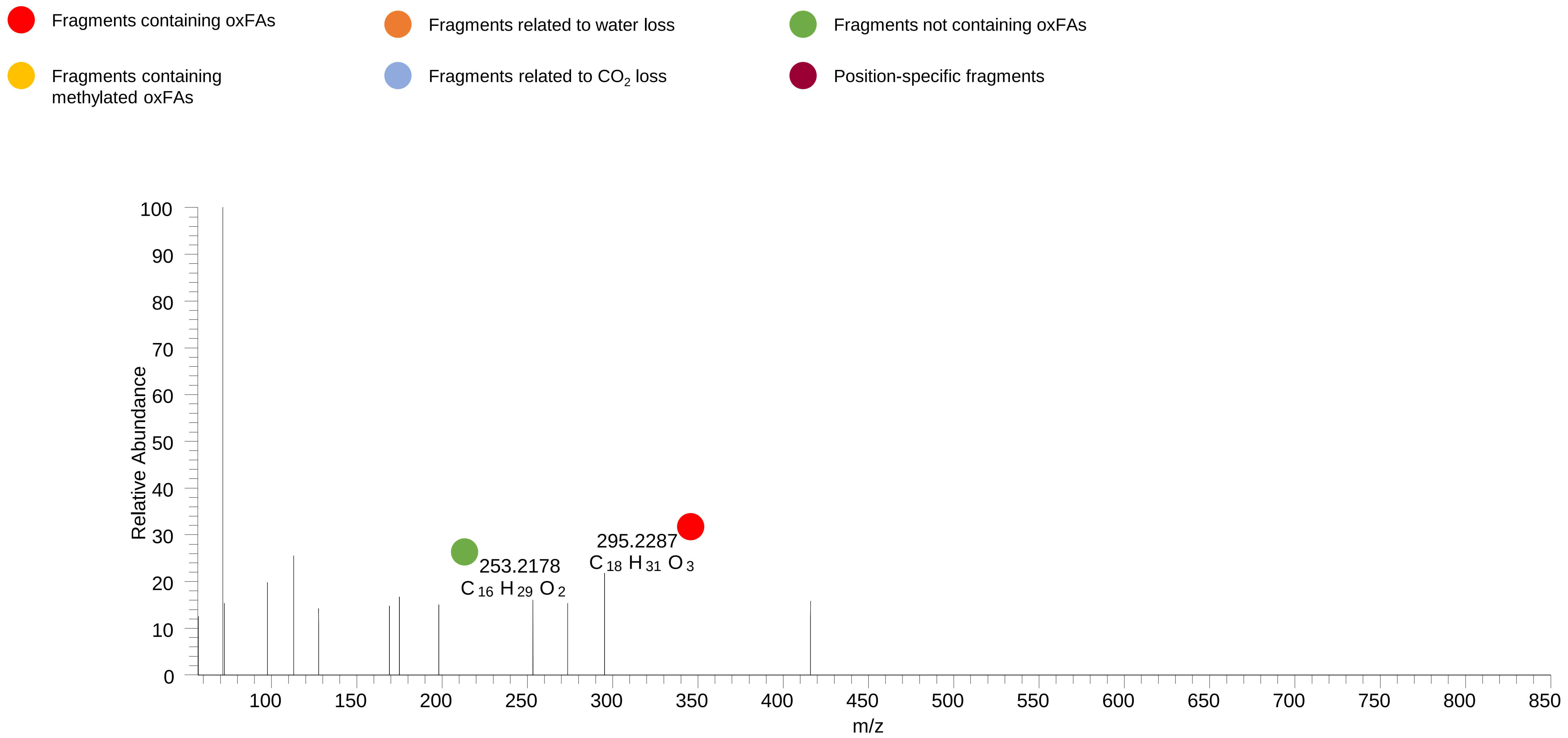

# PC(16:0_18:2<OH>)
# RT 12.6

**[oxPC+HCOO]- XIC 818.5553**

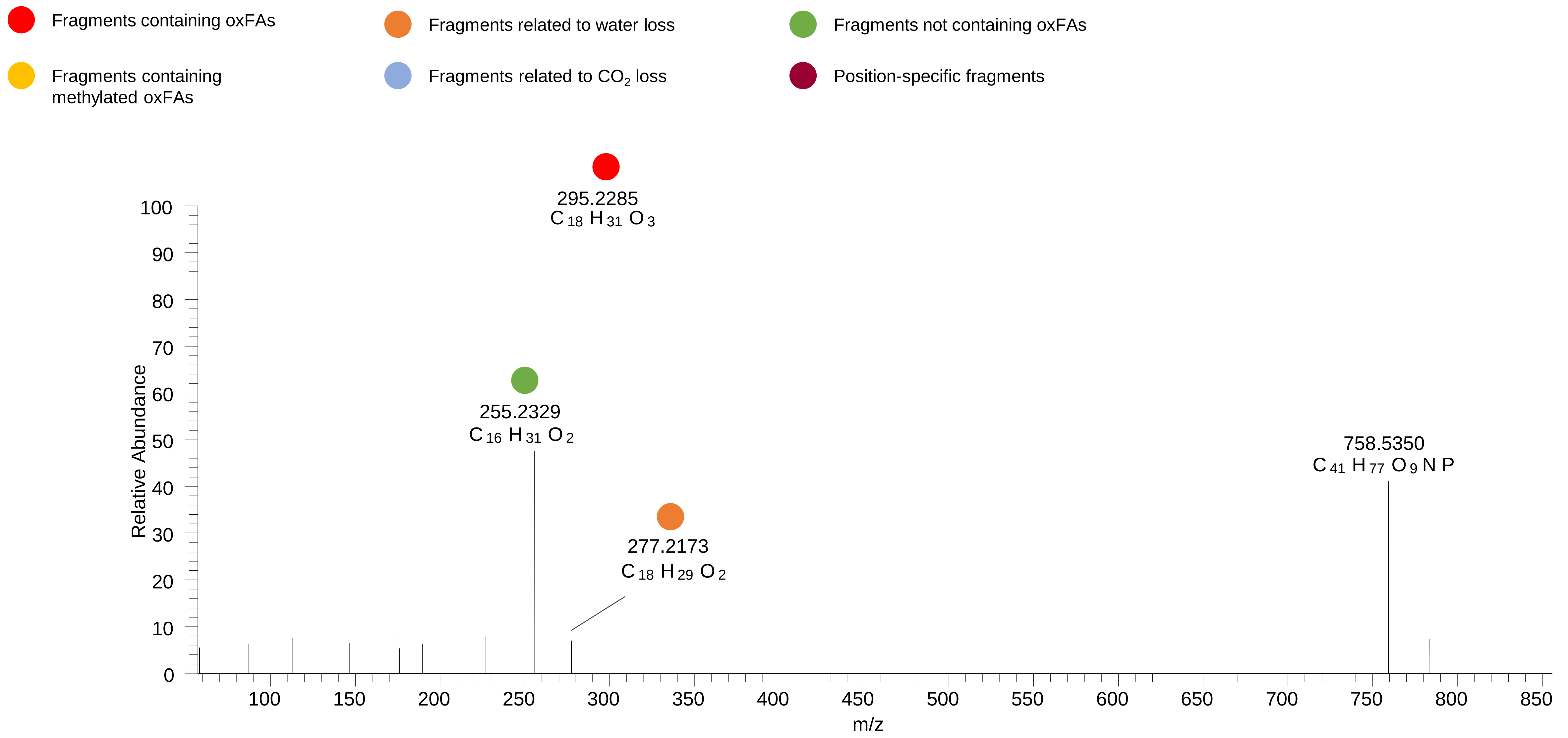

# PC(16:0_18:2<OH{13}>)
# RT 13.4

**[oxPC+HCOO]- XIC 818.5553**

- Fragments containing oxFAs
- Fragments related to water loss
- Fragments not containing oxFAs
- Fragments containing methylated oxFAs
- Fragments related to $CO_2$ loss
- Position-specific fragments

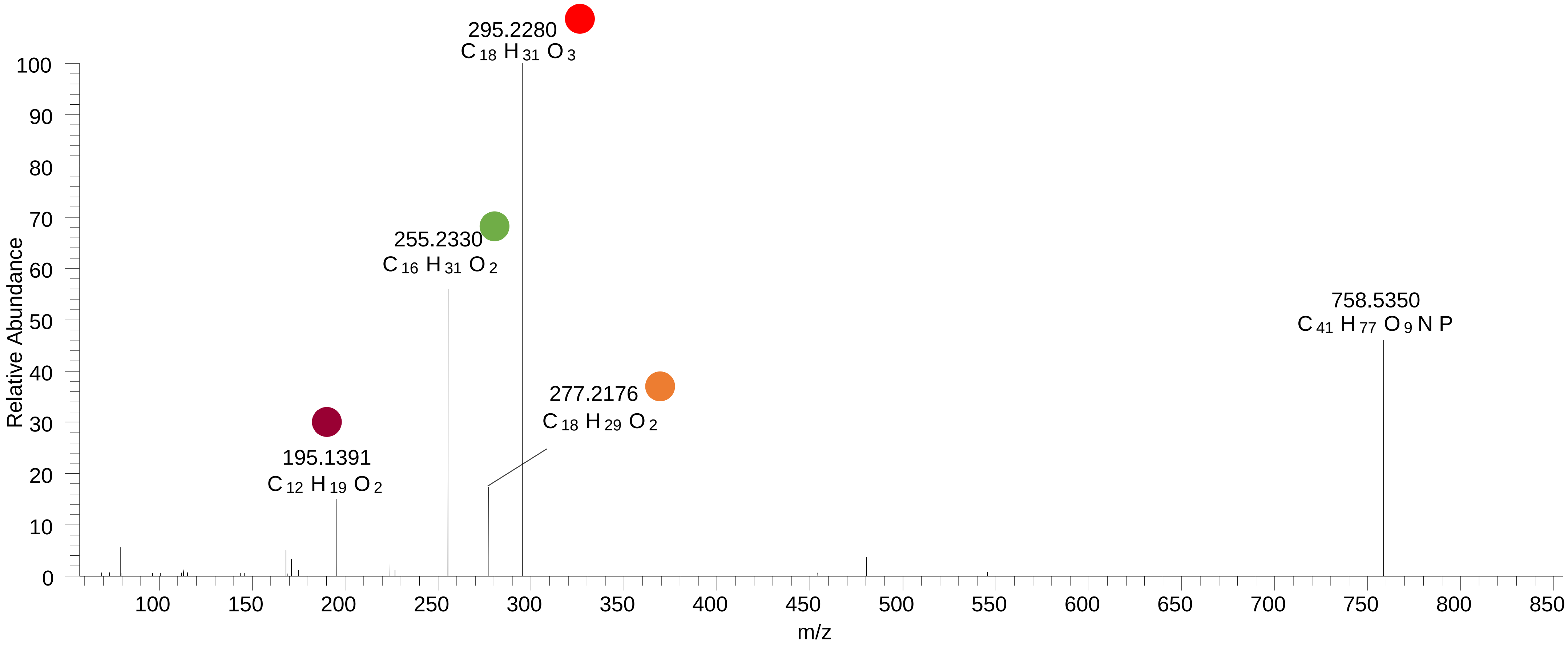

# PC(16:0_18:2<OH{9}>)
# RT 13.6

**[oxPC+HCOO]- XIC 818.5553**

- Fragments containing oxFAs
- Fragments related to water loss
- Fragments not containing oxFAs
- Fragments containing methylated oxFAs
- Fragments related to $CO_2$ loss
- Position-specific fragments

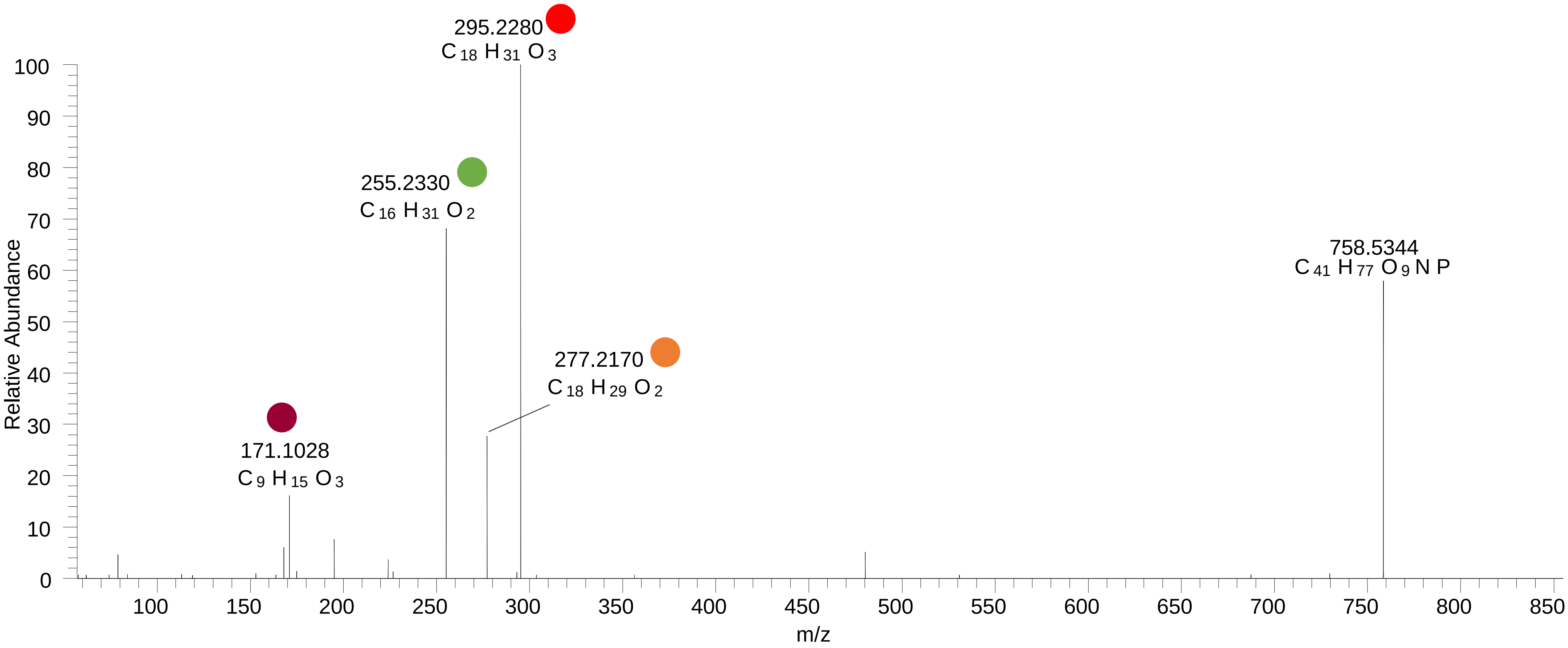

# PC(16:0<O>_18:2)
# RT 15.7

**[oxPC+HCOO]-   XIC 818.5553**

- Fragments containing oxFAs
- Fragments related to water loss
- Fragments not containing oxFAs
- Fragments containing methylated oxFAs
- Fragments related to $CO_2$ loss
- Position-specific fragments

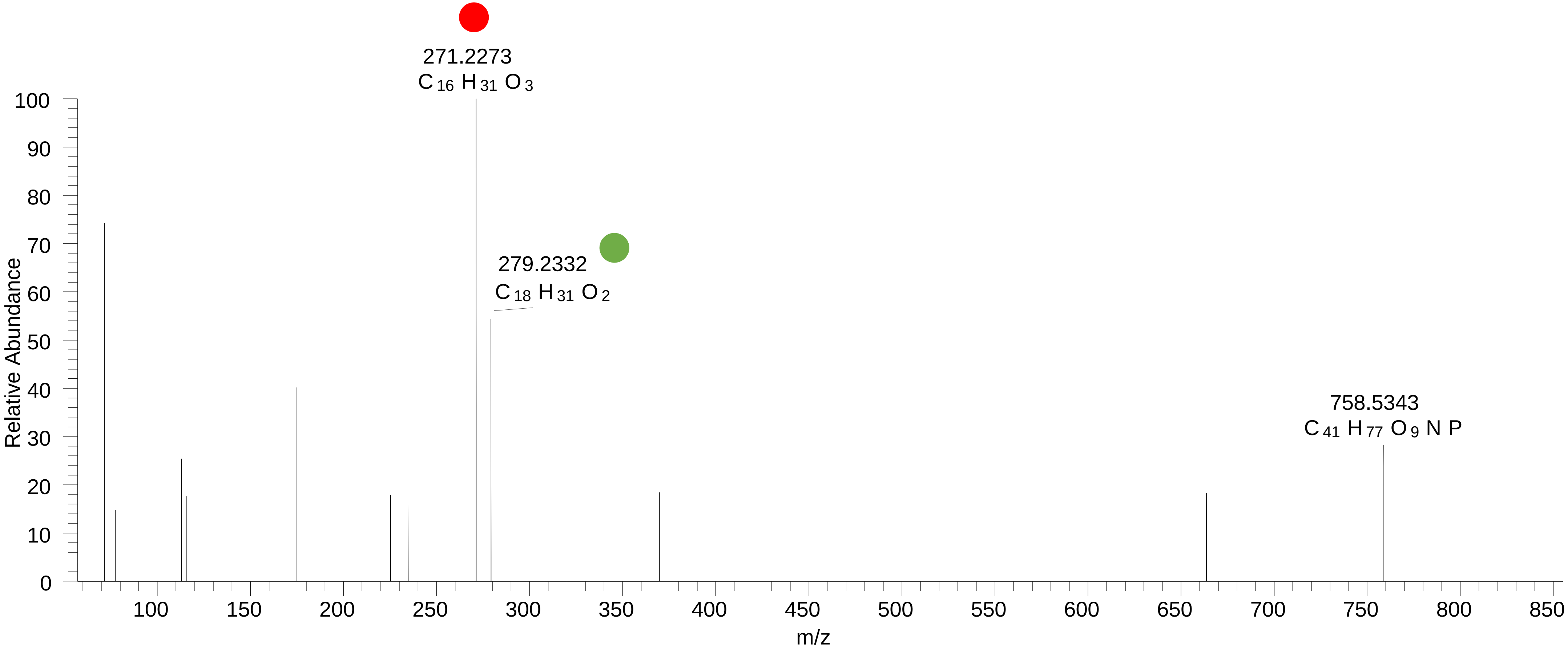

# PC(16:0_18:1<O>)
# RT 14.4

**[oxPC+HCOO]- XIC 820.5709**

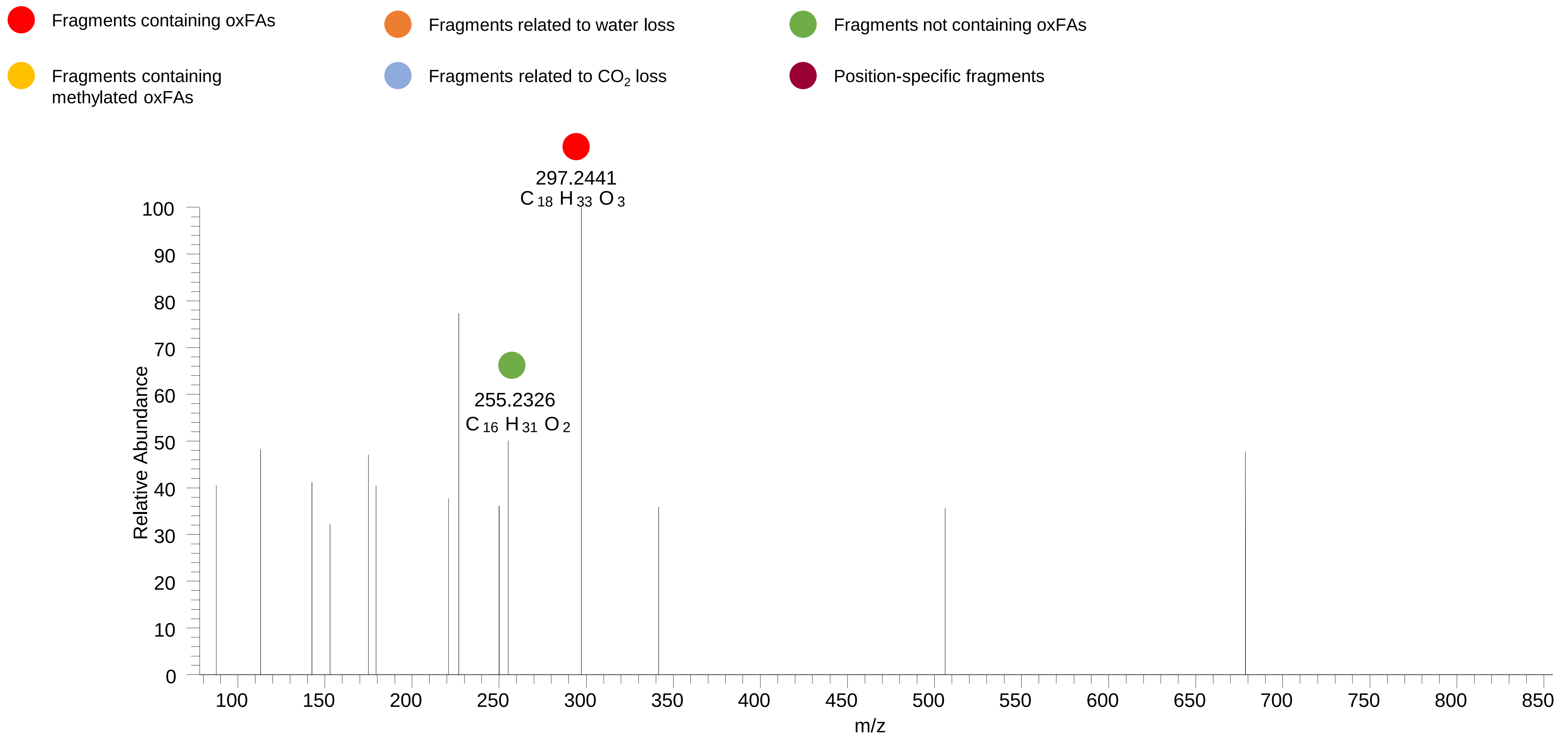

# PC(16:0<O>_18:1)
# RT 16.8-17.1

**[oxPC+HCOO]- XIC 820.5709**

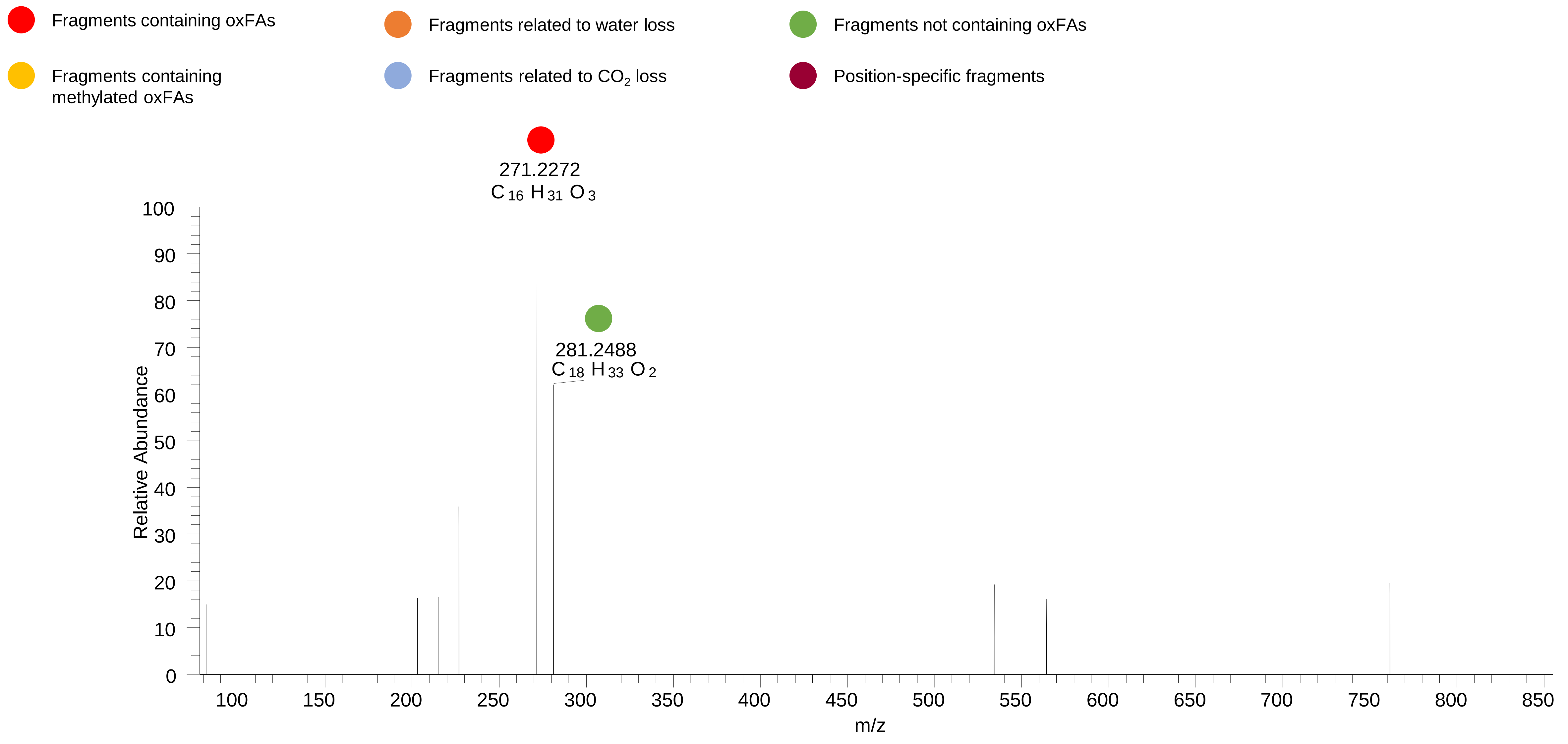

# PC(16:0_18:2<OH,O{9}>)
RT 11.6

[oxPC+HCOO]- XIC 834.5502

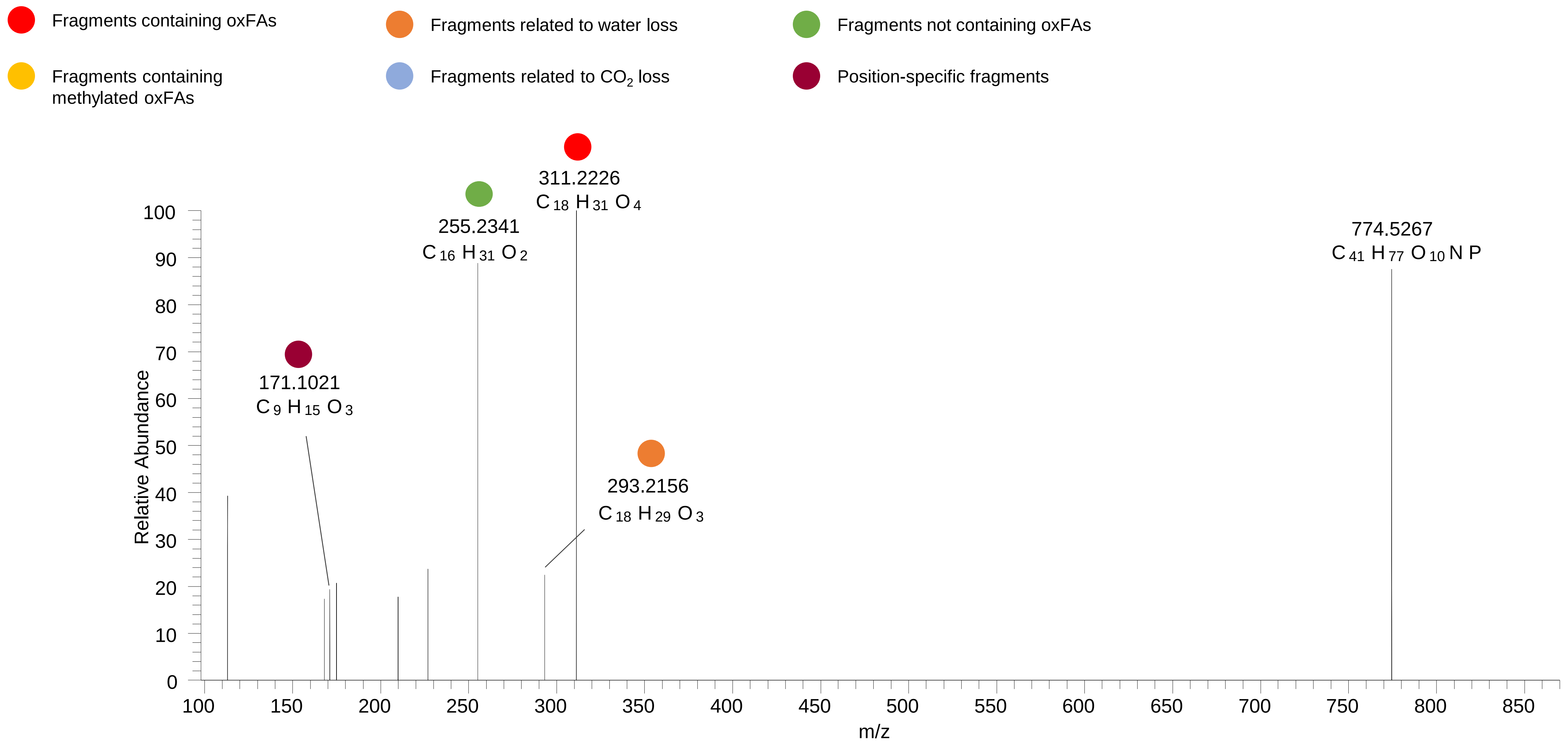

# PC(16:0_18:2<2O>)
# RT 12.0

**[oxPC+HCOO]- XIC 834.5502**

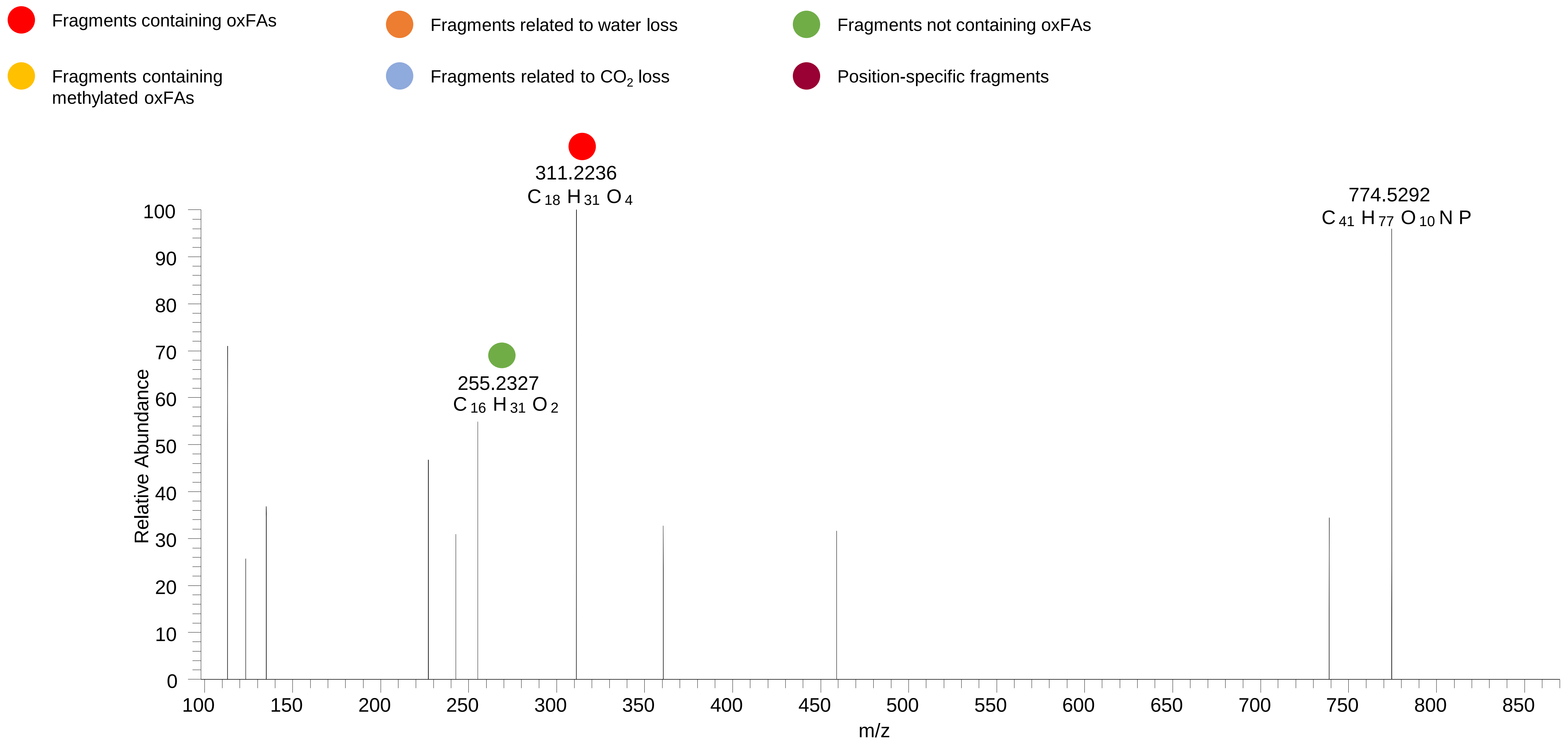

### PC(16:0_18:2<OOH>)
# RT 12.6

**[oxPC+HCOO]- XIC 834.5502**

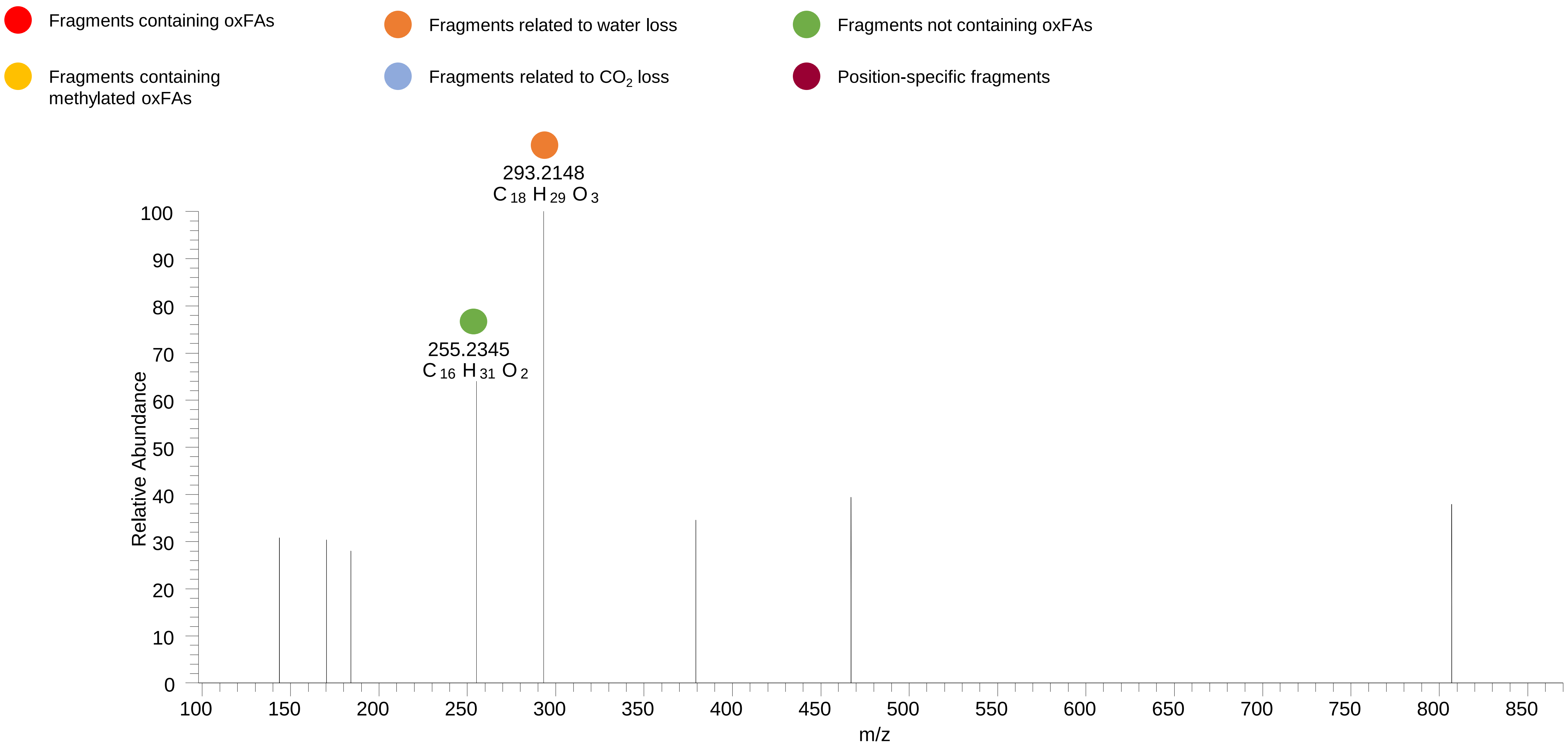

### PC(16:0_18:2<OOH{9}>)
### PC(16:0_18:2<OOH{13}>)
# RT 13.2-13.4

**[oxPC+HCOO]- XIC 834.5502**

- Fragments containing oxFAs
- Fragments related to water loss
- Fragments not containing oxFAs
- Fragments containing methylated oxFAs
- Fragments related to $CO_2$ loss
- Position-specific fragments

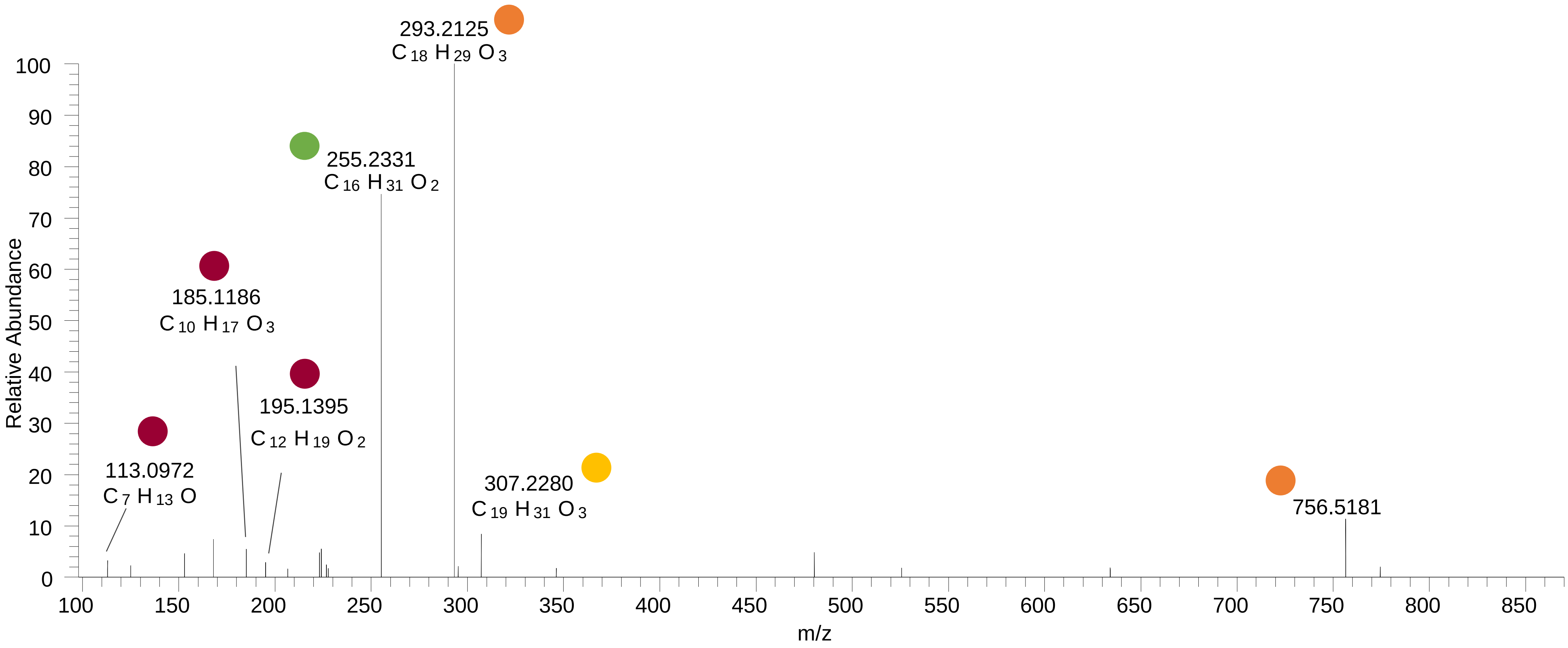

### PC(16:0_18:1<OOH>)
# RT 14.2-14.3

**[oxPC+HCOO]- XIC 836.5658**

Fragments containing oxFAs

Fragments related to water loss

Fragments not containing oxFAs

Fragments containing methylated oxFAs

Fragments related to $CO_2$ loss

Position-specific fragments

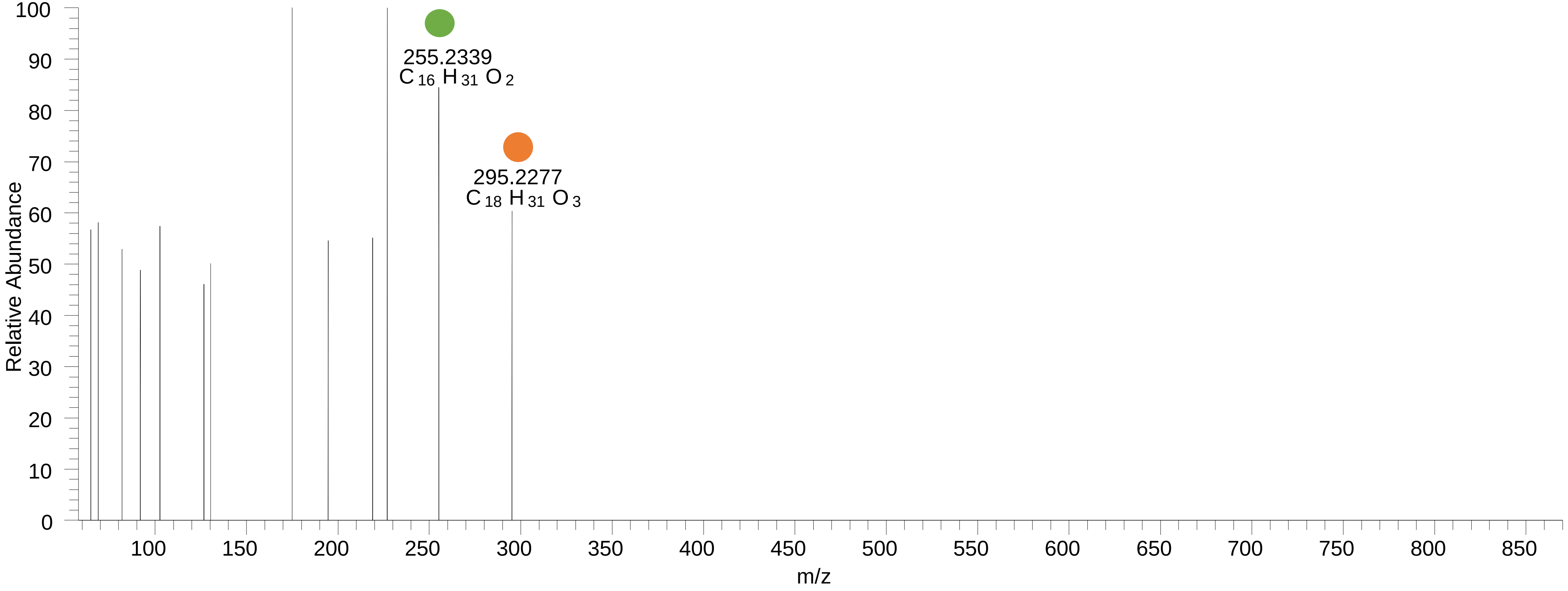

# PC(16:0_20:5<O>)
# RT 12.8

**[oxPC+HCOO]- XIC 840.5396**

- Fragments containing oxFAs
- Fragments related to water loss
- Fragments not containing oxFAs
- Fragments containing methylated oxFAs
- Fragments related to $CO_2$ loss
- Position-specific fragments

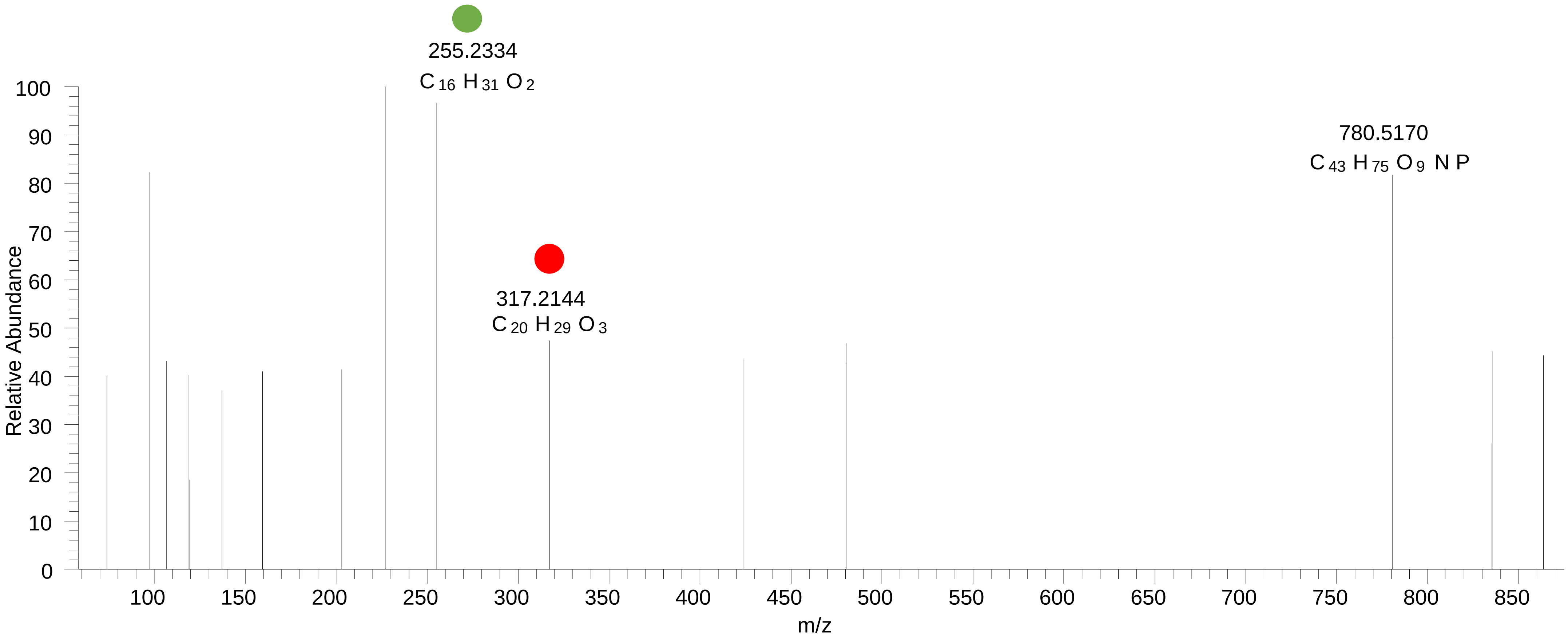

# PC(16:0_20:4<OH{13}>)
# RT 13.2

**[oxPC+HCOO]- XIC 842.5553**

- Fragments containing oxFAs
- Fragments related to water loss
- Fragments not containing oxFAs
- Fragments containing methylated oxFAs
- Fragments related to $CO_2$ loss
- Position-specific fragments

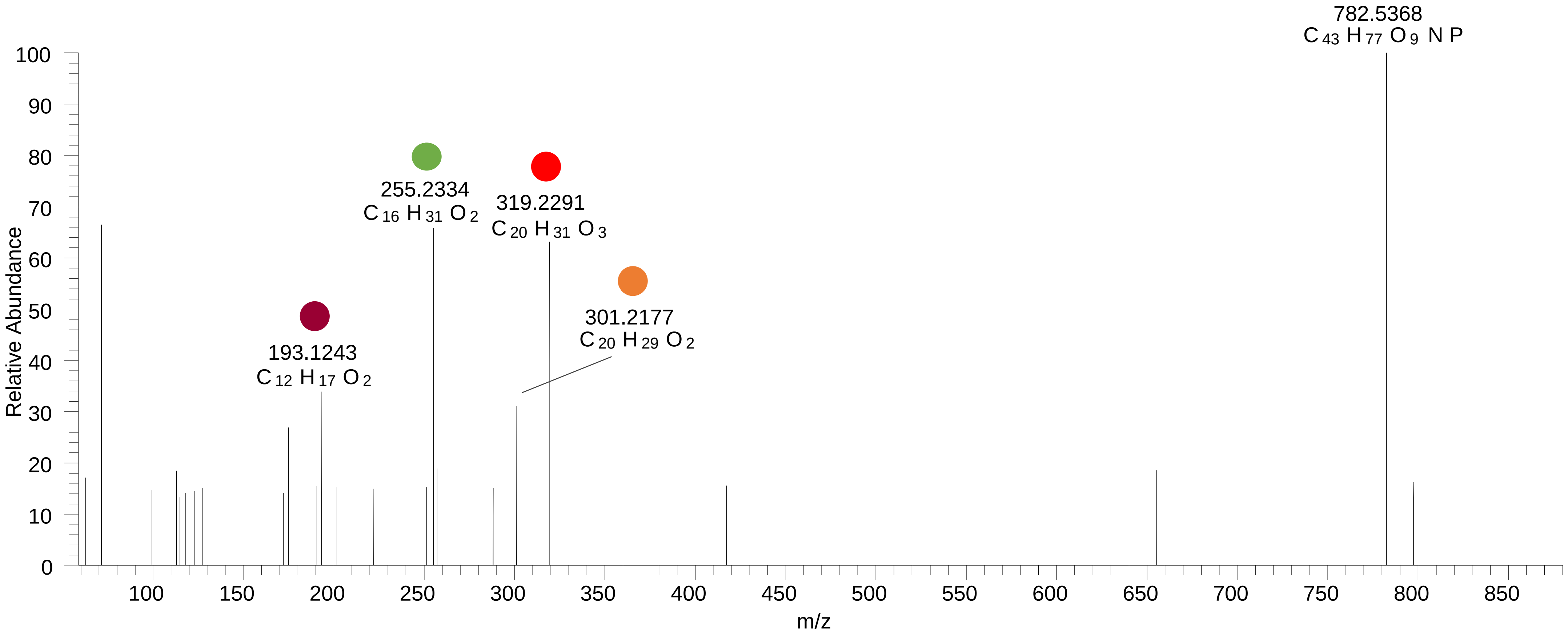

# PC(16:0_20:4<OH{15}>)
# RT 13.5

**[oxPC+HCOO]- XIC 842.5553**

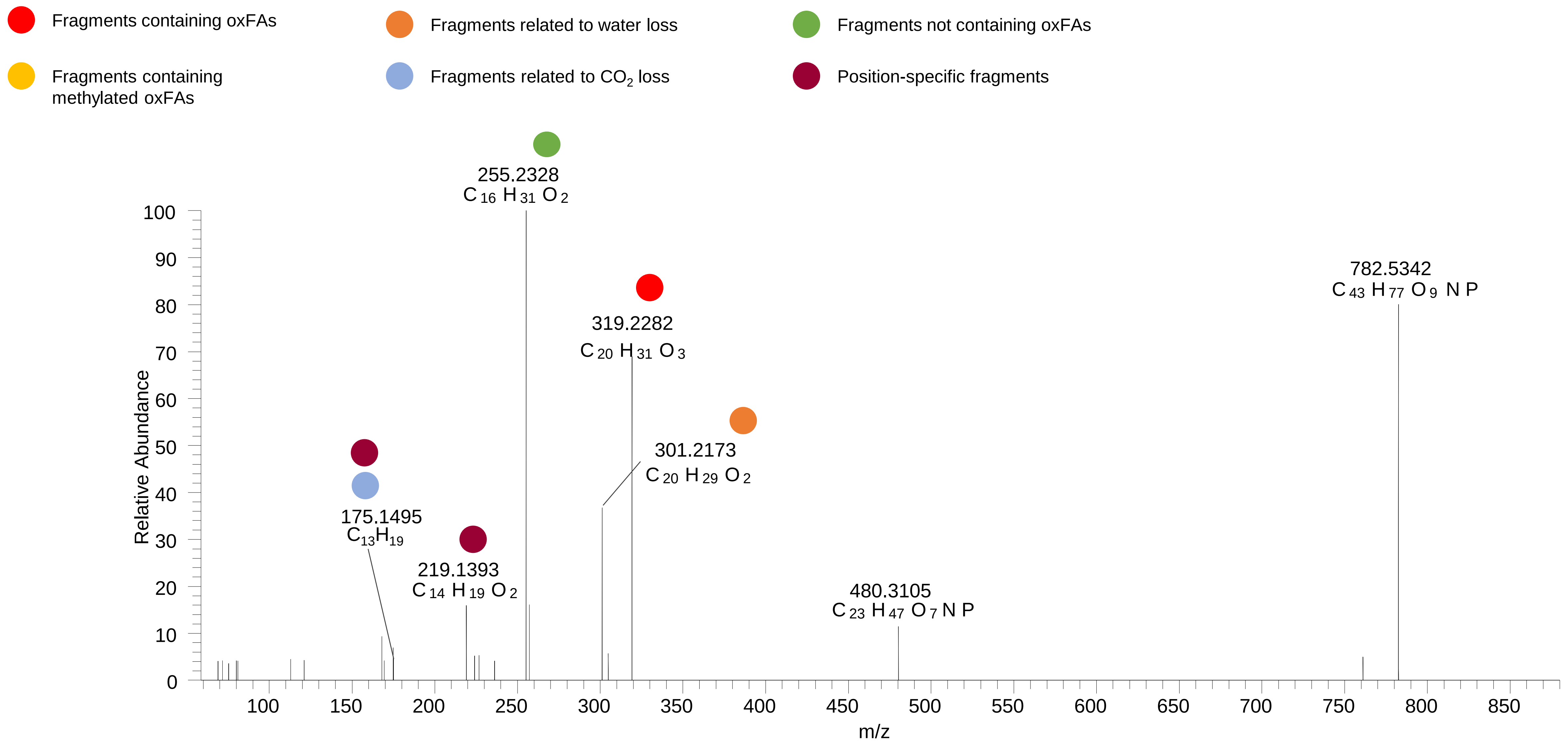

# PC(16:0_20:4<OH{11}>)
# RT 13.8

**[oxPC+HCOO]- XIC 842.5553**

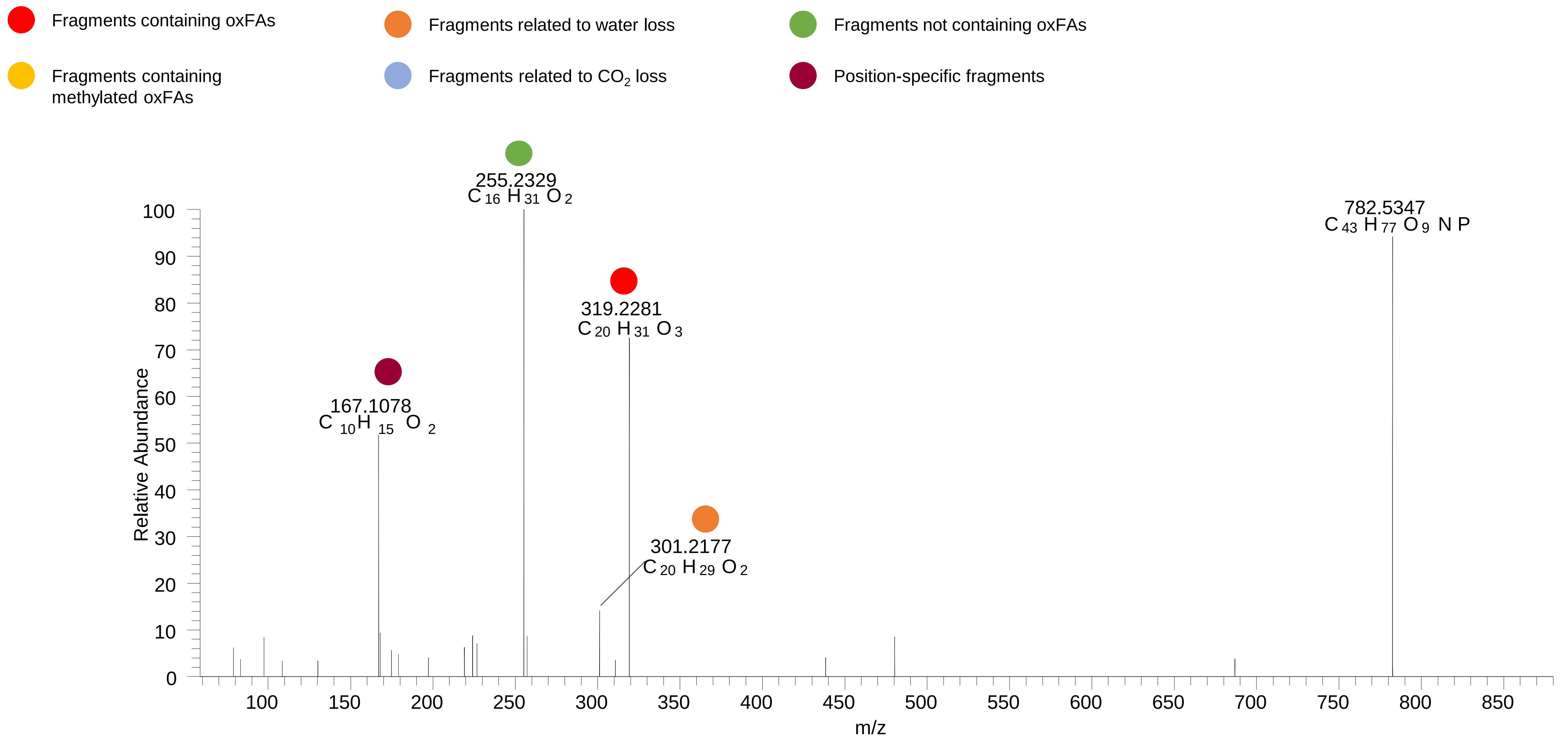

# PC(16:0_20:4<OH{11}>)
# PC(16:0_20:4<OH{12}>)
# RT 14.0

**[oxPC+HCOO]-   XIC 842.5553**

- Fragments containing oxFAs
- Fragments related to water loss
- Fragments not containing oxFAs
- Fragments containing methylated oxFAs
- Fragments related to $CO_2$ loss
- Position-specific fragments

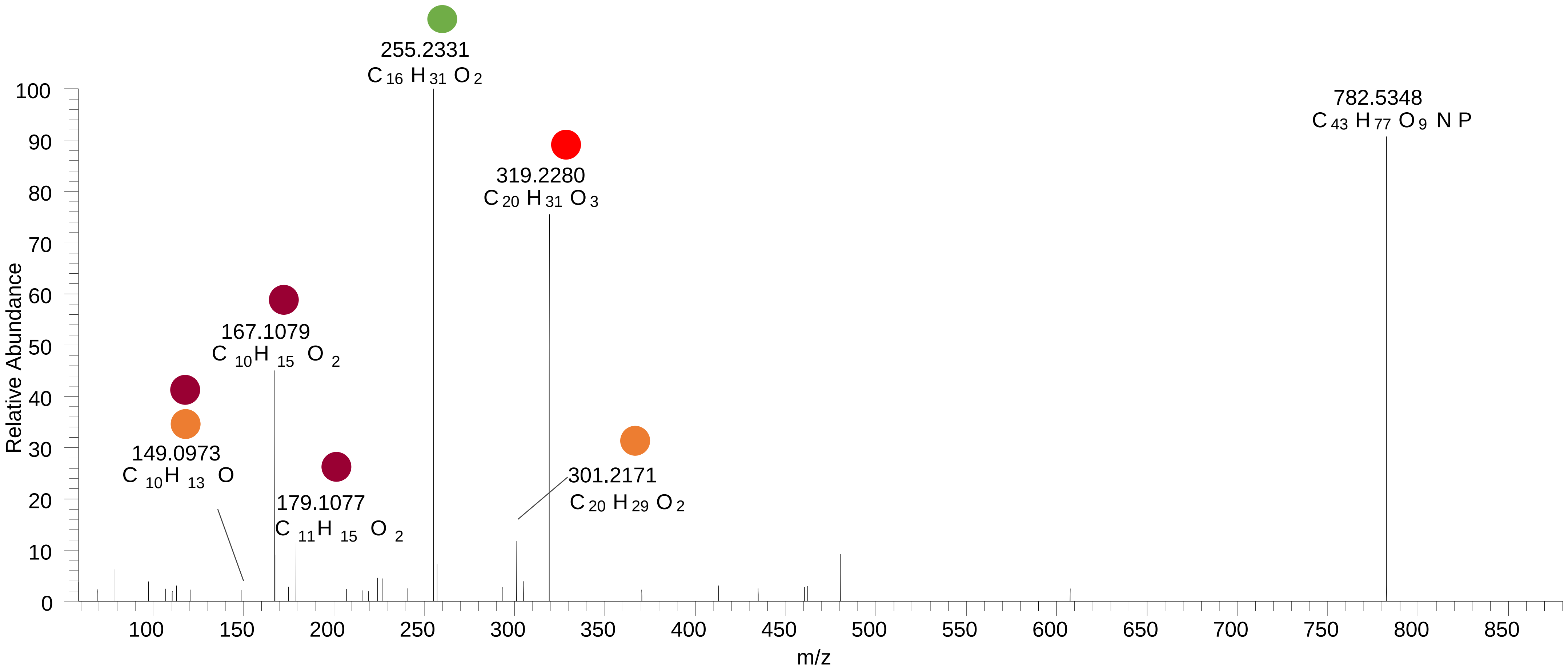

# PC(16:0_20:4<OH{8}>)
# PC(16:0_20:4<OH{7}>)
# RT 14.3-14.4

**[oxPC+HCOO]- XIC 842.5553**

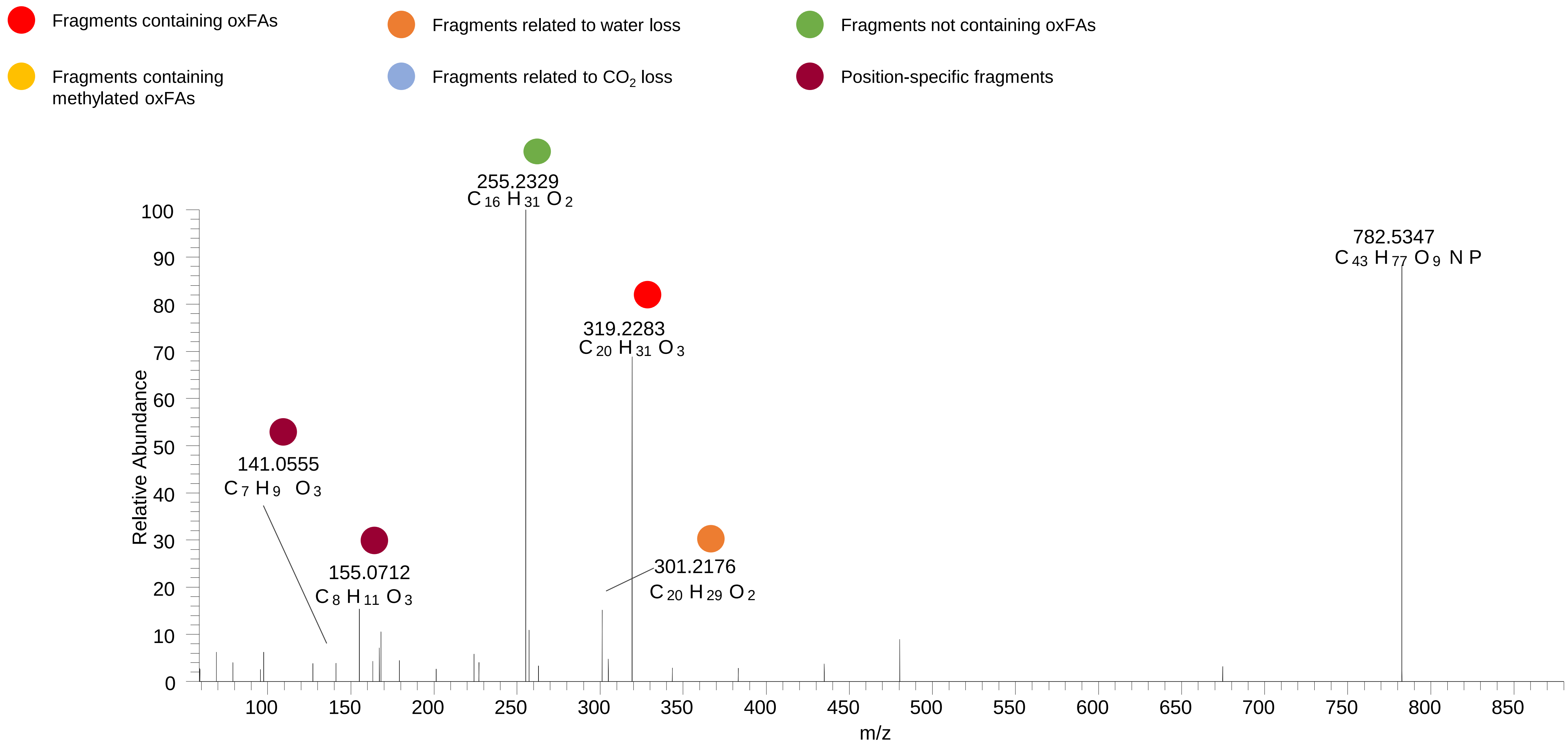

# PC(16:0_20:4<OH{5}>)
# RT 15.0

**[oxPC+HCOO]- XIC 842.5553**

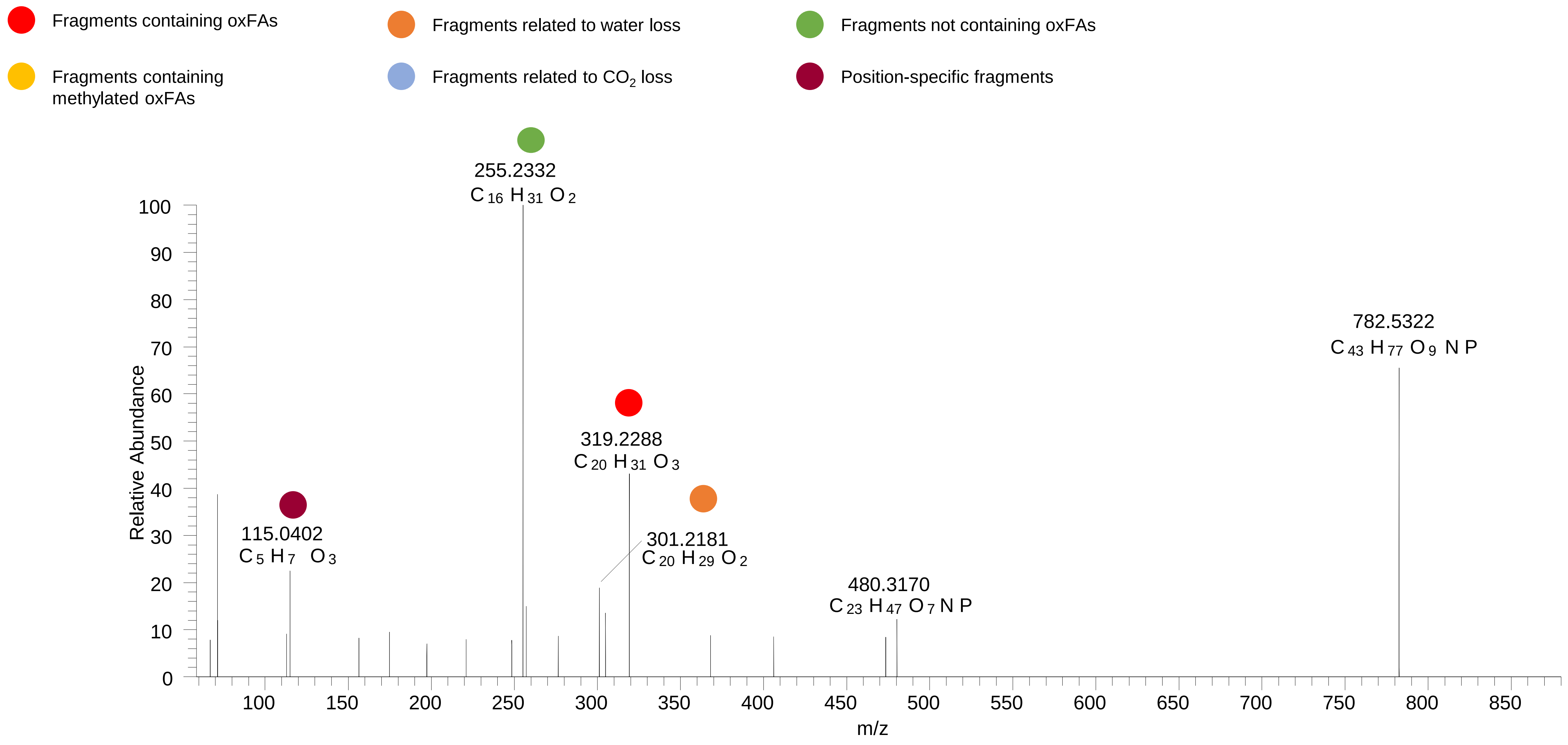

# PC(18:1_18:2<O>)
# RT 13.0

**[oxPC+HCOO]- XIC 844.5709**

Fragments containing oxFAs

Fragments related to water loss

Fragments not containing oxFAs

Fragments containing methylated oxFAs

Fragments related to $CO_2$ loss

Position-specific fragments

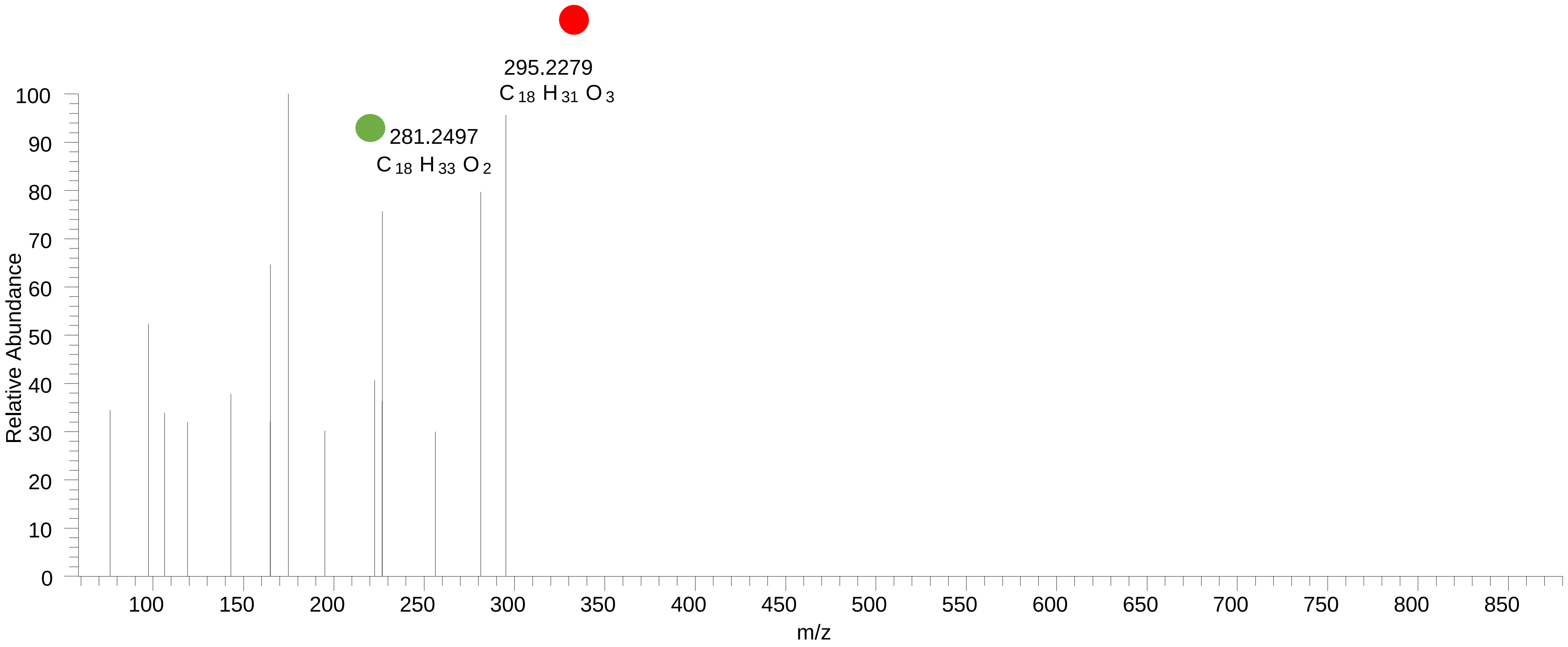

# PC(16:0_20:3<OH{11}>)
# PC(16:0_20:3<OH{12}>)
# RT 14.2-14.3

**[oxPC+HCOO]- XIC 844.5709**

- Fragments containing oxFAs
- Fragments related to water loss
- Fragments not containing oxFAs
- Fragments containing methylated oxFAs
- Fragments related to $CO_2$ loss
- Position-specific fragments

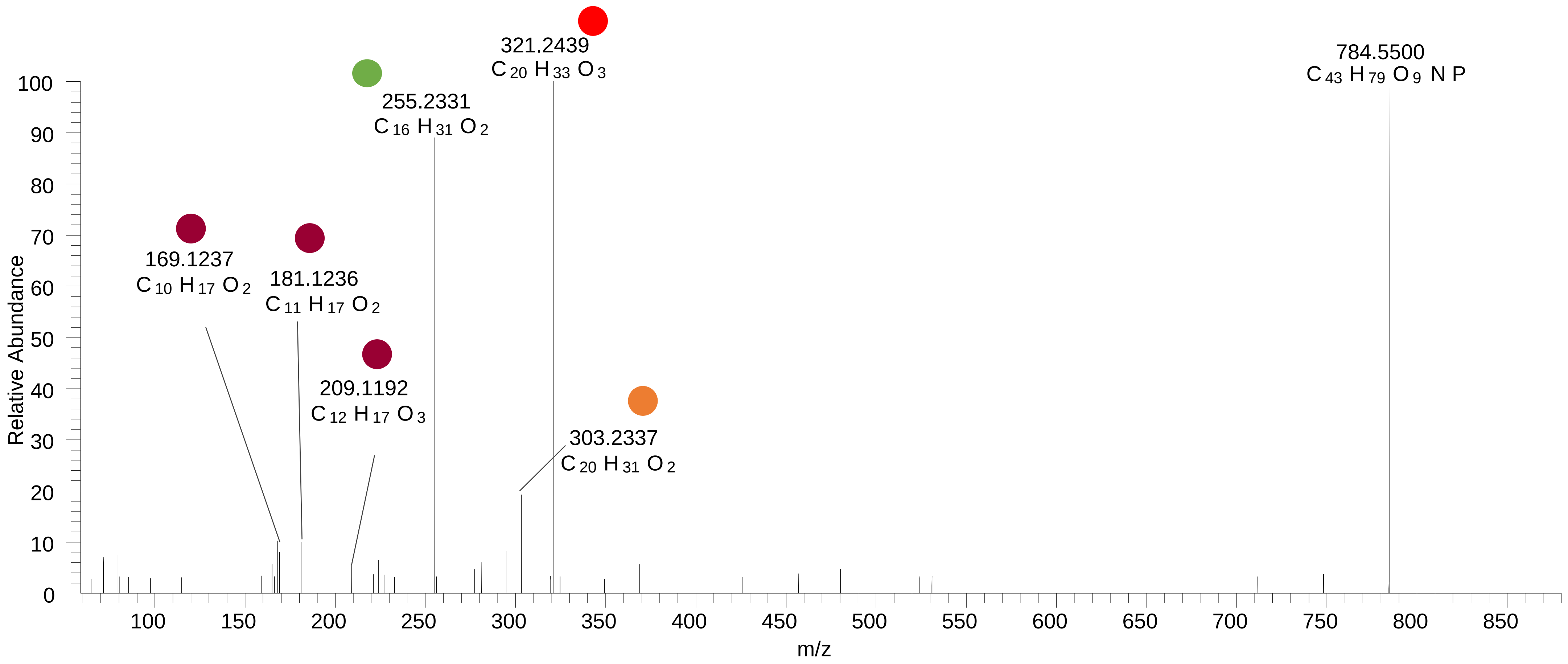

# PC(16:0_20:4<2O>)
# RT 11.2, 12.0, 12.6

**[oxPC+HCOO]- XIC 858.5502**

Fragments containing oxFAs

Fragments related to water loss

Fragments not containing oxFAs

Fragments containing methylated oxFAs

Fragments related to $CO_2$ loss

Position-specific fragments

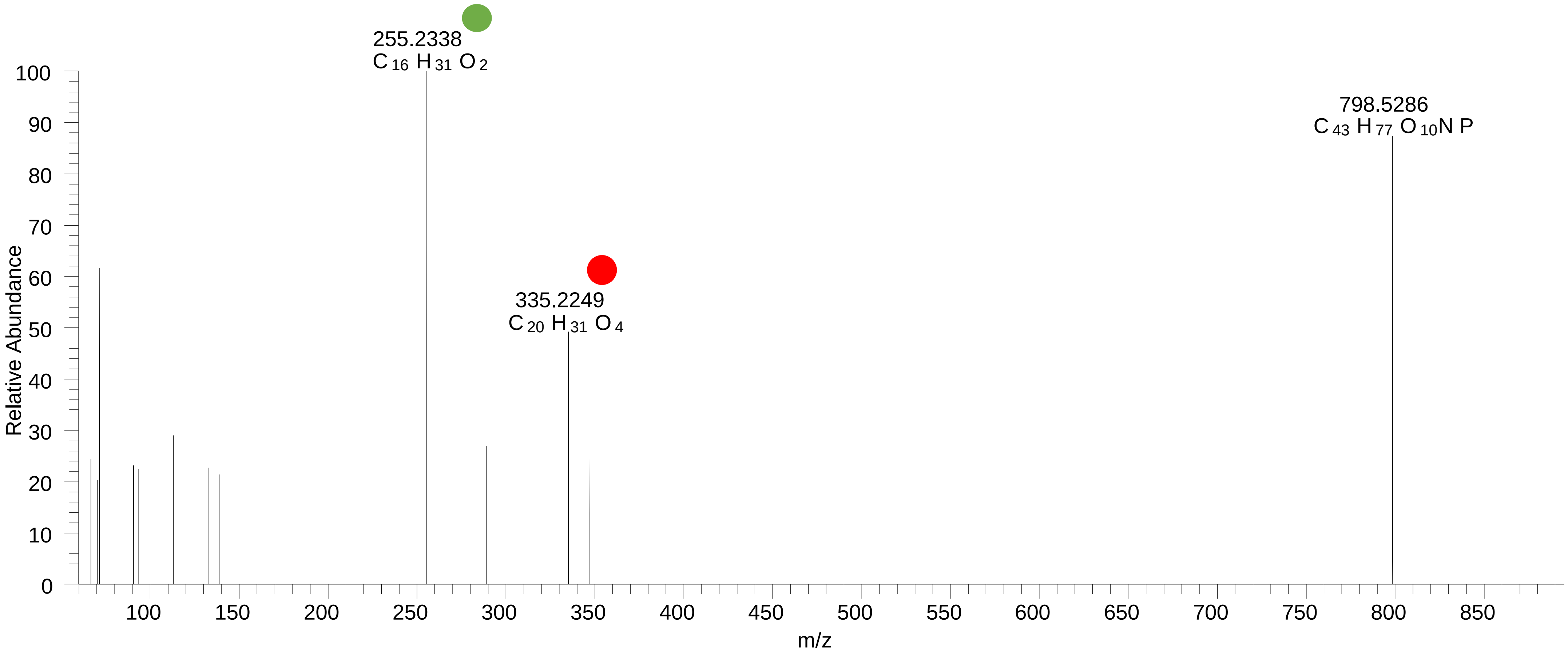

### PC(16:0_20:4<OOH{15}>)
# RT 13.6

**[oxPC+HCOO]- XIC 858.5502**

Fragments containing oxFAs

Fragments related to water loss

Fragments not containing oxFAs

Fragments containing methylated oxFAs

Fragments related to $CO_2$ loss

Position-specific fragments

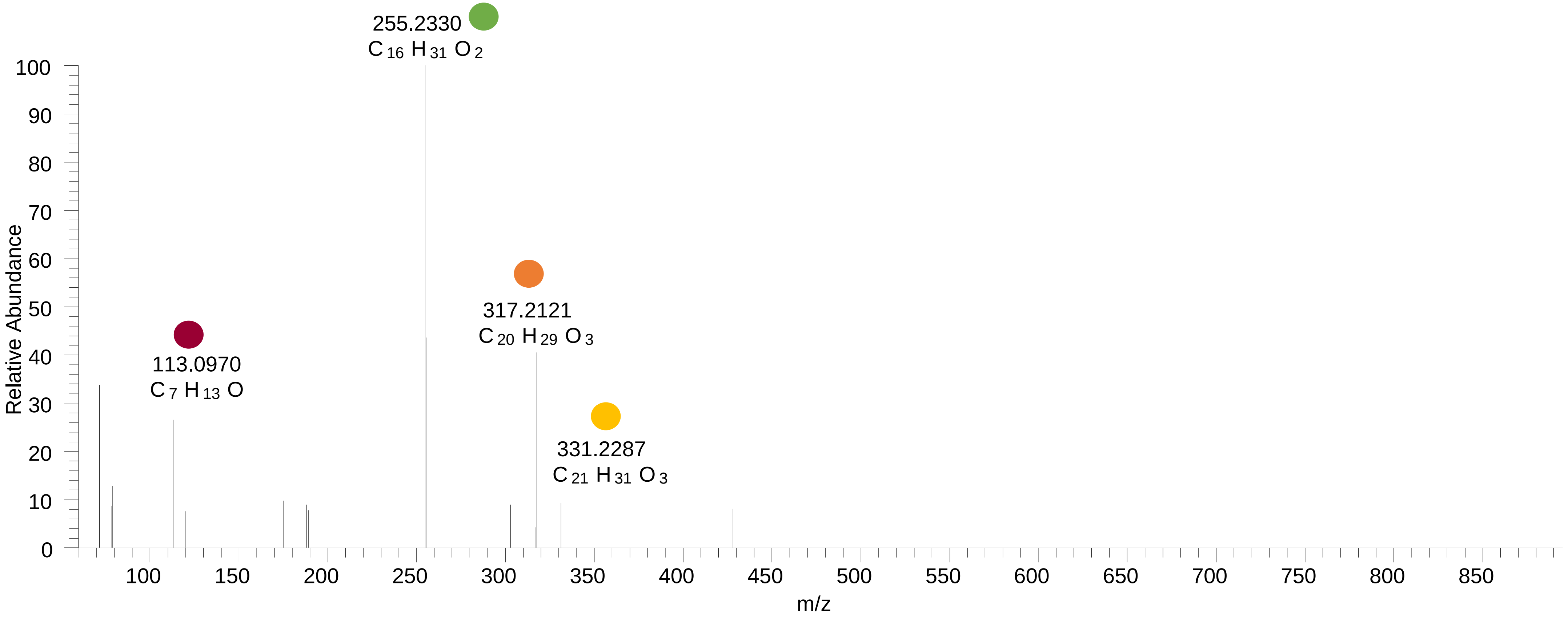

### PC(16:0_20:4<OOH{12}>)
# RT 13.9

**[oxPC+HCOO]- XIC 858.5502**

- Fragments containing oxFAs
- Fragments related to water loss
- Fragments not containing oxFAs
- Fragments containing methylated oxFAs
- Fragments related to $CO_2$ loss
- Position-specific fragments

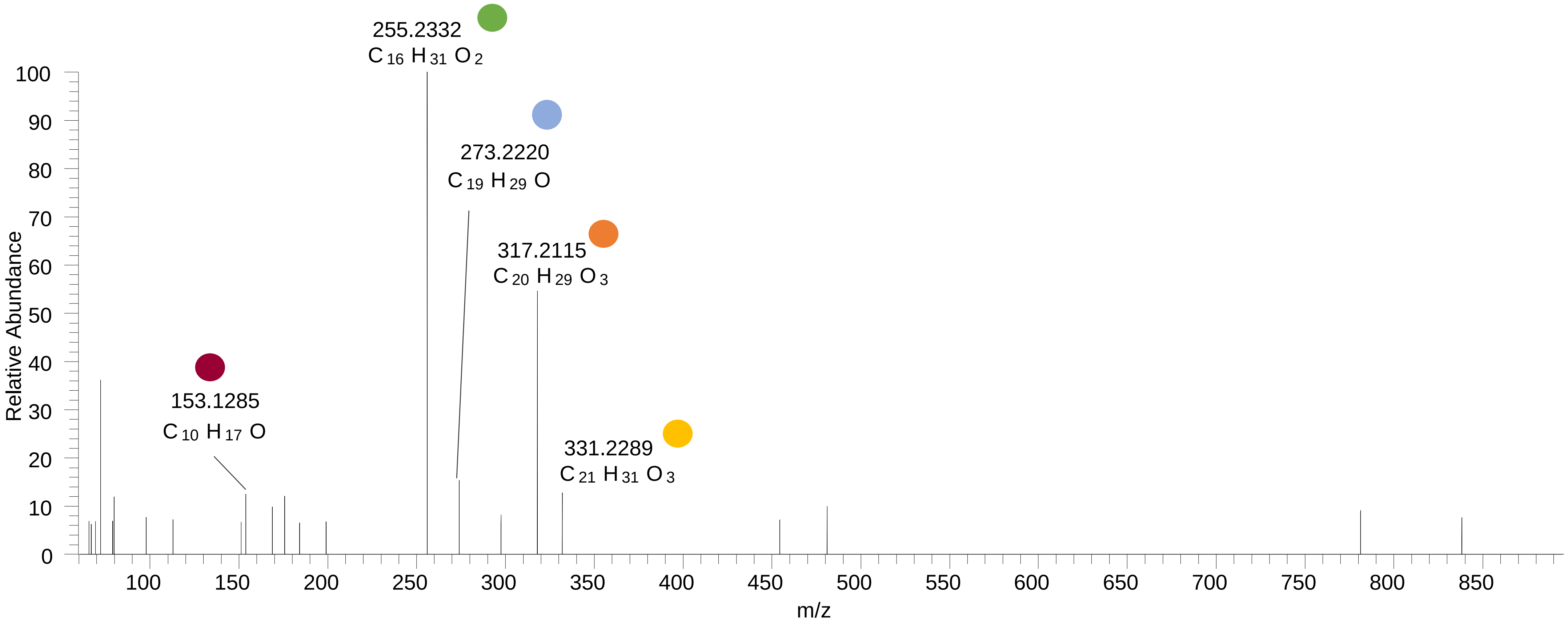

### PC(16:0_20:4<OOH{7}>)
# RT 14.4

**[oxPC+HCOO]- XIC 858.5502**

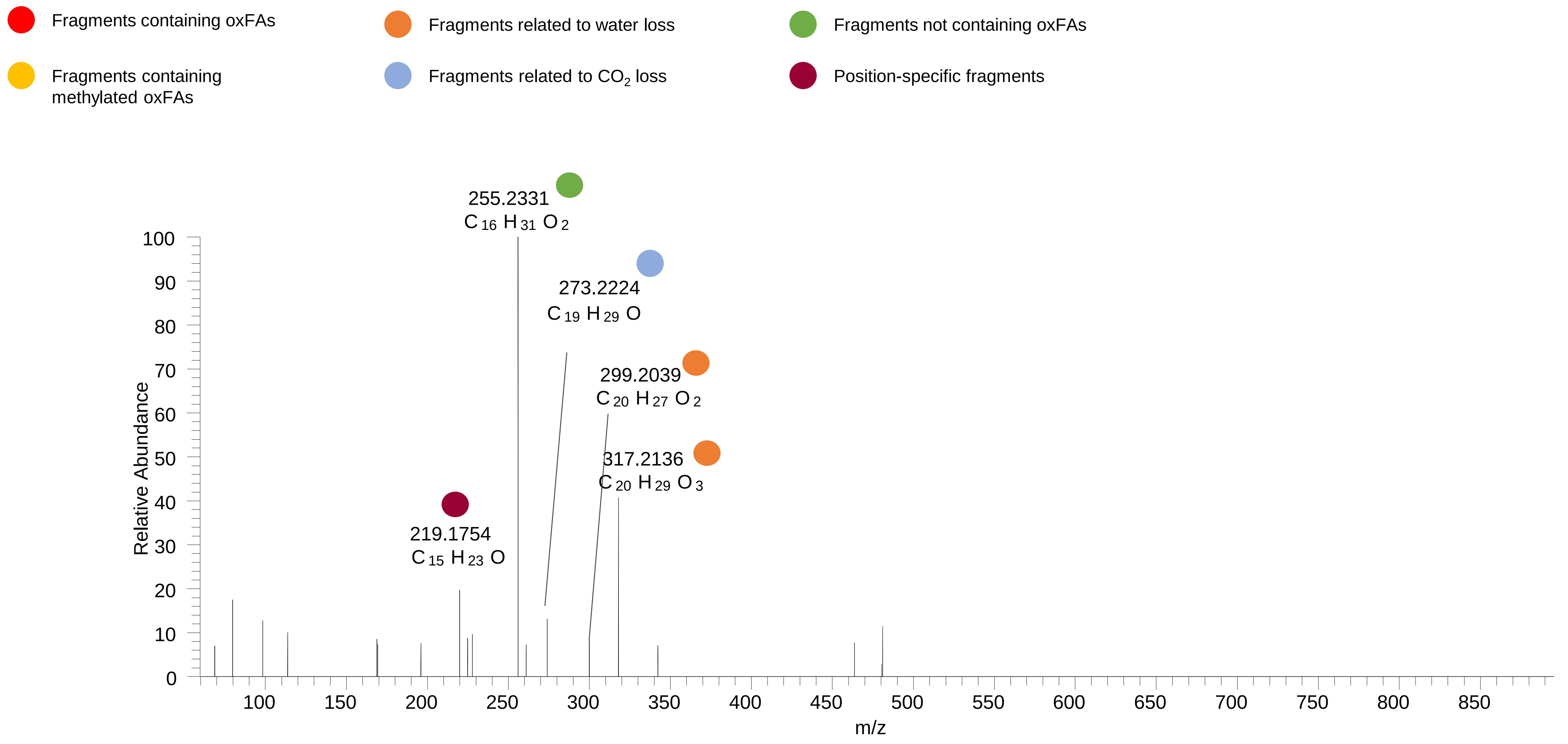

### PC(18:1_18:2<OOH>)
### PC(16:0_20:3<OOH>)
# RT 14.1

**[oxPC+HCOO]-   XIC 860.5658**

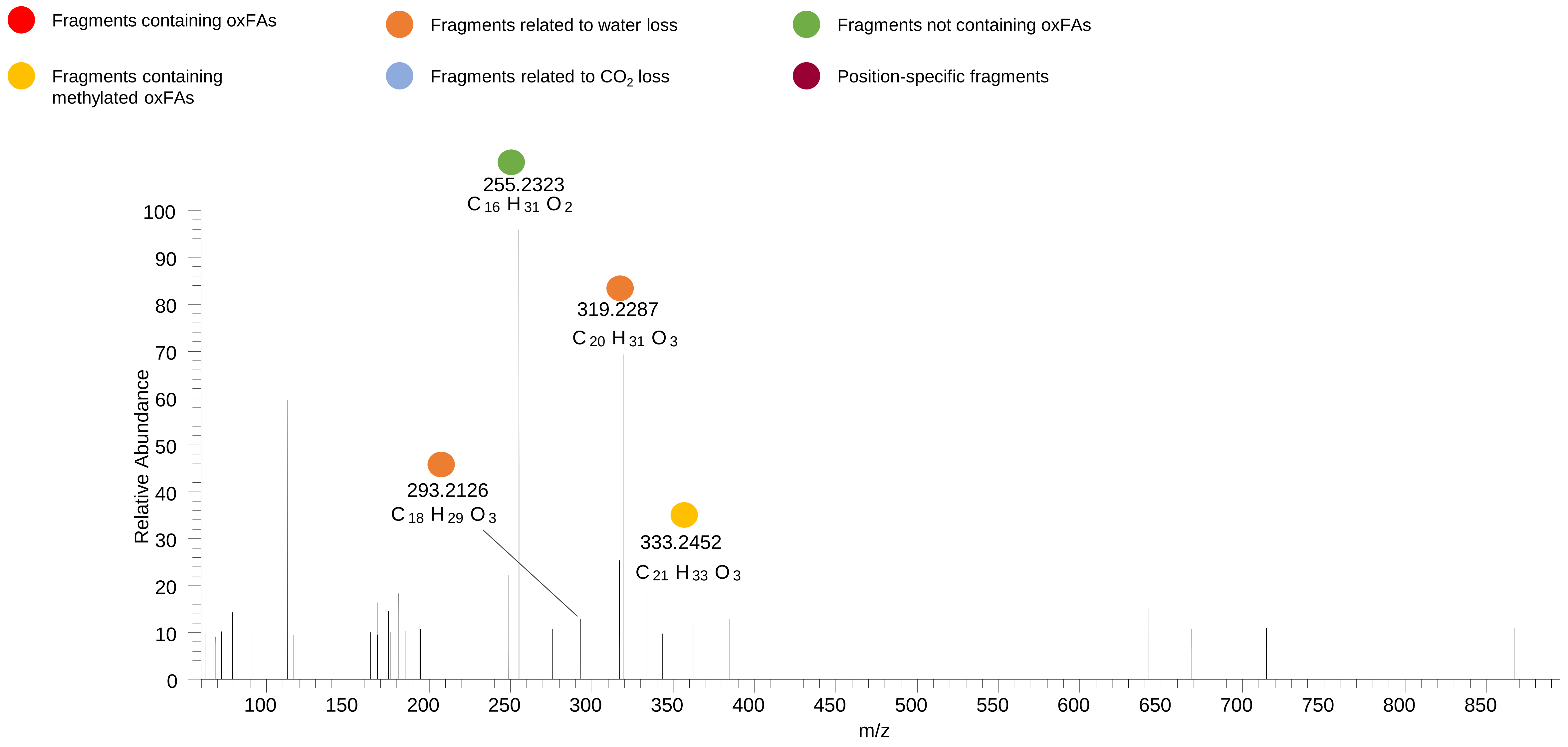

### PC(16:0_20:3<OOH>)
# RT 14.2

**[oxPC+HCOO]- XIC 860.5658**

Fragments containing oxFAs

Fragments related to water loss

Fragments not containing oxFAs

Fragments containing methylated oxFAs

Fragments related to $CO_2$ loss

Position-specific fragments

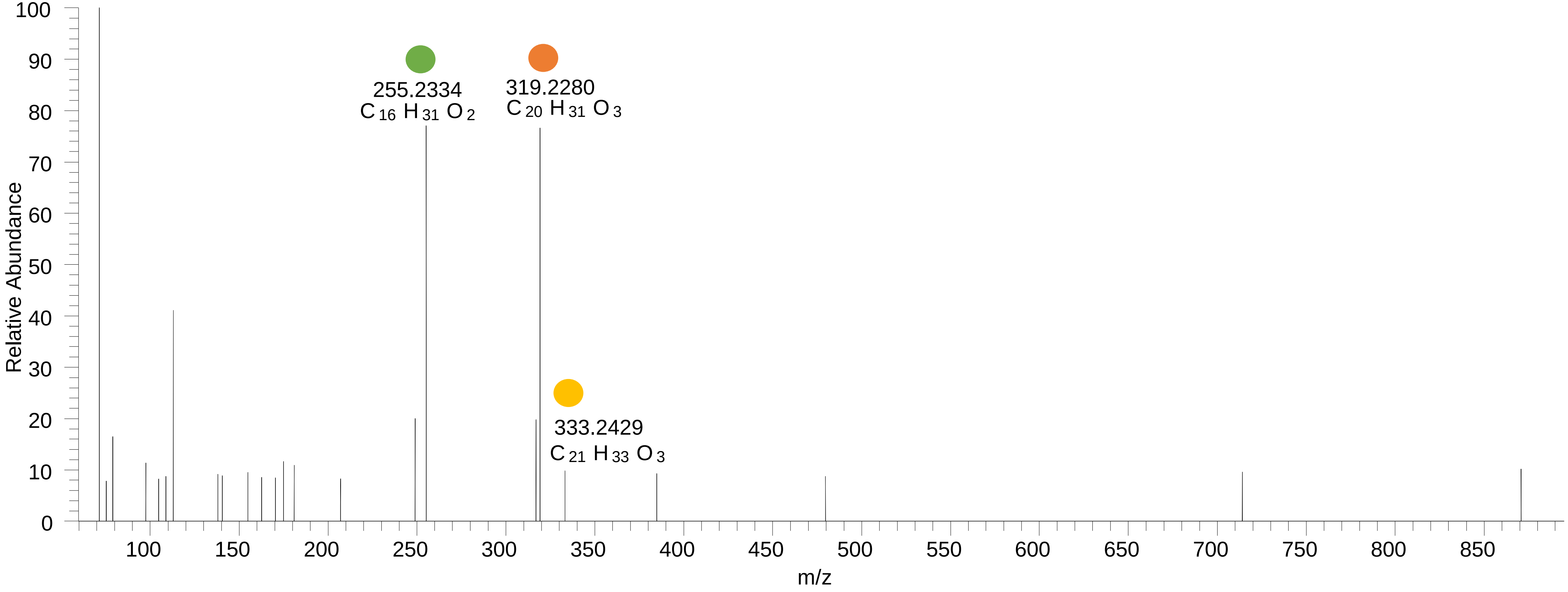

# PC(18:0_18:2<2O>)
# RT 13.6

**[oxPC+HCOO]- XIC 862.5815**

- Fragments containing oxFAs
- Fragments related to water loss
- Fragments not containing oxFAs
- Fragments containing methylated oxFAs
- Fragments related to $CO_2$ loss
- Position-specific fragments

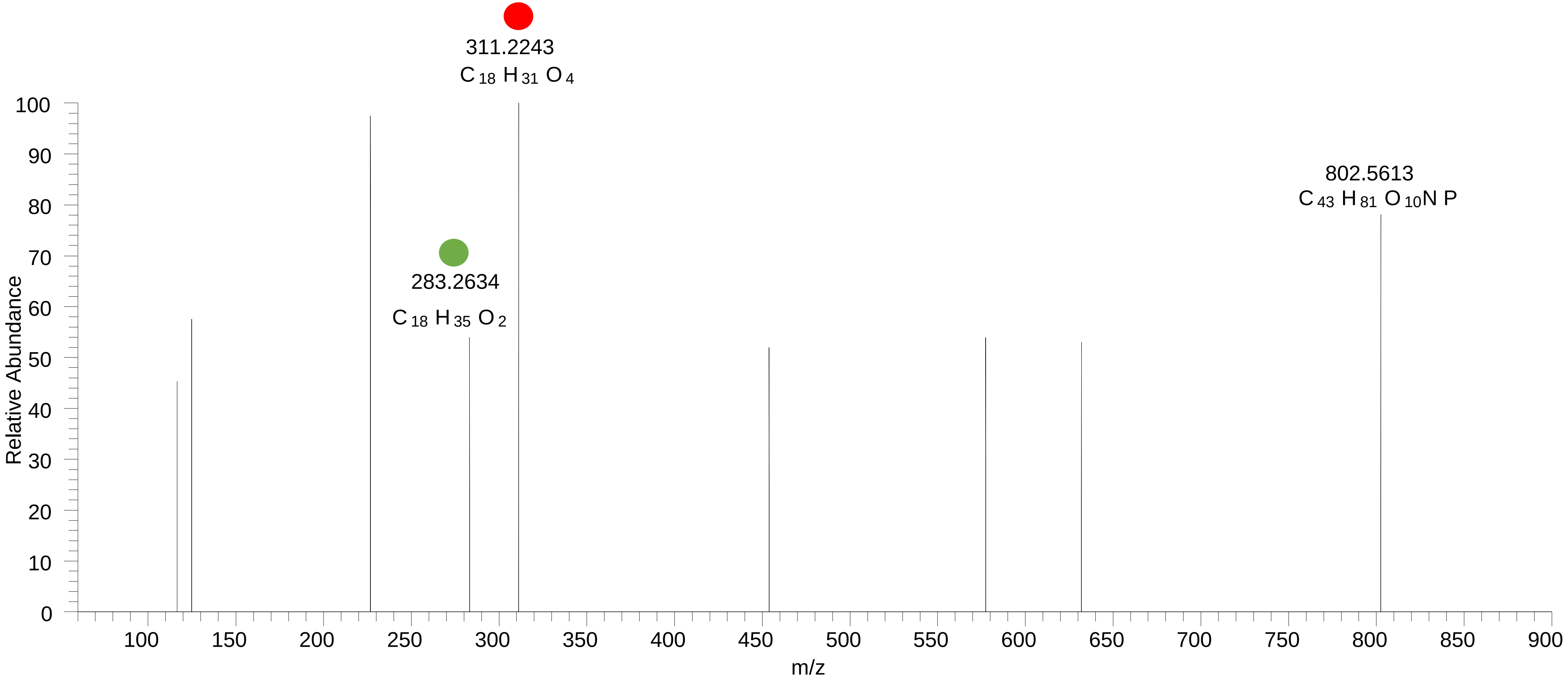

### PC(18:0_18:2<OOH>)
# RT 14.5

**[oxPC+HCOO]- XIC 862.5815**

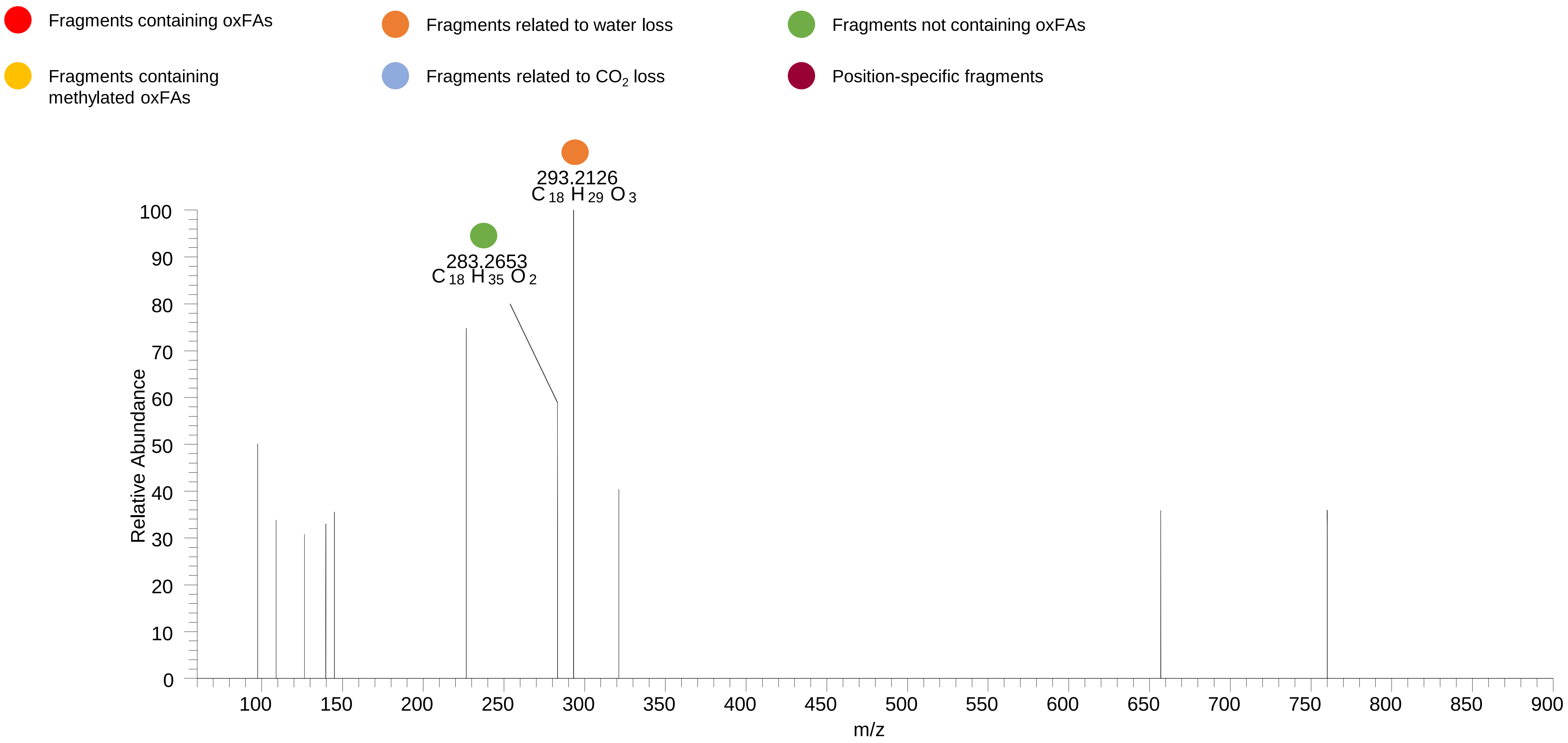

### PC(18:0_18:2<OOH{13}>)
# RT 15.0

**[oxPC+HCOO]-   XIC 862.5815**

- Fragments containing oxFAs
- Fragments related to water loss
- Fragments not containing oxFAs
- Fragments containing methylated oxFAs
- Fragments related to $CO_2$ loss
- Position-specific fragments

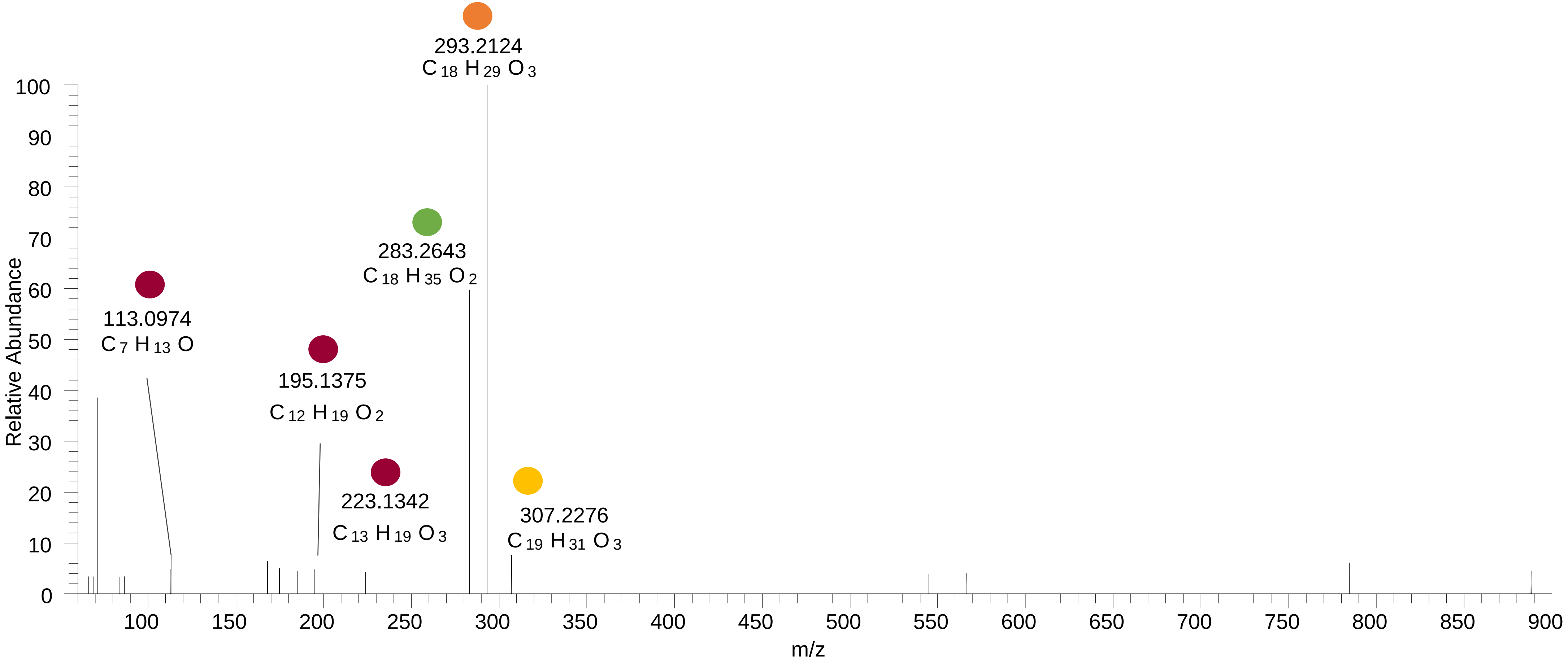

### PC(18:0_18:2<OOH{9}>)

# RT 15.1

**[oxPC+HCOO]- XIC 862.5815**

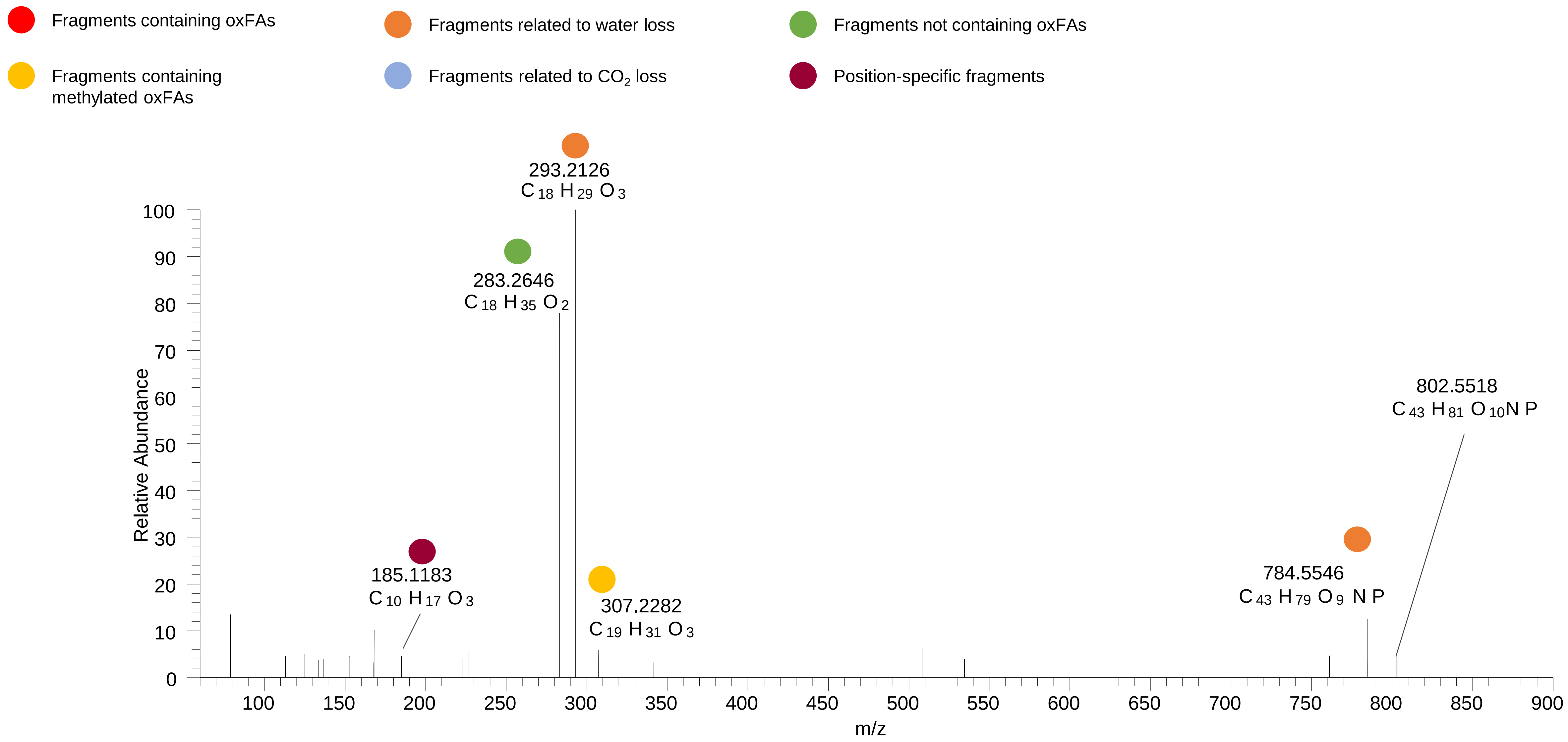

### PC(18:0_18:2<OOH>)
# RT 15.5

**[oxPC+HCOO]- XIC 862.5815**

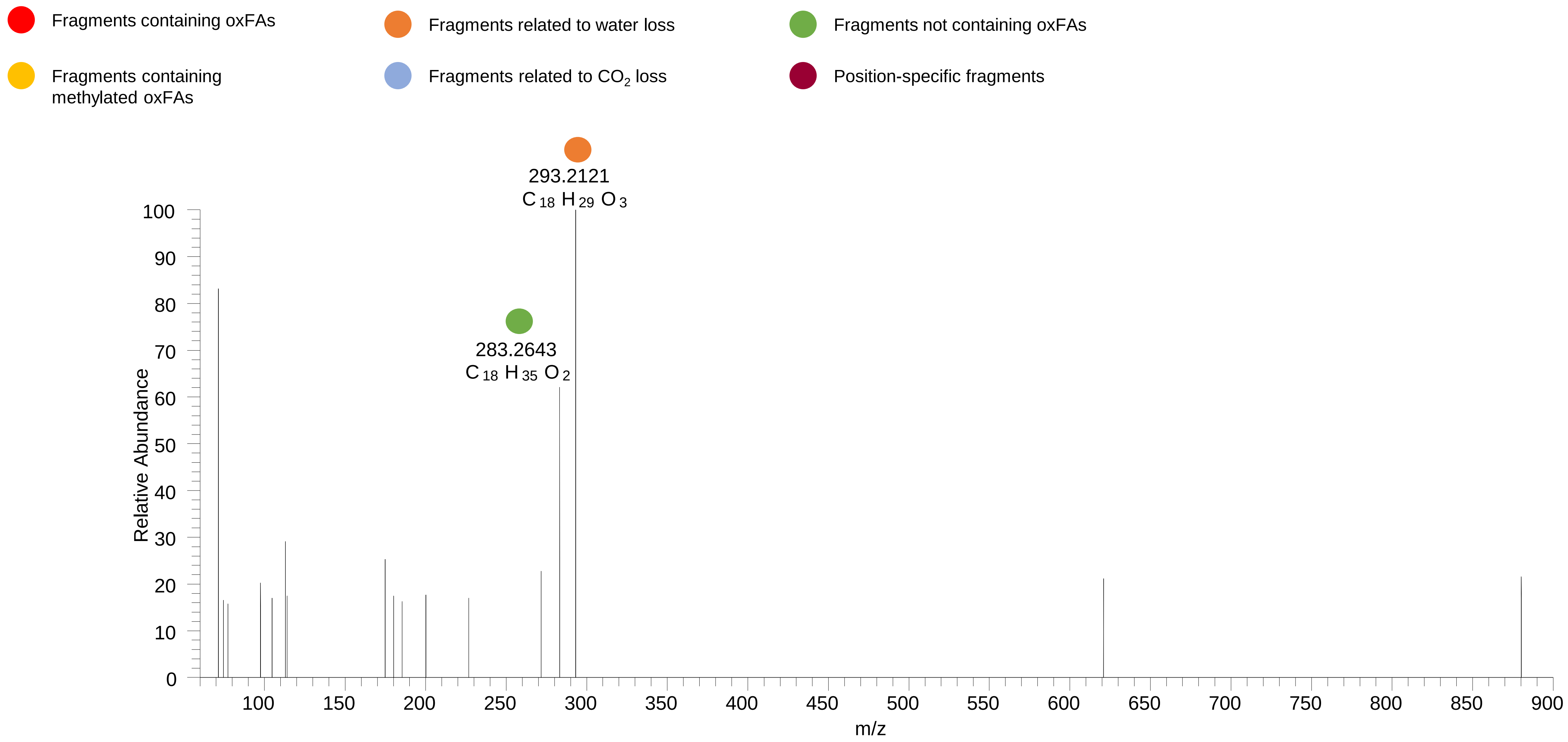

# PC(16:0_22:6<O>)
# RT 13.1

**[oxPC+HCOO]- XIC 866.5553**

- Fragments containing oxFAs
- Fragments related to water loss
- Fragments not containing oxFAs
- Fragments containing methylated oxFAs
- Fragments related to $CO_2$ loss
- Position-specific fragments

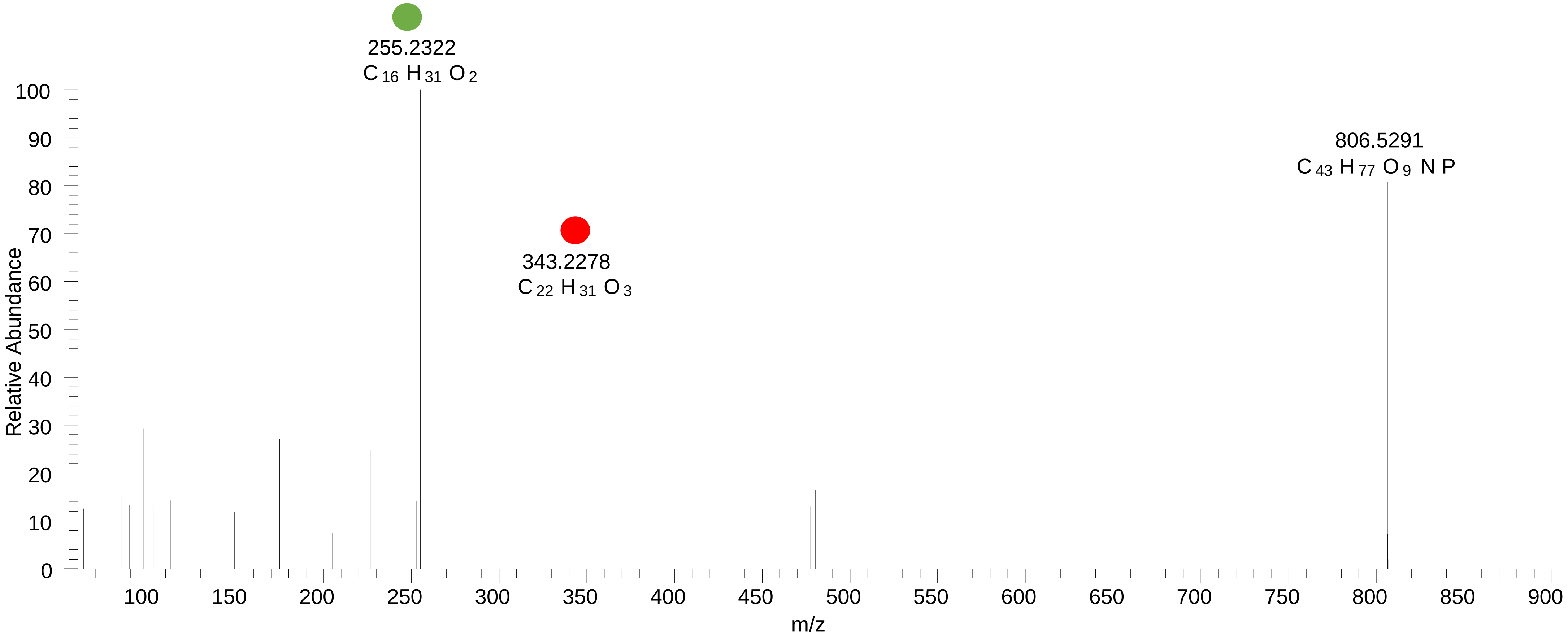

# PC(16:0_22:6<OH{16}>)
# RT 13.4

**[oxPC+HCOO]- XIC 866.5553**

- Fragments containing oxFAs
- Fragments related to water loss
- Fragments not containing oxFAs
- Fragments containing methylated oxFAs
- Fragments related to $CO_2$ loss
- Position-specific fragments

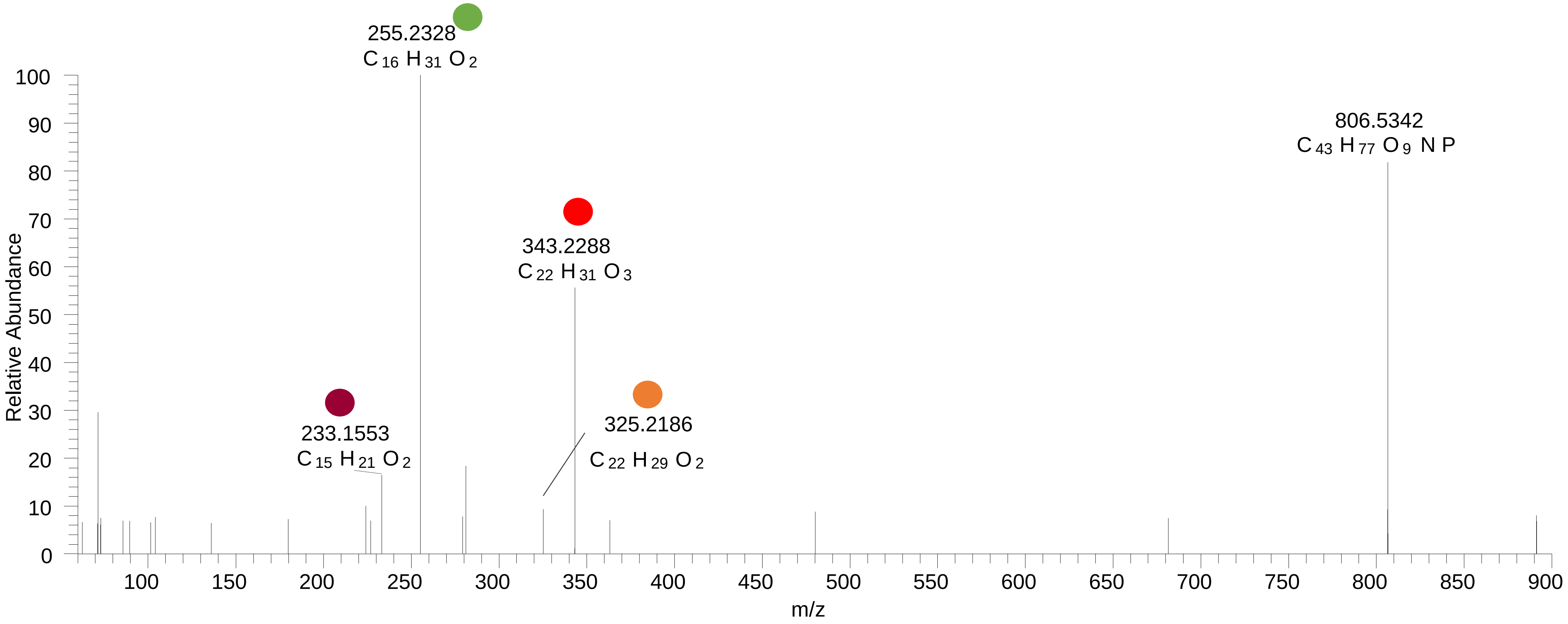

# PC(16:0_22:6<O>)
# RT 14.4

**[oxPC+HCOO]- XIC 866.5553**

- Fragments containing oxFAs
- Fragments related to water loss
- Fragments not containing oxFAs
- Fragments containing methylated oxFAs
- Fragments related to $CO_2$ loss
- Position-specific fragments

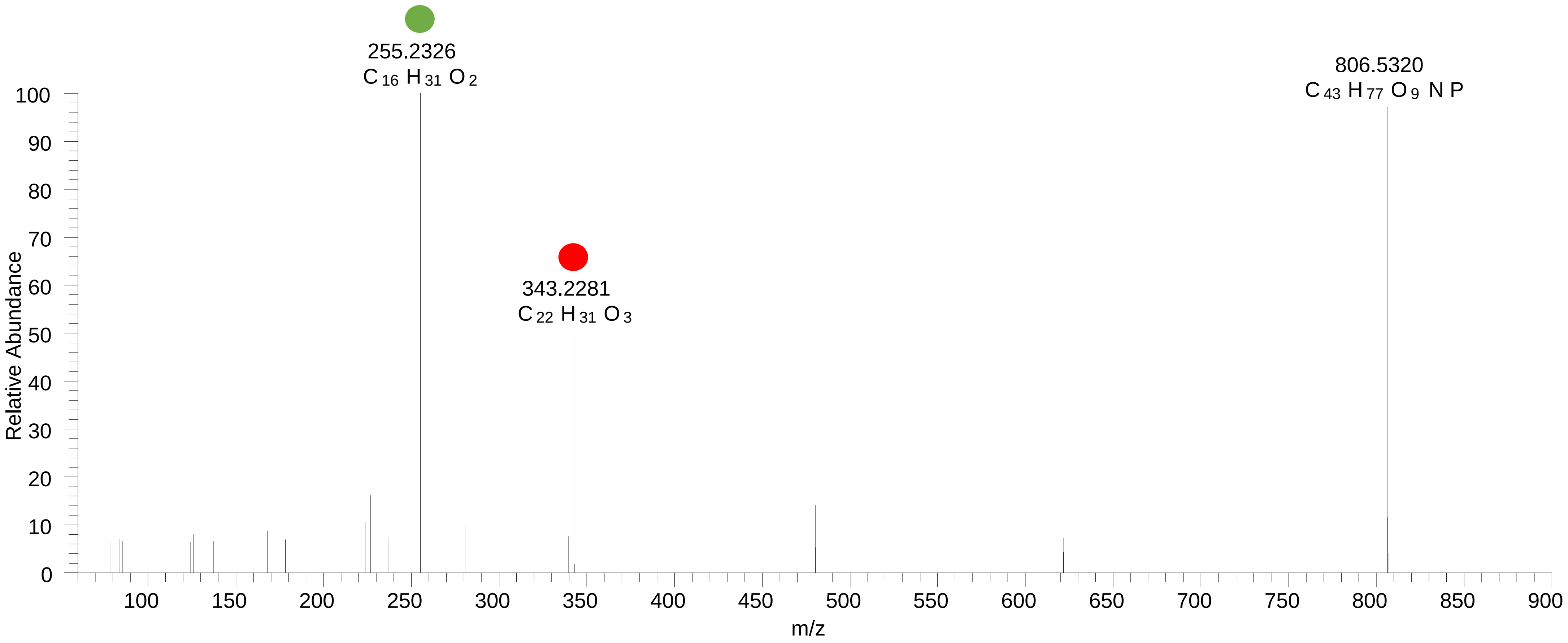

# PC(16:0_22:5<OH>)
# PC(18:1_20:4<O>)
# RT 13.7 -14.6

**[oxPC+HCOO]- XIC 868.5709**

- Fragments containing oxFAs
- Fragments related to water loss
- Fragments not containing oxFAs
- Fragments containing methylated oxFAs
- Fragments related to $CO_2$ loss
- Position-specific fragments

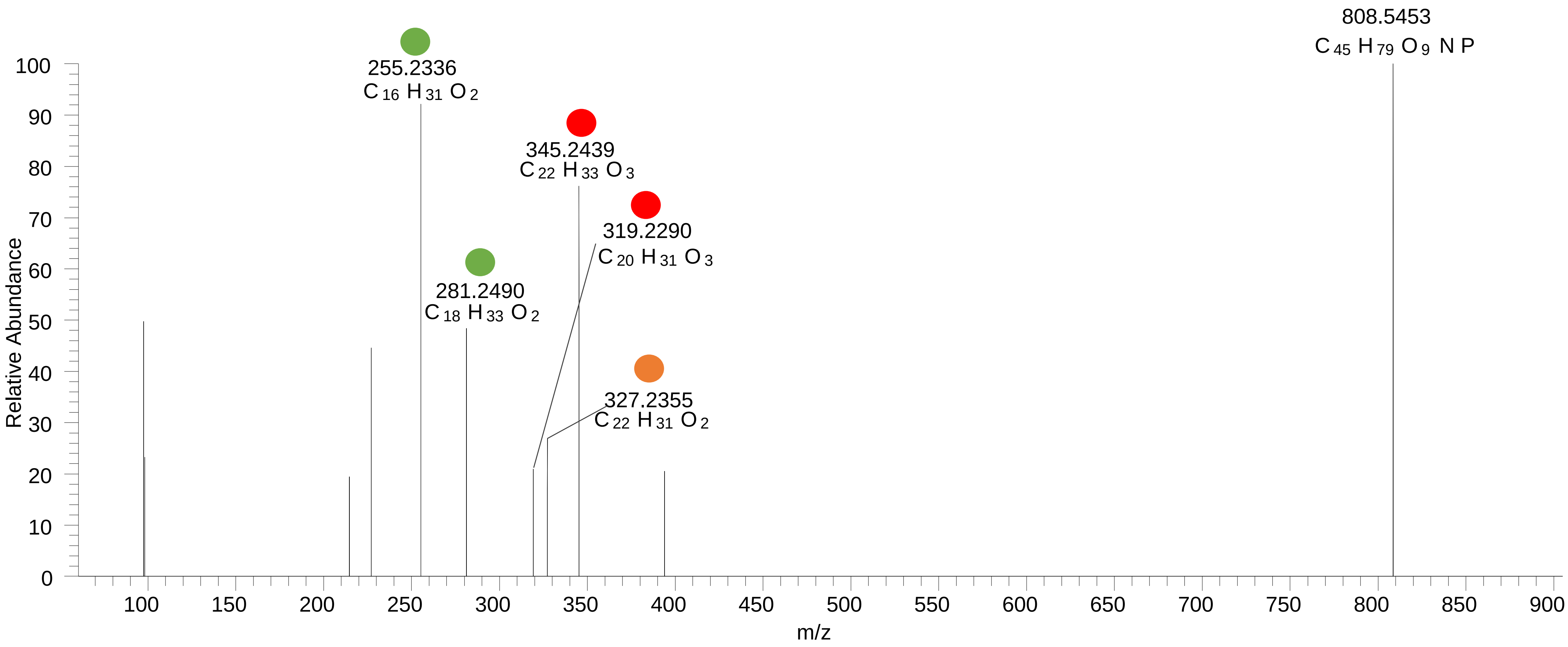

# PC(18:1_20:4<O>)
# RT 14.9

**[oxPC+HCOO]- XIC 868.5709**

Fragments containing oxFAs

Fragments related to water loss

Fragments not containing oxFAs

Fragments containing methylated oxFAs

Fragments related to $CO_2$ loss

Position-specific fragments

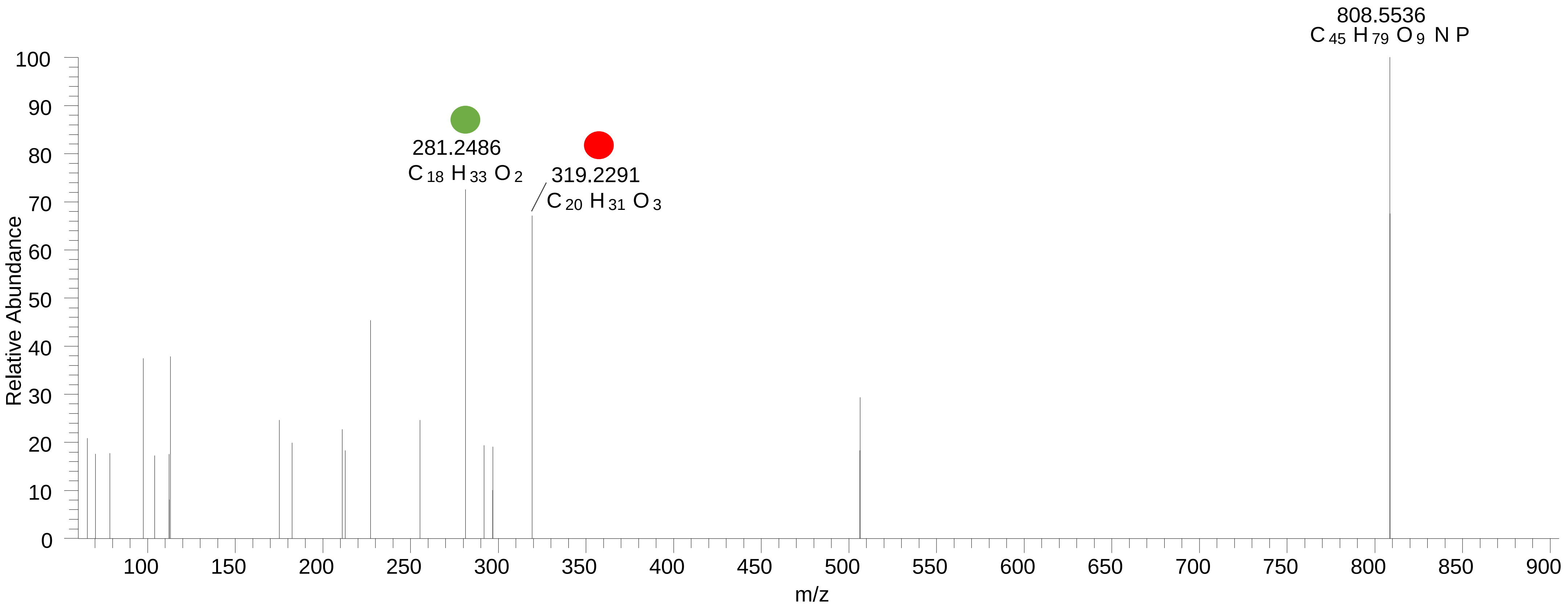

# PC(18:1_20:3<O>)
# RT 14.5

**[oxPC+HCOO]- XIC 870.5866**

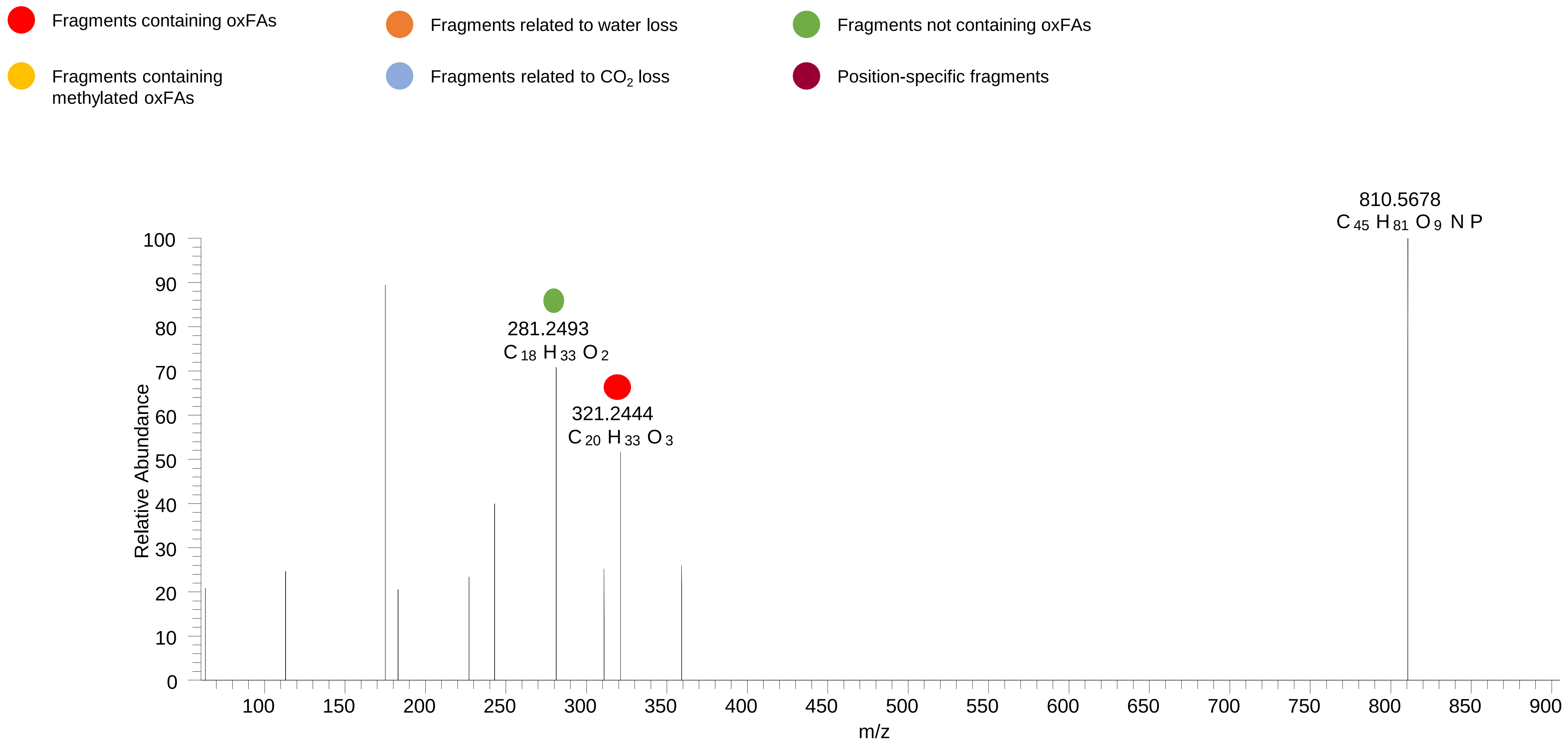

# PC(18:0_20:4<OH{15}>)
# RT 15.3

**[oxPC+HCOO]- XIC 870.5866**

- Fragments containing oxFAs
- Fragments related to water loss
- Fragments not containing oxFAs
- Fragments containing methylated oxFAs
- Fragments related to $CO_2$ loss
- Position-specific fragments

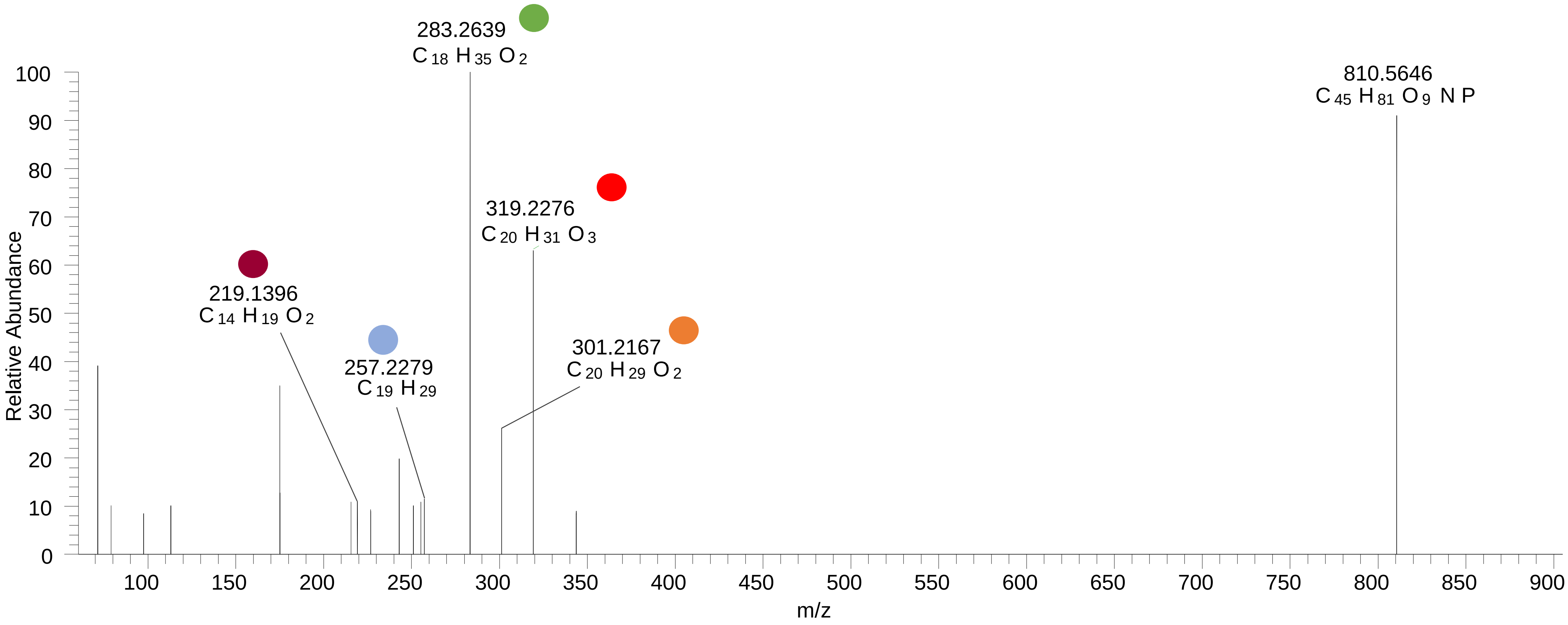

# PC(18:0_20:4<OH{11}>)
# RT 15.7

**[oxPC+HCOO]-   XIC 870.5866**

- Fragments containing oxFAs
- Fragments related to water loss
- Fragments not containing oxFAs
- Fragments containing methylated oxFAs
- Fragments related to $CO_2$ loss
- Position-specific fragments

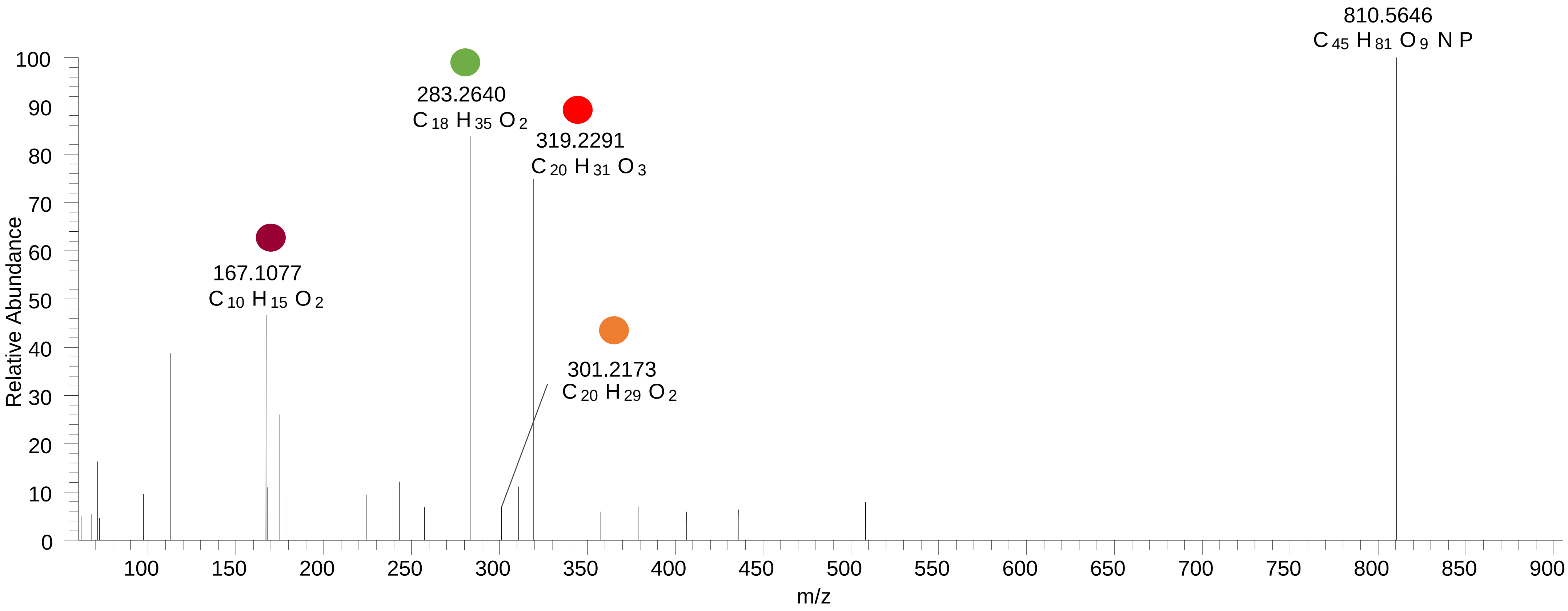

PC(18:0_20:4<OH{12}>)
PC(18:0_20:4<OH{9}>)
PC(18:0_20:4<OH{8}>)
RT 16.2

**[oxPC+HCOO]- XIC 870.5866**

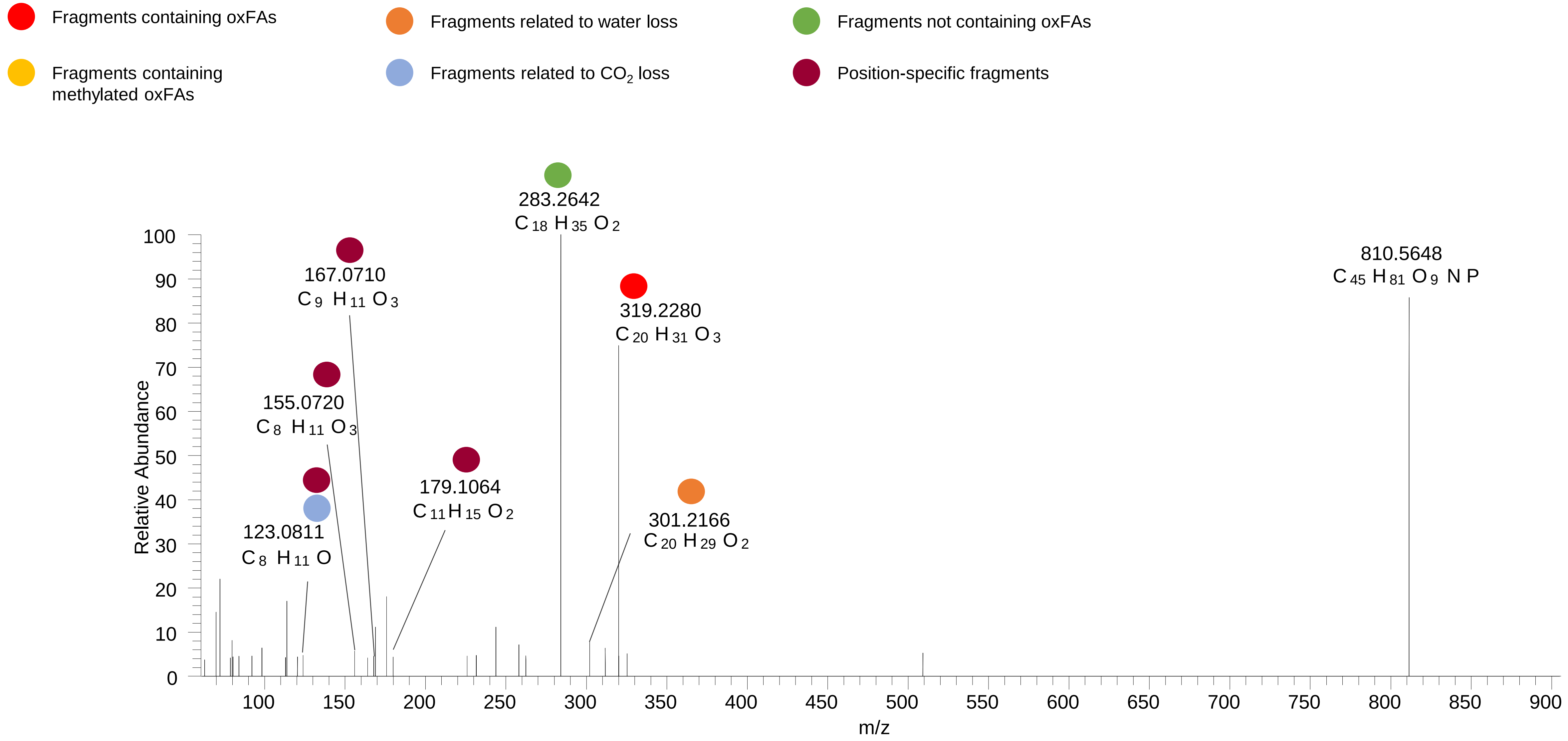

# PC(18:0_20:4<O>)
# RT 16.7

**[oxPC+HCOO]- XIC 870.5866**

Fragments containing oxFAs

Fragments related to water loss

Fragments not containing oxFAs

Fragments containing methylated oxFAs

Fragments related to $CO_2$ loss

Position-specific fragments

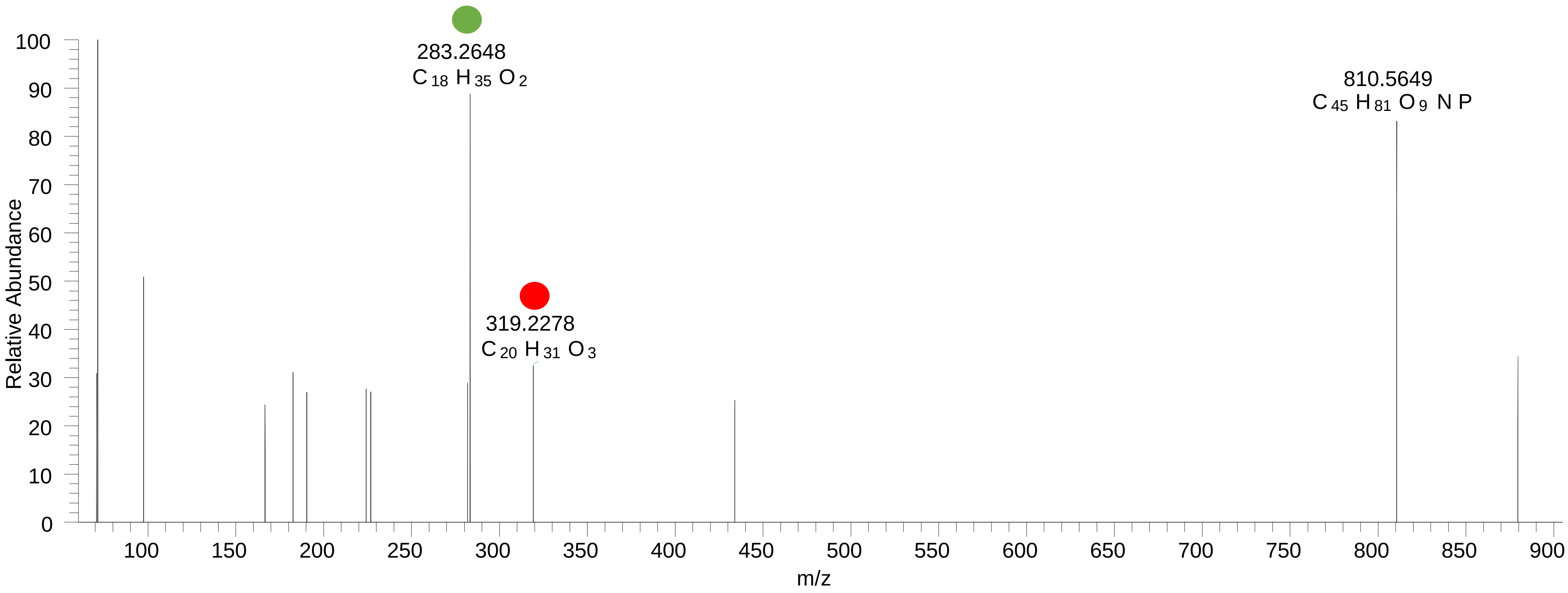

# PC(16:0_22:6<2O>)
# RT 10.9-13.4

**[oxPC+HCOO]- XIC 882.5502**

- Fragments containing oxFAs
- Fragments related to water loss
- Fragments not containing oxFAs
- Fragments containing methylated oxFAs
- Fragments related to $CO_2$ loss
- Position-specific fragments

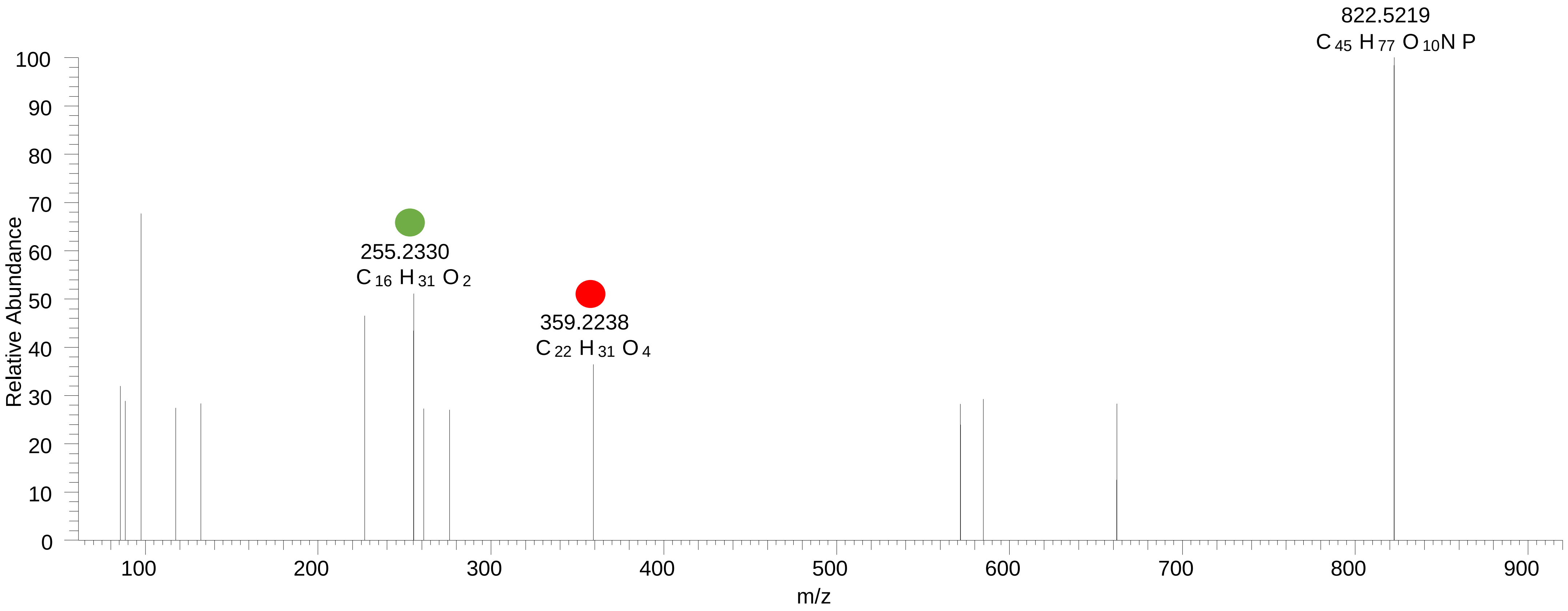

### PC(16:0_22:5<OOH>)
# RT 12.4

**[oxPC+HCOO]-   XIC 884.5658**

- Fragments containing oxFAs
- Fragments related to water loss
- Fragments not containing oxFAs
- Fragments containing methylated oxFAs
- Fragments related to $CO_2$ loss
- Position-specific fragments

T: FTMS - p ESI d Full ms2 884.5670@hcd30.00 [61.3333-920.0000]

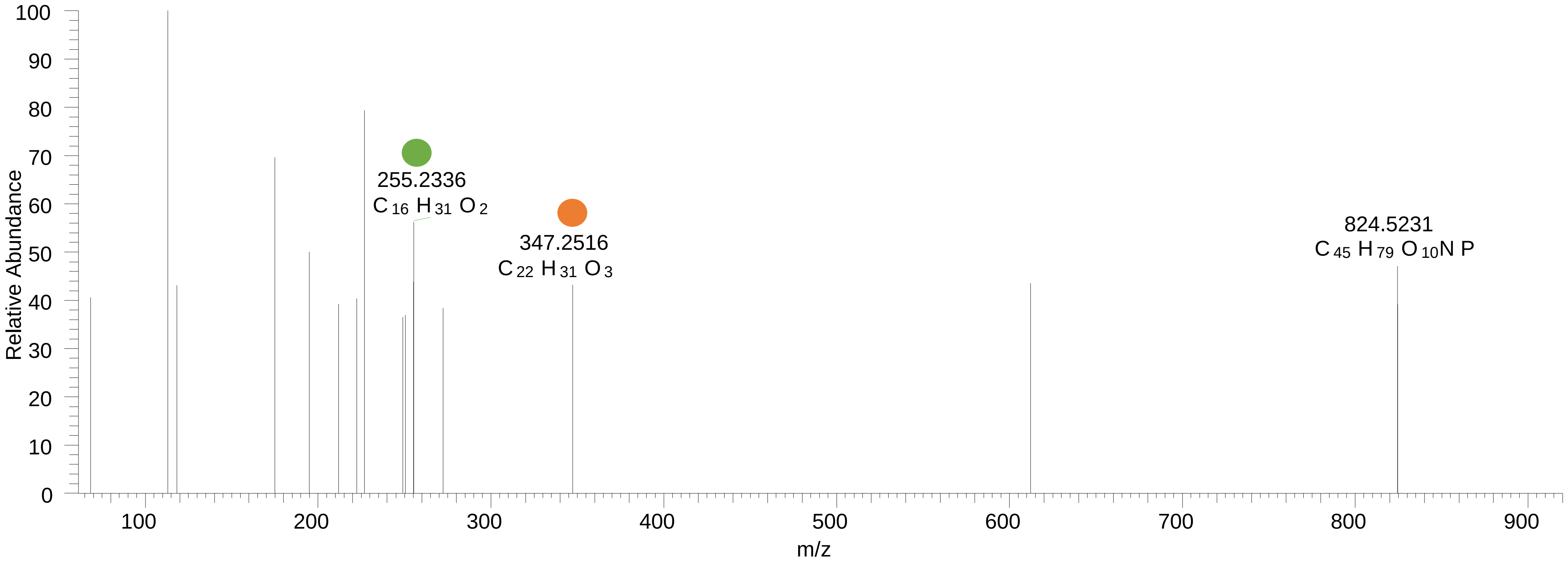

# PC(18:0_20:4<2O>)
# RT 13.0

**[oxPC+HCOO]- XIC 886.5815**

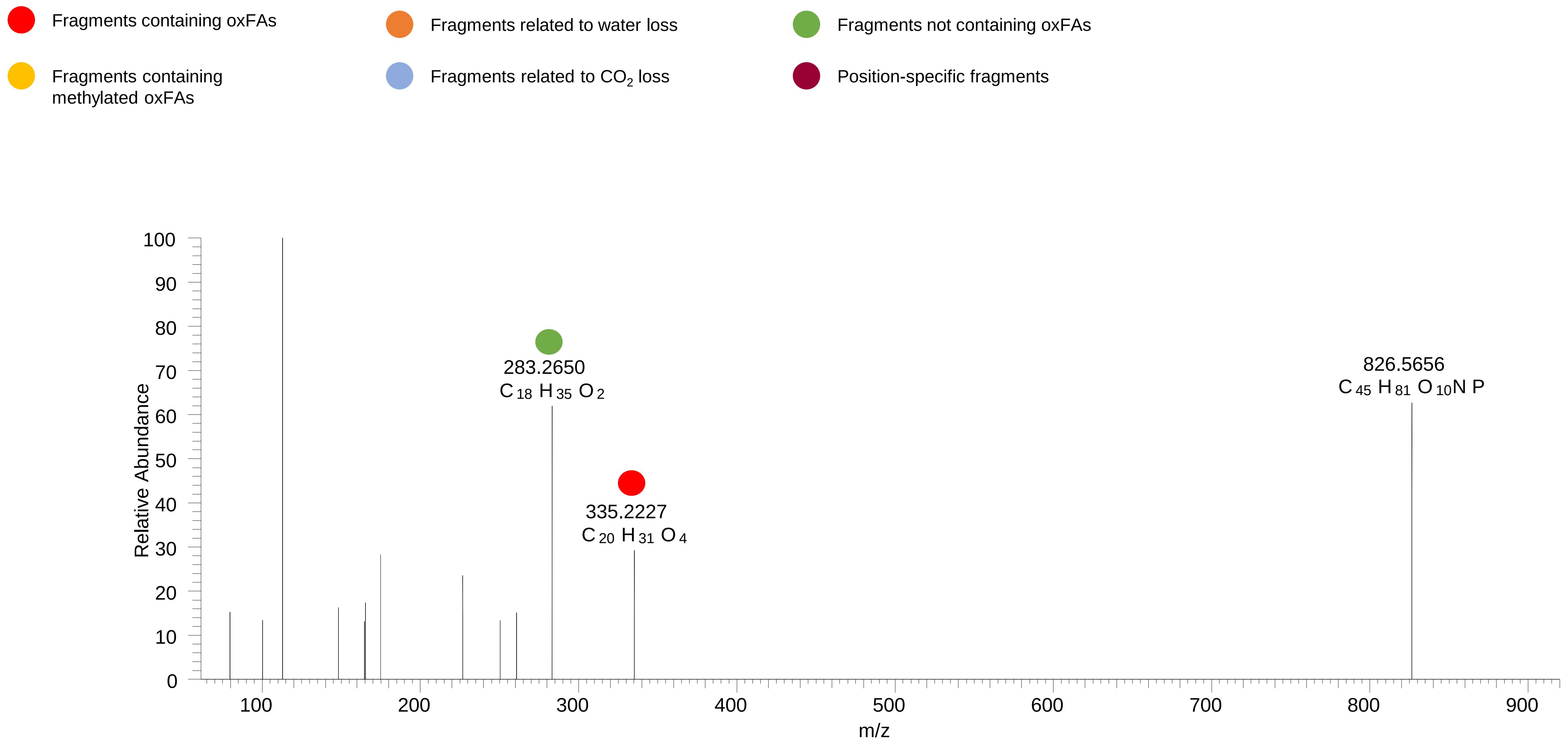

### PC(18:0_20:4<OOH>)
# RT 14.4 -15.3

**[oxPC+HCOO]- XIC 886.5815**

- Fragments containing oxFAs
- Fragments related to water loss
- Fragments not containing oxFAs
- Fragments containing methylated oxFAs
- Fragments related to $CO_2$ loss
- Position-specific fragments

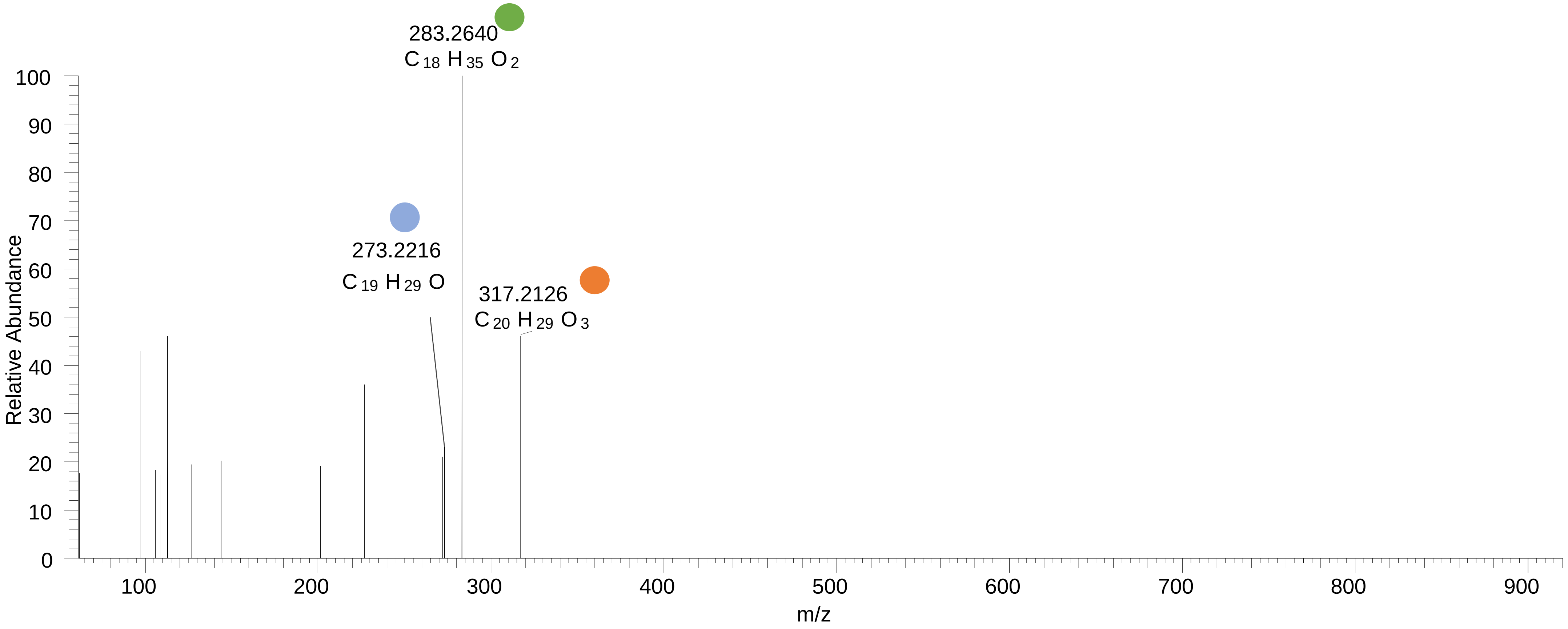

### PC(18:0_20:4<OOH{7}>)
# RT 16.0

**[oxPC+HCOO]- XIC 886.5815**

- Fragments containing oxFAs
- Fragments related to water loss
- Fragments not containing oxFAs
- Fragments containing methylated oxFAs
- Fragments related to $CO_2$ loss
- Position-specific fragments

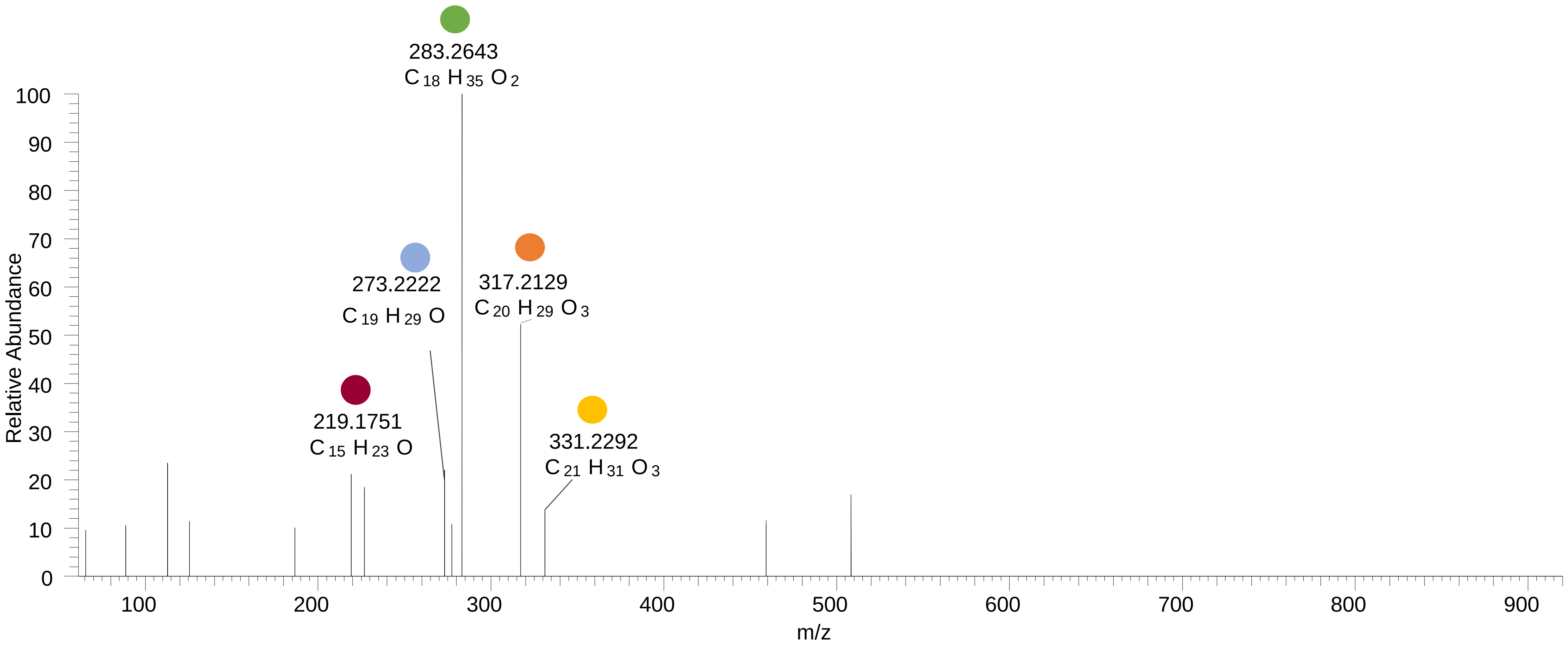

### PC(18:0_20:4<OOH>)
# RT 16.4

**[oxPC+HCOO]- XIC 886.5815**

Fragments containing oxFAs

Fragments related to water loss

Fragments not containing oxFAs

Fragments containing methylated oxFAs

Fragments related to $CO_2$ loss

Position-specific fragments

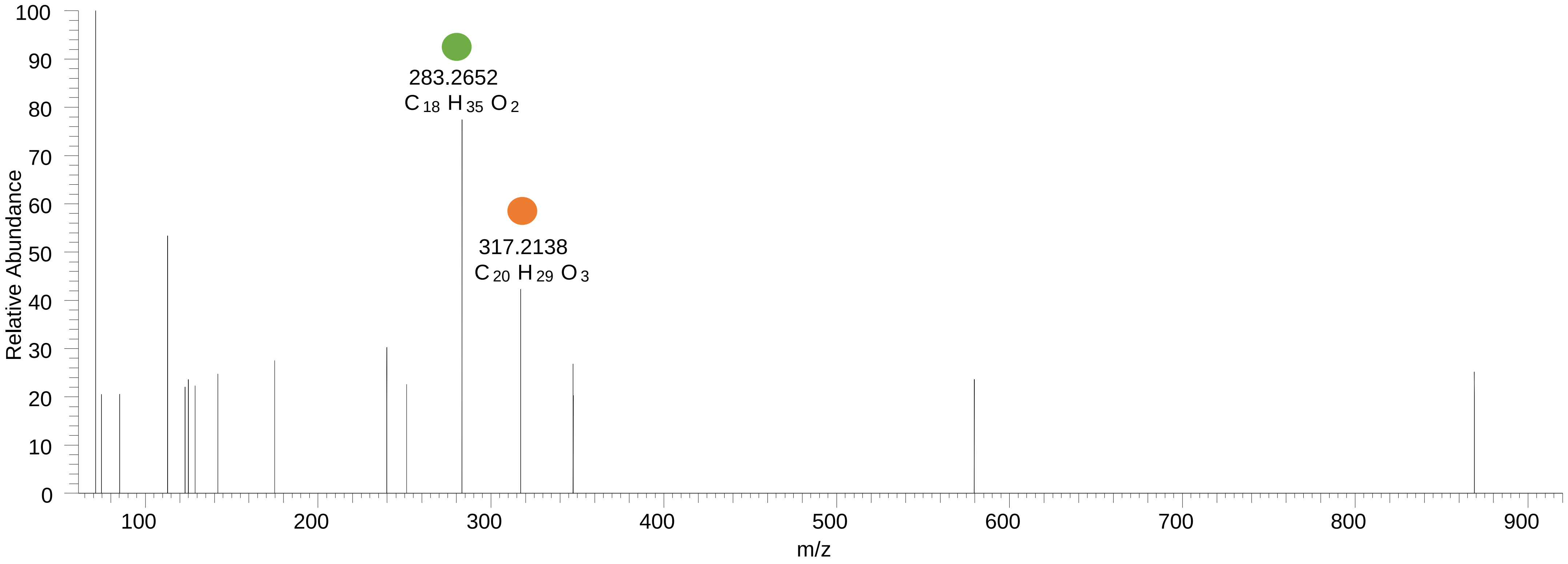

### PC(18:0_20:3<OOH>)
# RT 15.8

**[oxPC+HCOO]- XIC 888.5971**

Fragments containing oxFAs

Fragments related to water loss

Fragments not containing oxFAs

Fragments containing methylated oxFAs

Fragments related to $CO_2$ loss

Position-specific fragments

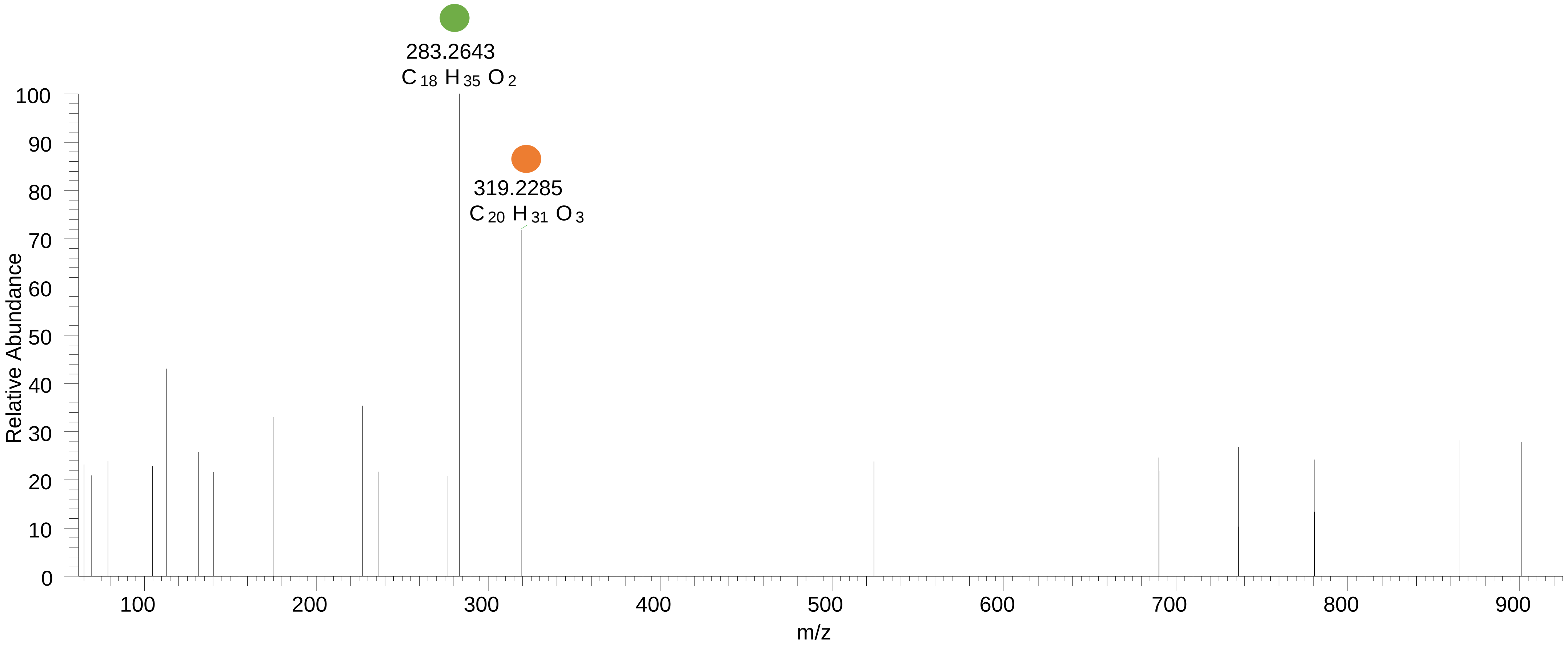

# PC(18:0_22:6<O>)
# RT 15.1-15.8

**[oxPC+HCOO]-   XIC 894.5866**

- Fragments containing oxFAs
- Fragments related to water loss
- Fragments not containing oxFAs
- Fragments containing methylated oxFAs
- Fragments related to $CO_2$ loss
- Position-specific fragments

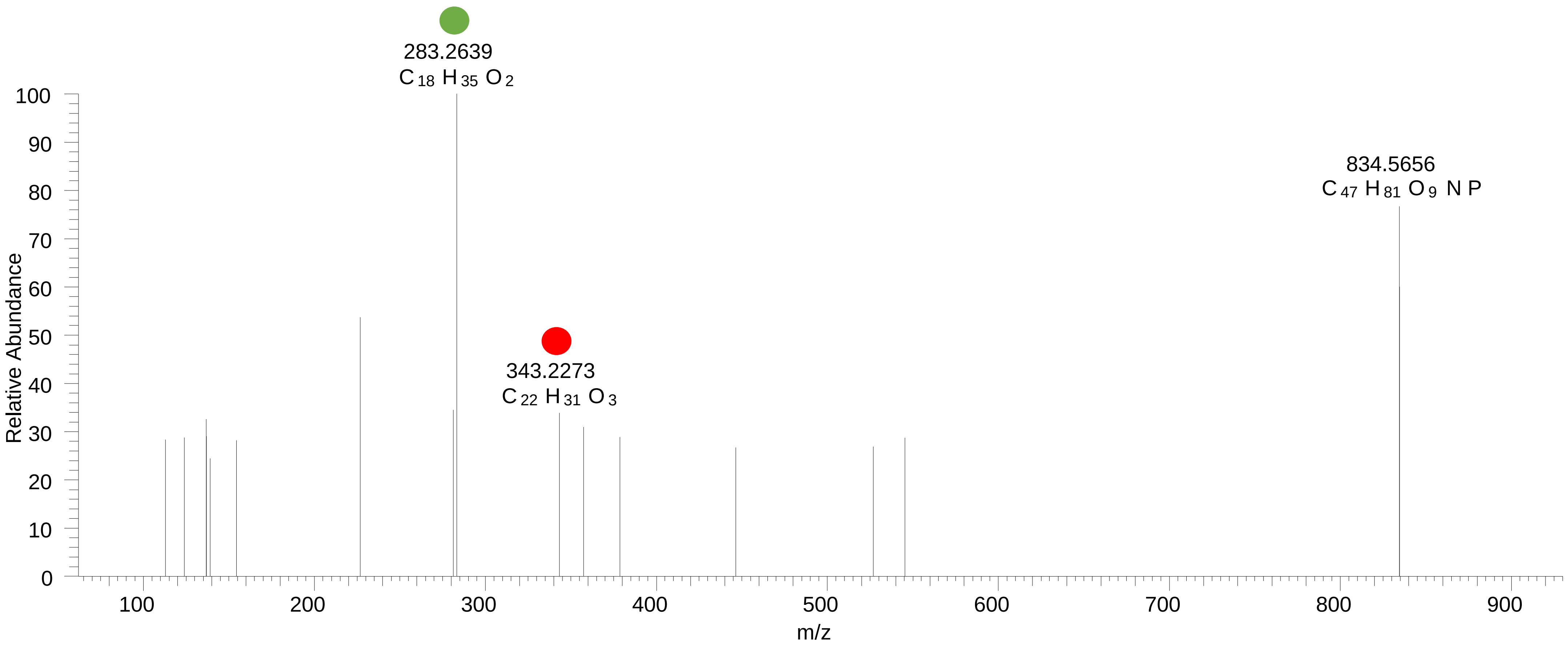
