## Supplementary material for "Epilipidomics platform for holistic profiling of oxidized complex lipids in blood plasma of obese individuals": FileS3

**Files S3.** MS2 spectra acquired using stDDA and used for the annotation of oxidized CE (oxCE) species in group pooled blood plasma samples of obese non-diabetic (OND) and obese with type 2 diabetes (OT2D) individuals. Structure-related fragmentation ions are colour-codes according to the legend provided. Annotated lipids for each group pool are sorted by their precursor *m/z*.

### OND_pool

### CE(5:0<oxo{5}>)
# RT 18.4

[oxCE+Na]+ XIC 507.3809 NL: 1.33E5

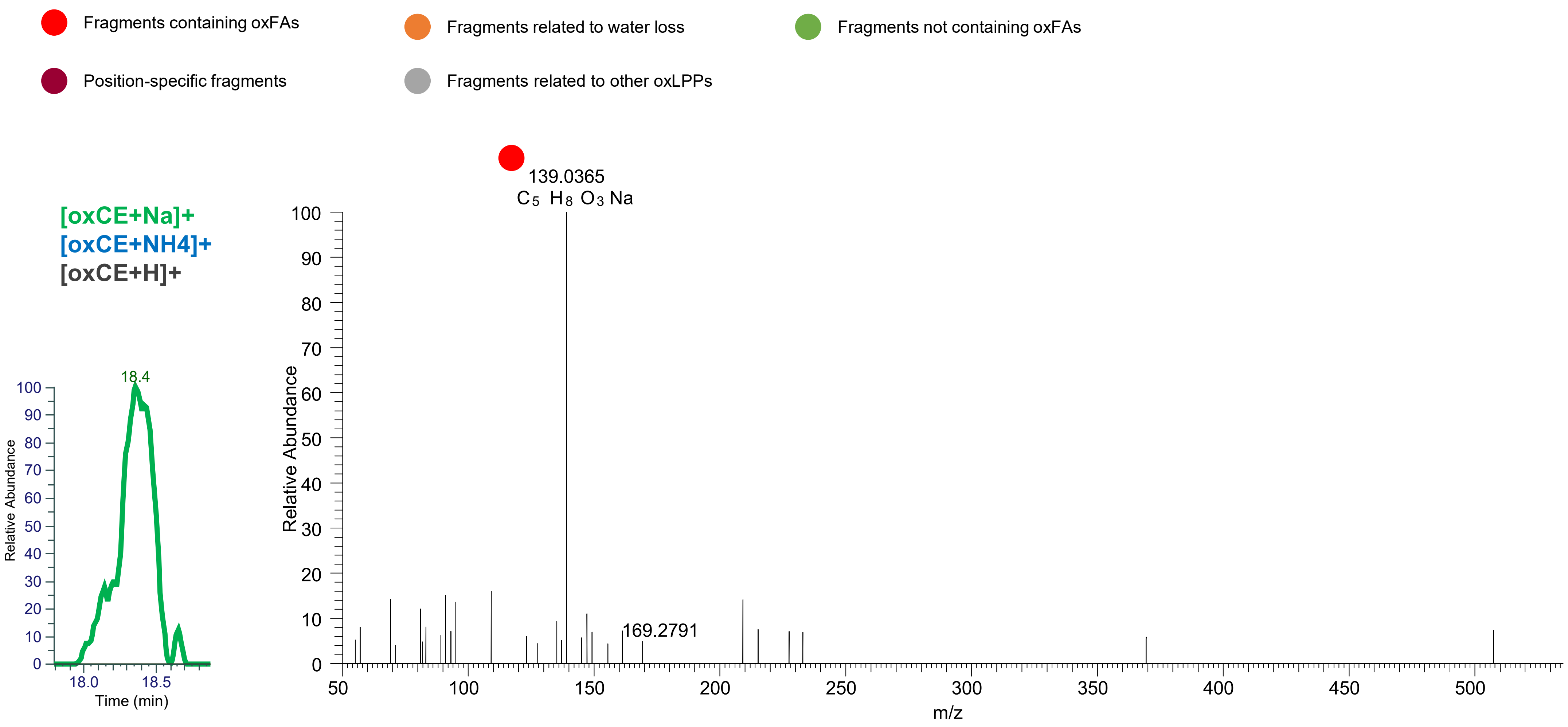

### CE(7:1<oxo{7}>)
# RT 20.7

**[oxCE+Na]+** **XIC 533.3965 NL: 6.88E4**

- Fragments containing oxFAs
- Fragments related to water loss
- Fragments not containing oxFAs
- Position-specific fragments
- Fragments related to other oxLPPs

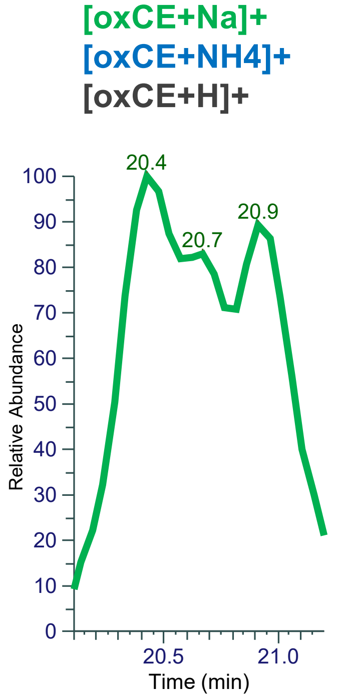


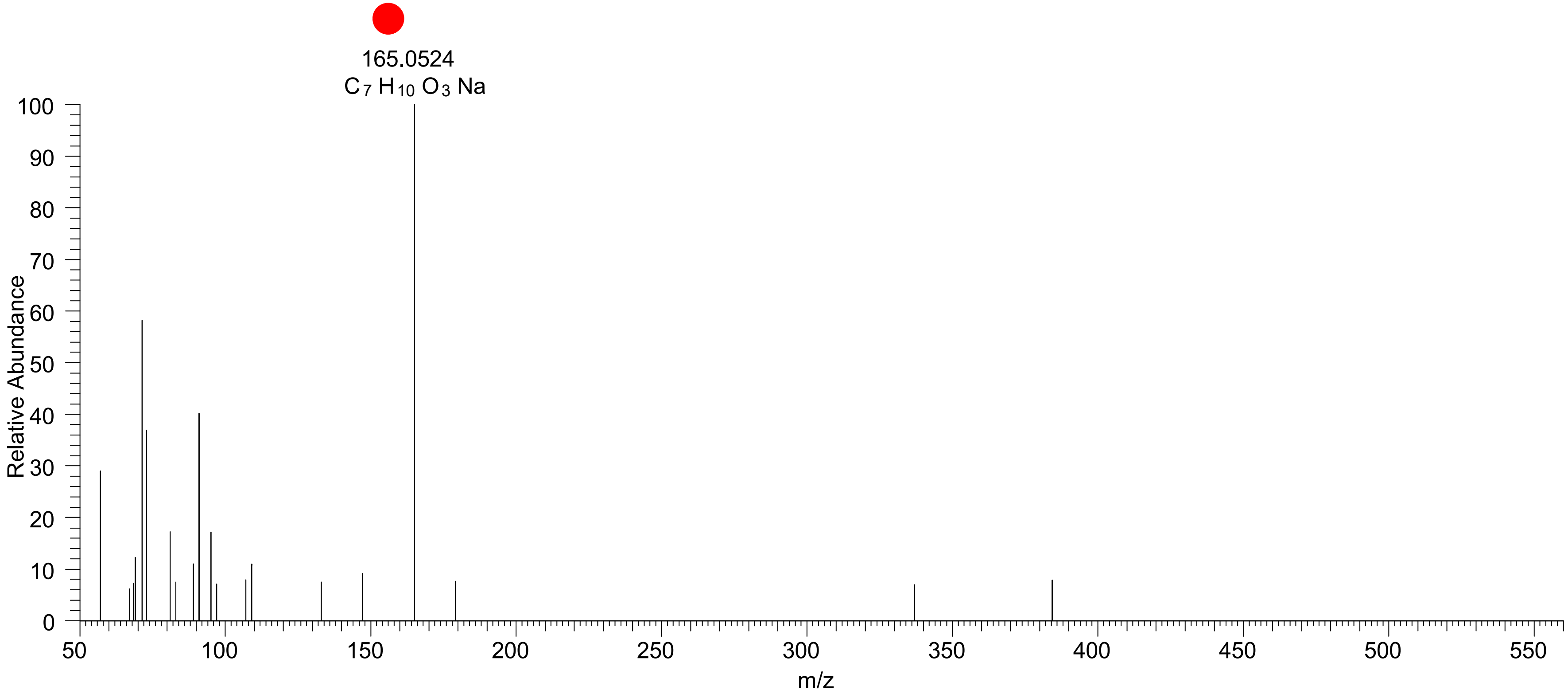

### CE(7:0<oxo{7}>)
# RT 18.7

**[oxCE+Na]+** **XIC 535.4122 NL: 7.18E4**

- Fragments containing oxFAs
- Fragments related to water loss
- Fragments not containing oxFAs
- Position-specific fragments
- Fragments related to other oxLPPs

**[oxCE+Na]+**
**[oxCE+NH4]+**
**[oxCE+H]+**

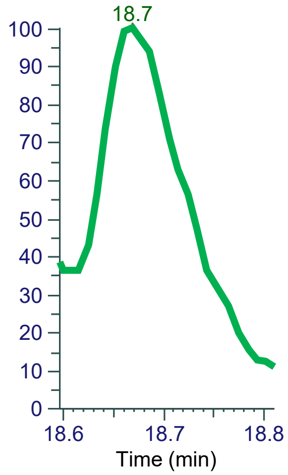


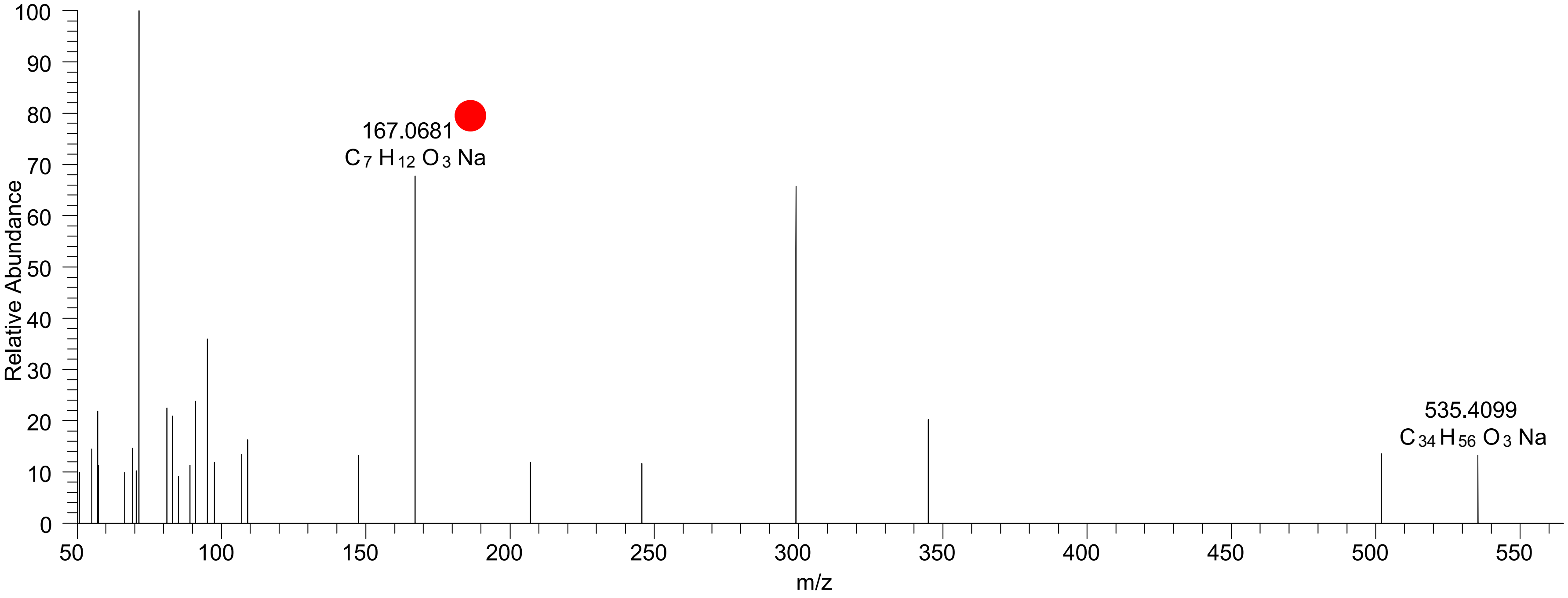

### CE(9:0<oxo{9}>)
# RT 20.9

**[oxCE+Na]+** **XIC 563.4435 NL: 8.78E5**

- Fragments containing oxFAs
- Fragments related to water loss
- Fragments not containing oxFAs
- Position-specific fragments
- Fragments related to other oxLPPs

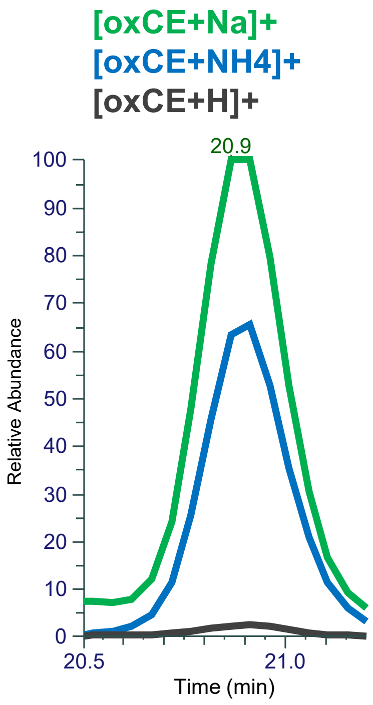


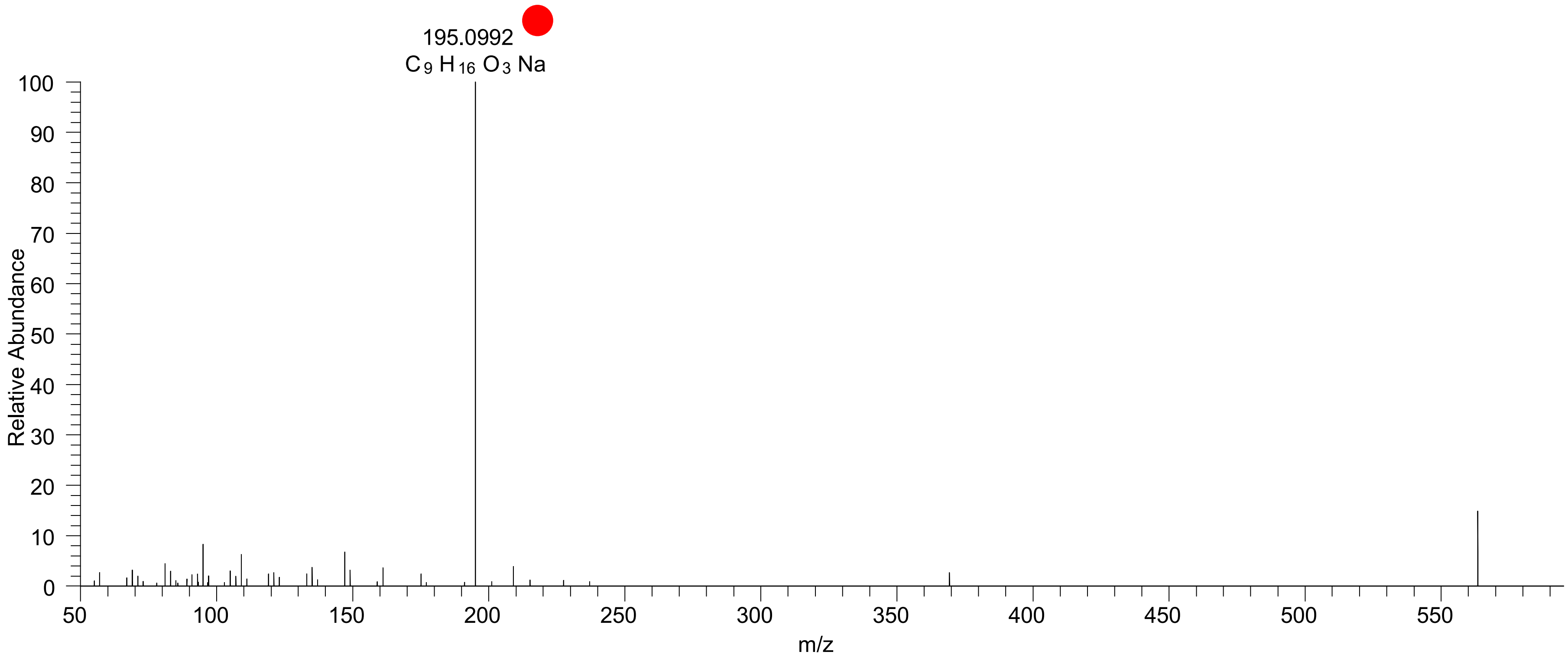

### CE(10:2<oxo{10}>)
# RT 20.8

[oxCE+Na]+ XIC 573.4278 NL: 2.02E5

Fragments containing oxFAs
Fragments related to water loss
Fragments not containing oxFAs
Position-specific fragments
Fragments related to other oxLPPs

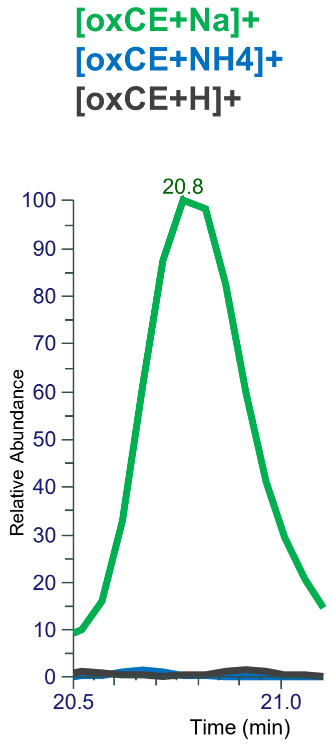


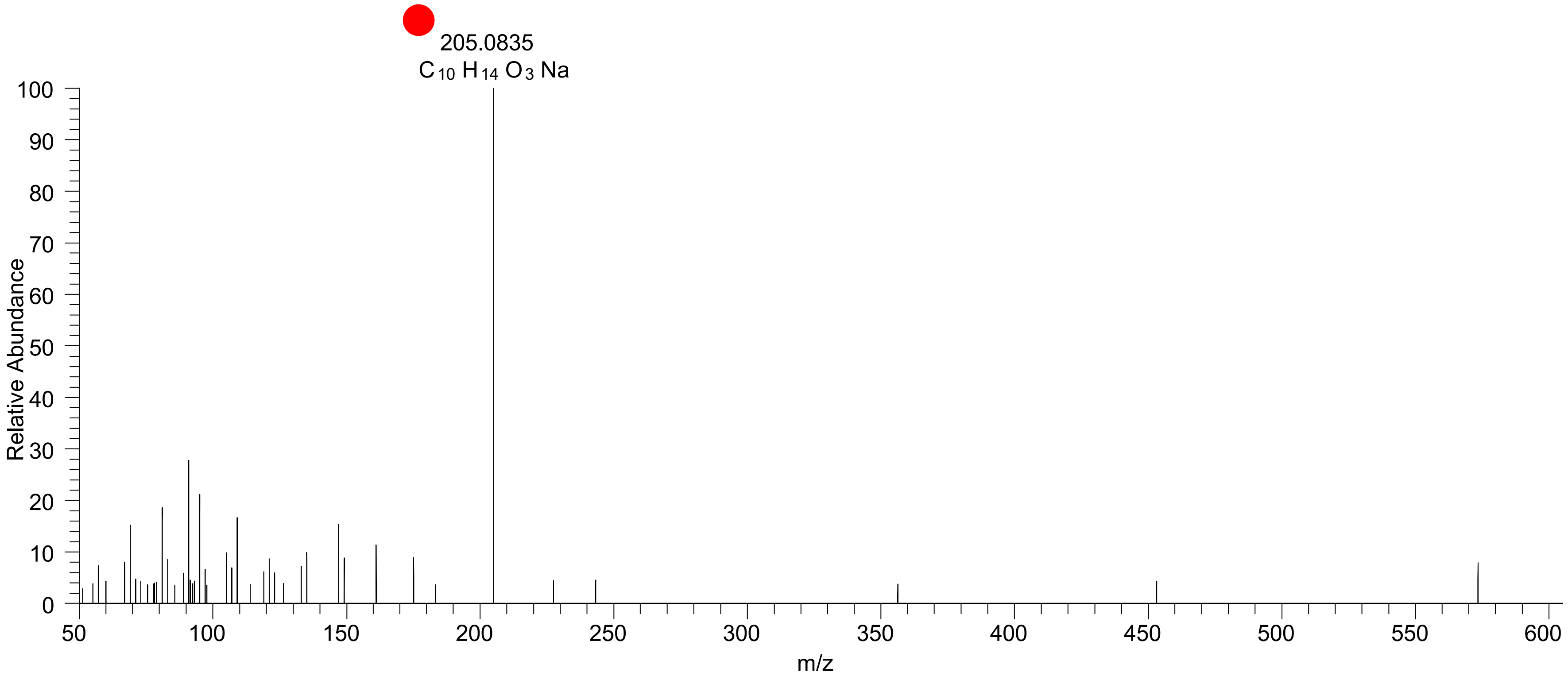

### CE(10:1<oxo{10}>)
# RT 20.7

**[oxCE+Na]+** **XIC 575.4440 NL: 1.58E4**

- Fragments containing oxFAs
- Fragments related to water loss
- Fragments not containing oxFAs
- Position-specific fragments
- Fragments related to other oxLPPs

**[oxCE+Na]+**
**[oxCE+NH4]+**
**[oxCE+H]+**

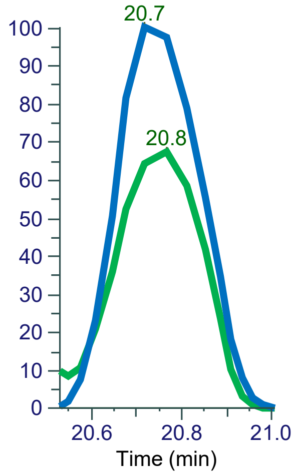


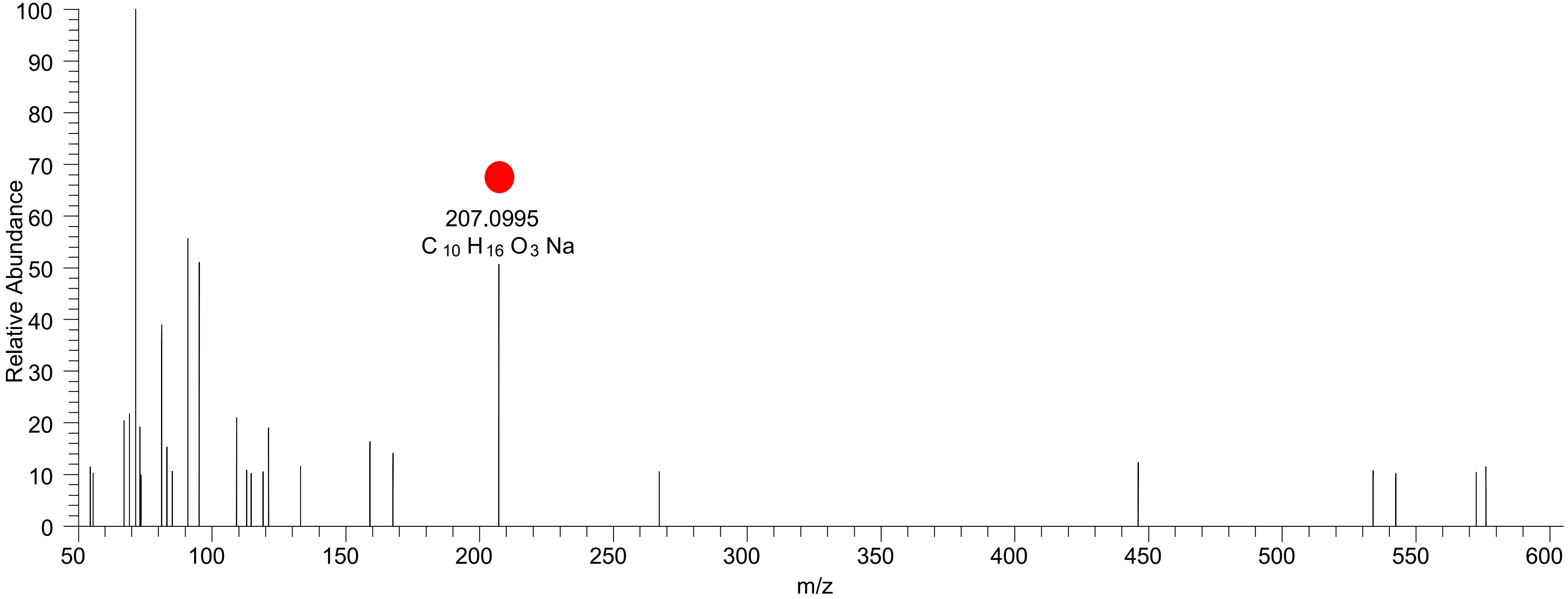

### CE(11:2<oxo{11}>)
# RT 20.3

**[oxCE+Na]+** **XIC 587.4440 NL: 3.04E4**

- Fragments containing oxFAs
- Fragments related to water loss
- Fragments not containing oxFAs
- Position-specific fragments
- Fragments related to other oxLPPs

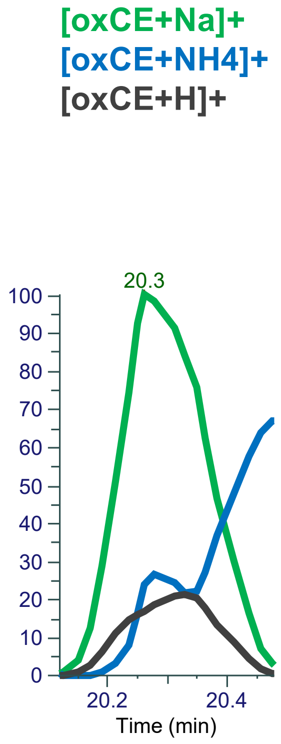


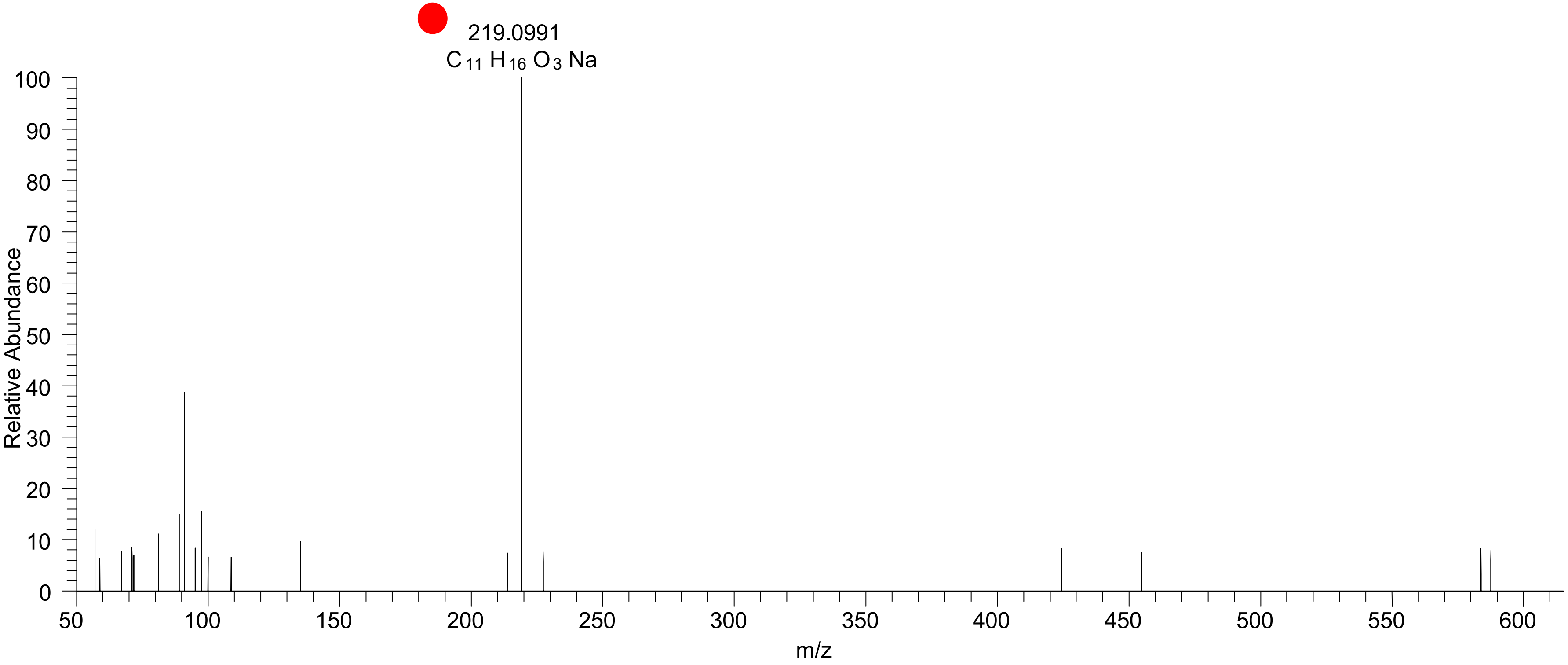

### CE(11:1<oxo{11}>)
# RT 21.4

**[oxCE+Na]+** **XIC 589.4591 NL: 1.12E5**

- Fragments containing oxFAs
- Fragments related to water loss
- Fragments not containing oxFAs
- Position-specific fragments
- Fragments related to other oxLPPs

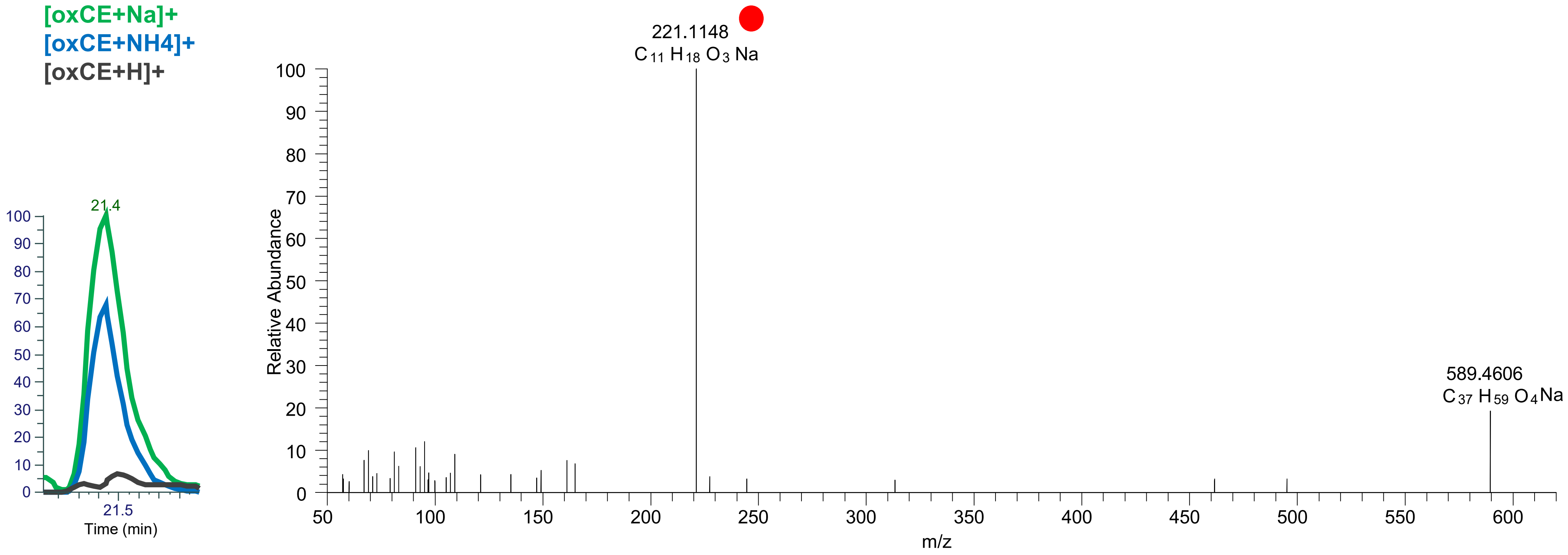

### CE(10:1<COOH{10}>)
# RT 19.8

[oxCE+Na]+ XIC 591.4384 NL: 2.06E4

Fragments containing oxFAs

Fragments related to water loss

Fragments not containing oxFAs

Position-specific fragments

Fragments related to other oxLPPs

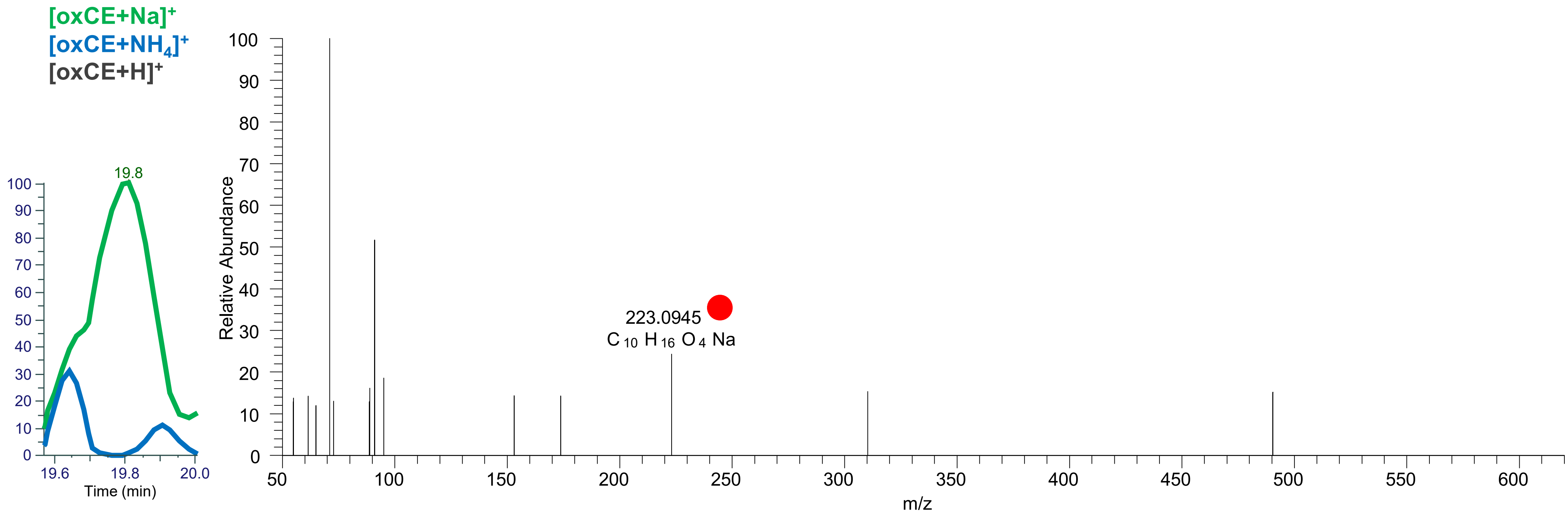

### CE(12:2<COOH{12}>)
# RT 20.6

**[oxCE+Na]+** **XIC 617.4540 NL: 1.48E4**

- Fragments containing oxFAs
- Fragments related to water loss
- Fragments not containing oxFAs
- Position-specific fragments
- Fragments related to other oxLPPs

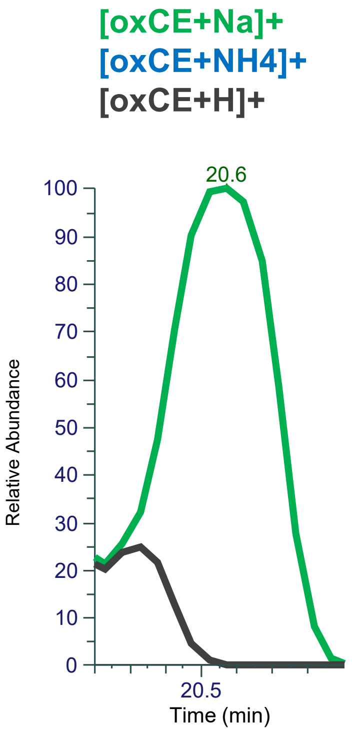


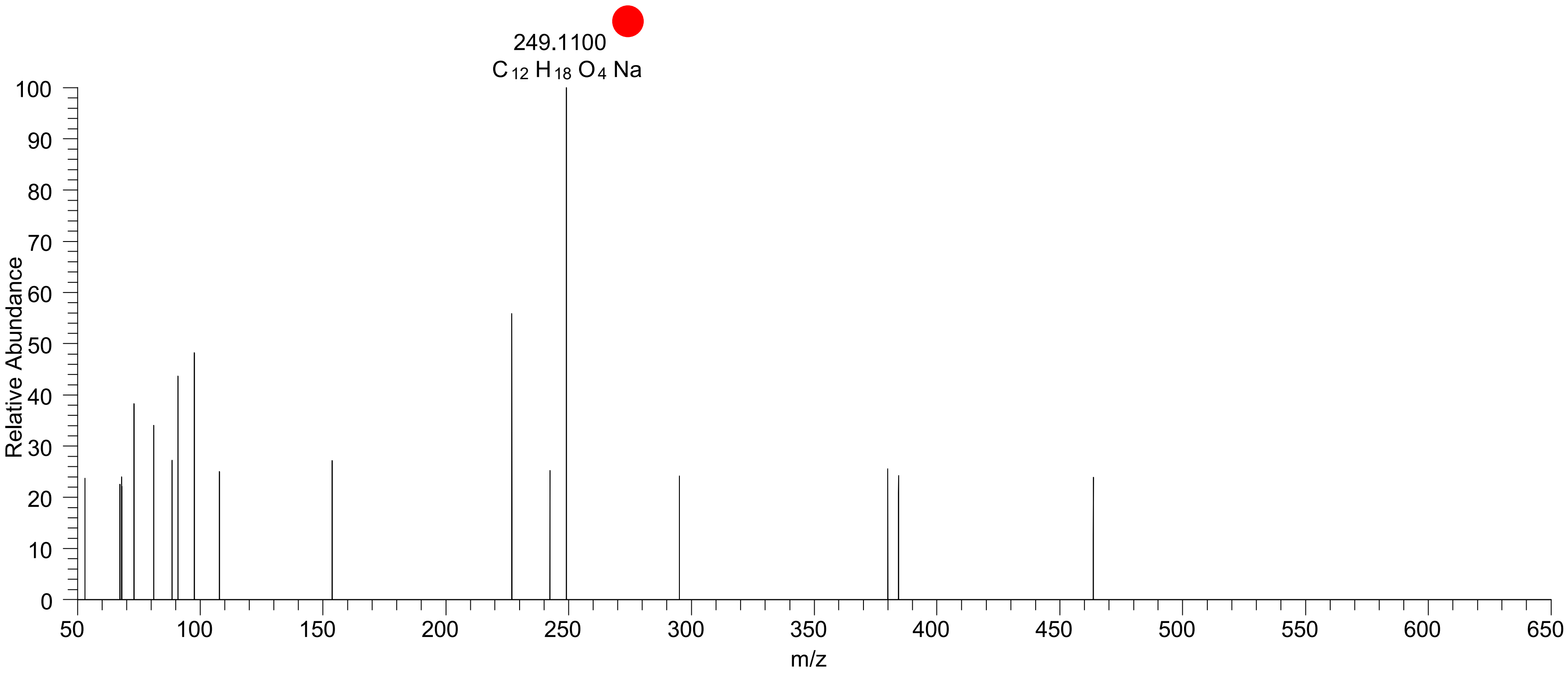

### CE(12:1<COOH{12}>)
# RT 20.3

[oxCE+Na]+ XIC 619.4697 NL: 3.86E4

Fragments containing oxFAs
Fragments related to water loss
Fragments not containing oxFAs
Position-specific fragments
Fragments related to other oxLPPs

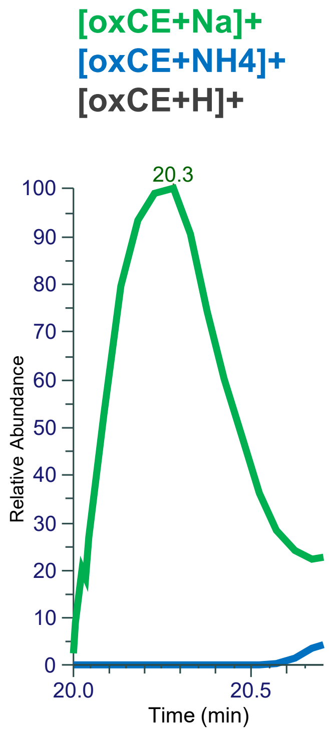


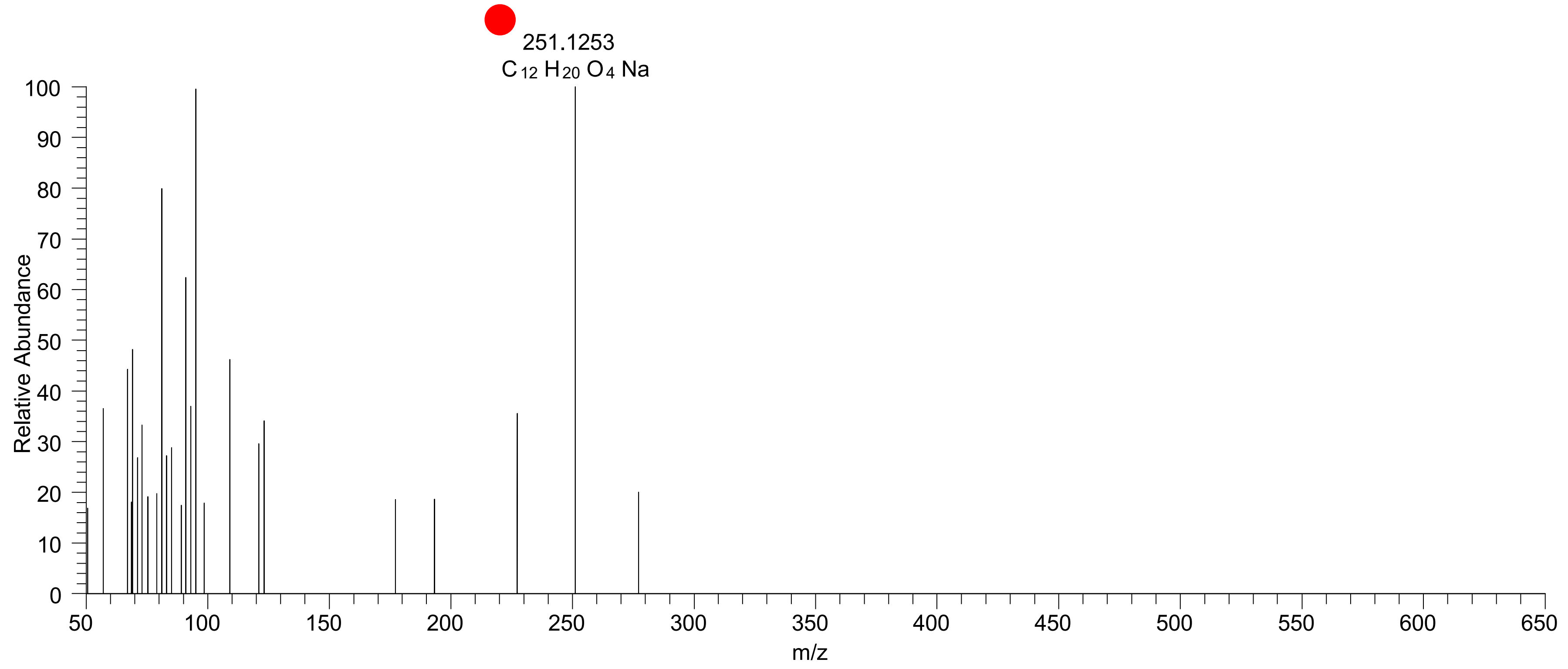

# CE(18:3<O>)
# RT 22.6

**[oxCE+Na]+** **XIC 685.5530 NL: 6.866E4**

- Fragments containing oxFAs
- Fragments related to water loss
- Fragments not containing oxFAs
- Position-specific fragments
- Fragments related to other oxLPPs

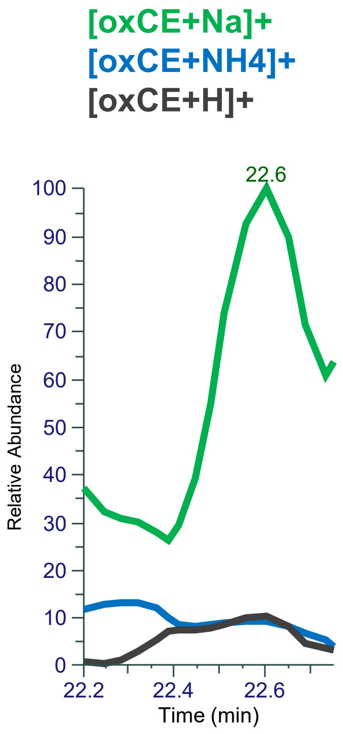


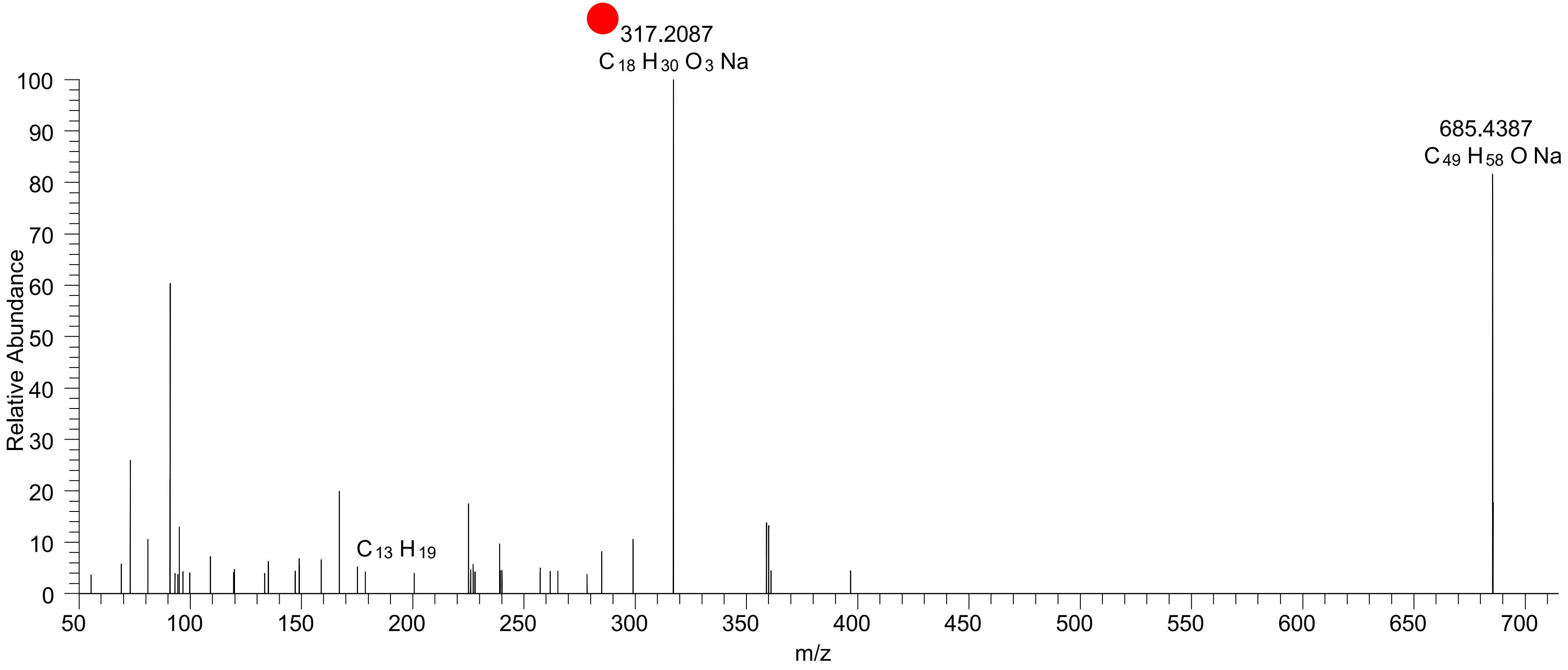

# CE(18:3<OH>)
# RT 23.3

**[oxCE+Na]+** **XIC 685.5530 NL: 1.89E6**

- Fragments containing oxFAs
- Fragments related to water loss
- Fragments not containing oxFAs
- Position-specific fragments
- Fragments related to other oxLPPs

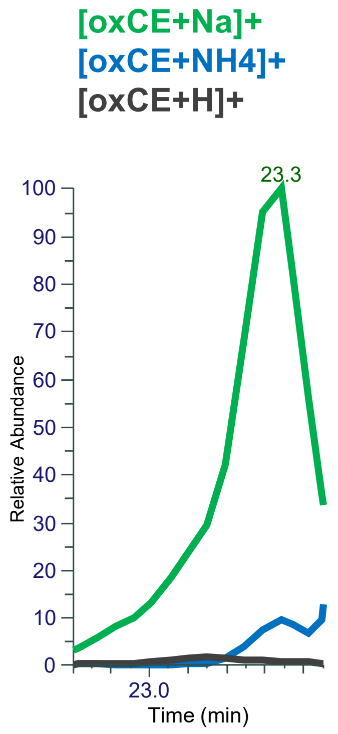


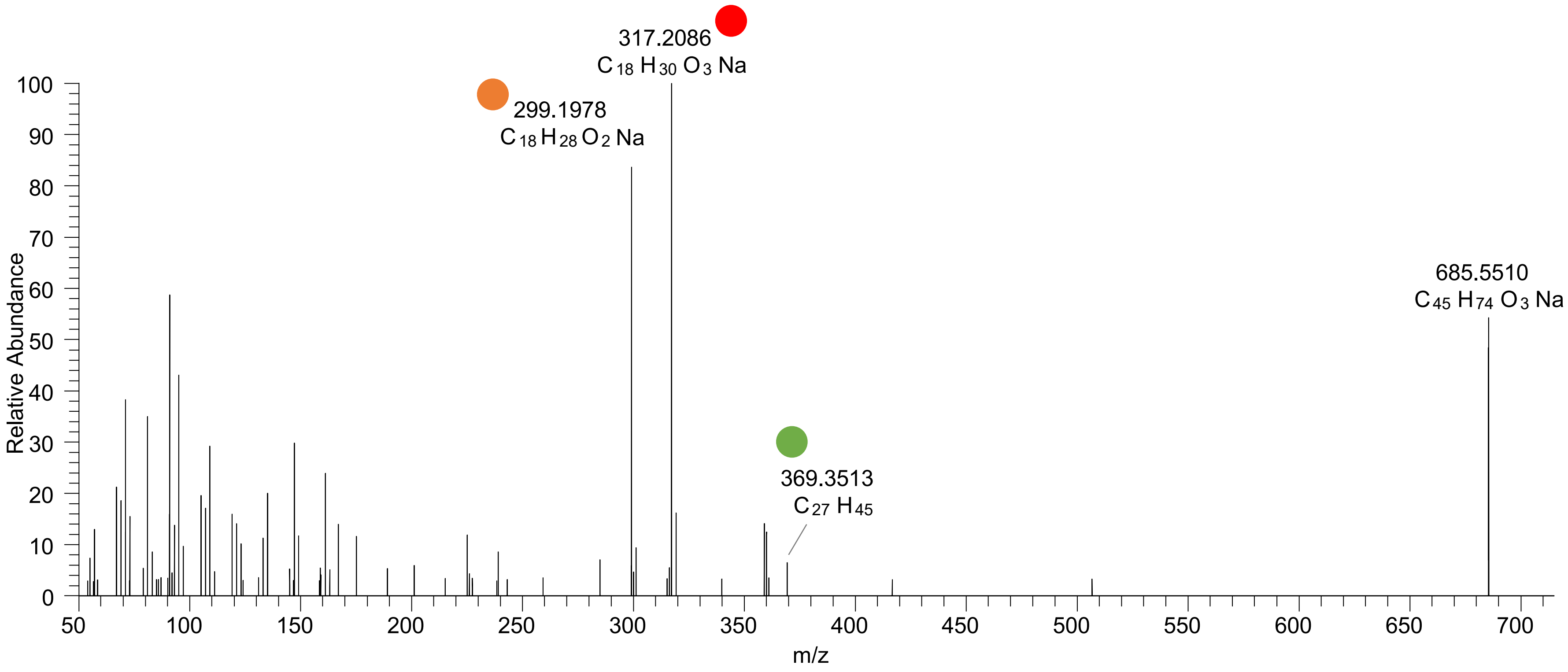

# CE(18:3<O>)
# RT 24.1

**[oxCE+Na]+** **XIC 685.5530 NL: 7.54E6**

- Fragments containing oxFAs
- Fragments related to water loss
- Fragments not containing oxFAs
- Position-specific fragments
- Fragments related to other oxLPPs

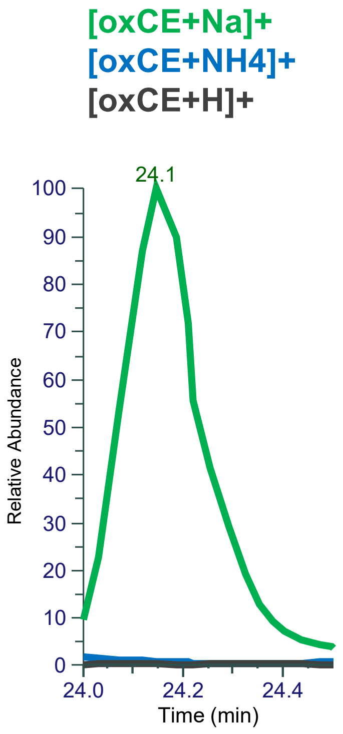


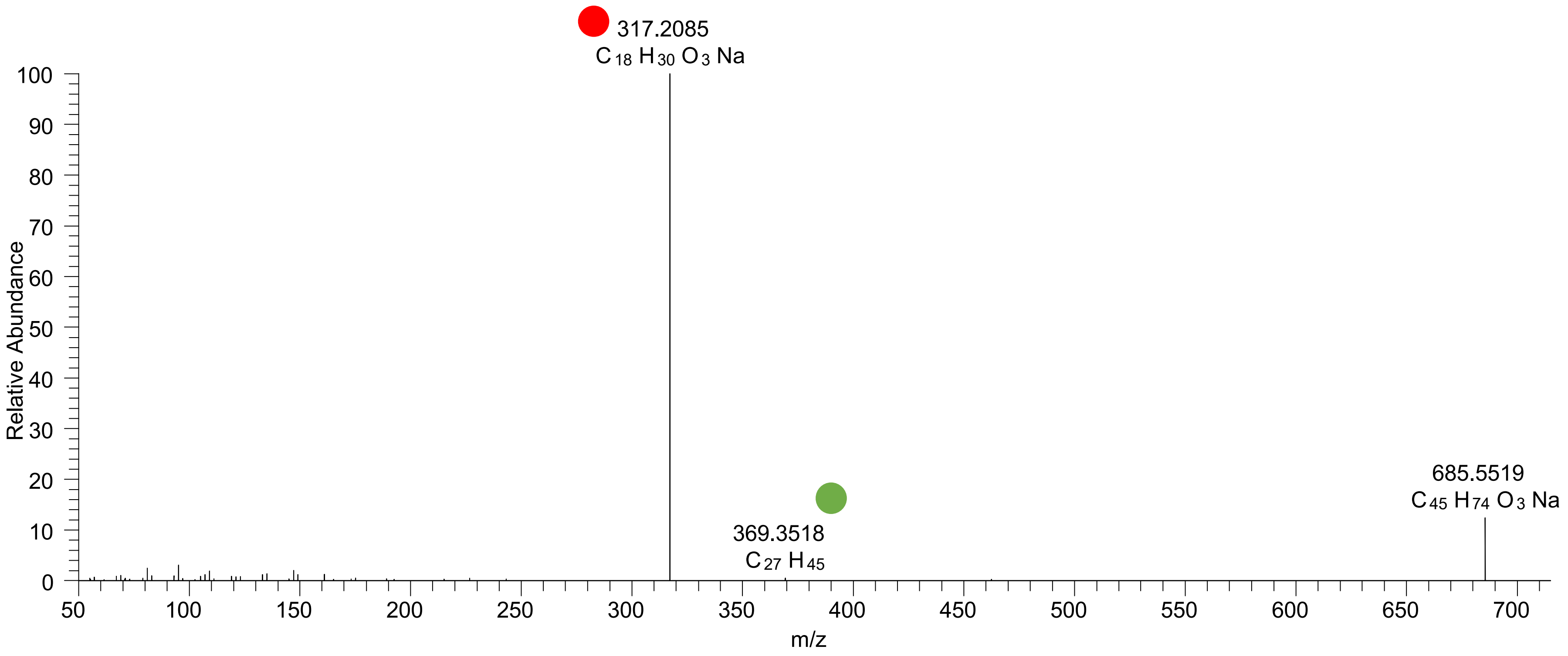

# CE(18:3<OH>)
# RT 25.0

**[oxCE+Na]+** **XIC 685.5530 NL: 3.68E5**

- Fragments containing oxFAs
- Fragments related to water loss
- Fragments not containing oxFAs
- Position-specific fragments
- Fragments related to other oxLPPs

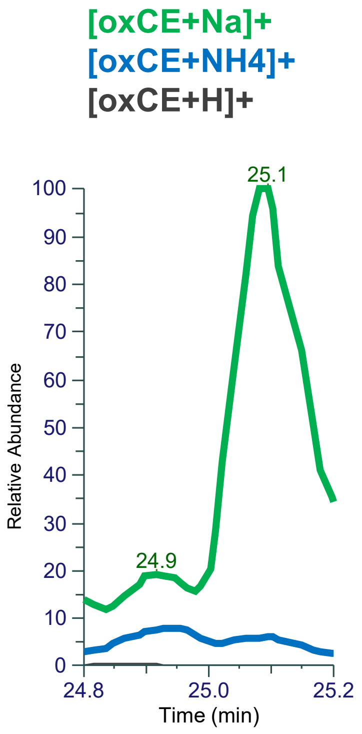


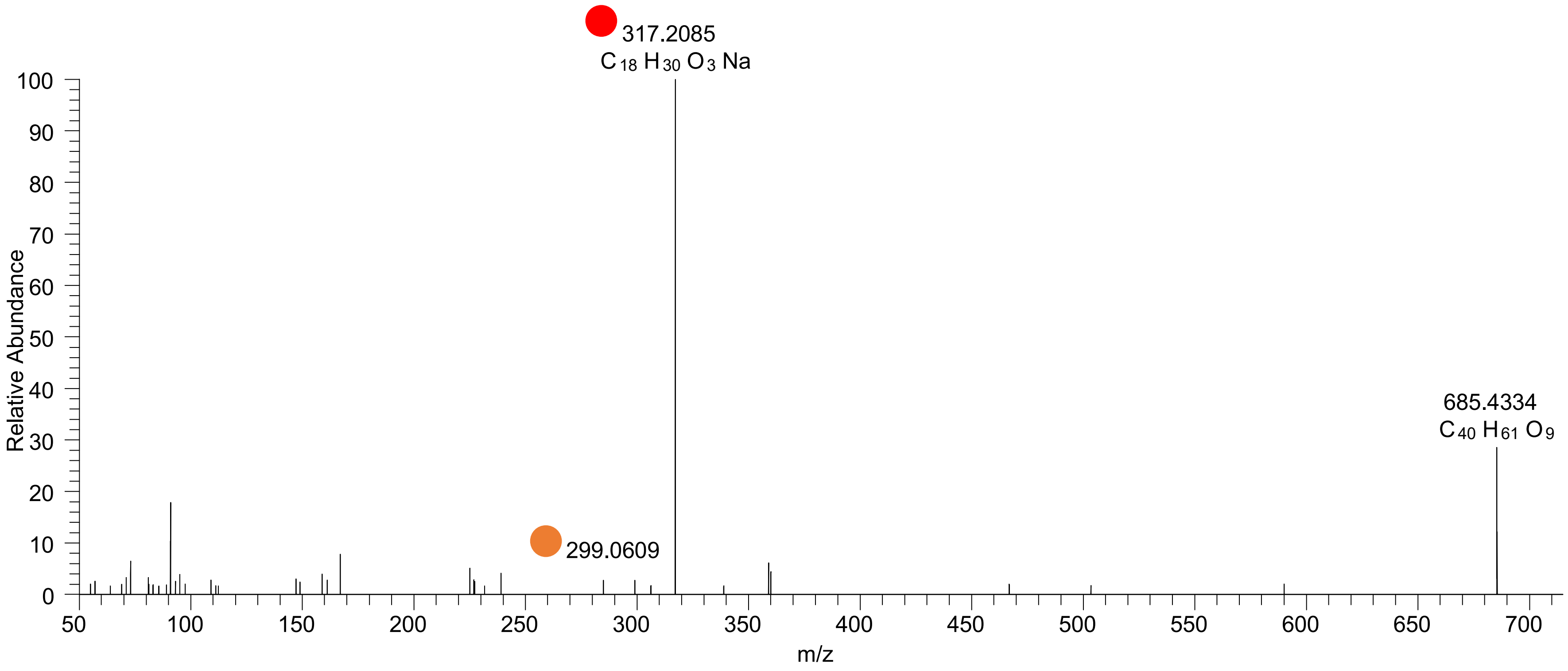

# CE(18:2<OH>)
# RT 23.5, 23.8

**[oxCE+Na]+**    **XIC 687.5687 NL: 1.71E7**

- Fragments containing oxFAs
- Fragments related to water loss
- Fragments not containing oxFAs
- Position-specific fragments
- Fragments related to other oxLPPs

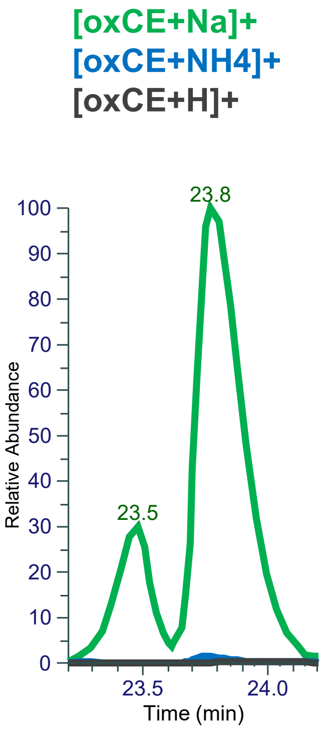


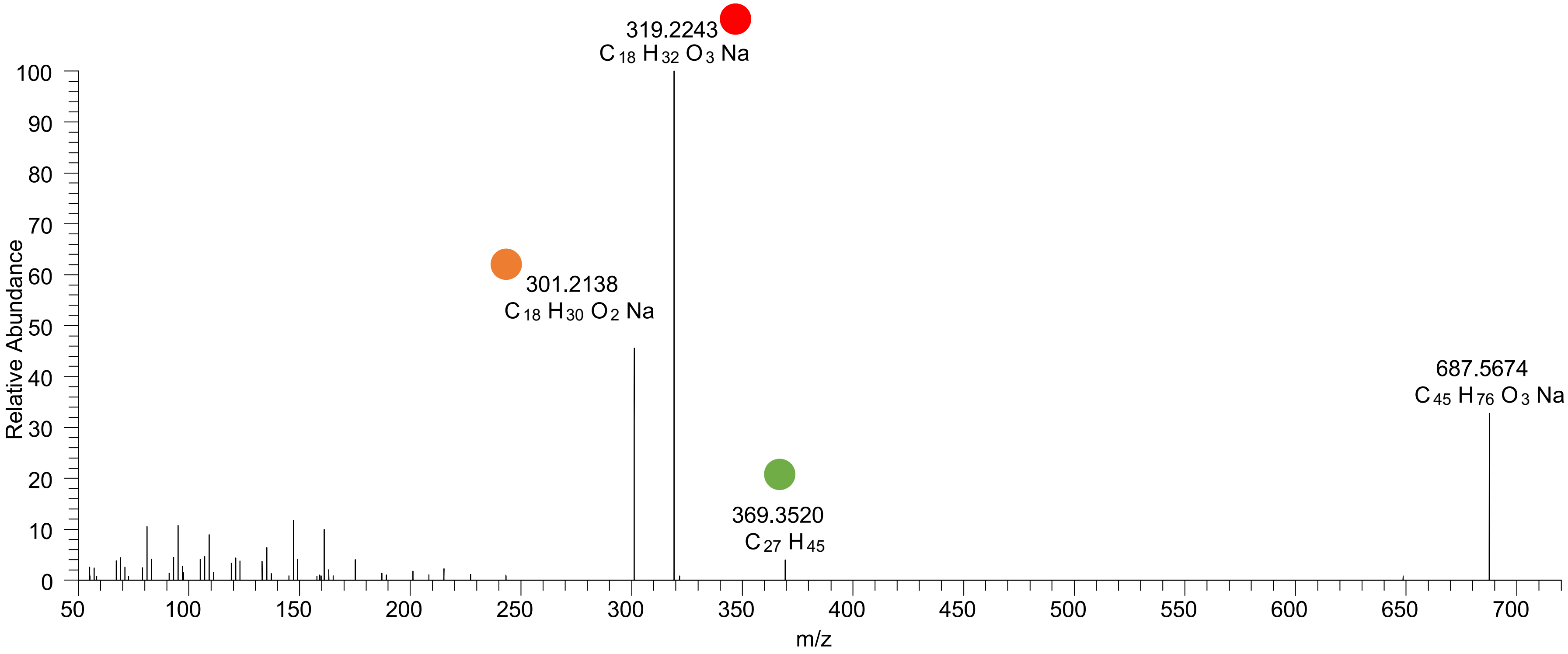

# CE(18:2<O>)
# RT 24.6, 25.2

[oxCE+Na]+ XIC 687.5687 NL: 3.03E5

Fragments containing oxFAs · Fragments not containing oxFAs · Fragments relative to water loss · Specific position fragments · Fragments relative to other oxLPPs

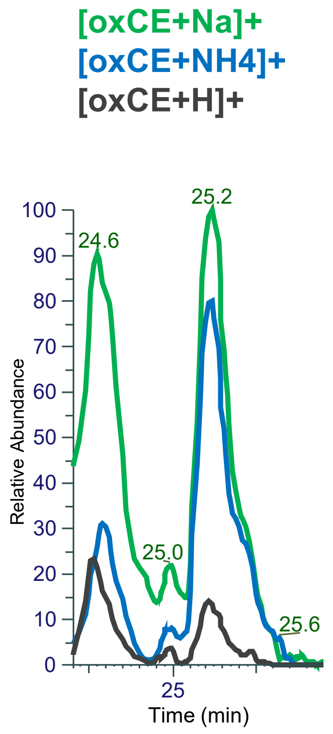


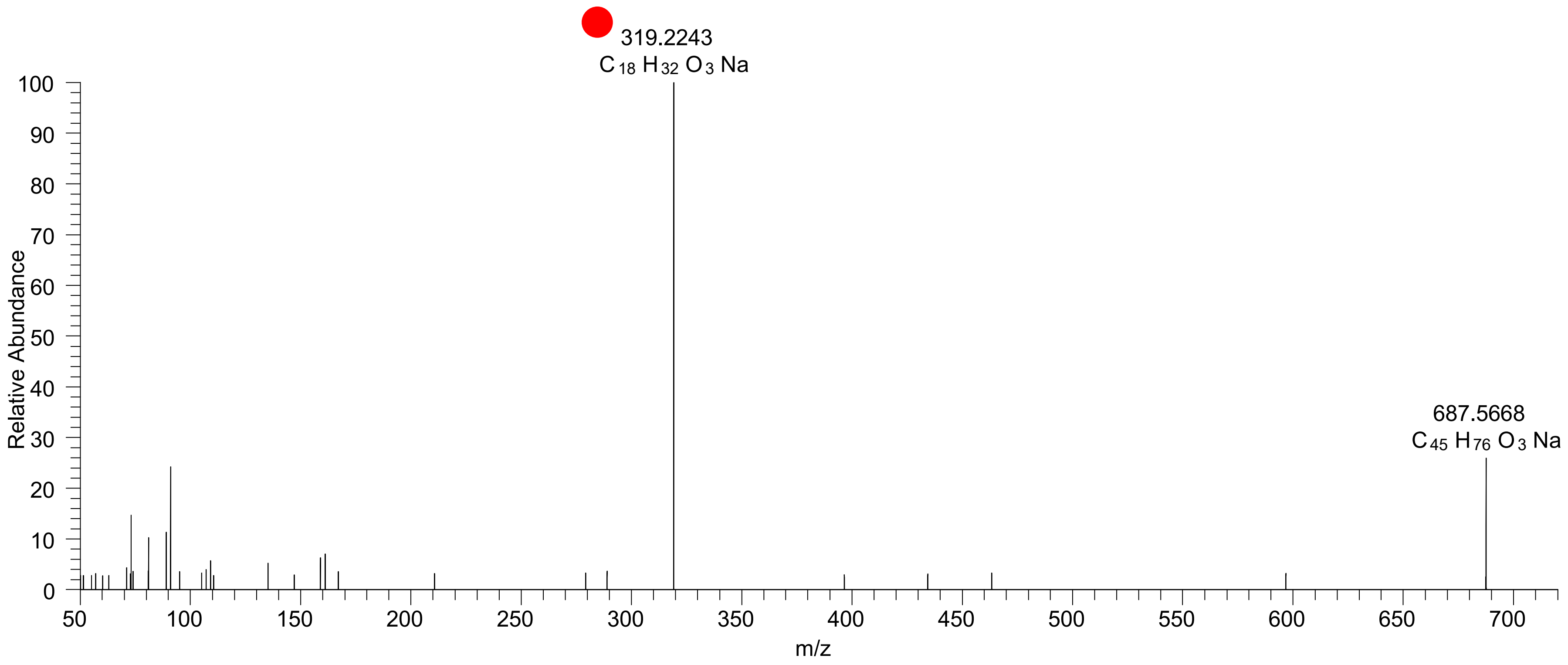

# CE(18:1<OH>)
# RT 24.2

**[oxCE+Na]+** **XIC 689.5843 NL: 5.26E5**

Fragments containing oxFAs · Fragments not containing oxFAs · Fragments relative to water loss · Specific position fragments · Fragments relative to other oxLPPs

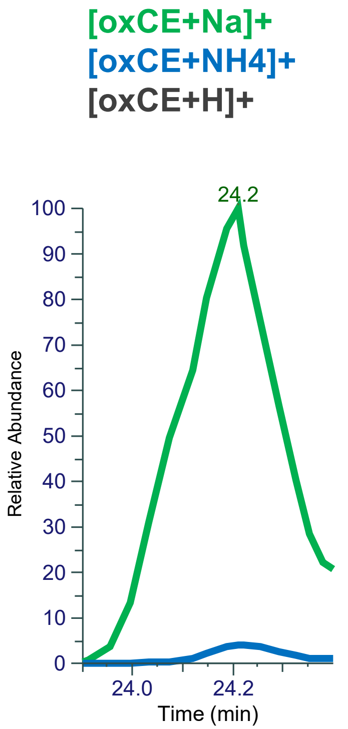


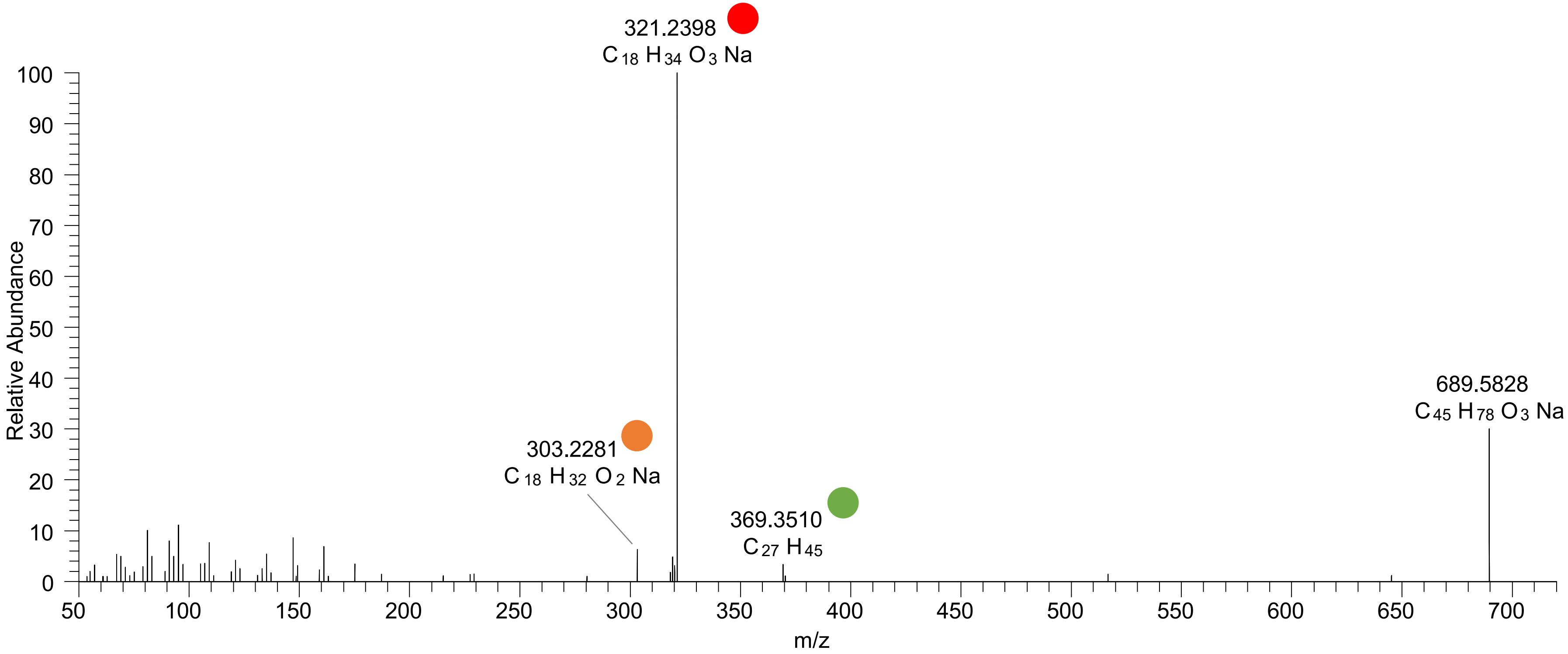

# CE(18:0<ep>)
# RT 25.9

**[oxCE+Na]+** **XIC 689.5843 NL: 4.79E4**

- Fragments containing oxFAs
- Fragments related to water loss
- Fragments not containing oxFAs
- Position-specific fragments
- Fragments related to other oxLPPs

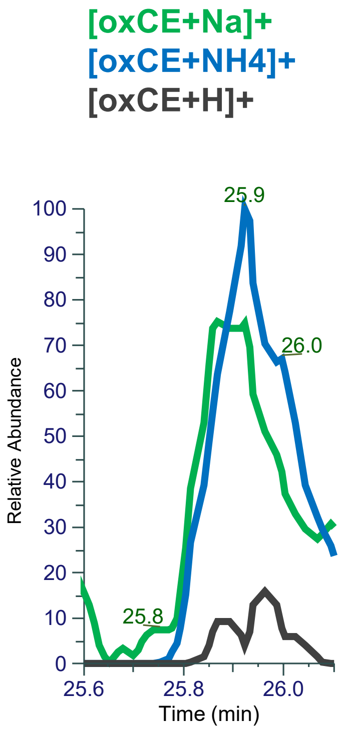


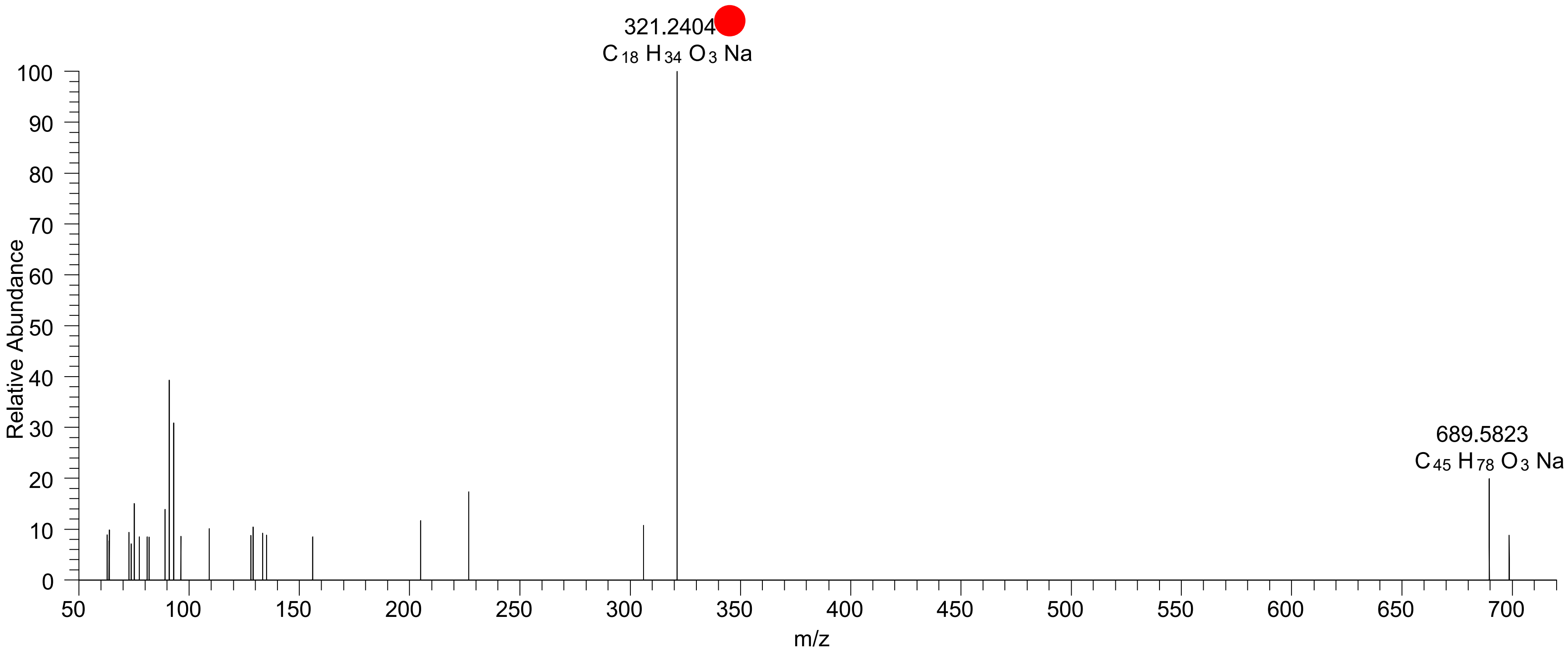

# CE(18:3<2OH>)
# RT 20.6

**[oxCE+Na]+** **XIC 701.5479 NL: 2.38E5**

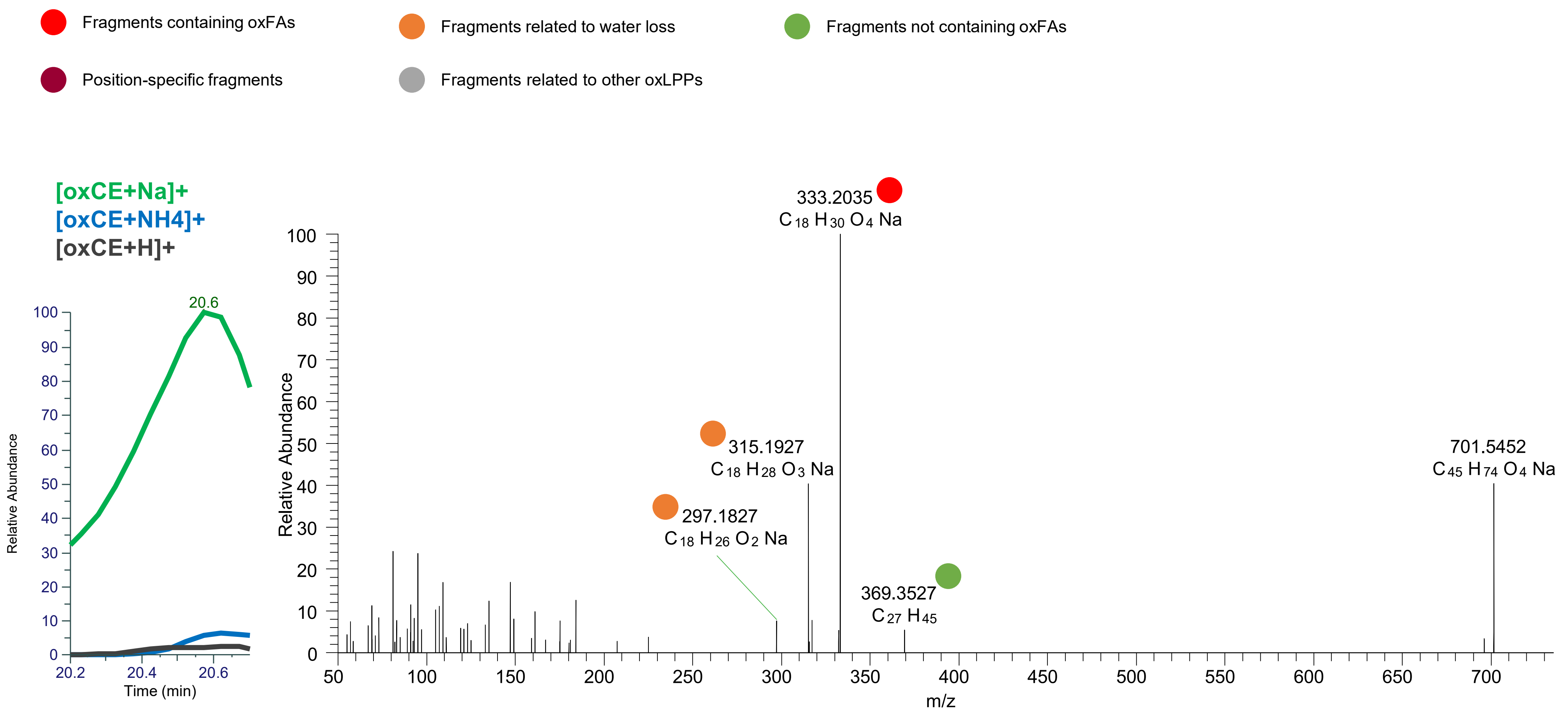

# CE(18:3<OH,O>)
# RT 21.3

**[oxCE+Na]+** **XIC 701.5479 NL: 4.10E5**

- Fragments containing oxFAs
- Fragments related to water loss
- Fragments not containing oxFAs
- Position-specific fragments
- Fragments related to other oxLPPs

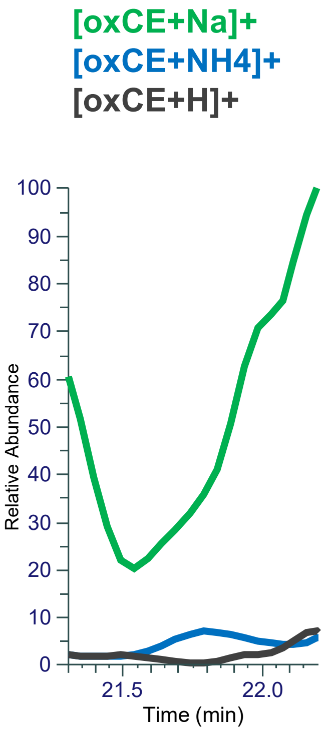


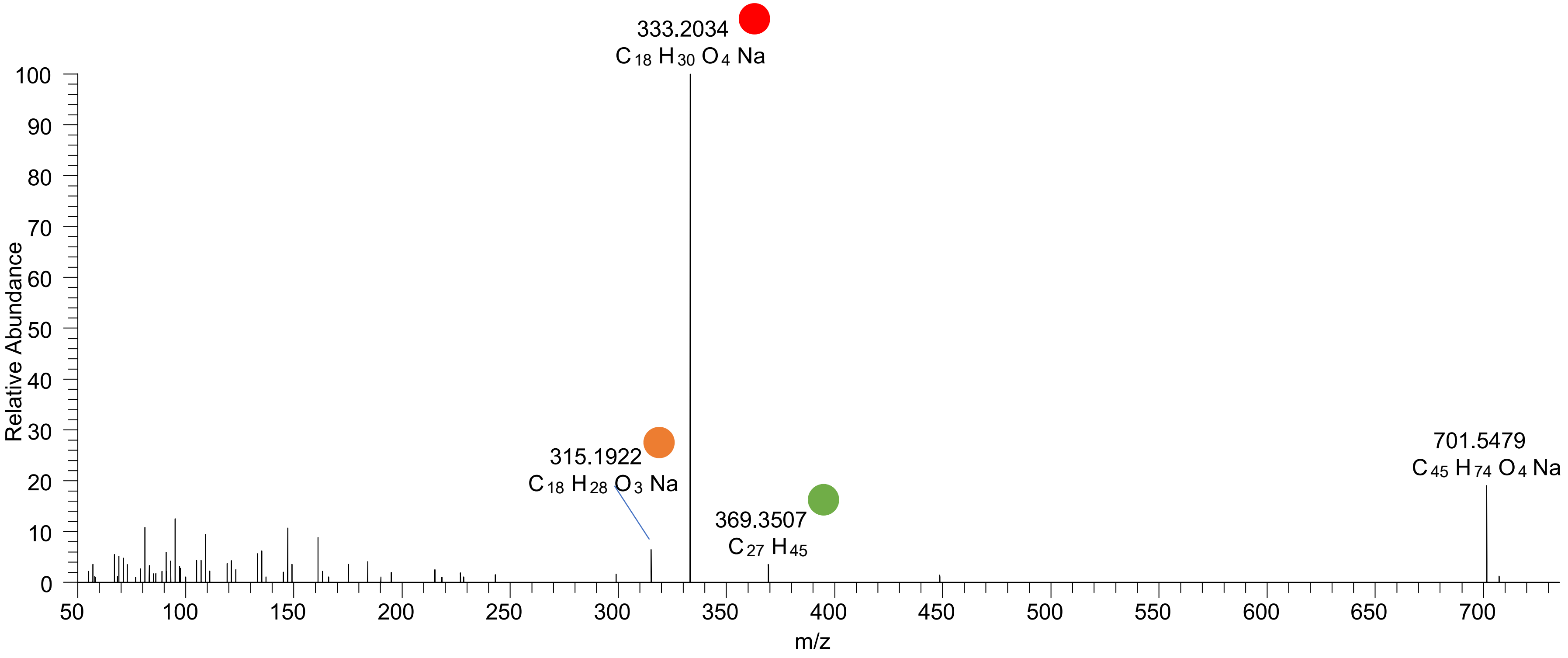

# CE(18:3<OH,O>)
# RT 22.6

**[oxCE+Na]+** **XIC 701.5479 NL: 8.42E5**

- Fragments containing oxFAs
- Fragments related to water loss
- Fragments not containing oxFAs
- Position-specific fragments
- Fragments related to other oxLPPs

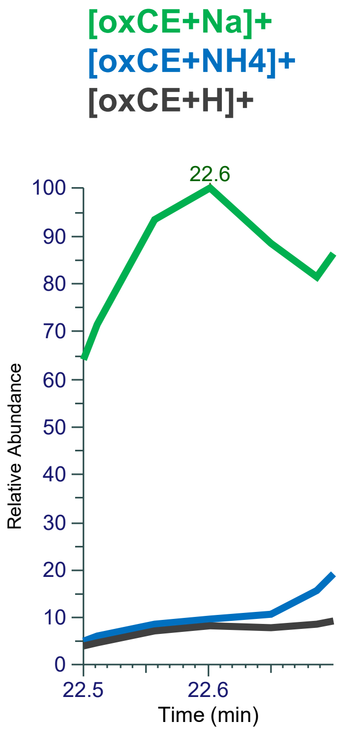


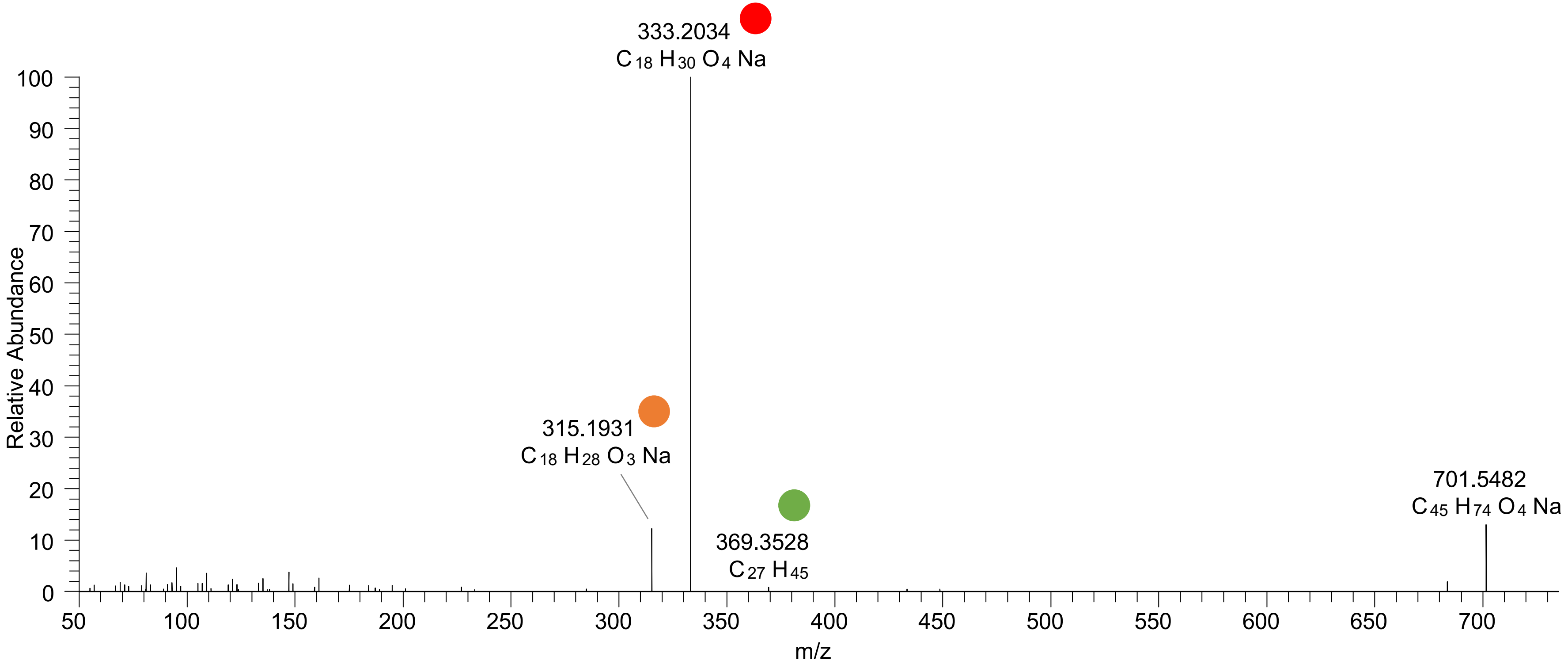

# CE(18:3<2O>)
# RT 22.8

**[oxCE+Na]+ XIC 701.5479 NL: 9.97E5**

- Fragments containing oxFAs
- Fragments related to water loss
- Fragments not containing oxFAs
- Position-specific fragments
- Fragments related to other oxLPPs

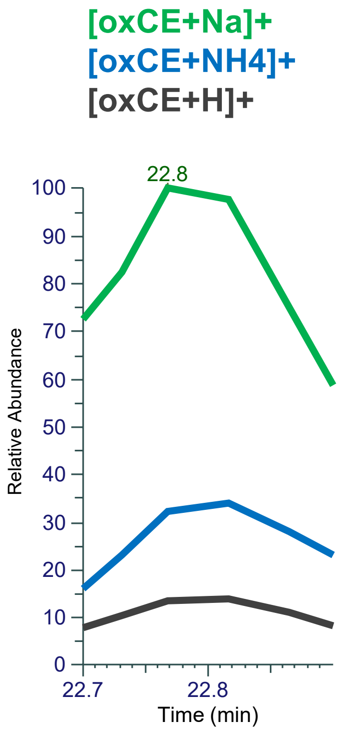


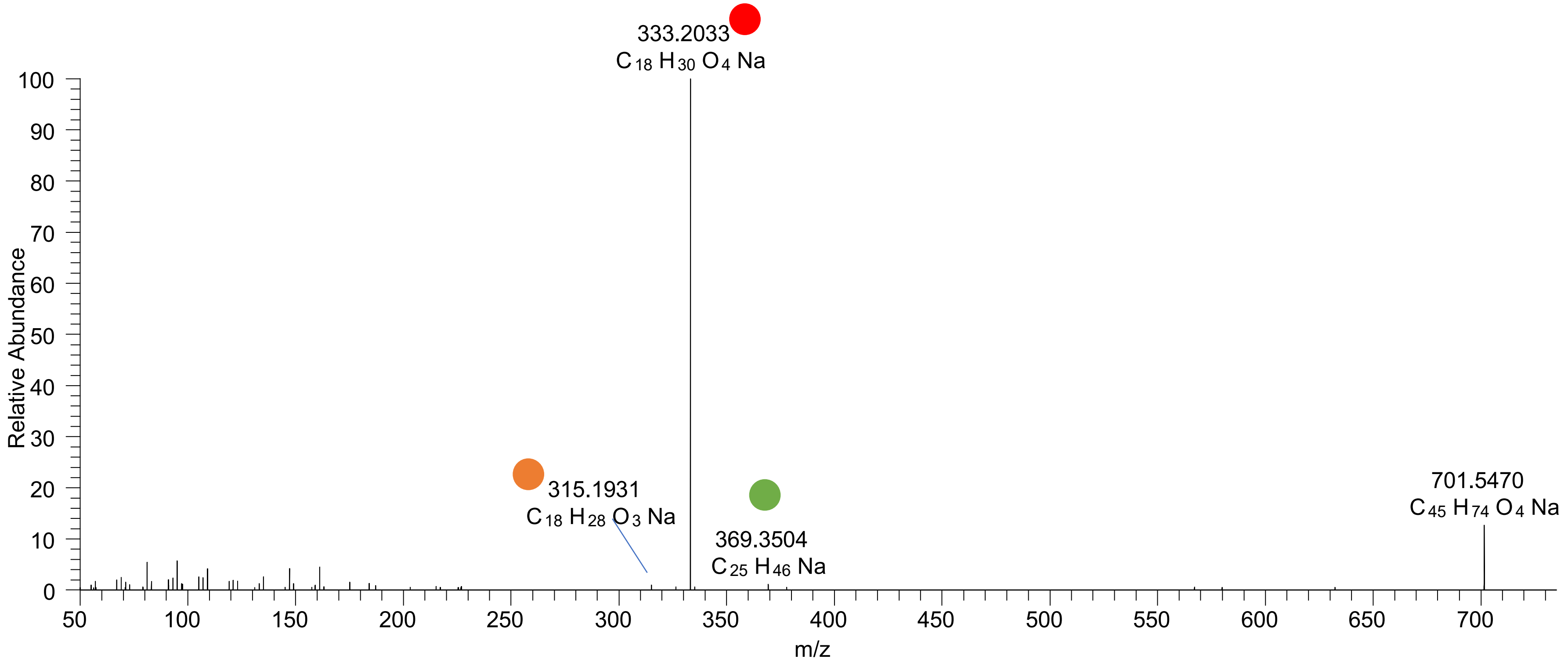

### CE(18:3<OOH{10}>)
### CE(18:3<OOH{13}>)
# RT 23.0-23.2

**[oxCE+Na]+** **XIC 701.5479 NL: 1.11E6**

Fragments containing oxFAs
Fragments related to water loss
Fragments not containing oxFAs
Position-specific fragments
Fragments related to other oxLPPs

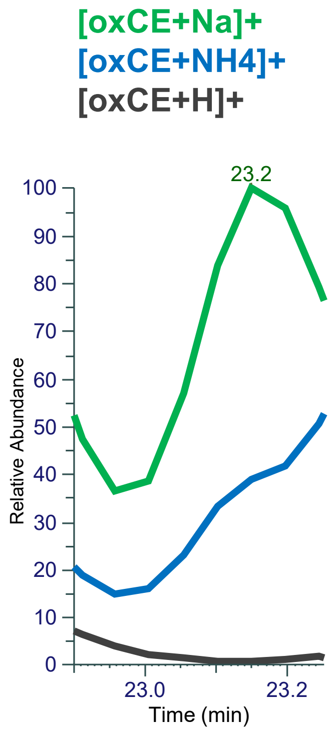


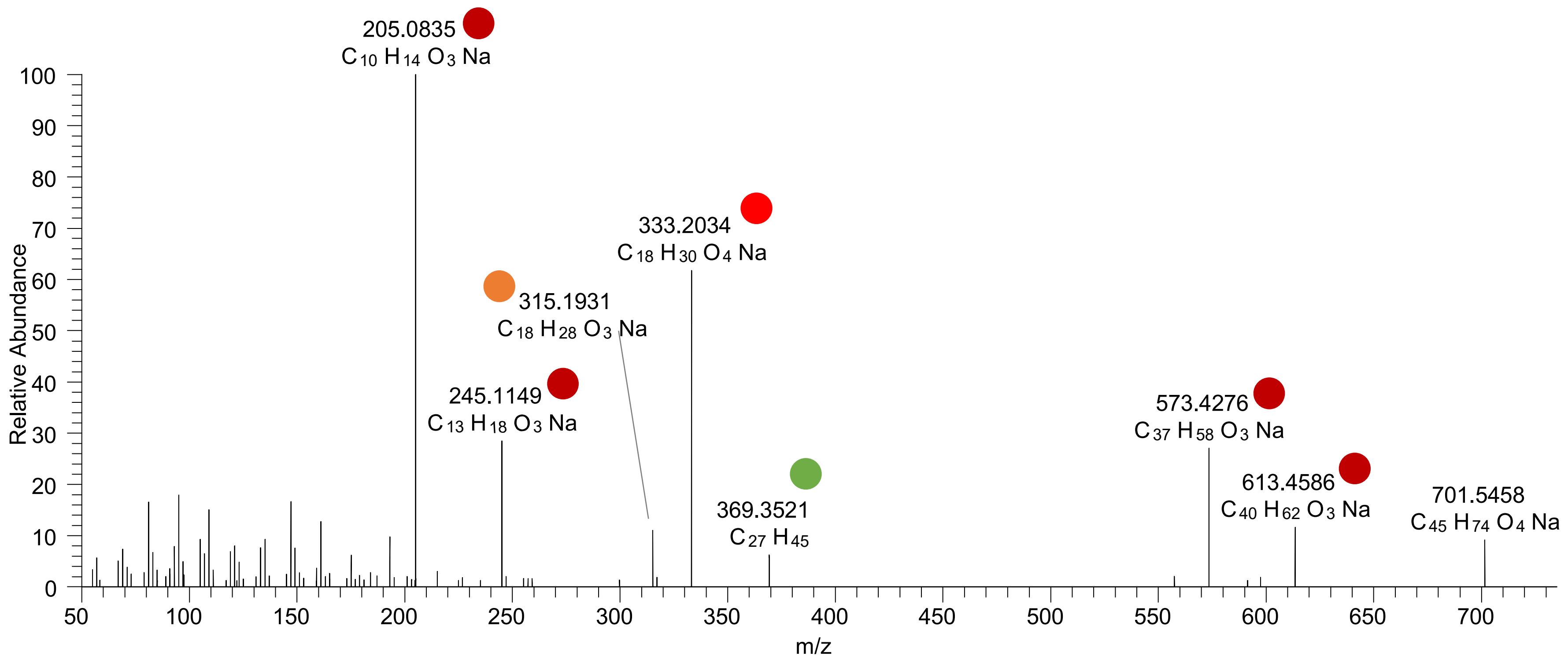

### CE(18:3<OOH{13}>)
### CE(18:3<OOH{16}>)
# RT 23.3

**[oxCE+Na]+** **XIC 701.5479 NL: 6.80E5**

- Fragments containing oxFAs
- Fragments related to water loss
- Fragments not containing oxFAs
- Position-specific fragments
- Fragments related to other oxLPPs

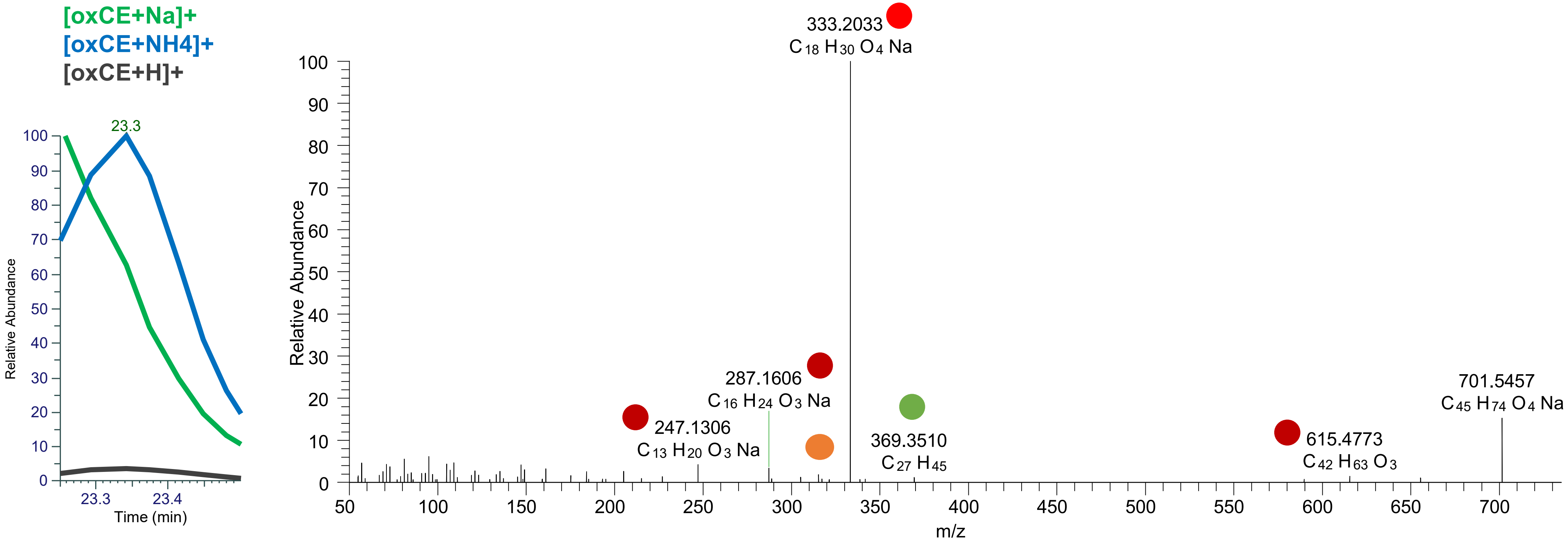

# CE(18:2<2OH>)
# RT 21.1

**[oxCE+Na]+** **XIC 703.5636 NL: 2.03E5**

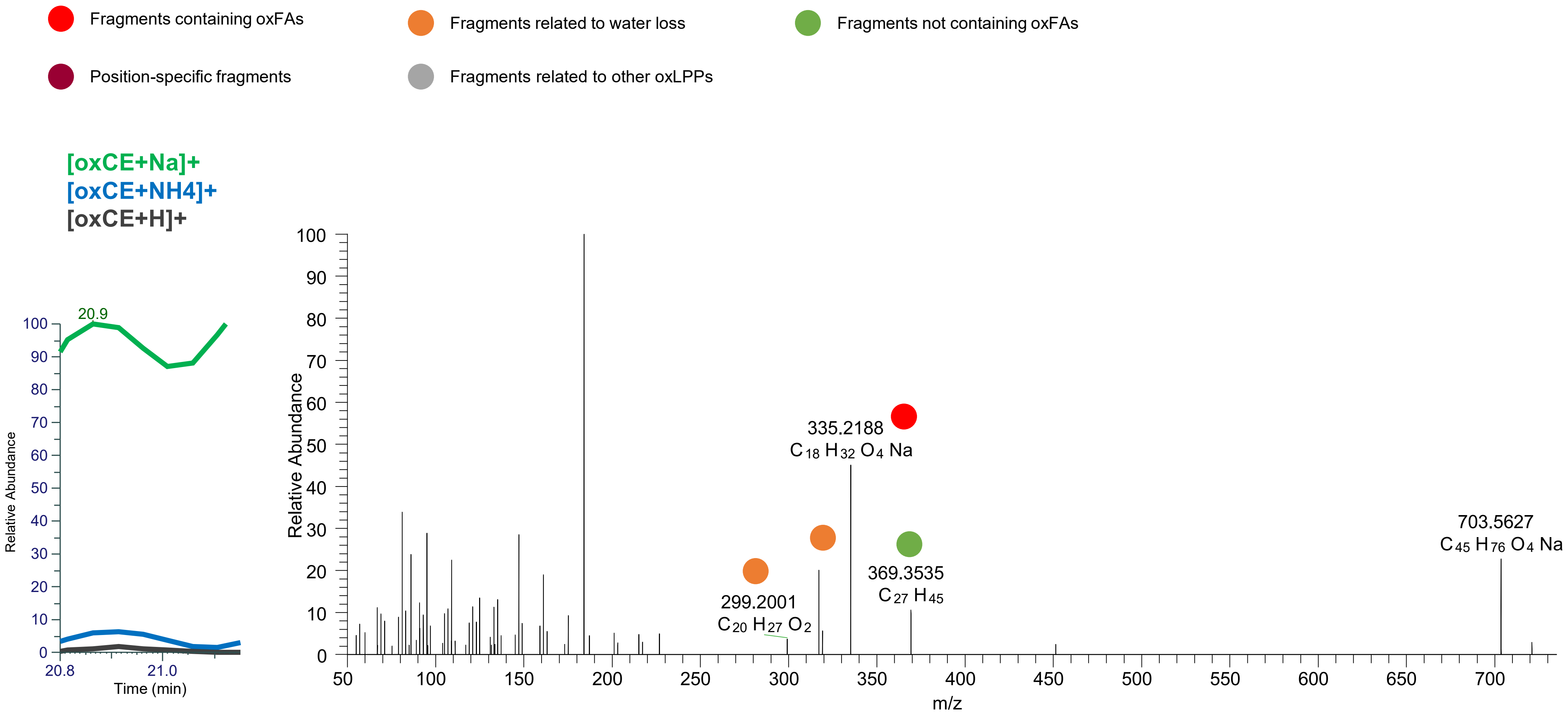

# CE(18:2<OH,O>)
# RT 21.6

**[oxCE+Na]+** **XIC 703.5636 NL: 2.02E5**

- Fragments containing oxFAs
- Fragments related to water loss
- Fragments not containing oxFAs
- Position-specific fragments
- Fragments related to other oxLPPs

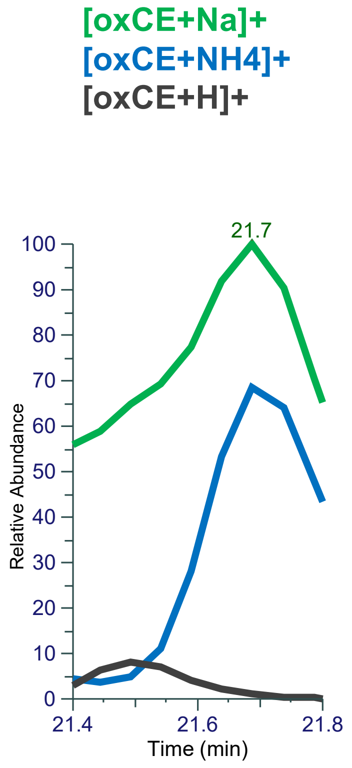


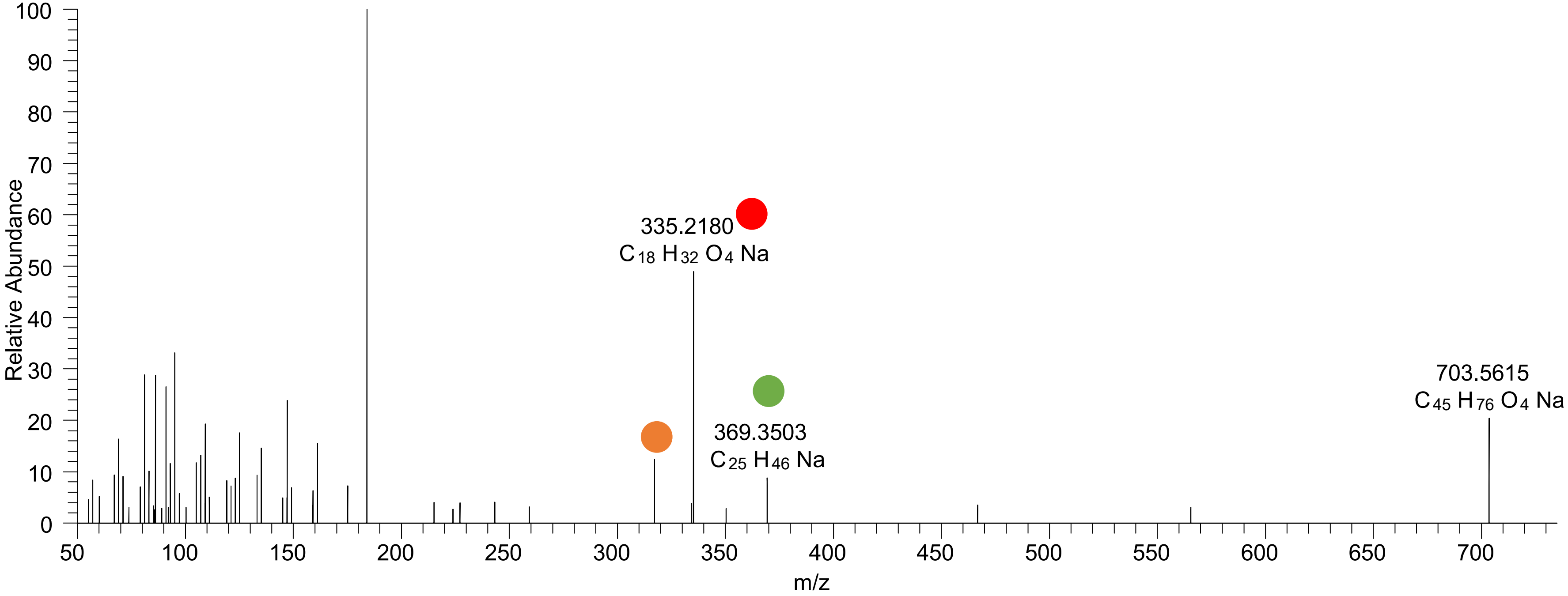

# CE(18:2<OH{9},O{10}>)
# RT 22.3

**[oxCE+Na]+** **XIC 703.5636 NL: 2.28E6**

Fragments containing oxFAs
Fragments related to water loss
Fragments not containing oxFAs
Position-specific fragments
Fragments related to other oxLPPs

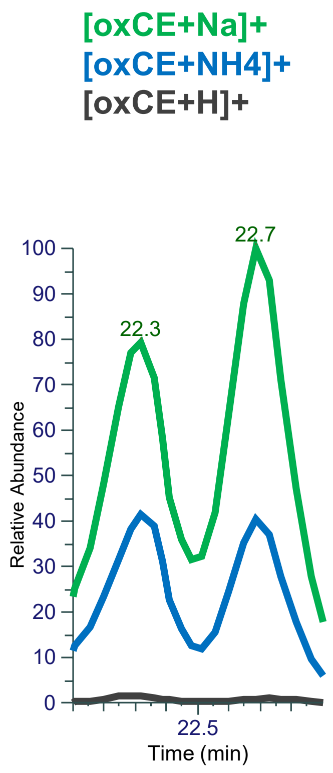


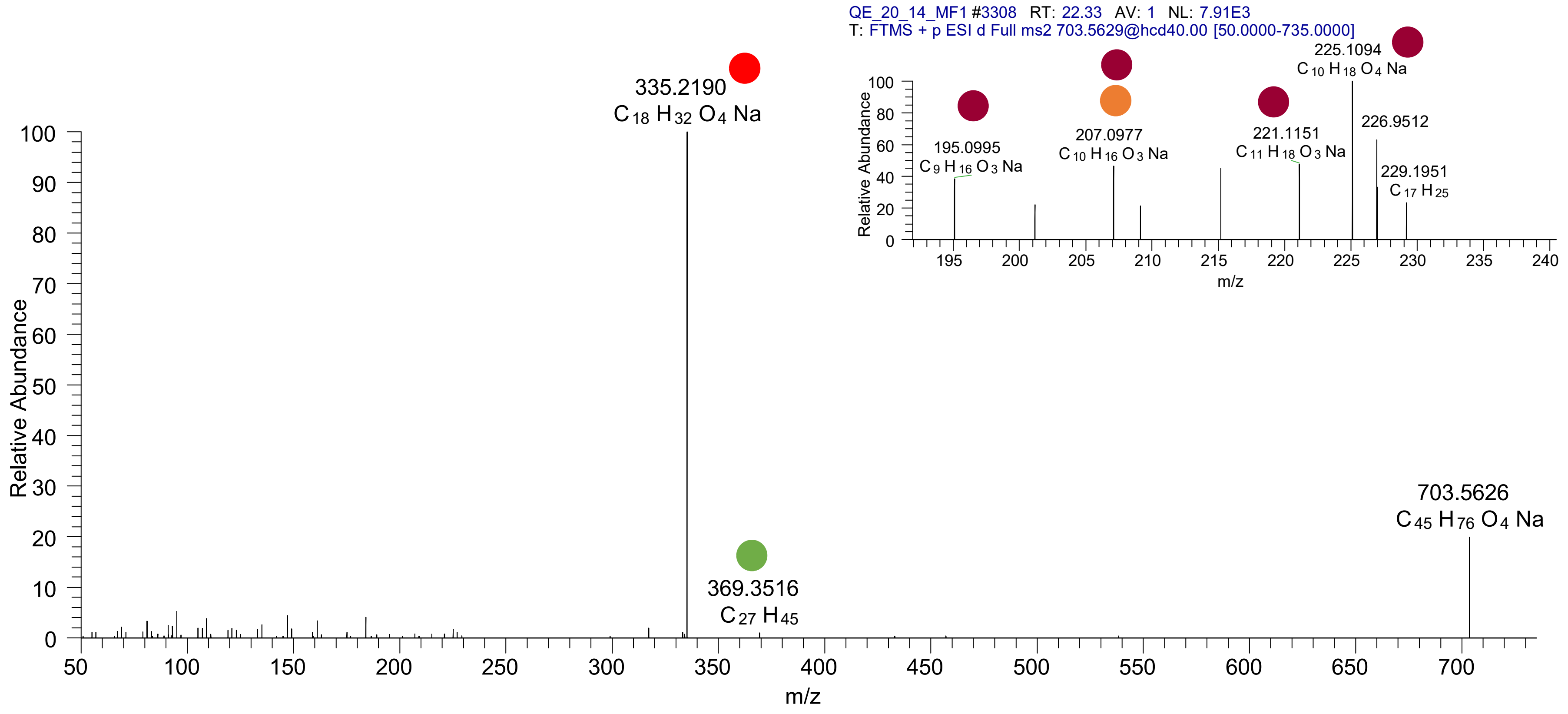

CE(18:2<OOH{10}>)
CE(18:2<OOH{11}>)
CE(18:2<OOH{13}>)
CE(18:2<OOH{15}>)
RT 23.3

**[oxCE+Na]+** **XIC 703.5636 NL: 3.61E6**

- Fragments containing oxFAs
- Fragments related to water loss
- Fragments not containing oxFAs
- Position-specific fragments
- Fragments related to other oxLPPs

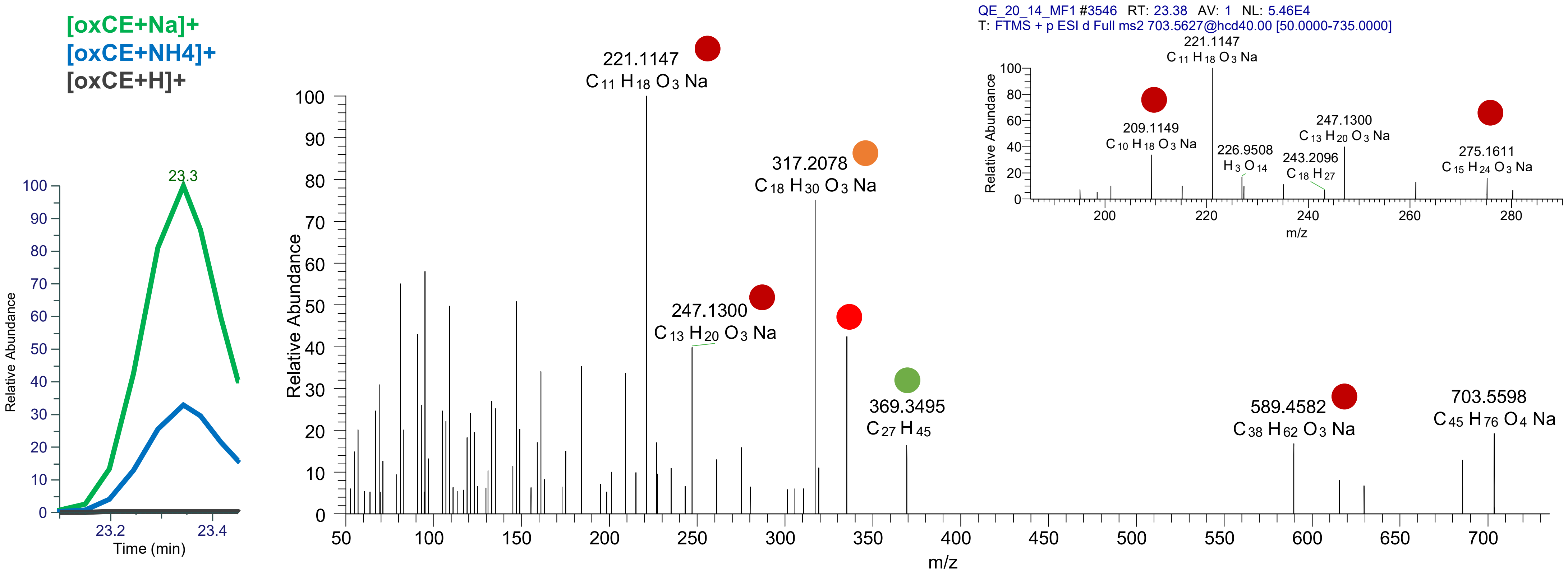

### CE(18:2<OOH{9}>)
### CE(18:2<OOH{13}>)
# RT 23.6

**[oxCE+Na]+** **XIC 703.5636 NL: 1.35E7**

- Fragments containing oxFAs
- Fragments related to water loss
- Fragments not containing oxFAs
- Position-specific fragments
- Fragments related to other oxLPPs

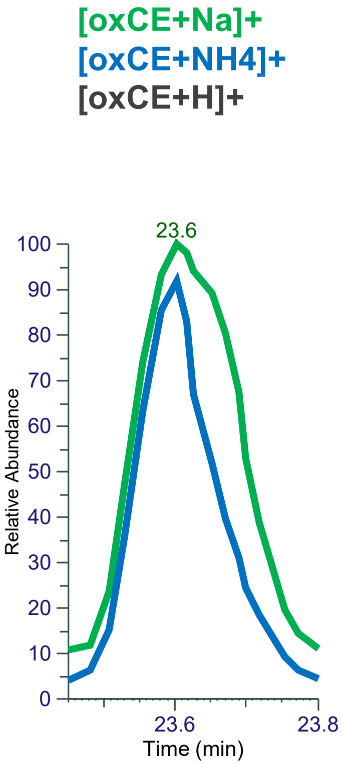


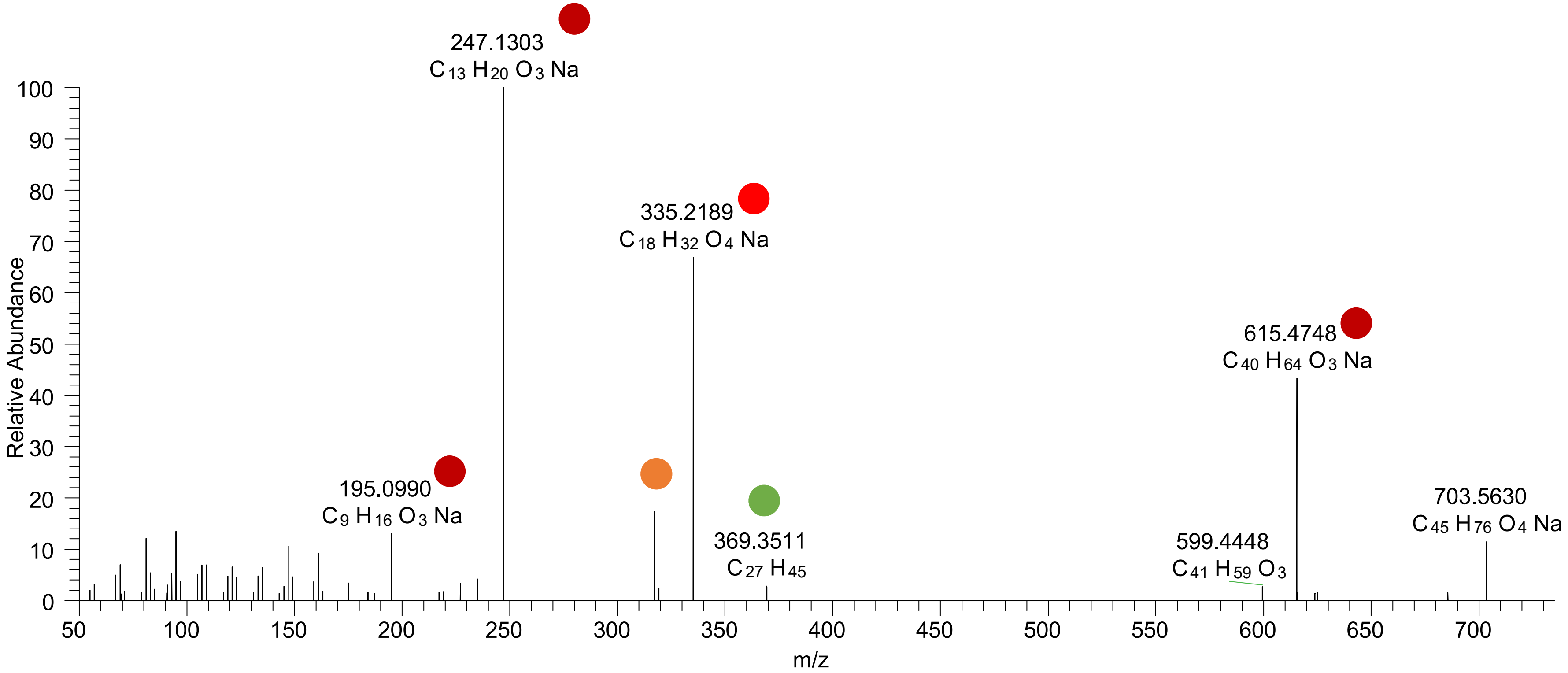

# CE(18:0<ep,O>)
# RT 23.2

**[oxCE+Na]+** **XIC 705.5792 NL: 7.39E3**

- Fragments containing oxFAs
- Fragments related to water loss
- Fragments not containing oxFAs
- Position-specific fragments
- Fragments related to other oxLPPs

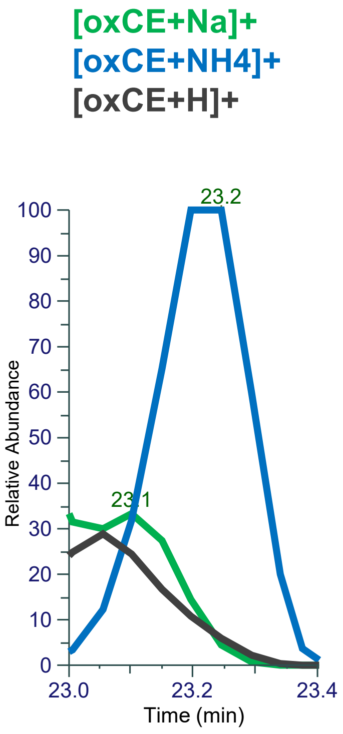


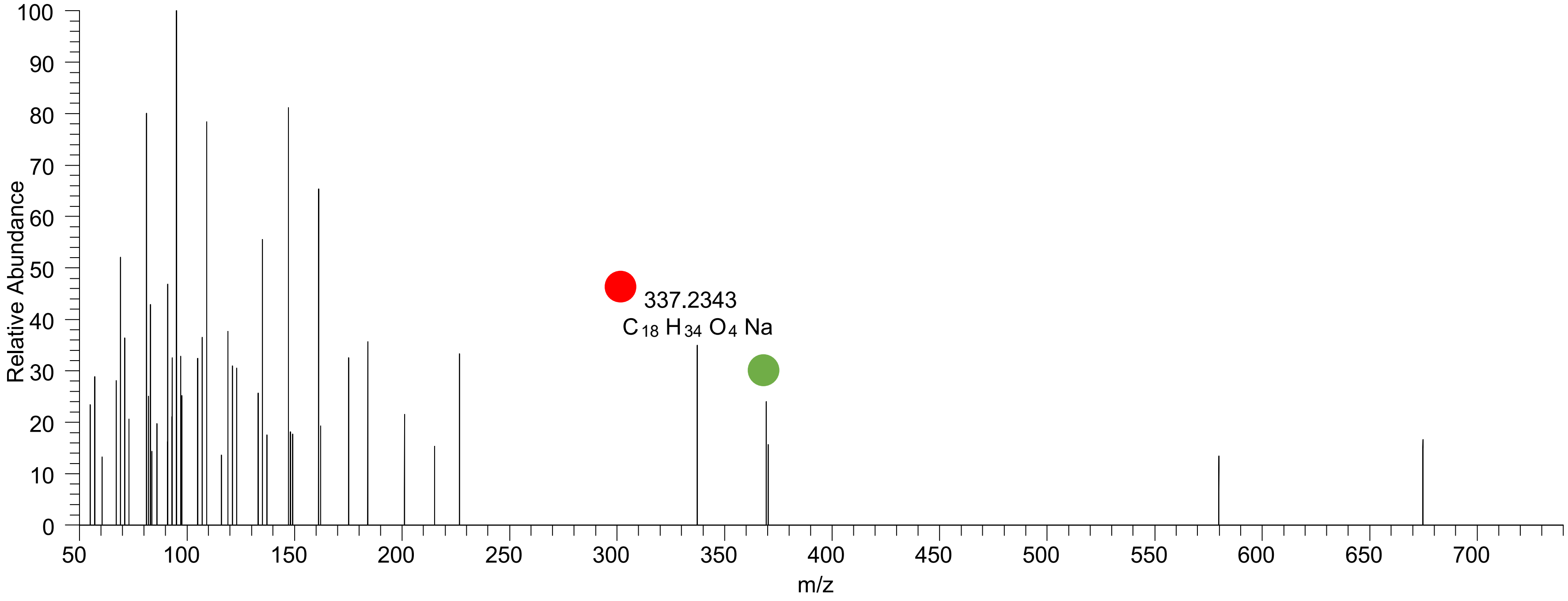

### CE(18:1<OOH{11}>)
### CE(18:1<OOH{13}>)
# RT 24.0

**[oxCE+Na]+** **XIC 705.5792 NL: 4.64E5**

- Fragments containing oxFAs
- Fragments related to water loss
- Fragments not containing oxFAs
- Position-specific fragments
- Fragments related to other oxLPPs

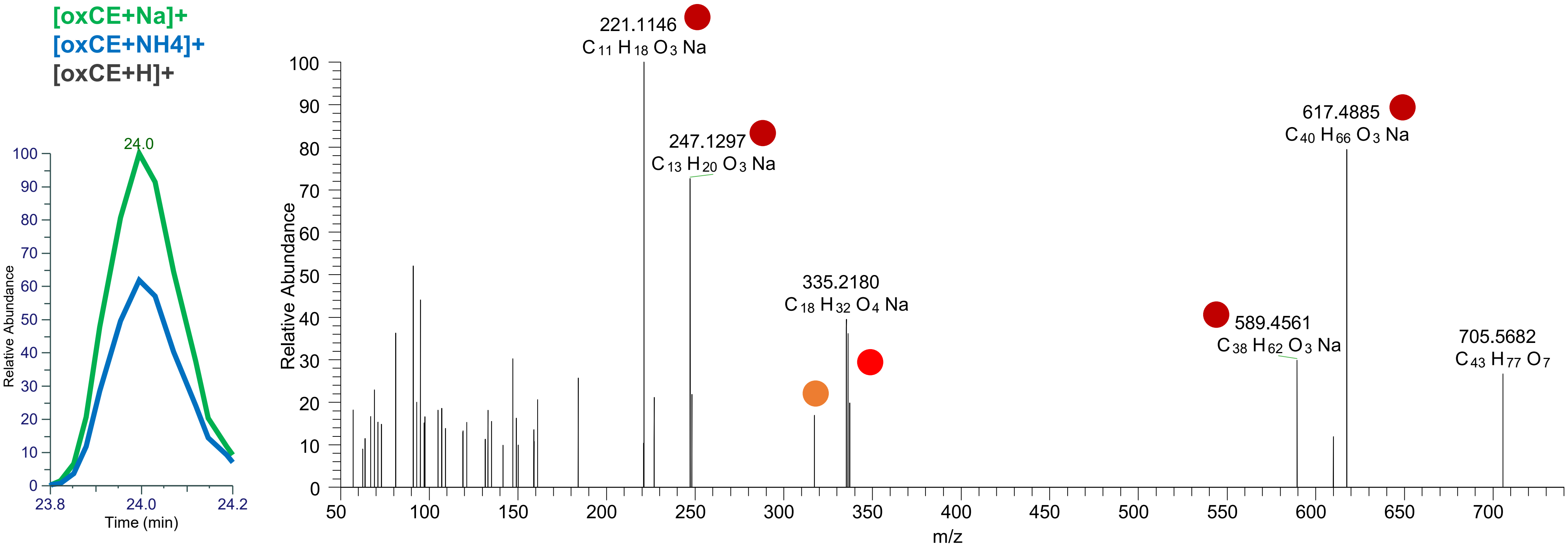

### CE(18:1<OOH{10}>)
### CE(18:1<OOH{12}>)
# RT 24.1

**[oxCE+Na]+** **XIC 705.5792 NL: 4.64E5**

- Fragments containing oxFAs
- Fragments related to water loss
- Fragments not containing oxFAs
- Position-specific fragments
- Fragments related to other oxLPPs

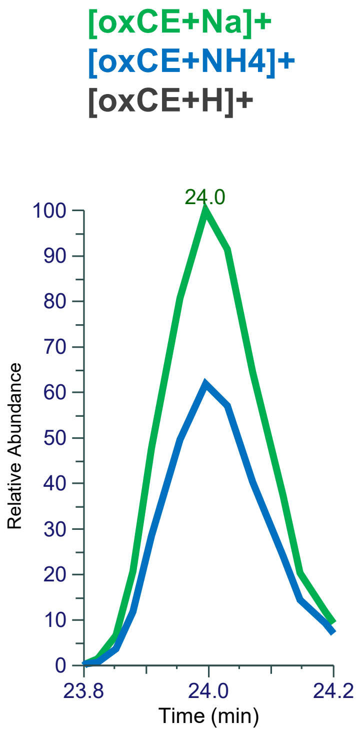


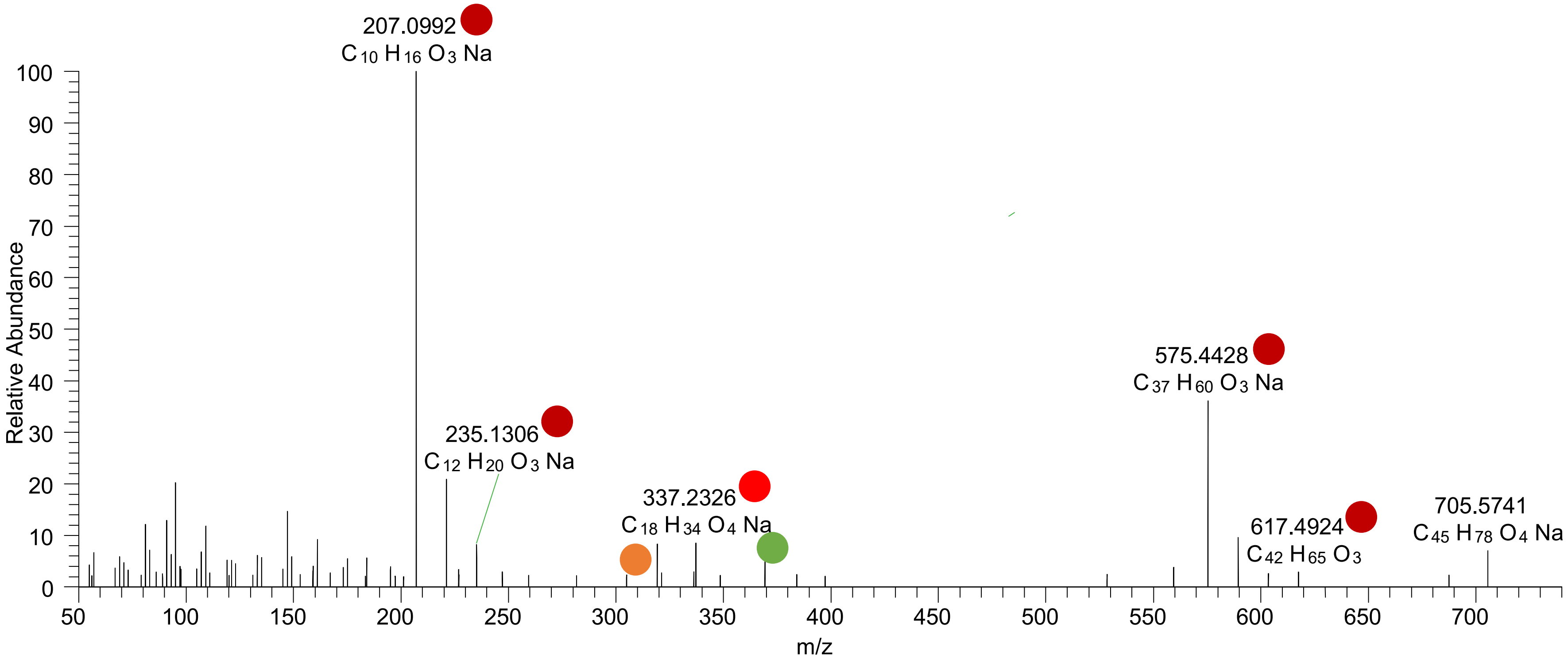

# CE(20:5<OH>)
# RT 23.1

**[oxCE+Na]+** **XIC 709.5530 NL: 1.44E6**

- Fragments containing oxFAs
- Fragments related to water loss
- Fragments not containing oxFAs
- Position-specific fragments
- Fragments related to other oxLPPs

**[oxCE+Na]+**
**[oxCE+NH4]+**
**[oxCE+H]+**

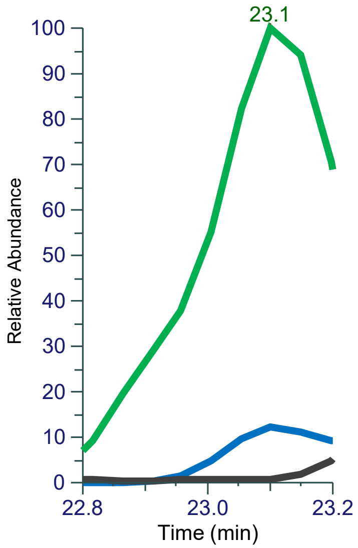


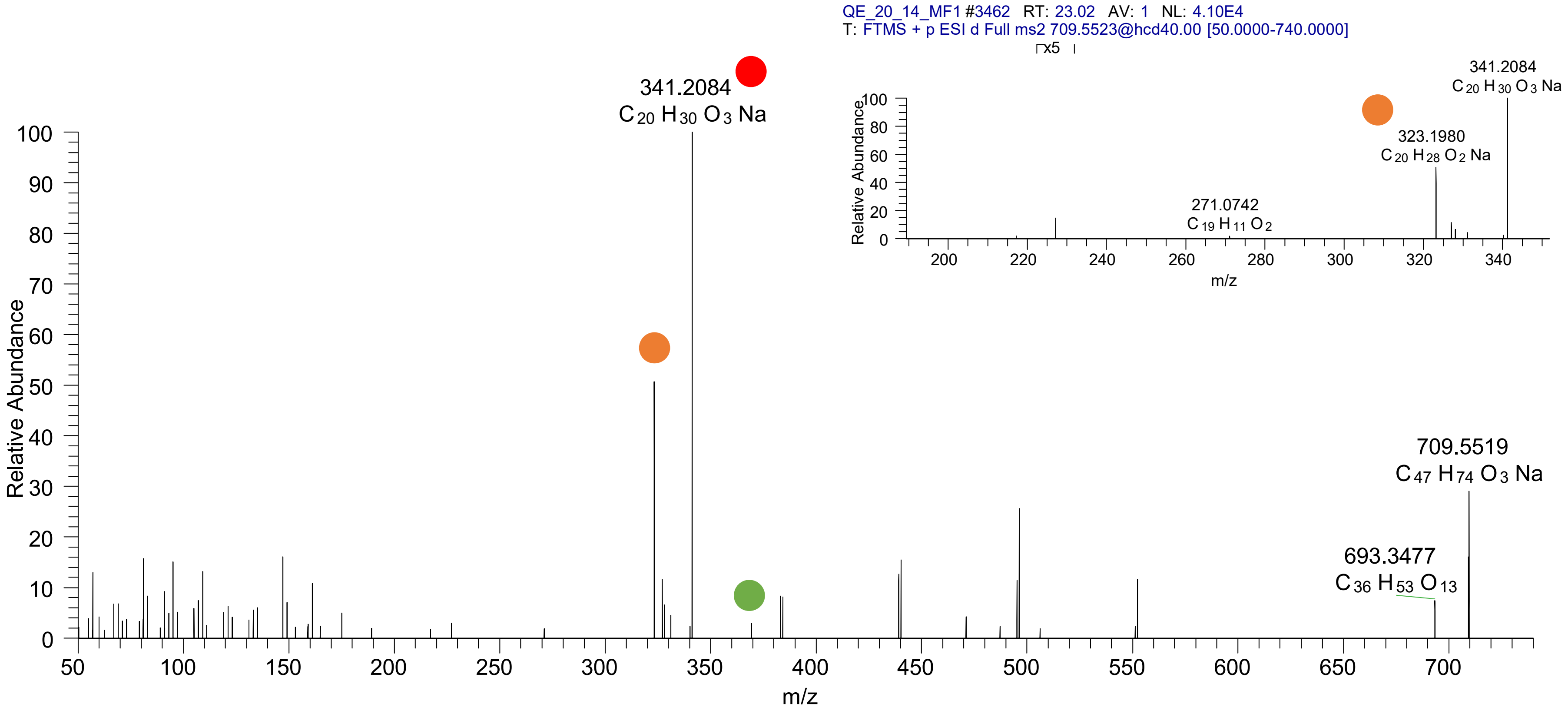

# CE(20:4<OH{9}>)
# RT 23.6

**[oxCE+Na]+** **XIC 711.5687 NL: 2.94E6**

- Fragments containing oxFAs
- Fragments related to water loss
- Fragments not containing oxFAs
- Position-specific fragments
- Fragments related to other oxLPPs

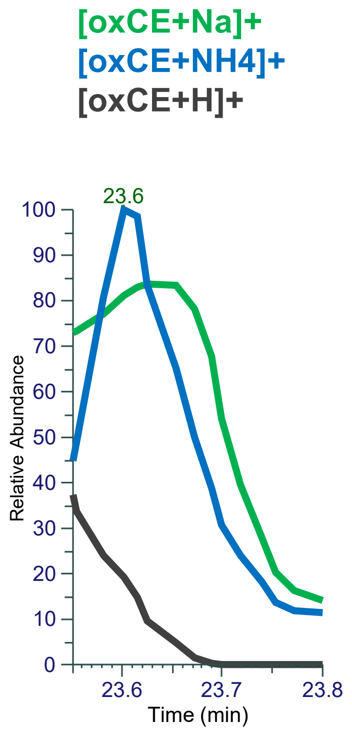


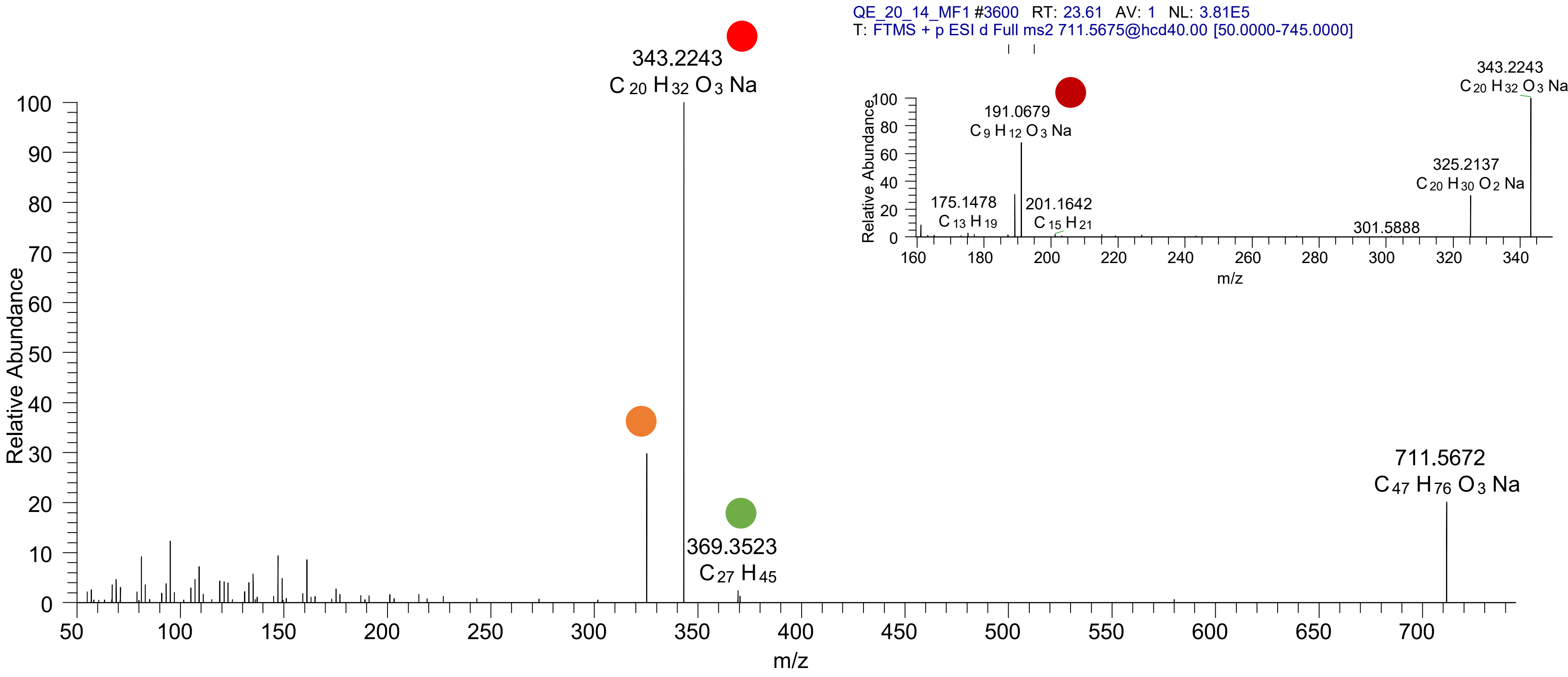

### CE(20:3<oxo>)
# RT 24.6

**[oxCE+Na]+** **XIC 711.5687 NL: 9.11E4**

- Fragments containing oxFAs
- Fragments related to water loss
- Fragments not containing oxFAs
- Position-specific fragments
- Fragments related to other oxLPPs

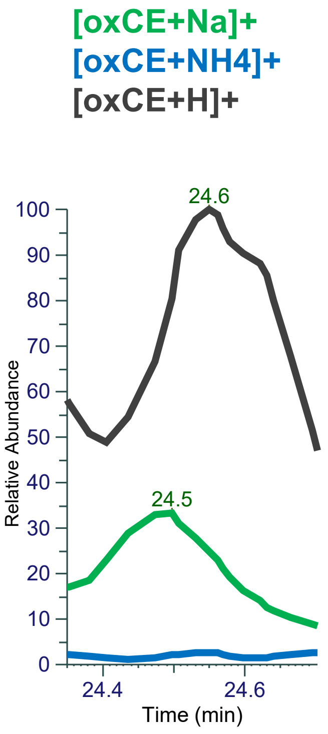


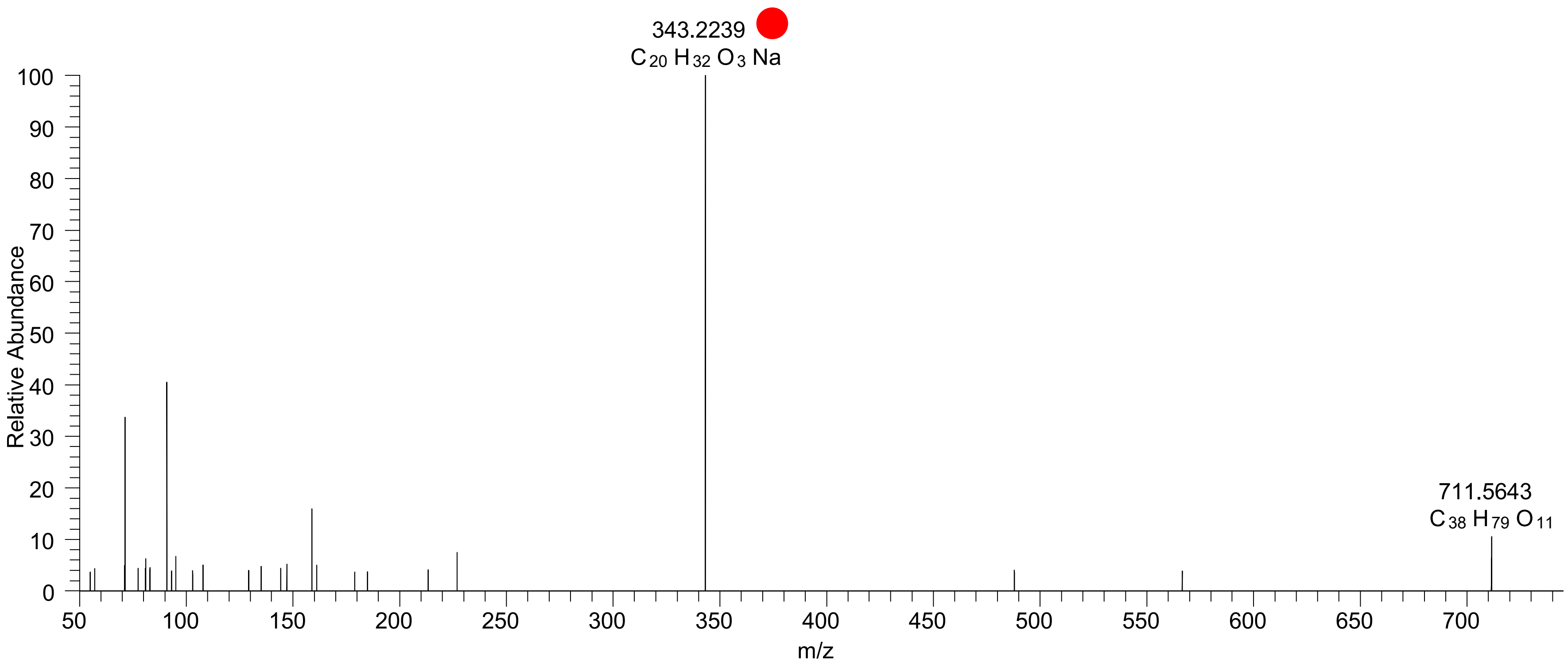

# CE(20:3<OH>)
# RT 24.0

[oxCE+Na]+ XIC 713.5849 NL: 1.67E6

Fragments containing oxFAs
Fragments related to water loss
Fragments not containing oxFAs
Position-specific fragments
Fragments related to other oxLPPs

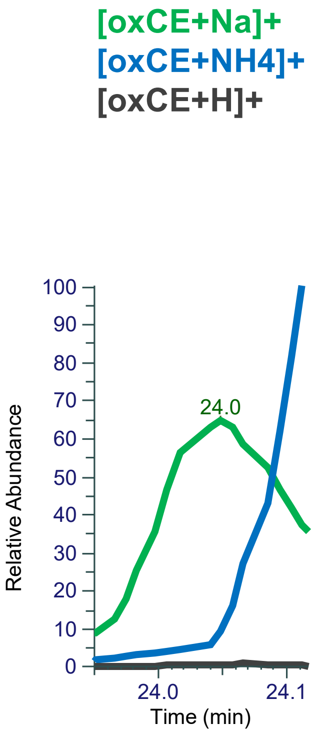


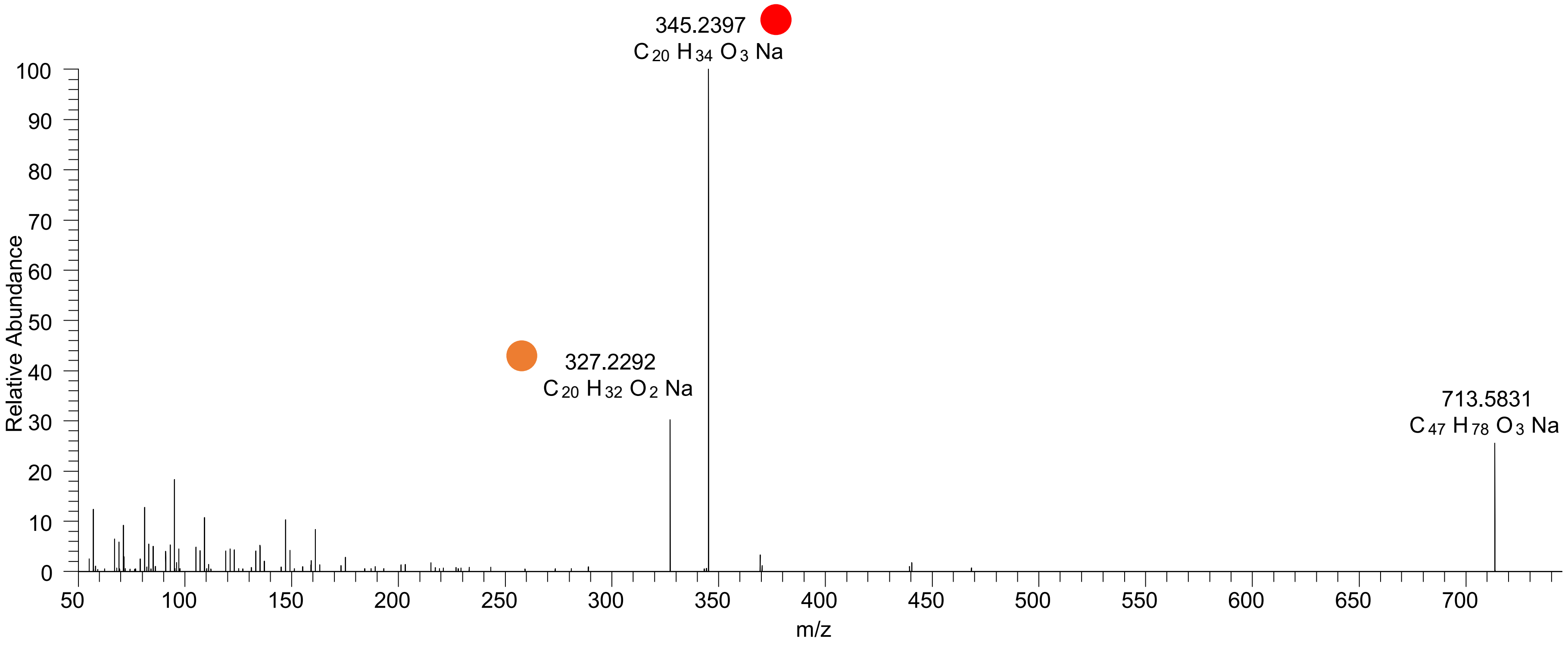

# CE(18:3<2OH,O>)
# RT 20.2

**[oxCE+Na]+** **XIC 717.5428 NL: 1.36E5**

- Fragments containing oxFAs
- Fragments related to water loss
- Fragments not containing oxFAs
- Position-specific fragments
- Fragments related to other oxLPPs

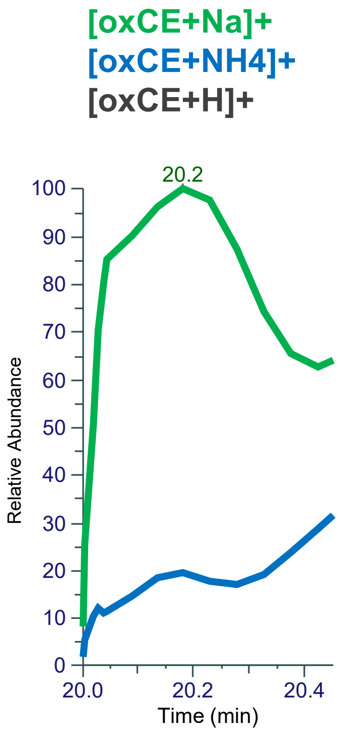


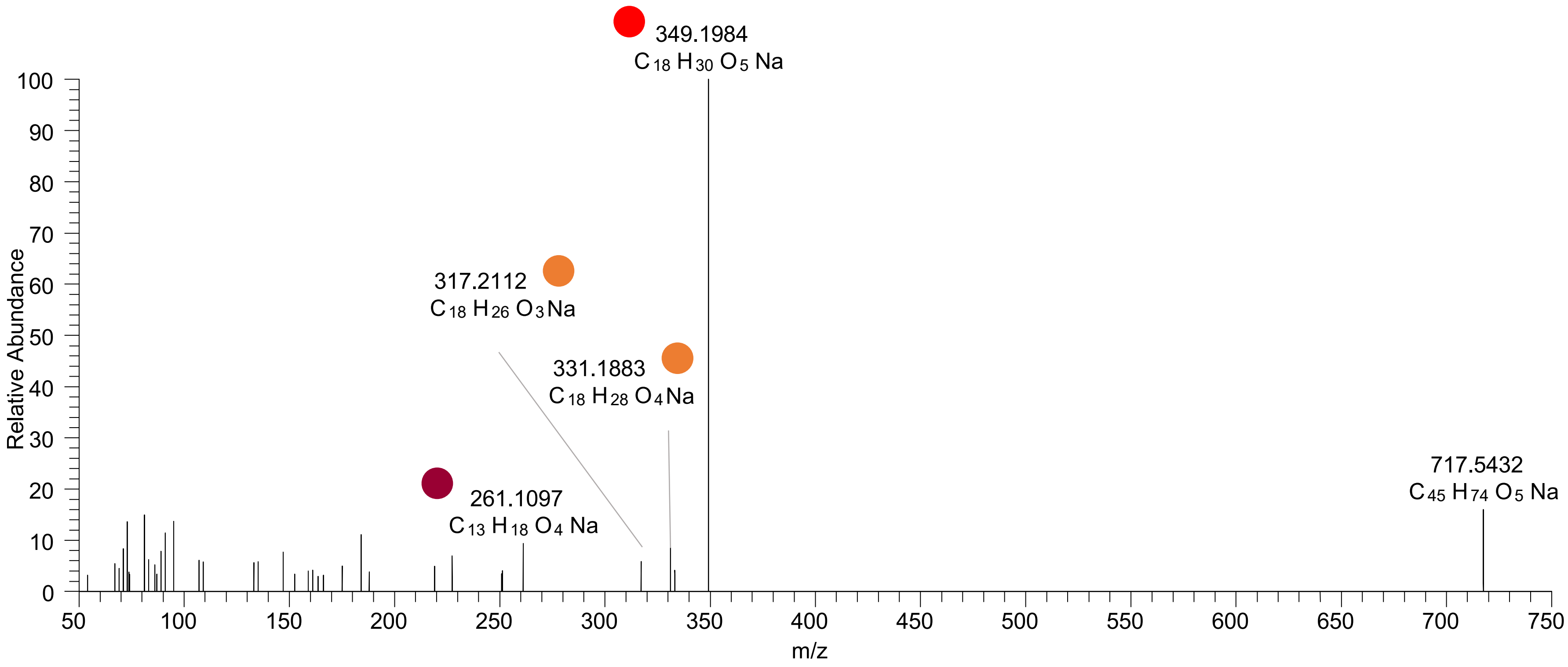

# CE(18:3<3O>)
# RT 20.7

**[oxCE+Na]+** **XIC 717.5428 NL: 1.03E5**

Fragments containing oxFAs

Fragments related to water loss

Fragments not containing oxFAs

Position-specific fragments

Fragments related to other oxLPPs

[oxCE+Na]+
[oxCE+NH4]+
[oxCE+H]+

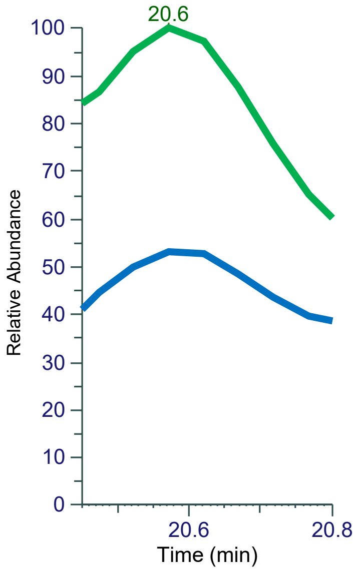


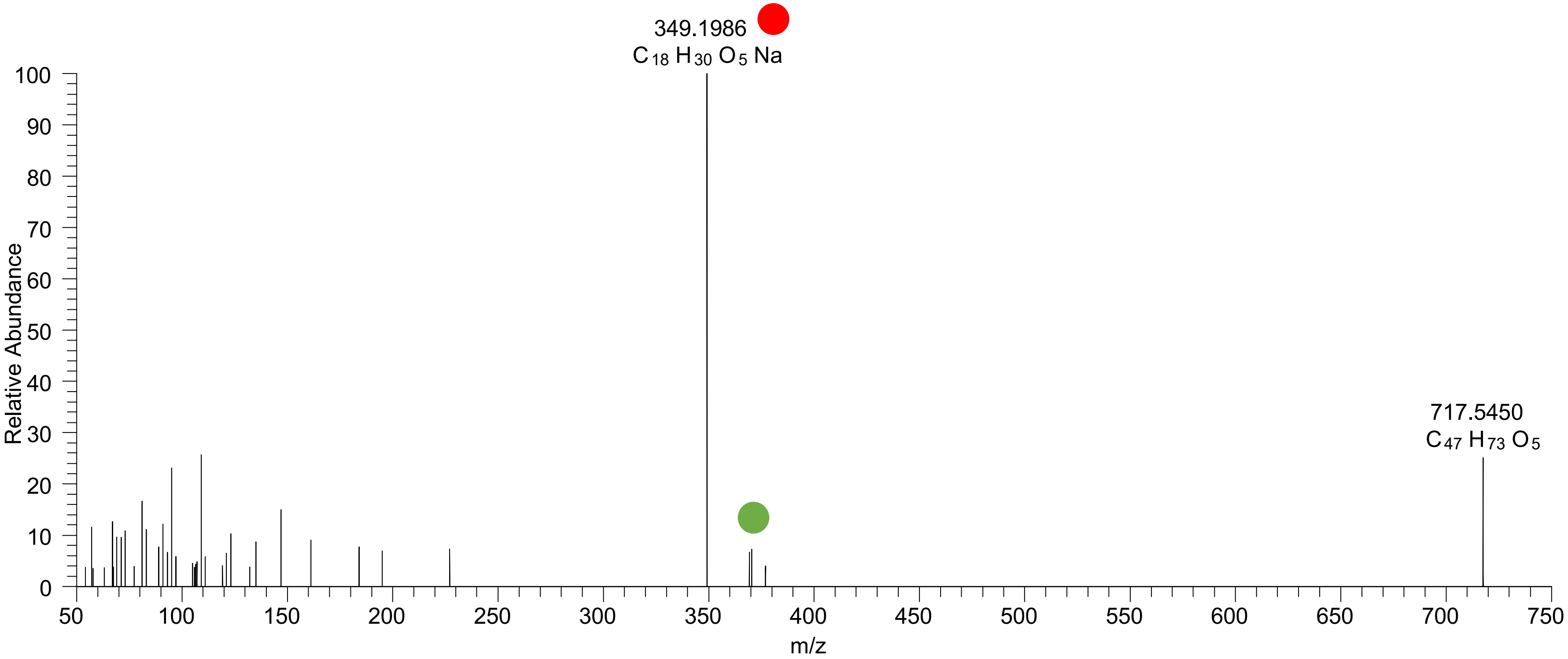

# CE(18:3<OH,2O>)
# RT 21.9

**[oxCE+Na]+** **XIC 717.5428 NL: 6.14E4**

Fragments containing oxFAs
Fragments related to water loss
Fragments not containing oxFAs
Position-specific fragments
Fragments related to other oxLPPs

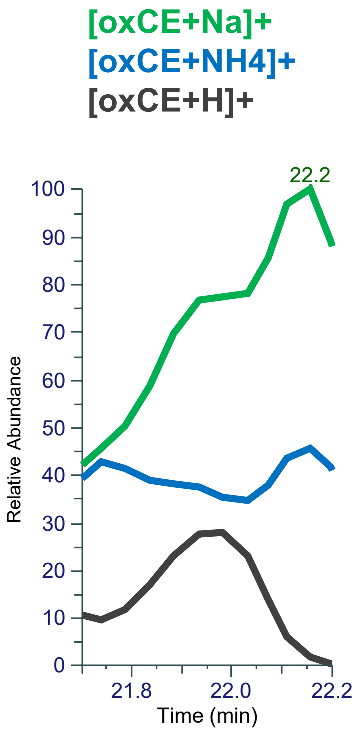


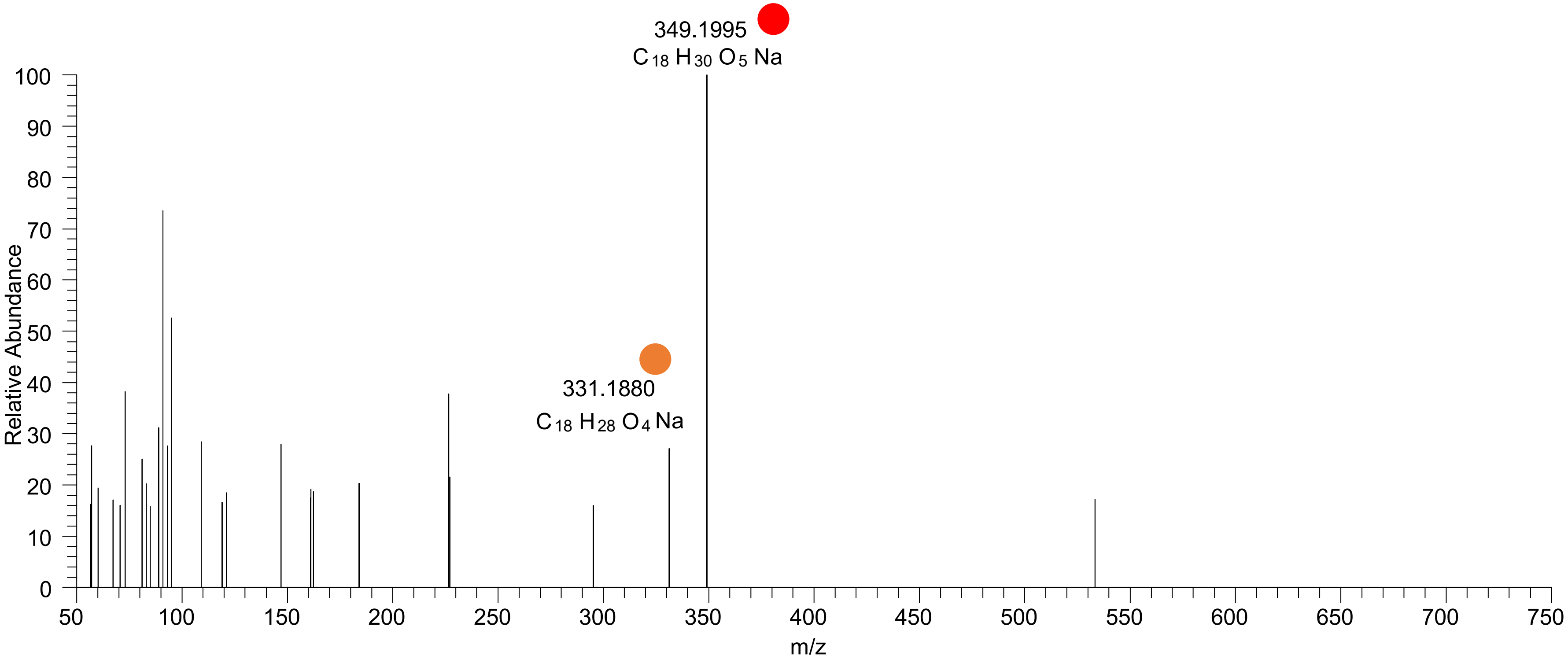

# CE(18:3<3O>)
# RT 23.3

**[oxCE+Na]+** **XIC 717.5428 NL: 1.01E5**

- Fragments containing oxFAs
- Fragments related to water loss
- Fragments not containing oxFAs
- Position-specific fragments
- Fragments related to other oxLPPs

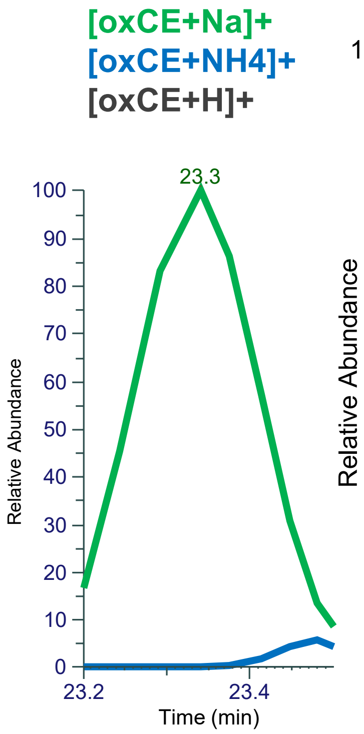


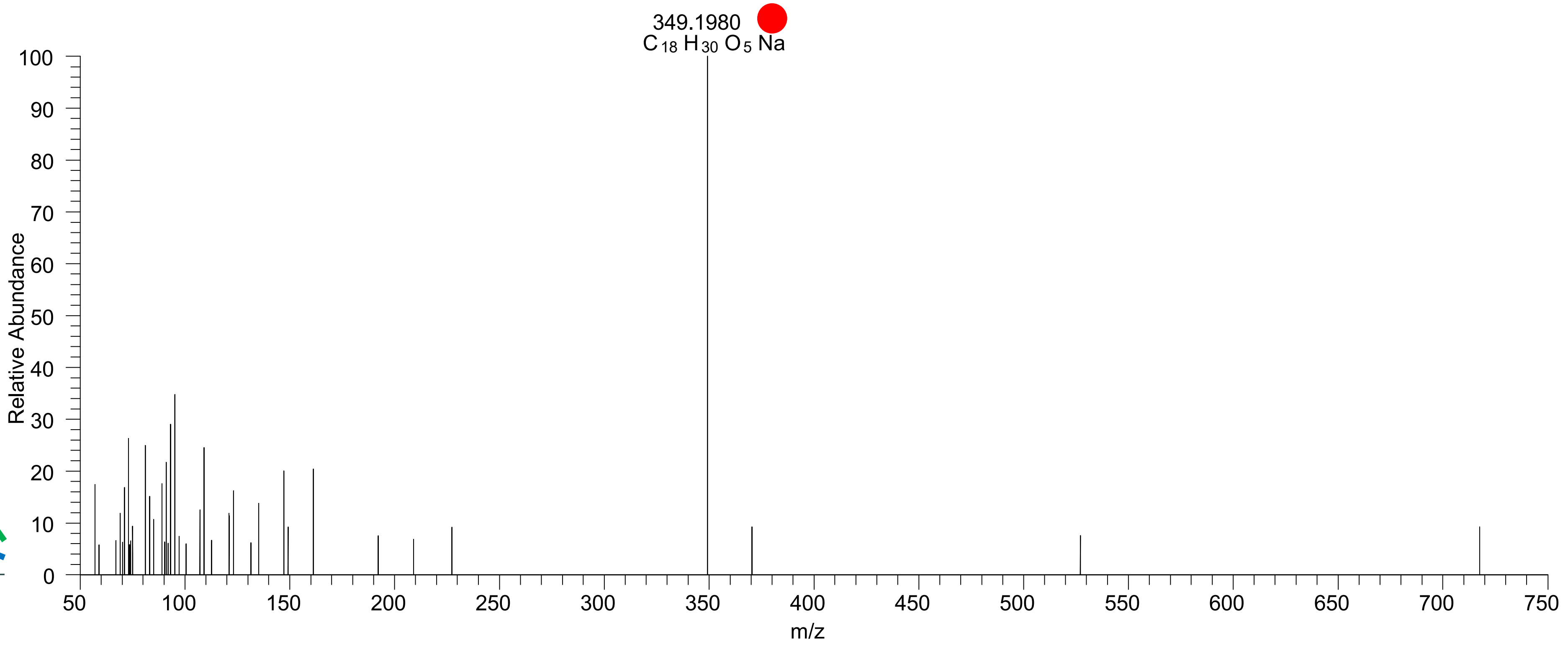

# CE(18:2<3O>)
# RT 21.4

**[oxCE+Na]+** **XIC 719.5585 NL: 1.05E5**

- Fragments containing oxFAs
- Fragments related to water loss
- Fragments not containing oxFAs
- Position-specific fragments
- Fragments related to other oxLPPs

**[oxCE+Na]+**
**[oxCE+NH4]+**
**[oxCE+H]+**

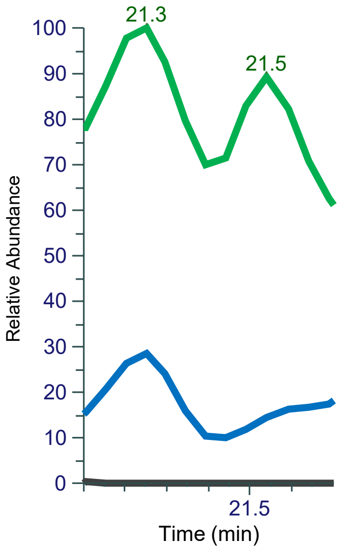


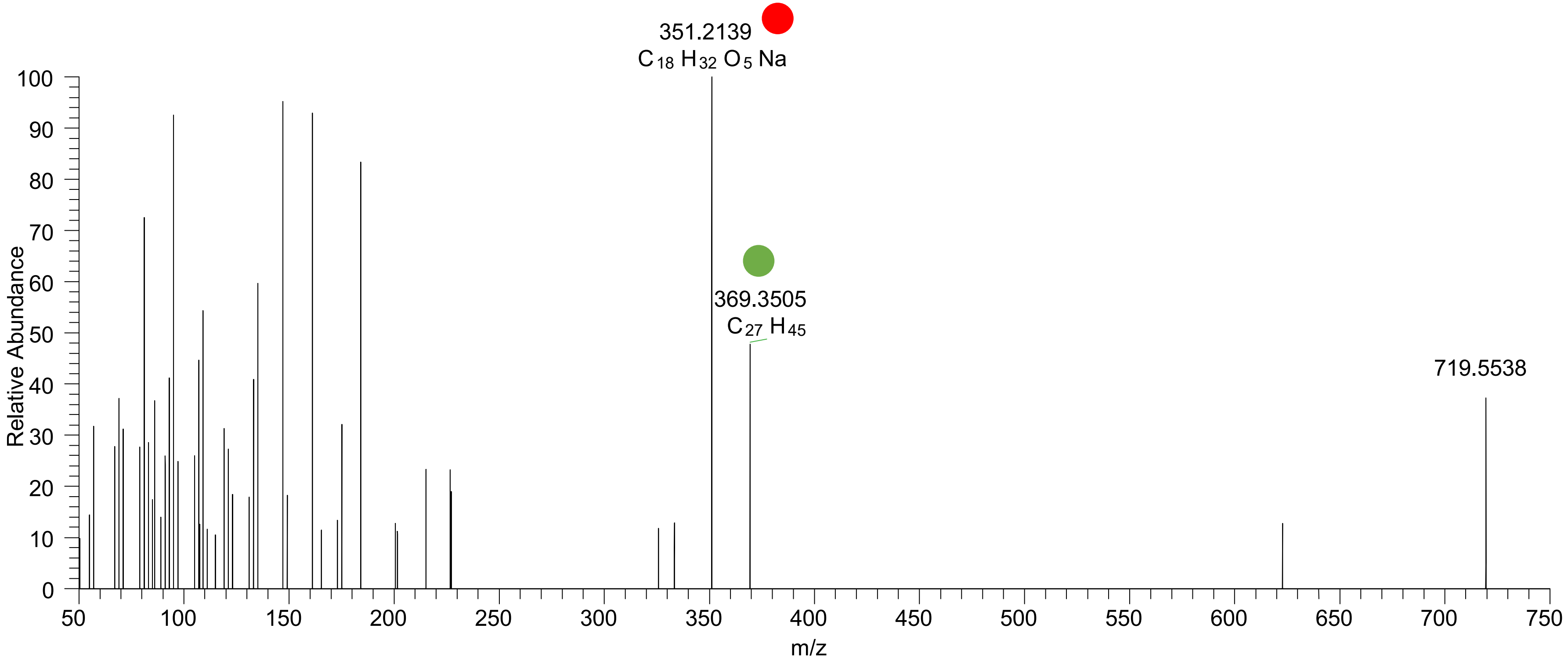

### CE(18:2<OH{9},OOH{13}>)
# RT 22.3

**[oxCE+Na]+** **XIC 719.5585 NL: 2.66E5**

- Fragments containing oxFAs
- Fragments related to water loss
- Fragments not containing oxFAs
- Position-specific fragments
- Fragments related to other oxLPPs

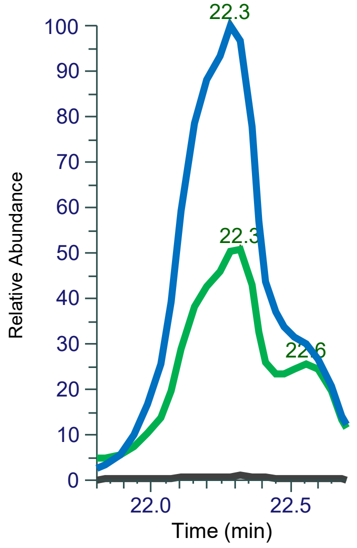


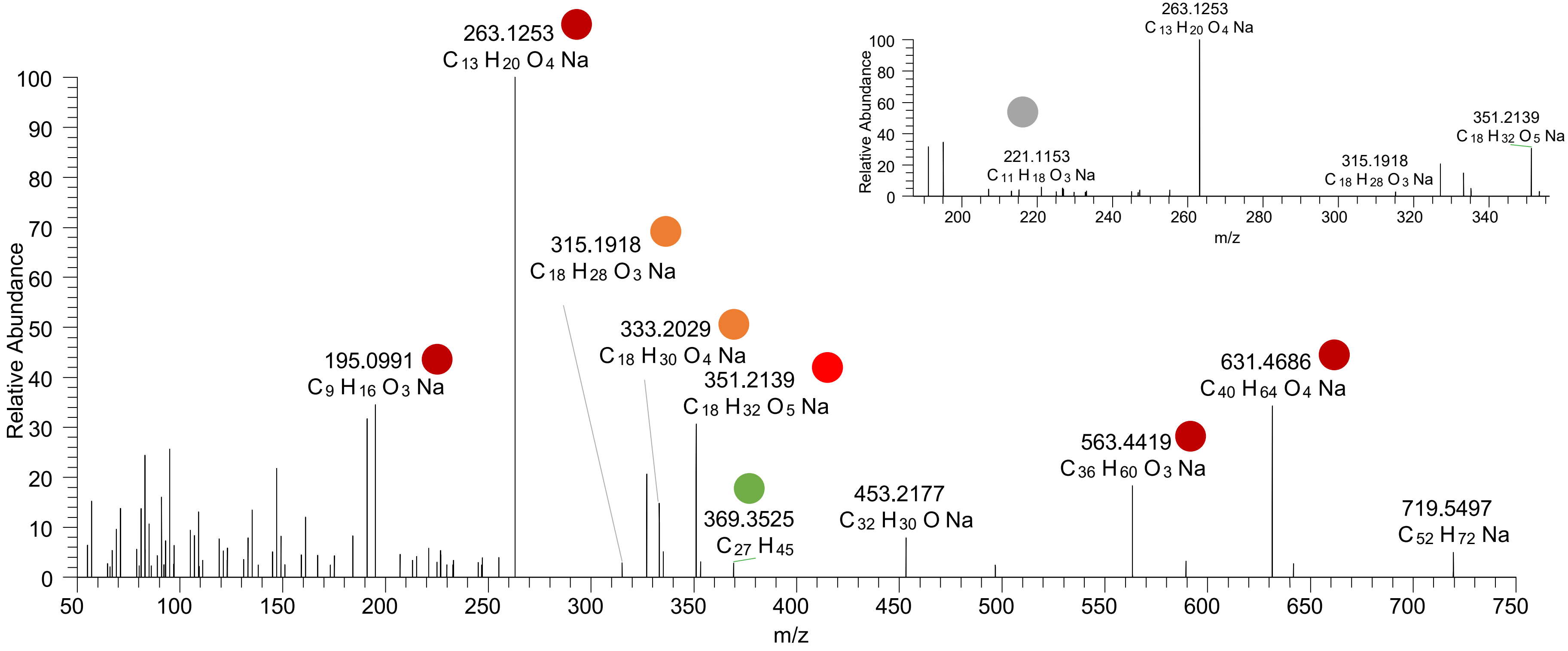

### CE(18:1<ep{10-11},OOH{13}>)
# RT 22.6

**[oxCE+Na]+** **XIC 719.5585 NL: 2.66E5**

- Fragments containing oxFAs
- Fragments related to water loss
- Fragments not containing oxFAs
- Position-specific fragments
- Fragments related to other oxLPPs

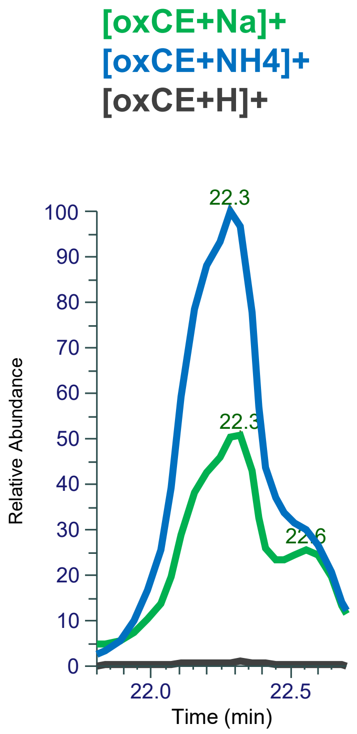


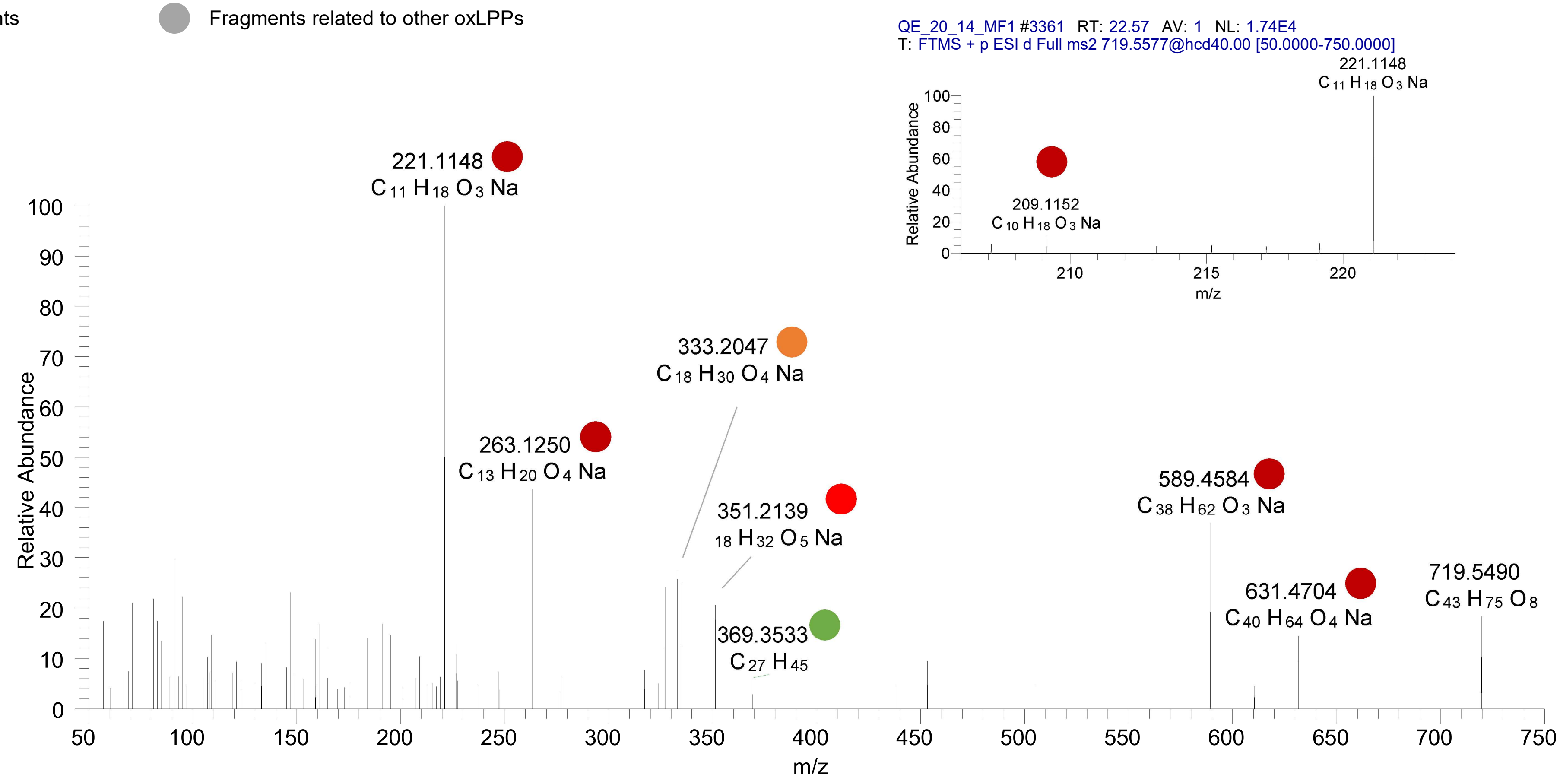

### CE(20:5<OH,oxo>)
# RT 20.6

**[oxCE+Na]+** **XIC 723.5323 NL: 8.34E4**

- Fragments containing oxFAs
- Fragments related to water loss
- Fragments not containing oxFAs
- Position-specific fragments
- Fragments related to other oxLPPs

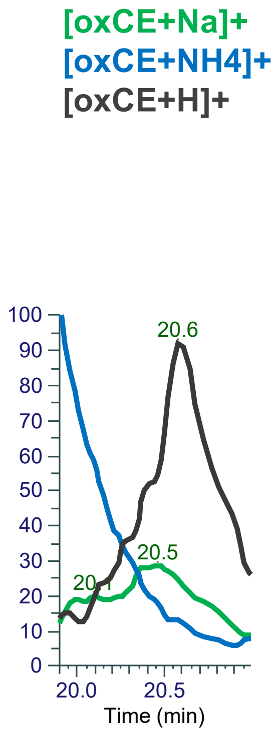


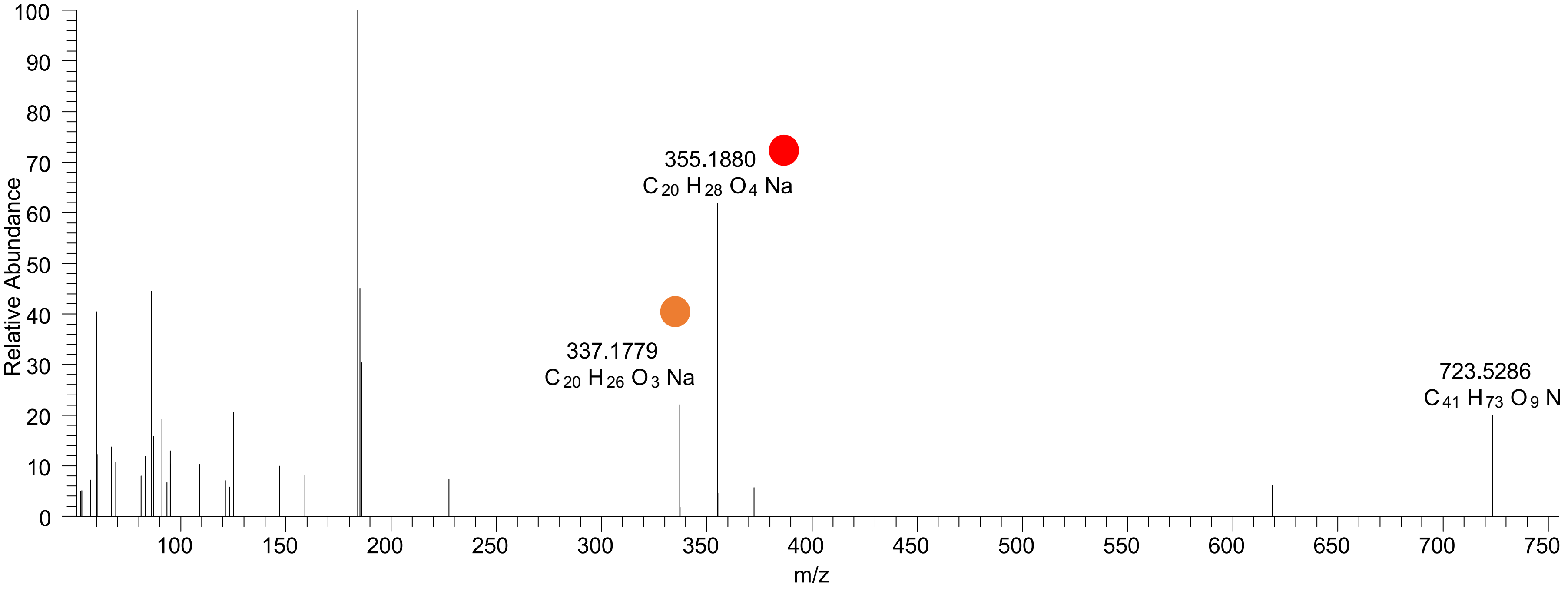

### CE(20:5<oxo,O>)
# RT 21.1-21.2

**[oxCE+Na]+** **XIC 723.5323 NL: 3.31E4**

- Fragments containing oxFAs
- Fragments related to water loss
- Fragments not containing oxFAs
- Position-specific fragments
- Fragments related to other oxLPPs

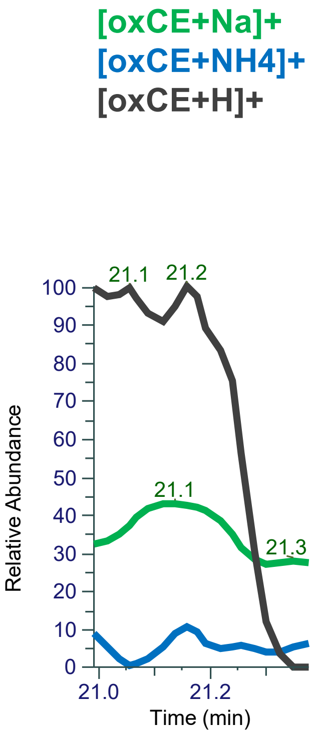


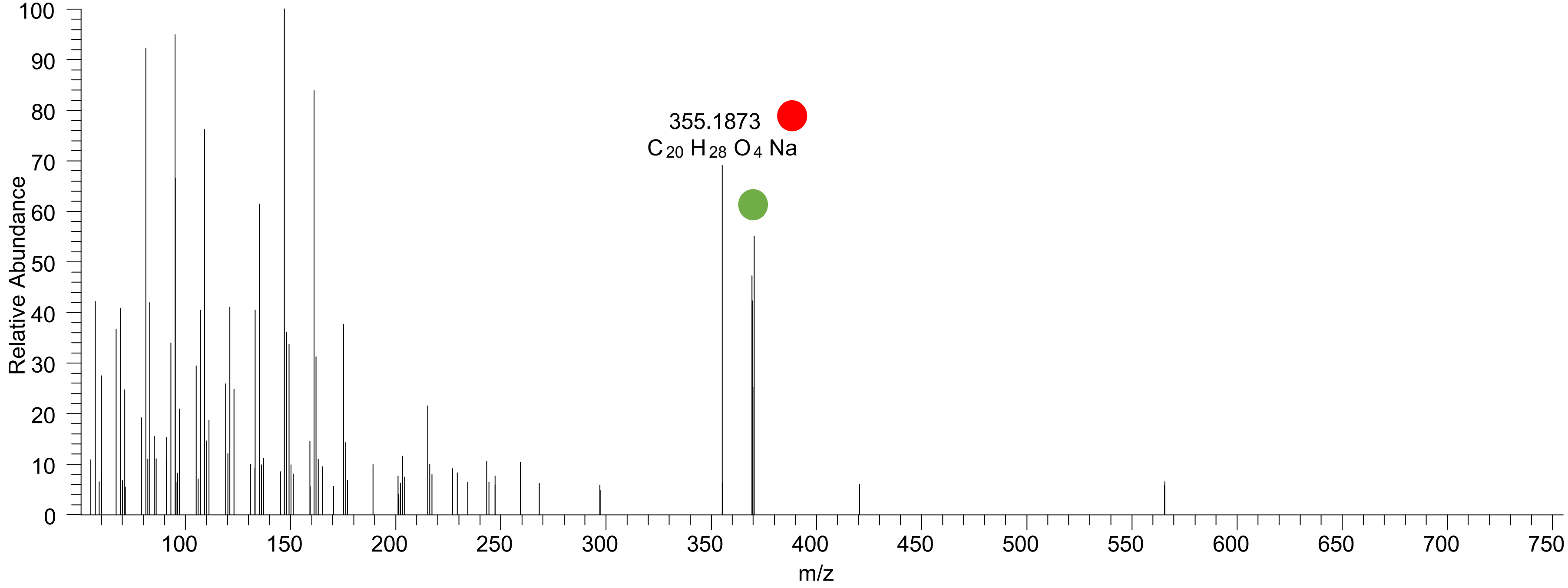

# CE(20:6<2O>)
# RT 22.5

[oxCE+Na]+ XIC 723.5323 NL: 4.84E4

Fragments containing oxFAs
Fragments related to water loss
Fragments not containing oxFAs
Position-specific fragments
Fragments related to other oxLPPs

[oxCE+Na]+
[oxCE+NH4]+
[oxCE+H]+

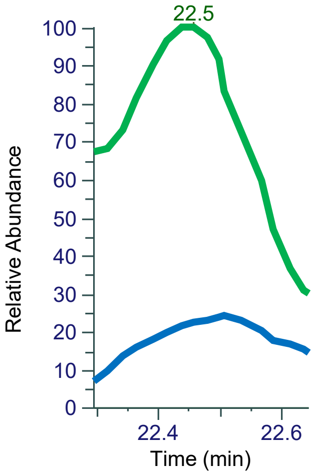


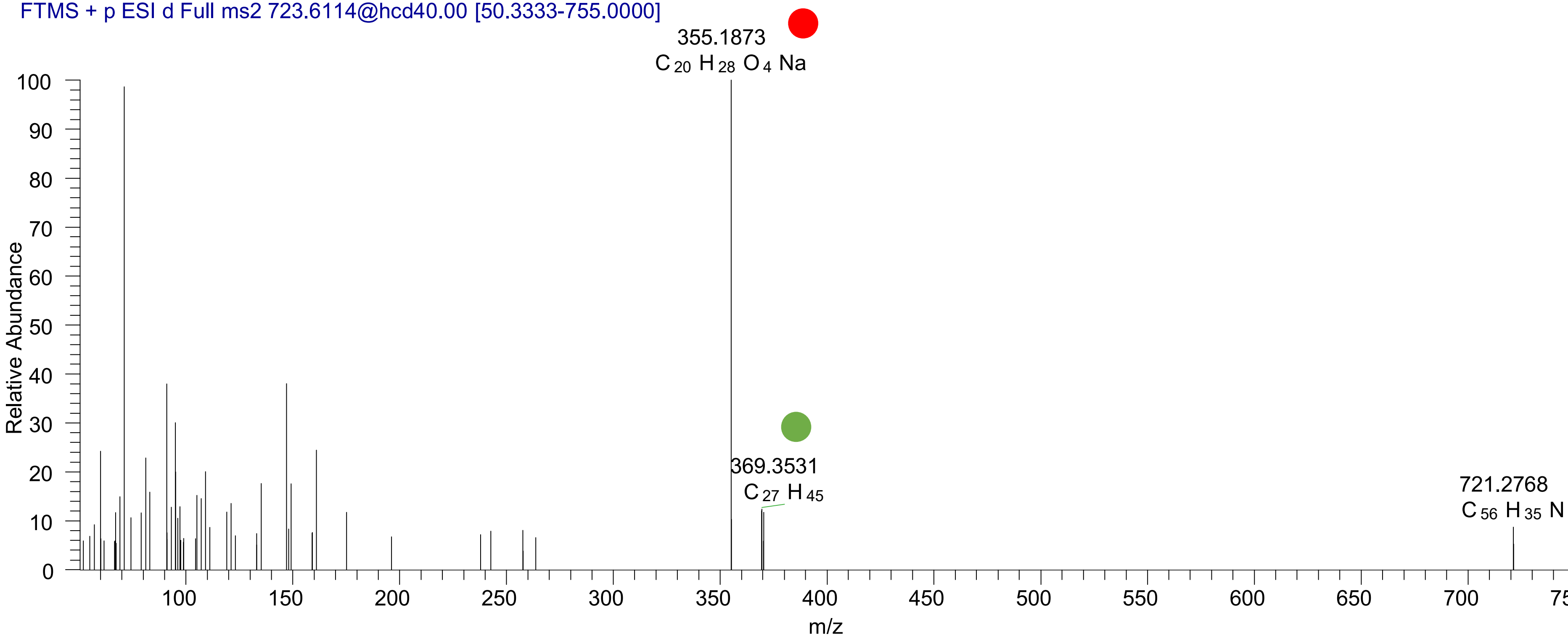

# CE(20:5<2OH{9,14}>)
# RT 20.4

[oxCE+Na]+ XIC 725.5479 NL: 1.03E6

- Fragments containing oxFAs
- Fragments related to water loss
- Fragments not containing oxFAs
- Position-specific fragments
- Fragments related to other oxLPPs

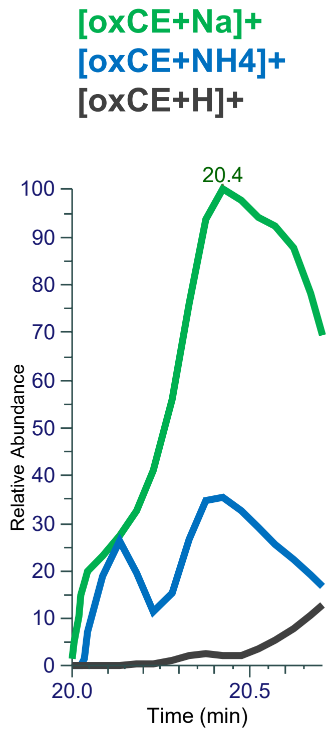


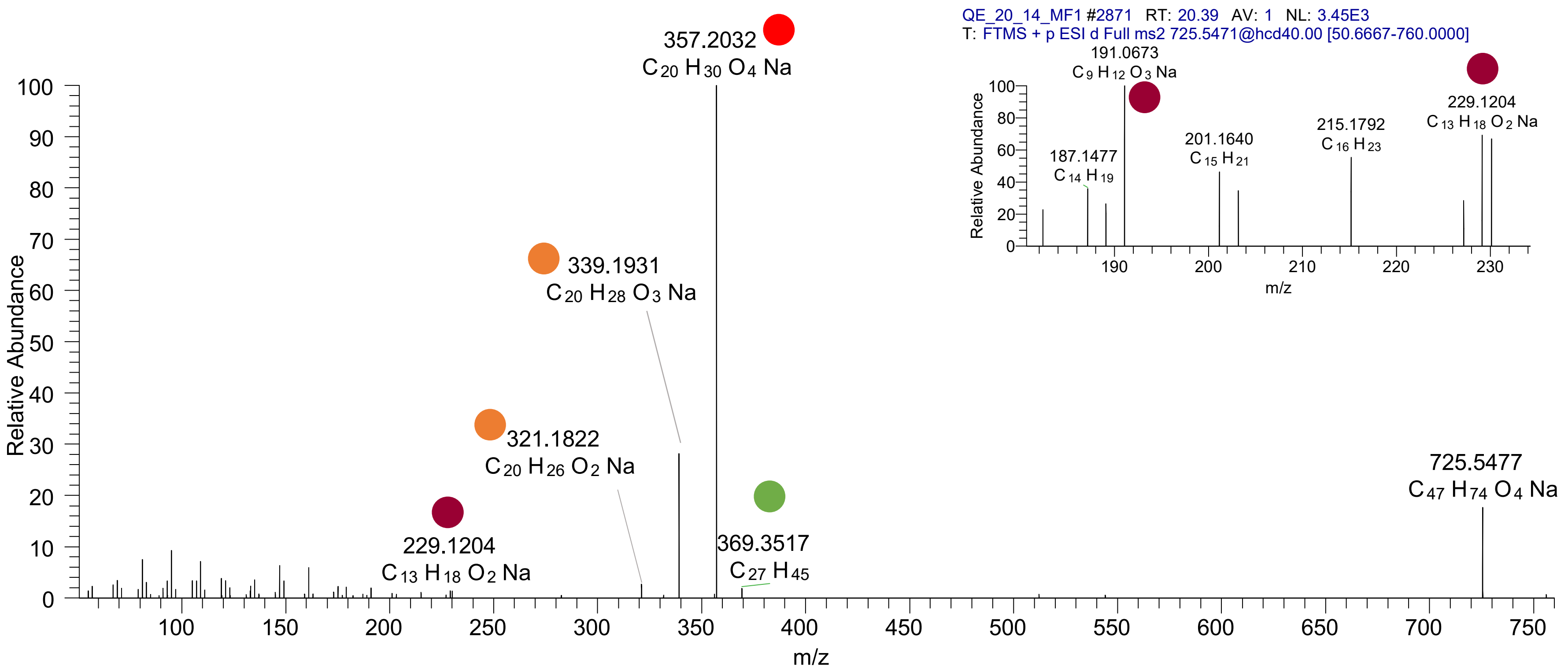

# CE(20:5<2OH{9,12}>)
# RT 20.6

**[oxCE+Na]+** **XIC 725.5479 NL: 1.03E6**

- Fragments containing oxFAs
- Fragments related to water loss
- Fragments not containing oxFAs
- Position-specific fragments
- Fragments related to other oxLPPs

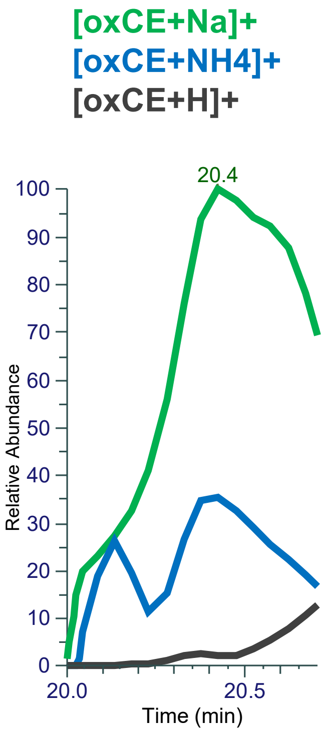


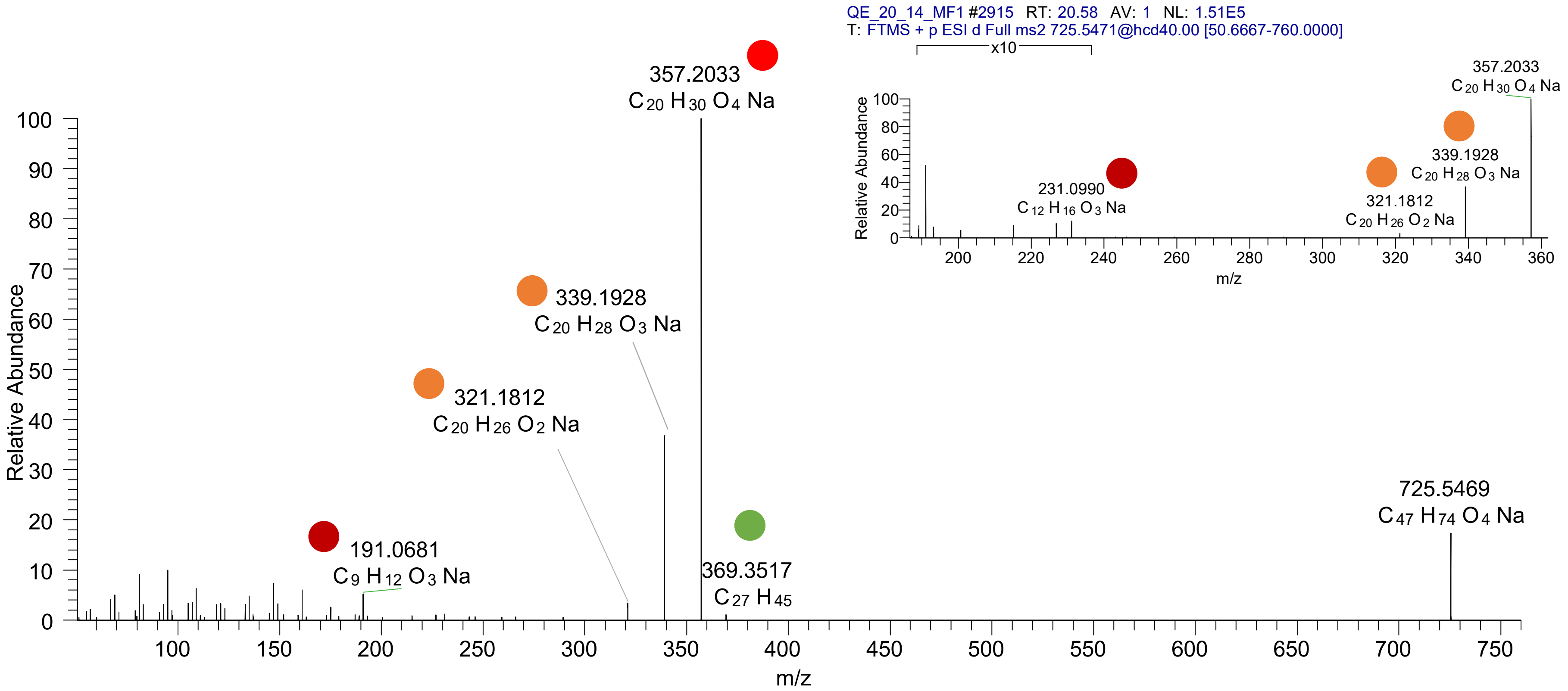

CE(20:5<OOH{9}>)
CE(20:5<OOH{12}>)
CE(20:5<OOH{15}>)
CE(20:5<OOH{18}>)
RT 23.0

**[oxCE+Na]+** **XIC 725.5479 NL: 6.18E5**

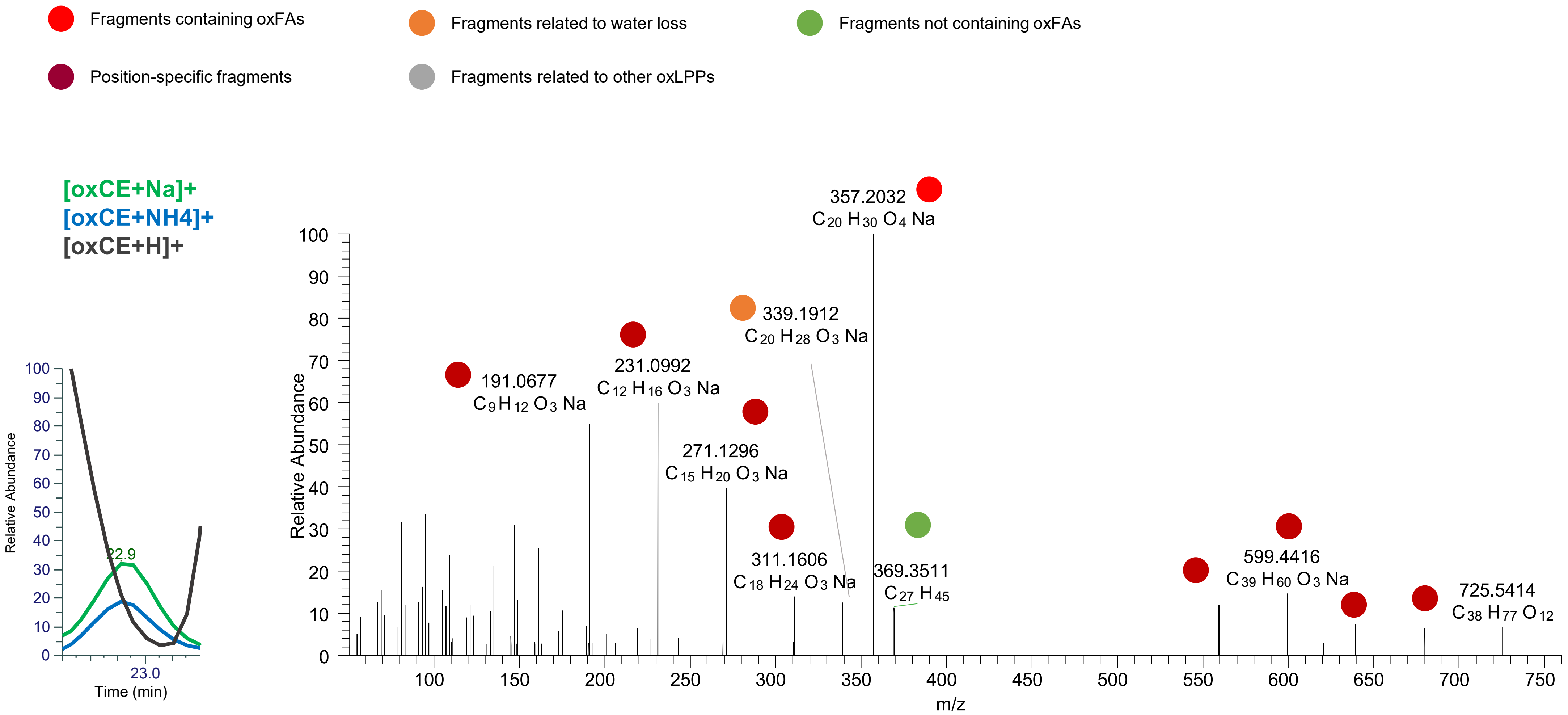

# CE(20:4<2OH{9,14}>)
# RT 21.1

**[oxCE+Na]+** **XIC 727.5636 NL: 3.04E6**

Fragments containing oxFAs
Fragments not containing oxFAs
Fragments relative to water loss
Specific position fragments
Fragments relative to other oxLPPs

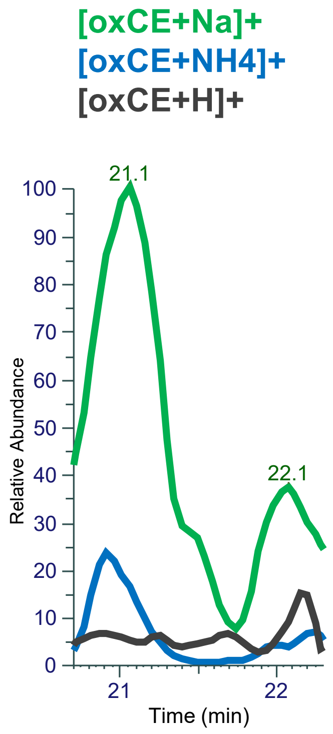


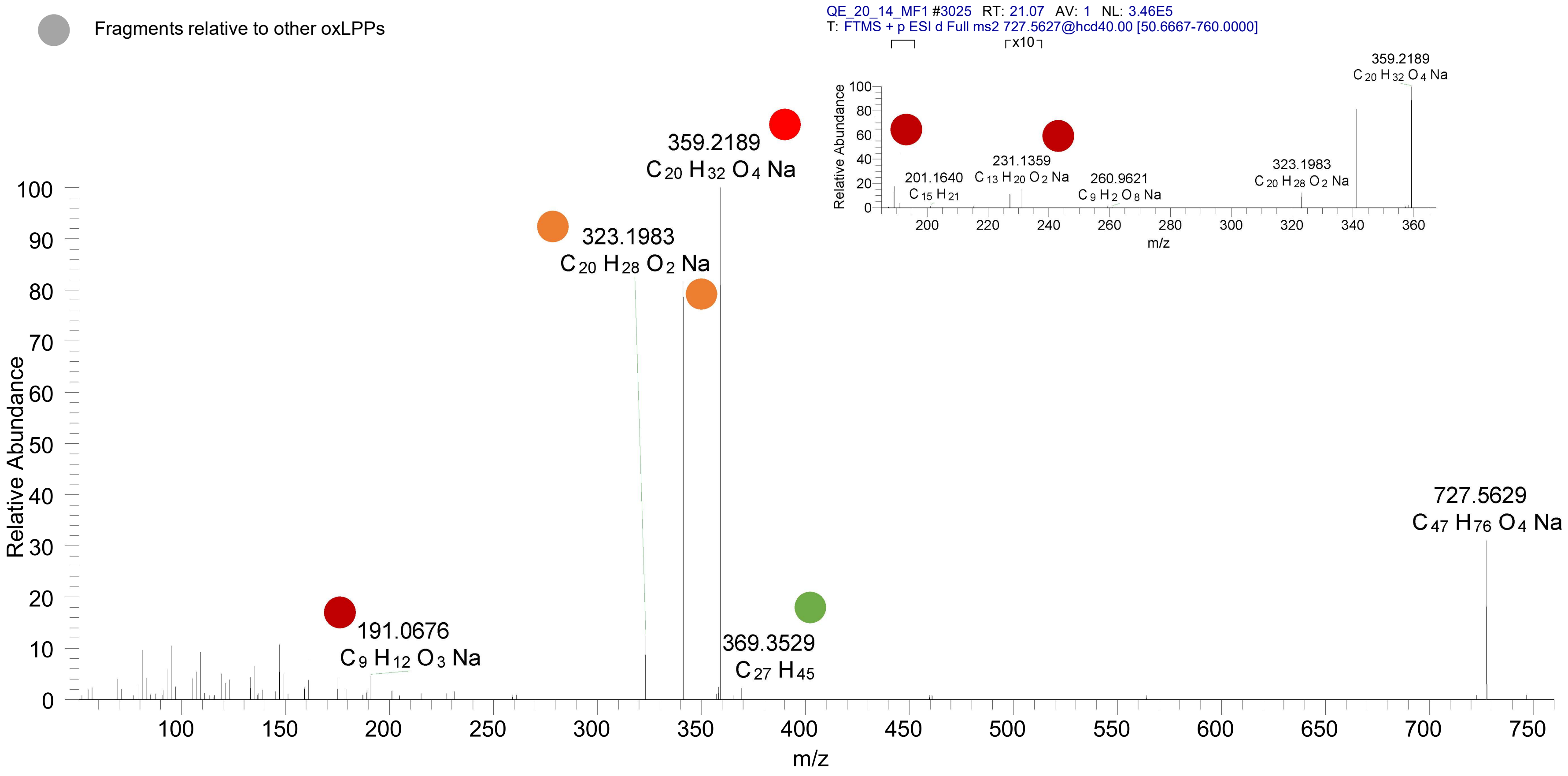

### CE(20:3<OH,oxo{11}>)
# RT 22.7

**[oxCE+Na]+** **XIC 727.5636 NL: 6.27E5**

- Fragments containing oxFAs
- Fragments related to water loss
- Fragments not containing oxFAs
- Position-specific fragments
- Fragments related to other oxLPPs

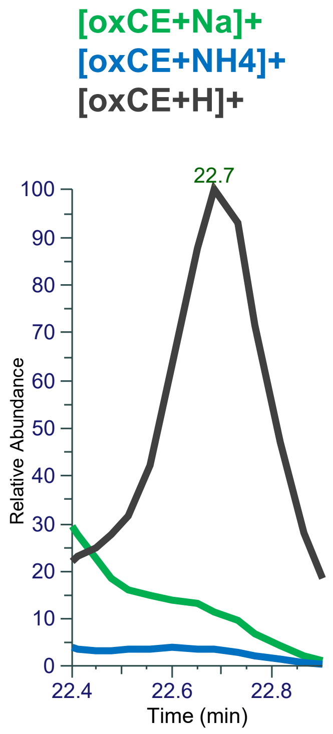


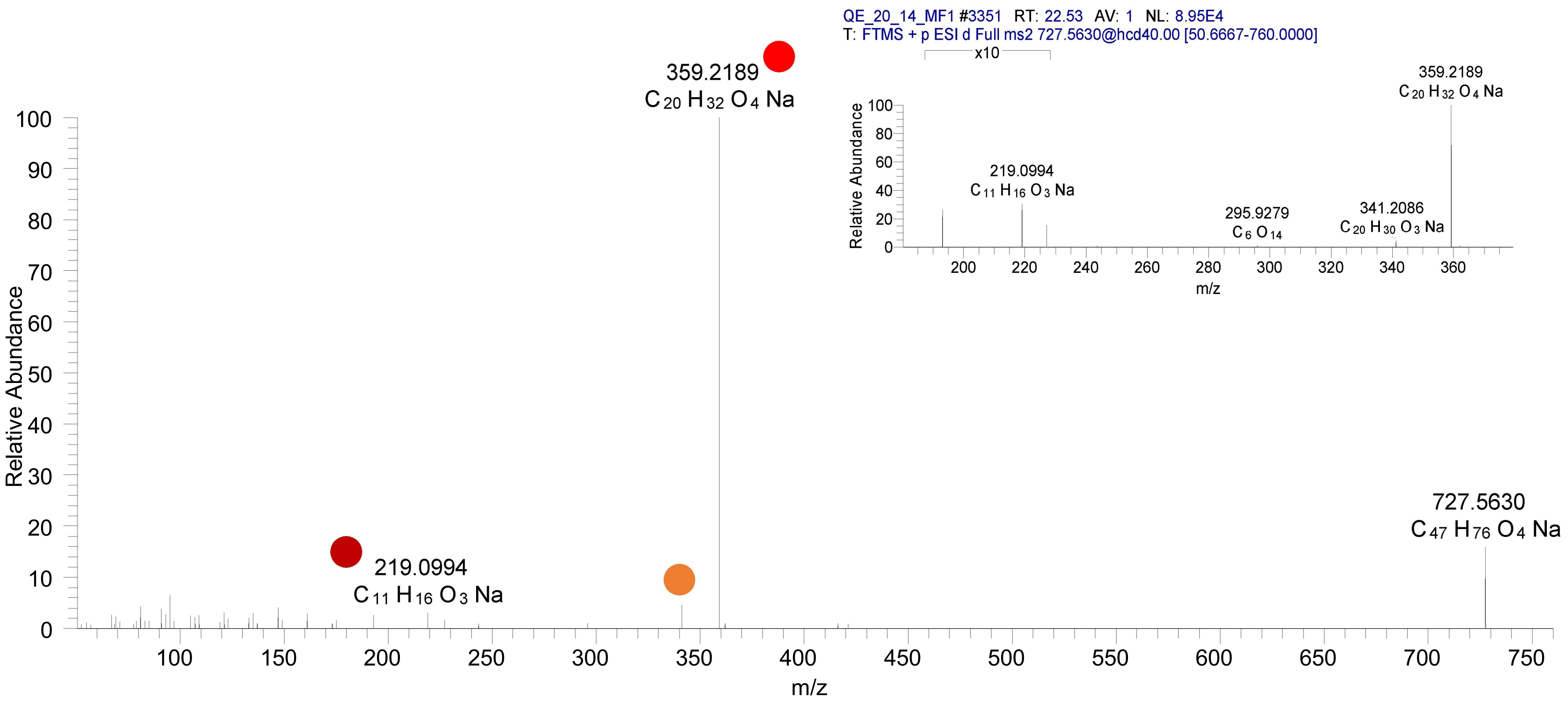

# CE(20:3<OH,ep>)
# RT 23.1

**[oxCE+Na]+** **XIC 727.5636 NL: 5.36E5**

Fragments containing oxFAs

Fragments related to water loss

Fragments not containing oxFAs

Position-specific fragments

Fragments related to other oxLPPs

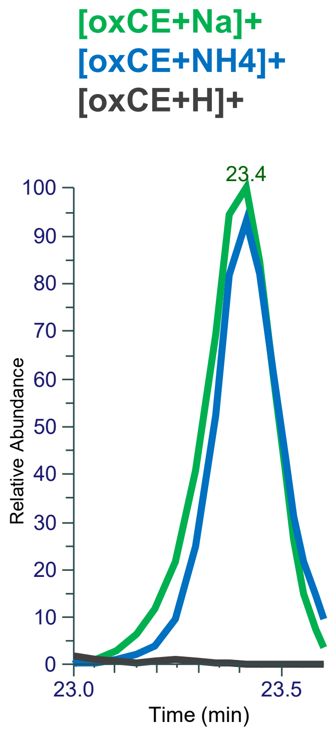


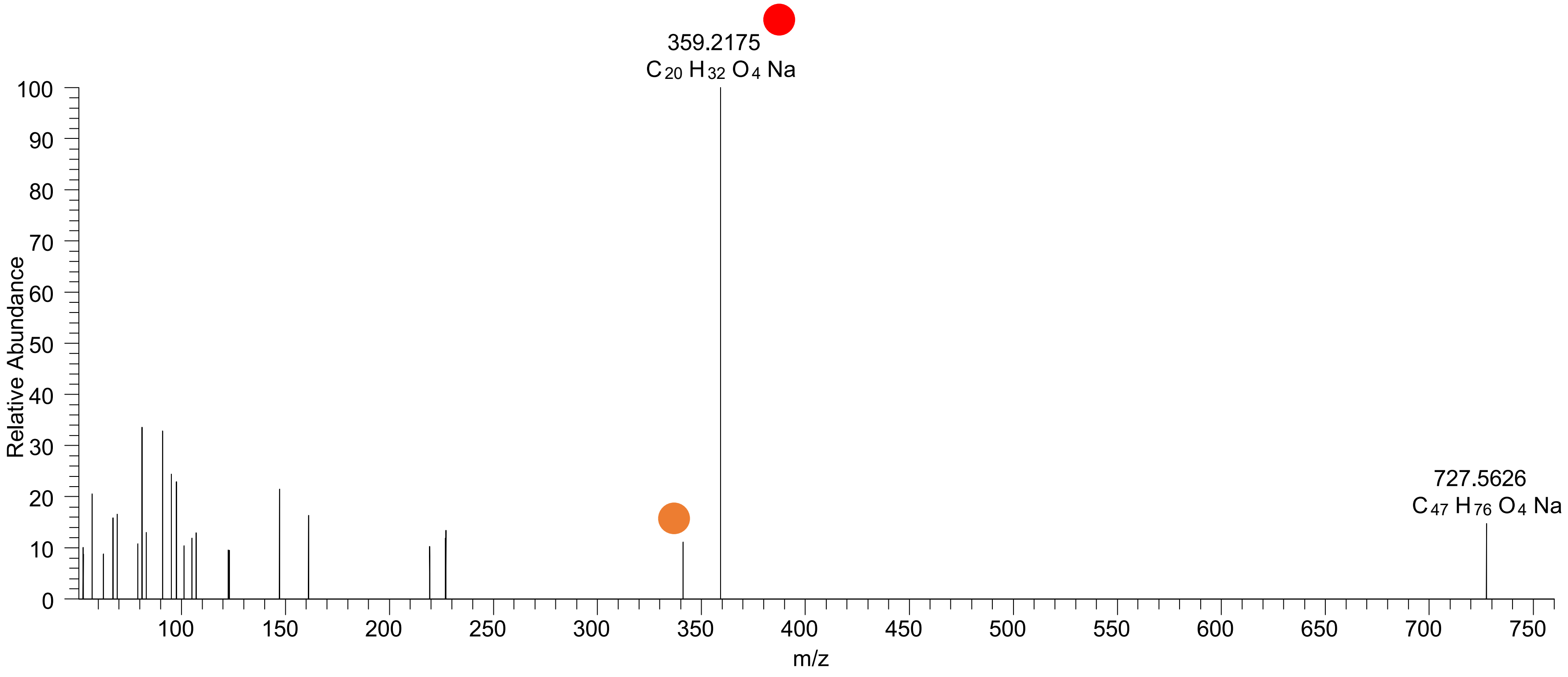

CE(20:4<OOH{9}>)
CE(20:4<OOH{10}>)
CE(20:4<OOH{12}>)
CE(20:4<OOH{15}>)
CE(20:4<OOH{13}>)
RT 23.3

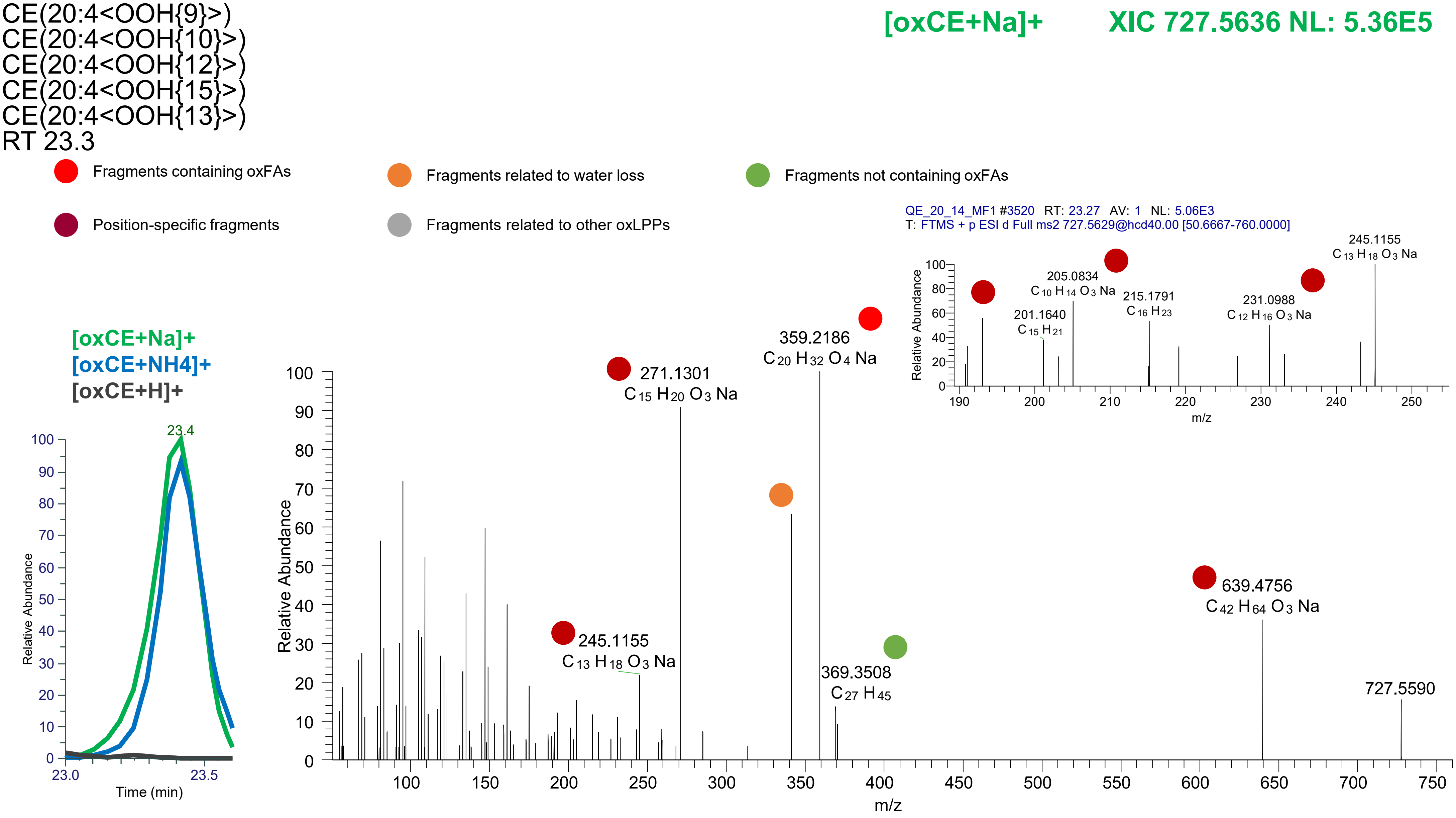

CE(20:4<OOH{9}>)
CE(20:4<OOH{11}>)
CE(20:4<OOH{12}>)
CE(20:4<OOH{15}>)
RT 23.4

**[oxCE+Na]+** **XIC 727.5636 NL: 5.36E5**

- Fragments containing oxFAs
- Fragments related to water loss
- Fragments not containing oxFAs
- Position-specific fragments
- Fragments related to other oxLPPs

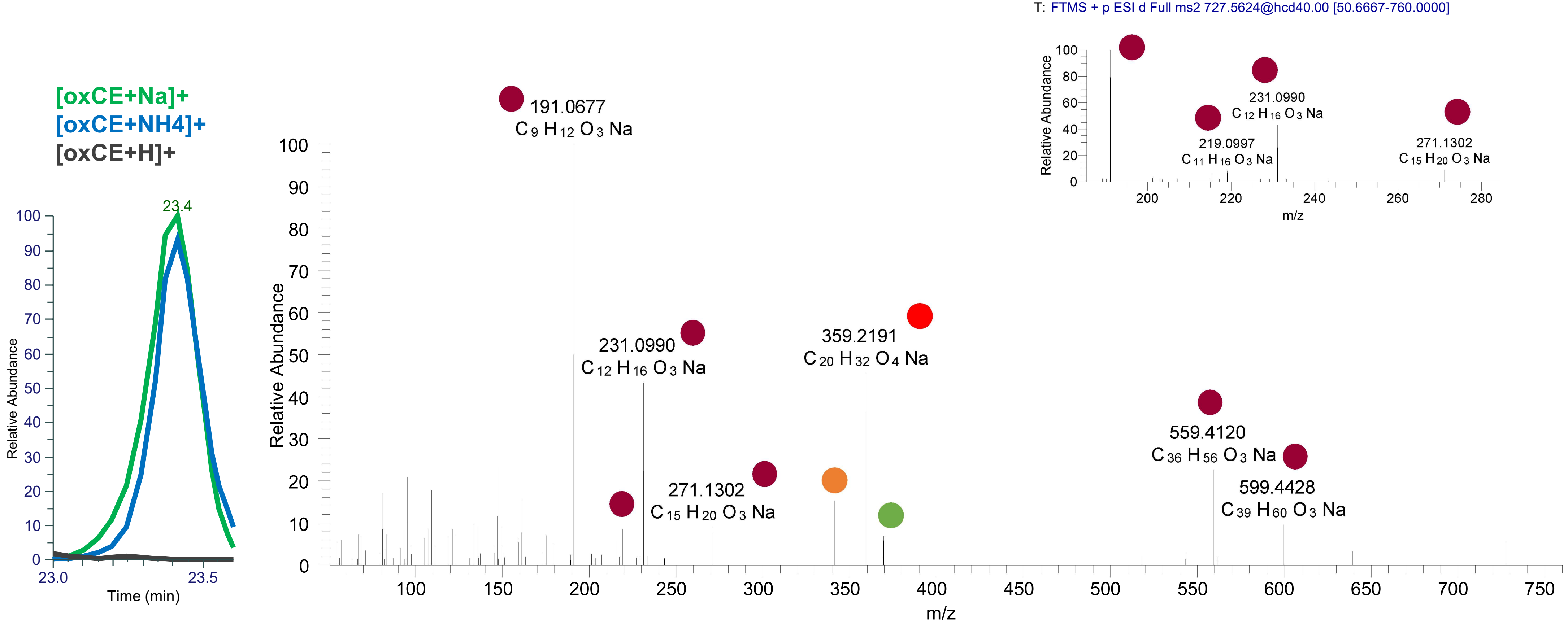

# CE(20:3<OH{9},OH>)
# RT 21.5

**[oxCE+Na]+** **XIC 729.5798 NL: 1.06E5**

- Fragments containing oxFAs
- Fragments related to water loss
- Fragments not containing oxFAs
- Position-specific fragments
- Fragments related to other oxLPPs

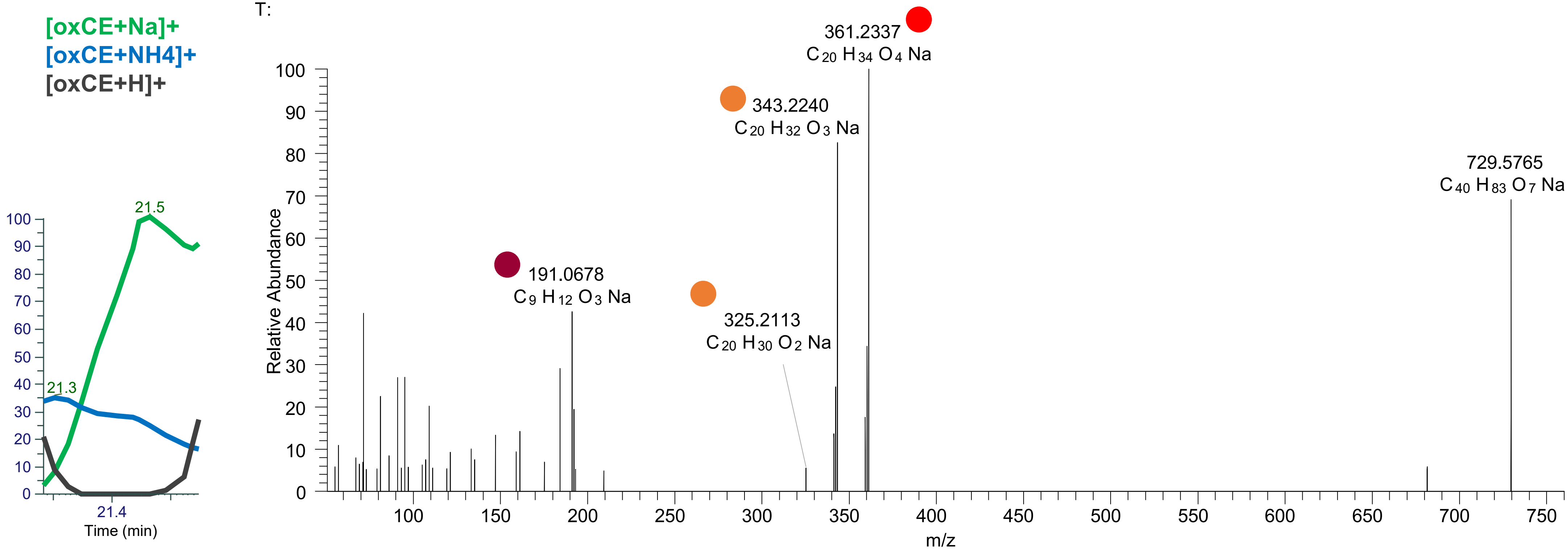

# CE(20:2<OH,ep>)
# RT 22.8

**[oxCE+Na]+** **XIC 729.5798 NL: 9.45E4**

- Fragments containing oxFAs
- Fragments related to water loss
- Fragments not containing oxFAs
- Position-specific fragments
- Fragments related to other oxLPPs

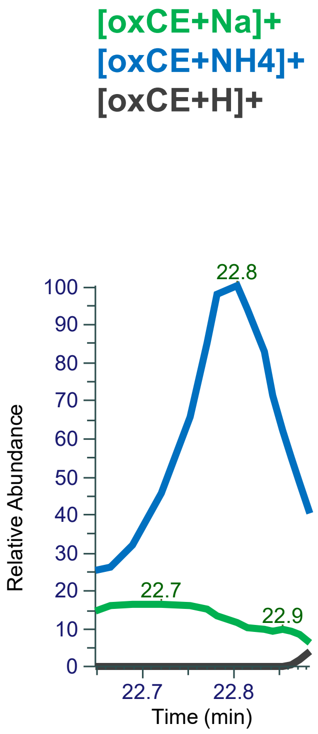


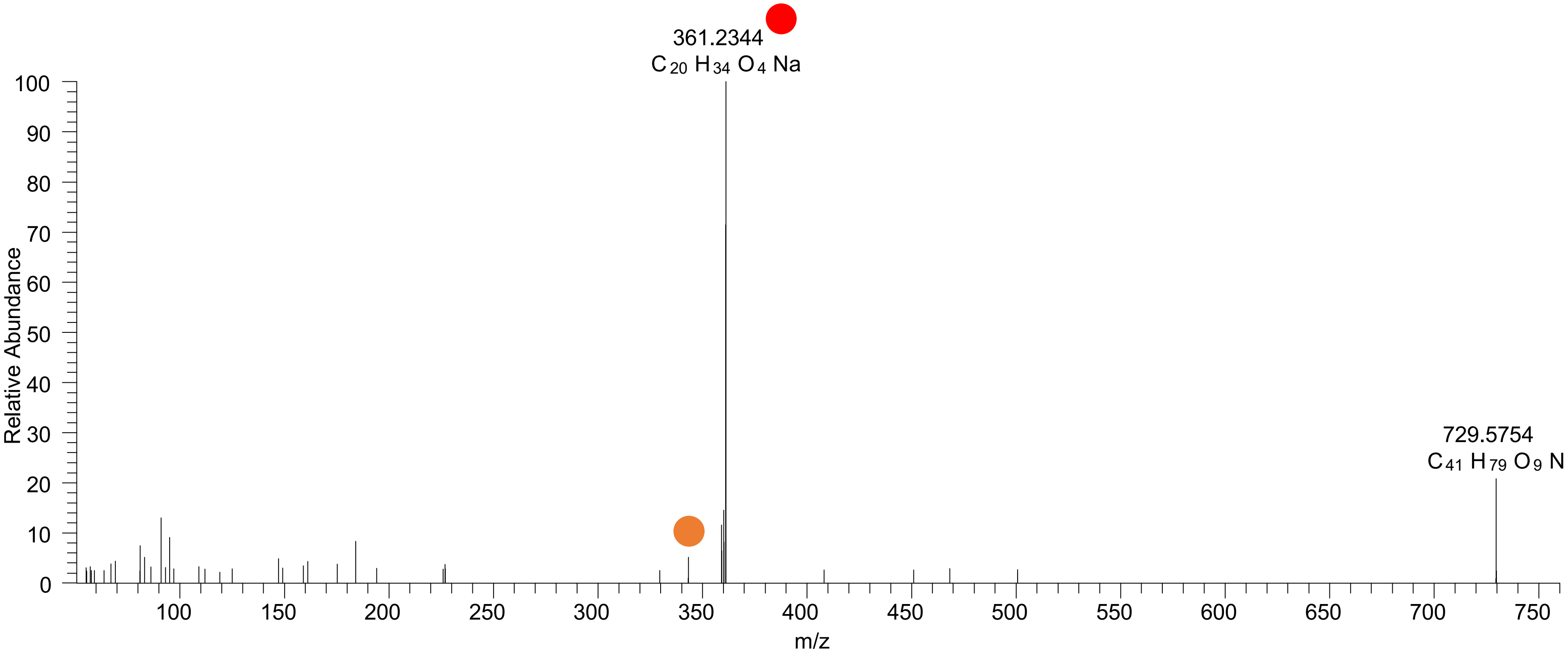

### CE(20:3<OOH{15}>)
### CE(20:3<OOH{12}>)
# RT 23.8

**[oxCE+Na]+** **XIC 729.5798 NL: 9.80E5**

- Fragments containing oxFAs
- Fragments related to water loss
- Fragments not containing oxFAs
- Position-specific fragments
- Fragments related to other oxLPPs

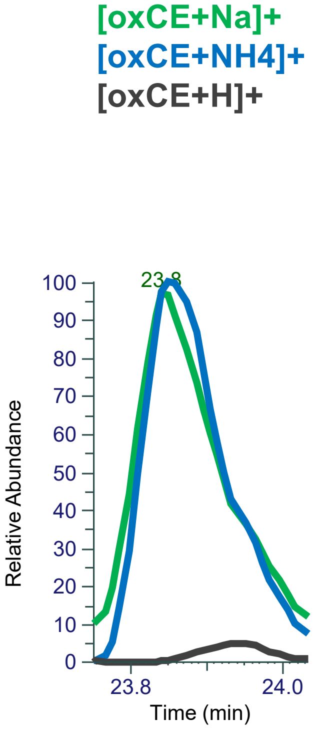


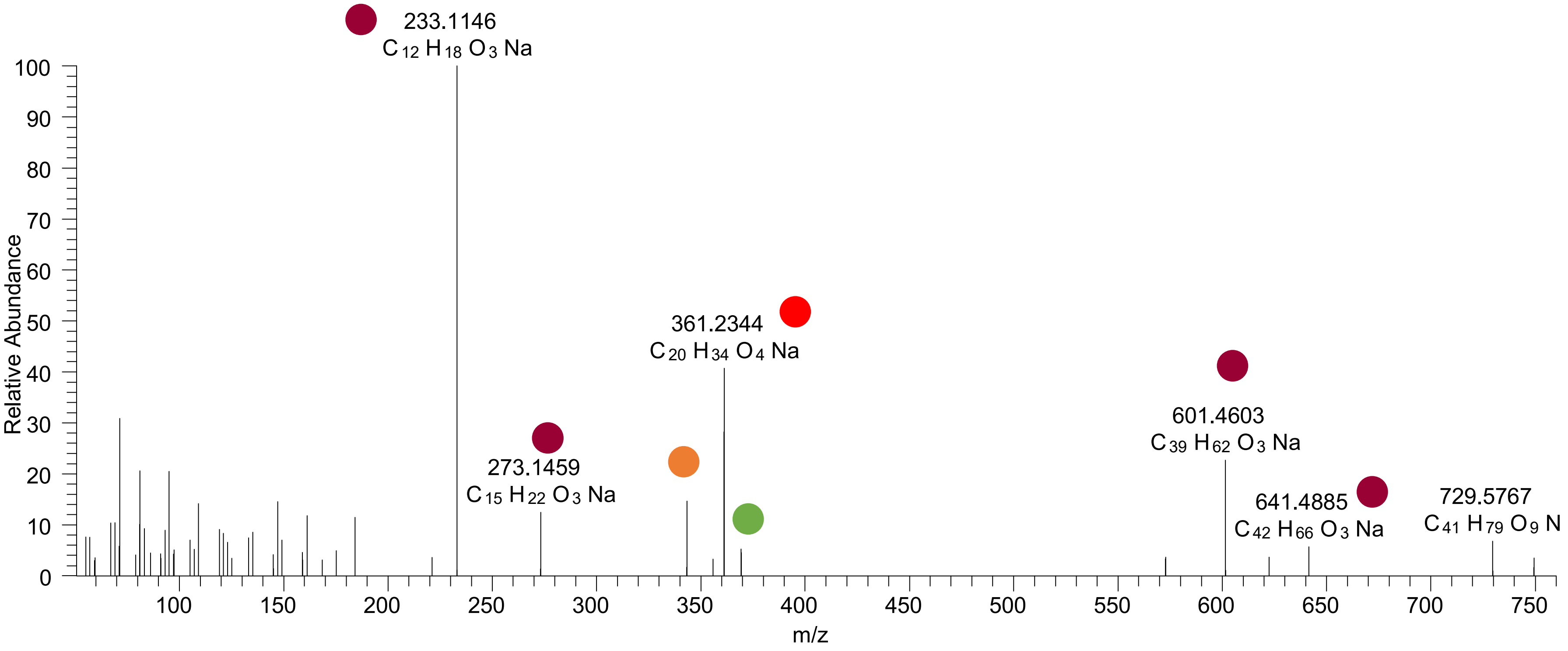

# CE(20:2<2O{12}>)
# RT 24.3

[oxCE+Na]+ XIC 731.5949 NL: 2.63E4

- Fragments containing oxFAs
- Fragments related to water loss
- Fragments not containing oxFAs
- Position-specific fragments
- Fragments related to other oxLPPs

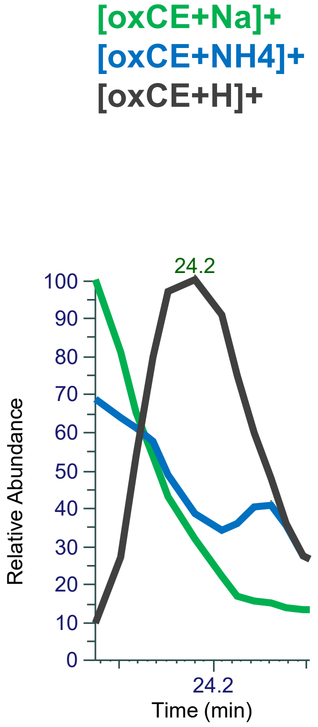


FTMS + p ESI d Full ms2 731.5939@hcd40.00 [51.0000-765.0000]

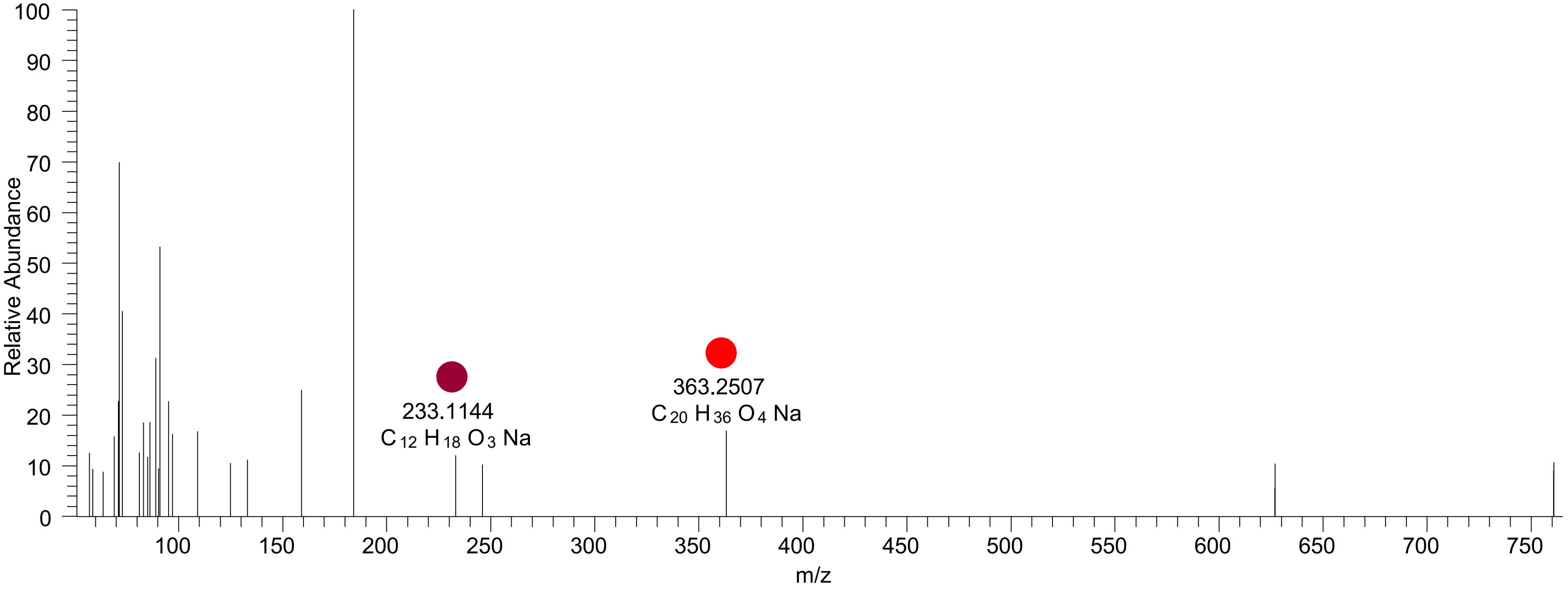

# CE(20:2<2O{15}>)
# RT 24.4

**[oxCE+Na]$^{+}$** **XIC 731.5949 NL: 5.13E4**

- Fragments containing oxFAs
- Fragments related to water loss
- Fragments not containing oxFAs
- Position-specific fragments
- Fragments related to other oxLPPs

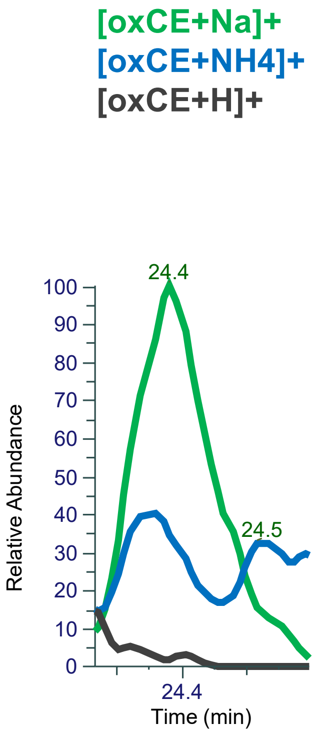


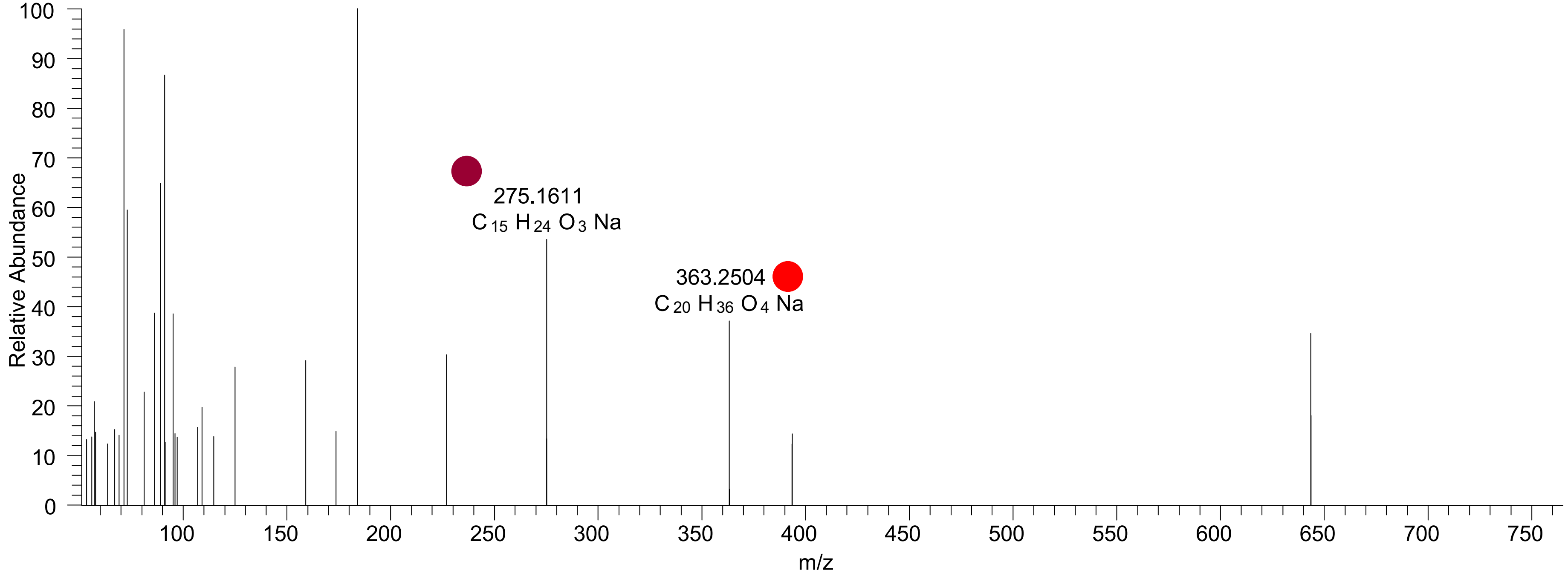

# CE(22:6<OH>)
# RT 23.4

**[oxCE+Na]+** **XIC 735.5687 NL: 4.31E5**

- Fragments containing oxFAs
- Fragments related to water loss
- Fragments not containing oxFAs
- Position-specific fragments
- Fragments related to other oxLPPs

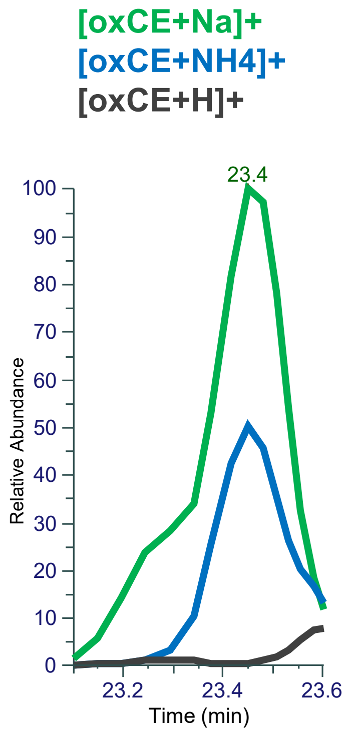


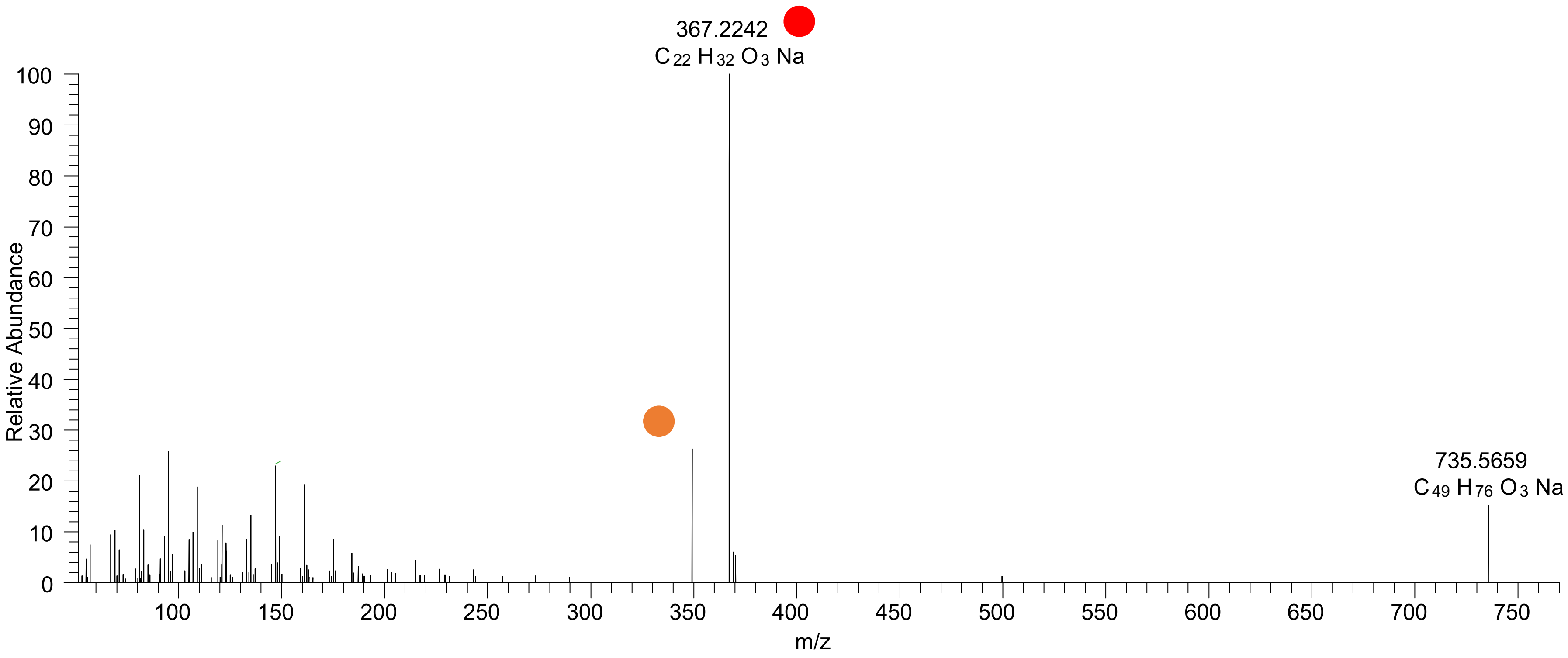

# CE(20:5<2OH,O>)
# RT 18.1

**[oxCE+Na]+** **XIC 741.5428 NL: 6.17E4**

- Fragments containing oxFAs
- Fragments related to water loss
- Fragments not containing oxFAs
- Position-specific fragments
- Fragments related to other oxLPPs

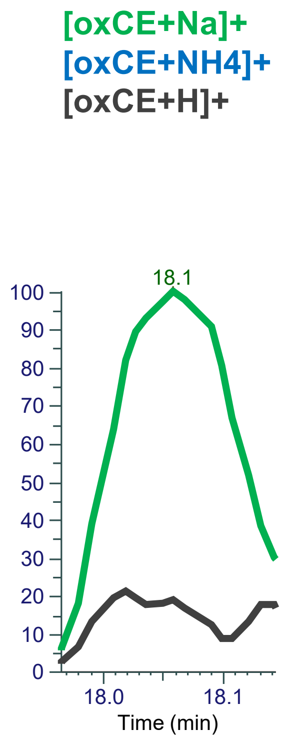


FTMS + p ESI d Full ms2 741.5415@hcd40.00 [51.6667-775.0000]

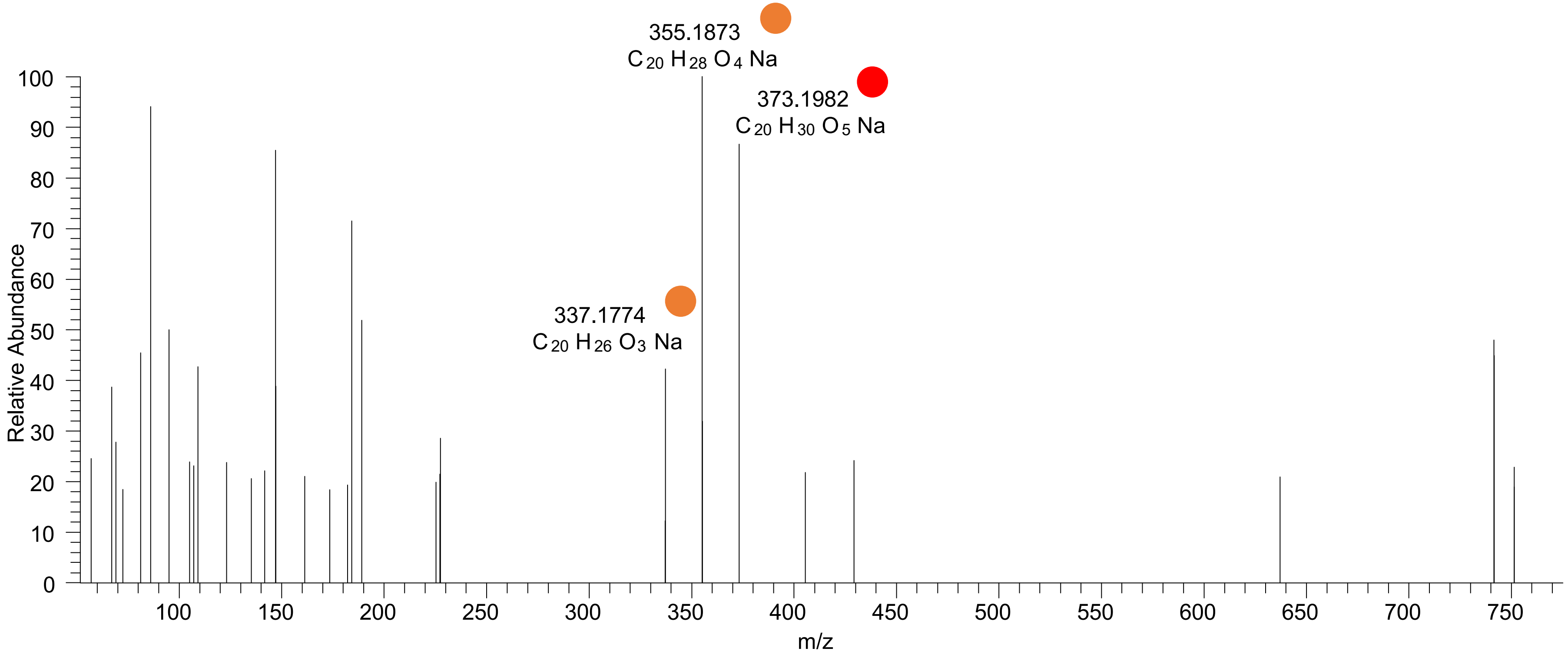

# CE(20:5<2OH,O>)
# RT 19.3

**[oxCE+Na]+** **XIC 741.5428 NL: 1.13E5**

Fragments containing oxFAs

Fragments related to water loss

Fragments not containing oxFAs

Position-specific fragments

Fragments related to other oxLPPs

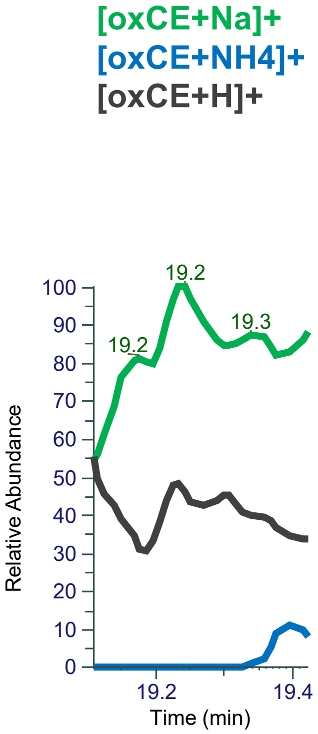


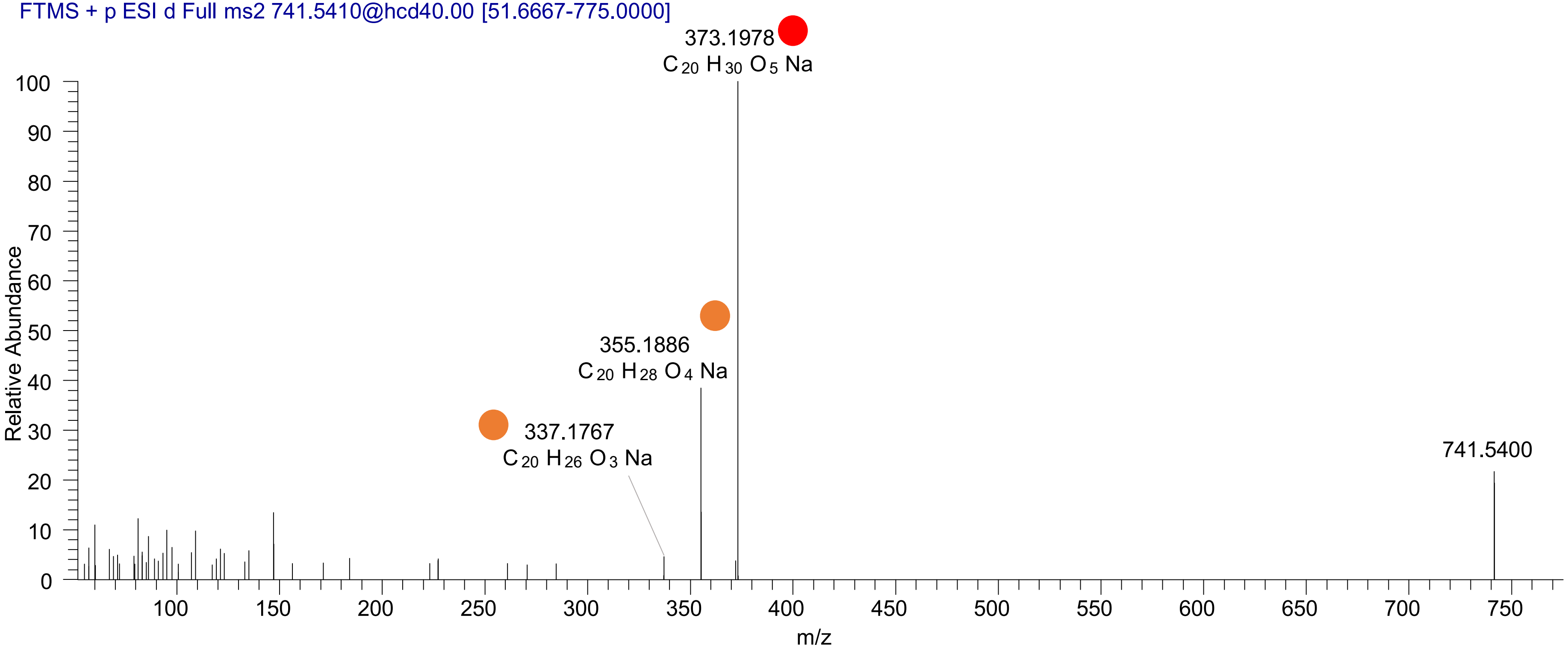

### CE(20:5<O,OOH{15}>)
### CE(20:5<O,OOH{9}>)
# RT 20.2

**[oxCE+Na]+** **XIC 741.5428 NL: 1.13E5**

- Fragments containing oxFAs
- Fragments related to water loss
- Fragments not containing oxFAs
- Position-specific fragments
- Fragments related to other oxLPPs

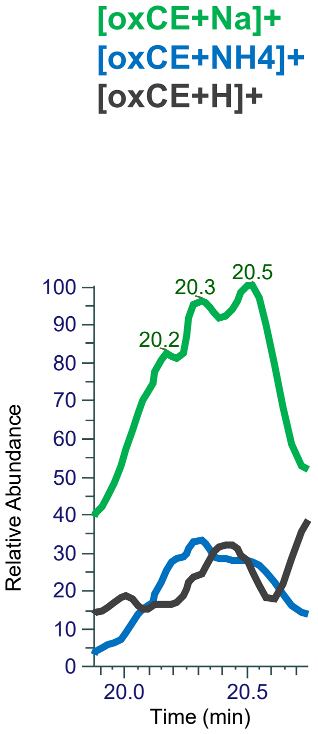


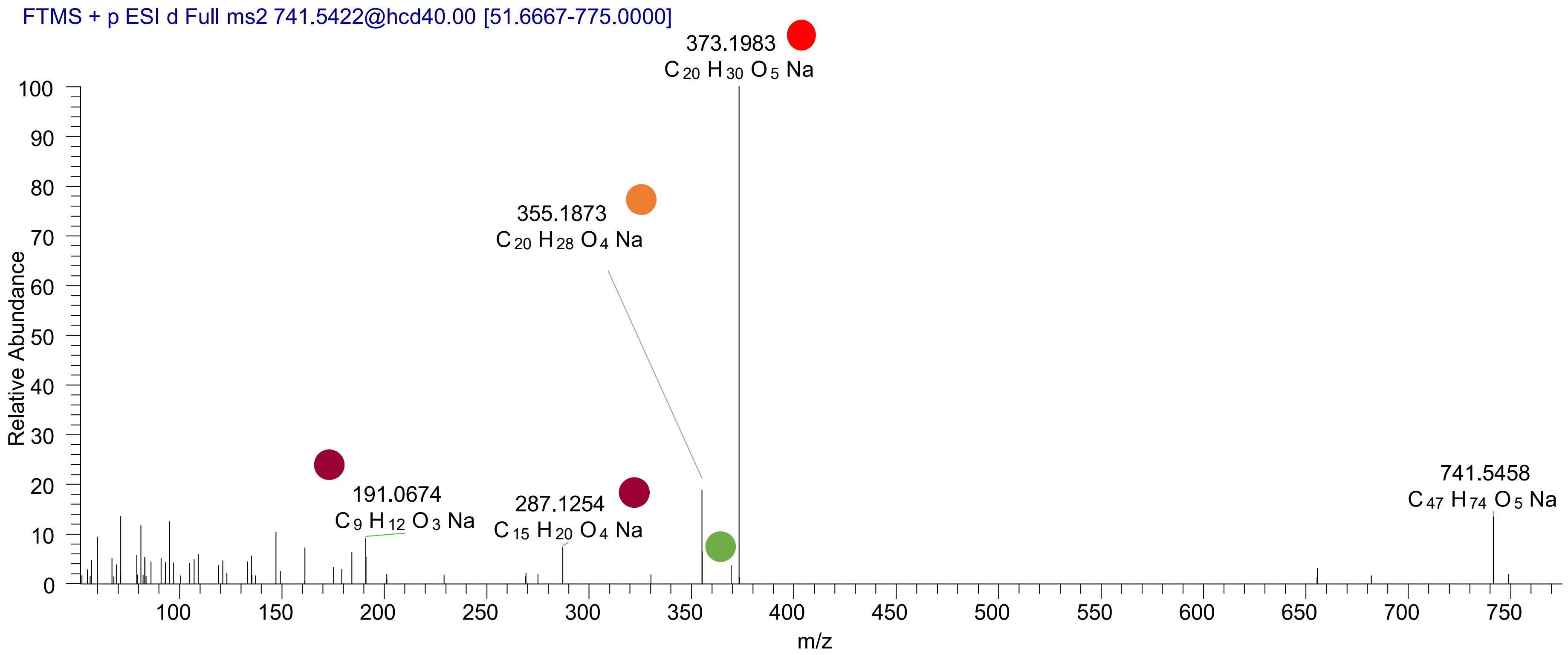

### CE(20:5<OH,OOH{15}>)
# RT 20.5

**[oxCE+Na]+** **XIC 741.5428 NL: 1.13E5**

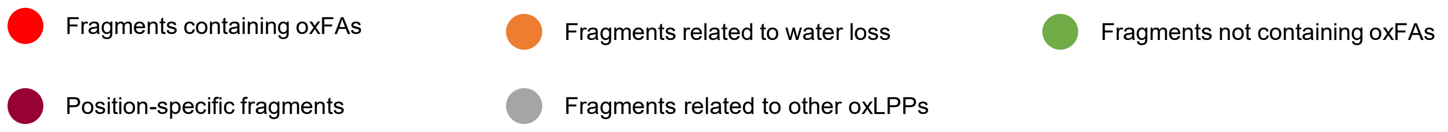


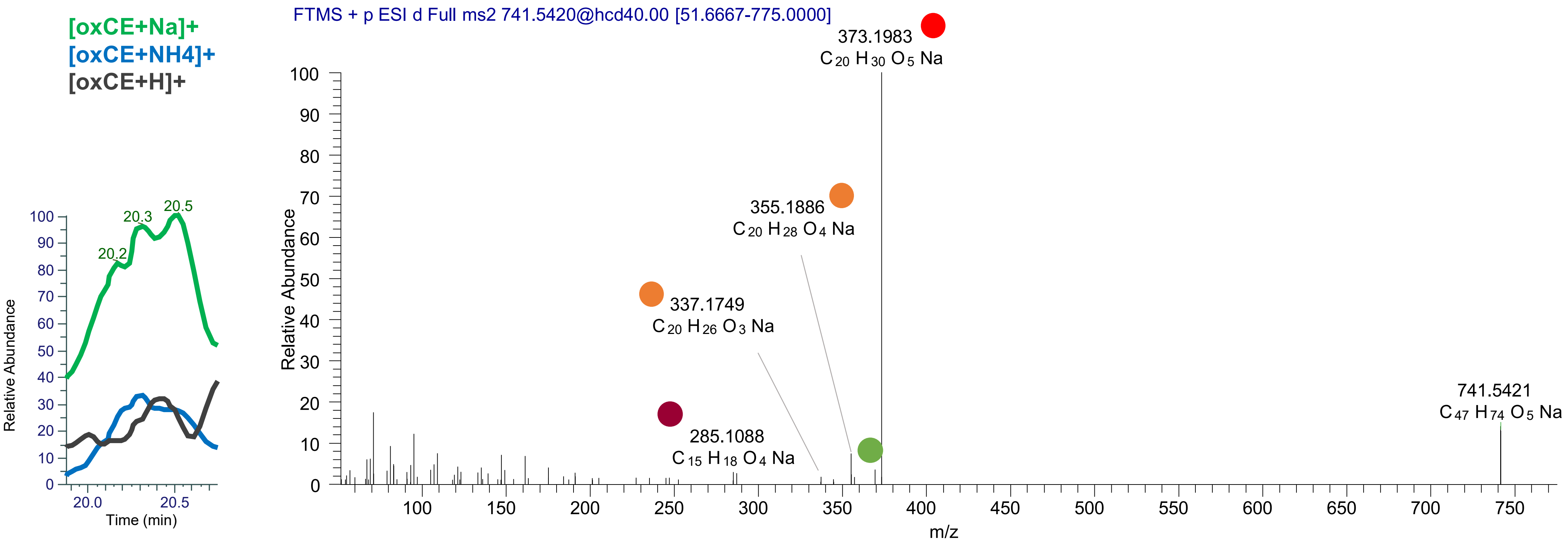

# CE(20:5<OH,2O{14}>)
# RT 21.1

**[oxCE+Na]+** **XIC 741.5428 NL: 2.59E5**

- Fragments containing oxFAs
- Fragments related to water loss
- Fragments not containing oxFAs
- Position-specific fragments
- Fragments related to other oxLPPs

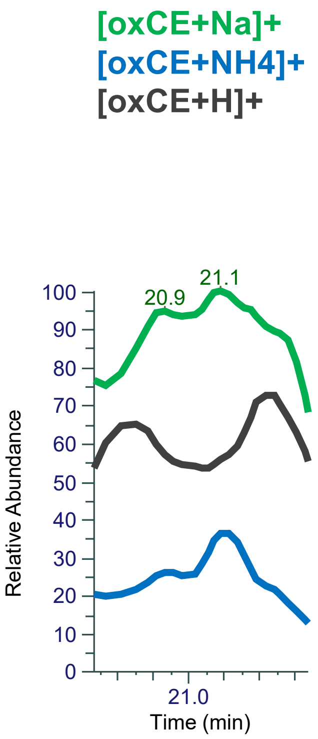


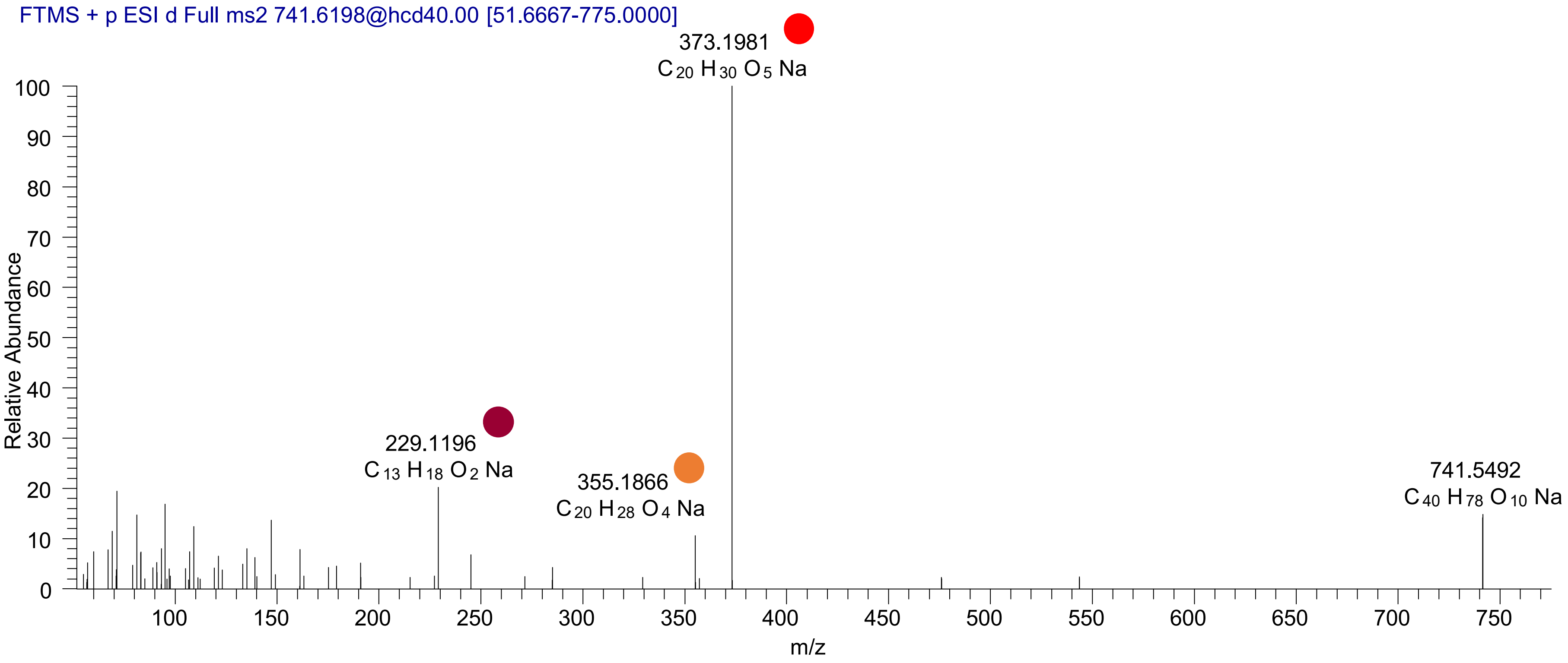

### CE(20:4<oxo,OOH{12}>)
# RT 21.2

**[oxCE+Na]+** **XIC 741.5428 NL: 2.59E5**

- Fragments containing oxFAs
- Fragments related to water loss
- Fragments not containing oxFAs
- Position-specific fragments
- Fragments related to other oxLPPs

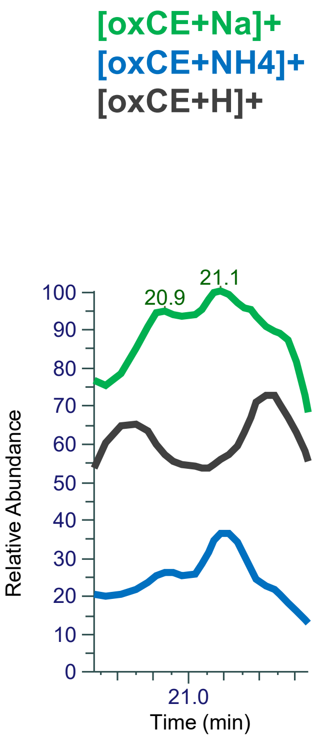


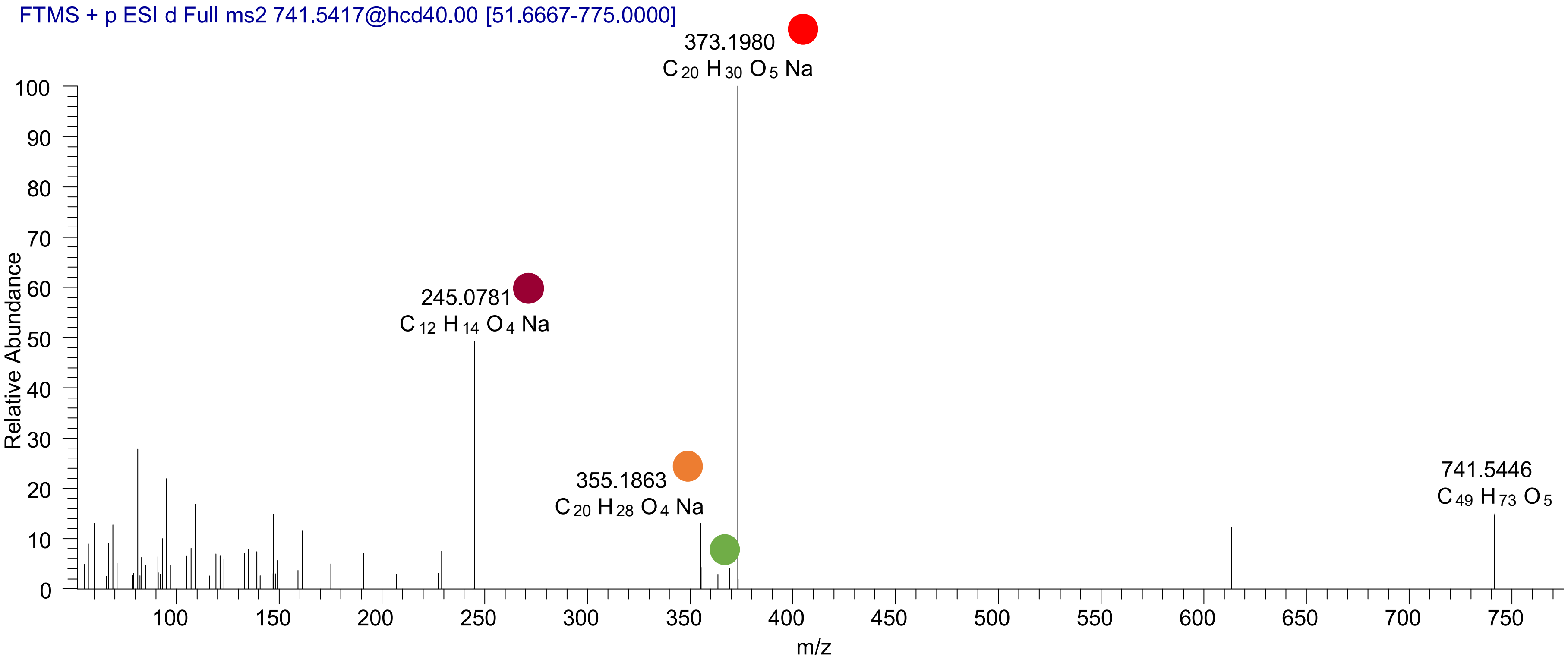

# CE(20:4<2OH,O>)
# RT 19.5

**[oxCE+Na]+** **XIC 743.5585 NL: 1.52E5**

- Fragments containing oxFAs
- Fragments related to water loss
- Fragments not containing oxFAs
- Position-specific fragments
- Fragments related to other oxLPPs

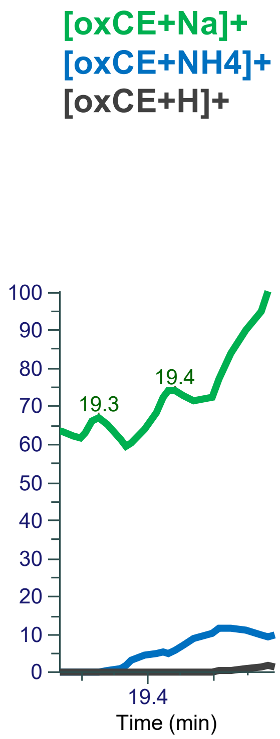


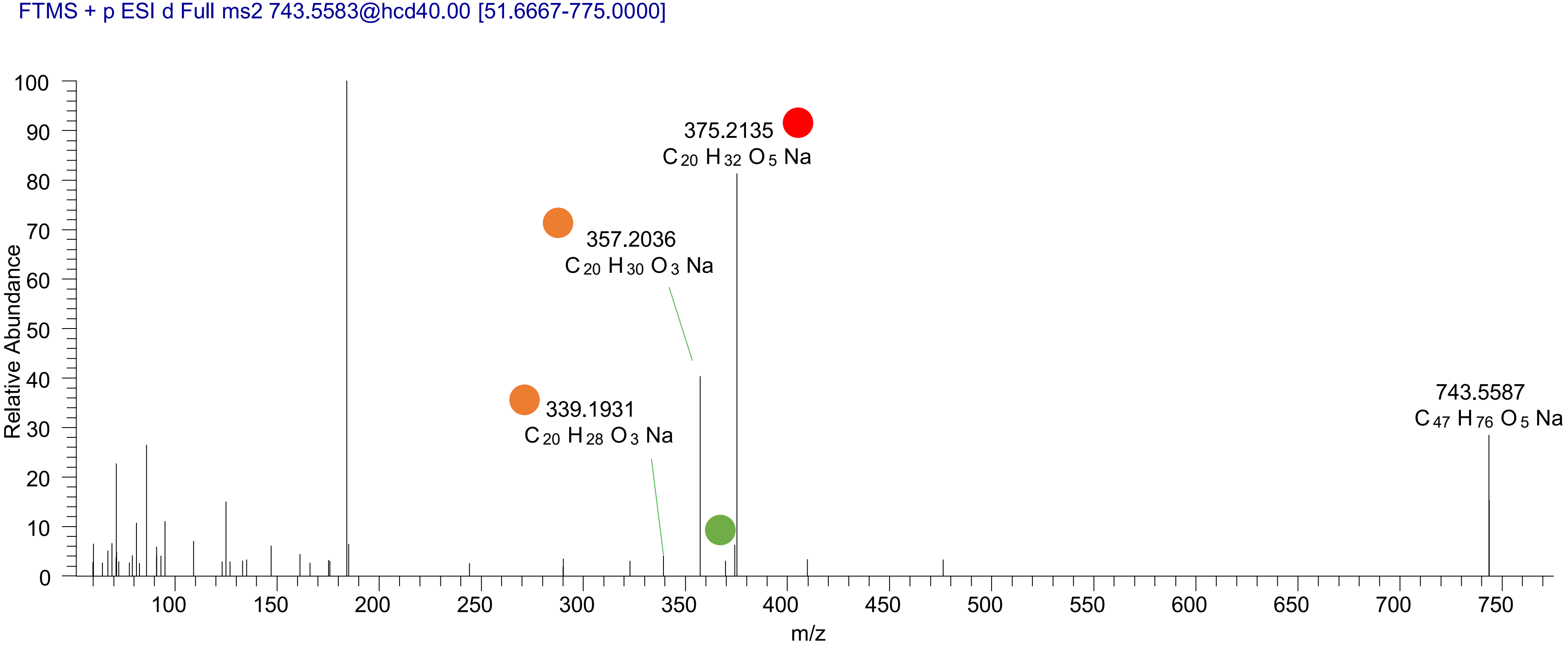

# CE(20:4<OH,2O>)
# RT 20.0

[oxCE+Na]+ XIC 743.5585 NL: 6.33E5

Fragments containing oxFAs
Fragments related to water loss
Fragments not containing oxFAs
Position-specific fragments
Fragments related to other oxLPPs

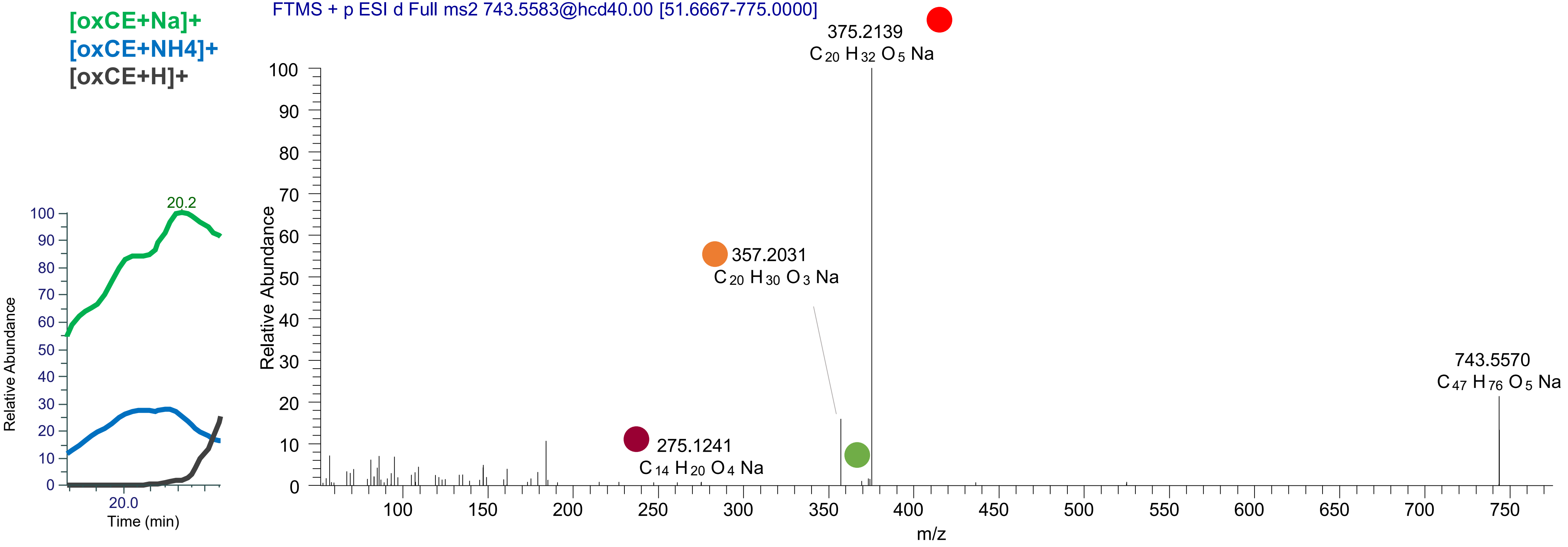

### CE(20:4<OH,OOH{15}>)
# RT 20.2

[oxCE+Na]+ XIC 743.5585 NL: 6.33E5

Fragments containing oxFAs

Fragments related to water loss

Fragments not containing oxFAs

Position-specific fragments

Fragments related to other oxLPPs

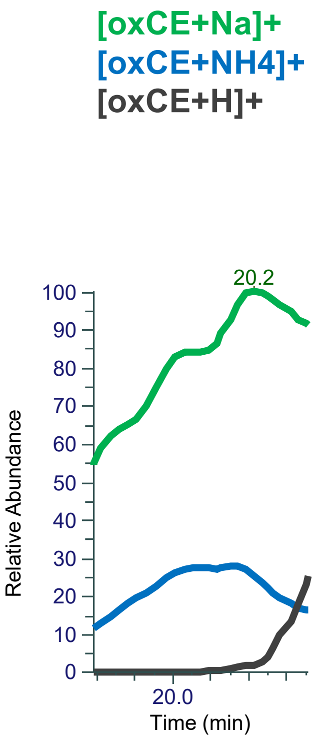


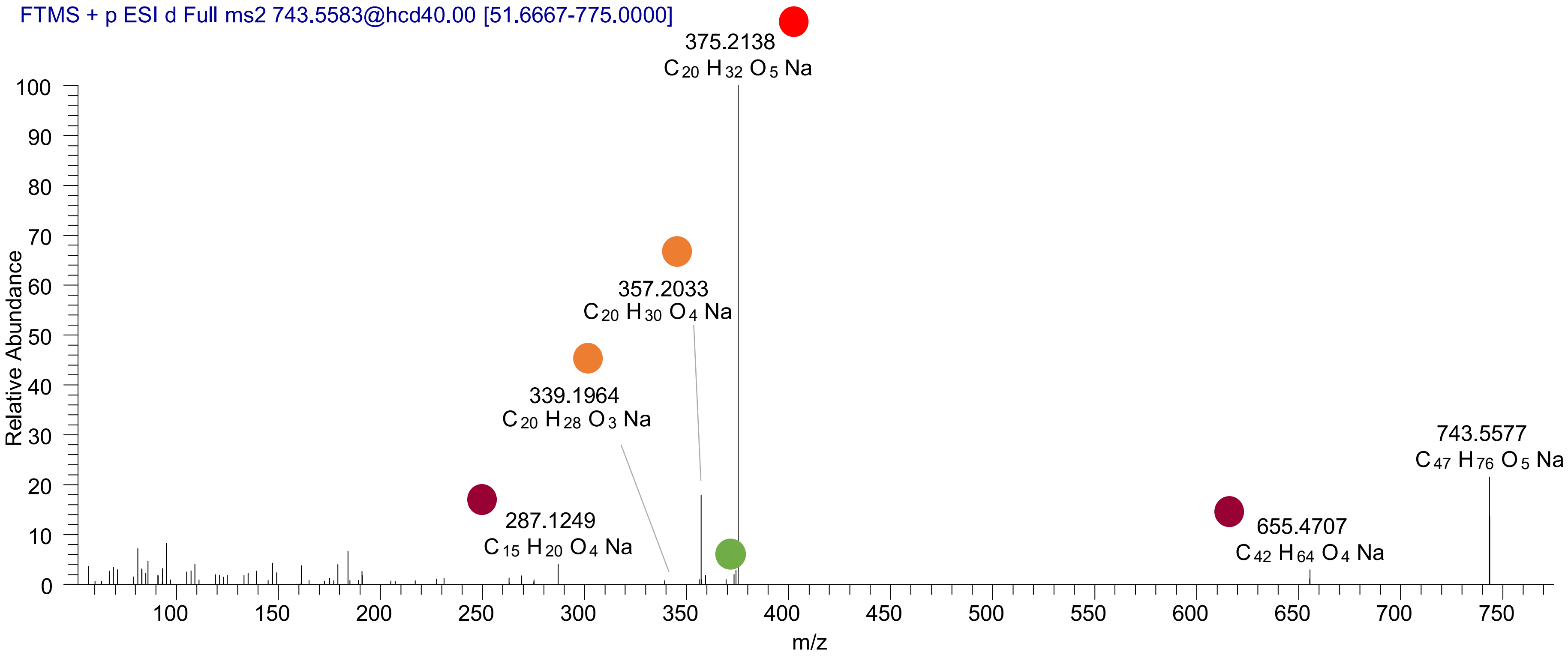

### CE(20:4<OH,OOH{15}>)
### CE(20:4<OH,OOH{12}>)
### CE(20:4<OH,OOH{9}>)
# RT 20.7

**[oxCE+Na]+** **XIC 743.5585 NL: 1.61E6**

- Fragments containing oxFAs
- Fragments related to water loss
- Fragments not containing oxFAs
- Position-specific fragments
- Fragments related to other oxLPPs

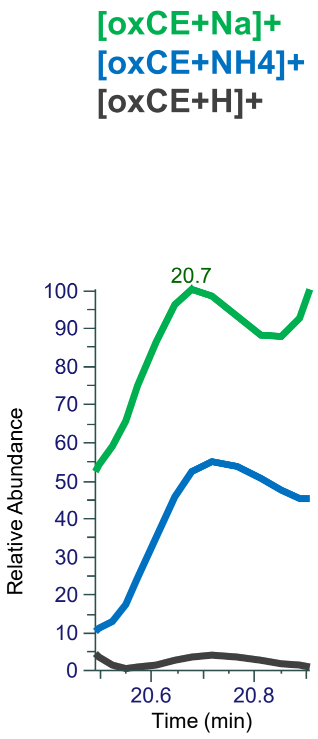


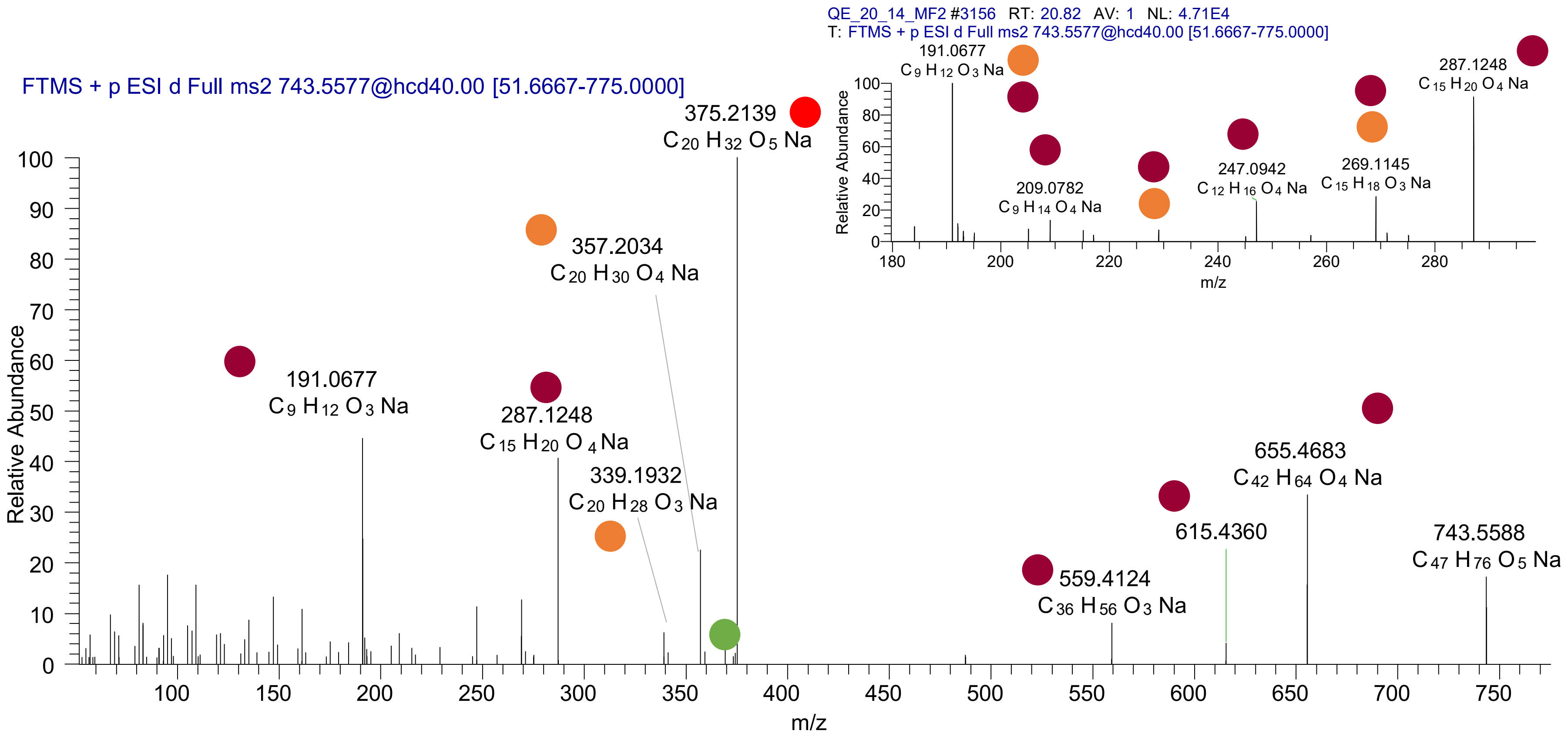

### CE(20:3<ep{11-12},OOH{9}>) RT 21.2

**[oxCE+Na]+  XIC 743.5585 NL: 1.87E6**

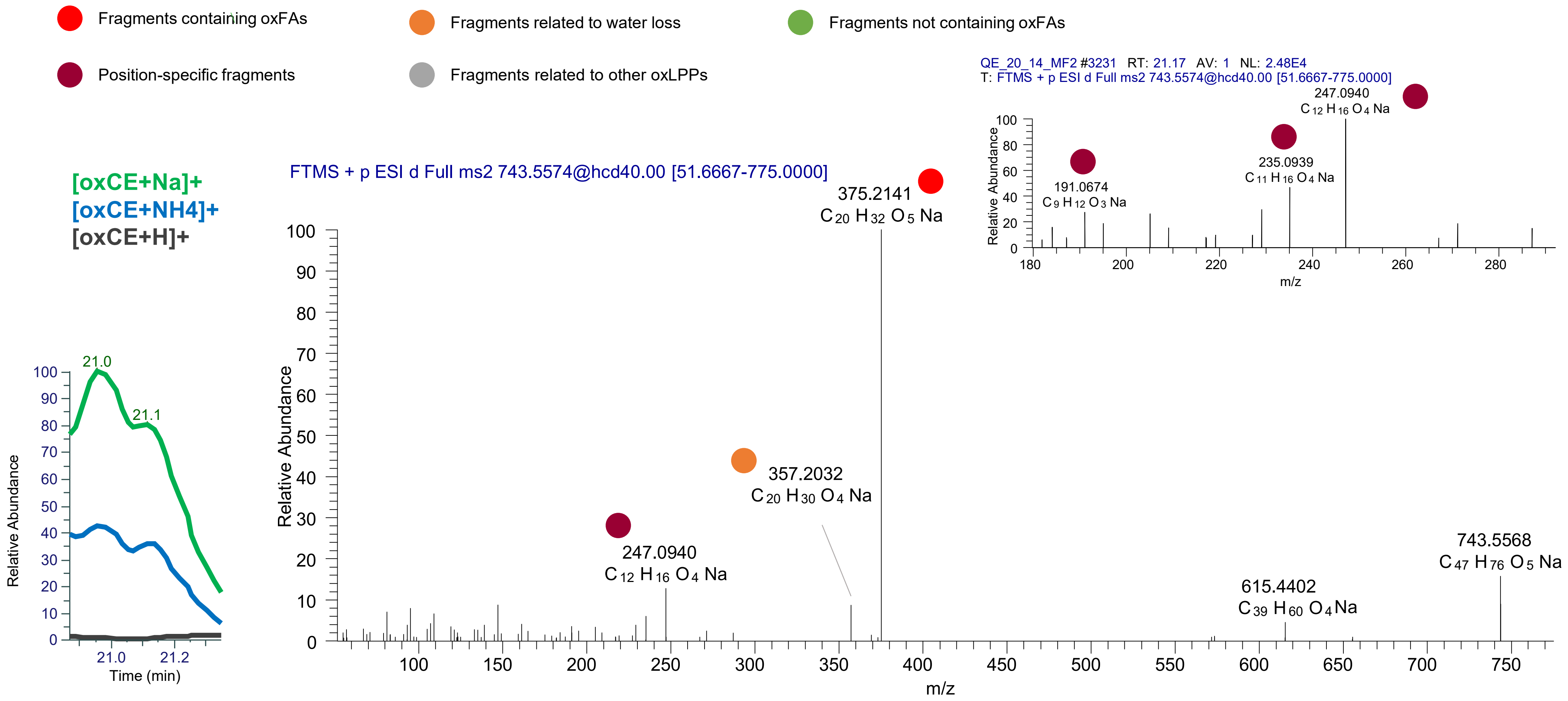

CE(20:3<ep,OOH{16}>)
CE(20:3<ep,OOH{15}>)
CE(20:3<ep,OOH{11}>)
CE(20:3<ep,OOH{9}>)
RT 21.5

**[oxCE+Na]+** **XIC 743.5585 NL: 4.81E5**

- Fragments containing oxFAs
- Fragments related to water loss
- Fragments not containing oxFAs
- Position-specific fragments
- Fragments related to other oxLPPs

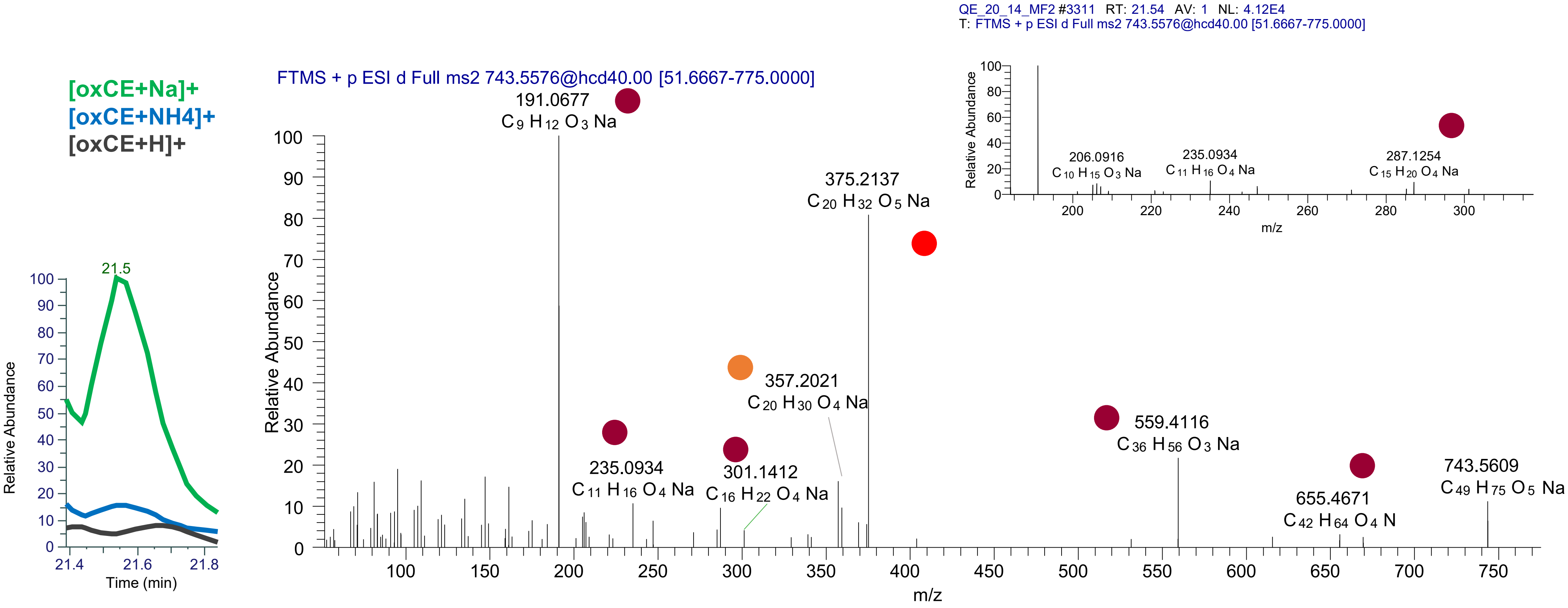

### CE(20:3<oxo,OOH{12}>)
### CE(20:3<oxo,OOH{15}>)
# RT 22.1

[oxCE+Na]+ XIC 743.5585 NL: 1.51E5

- Fragments containing oxFAs
- Fragments related to water loss
- Fragments not containing oxFAs
- Position-specific fragments
- Fragments related to other oxLPPs

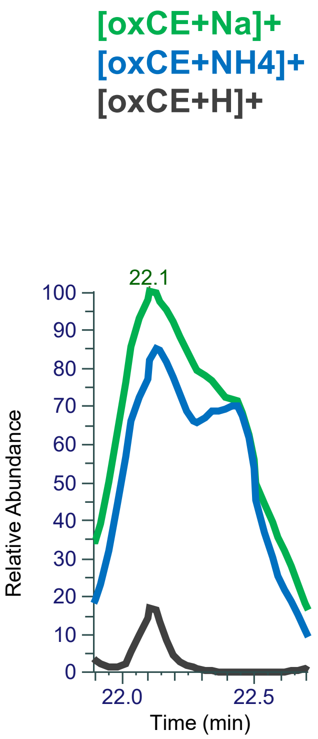


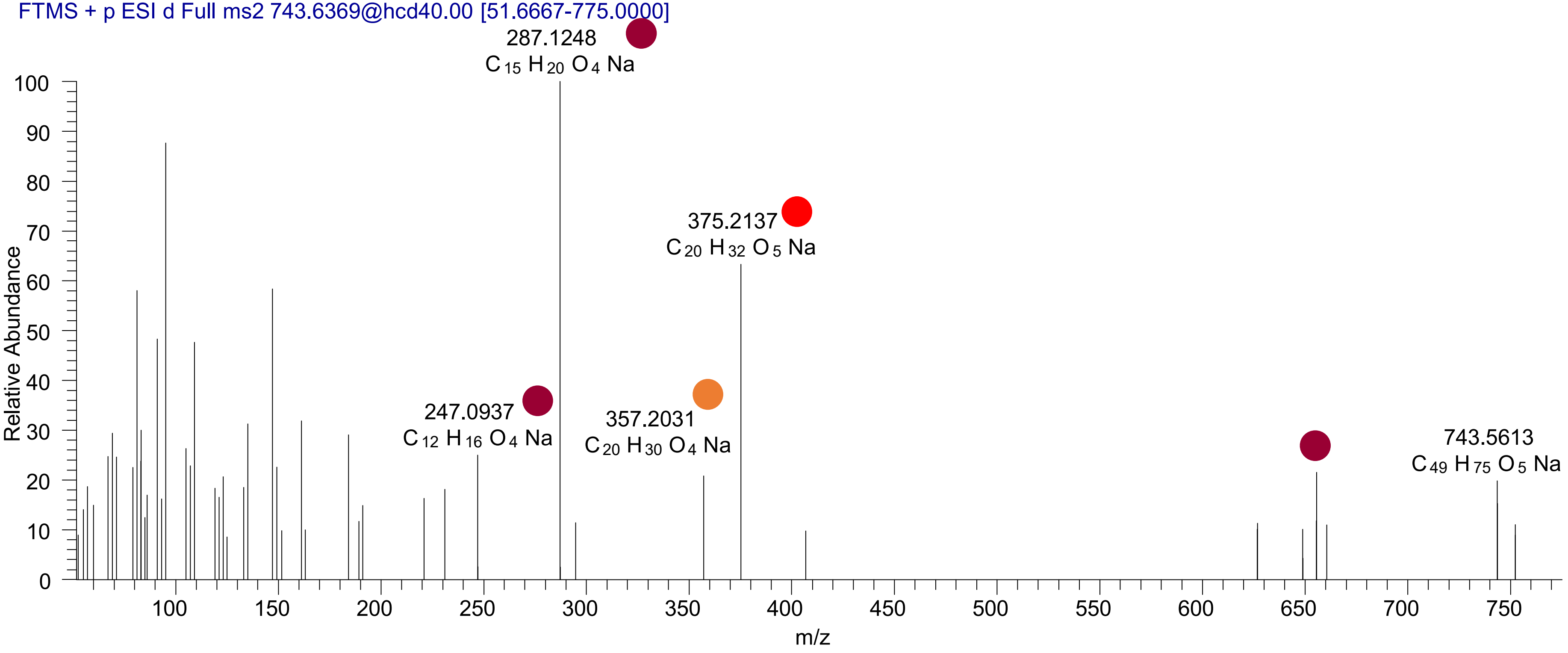

# CE(20:2<ep,2O>)
# RT 20.7

**[oxCE+Na]+** **XIC 745.5741 NL: 3.39E4**

- Fragments containing oxFAs
- Fragments related to water loss
- Fragments not containing oxFAs
- Position-specific fragments
- Fragments related to other oxLPPs

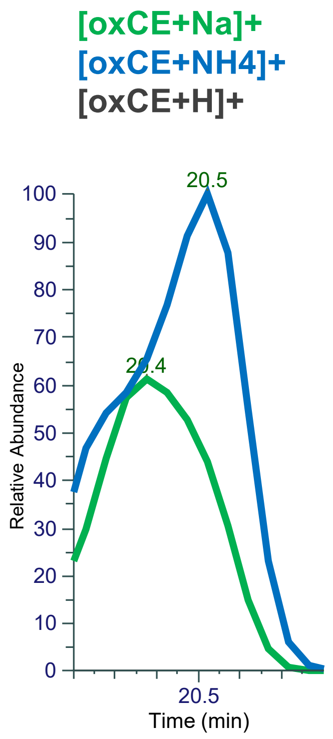


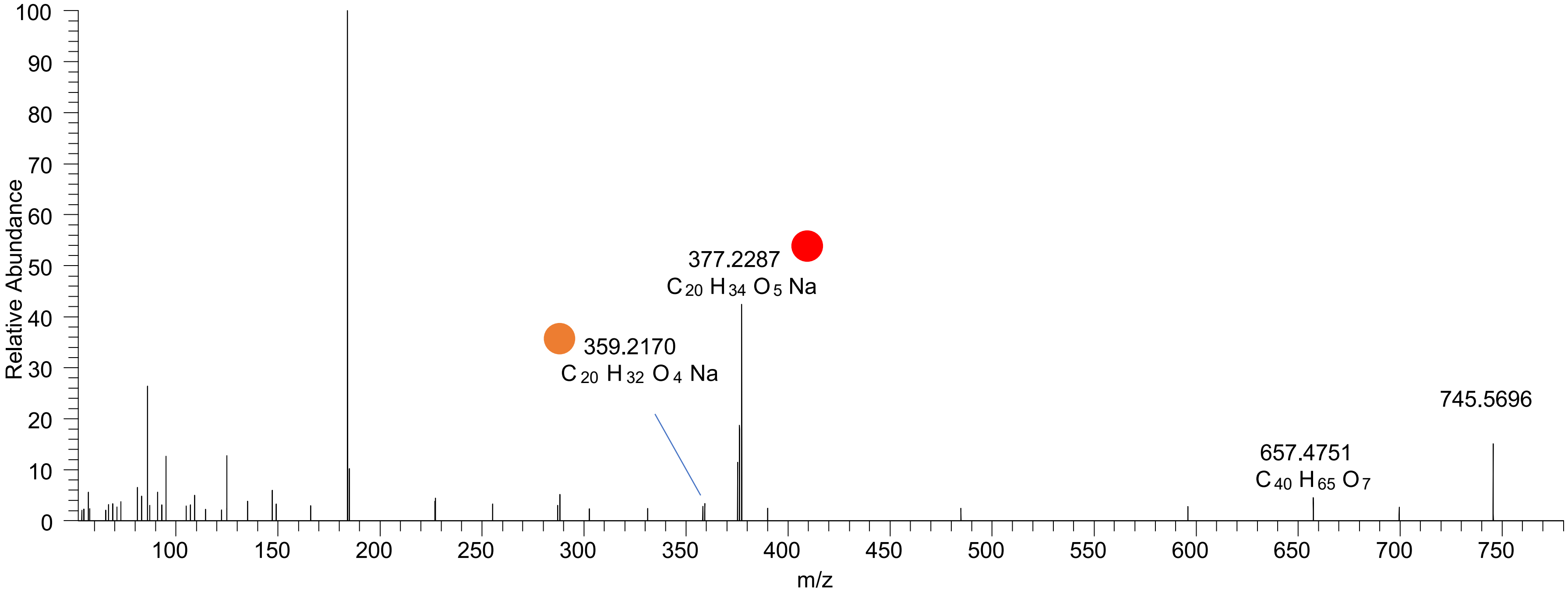

### CE(20:3<OOH{13},O>)
# RT 21.4

**[oxCE+Na]+** **XIC 745.5741 NL: 7.81E4**

- Fragments containing oxFAs
- Fragments related to water loss
- Fragments not containing oxFAs
- Position-specific fragments
- Fragments related to other oxLPPs

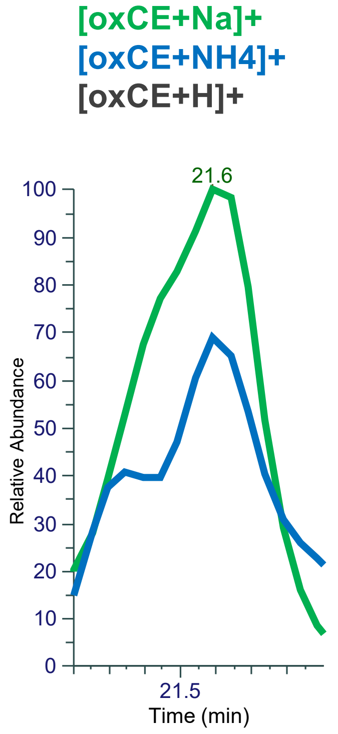


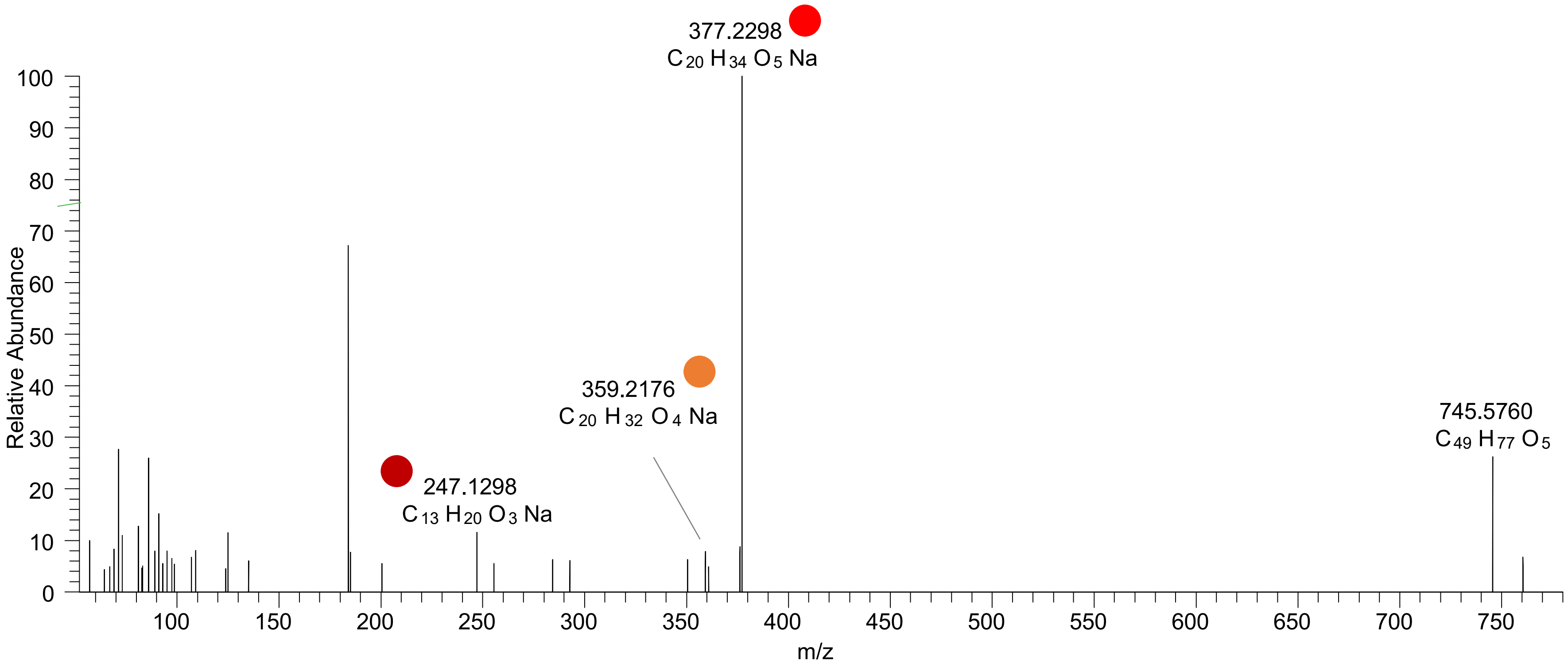

# CE(22:6<2OH>)
# RT 20.7

**[oxCE+Na]+** **XIC 751.5636 NL: 1.87E5**

- Fragments containing oxFAs
- Fragments related to water loss
- Fragments not containing oxFAs
- Position-specific fragments
- Fragments related to other oxLPPs

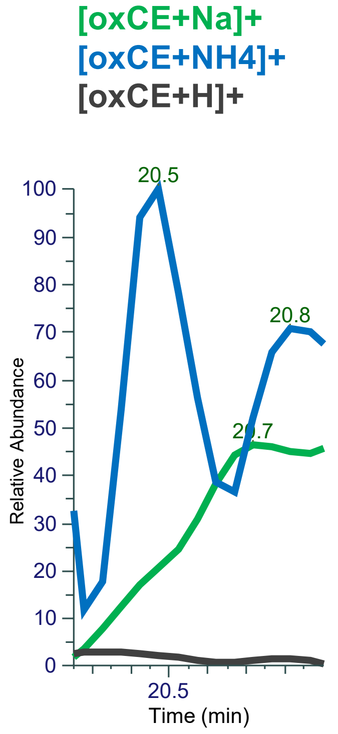


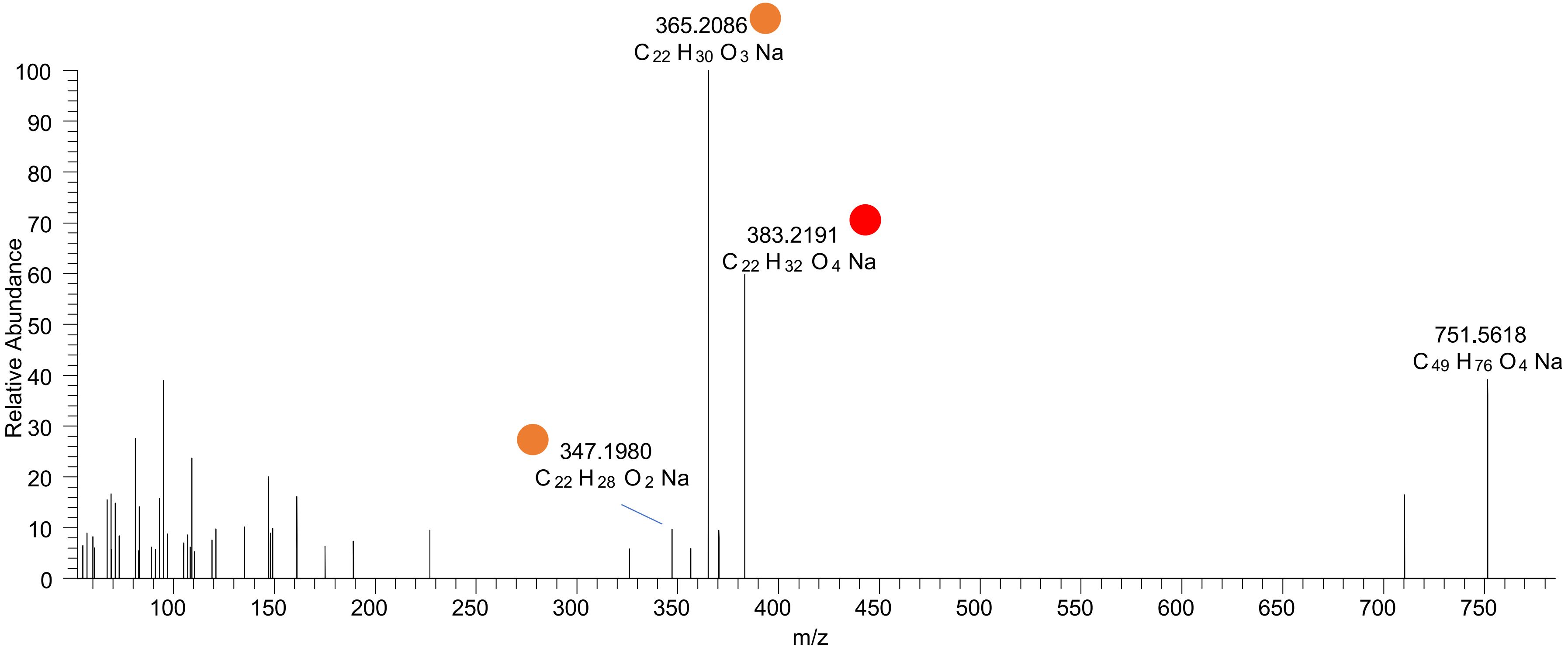

### CE(22:5<oxo,OH>)
# RT 21.5-21.9

**[oxCE+Na]+** **XIC 751.5636 NL: 7.06E4**

Fragments containing oxFAs
Fragments related to water loss
Fragments not containing oxFAs
Position-specific fragments
Fragments related to other oxLPPs

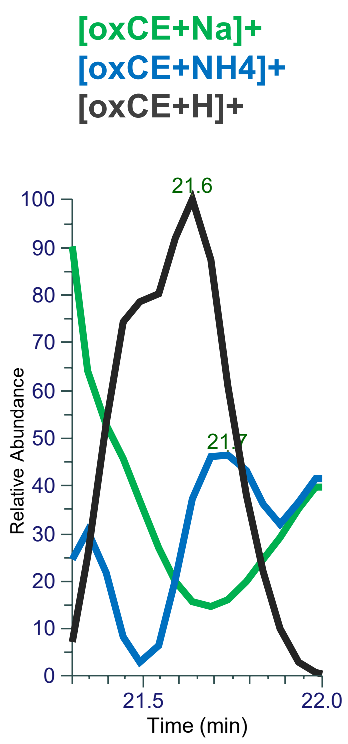


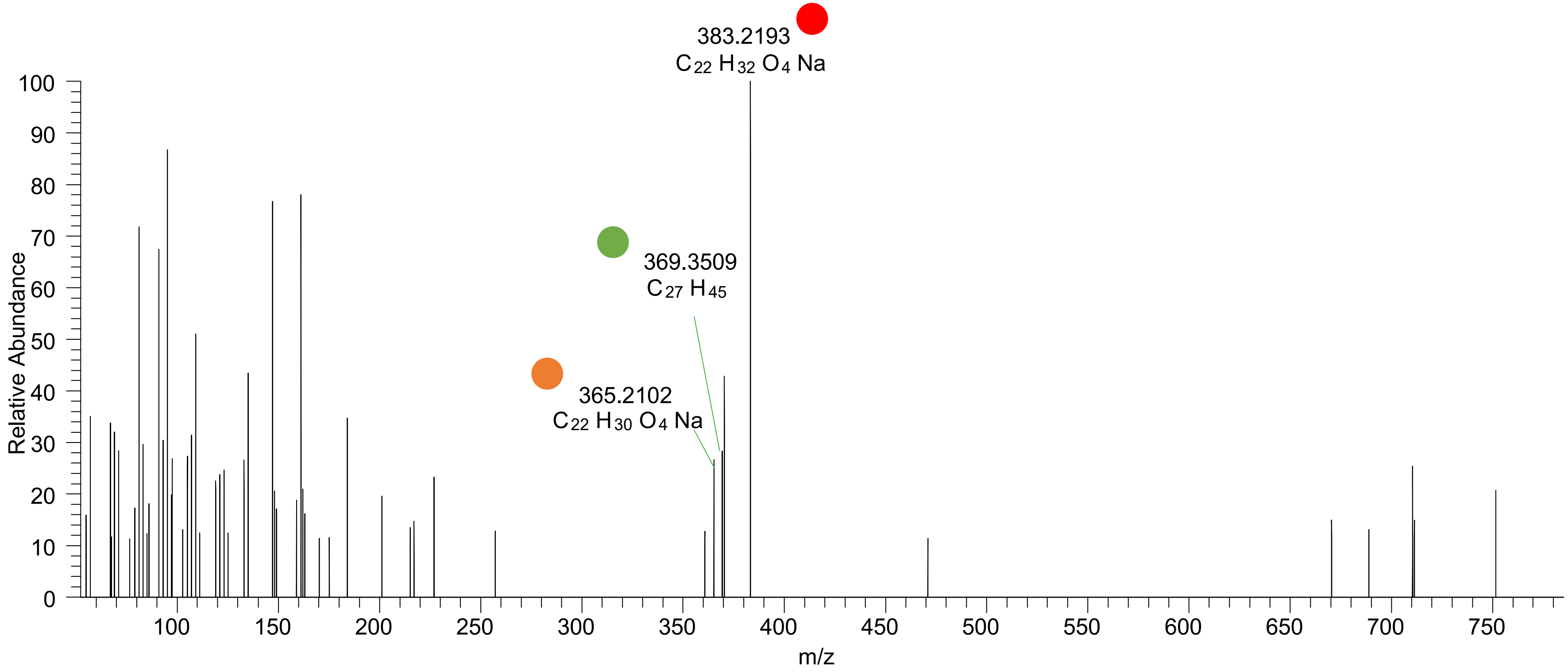

CE(22:6<OOH{8}>)
CE(22:6<OOH{11}>)
CE(22:6<OOH{14}>)
CE(22:6<OOH{17}>)
RT 23.3

**[oxCE+Na]+** **XIC 751.5636 NL: 2.27E5**

Fragments containing oxFAs
Fragments related to water loss
Fragments not containing oxFAs
Position-specific fragments
Fragments related to other oxLPPs

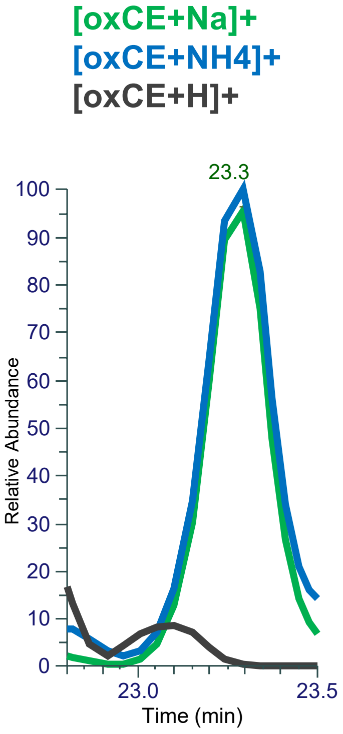


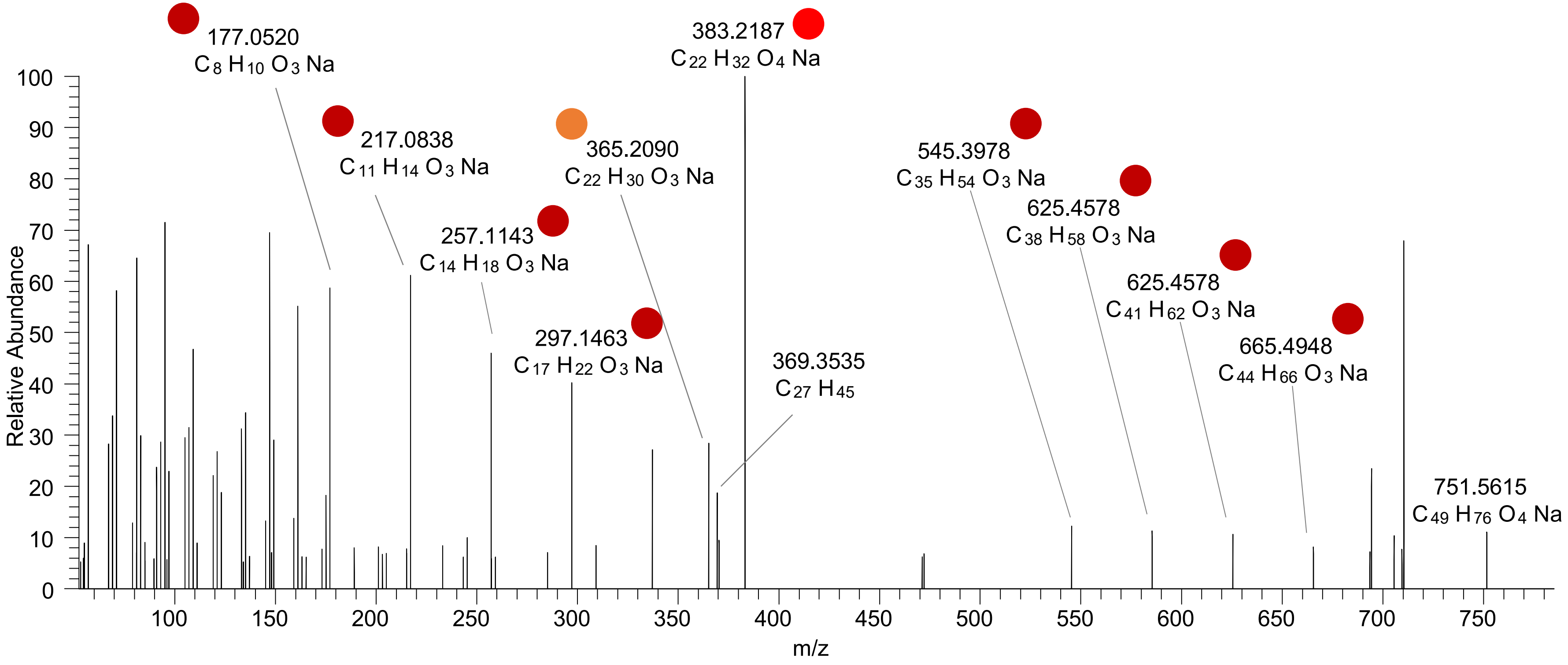

# CE(22:4<OH,ep>)
# RT 21.4

**[oxCE+Na]$^+$** **XIC 753.5792 NL: 3.91E4**

- Fragments containing oxFAs
- Fragments related to water loss
- Fragments not containing oxFAs
- Position-specific fragments
- Fragments related to other oxLPPs

[oxCE+Na]+
[oxCE+NH4]+
[oxCE+H]+

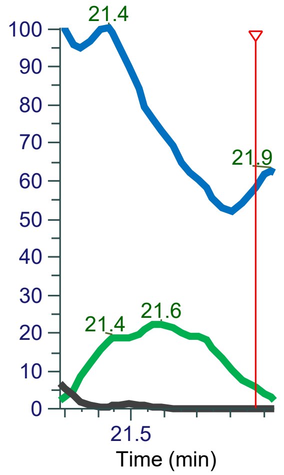


FTMS + p ESI d Full ms2 753.5786@hcd40.00 [52.3333-785.0000]

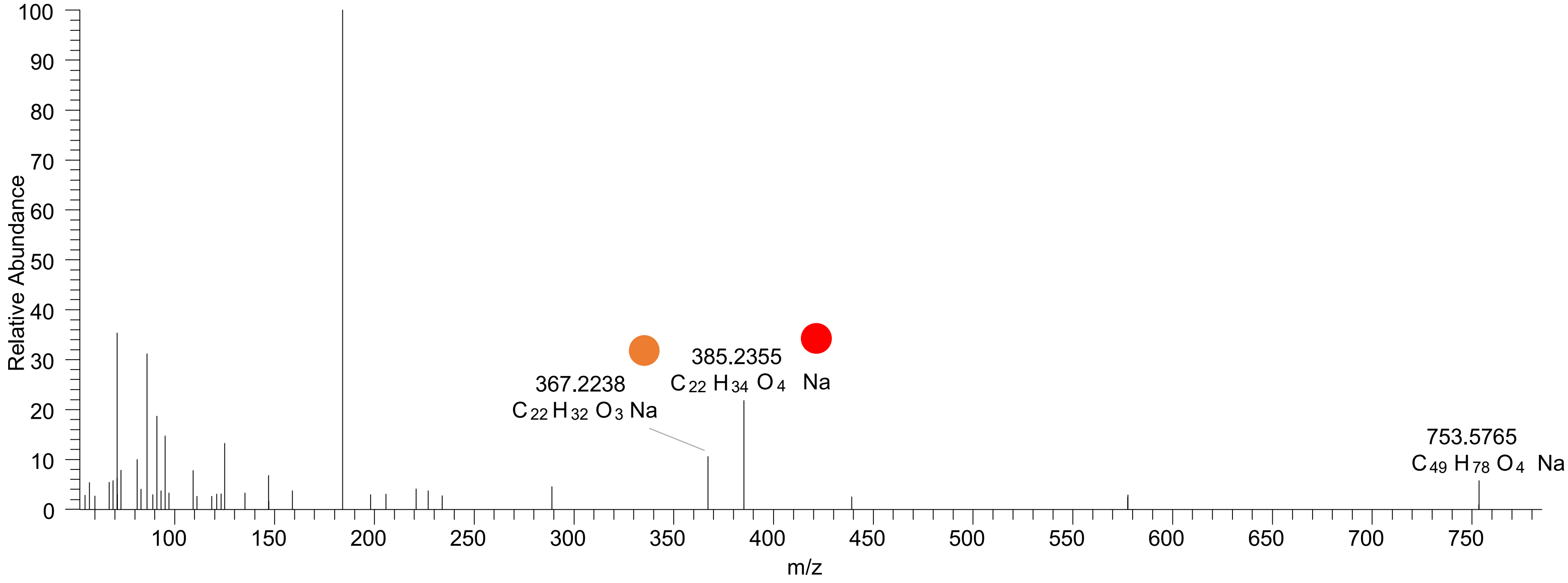

### OT2D_pool

### CE(5:0<oxo{5}>)
# RT 18.4

**[oxCE+Na]+** **XIC 507.3809 NL: 1.36E5**

- Fragments containing oxFAs
- Fragments related to water loss
- Fragments not containing oxFAs
- Position-specific fragments
- Fragments related to other oxLPPs

**[oxCE+Na]+**
**[oxCE+NH4]+**
**[oxCE+H]+**

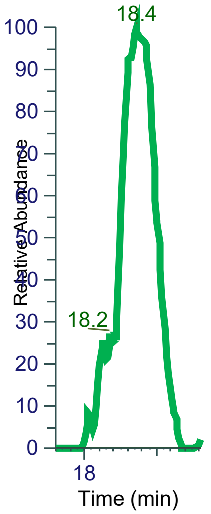


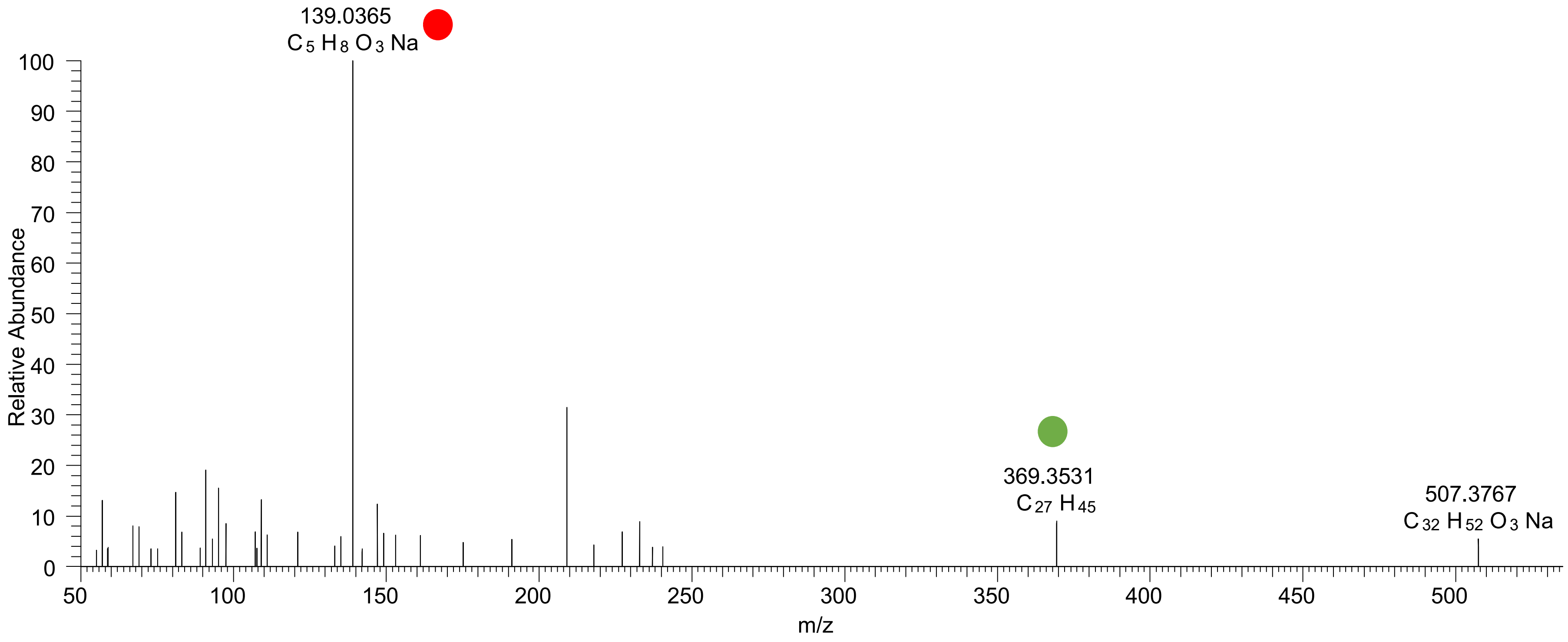

### CE(9:0<oxo{9}>)
# RT 20.9

**[oxCE+Na]+** **XIC 563.4435 NL: 8.52E5**

Fragments containing oxFAs
Fragments related to water loss
Fragments not containing oxFAs
Position-specific fragments
Fragments related to other oxLPPs

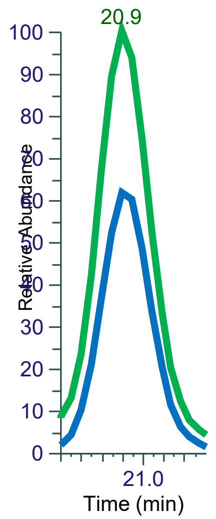


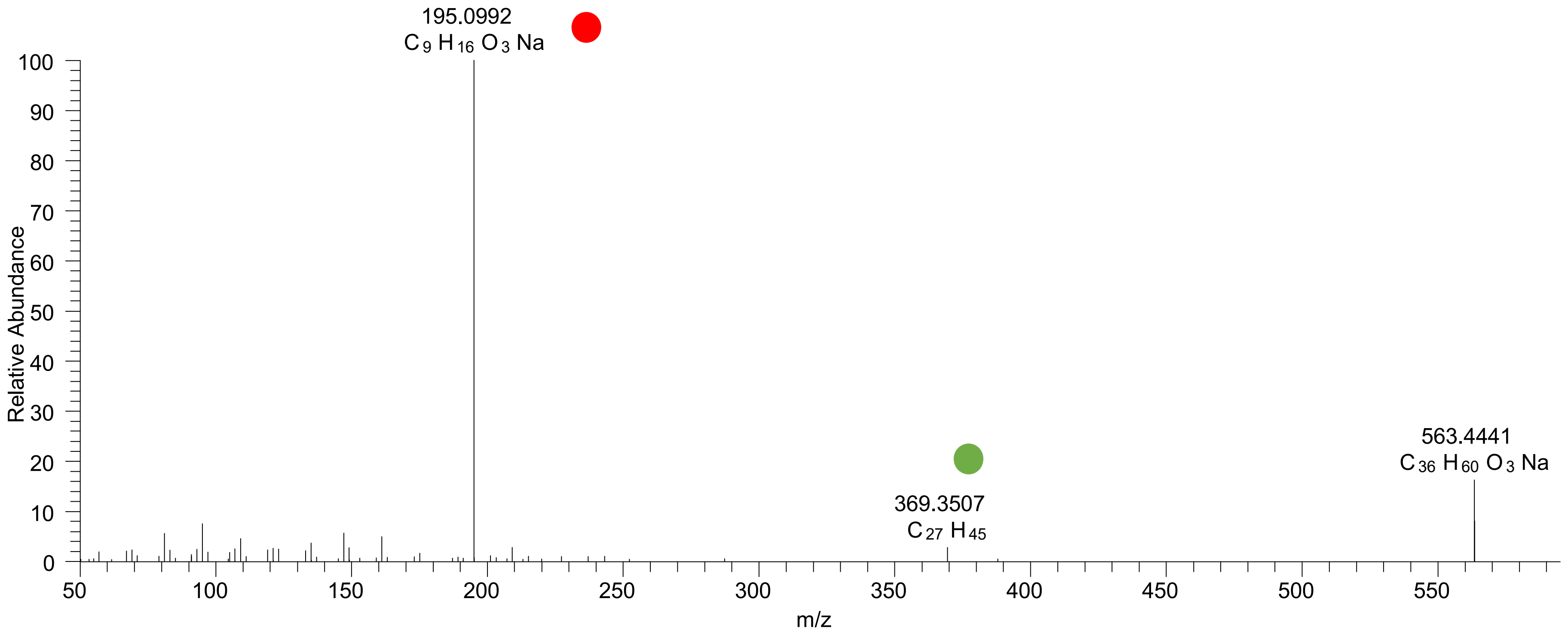

### CE(8:0<COOH{8}>)
# RT 22.1

**[oxCE+Na]+** **XIC 565.4227 NL: 2.69E4**

Fragments containing oxFAs

Fragments related to water loss

Fragments not containing oxFAs

Position-specific fragments

Fragments related to other oxLPPs

**[oxCE+Na]+**
**[oxCE+NH4]+**
**[oxCE+H]+**

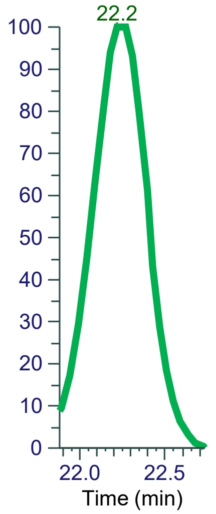


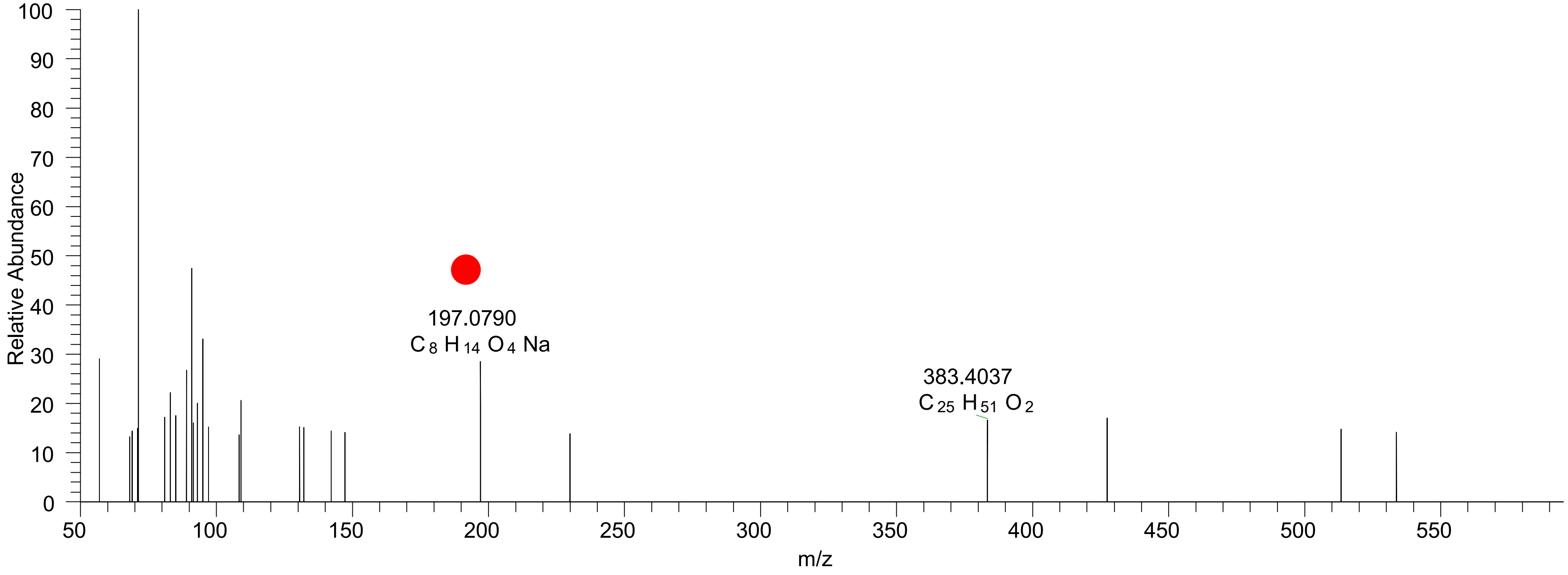

### CE(10:2<oxo{10}>)
# RT 20.8

**[oxCE+Na]+** **XIC 573.4278 NL: 2.19E5**

Fragments containing oxFAs
Fragments related to water loss
Fragments not containing oxFAs
Position-specific fragments
Fragments related to other oxLPPs

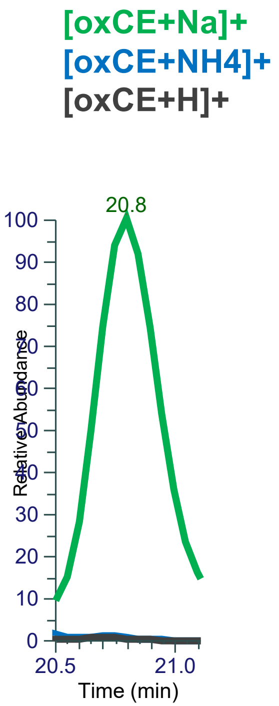


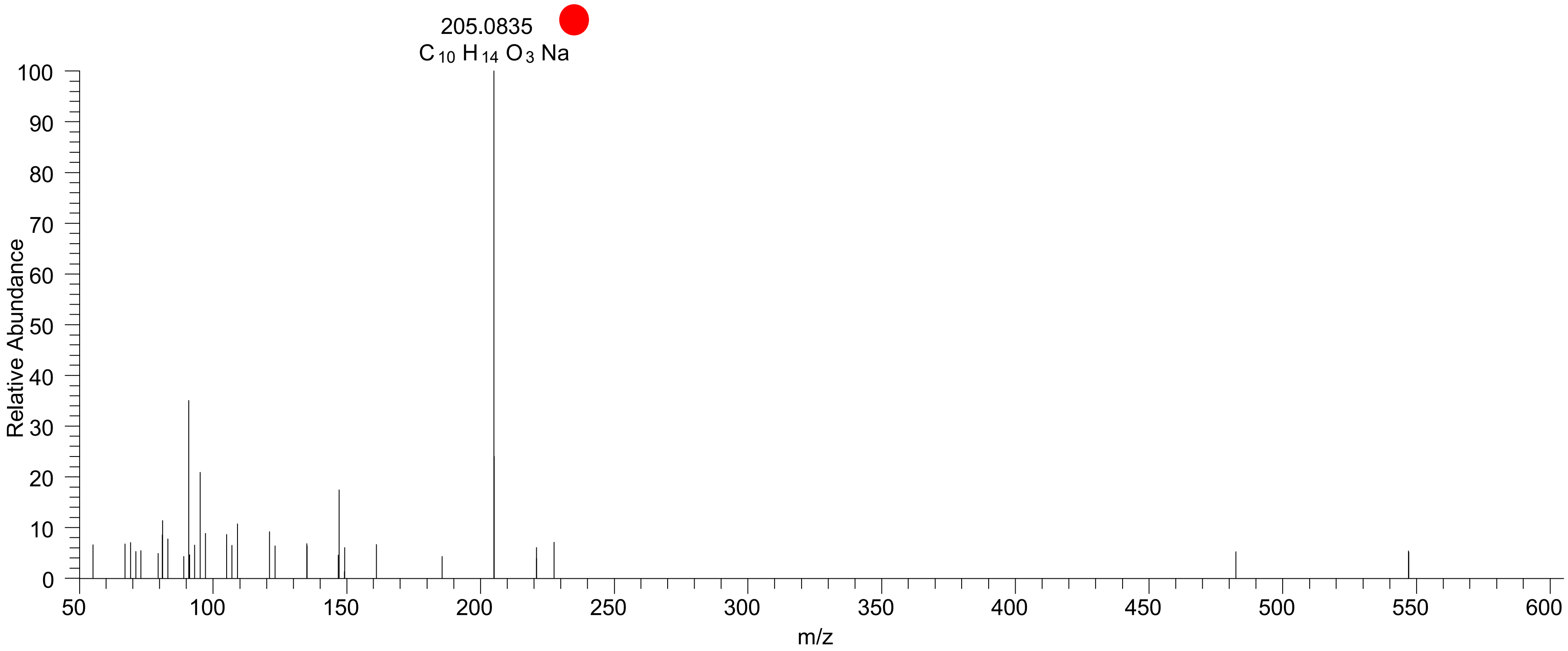

### CE(11:2<oxo{11}>)
# RT 20.3

[oxCE+Na]+ XIC 587.4435 NL: 2.90E4

Fragments containing oxFAs
Fragments related to water loss
Fragments not containing oxFAs
Position-specific fragments
Fragments related to other oxLPPs

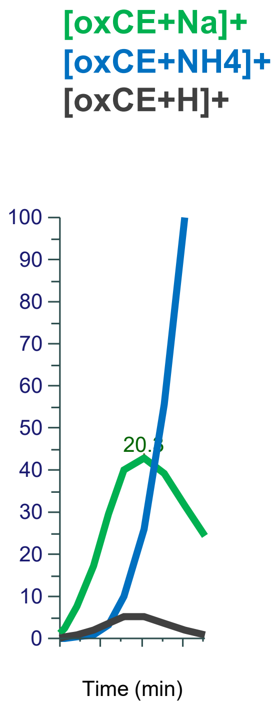


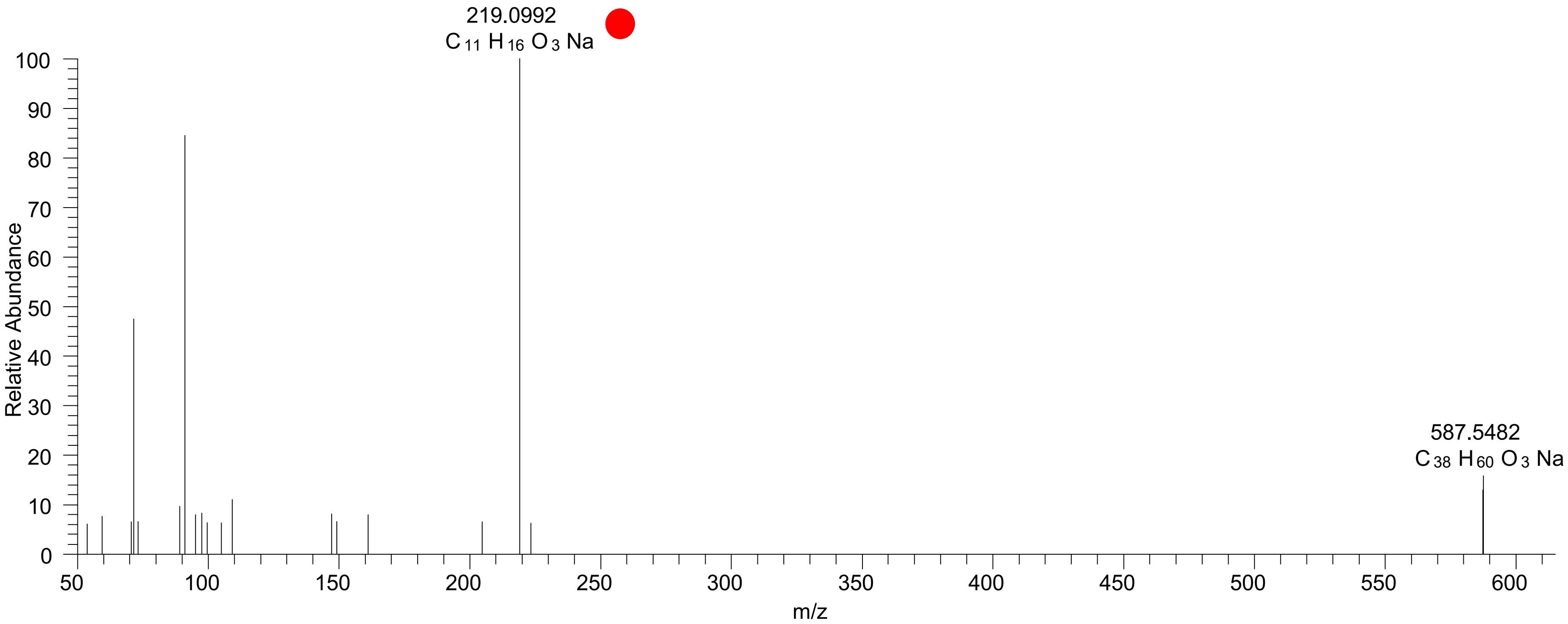

### CE(11:3<COOH{11}>)
# RT 21.1

**[oxCE+Na]+** **XIC 601.4227 NL: 2.04E4**

- Fragments containing oxFAs
- Fragments related to water loss
- Fragments not containing oxFAs
- Position-specific fragments
- Fragments related to other oxLPPs

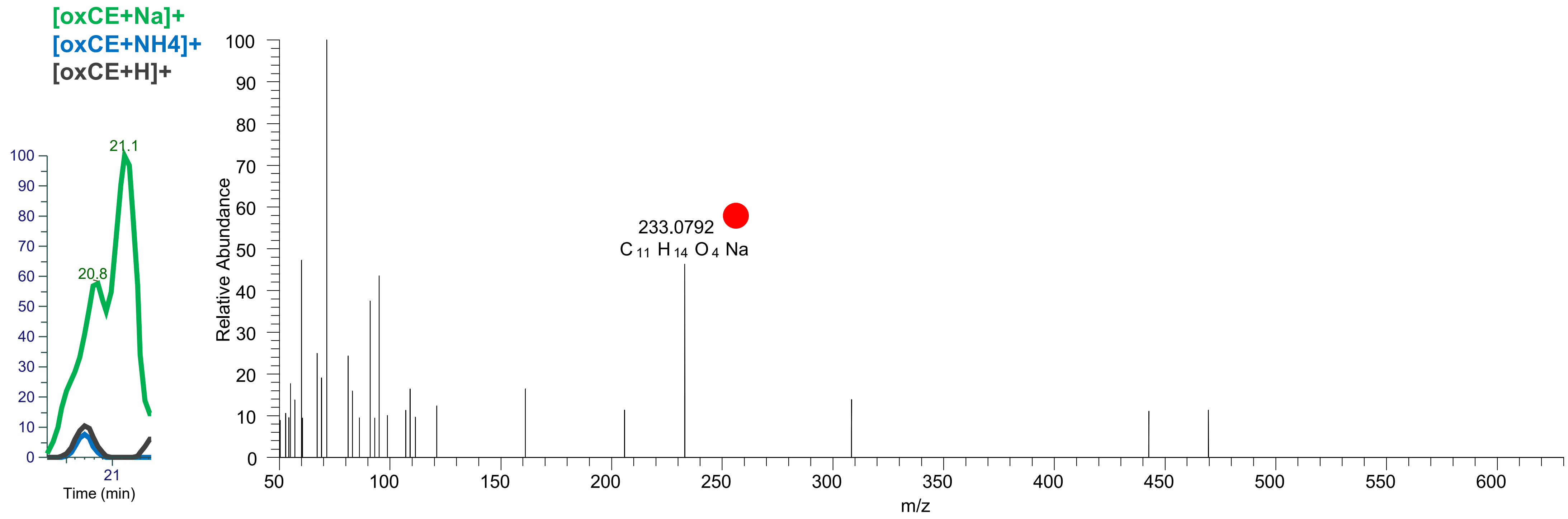

### CE(12:1<oxo{12}>)
# RT 22.1

[oxCE+Na]+ XIC 603.4753 NL: 1.98E4

Fragments containing oxFAs
Fragments related to water loss
Fragments not containing oxFAs
Position-specific fragments
Fragments related to other oxLPPs

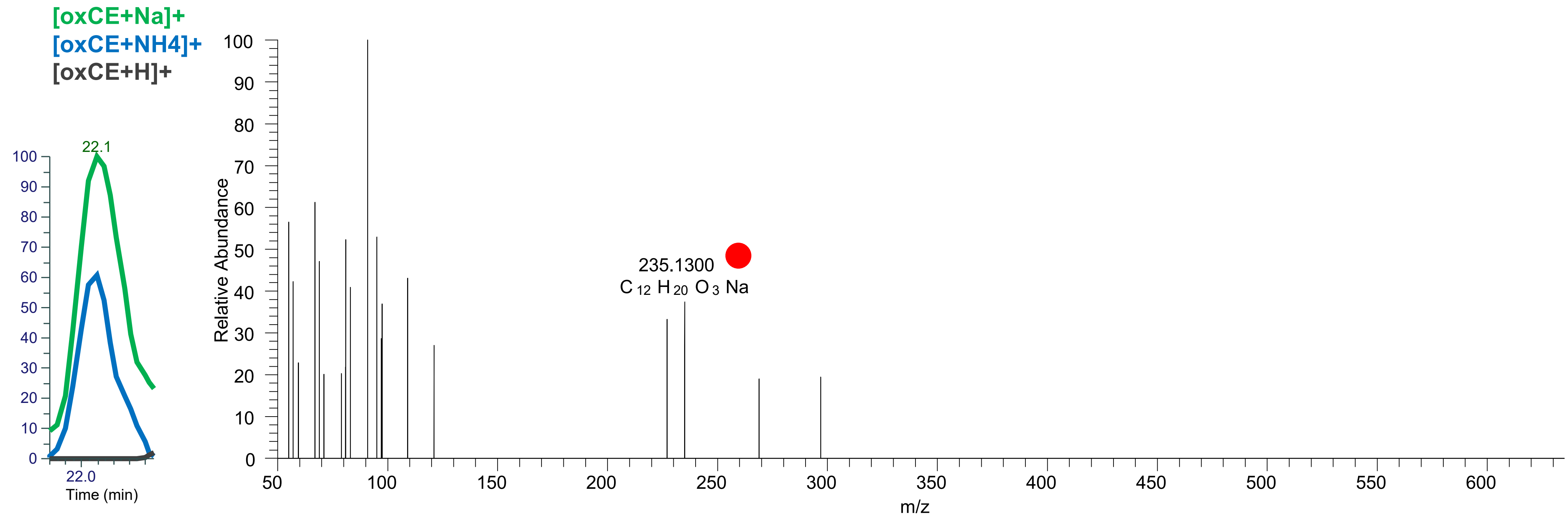

### CE(12:2<COOH{12}>)
# RT 20.5

**[oxCE+Na]+** **XIC 617.4541 NL: 1.26E4**

- Fragments containing oxFAs
- Fragments related to water loss
- Fragments not containing oxFAs
- Position-specific fragments
- Fragments related to other oxLPPs

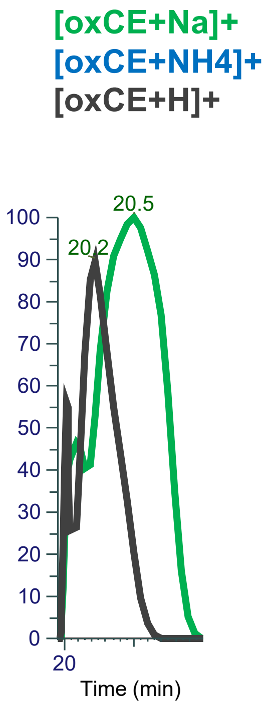


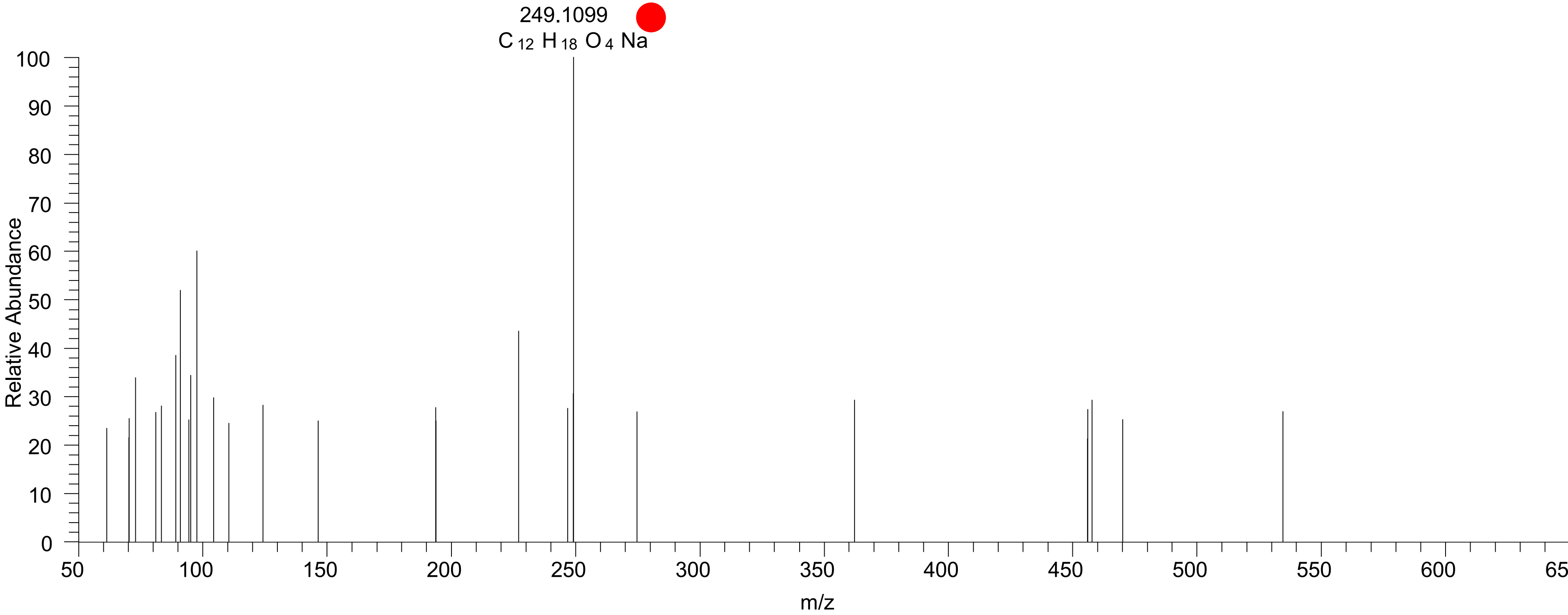

### CE(12:1<COOH{12}>)
# RT 20.3

**[oxCE+Na]+** **XIC 619.4697 NL: 3.73E4**

Fragments containing oxFAs
Fragments related to water loss
Fragments not containing oxFAs
Position-specific fragments
Fragments related to other oxLPPs

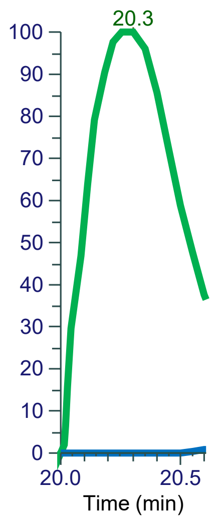


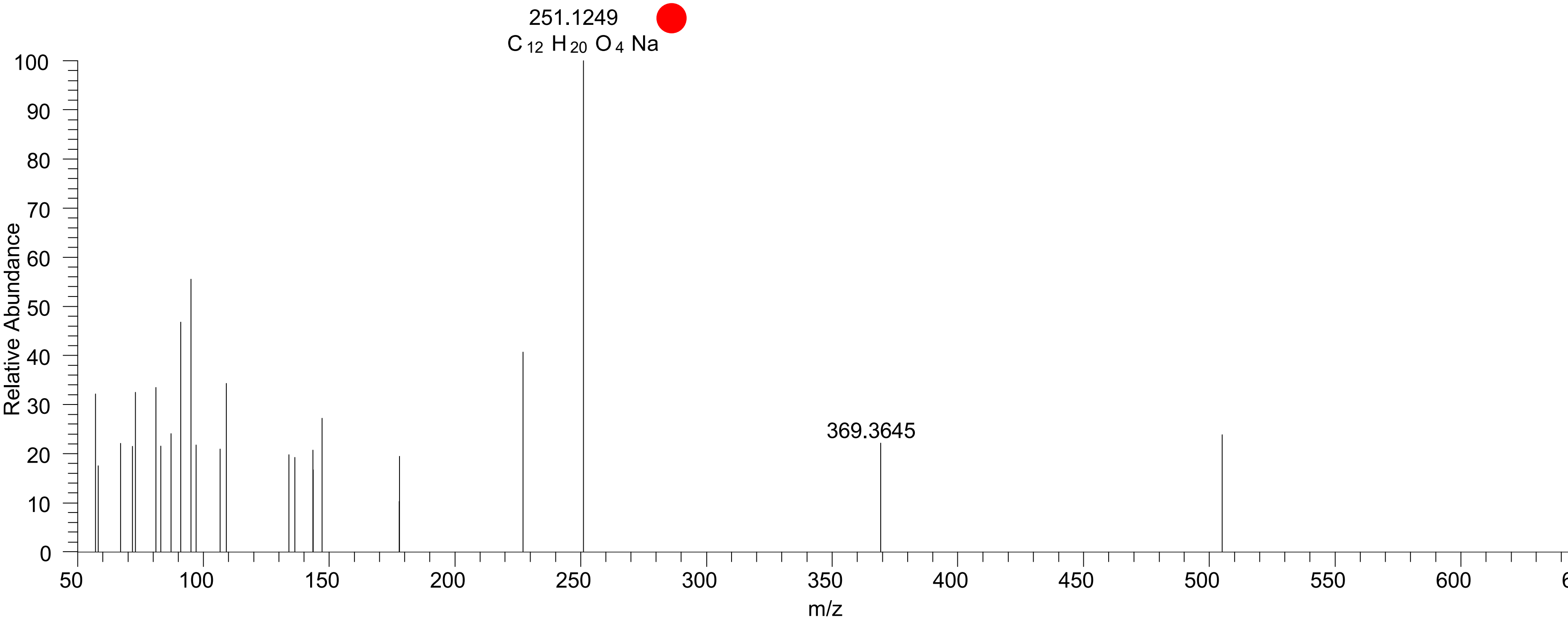

# CE(18:3<OH>)
# RT 23.3

**[oxCE+Na]+** **XIC 685.5531 NL: 1.51E6**

- Fragments containing oxFAs
- Fragments related to water loss
- Fragments not containing oxFAs
- Position-specific fragments
- Fragments related to other oxLPPs

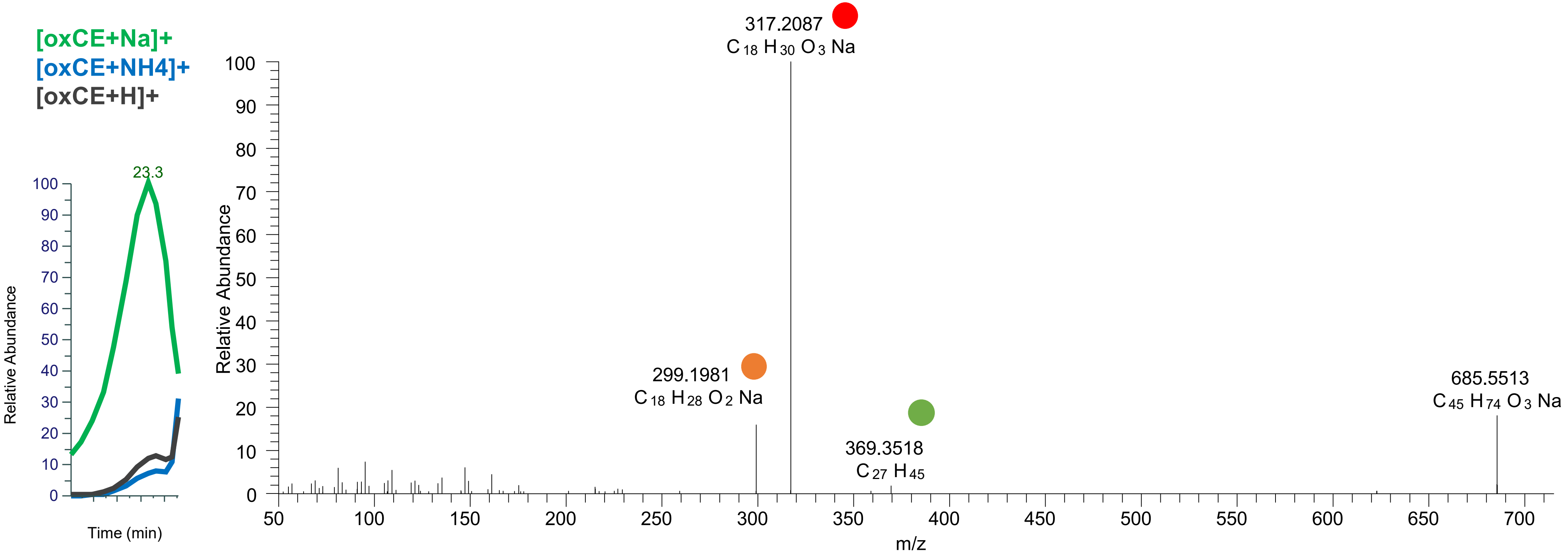

# CE(18:2<ep>)
# RT 23.9

[oxCE+Na]+ XIC 685.5531 NL: 8.84E5

Fragments containing oxFAs
Fragments related to water loss
Fragments not containing oxFAs
Position-specific fragments
Fragments related to other oxLPPs

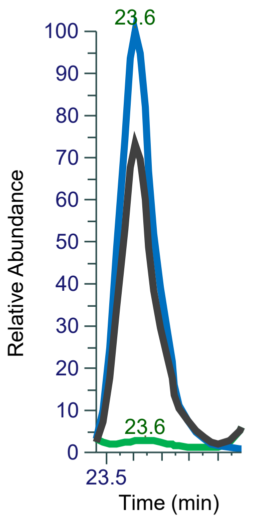


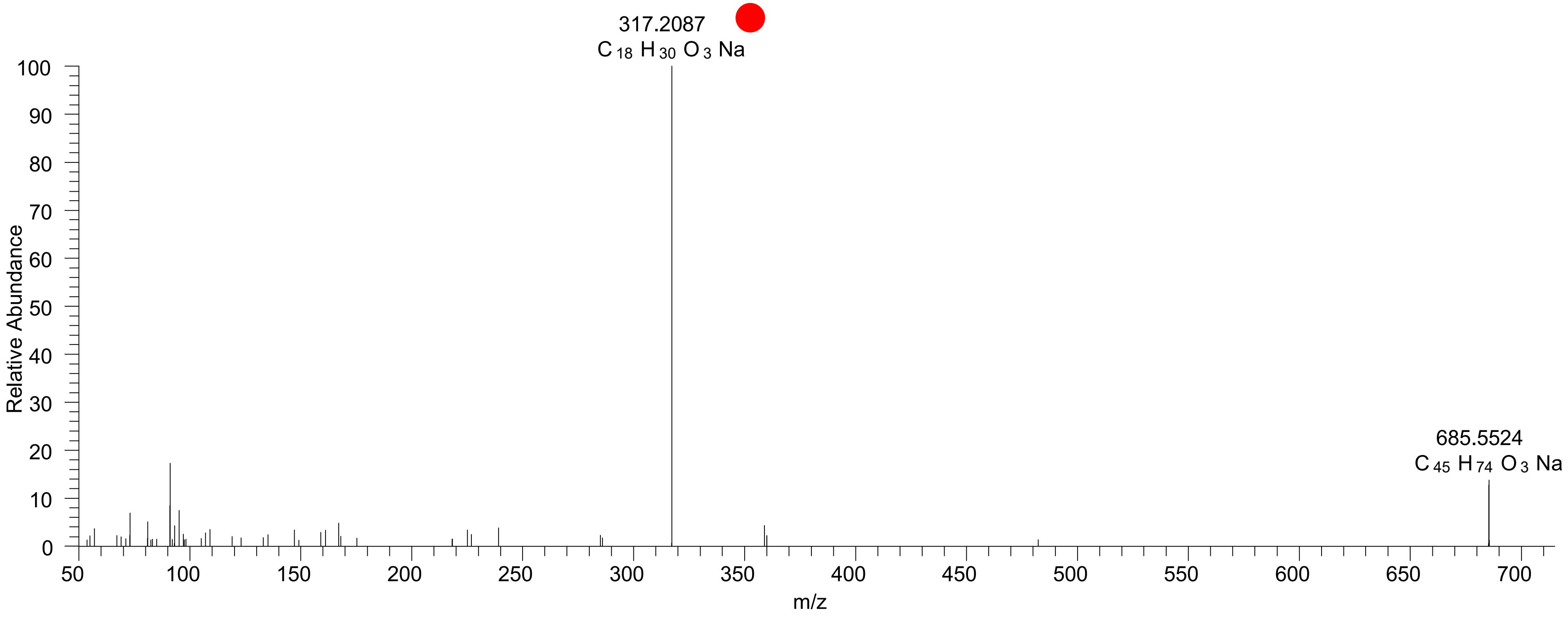

### CE(18:2<oxo>)
# RT 24.2

**[oxCE+Na]+** **XIC 685.5531 NL: 5.99E6**

- Fragments containing oxFAs
- Fragments related to water loss
- Fragments not containing oxFAs
- Position-specific fragments
- Fragments related to other oxLPPs

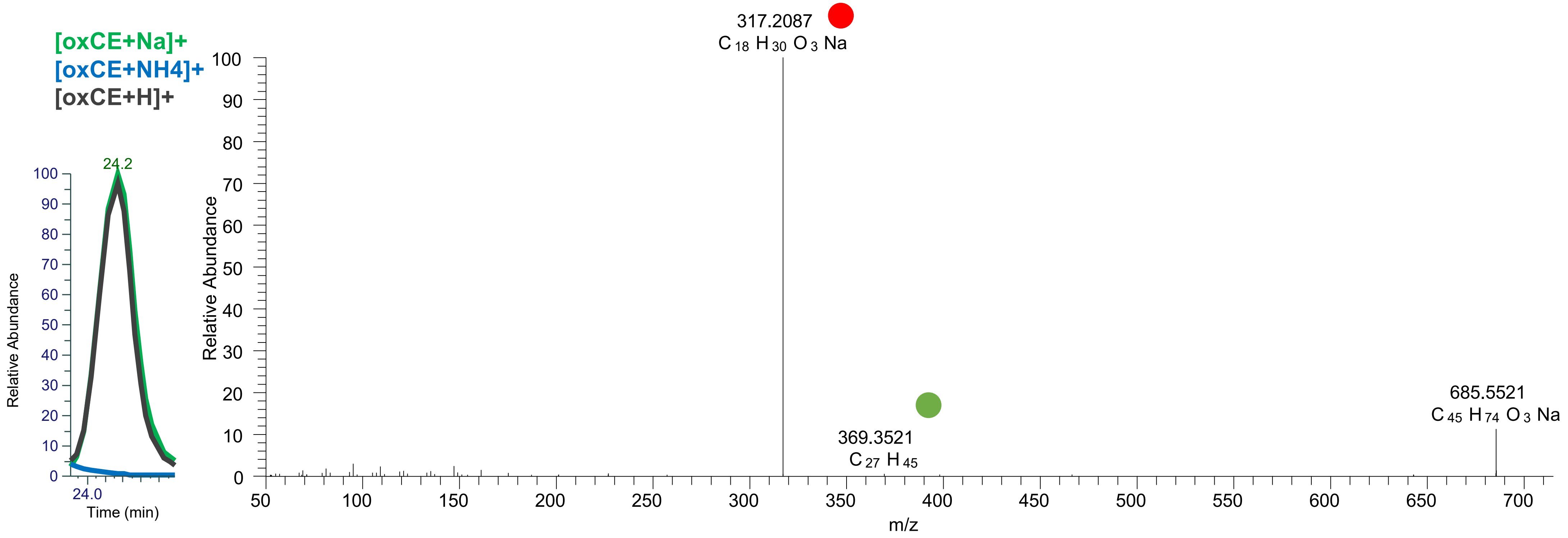

# CE(18:2<OH>)
# RT 23.5

**[oxCE+Na]+** **XIC 687.5687 NL: 4.00E6**

- Fragments containing oxFAs
- Fragments related to water loss
- Fragments not containing oxFAs
- Position-specific fragments
- Fragments related to other oxLPPs

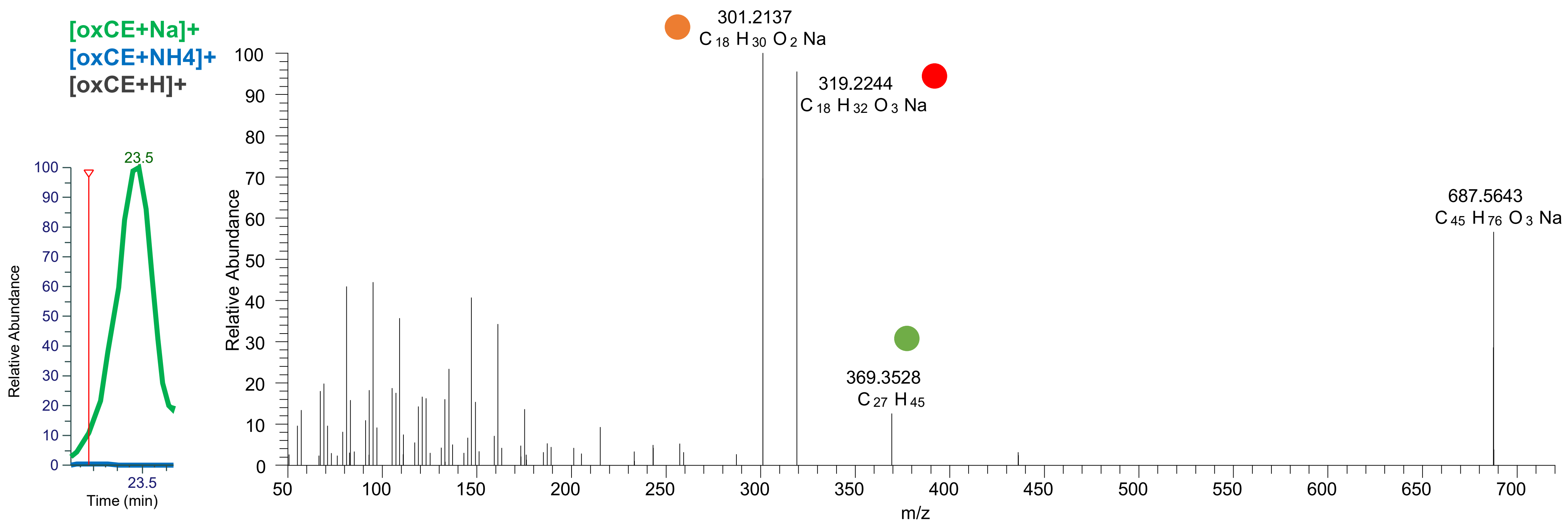

# CE(18:2<OH>)
# RT 23.8

**[oxCE+Na]+** **XIC 687.5687 NL: 1.62E7**

- Fragments containing oxFAs
- Fragments related to water loss
- Fragments not containing oxFAs
- Position-specific fragments
- Fragments related to other oxLPPs

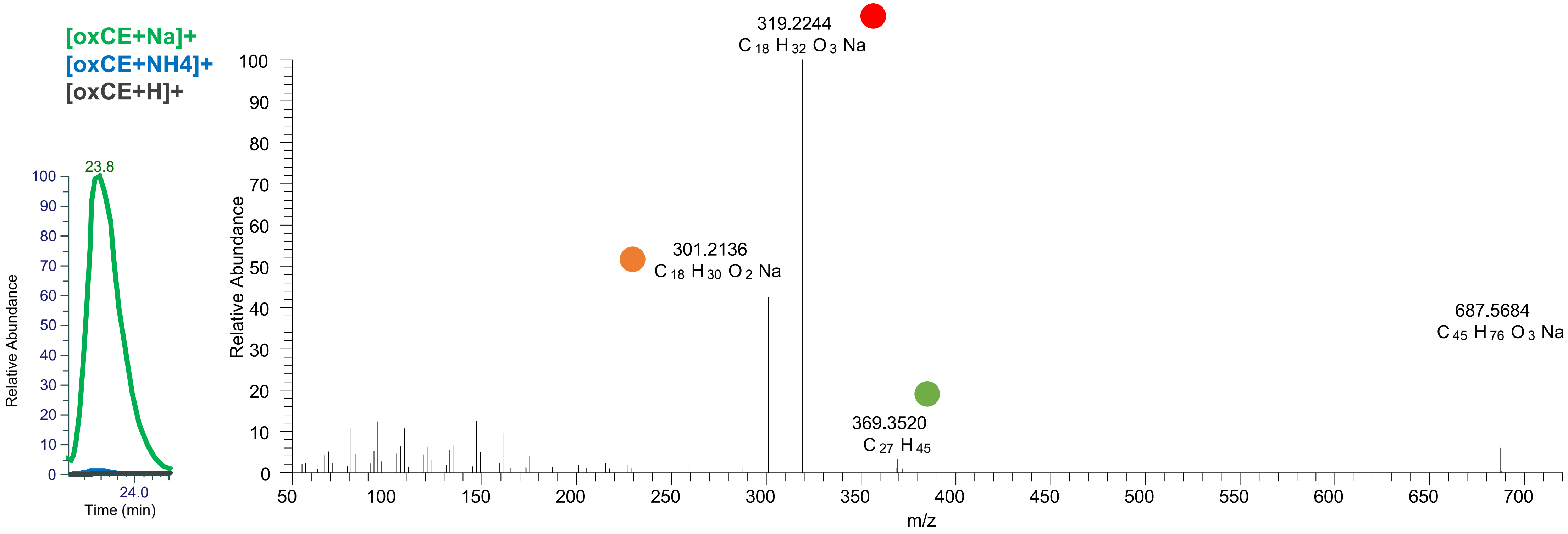

# CE(18:1<ep>)
# RT 25.2

**[oxCE+Na]+** **XIC 687.5687 NL: 3.37E5**

- Fragments containing oxFAs
- Fragments related to water loss
- Fragments not containing oxFAs
- Position-specific fragments
- Fragments related to other oxLPPs

[oxCE+Na]+
[oxCE+NH4]+
[oxCE+H]+

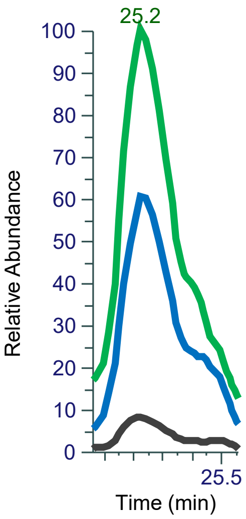


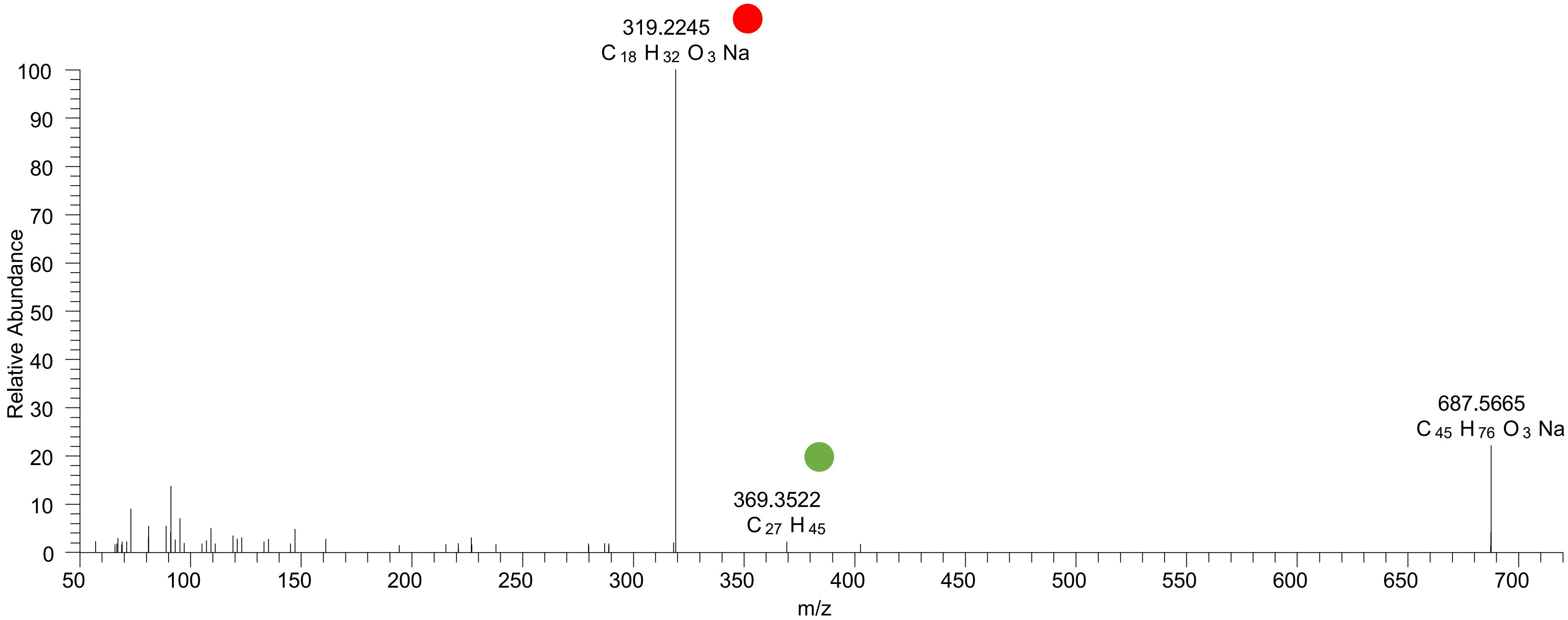

# CE(18:1<OH>)
# RT 24.2

**[oxCE+Na]+** **XIC 689.5843 NL: 4.81E5**

- Fragments containing oxFAs
- Fragments related to water loss
- Fragments not containing oxFAs
- Position-specific fragments
- Fragments related to other oxLPPs

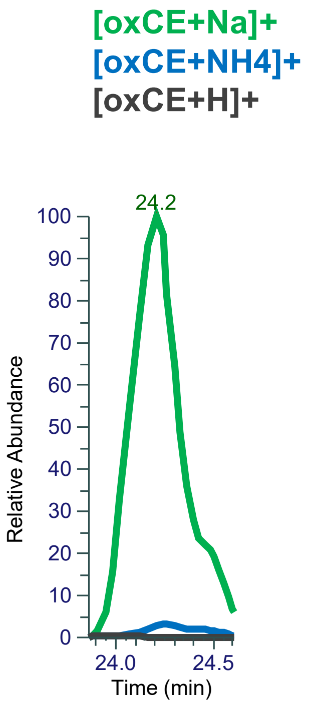


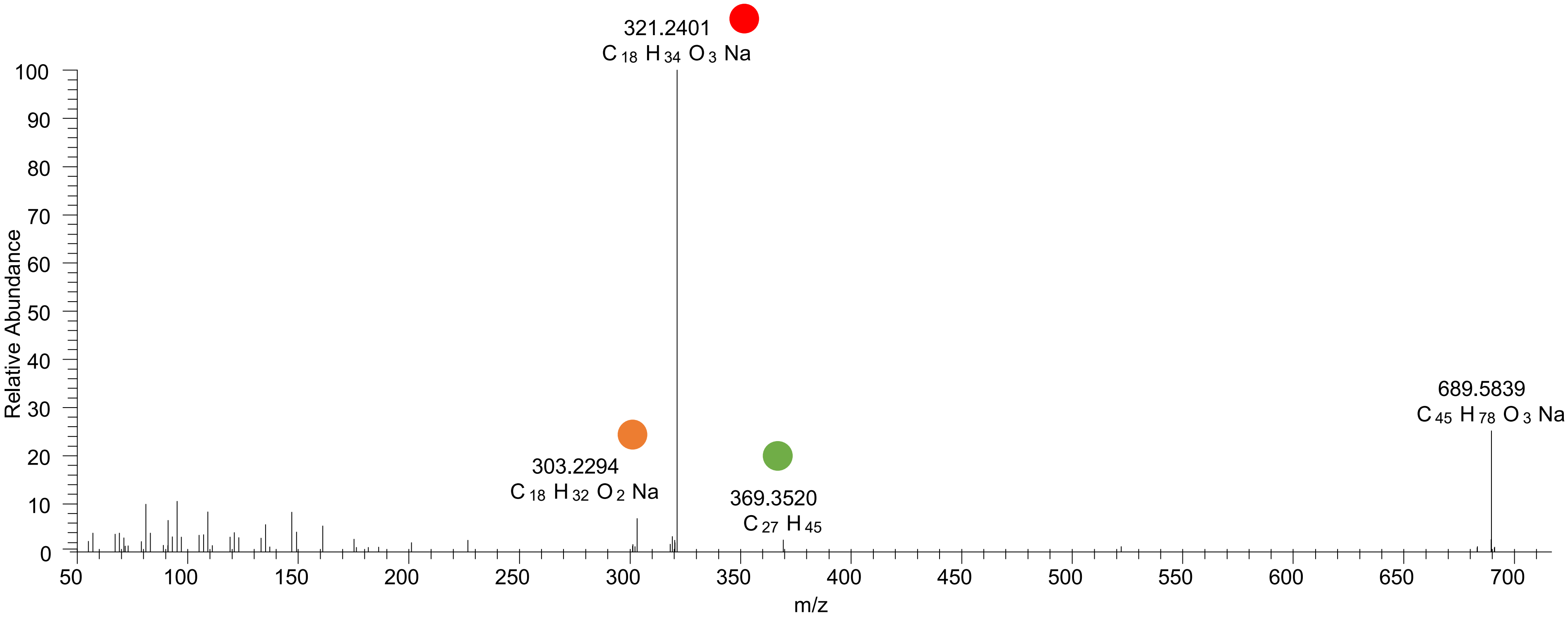

# CE(18:3<2OH>)
# RT 20.6

**[oxCE+Na]+** **XIC 701.5480 NL: 2.15E5**

- Fragments containing oxFAs
- Fragments related to water loss
- Fragments not containing oxFAs
- Position-specific fragments
- Fragments related to other oxLPPs

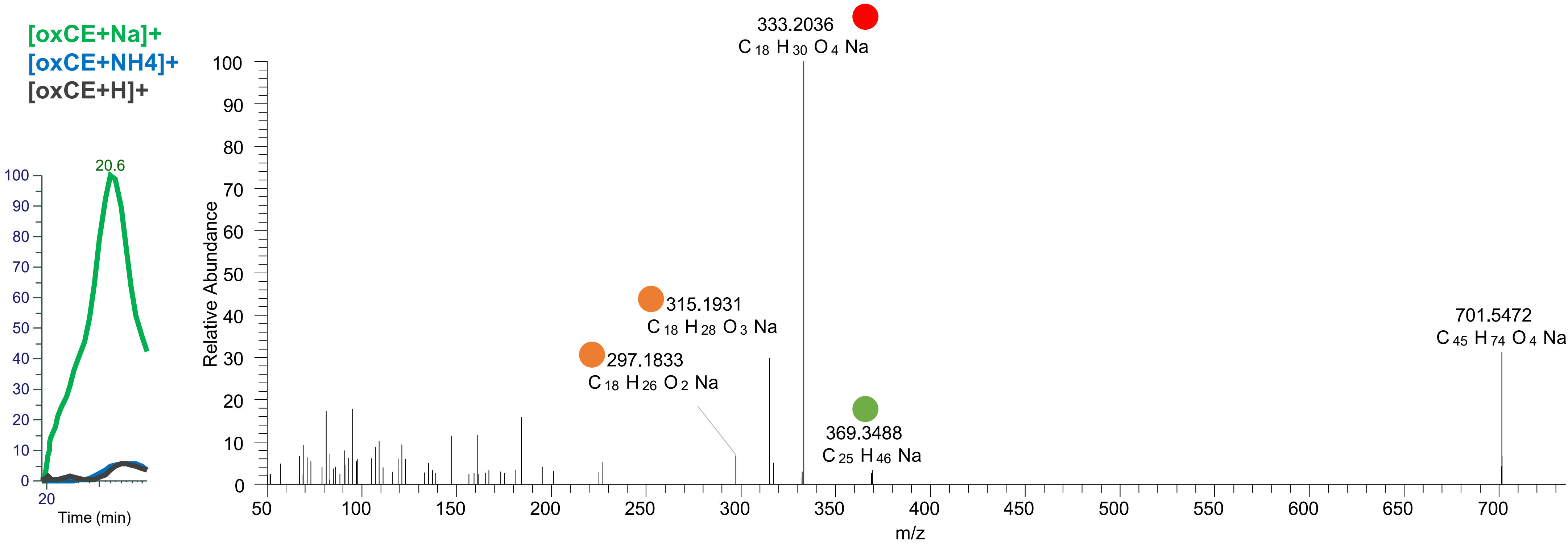

# CE(18:3<OH,O>)
# RT 21.3

[oxCE+Na]+ XIC 701.5480 NL: 2.82E5

Fragments containing oxFAs
Fragments related to water loss
Fragments not containing oxFAs
Position-specific fragments
Fragments related to other oxLPPs

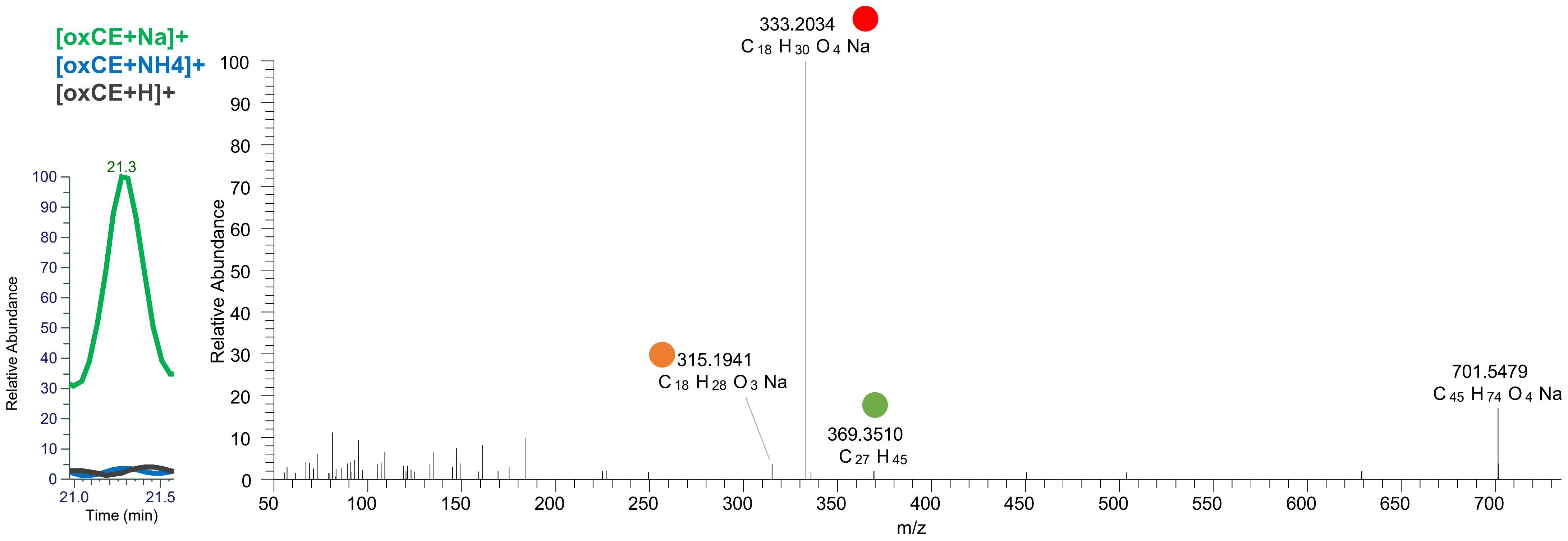

# CE(18:3<OH,O>)
# RT 21.8

**[oxCE+Na]+** **XIC 701.5480 NL: 2.70E5**

Fragments containing oxFAs

Fragments related to water loss

Fragments not containing oxFAs

Position-specific fragments

Fragments related to other oxLPPs

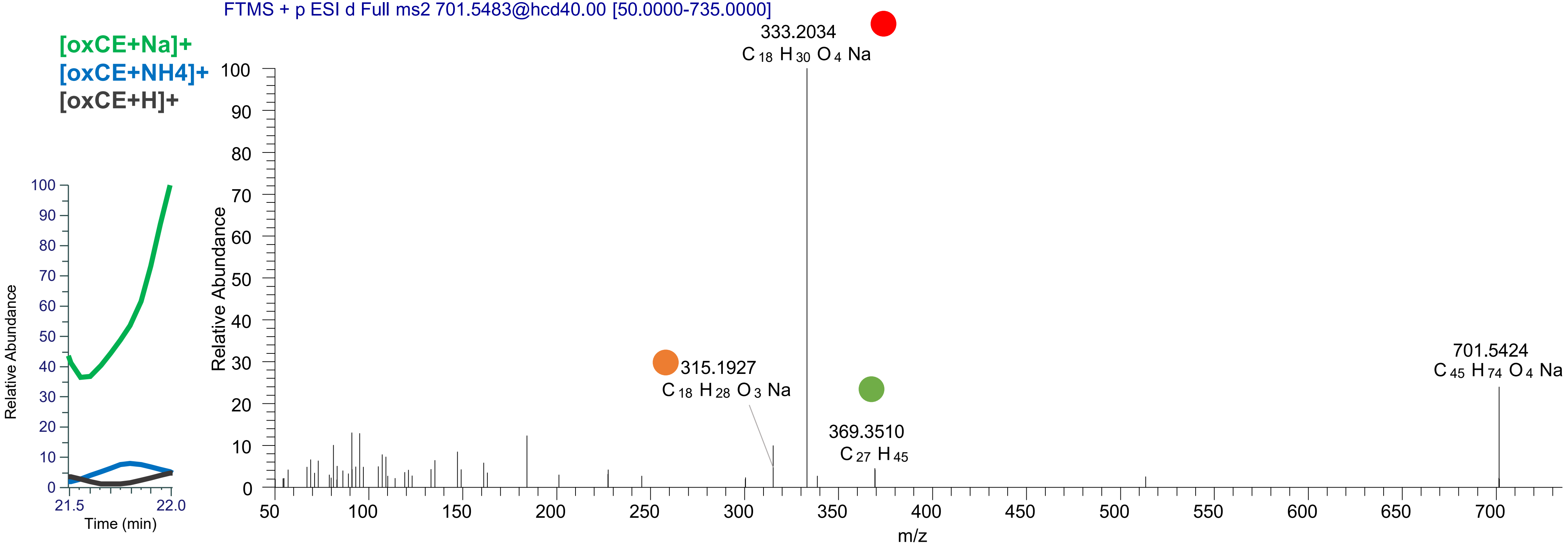

# CE(18:3<OH,O>)
# RT 22.2

**[oxCE+Na]+** **XIC 701.5480 NL: 3.79E5**

- Fragments containing oxFAs
- Fragments related to water loss
- Fragments not containing oxFAs
- Position-specific fragments
- Fragments related to other oxLPPs

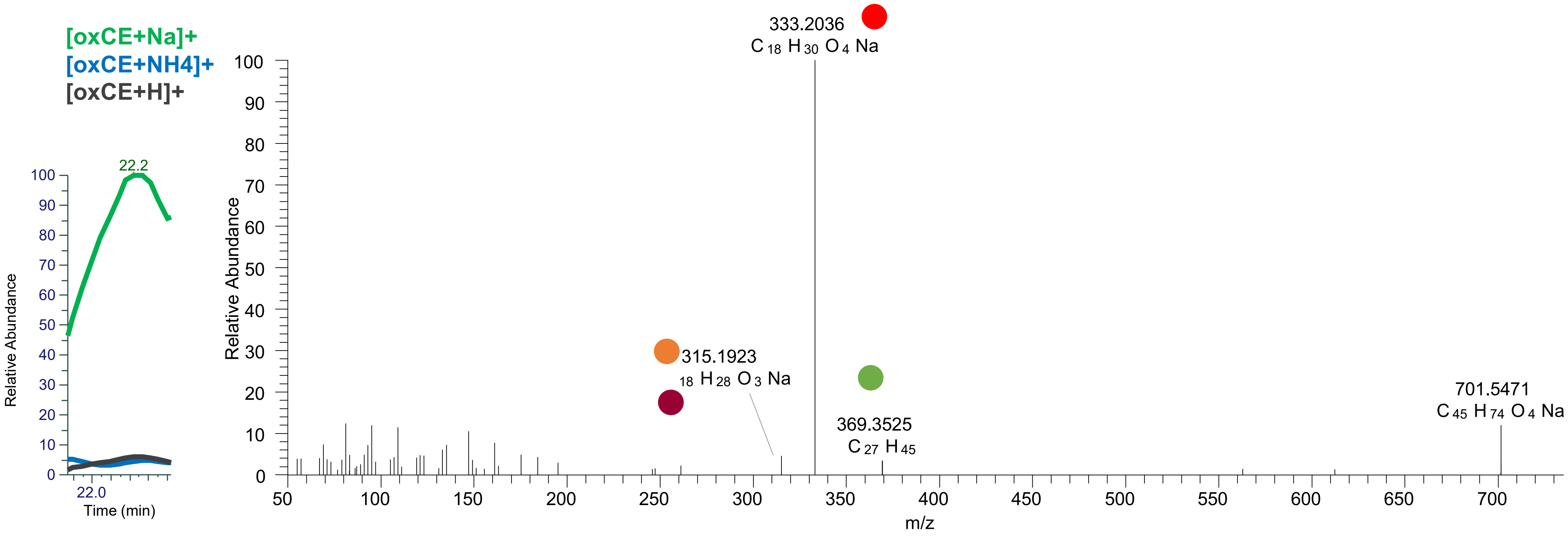

# CE(18:3<OH,O{9}>)
# RT 22.6

[oxCE+Na]+ XIC 701.5480 NL: 6.45E5

Fragments containing oxFAs
Fragments related to water loss
Fragments not containing oxFAs
Position-specific fragments
Fragments related to other oxLPPs

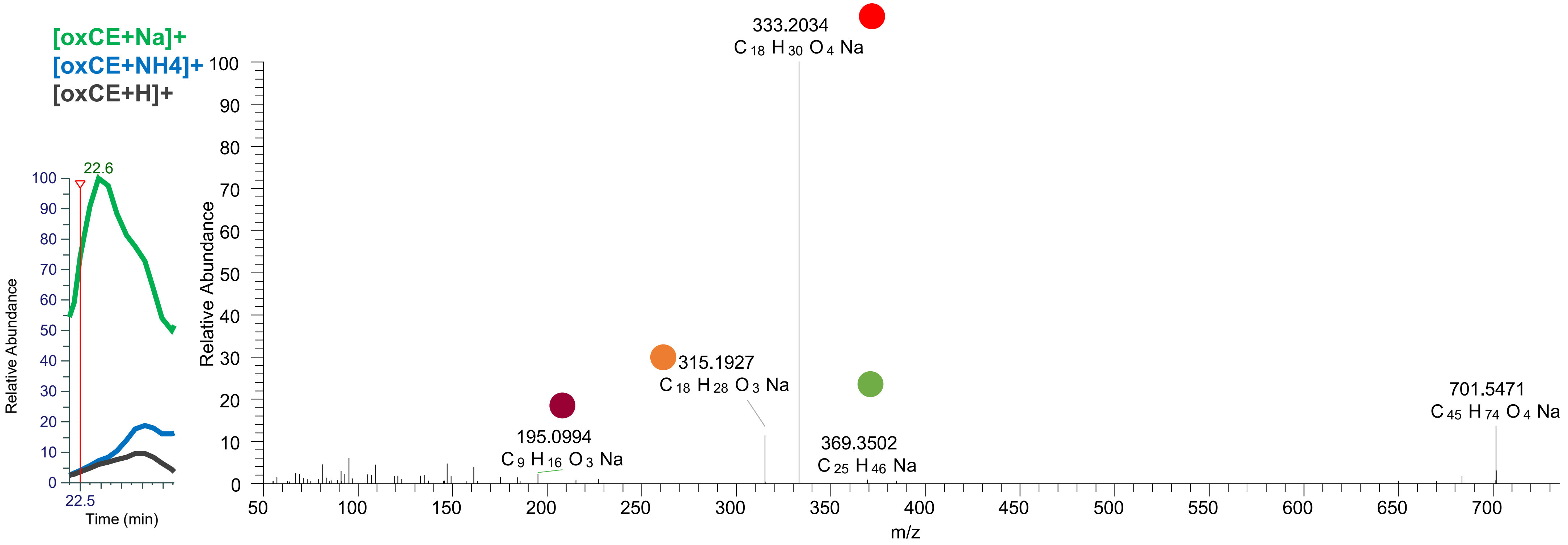

### CE(18:3<OOH{10}>)
### CE(18:3<OOH{13}>)
# RT 23.2

**[oxCE+Na]+** **XIC 701.5480 NL: 8.59E5**

Fragments containing oxFAs
Fragments related to water loss
Fragments not containing oxFAs
Position-specific fragments
Fragments related to other oxLPPs

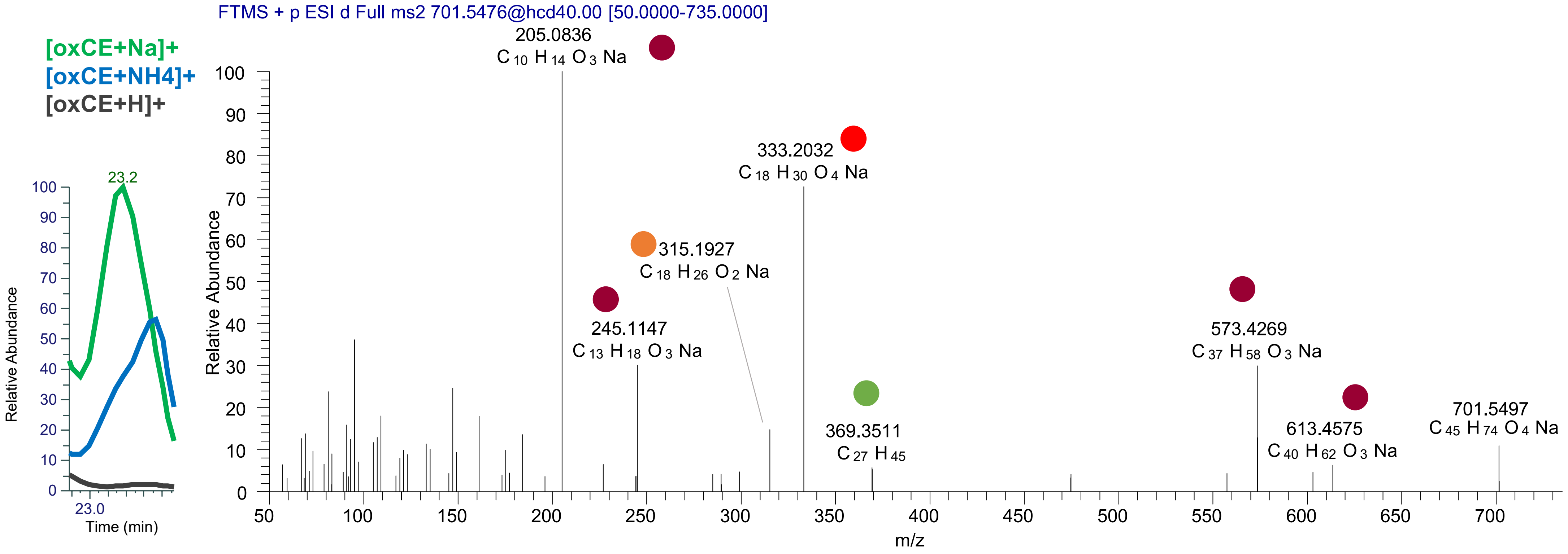

### CE(18:3<OOH{10}>)
### CE(18:3<OOH{13}>)
### CE(18:3<OOH{16}>)
# RT 23.3

**[oxCE+Na]+** **XIC 701.5480 NL: 8.59E5**

- Fragments containing oxFAs
- Fragments related to water loss
- Fragments not containing oxFAs
- Position-specific fragments
- Fragments related to other oxLPPs

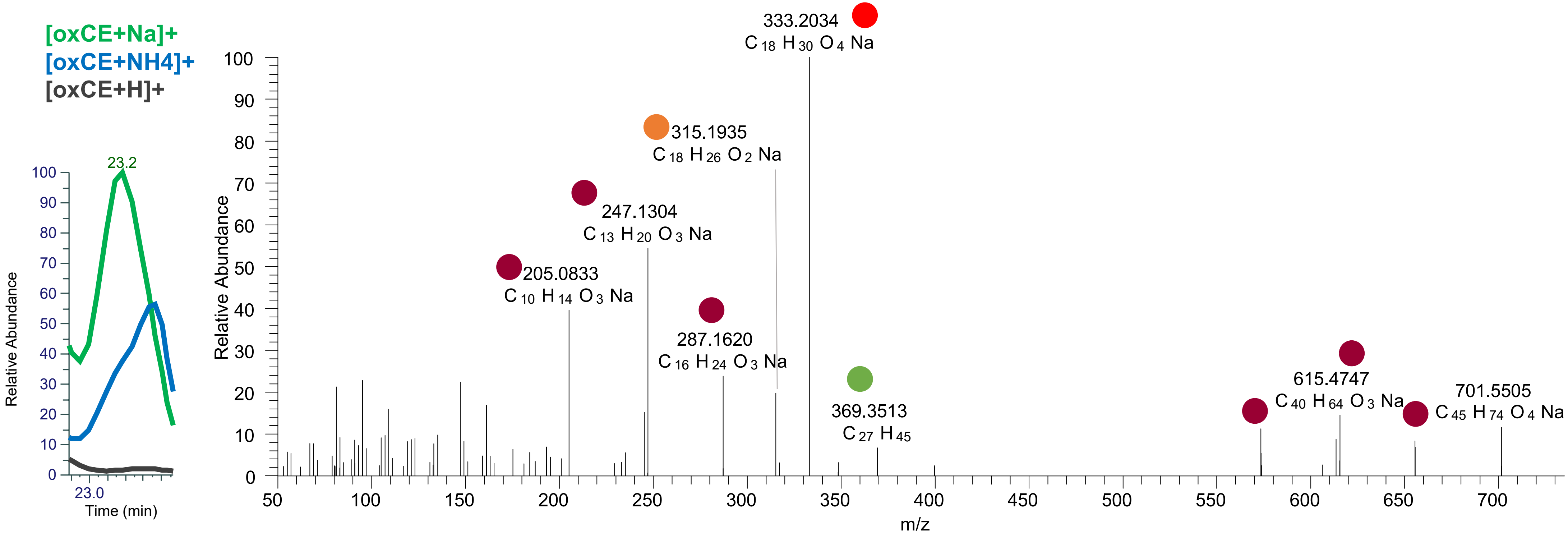

# CE(18:2<2OH>)
# RT 21.2

**[oxCE+Na]+** **XIC 703.5636 NL: 2.68E5**

- Fragments containing oxFAs
- Fragments related to water loss
- Fragments not containing oxFAs
- Position-specific fragments
- Fragments related to other oxLPPs

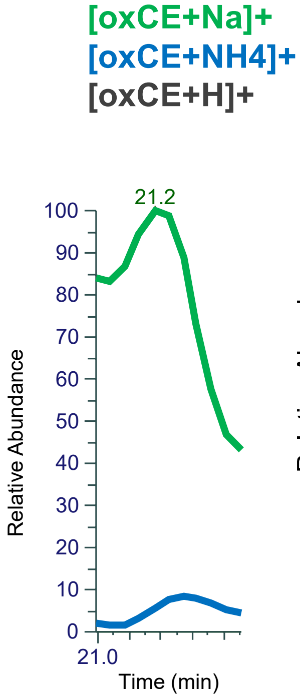


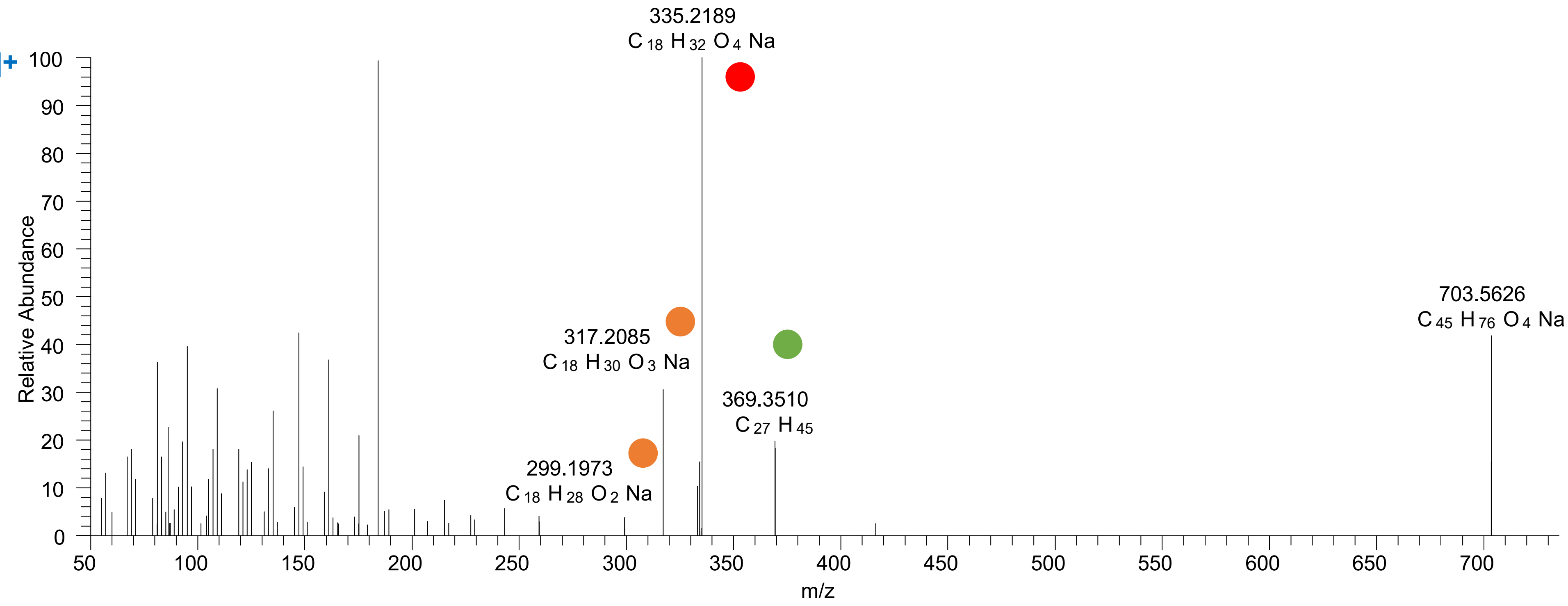

# CE(18:1<OH,ep>)
# RT 21.7

**[oxCE+Na]+** **XIC 703.5636 NL: 1.77E5**

- Fragments containing oxFAs
- Fragments related to water loss
- Fragments not containing oxFAs
- Position-specific fragments
- Fragments related to other oxLPPs

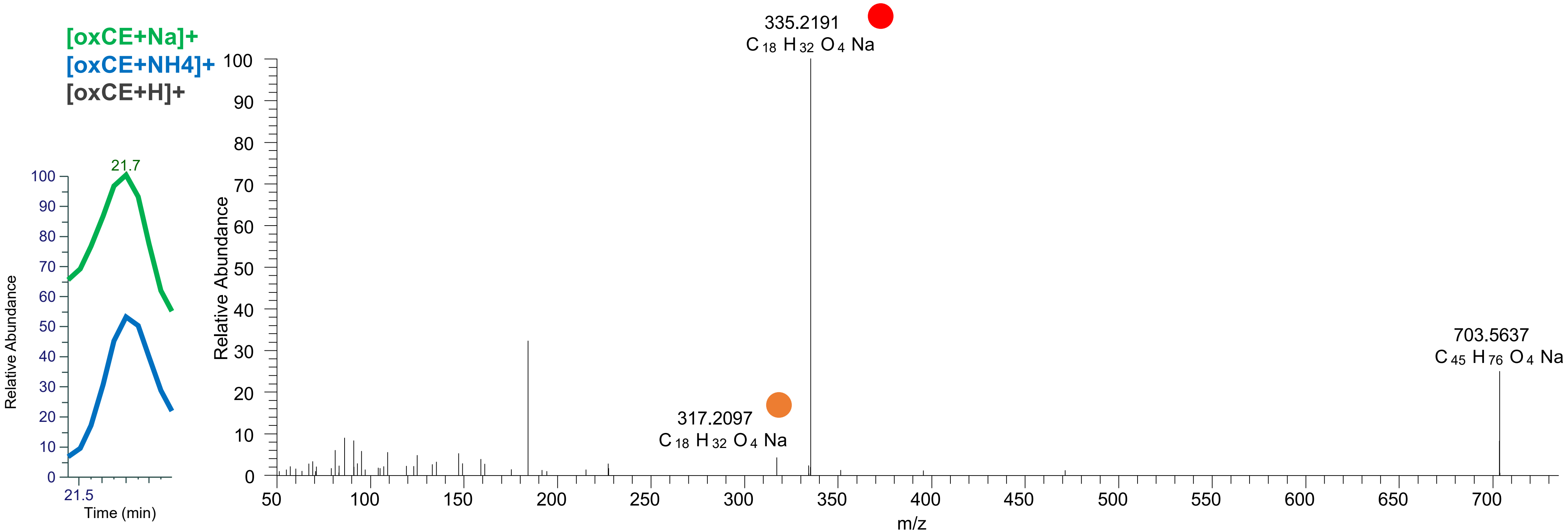

# CE(18:1<OH,ep>)
# RT 22.2

[oxCE+Na]+ XIC 703.5636 NL: 1.45E6

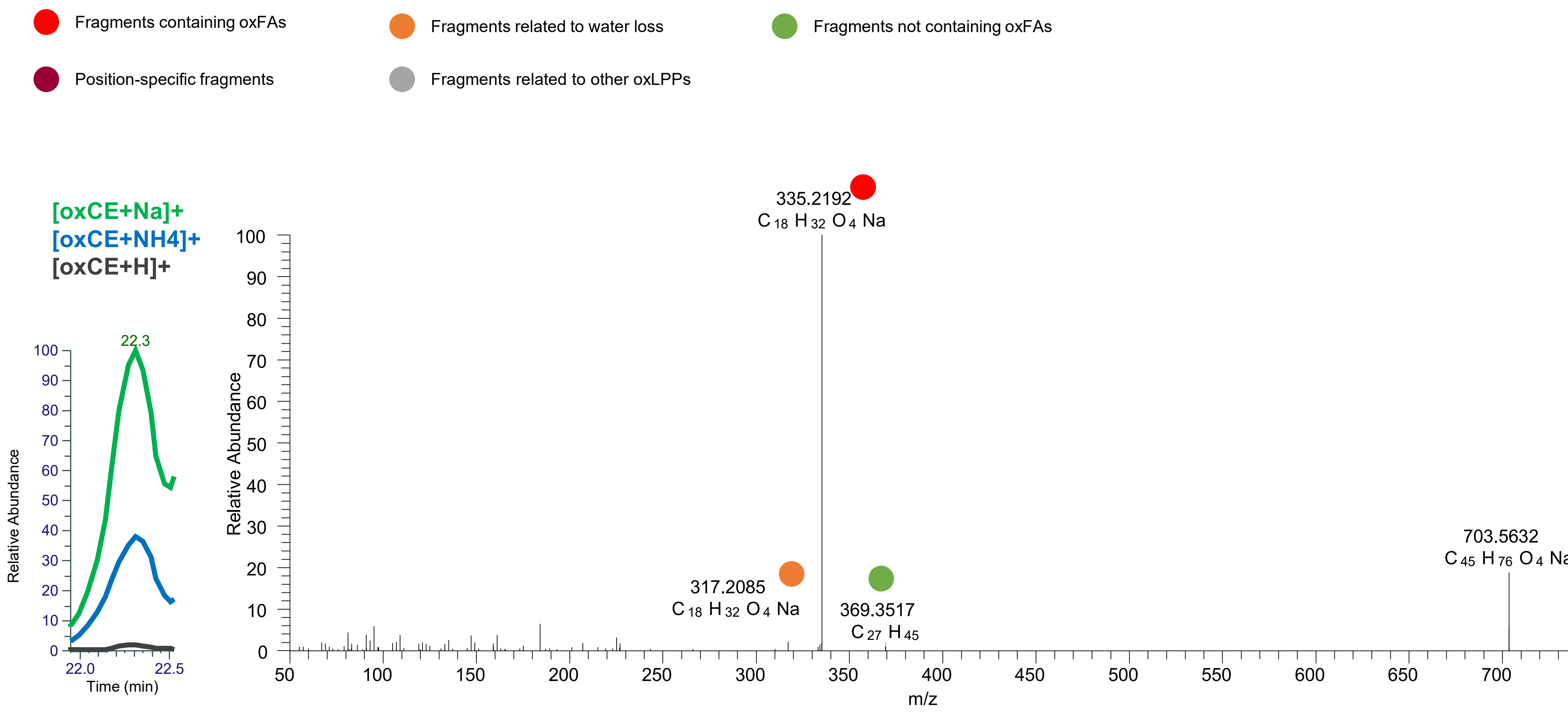

# CE(18:0<2ep>)
# RT 22.7

**[oxCE+Na]+** **XIC 703.5636 NL: 1.72E6**

- Fragments containing oxFAs
- Fragments related to water loss
- Fragments not containing oxFAs
- Position-specific fragments
- Fragments related to other oxLPPs

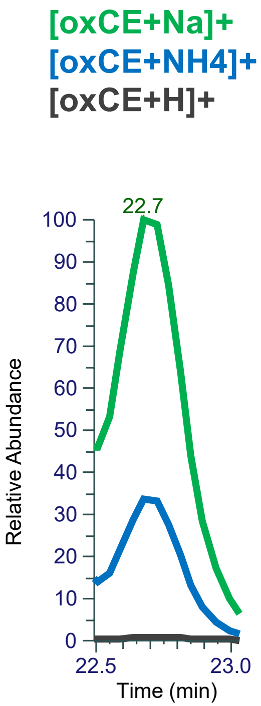


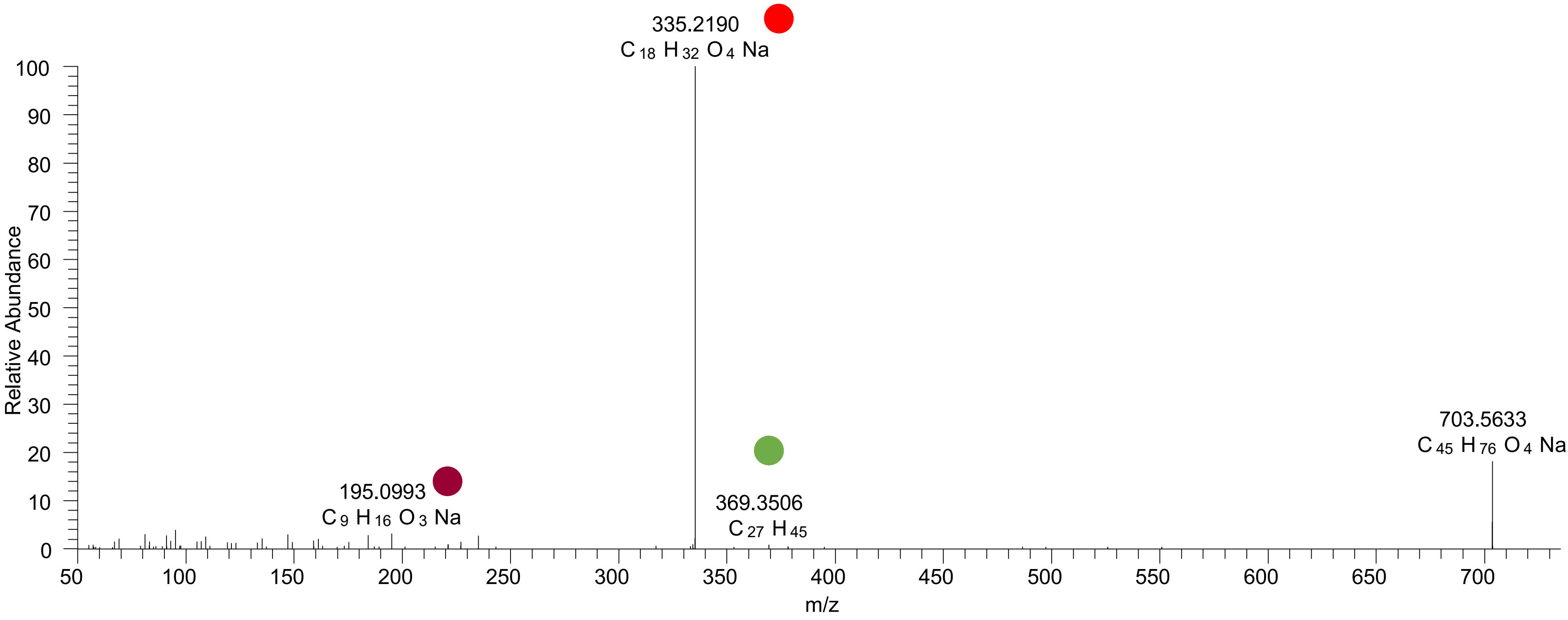

### CE(18:2<OOH{11}>)
### CE(18:2<OOH{13}>)
# RT 23.4

**[oxCE+Na]+** **XIC 703.5636 NL: 2.92E6**

- Fragments containing oxFAs
- Fragments related to water loss
- Fragments not containing oxFAs
- Position-specific fragments
- Fragments related to other oxLPPs

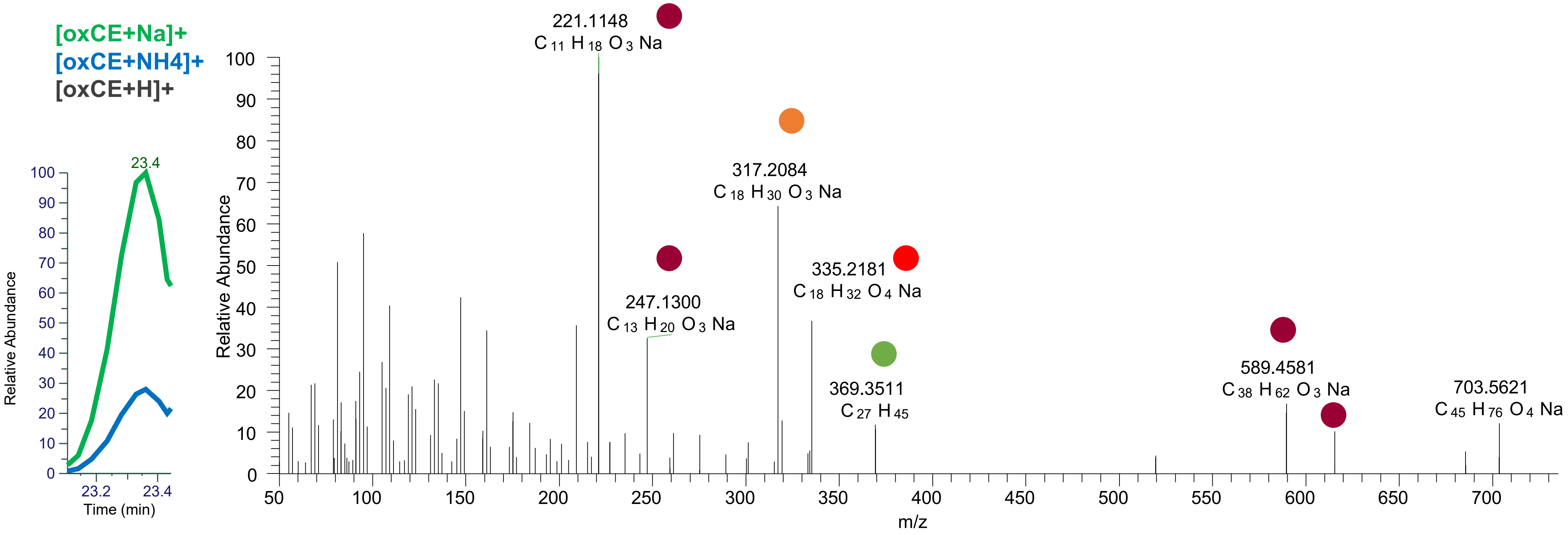

### CE(18:2<OOH{9}>)
### CE(18:2<OOH{13}>)
# RT 23.6

**[oxCE+Na]+**        **XIC 703.5636 NL: 1.36E7**

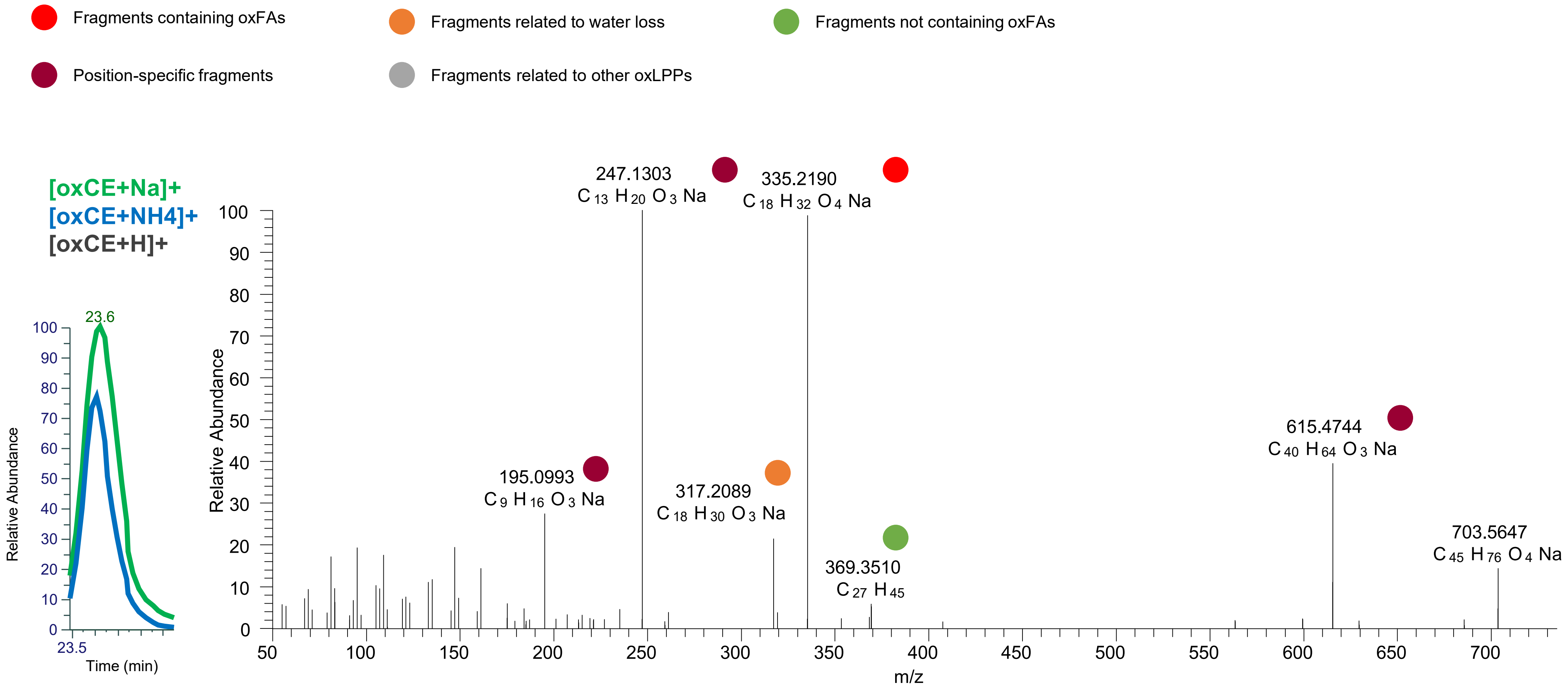

# CE(18:1<2O>)
# RT 23.1

**[oxCE+Na]+** **XIC 705.5792 NL: 1.36E4**

Fragments containing oxFAs
Fragments related to water loss
Fragments not containing oxFAs
Position-specific fragments
Fragments related to other oxLPPs

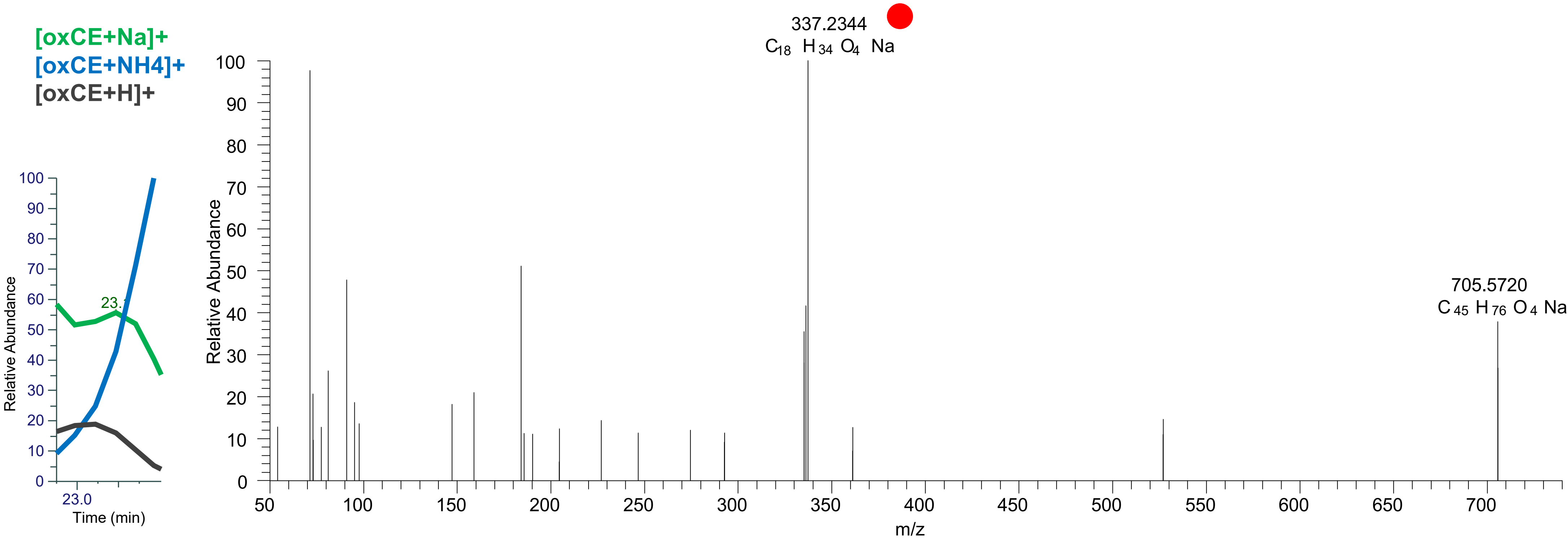

CE(18:1<OOH{10}>)
CE(18:1<OOH{11}>)
CE(18:1<OOH{12}>)
CE(18:1<OOH{13}>)
RT 24.0

[oxCE+Na]+ XIC 705.5792 NL: 4.12E5

- Fragments containing oxFAs
- Fragments related to water loss
- Fragments not containing oxFAs
- Position-specific fragments
- Fragments related to other oxLPPs
- Fragments containing oxChol

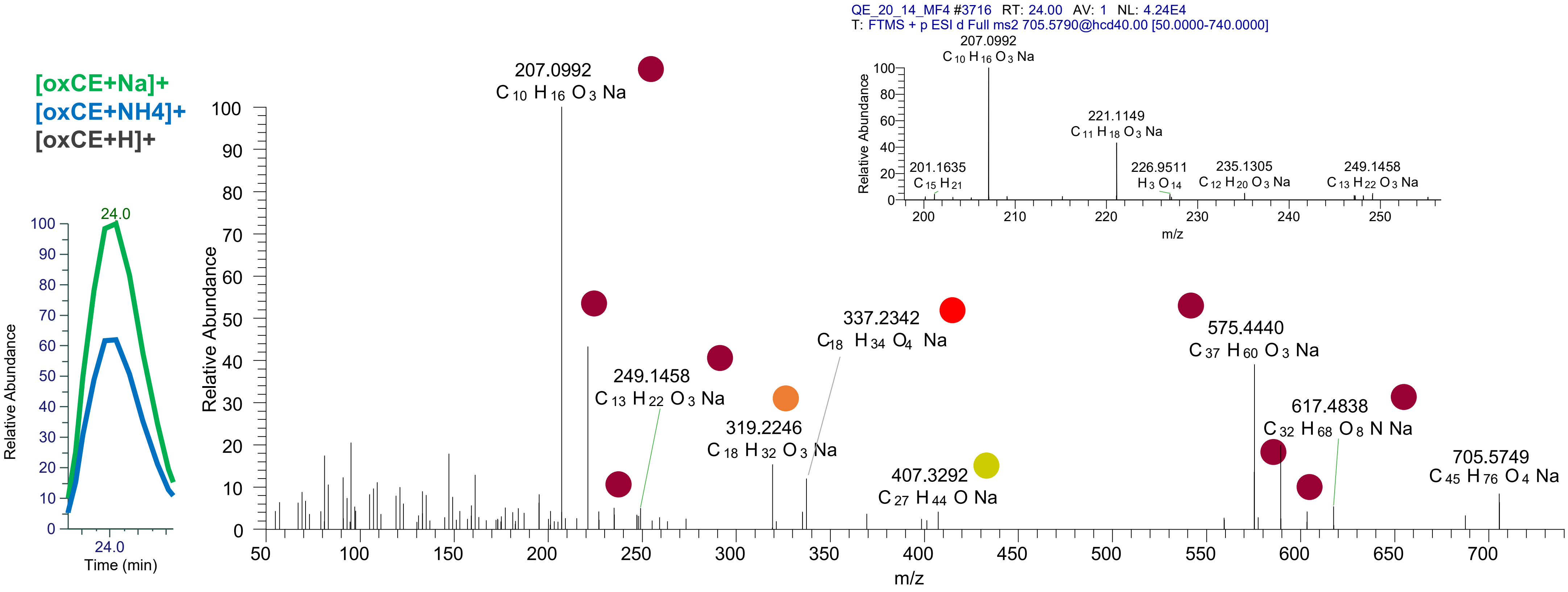

# CE(20:5<OH>)
# RT 23.1

**[oxCE+Na]+** **XIC 709.5530 NL: 9.57E5**

- Fragments containing oxFAs
- Fragments related to water loss
- Fragments not containing oxFAs
- Position-specific fragments
- Fragments related to other oxLPPs
- Fragments containing oxChol

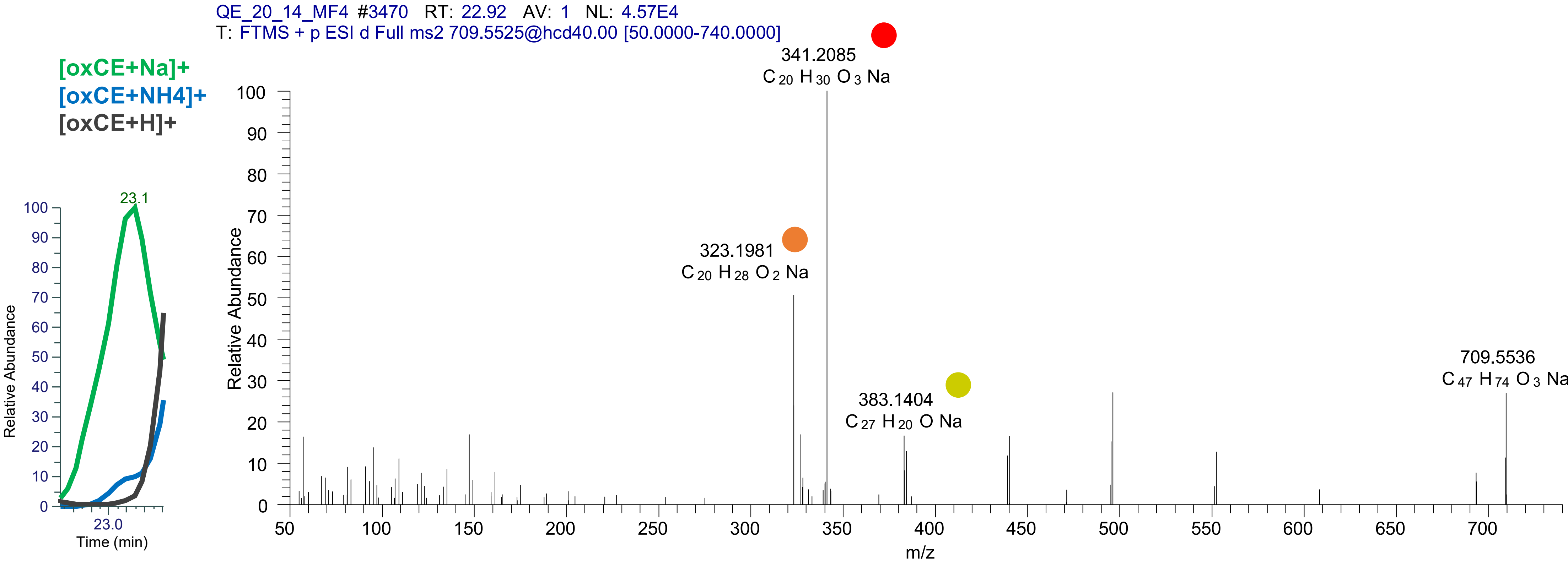

# CE(20:4<OH>)
# RT 23.6

**[oxCE+Na]+** **XIC 711.5687 NL: 2.94E6**

- Fragments containing oxFAs
- Fragments related to water loss
- Fragments not containing oxFAs
- Position-specific fragments
- Fragments related to other oxLPPs

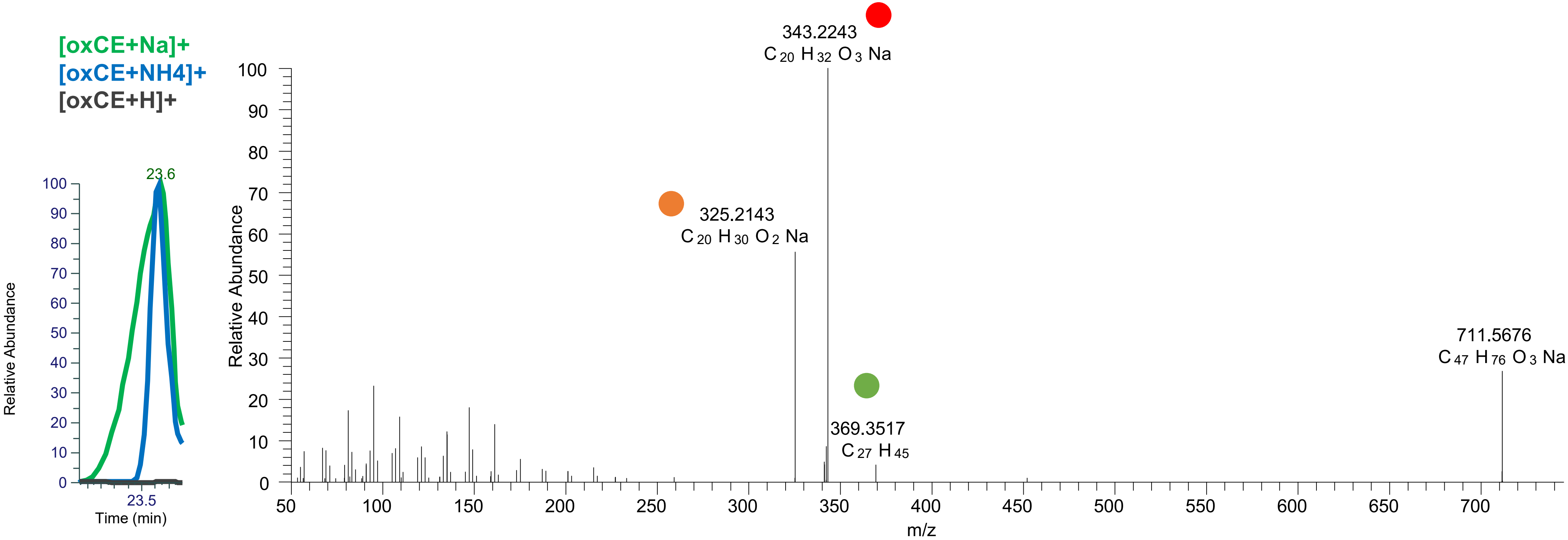

### CE(20:3<oxo>)
# RT 24.2

**[oxCE+Na]+** **XIC 711.5687 NL: 1.30E5**

- Fragments containing oxFAs
- Fragments related to water loss
- Fragments not containing oxFAs
- Position-specific fragments
- Fragments related to other oxLPPs

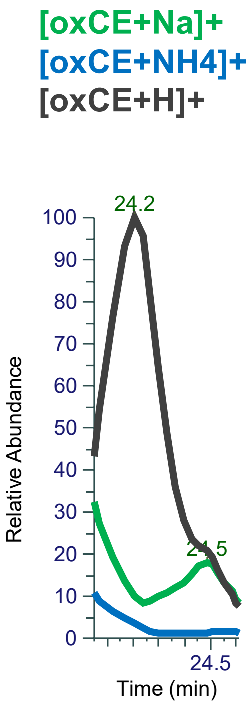


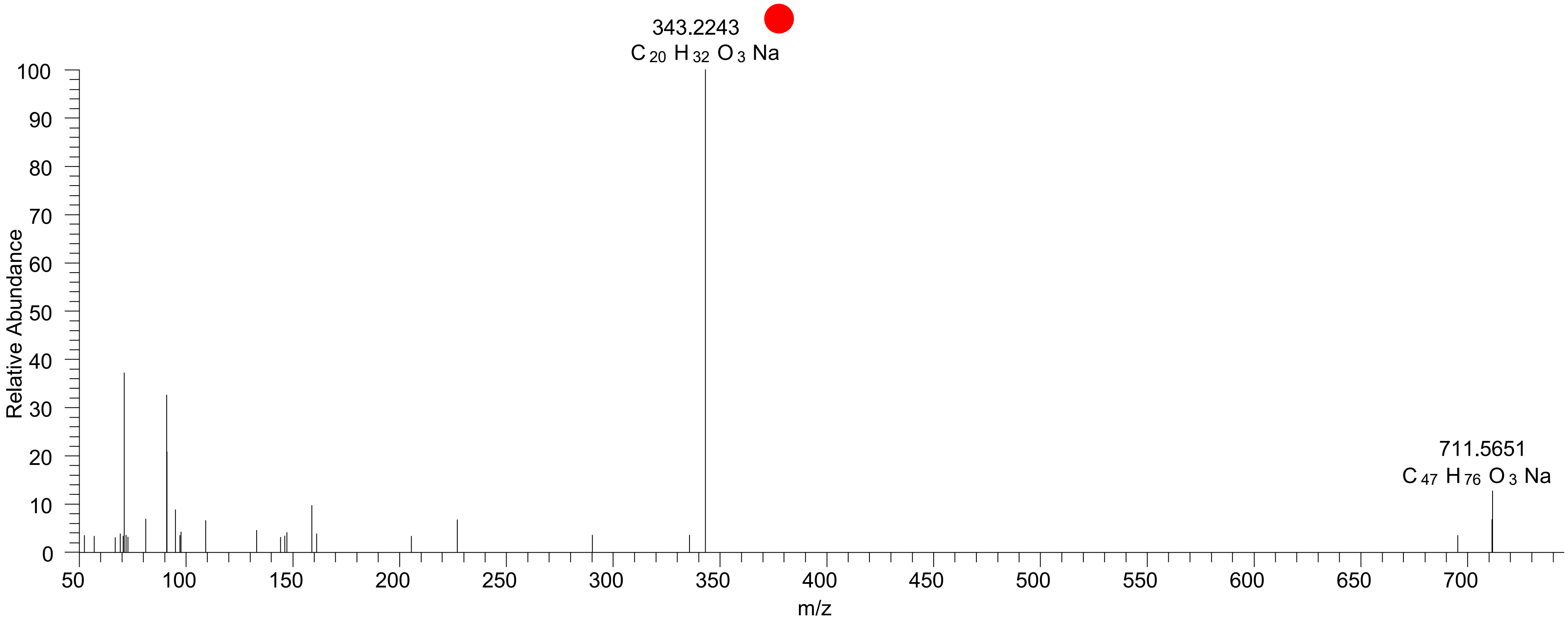

# CE(20:4<O>)
# RT 25.0

[oxCE+Na]+ XIC 711.5687 NL: 5.25E4

Fragments containing oxFAs
Fragments related to water loss
Fragments not containing oxFAs
Position-specific fragments
Fragments related to other oxLPPs

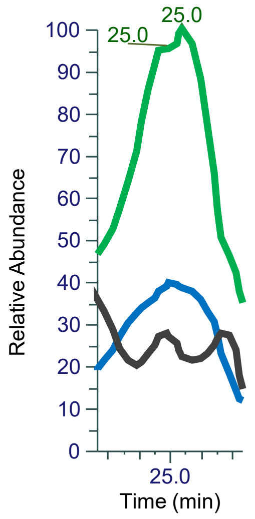


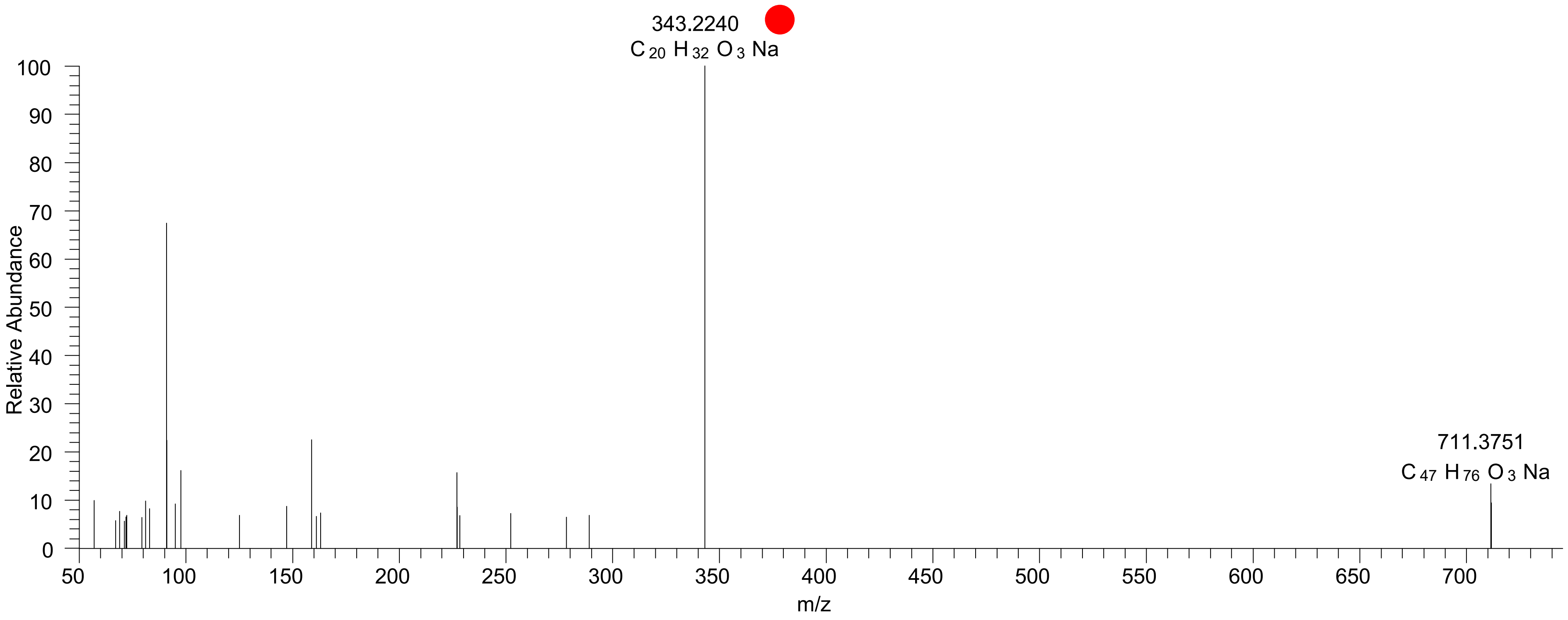

# CE(20:3<OH>)
# RT 24.2

**[oxCE+Na]+** **XIC 713.5843 NL: 1.43E6**

- Fragments containing oxFAs
- Fragments related to water loss
- Fragments not containing oxFAs
- Position-specific fragments
- Fragments related to other oxLPPs

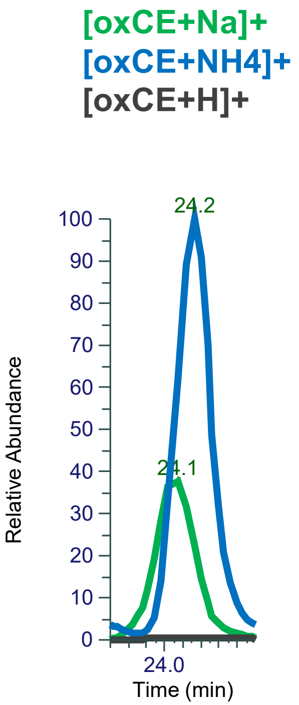


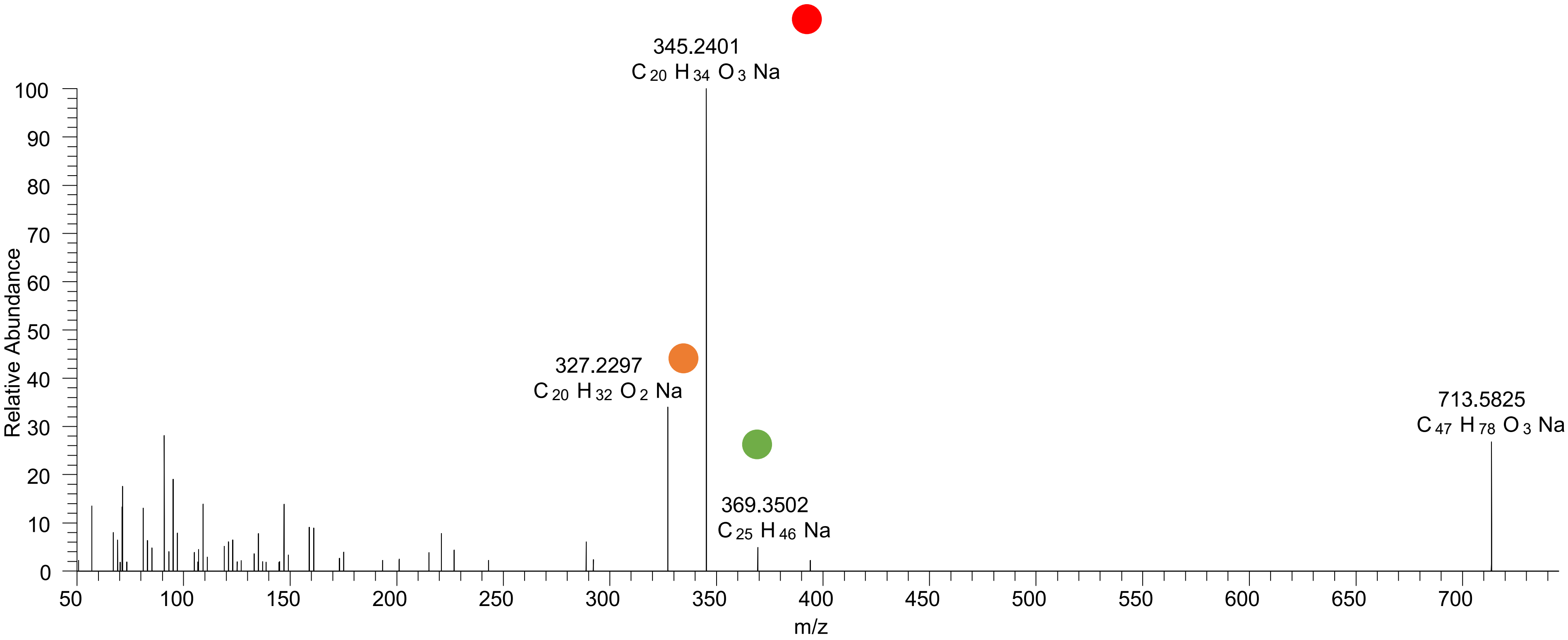

# CE(20:2<OH>)
# RT 24.4

**[oxCE+Na]$^+$** **XIC 715.6005 NL: 4.37E4**

- Fragments containing oxFAs
- Fragments related to water loss
- Fragments not containing oxFAs
- Position-specific fragments
- Fragments related to other oxLPPs

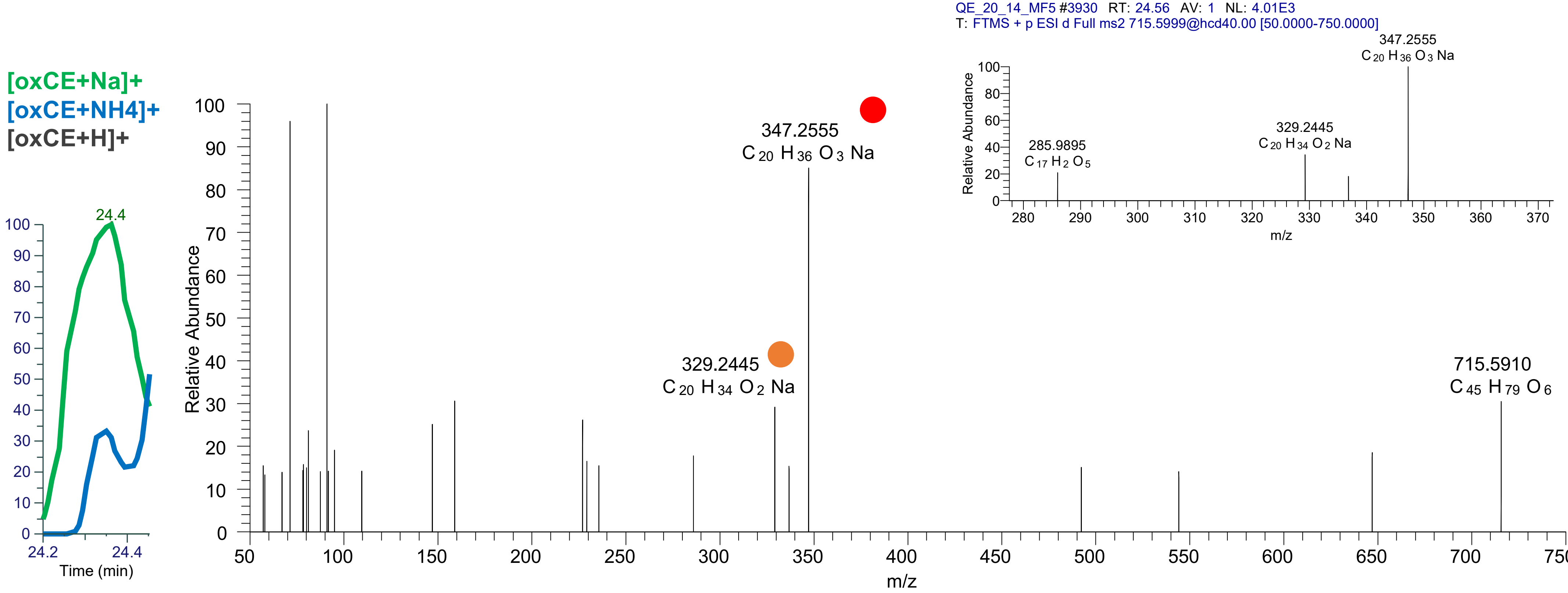

# CE(20:1<ep>)
# RT 24.8

[oxCE+Na]$^{+}$ XIC 715.6005 NL: 4.37E4

Fragments containing oxFAs
Fragments related to water loss
Fragments not containing oxFAs
Position-specific fragments
Fragments related to other oxLPPs

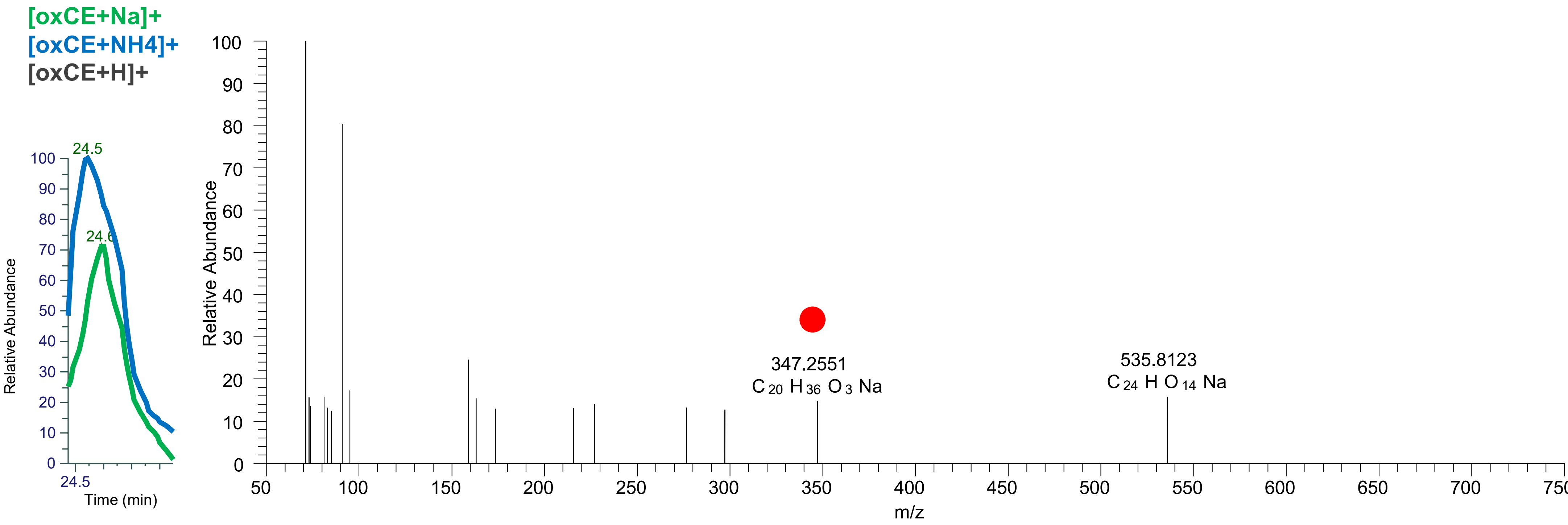

### CE(18:3<OH,OOH{13}>)
# RT 20.3

**[oxCE+Na]+** **XIC 717.5434 NL: 1.19E5**

- Fragments containing oxFAs
- Fragments related to water loss
- Fragments not containing oxFAs
- Position-specific fragments
- Fragments related to other oxLPPs

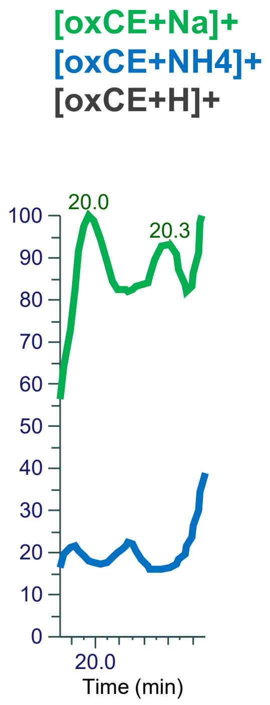


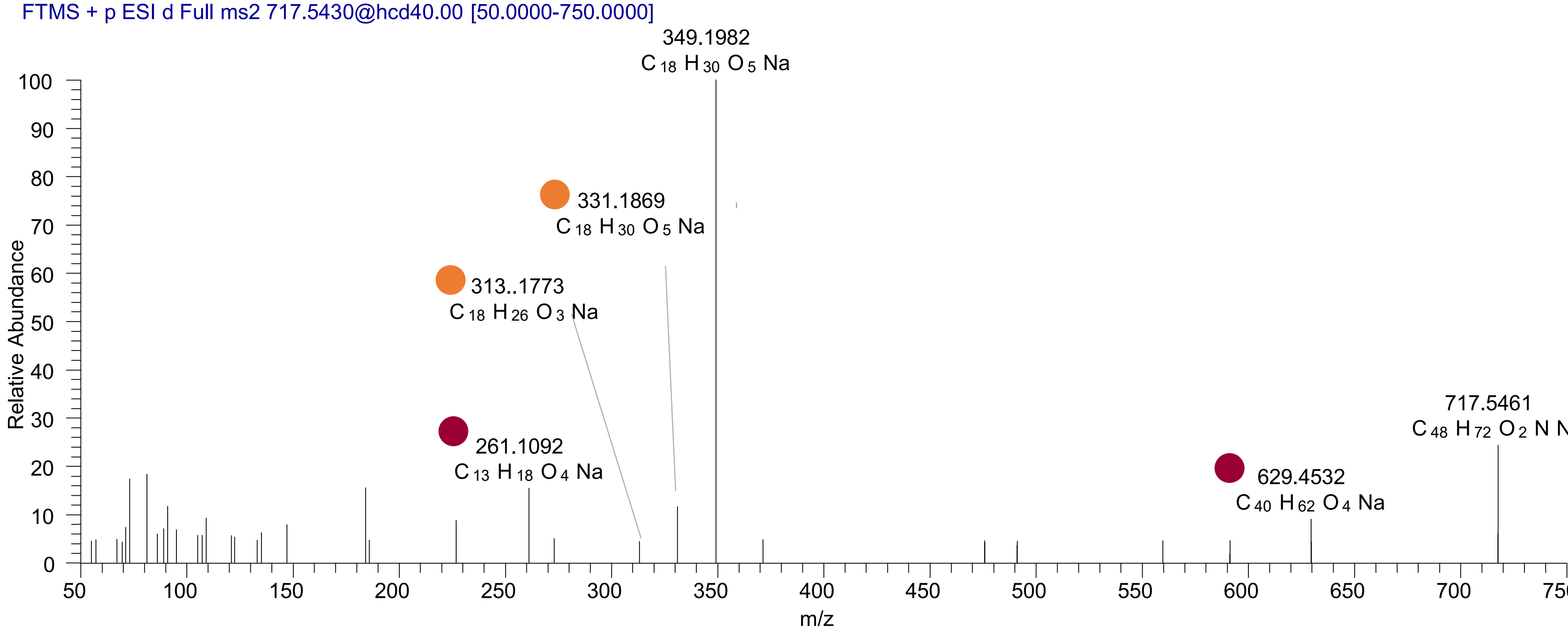

# CE(18:2<3O>)
# RT 20.8

[oxCE+Na]+ XIC 719.5590 NL: 1.45E5

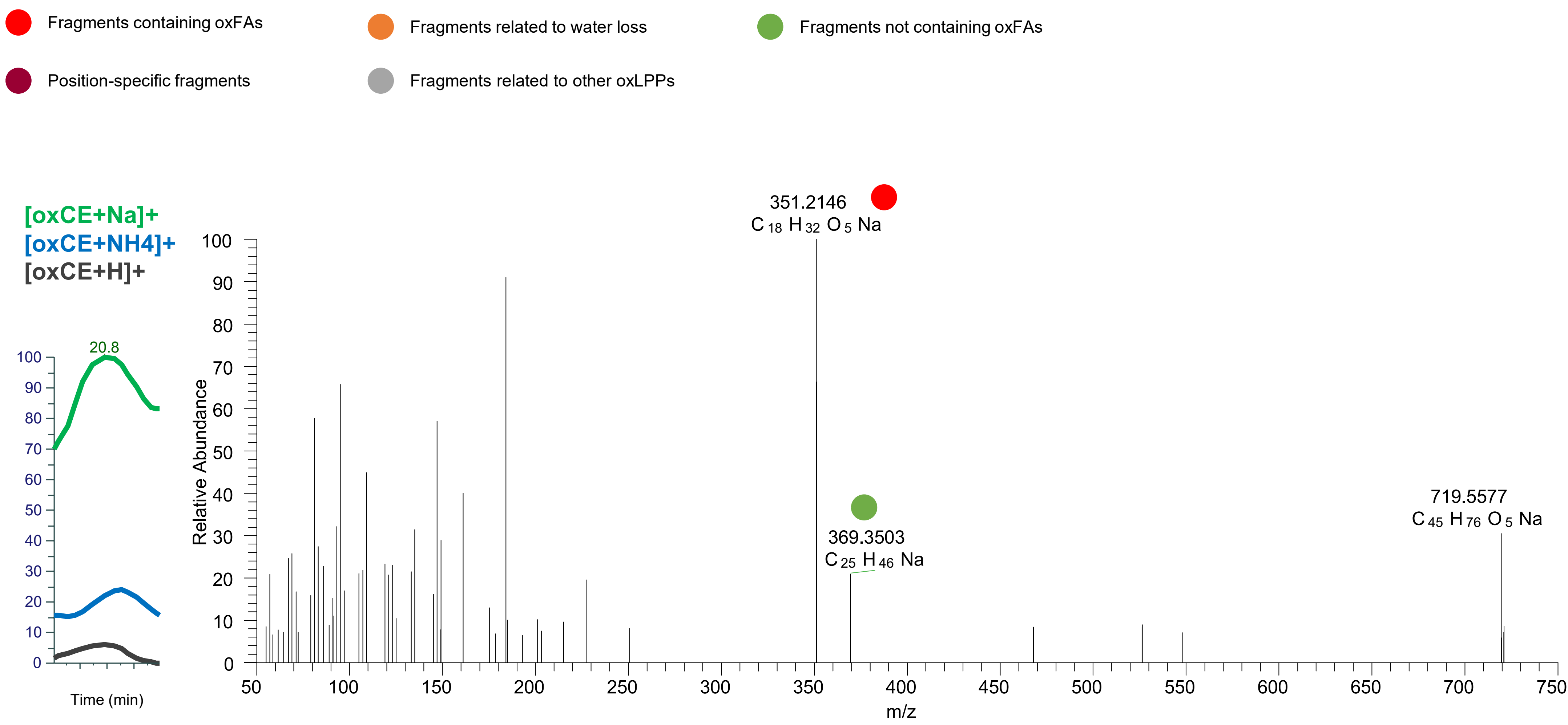

### CE(18:2<O,OOH{13}>)
# RT 22.0

**[oxCE+Na]+** **XIC 719.5590 NL: 3.77E5**

- Fragments containing oxFAs
- Fragments related to water loss
- Fragments not containing oxFAs
- Position-specific fragments
- Fragments related to other oxLPPs

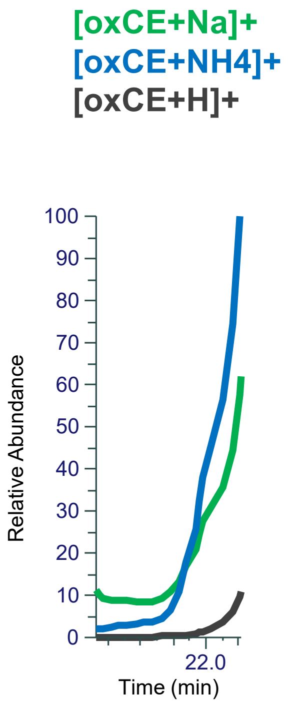


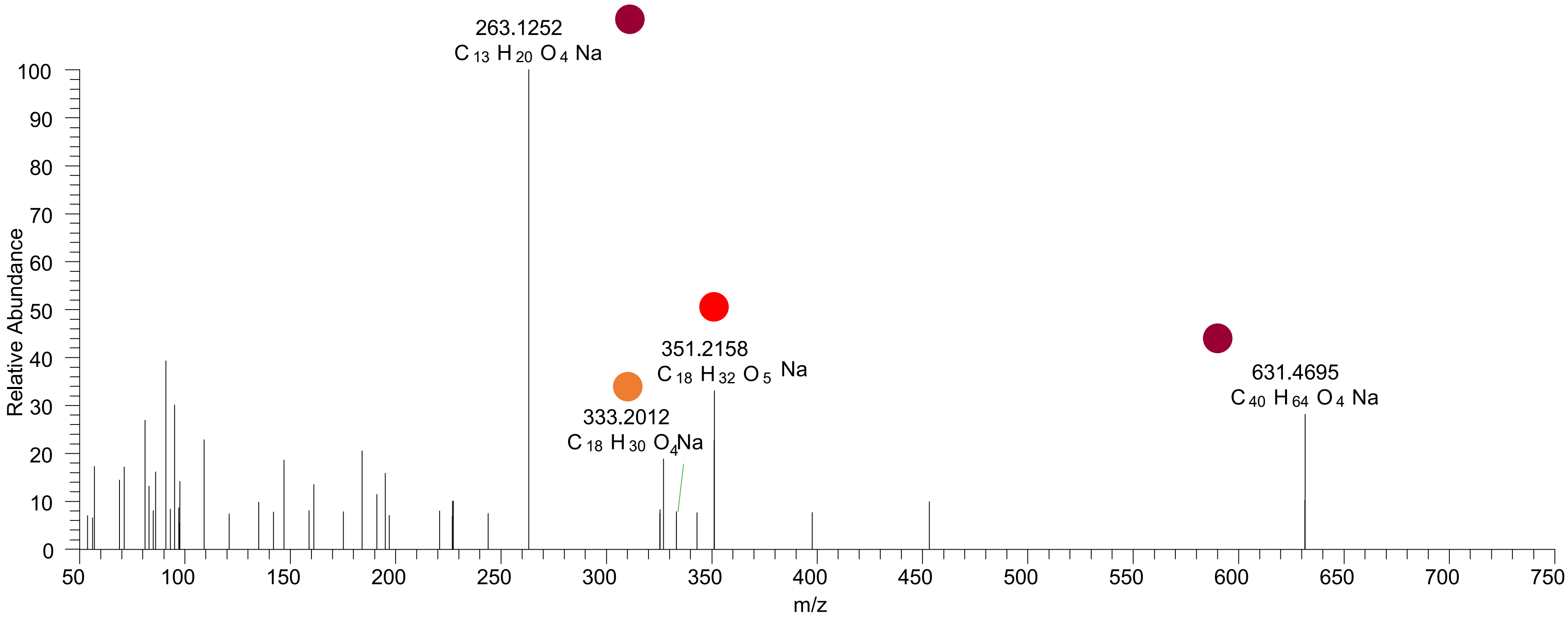

### CE(18:2<O{11},OOH{13}>)
### CE(18:2<O{Chol},OOH{9}>)
# RT 22.2

**[oxCE+Na]+** **XIC 719.5590 NL: 8.63E5**

- Fragments containing oxFAs
- Fragments related to water loss
- Fragments not containing oxFAs
- Position-specific fragments
- Fragments related to other oxLPPs
- Fragments containing oxChol

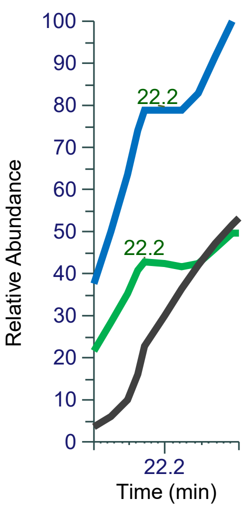


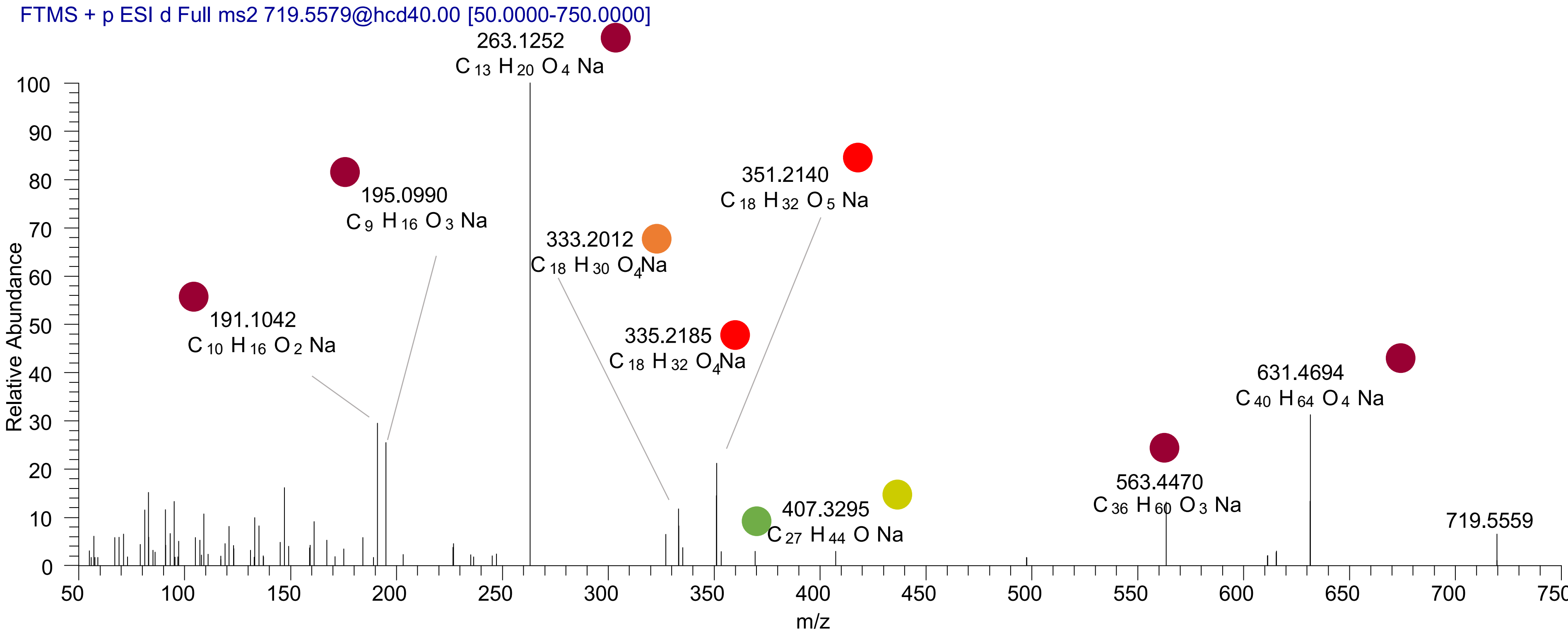

### CE(18:2<O{11},OOH{13}>)
# RT 22.5

**[oxCE+Na]+ XIC 719.5590 NL: 8.72E5**

- Fragments containing oxFAs
- Fragments related to water loss
- Fragments not containing oxFAs
- Position-specific fragments
- Fragments related to other oxLPPs

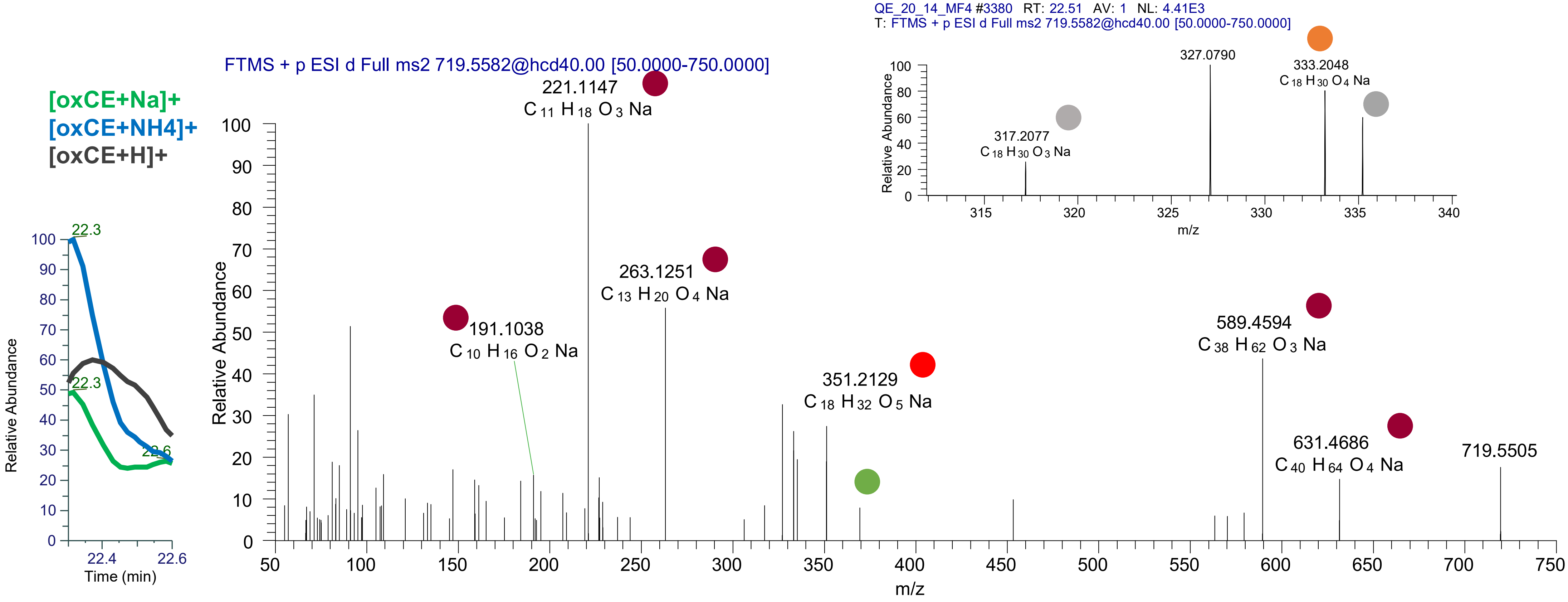

# CE(20:4<OH,ep>)
# RT 19.9

**[oxCE+Na]+** **XIC 723.5323 NL: 5.69E4**

- Fragments containing oxFAs
- Fragments related to water loss
- Fragments not containing oxFAs
- Position-specific fragments
- Fragments related to other oxLPPs

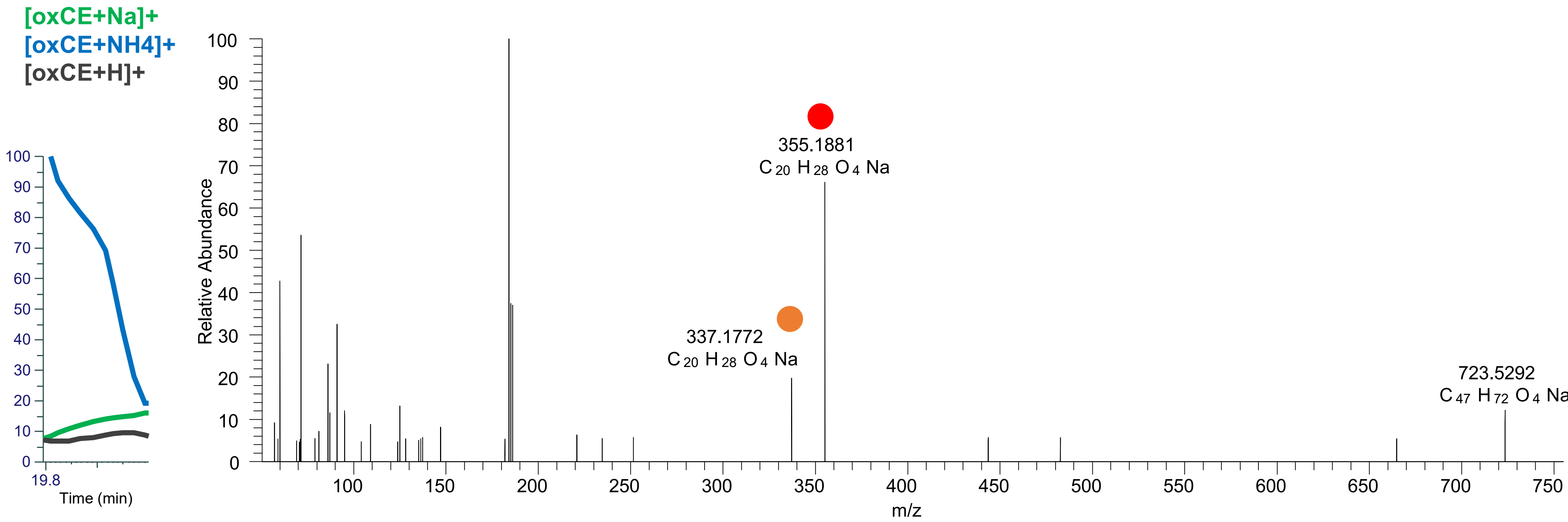

### CE(20:4<O,oxo>)
# RT 20.6

**[oxCE+Na]+** **XIC 723.5323 NL: 9.12E4**

Fragments containing oxFAs

Fragments related to water loss

Fragments not containing oxFAs

Position-specific fragments

Fragments related to other oxLPPs

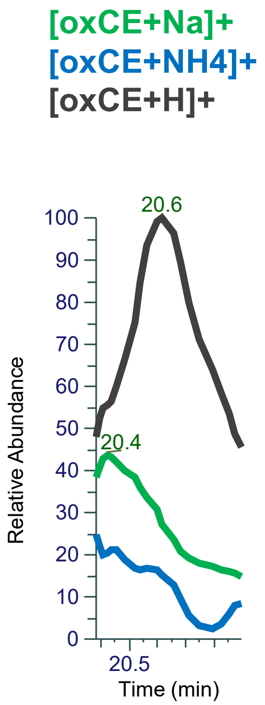


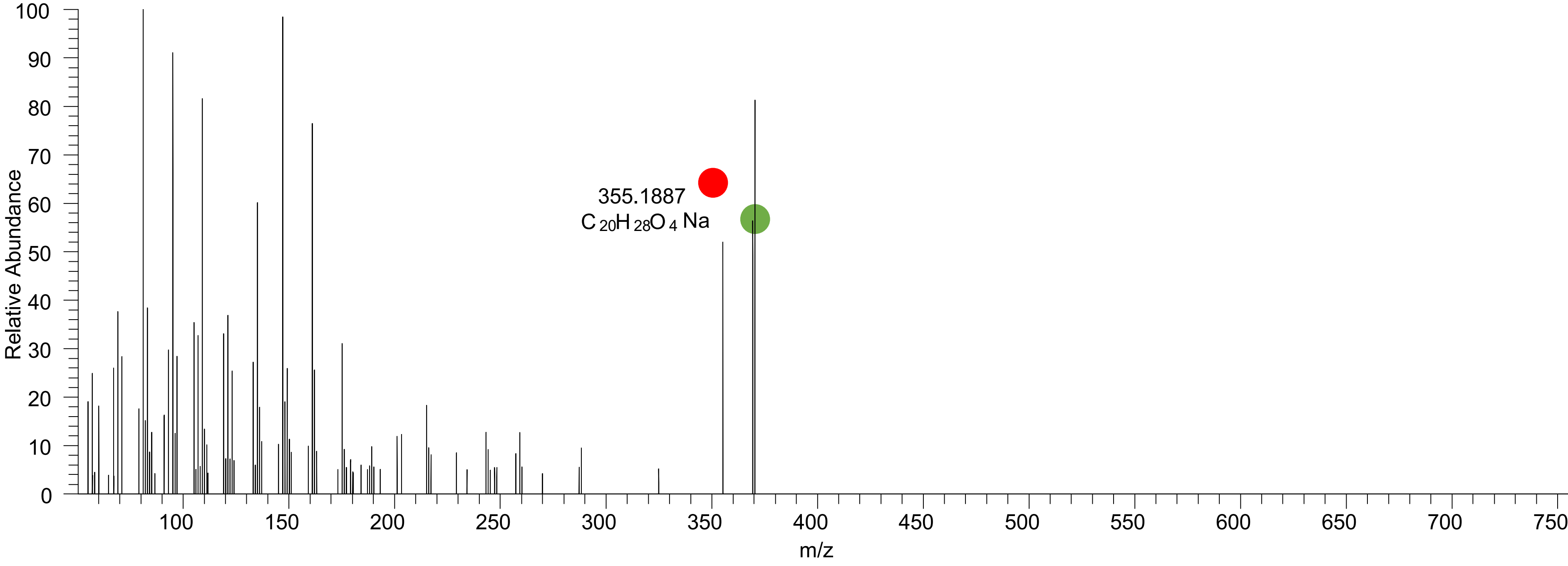

### CE(20:3<2oxo>)
# RT 22.0

[oxCE+Na]+ XIC 723.5323 NL: 1.07E5

Fragments containing oxFAs
Fragments related to water loss
Fragments not containing oxFAs
Position-specific fragments
Fragments related to other oxLPPs

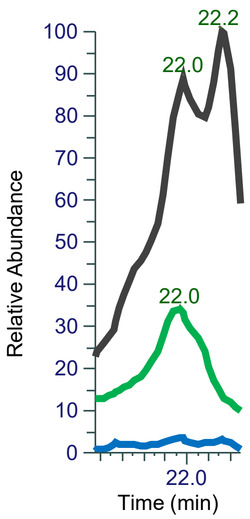


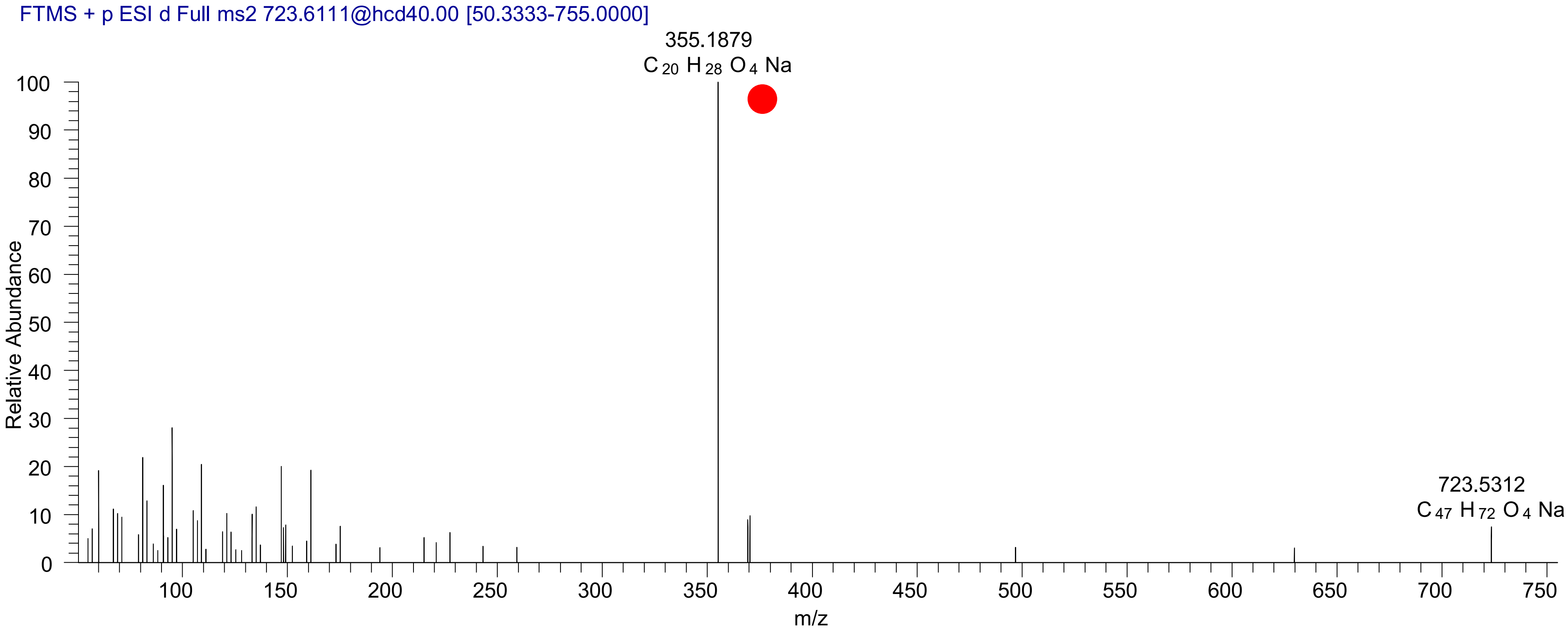

# CE(20:5<2OH>)
# RT 20.4

[oxCE+Na]+ XIC 725.5479 NL: 7.31E5

- Fragments containing oxFAs
- Fragments related to water loss
- Fragments not containing oxFAs
- Position-specific fragments
- Fragments related to other oxLPPs

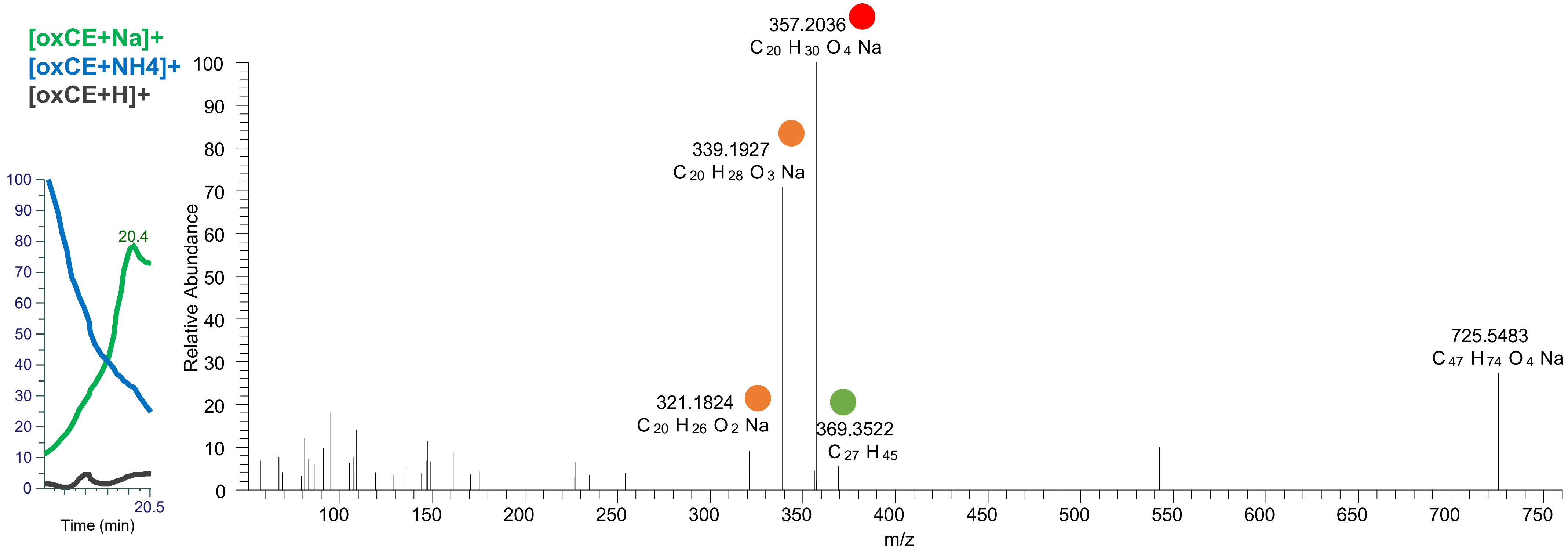

# CE(20:5<OH{9},OH>)
# RT 20.6

**[oxCE+Na]+ XIC 725.5479 NL: 7.57E5**

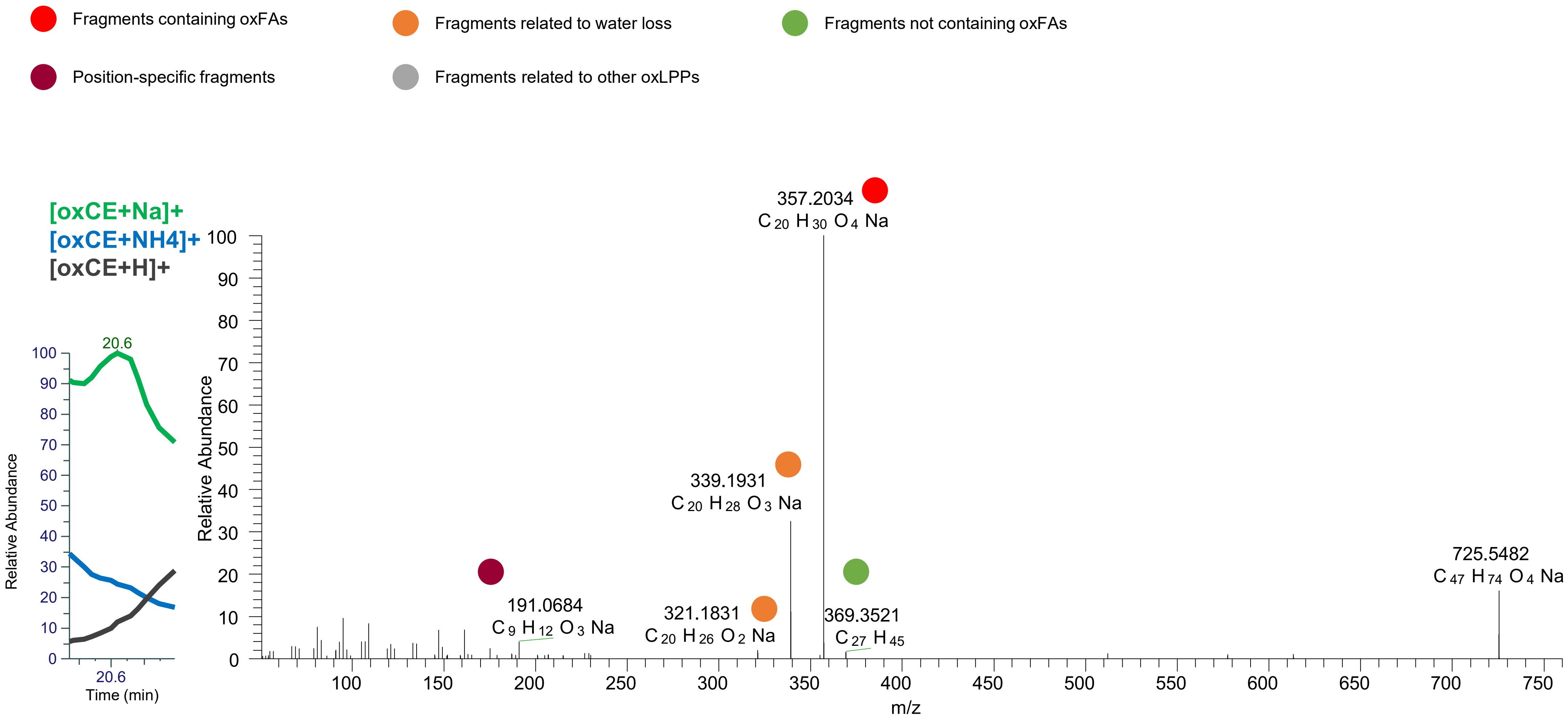

### CE(20:5<OOH{9}>)
### CE(20:5<OOH{12}>)
# RT 22.9

**[oxCE+Na]+** **XIC 725.5479 NL: 5.07E5**

- Fragments containing oxFAs
- Fragments related to water loss
- Fragments not containing oxFAs
- Position-specific fragments
- Fragments related to other oxLPPs

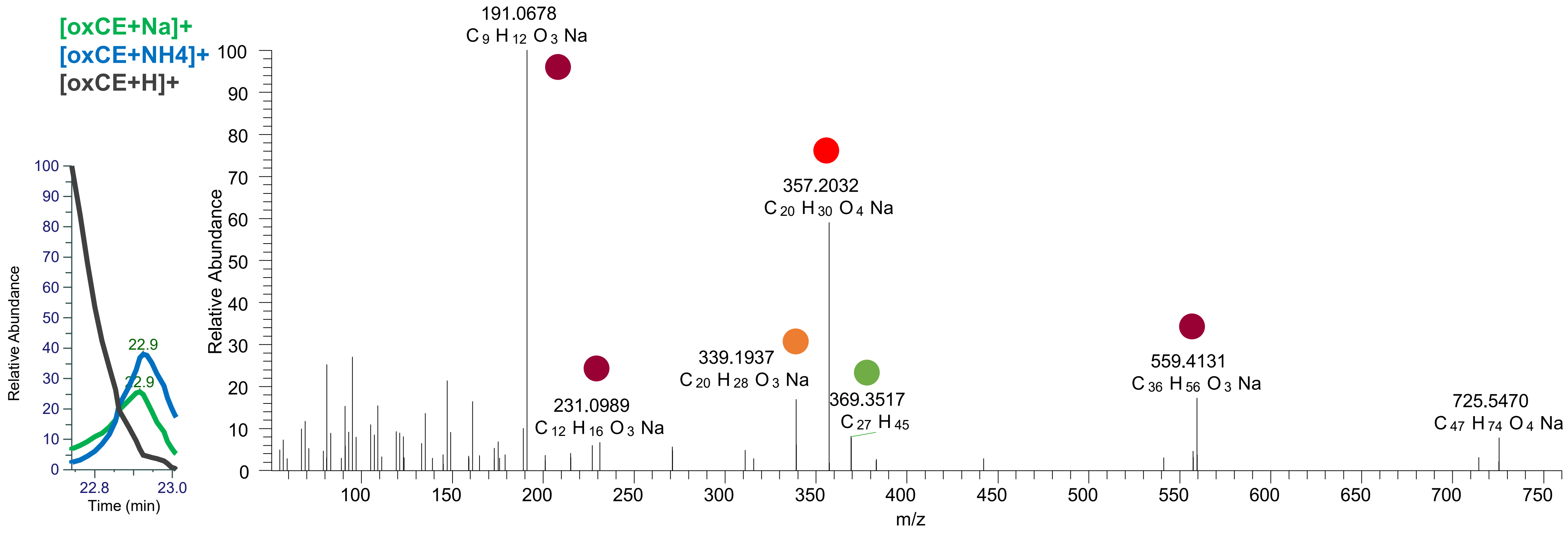

# CE(20:4<2OH>)
# RT 20.4

**[oxCE+Na]+ XIC 727.5636 NL: 2.84E5**

Fragments containing oxFAs
Fragments related to water loss
Fragments not containing oxFAs
Position-specific fragments
Fragments related to other oxLPPs

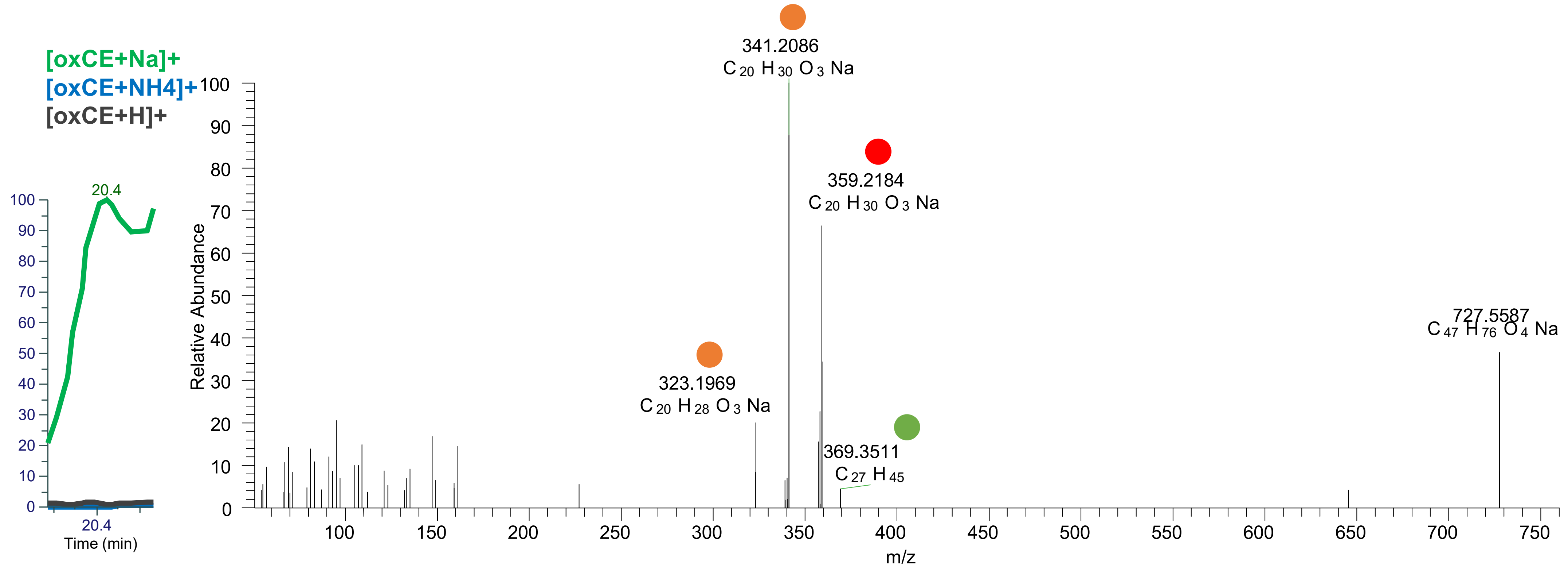

# CE(20:4<OH{9},OH>)
# RT 21.1

**[oxCE+Na]+ XIC 727.5636 NL: 3.36E6**

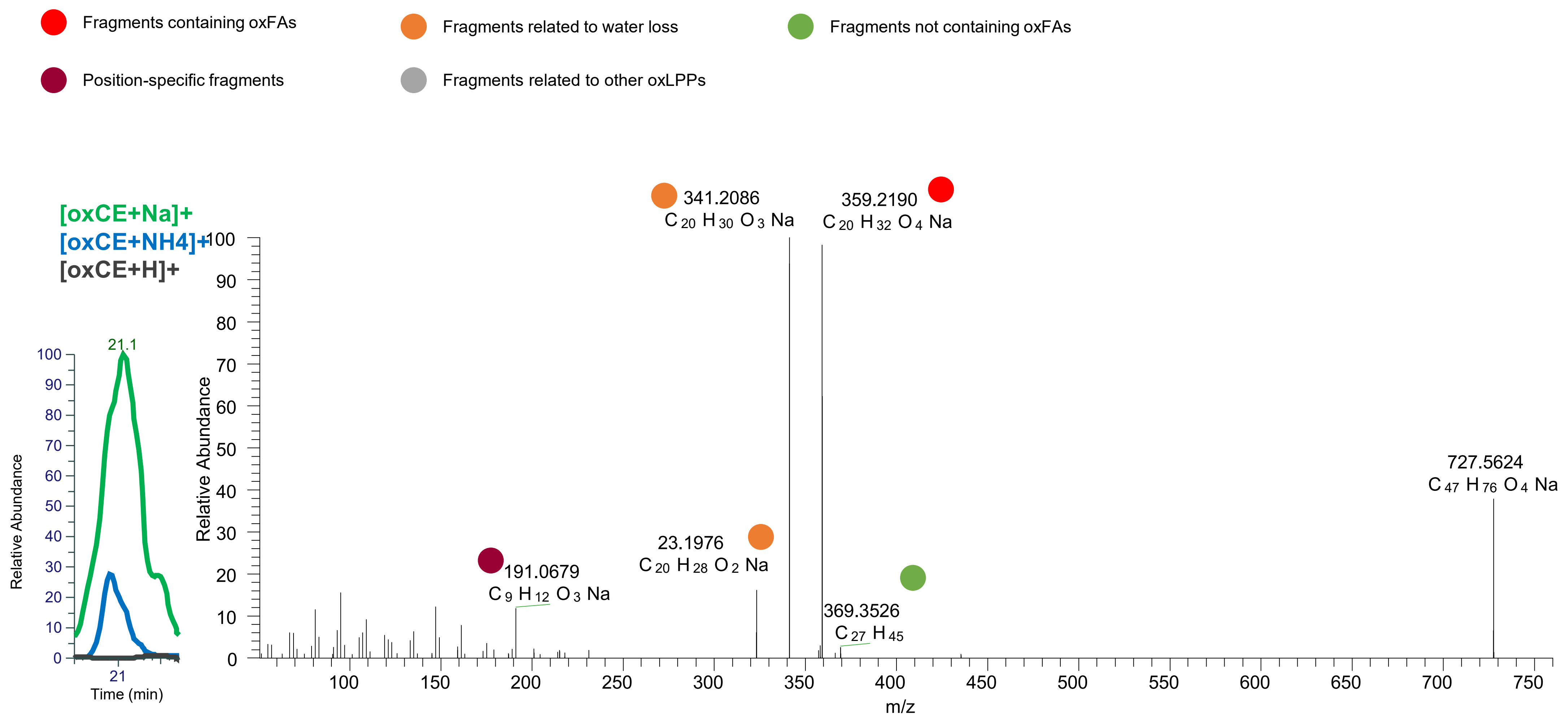

# CE(20:4<OH{9},OH>)
# RT 21.4

**[oxCE+Na]+** **XIC 727.5636 NL: 1.07E6**

- Fragments containing oxFAs
- Fragments related to water loss
- Fragments not containing oxFAs
- Position-specific fragments
- Fragments related to other oxLPPs

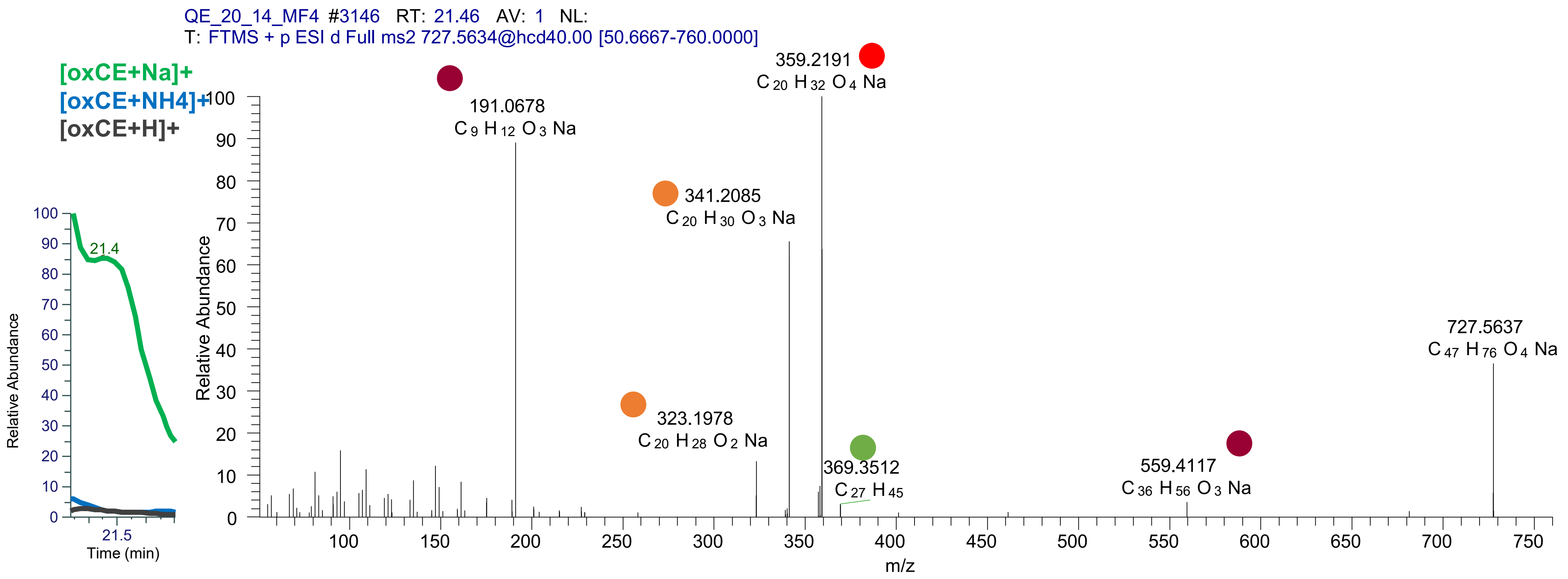

# CE(20:4<OH{9},OH>)
# RT 21.9

**[oxCE+Na]+** **XIC 727.5636 NL: 1.16E6**

- Fragments containing oxFAs
- Fragments related to water loss
- Fragments not containing oxFAs
- Position-specific fragments
- Fragments related to other oxLPPs

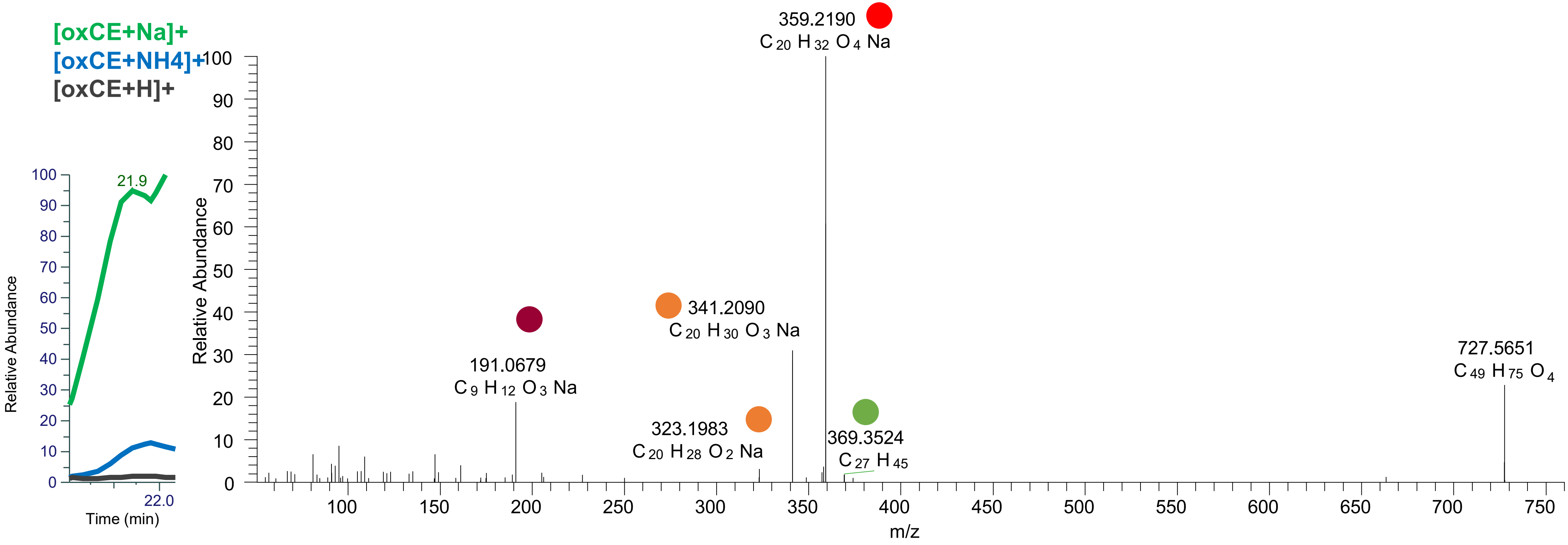

# CE(20:4<OH,O{10}>)
# RT 22.3

**[oxCE+Na]+** **XIC 727.5636 NL: 1.03E6**

- Fragments containing oxFAs
- Fragments related to water loss
- Fragments not containing oxFAs
- Position-specific fragments
- Fragments related to other oxLPPs

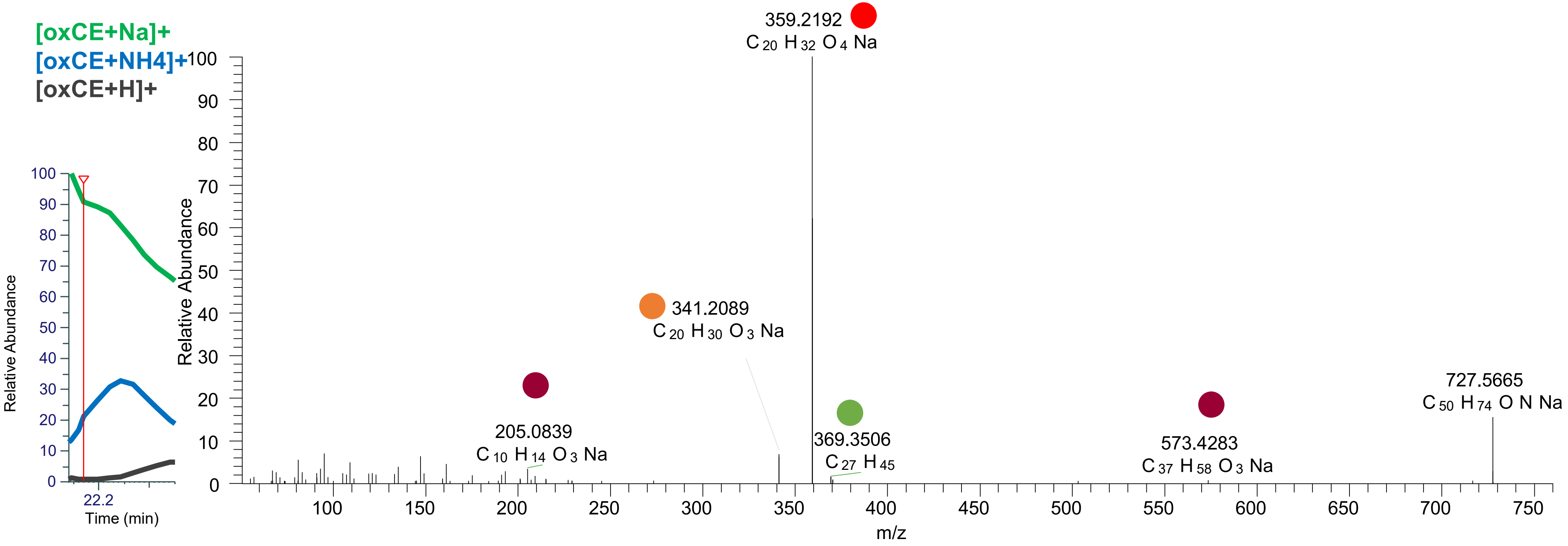

# CE(20:3<OH,O{9}>)
# RT 22.4

**[oxCE+Na]+** **XIC 727.5636 NL: 1.03E6**

- Fragments containing oxFAs
- Fragments related to water loss
- Fragments not containing oxFAs
- Position-specific fragments
- Fragments related to other oxLPPs

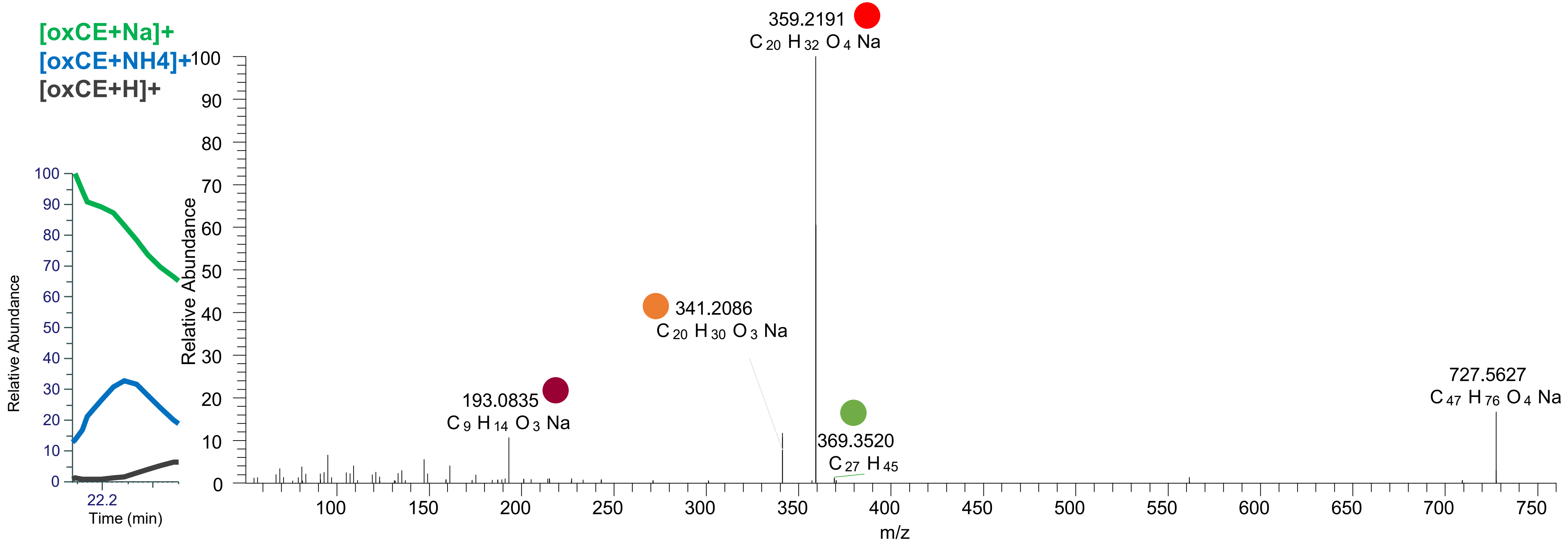

### CE(20:4<OOH{15}>)
### CE(20:4<OOH{13}>)
# RT 23.2

**[oxCE+Na]+** **XIC 727.5636 NL: 3.78E5**

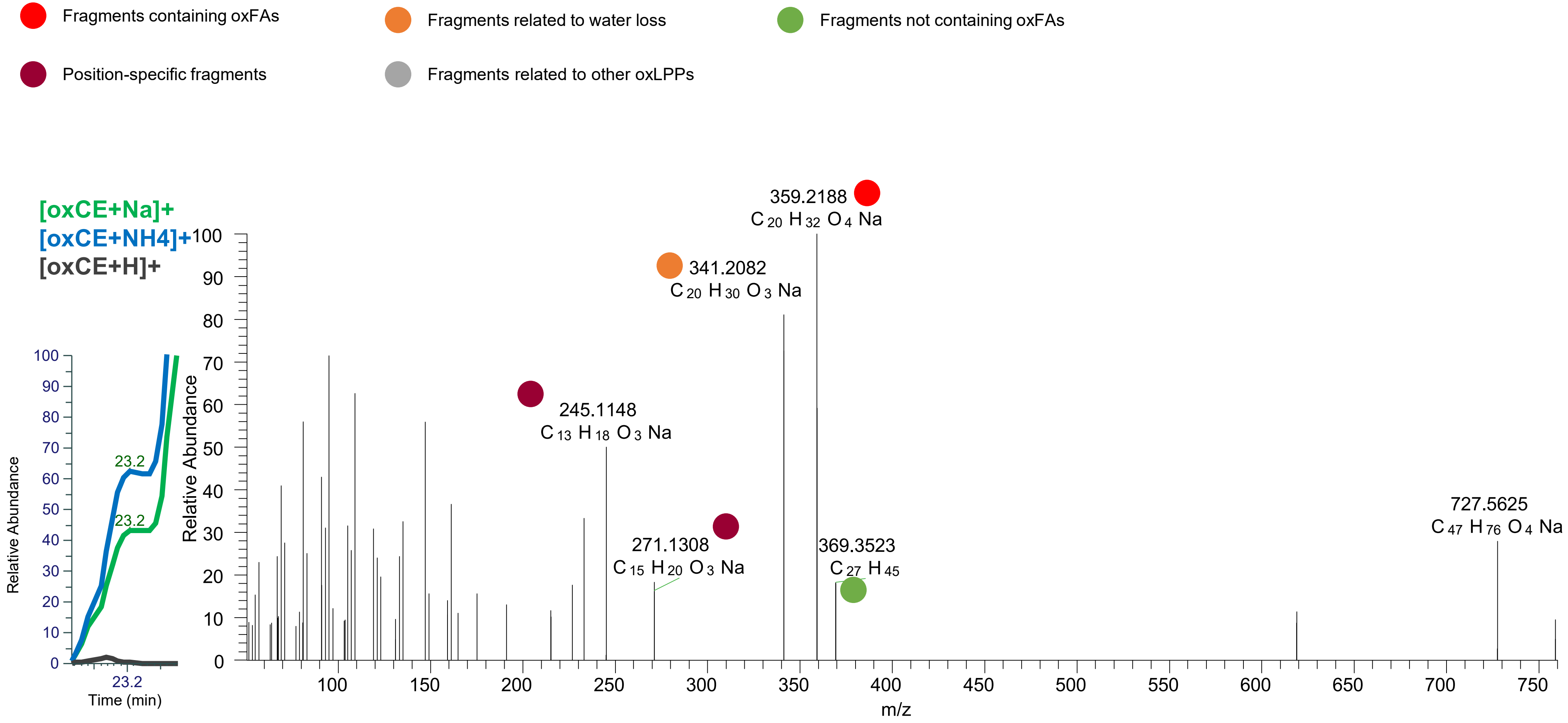

CE(20:4<OOH{15}>)
CE(20:4<OOH{12}>)
CE(20:4<OOH{9}>)
RT 23.4

[oxCE+Na]+ XIC 727.5636 NL: 6.07E6

- Fragments containing oxFAs
- Fragments related to water loss
- Fragments not containing oxFAs
- Position-specific fragments
- Fragments related to other oxLPPs

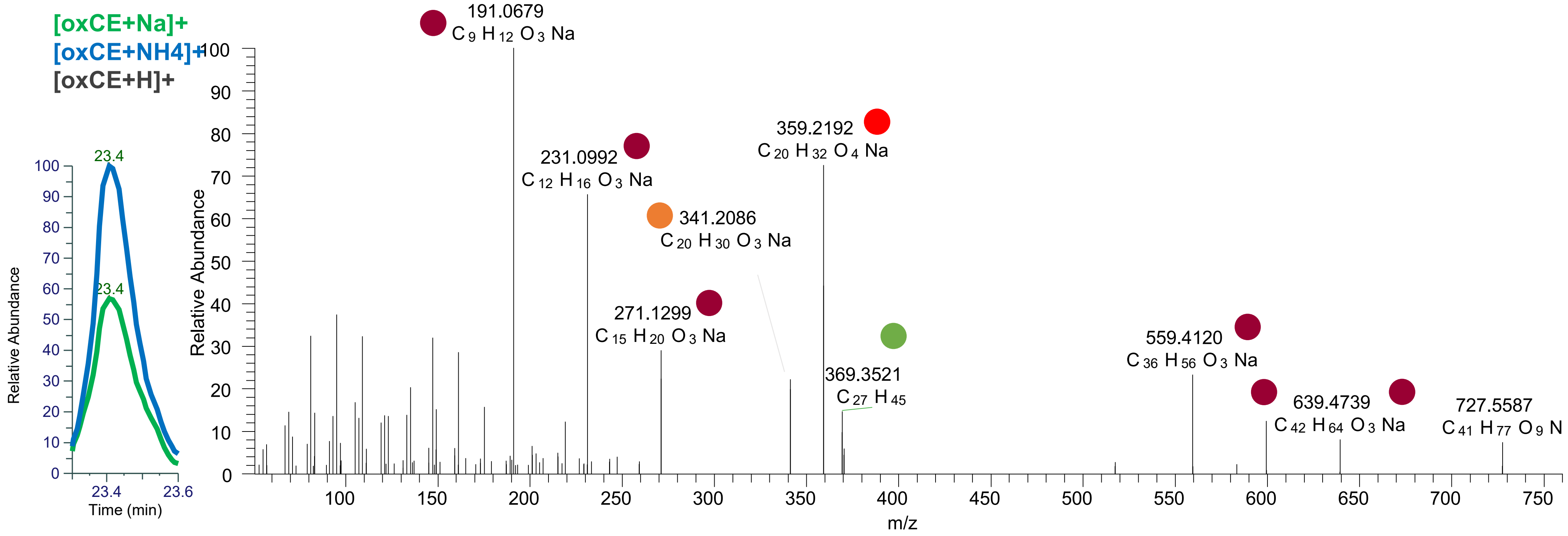

# CE(20:3<OH,O{9}>)
# RT 21.6

**[oxCE+Na]+** **XIC 729.5792 NL: 1.05E5**

Fragments containing oxFAs
Fragments related to water loss
Fragments not containing oxFAs
Position-specific fragments
Fragments related to other oxLPPs

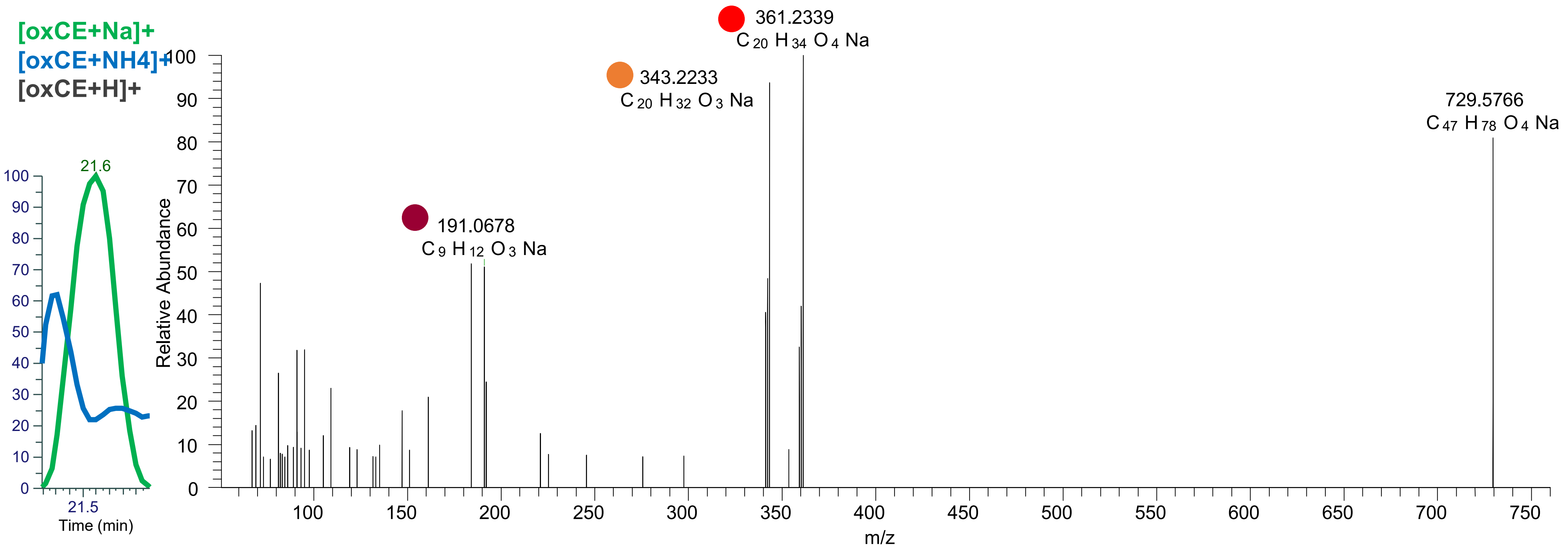

# CE(20:3<2OH>)
# RT 21.6

[oxCE+Na]+ XIC 729.5792 NL: 1.05E5

Fragments containing oxFAs
Fragments related to water loss
Fragments not containing oxFAs
Position-specific fragments
Fragments related to other oxLPPs

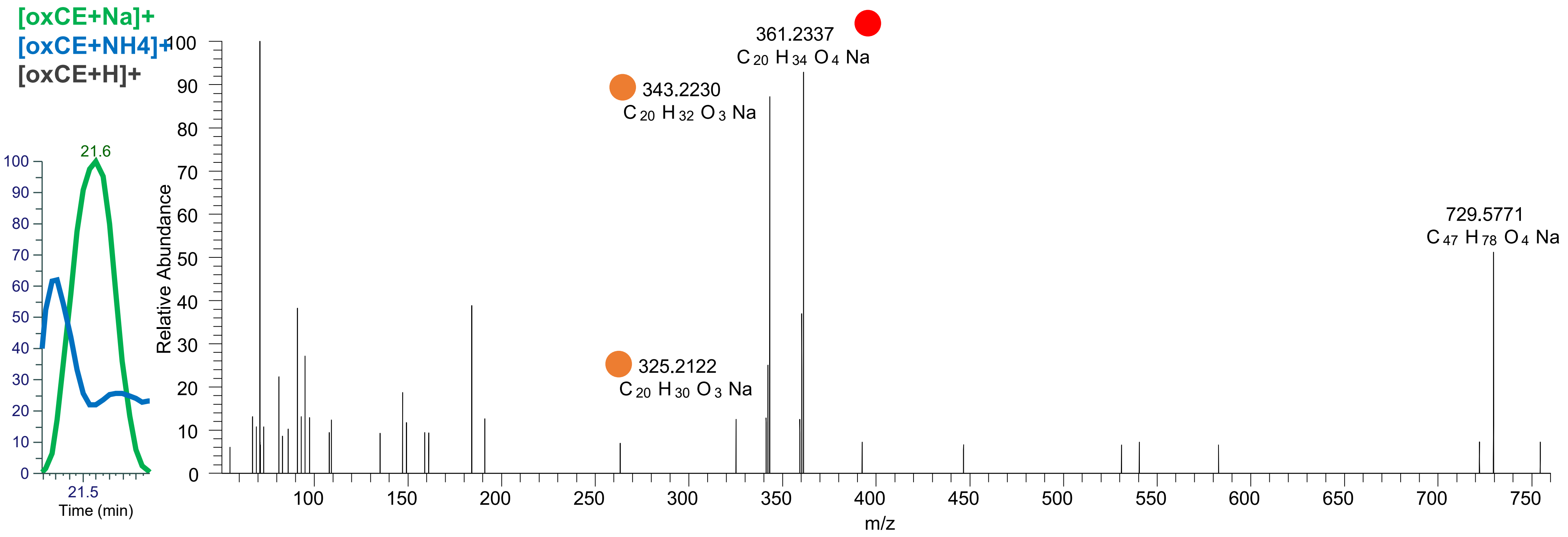

# CE(20:3<OH,O{9}>)
# RT 21.9

**[oxCE+Na]+** **XIC 729.5792 NL: 1.05E5**

- Fragments containing oxFAs
- Fragments related to water loss
- Fragments not containing oxFAs
- Position-specific fragments
- Fragments related to other oxLPPs

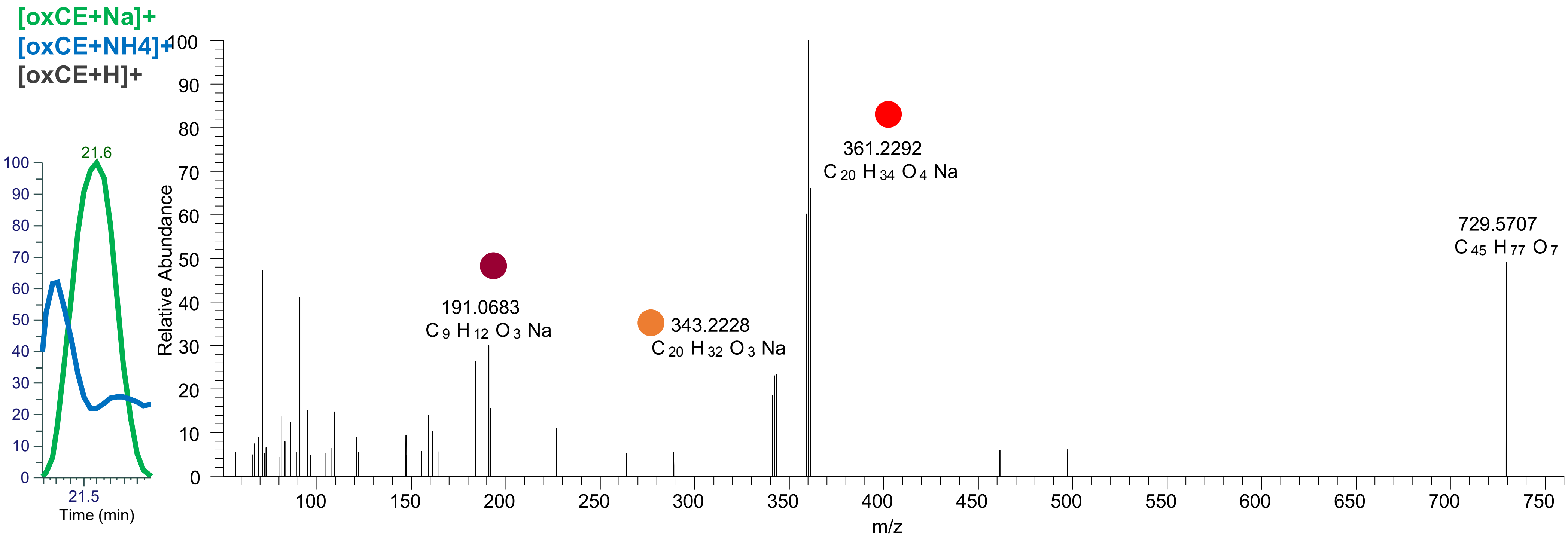

# CE(20:2<OH,ep>)
# RT 22.7

**[oxCE+Na]+** **XIC 729.5792 NL: 8.00E4**

- Fragments containing oxFAs
- Fragments related to water loss
- Fragments not containing oxFAs
- Position-specific fragments
- Fragments related to other oxLPPs

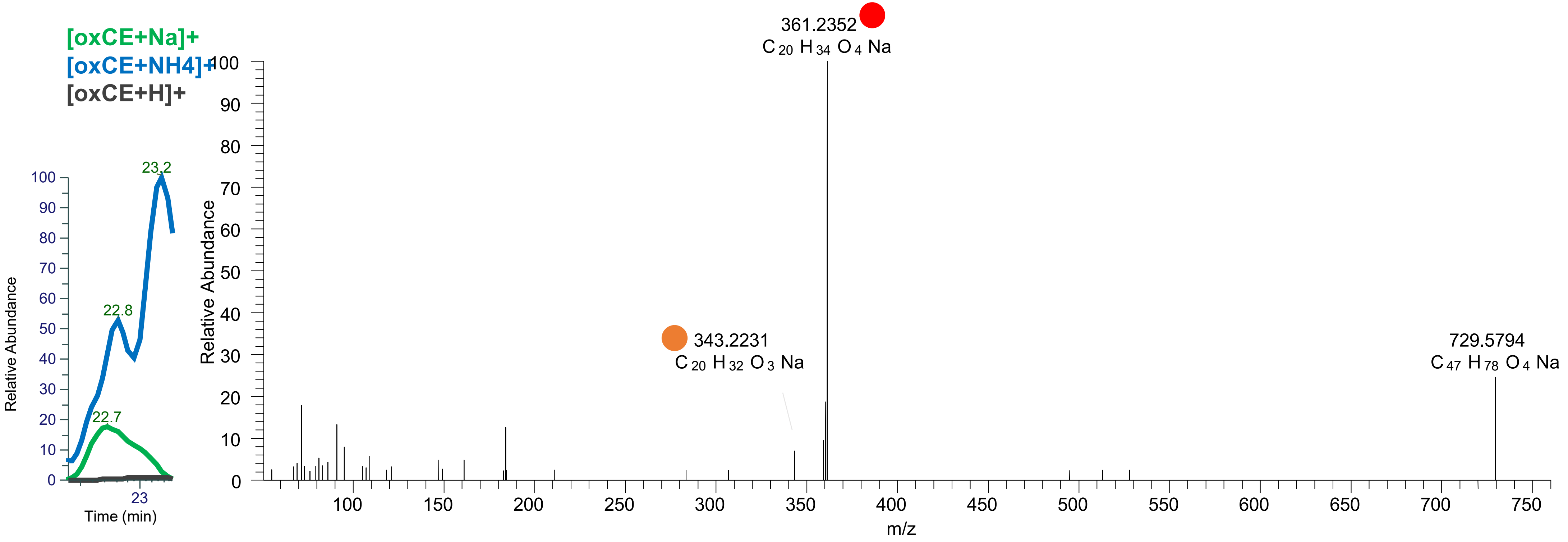

### CE(20:3<OOH{15}>)
### CE(20:3<OOH{11}>)
# RT 23.7

**[oxCE+Na]+** **XIC 729.5792 NL: 1.45E5**

- Fragments containing oxFAs
- Fragments related to water loss
- Fragments not containing oxFAs
- Position-specific fragments
- Fragments related to other oxLPPs

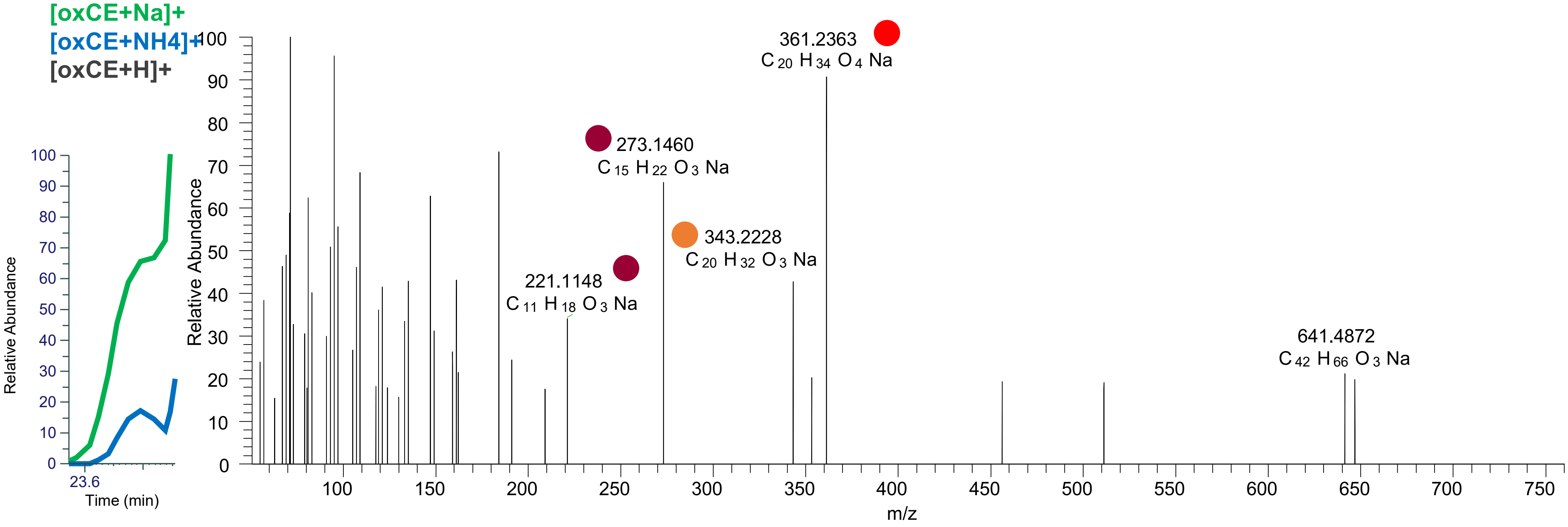

CE(20:3<OOH{15}>)
CE(20:3<OOH{12}>)
CE(20:3<OOH{11}>)
RT 23.9

[oxCE+Na]+ XIC 729.5792 NL: 8.16E5

Fragments containing oxFAs
Fragments related to water loss
Fragments not containing oxFAs
Position-specific fragments
Fragments related to other oxLPPs

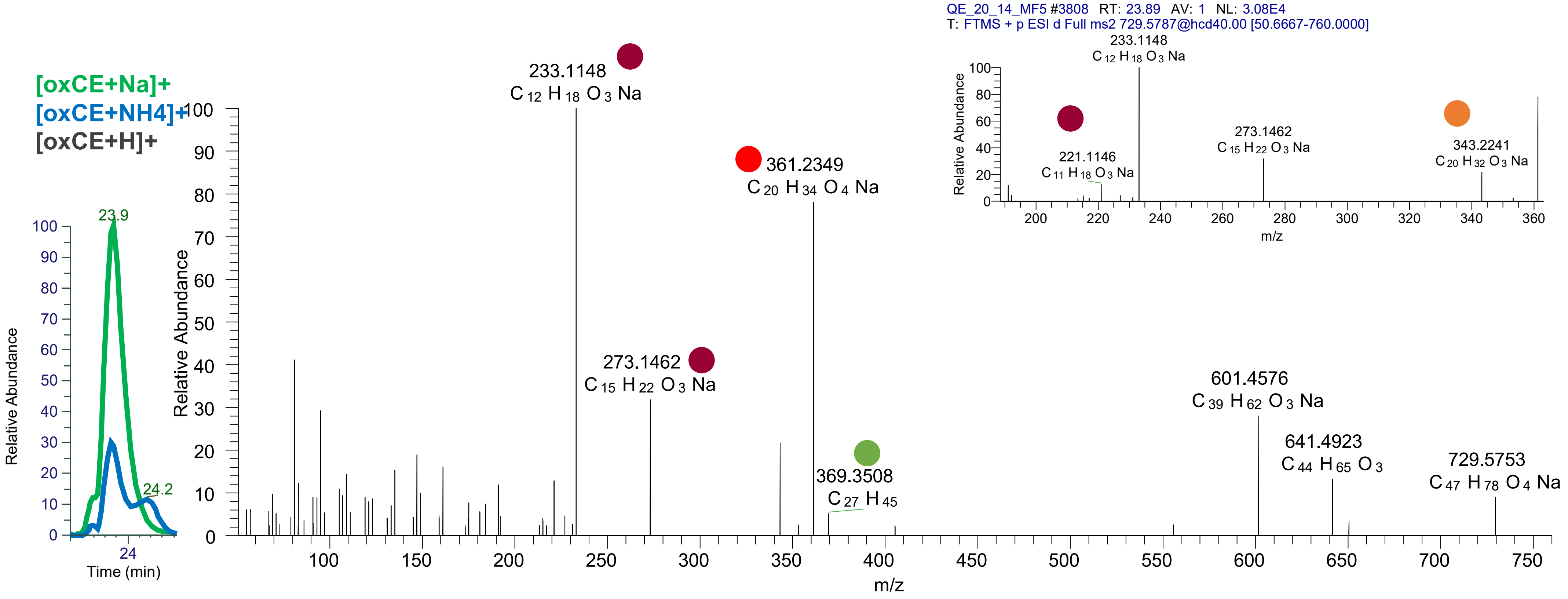

# CE(20:3<2O>)
# RT 24.2

[oxCE+Na]+ XIC 729.5792 NL: 8.16E5

Fragments containing oxFAs

Fragments related to water loss

Fragments not containing oxFAs

Position-specific fragments

Fragments related to other oxLPPs

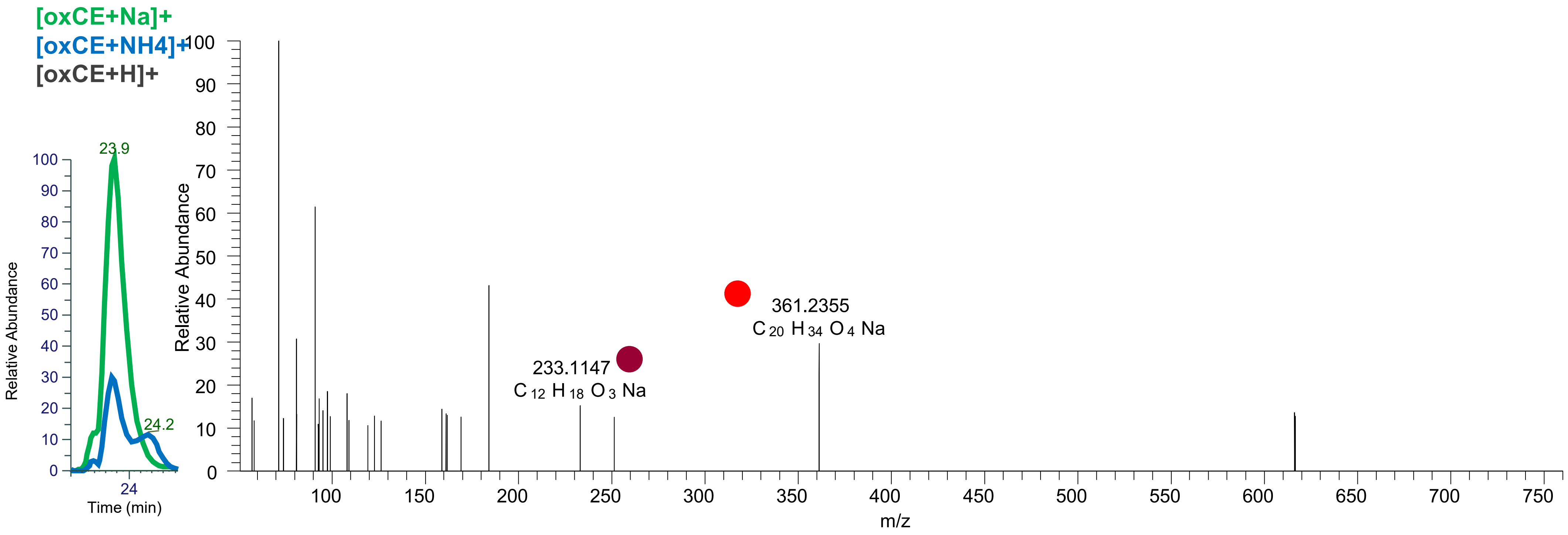

# CE(20:1<ep,O>)
# RT 24.5

**[oxCE+Na]+** **XIC 731.5949 NL: 2.81E4**

- Fragments containing oxFAs
- Fragments related to water loss
- Fragments not containing oxFAs
- Position-specific fragments
- Fragments related to other oxLPPs

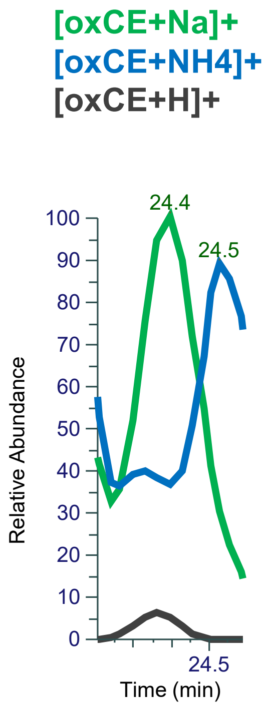


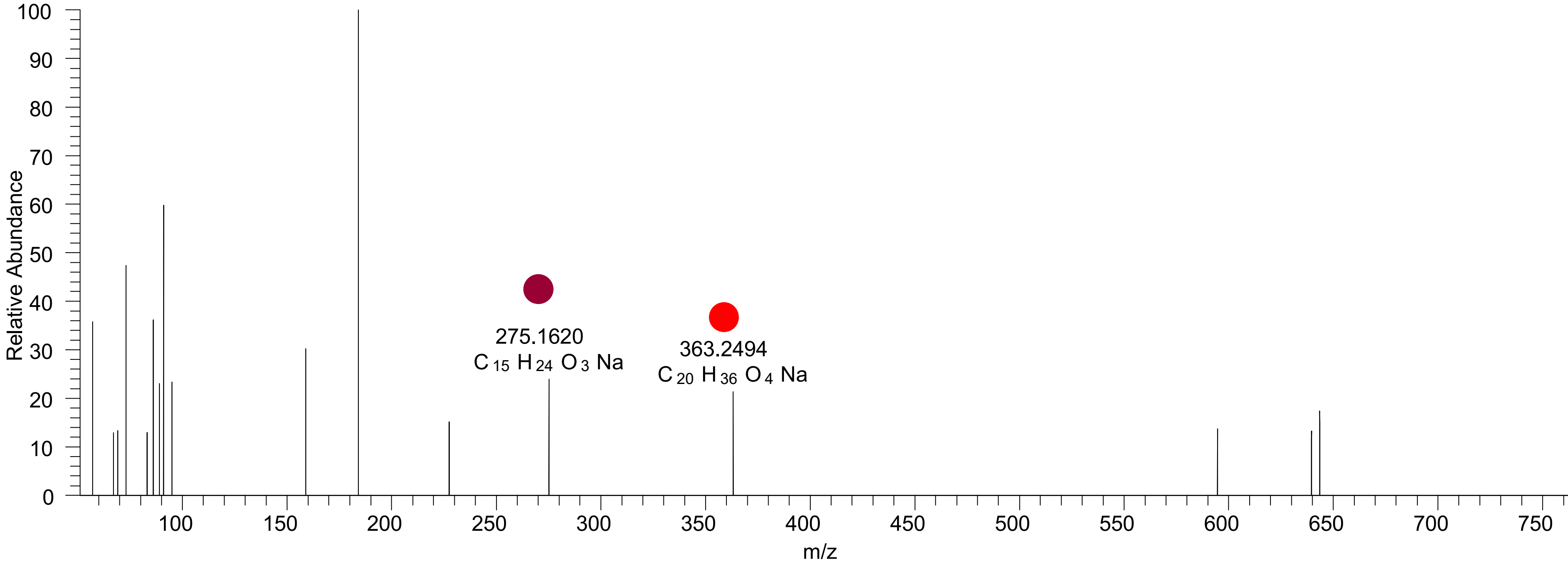

# CE(20:6<3O>)
# RT 20.4

**[oxCE+Na]+** **XIC 739.5272 NL: 2.51E4**

- Fragments containing oxFAs
- Fragments related to water loss
- Fragments not containing oxFAs
- Position-specific fragments
- Fragments related to other oxLPPs

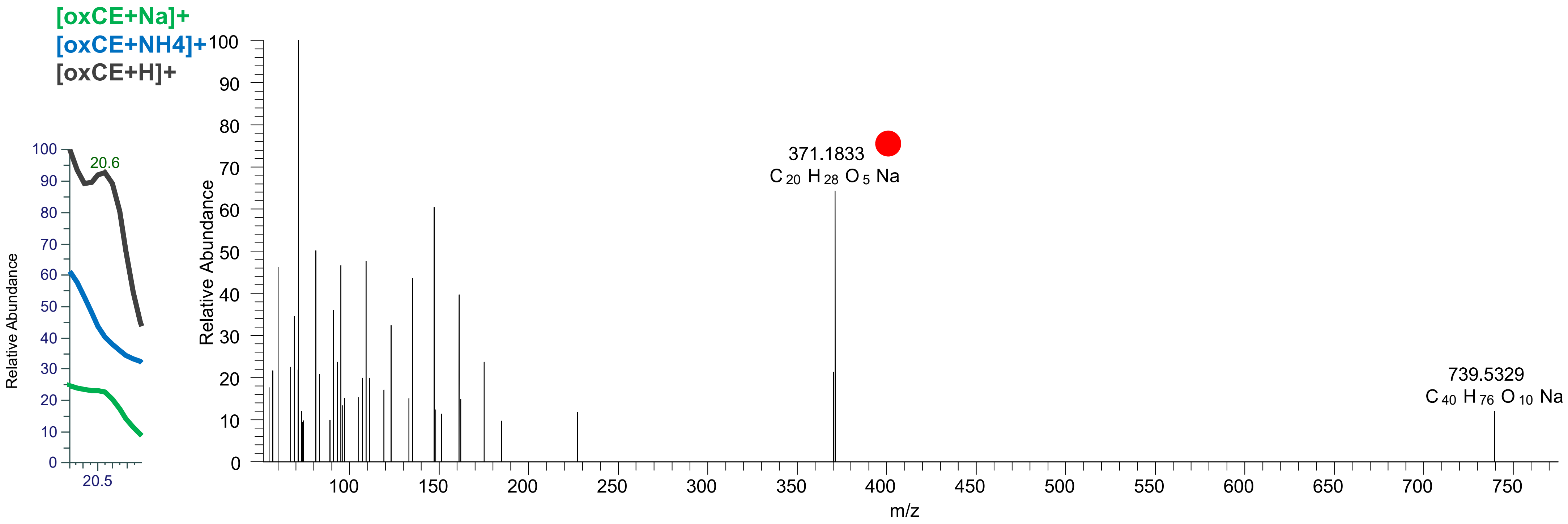

# CE(20:6<3O>)
# RT 21.6

**[oxCE+Na]+** **XIC 739.5272 NL: 8.20E4**

- Fragments containing oxFAs
- Fragments related to water loss
- Fragments not containing oxFAs
- Position-specific fragments
- Fragments related to other oxLPPs

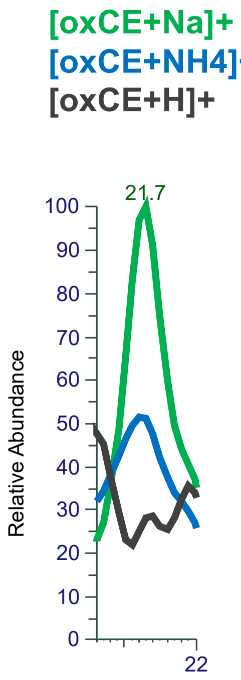


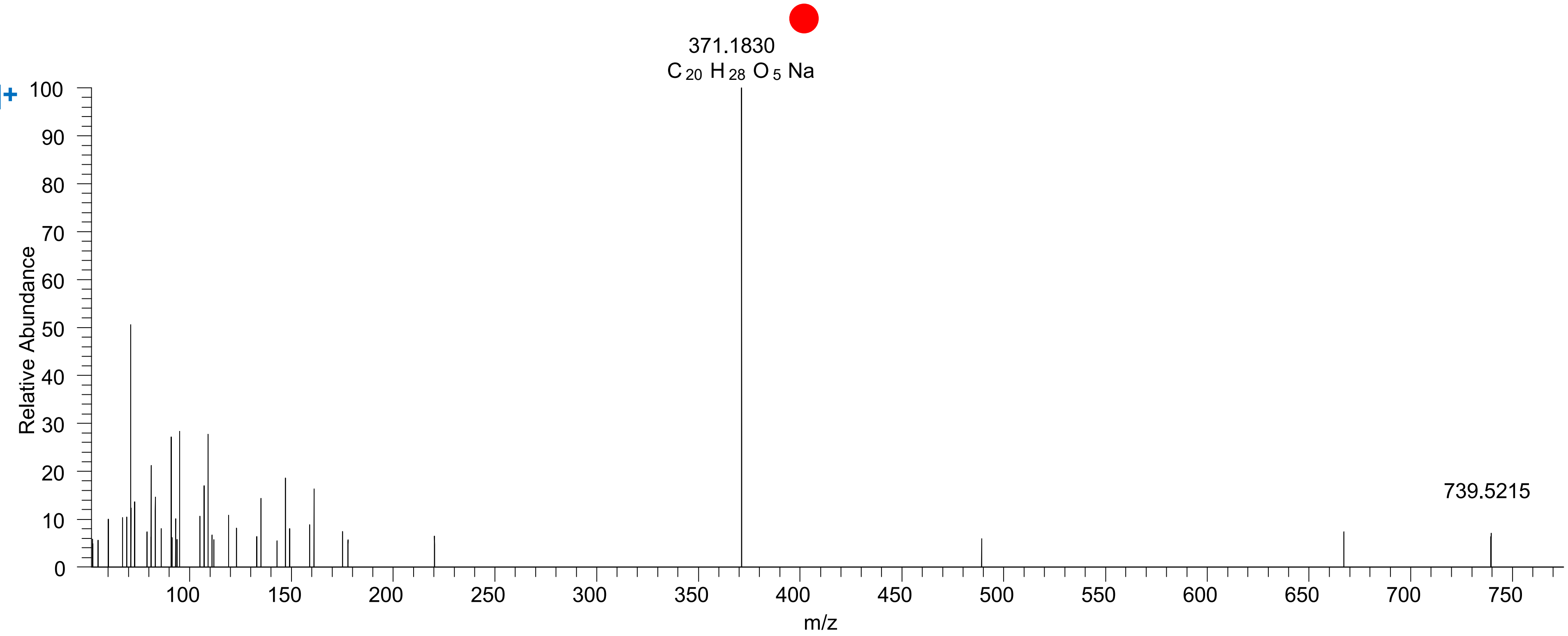

# CE(22:4<O>)
# RT 24.4

[oxCE+Na]+ XIC 739.6000 NL: 2.37E4

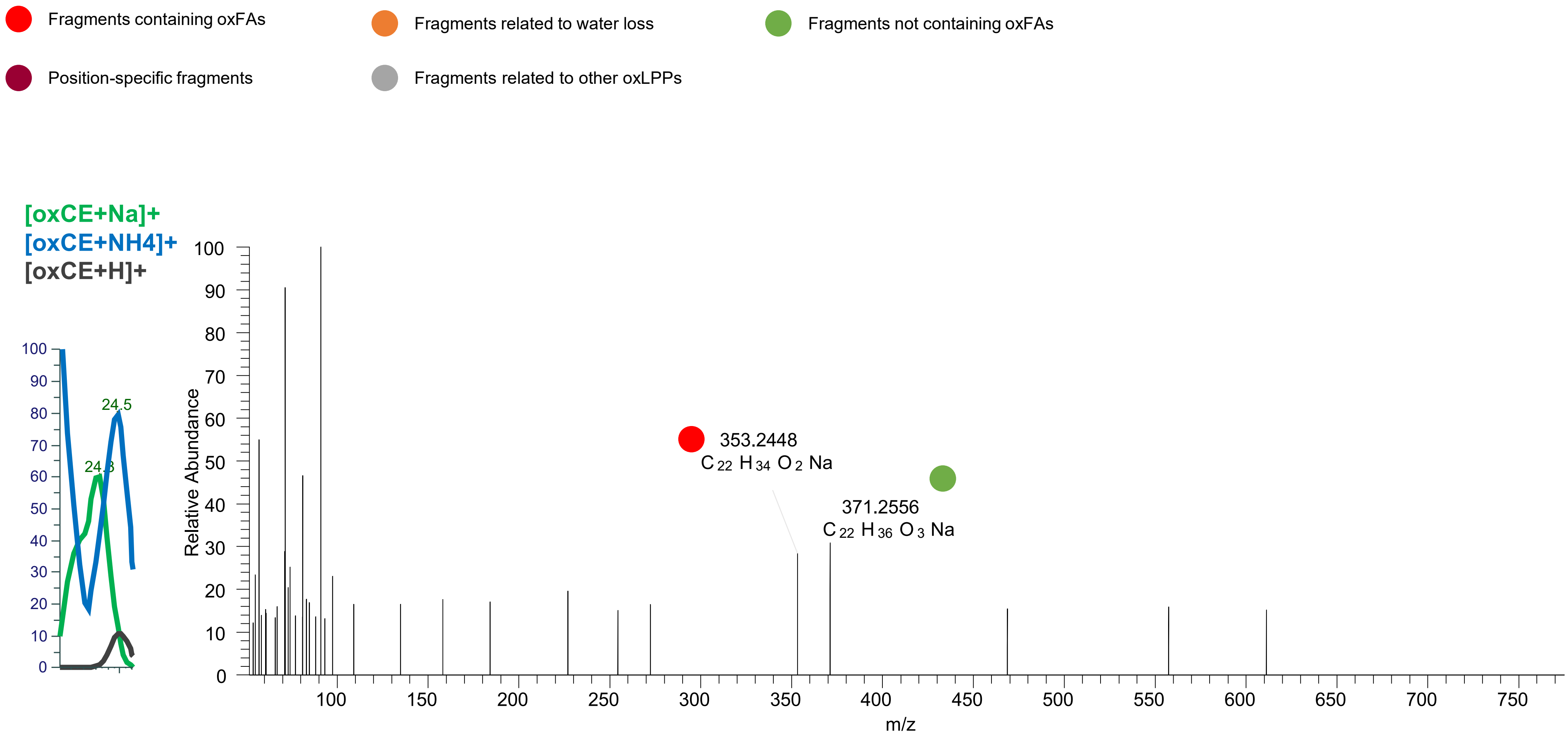

CE(20:5<OH,OOH{15}>)
CE(20:5<O,OOH{9}>)
RT 20.5

[oxCE+Na]+ XIC 741.5428 NL: 3.16E5

- Fragments containing oxFAs
- Fragments related to water loss
- Fragments not containing oxFAs
- Position-specific fragments
- Fragments related to other oxLPPs

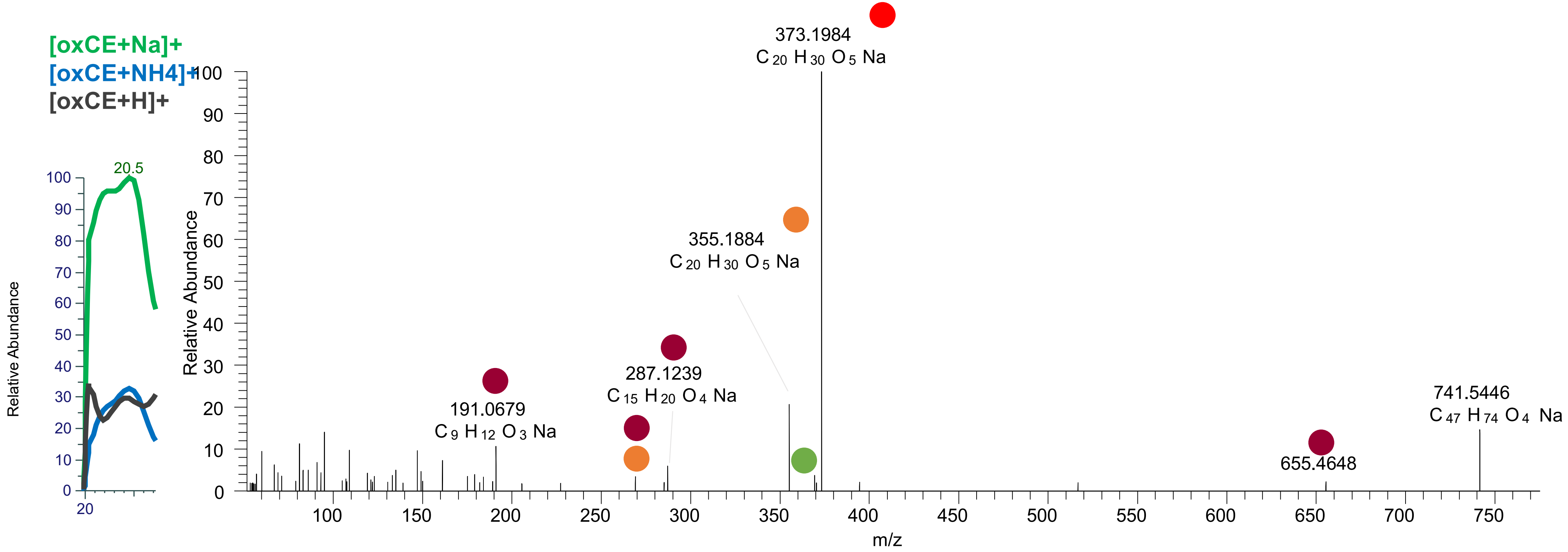

### CE(20:4<oxo,OH,O>)
# RT 21.1

**[oxCE+Na]+** **XIC 741.5428 NL: 2.18E5**

Fragments containing oxFAs

Fragments related to water loss

Fragments not containing oxFAs

Position-specific fragments

Fragments related to other oxLPPs

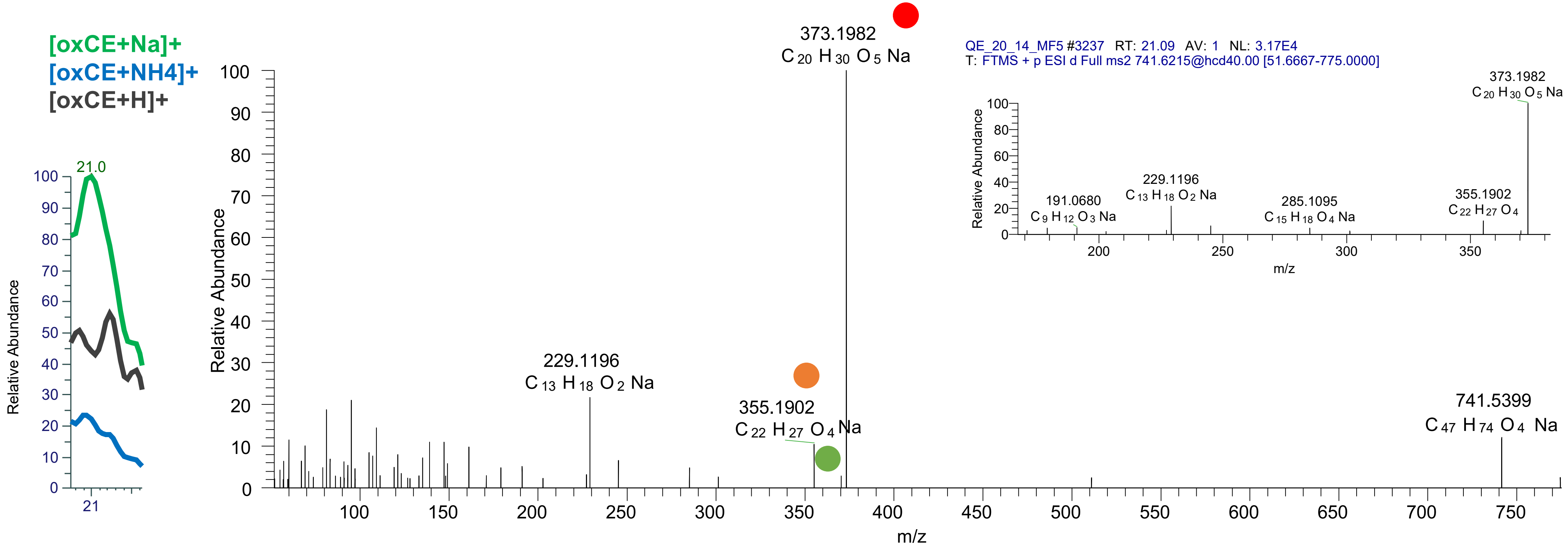

### CE(20:3<oxo,OH,O>)
# RT 21.9

**[oxCE+Na]+ XIC 741.5428 NL: 1.01E5**

Fragments containing oxFAs
Fragments related to water loss
Fragments not containing oxFAs
Position-specific fragments
Fragments related to other oxLPPs

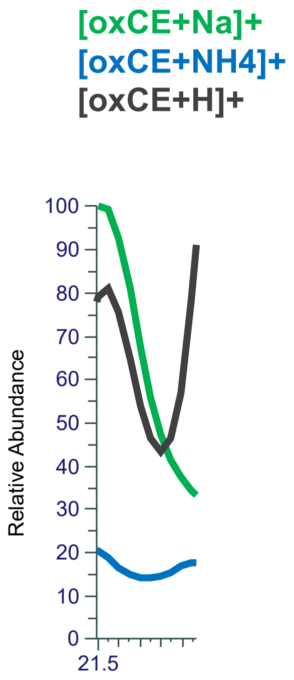


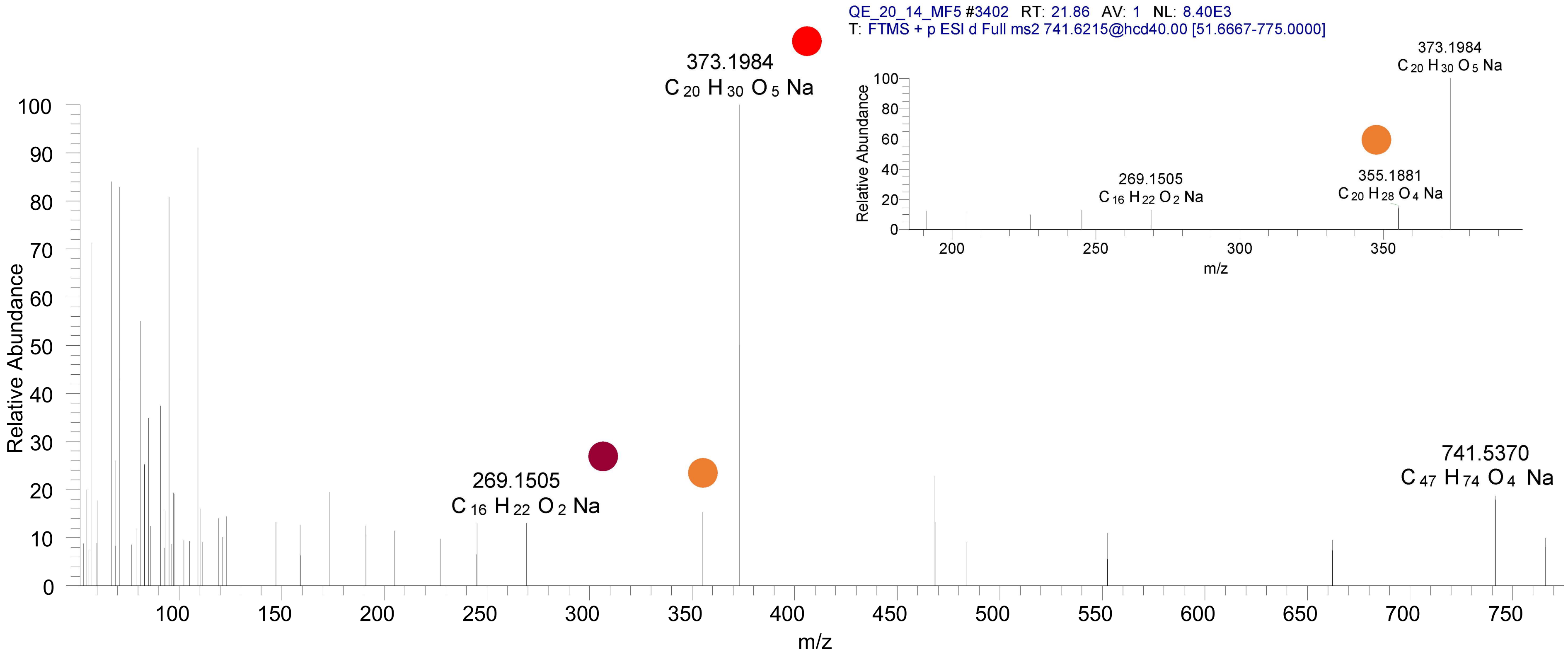

# CE(20:4<2OH,O>)
# RT 19.5

**[oxCE+Na]+** **XIC 743.5585 NL: 2.40E5**

- Fragments containing oxFAs
- Fragments related to water loss
- Fragments not containing oxFAs
- Position-specific fragments
- Fragments related to other oxLPPs

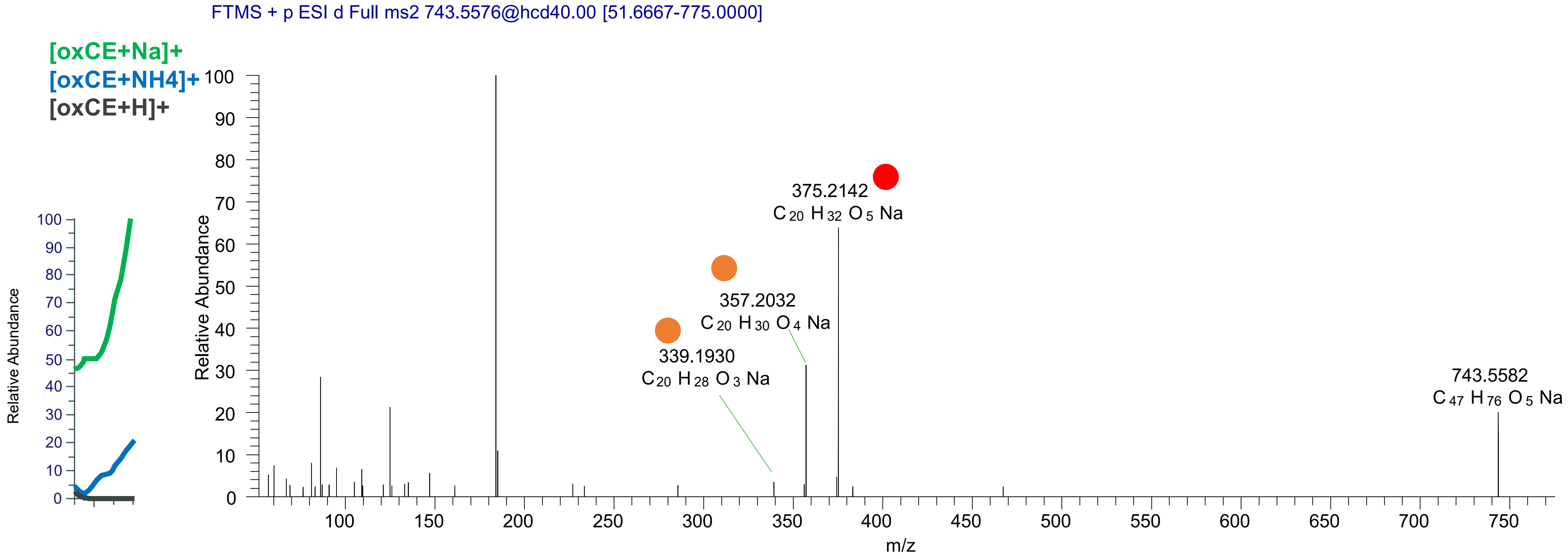

# CE(20:4<2OH,O>)
# RT 20.1

[oxCE+Na]+ XIC 743.5585 NL: 6.46E5

- Fragments containing oxFAs
- Fragments related to water loss
- Fragments not containing oxFAs
- Position-specific fragments
- Fragments related to other oxLPPs

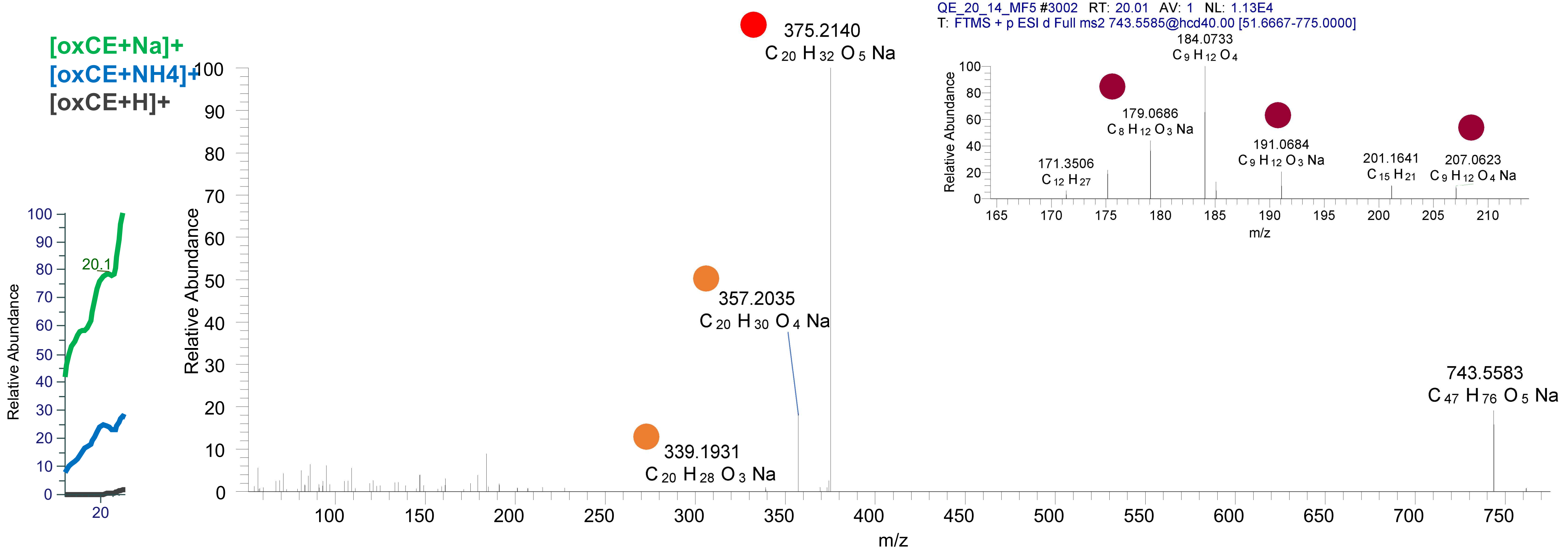

# CE(20:4<2OH,O>)
# RT 20.7

[oxCE+Na]+ XIC 743.5585 NL: 1.62E6

- Fragments containing oxFAs
- Fragments related to water loss
- Fragments not containing oxFAs
- Position-specific fragments
- Fragments related to other oxLPPs

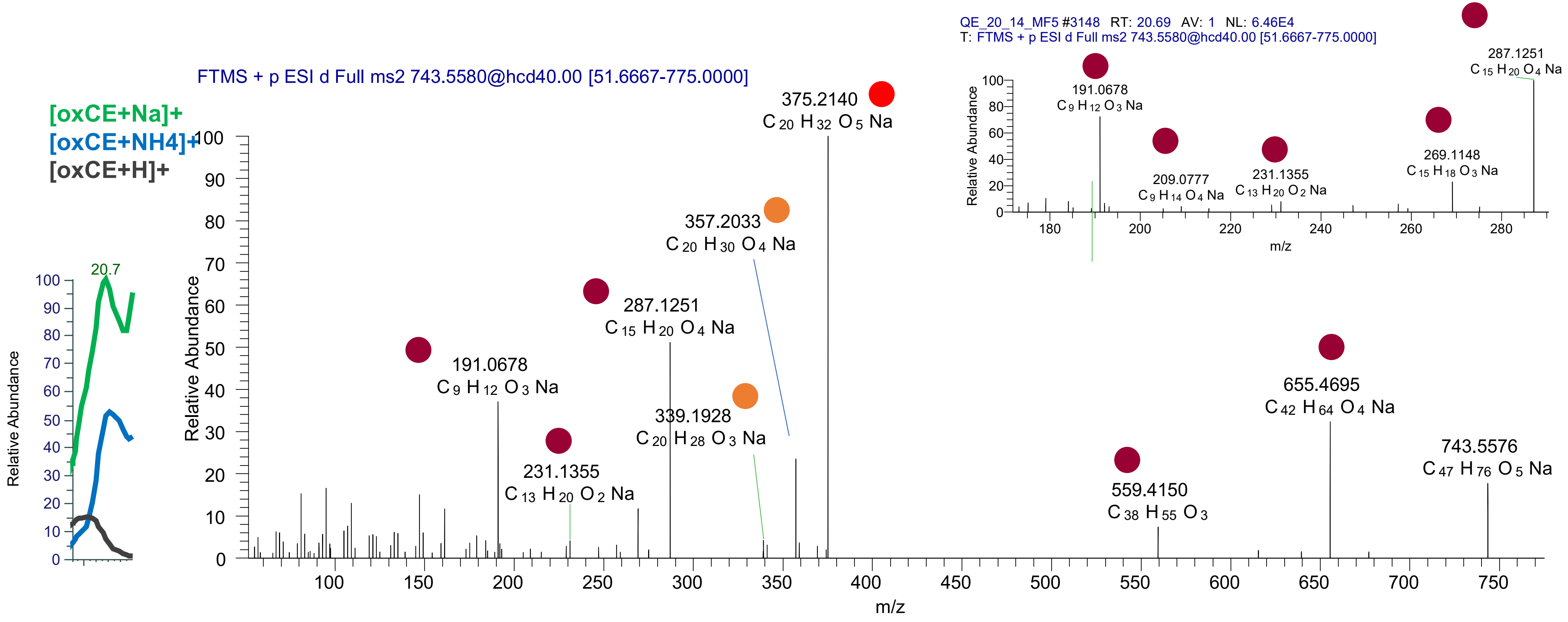

# CE(20:4<2OH,O>)
# RT 21.1

**[oxCE+Na]+** **XIC 743.5585 NL: 1.88E6**

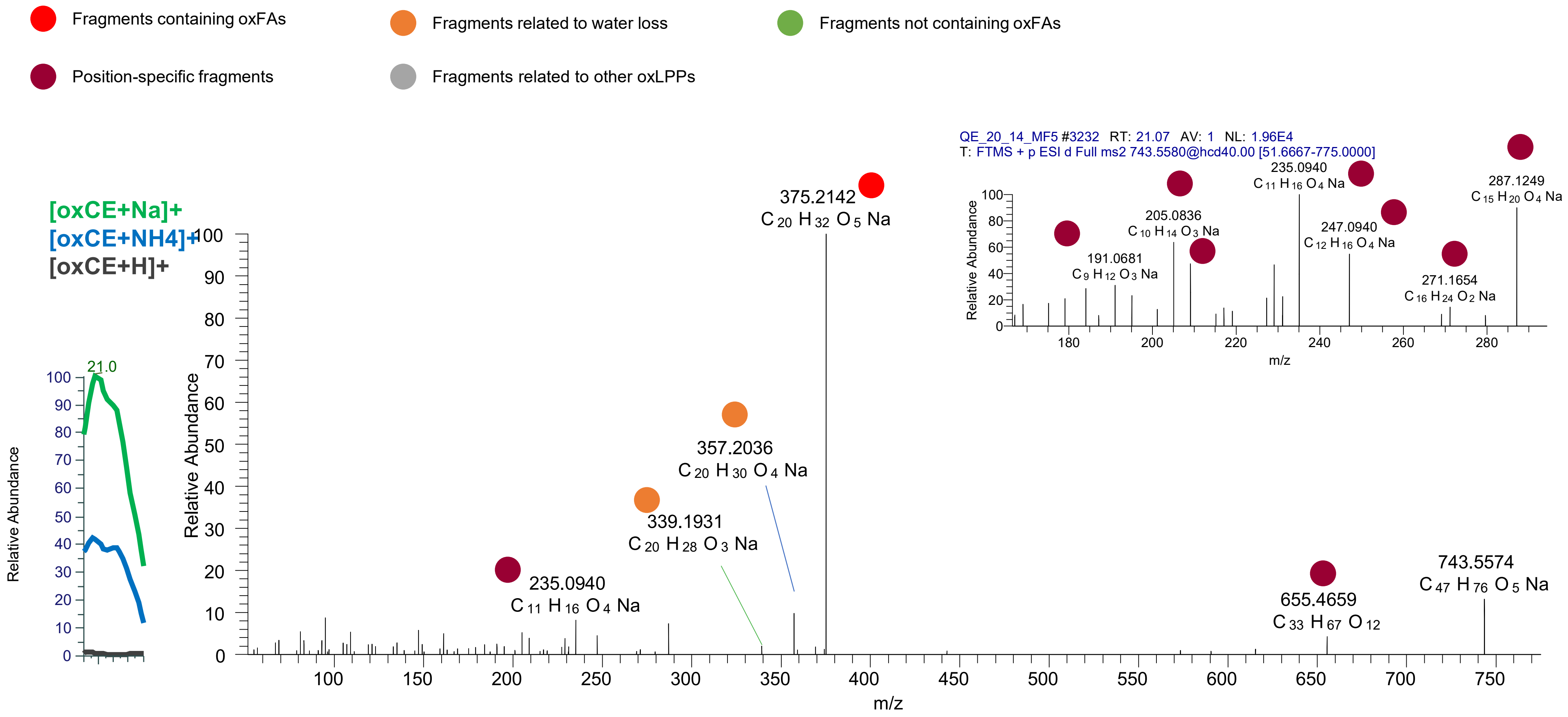

# CE(20:4<2OH,O>)
# RT 21.5

**[oxCE+Na]+        XIC 743.5585 NL: 4.29E5**

- Fragments containing oxFAs
- Fragments related to water loss
- Fragments not containing oxFAs
- Position-specific fragments
- Fragments related to other oxLPPs

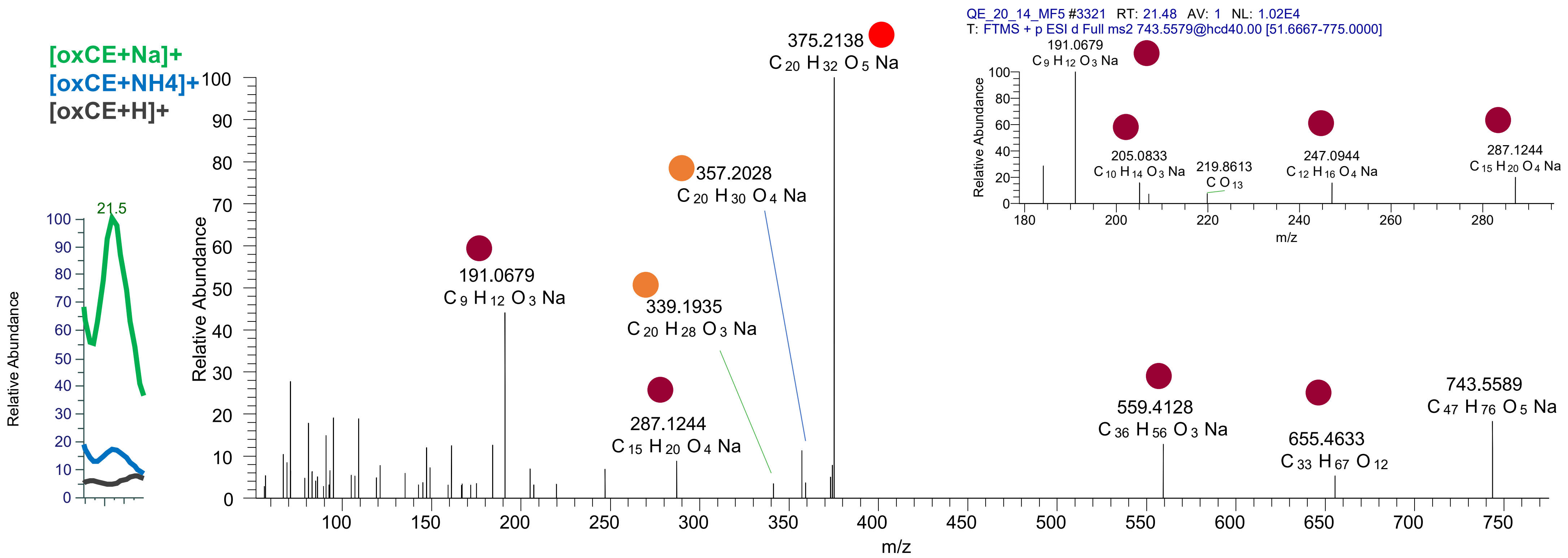

# CE(20:4<OH,2O>)
# RT 22.1

**[oxCE+Na]+ XIC 743.5585 NL: 1.49E5**

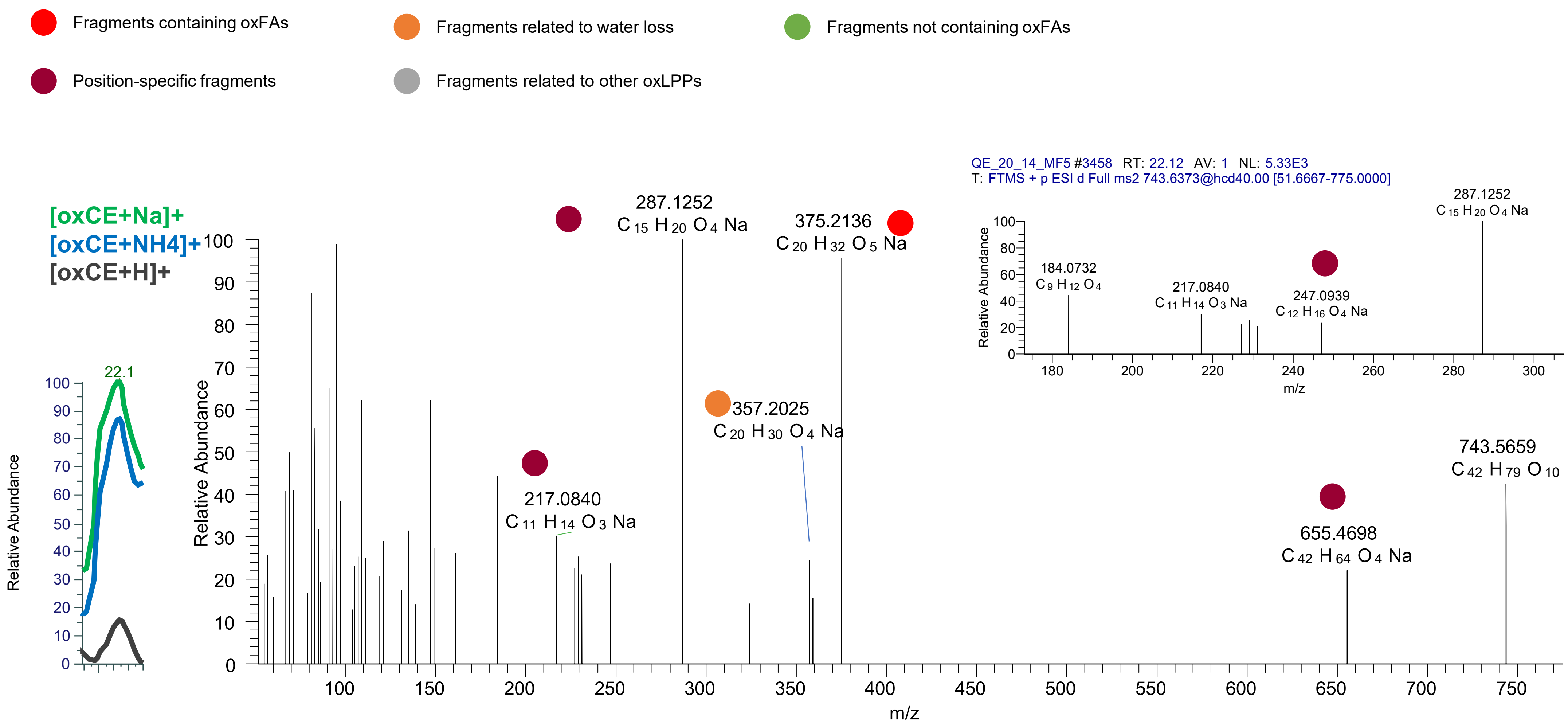

# CE(20:3<OH,ep,O>)
# RT 20.5

**[oxCE+Na]+**  **XIC 745.5741 NL: 5.43E4**

- Fragments containing oxFAs
- Fragments related to water loss
- Fragments not containing oxFAs
- Position-specific fragments
- Fragments related to other oxLPPs

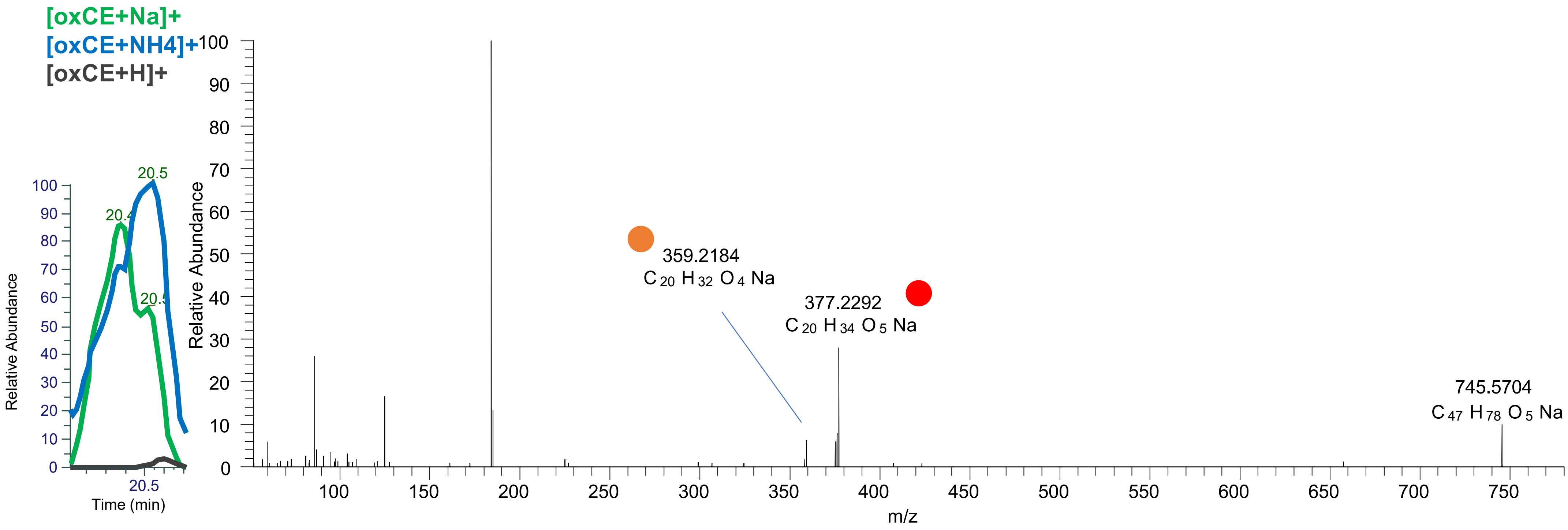

# CE(20:2<ep,OH,O>)
# RT 21.3

**[oxCE+Na]+** **XIC 745.5741 NL: 8.27E4**

- Fragments containing oxFAs
- Fragments related to water loss
- Fragments not containing oxFAs
- Position-specific fragments
- Fragments related to other oxLPPs

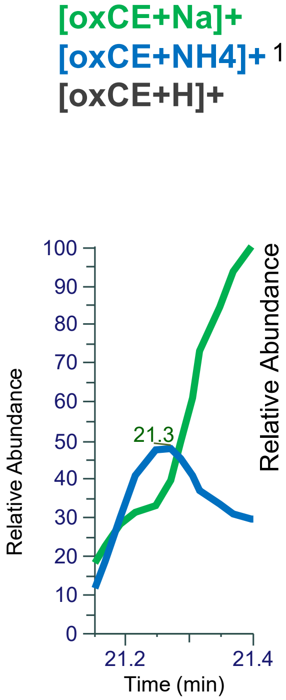


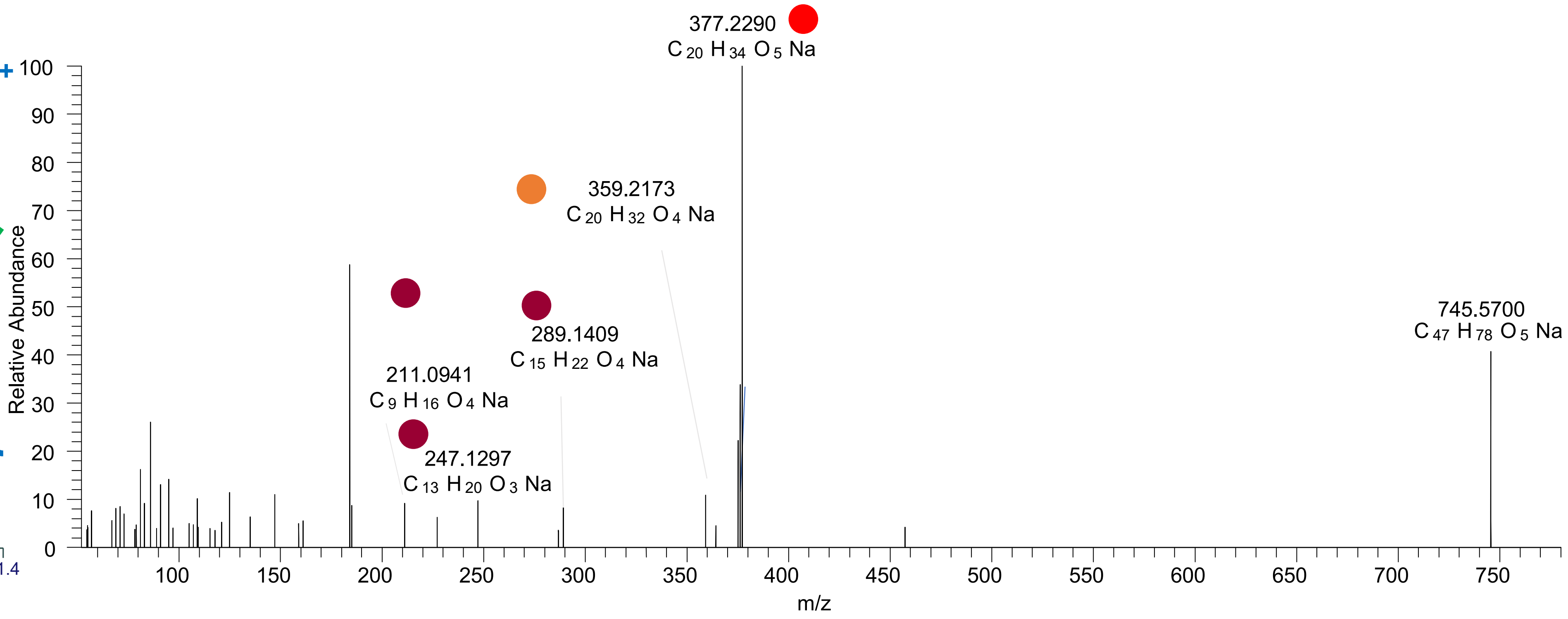

# CE(22:6<2OH>)
# RT 20.6

**[oxCE+Na]+** **XIC 751.5636 NL: 1.65E5**

- Fragments containing oxFAs
- Fragments related to water loss
- Fragments not containing oxFAs
- Position-specific fragments
- Fragments related to other oxLPPs

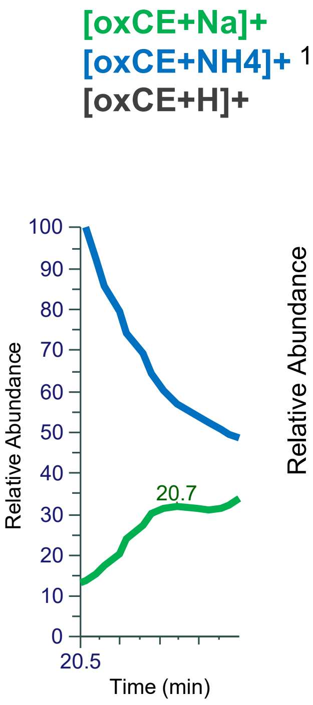


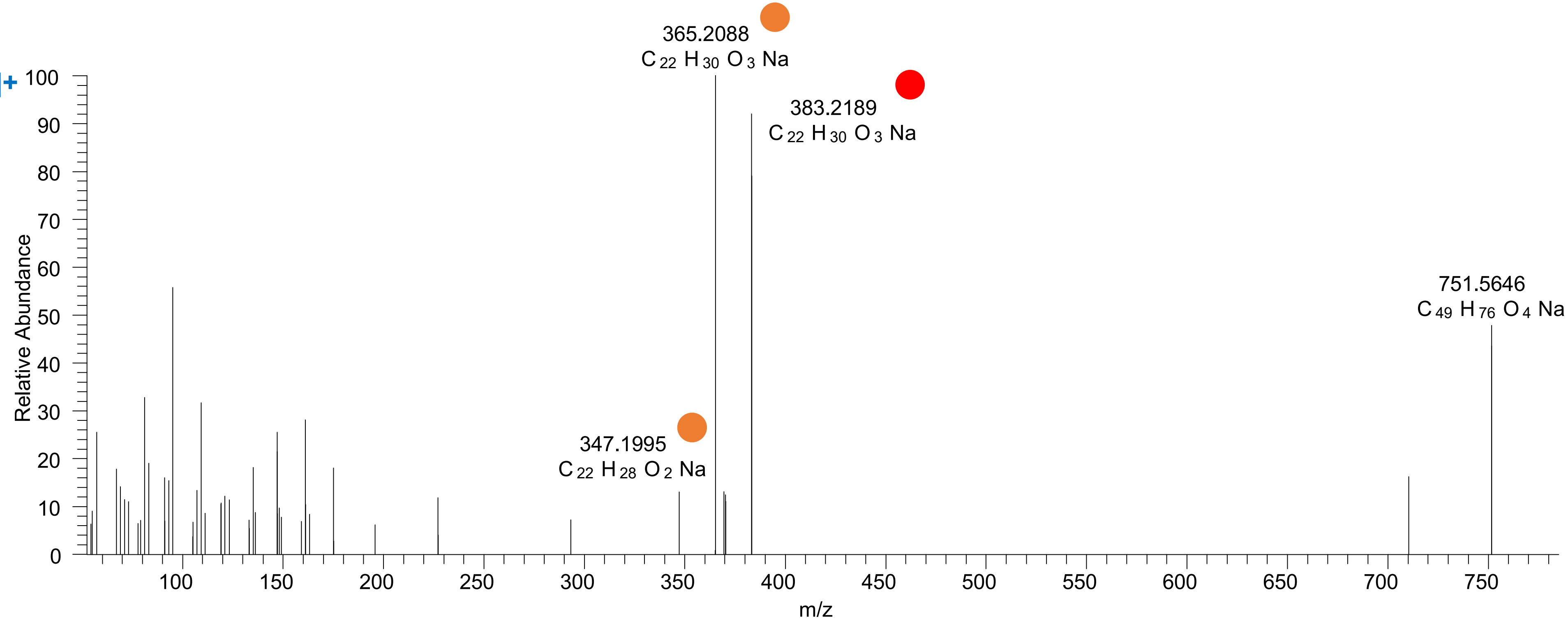

# CE(22:4<OH,ep>)
# RT 21.6-21.9

**[oxCE+Na]+** **XIC 753.5792 NL: 2.76E4**

- Fragments containing oxFAs
- Fragments related to water loss
- Fragments not containing oxFAs
- Position-specific fragments
- Fragments related to other oxLPPs

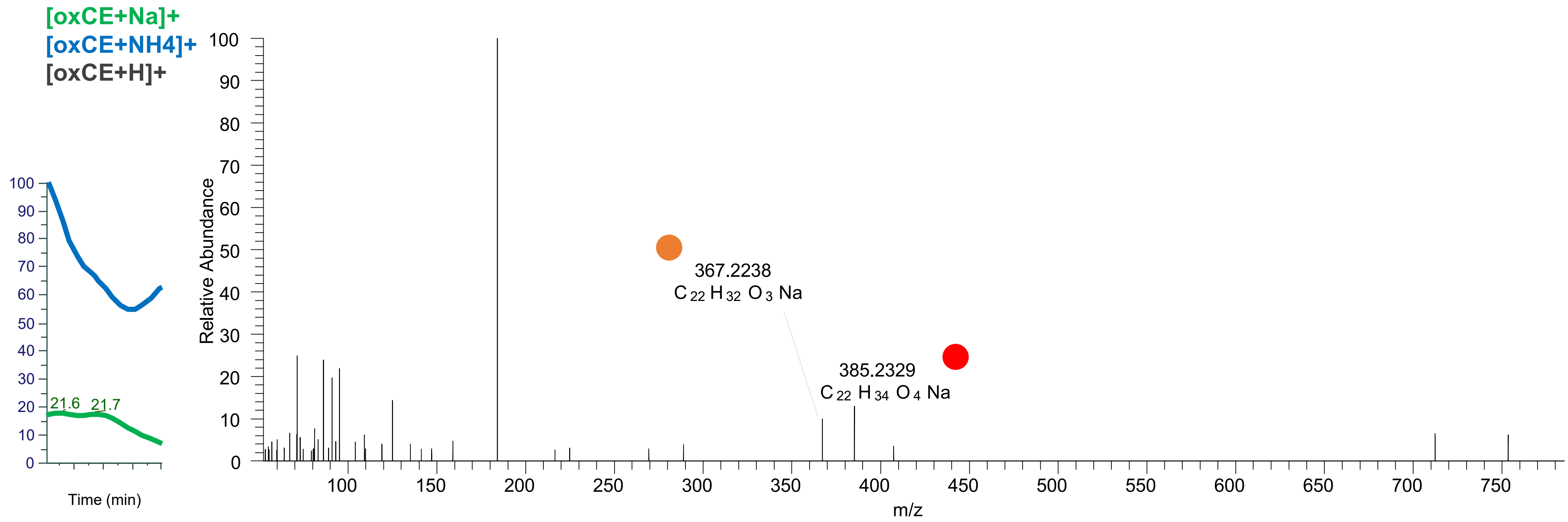

# CE(22:6<2OH,O>)
# RT 18.6

[oxCE+Na]+ XIC 767.5585 NL: 7.32E4

- Fragments containing oxFAs
- Fragments related to water loss
- Fragments not containing oxFAs
- Position-specific fragments
- Fragments related to other oxLPPs

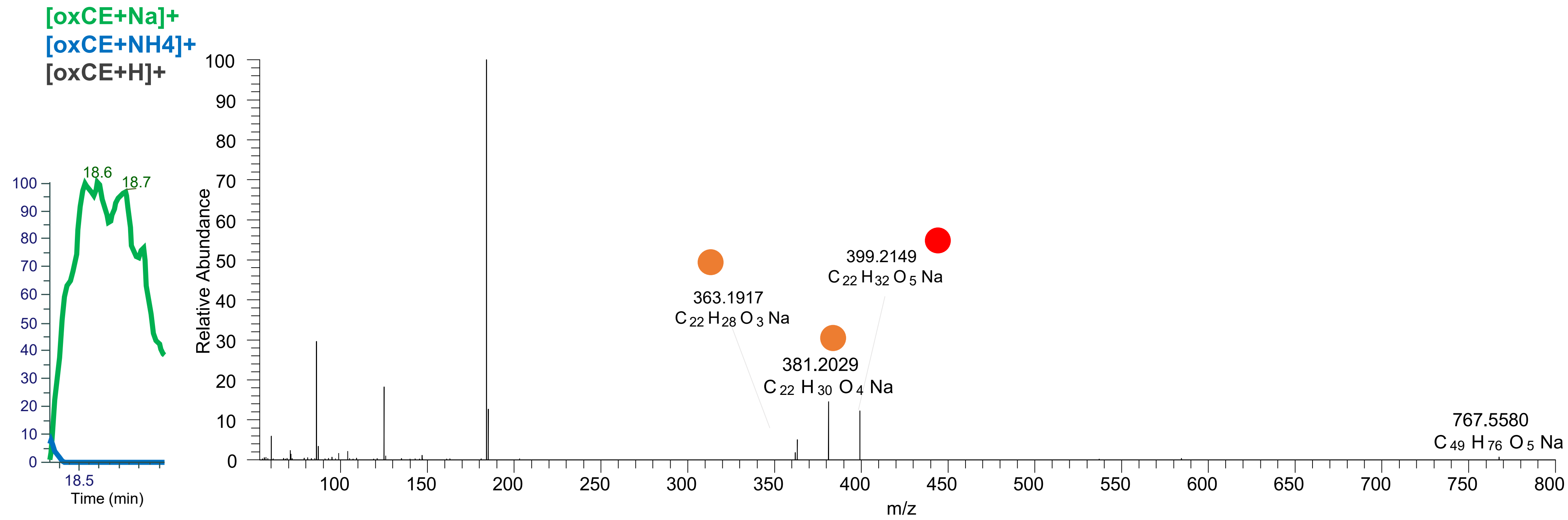

# CE(22:6<OH,2O>)
# RT 19.4

**[oxCE+Na]+** **XIC 767.5585 NL: 4.73E4**

- Fragments containing oxFAs
- Fragments related to water loss
- Fragments not containing oxFAs
- Position-specific fragments
- Fragments related to other oxLPPs

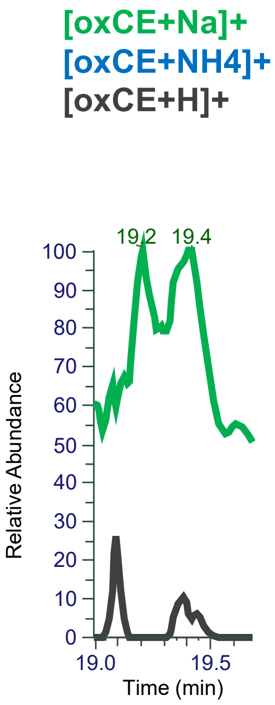


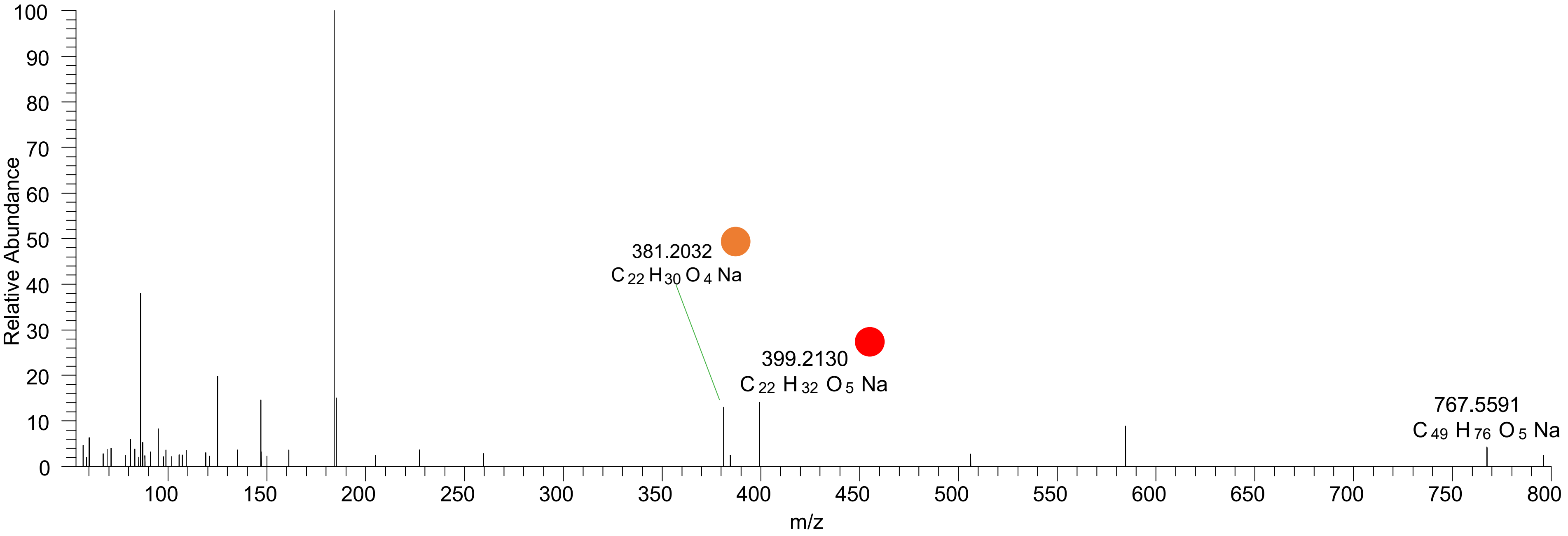

# CE(22:5<3O>)
# RT 21.8

**[oxCE+Na]+** **XIC 769.5741 NL: 7.38E3**

- Fragments containing oxFAs
- Fragments related to water loss
- Fragments not containing oxFAs
- Position-specific fragments
- Fragments related to other oxLPPs

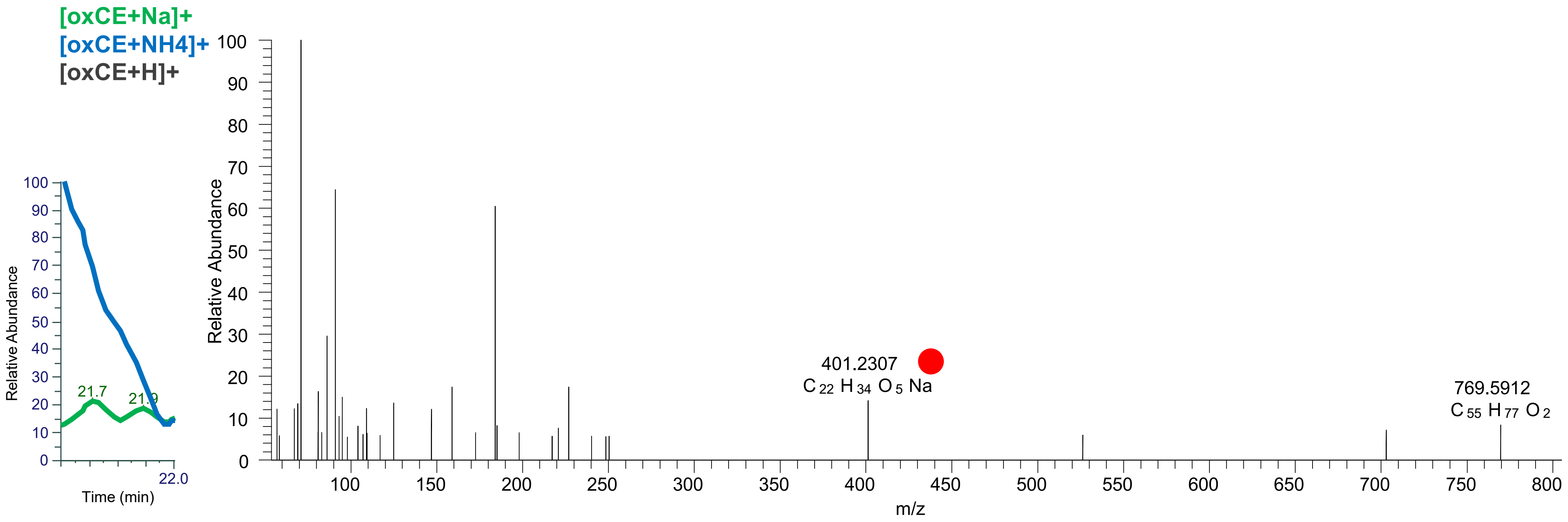
