## Supplementary material for "Epilipidomics platform for holistic profiling of oxidized complex lipids in blood plasma of obese individuals": FileS4

**Files S4.** MS2 spectra acquired using stDDA and used for the annotation of oxidized TG (oxTG) species in group pooled blood plasma samples of obese non-diabetic (OND) and obese with type 2 diabetes (OT2D) individuals. Structure-related fragmentation ions are colour-codes according to the legend provided. Annotated lipids for each group pool are sorted by their precursor *m/z*.

### OND_pool

# TG(16:1_18:1_16:0<OH>)
# RT 23.5

**[oxTG+Na]+** **XIC 869.7205 NL: 2.66E5**

- Fragments containing oxFAs
- Fragments related to water loss
- Fragments not containing oxFAs
- Fragments related to other oxLPPs
- Position-specific fragments
- Fragments related to FA loss
- Fragments related to oxFA loss

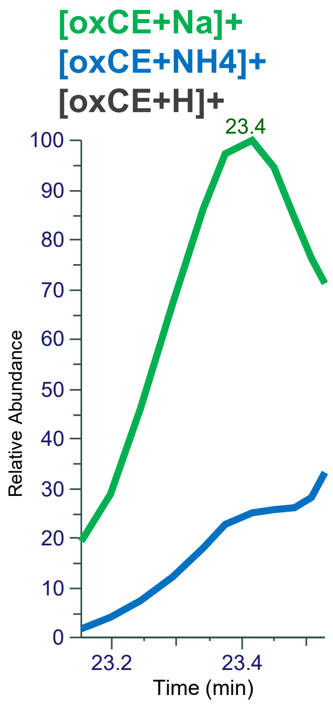


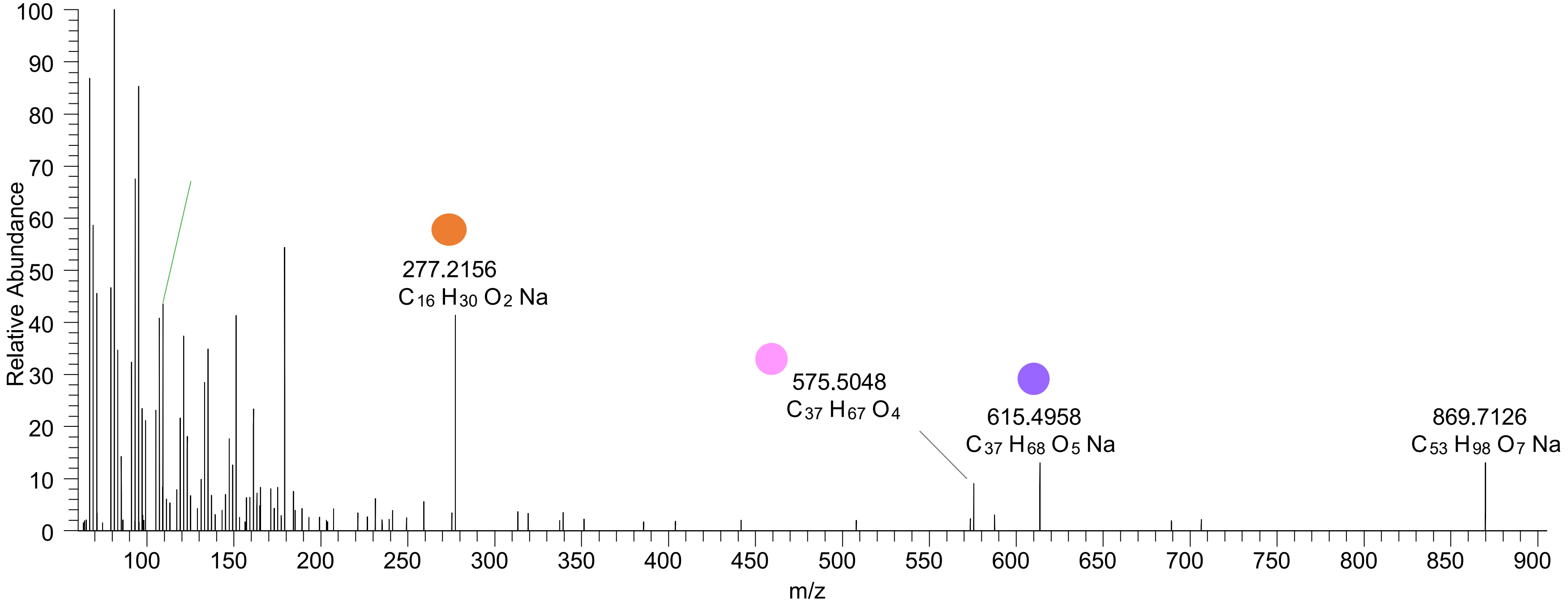

# TG(16:0_16:0_18:2<OH>)
# RT 23.7

**[oxTG+Na]+** **XIC 869.7205 NL: 2.66E5**

- Fragments containing oxFAs
- Fragments related to water loss
- Fragments not containing oxFAs
- Fragments related to other oxLPPs
- Position-specific fragments
- Fragments related to FA loss
- Fragments related to oxFA loss

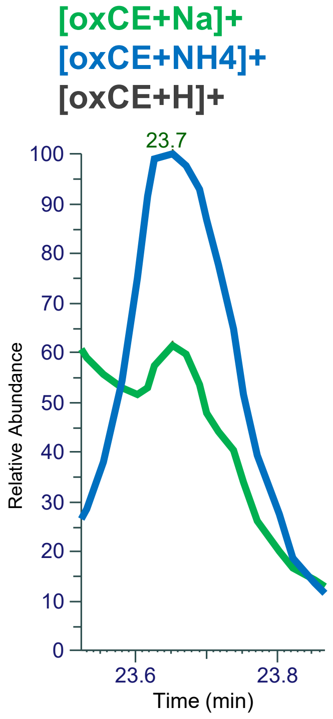


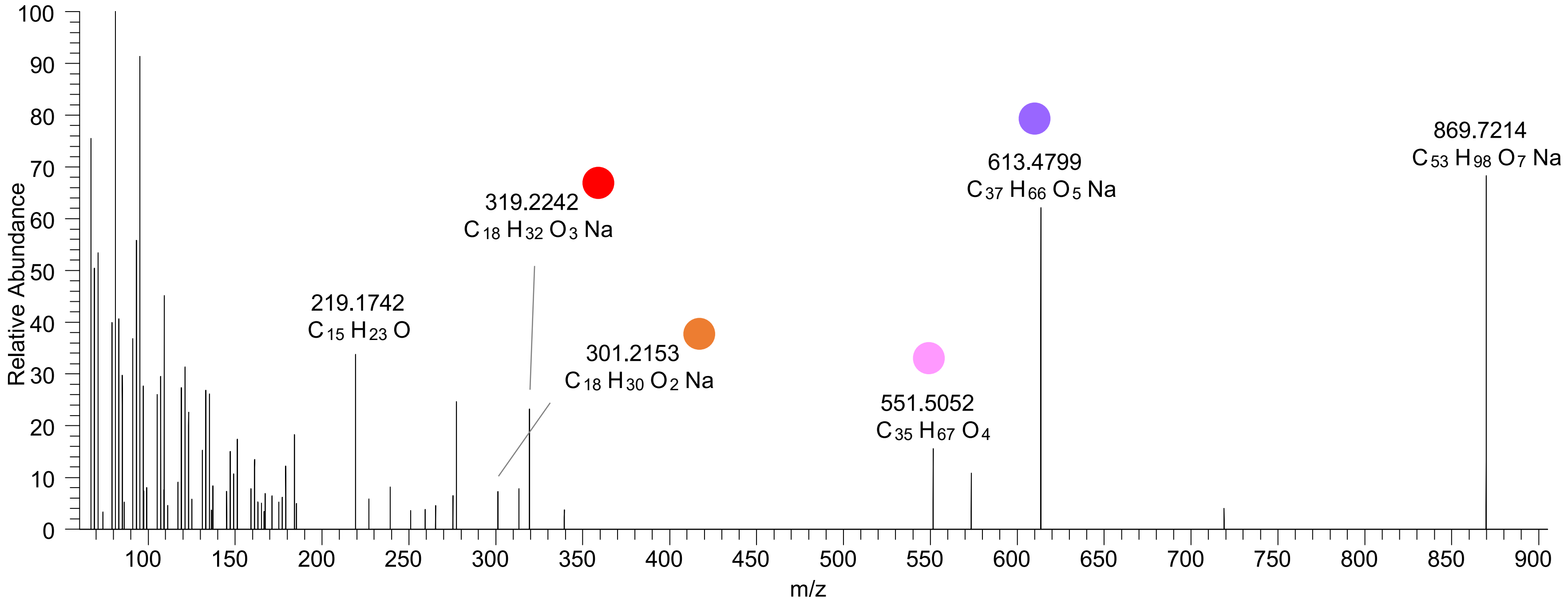

# TG(16:0_18:1_16:0<ep>)
# RT 24.1

**[oxTG+Na]+** **XIC 869.7205 NL: 5.49E4**

- Fragments containing oxFAs
- Fragments related to water loss
- Fragments not containing oxFAs
- Fragments related to other oxLPPs
- Position-specific fragments
- Fragments related to FA loss
- Fragments related to oxFA loss

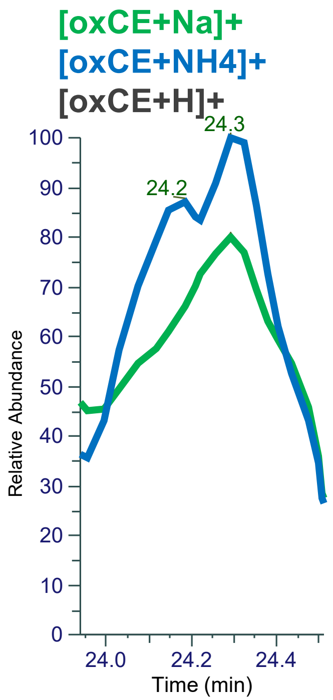


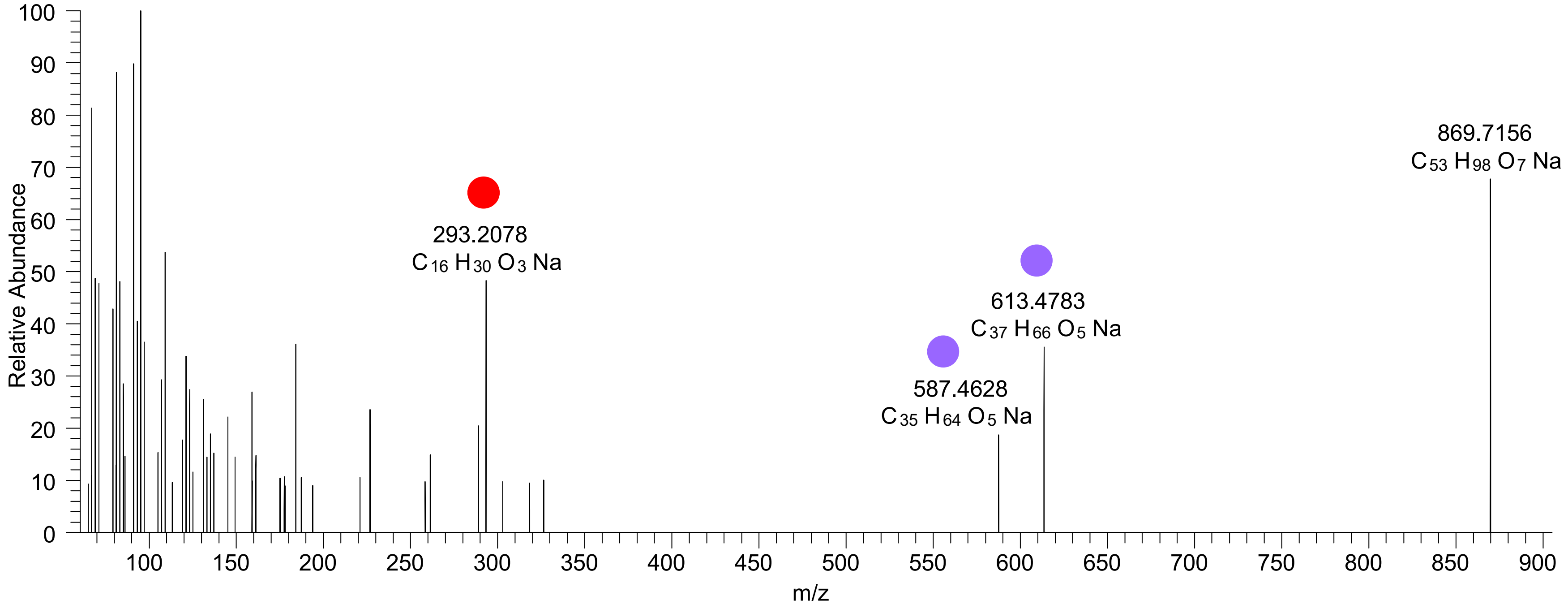

# TG(16:0_18:1_16:0<ep>)
# TG(16:0_16:0_18:1<ep>)
# RT 24.3-24.4

**[oxTG+Na]+** **XIC 869.7205 NL: 5.49E4**

- Fragments containing oxFAs
- Fragments related to water loss
- Fragments not containing oxFAs
- Fragments related to other oxLPPs
- Position-specific fragments
- Fragments related to FA loss
- Fragments related to oxFA loss

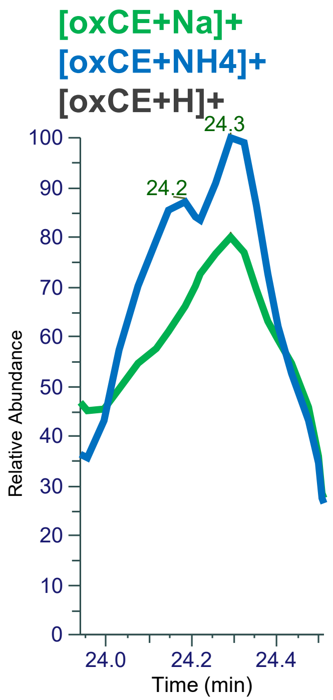


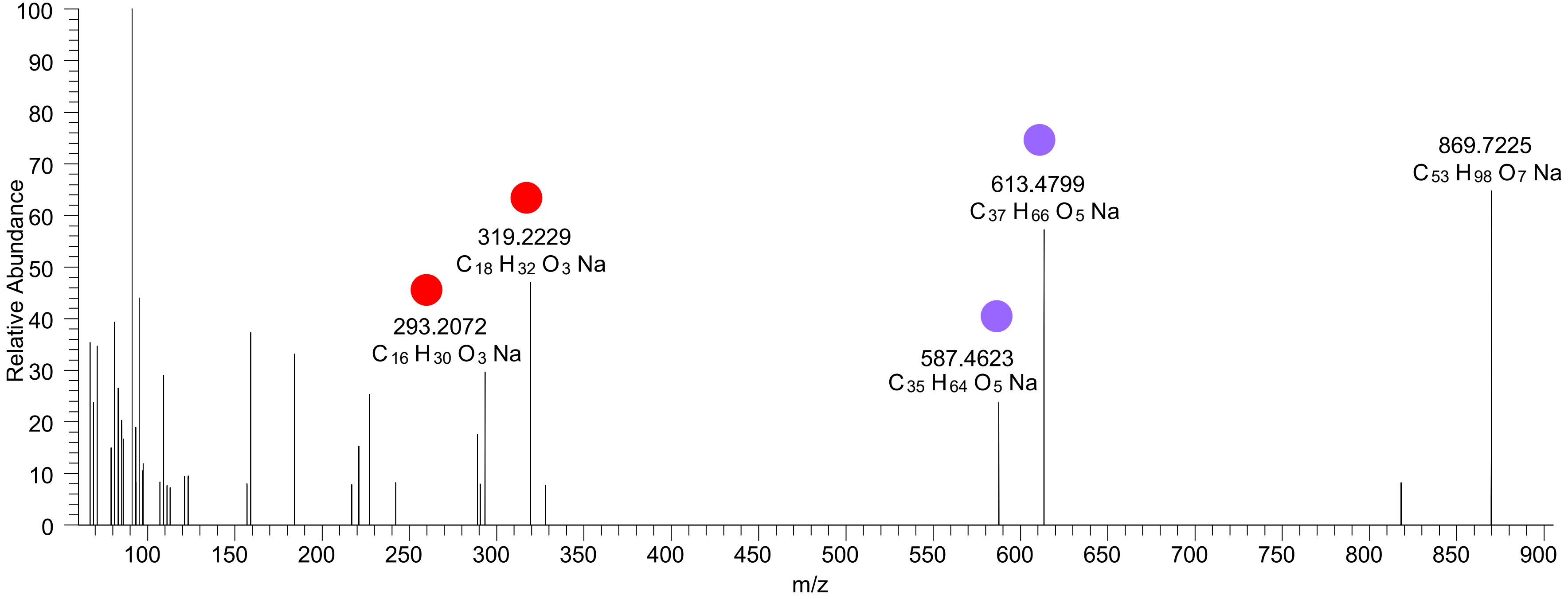

# TG(16:0_18:2_18:2<OH>)
# TG(16:1_18:1_18:2<OH>)
# RT 23.0-23.2

**[oxTG+Na]+** **XIC 893.7204 NL: 2.22E6**

- Fragments containing oxFAs
- Fragments related to water loss
- Fragments not containing oxFAs
- Fragments related to other oxLPPs
- Position-specific fragments
- Fragments related to FA loss
- Fragments related to oxFA loss

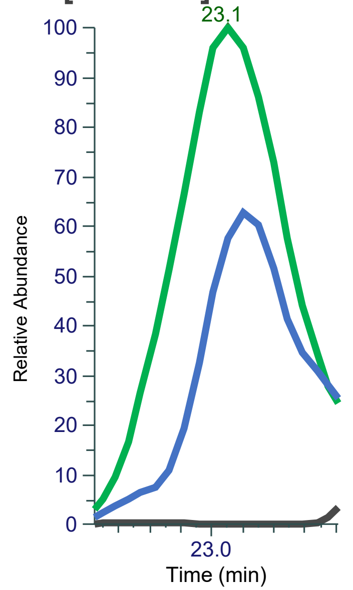


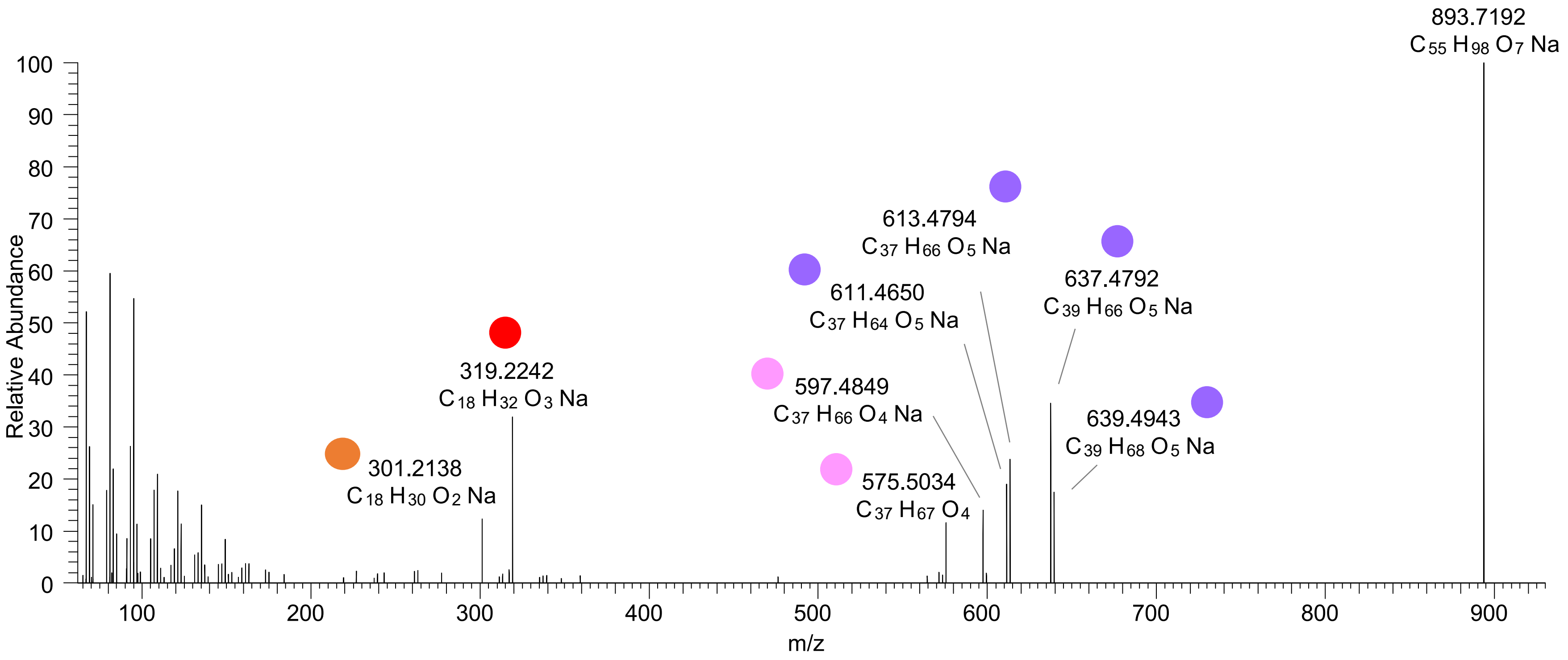

# TG(16:0_18:1_18:3<OH>)
# RT 23.4

**[oxTG+Na]+** **XIC 893.7204 NL: 2.22E6**

- Fragments containing oxFAs
- Fragments related to water loss
- Fragments not containing oxFAs
- Fragments related to other oxLPPs
- Position-specific fragments
- Fragments related to FA loss
- Fragments related to oxFA loss

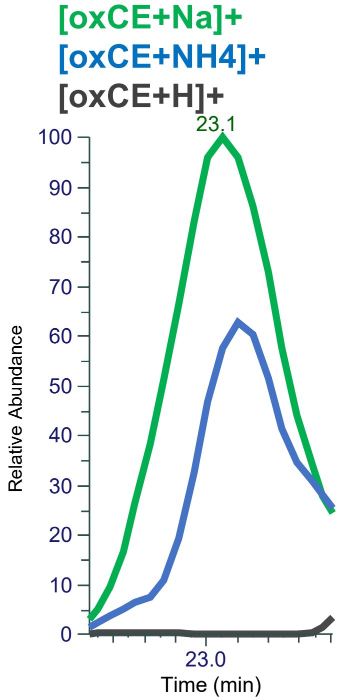


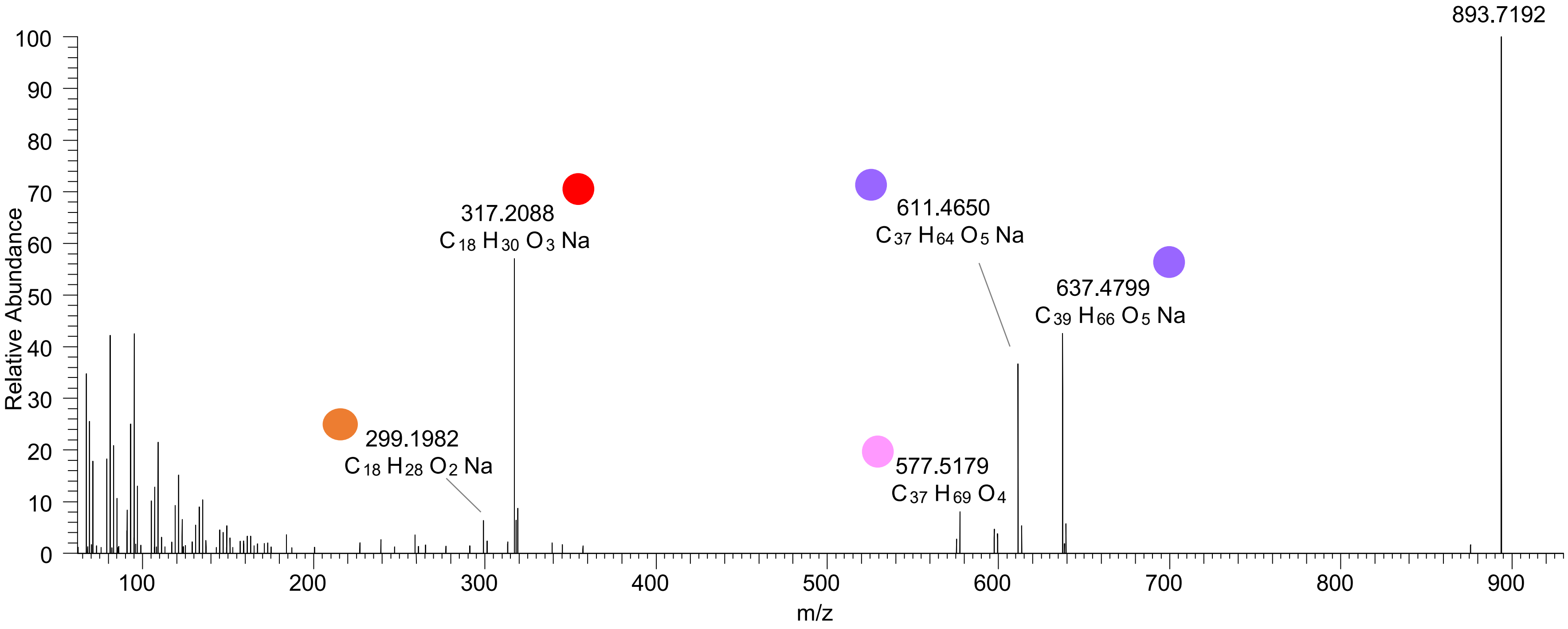

### TG(16:0_18:1_18:2<oxo>)
# RT 24.0

**[oxTG+Na]+** **XIC 893.7204 NL: 5.72E5**

- Fragments containing oxFAs
- Fragments related to water loss
- Fragments not containing oxFAs
- Fragments related to other oxLPPs
- Position-specific fragments
- Fragments related to FA loss
- Fragments related to oxFA loss

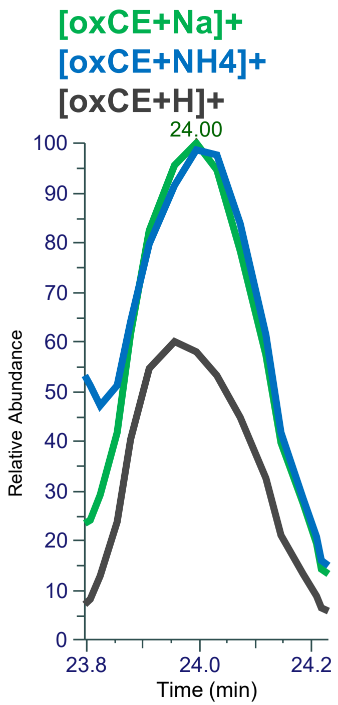


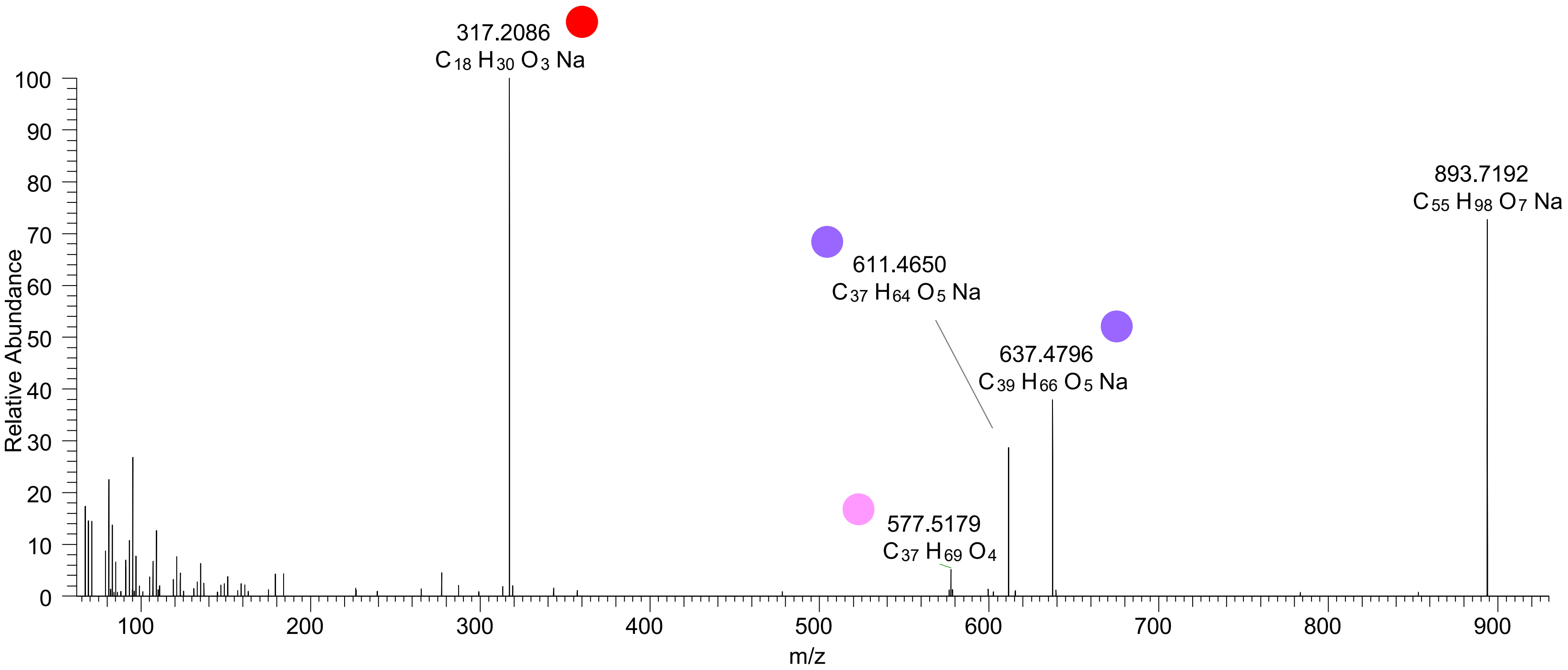

### TG(16:0_18:1_18:2<oxo>)
### TG(16:0_18:0_18:3<oxo>)
# RT 24.8

**[oxTG+Na]+** **XIC 893.7204 NL: 3.52E4**

- Fragments containing oxFAs
- Fragments related to water loss
- Fragments not containing oxFAs
- Fragments related to other oxLPPs
- Position-specific fragments
- Fragments related to FA loss
- Fragments related to oxFA loss

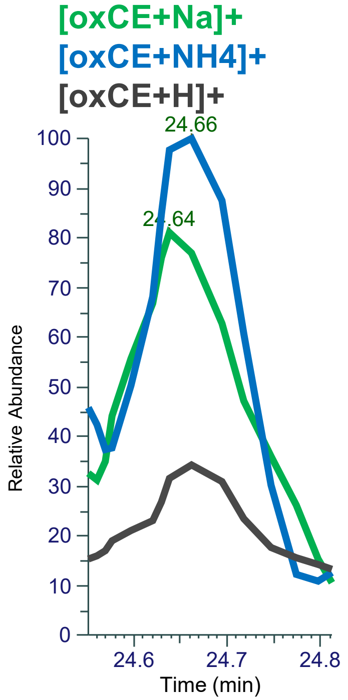


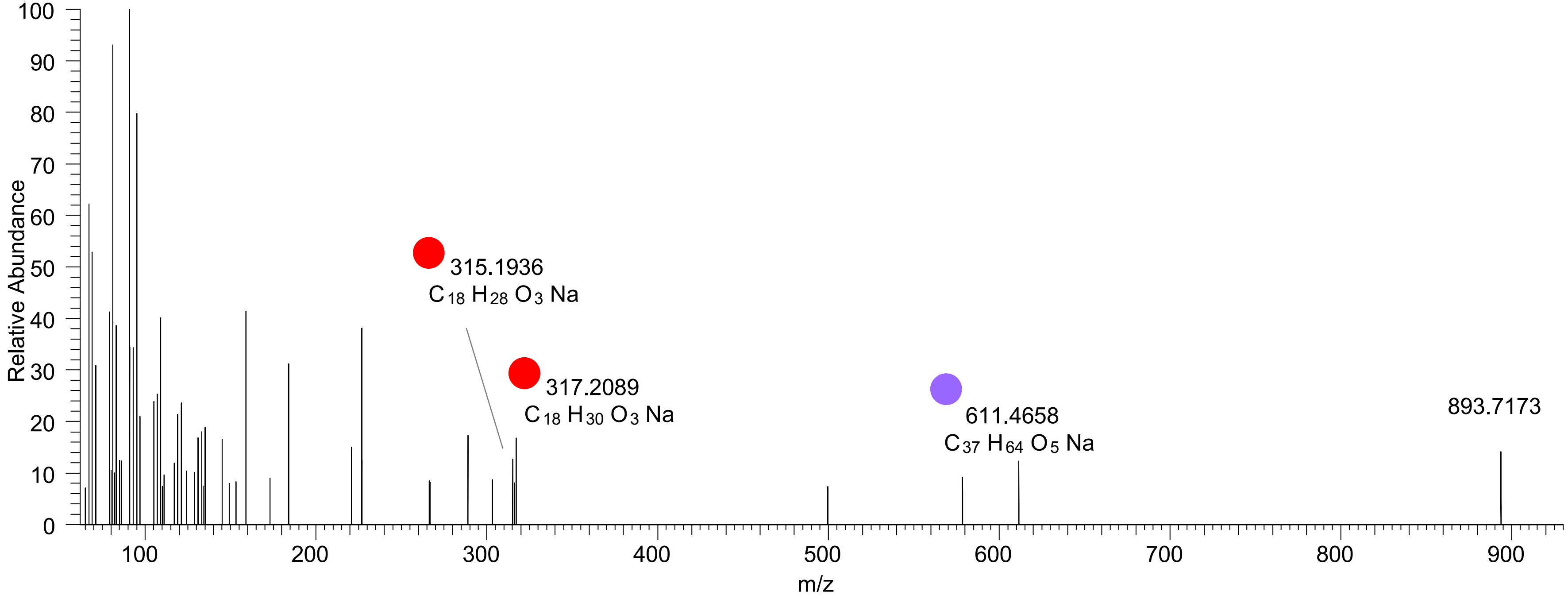

# TG(16:0_18:1_18:2<ep>)
# RT 25.4

**[oxTG+Na]+** **XIC 893.7204 NL: 2.02E5**

- Fragments containing oxFAs
- Fragments related to water loss
- Fragments not containing oxFAs
- Fragments related to other oxLPPs
- Position-specific fragments
- Fragments related to FA loss
- Fragments related to oxFA loss

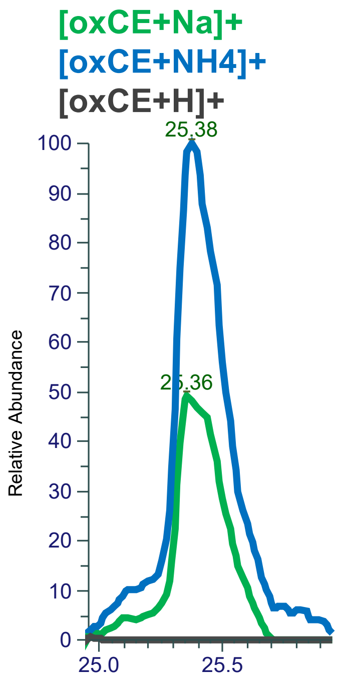


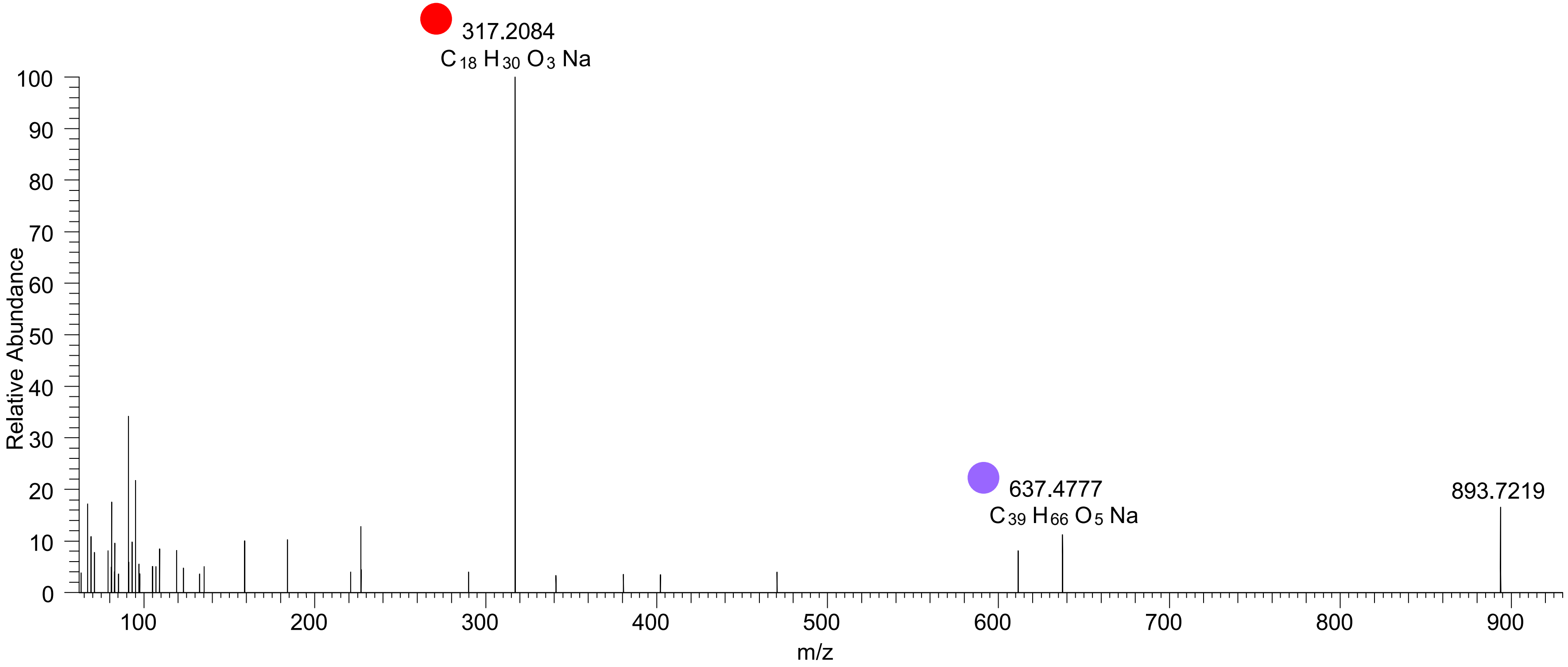

# TG(16:0_18:1_18:2<OH>)
# RT 23.4

**[oxTG+Na]+** **XIC 895.7361 NL: 7.26E5**

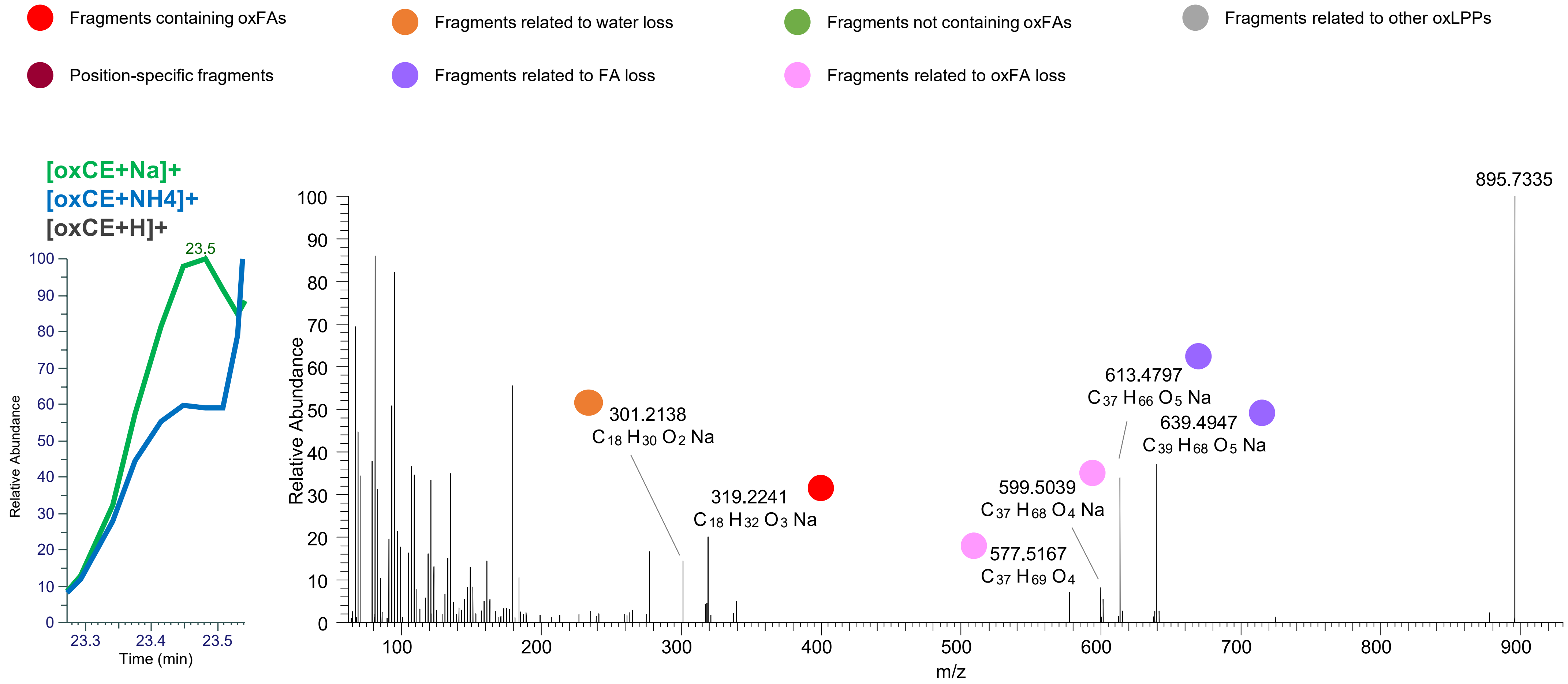

# TG(16:0_18:1_18:2<OH>)
# RT 23.7

**[oxTG+Na]+** **XIC 895.7361 NL: 2.27E6**

- Fragments containing oxFAs
- Fragments related to water loss
- Fragments not containing oxFAs
- Fragments related to other oxLPPs
- Position-specific fragments
- Fragments related to FA loss
- Fragments related to oxFA loss

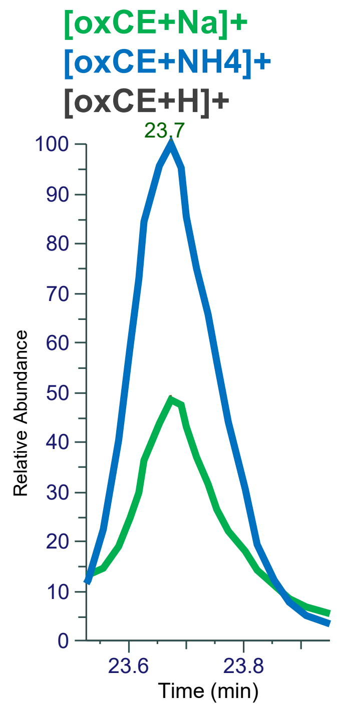


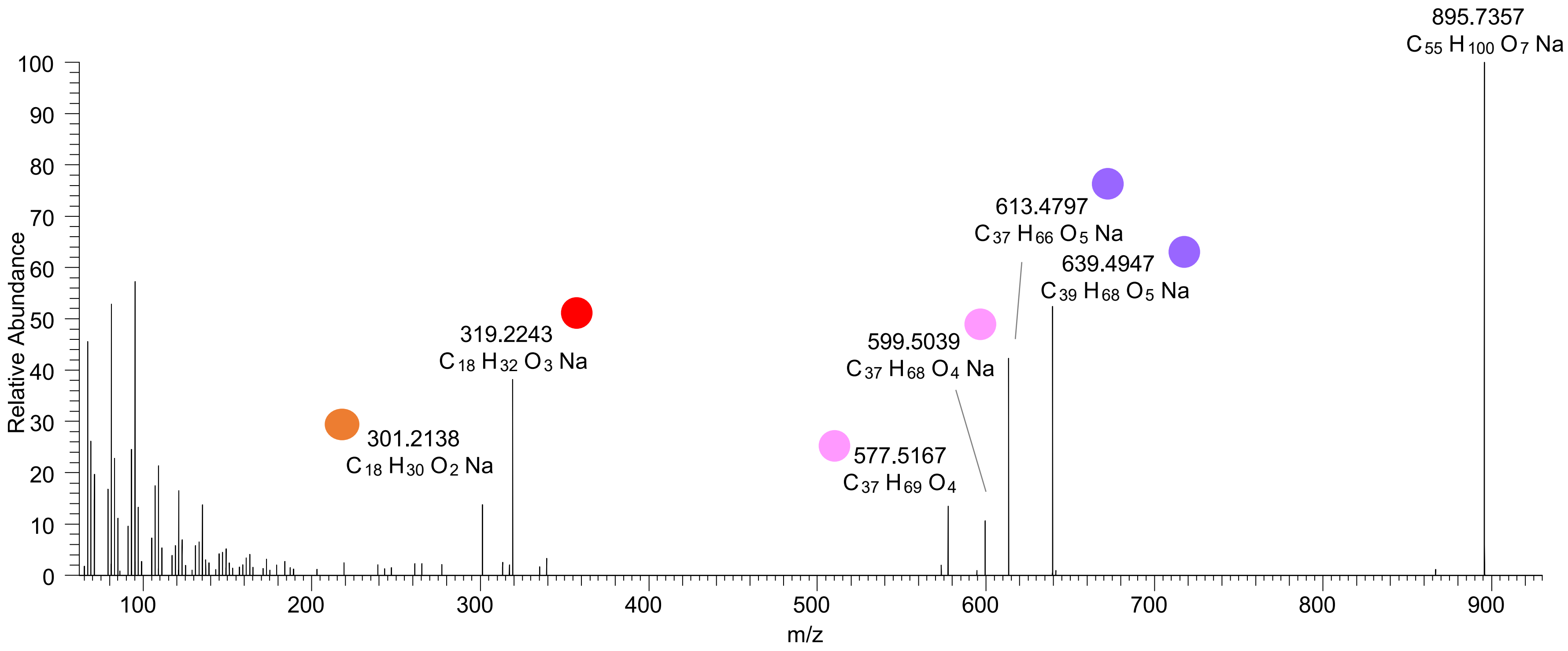

# TG(16:0_18:1_18:2<O>)
# TG(18:0_18:2_16:1<O>)
# RT 24.3

**[oxTG+Na]+** **XIC 895.7361 NL: 1.71E5**

- Fragments containing oxFAs
- Fragments related to water loss
- Fragments not containing oxFAs
- Fragments related to other oxLPPs
- Position-specific fragments
- Fragments related to FA loss
- Fragments related to oxFA loss

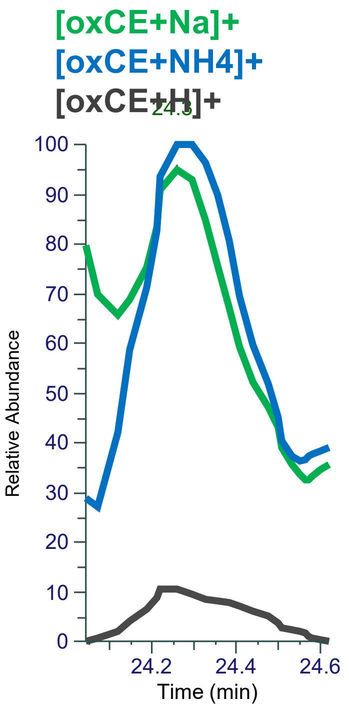


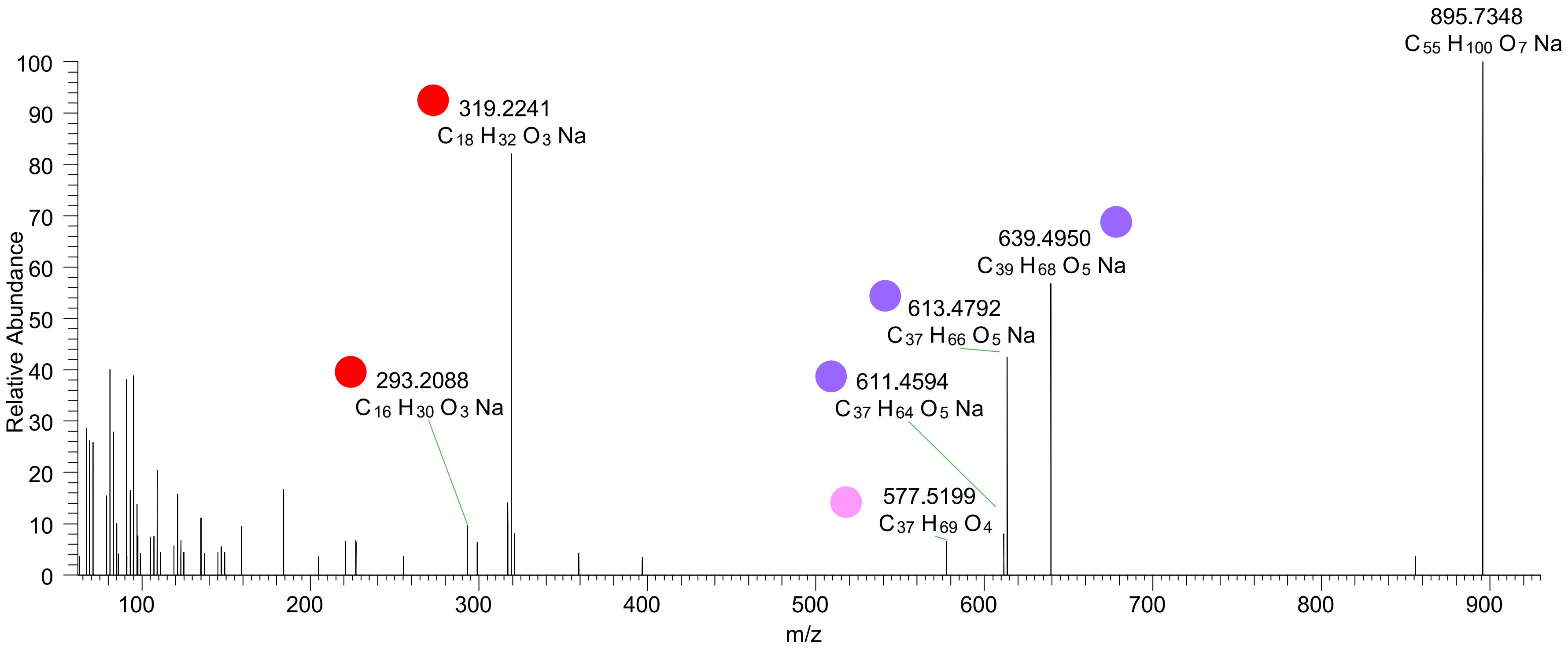

# TG(16:0_18:1_18:2<O>)
# RT 24.8

**[oxTG+Na]+** **XIC 895.7361 NL: 8.07E4**

- Fragments containing oxFAs
- Fragments related to water loss
- Fragments not containing oxFAs
- Fragments related to other oxLPPs
- Position-specific fragments
- Fragments related to FA loss
- Fragments related to oxFA loss

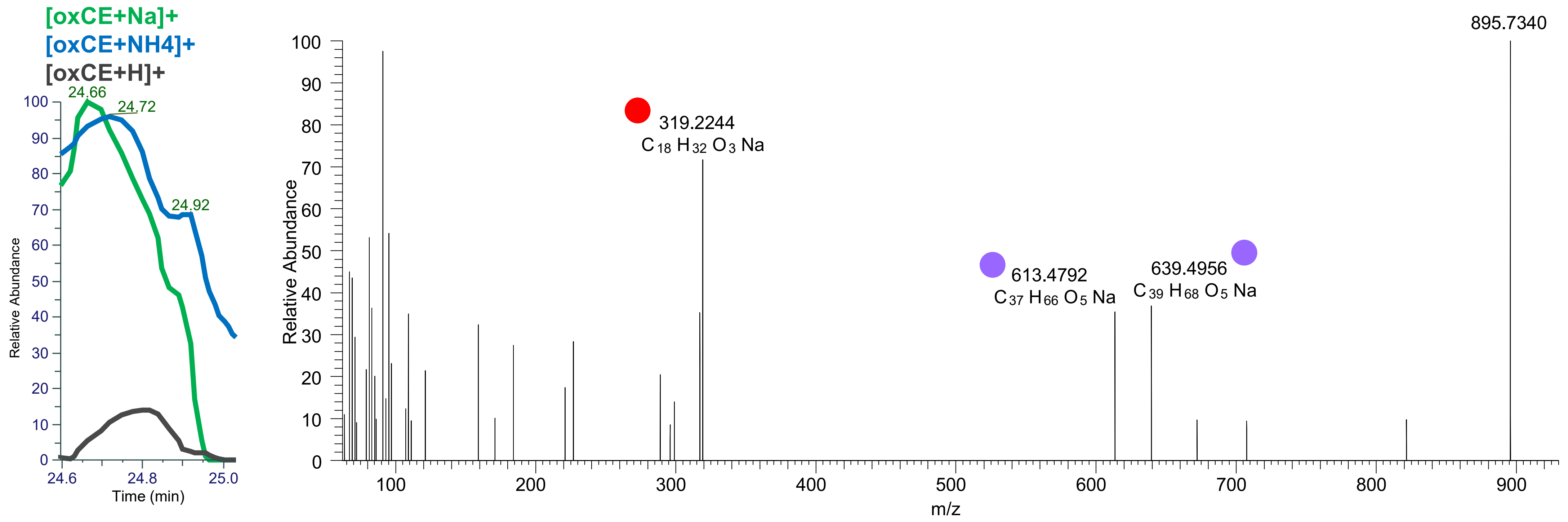

# TG(16:0_18:1_18:1<OH>)
# RT 24.1

**[oxTG+Na]+** **XIC 897.7517 NL: 4.14E5**

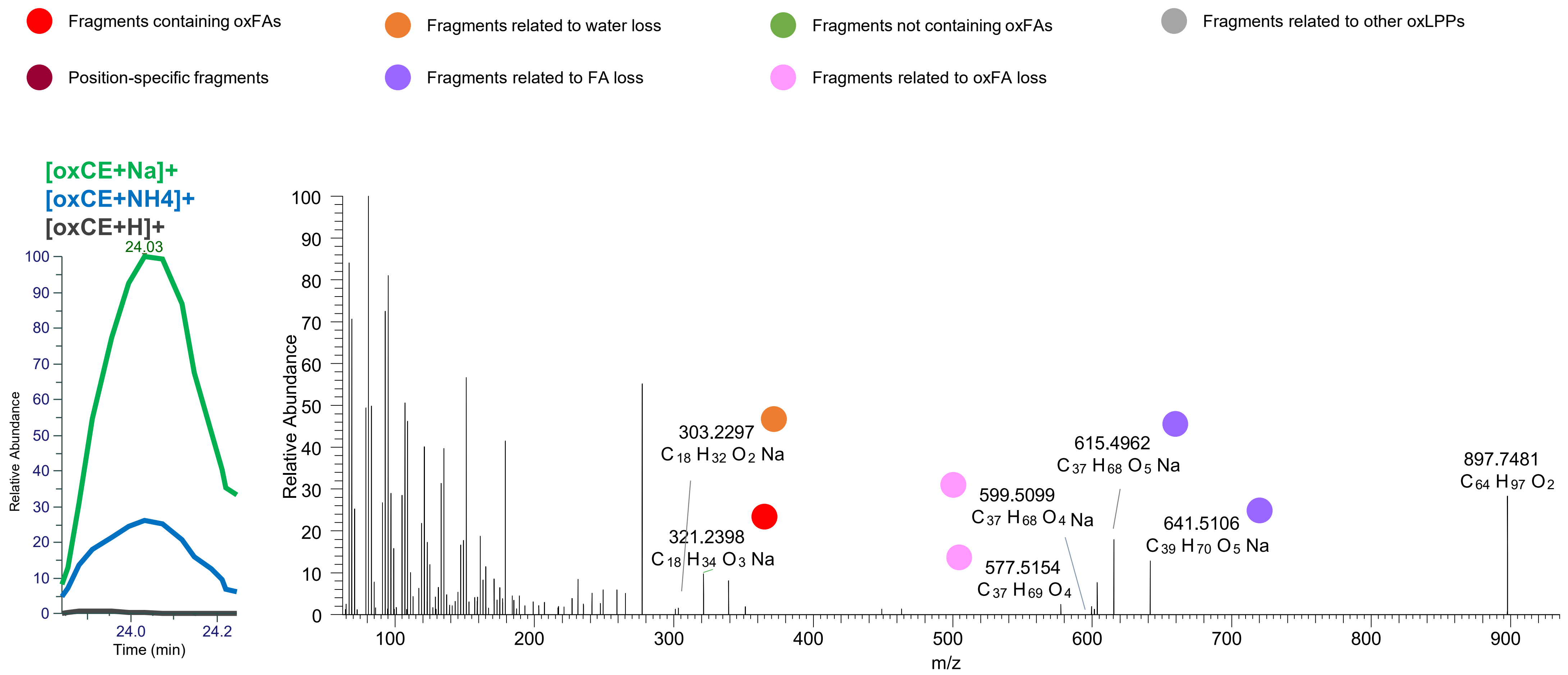

# TG(16:0_18:0_18:2<OH>)
# RT 24.4

**[oxTG+Na]+** **XIC 897.7517 NL: 1.35E5**

- Fragments containing oxFAs
- Fragments related to water loss
- Fragments not containing oxFAs
- Fragments related to other oxLPPs
- Position-specific fragments
- Fragments related to FA loss
- Fragments related to oxFA loss

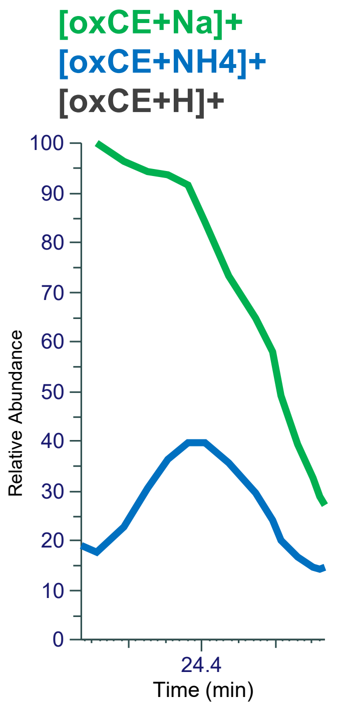


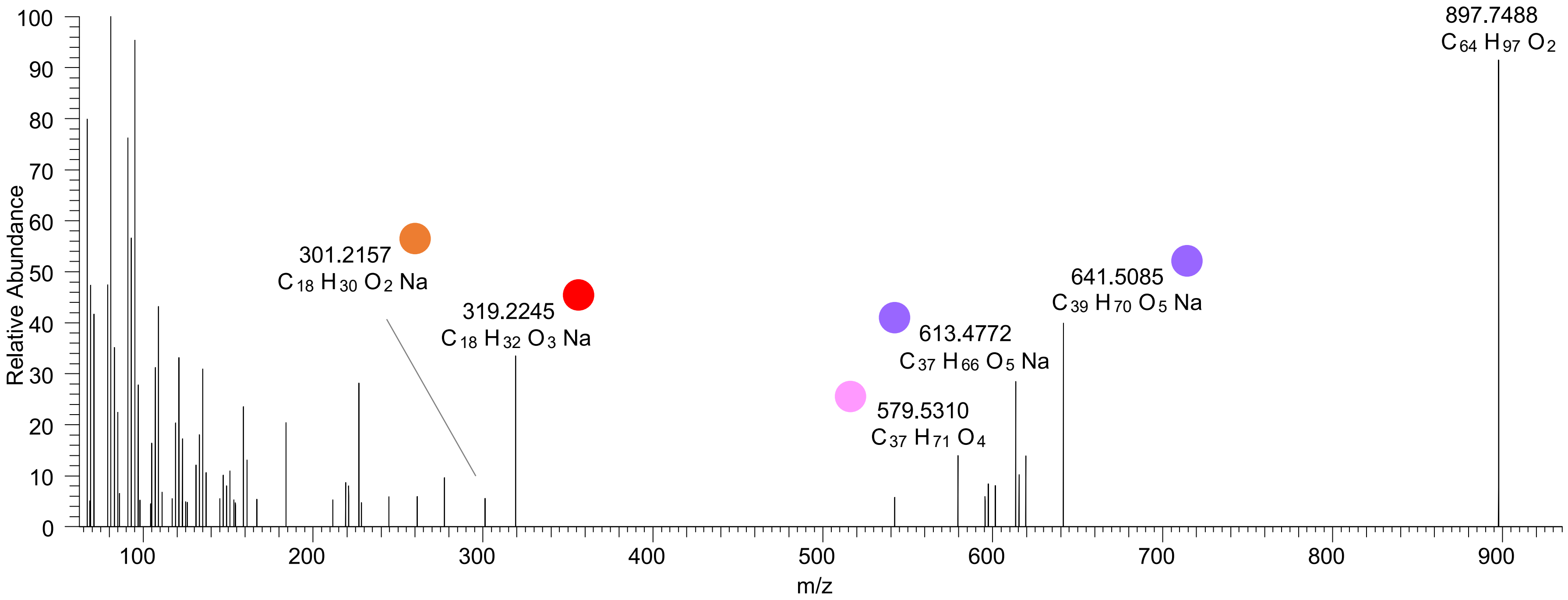

# TG(16:0_18:3_18:1<2OH>)
# RT 20.2

**[oxTG+Na]+** **XIC 909.7154 NL: 1.57E5**

- Fragments containing oxFAs
- Fragments related to water loss
- Fragments not containing oxFAs
- Fragments related to other oxLPPs
- Position-specific fragments
- Fragments related to FA loss
- Fragments related to oxFA loss

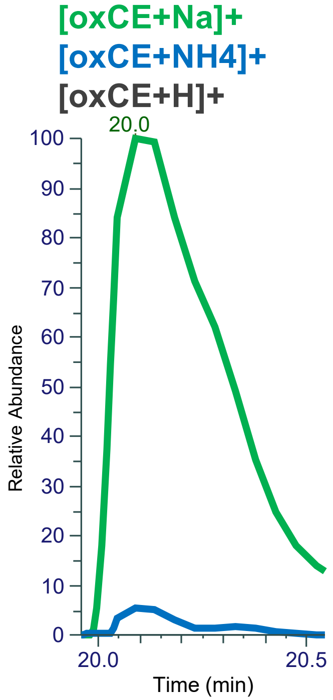


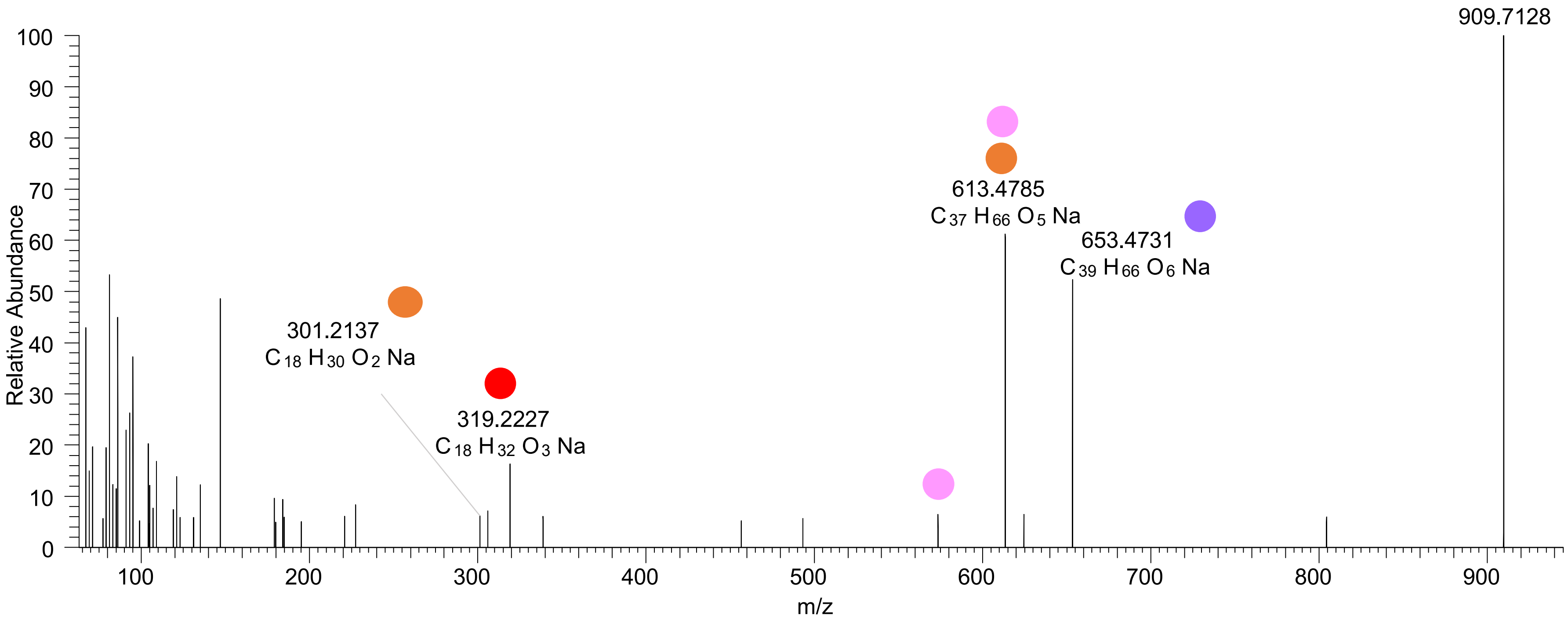

# TG(16:0_18:1_18:3<OH,O>)
# RT 21.6

**[oxTG+Na]+** **XIC 909.7154 NL: 9.65E4**

- Fragments containing oxFAs
- Fragments related to water loss
- Fragments not containing oxFAs
- Fragments related to other oxLPPs
- Position-specific fragments
- Fragments related to FA loss
- Fragments related to oxFA loss

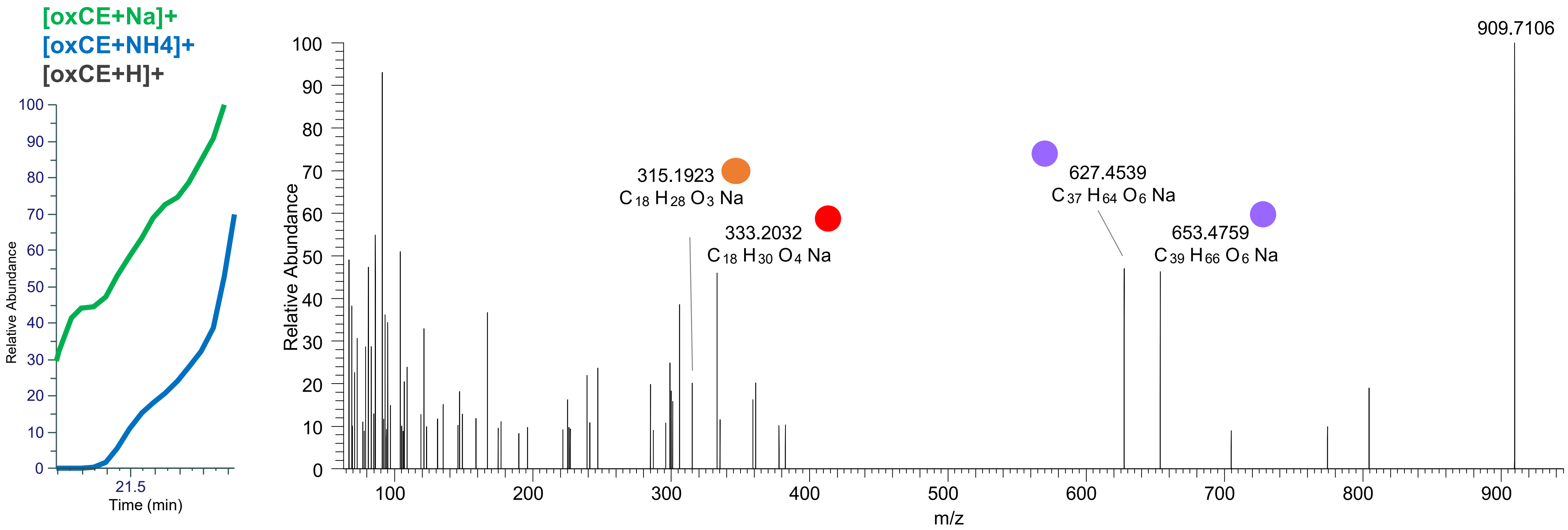

TG(16:1_18:1_18:2<2O>)
TG(16:0_18:1_18:3<2O>)
TG(16:0_18:2_18:2<2O>)
TG(16:0_16:0_20:4<2O>)
RT 22.1

**[oxTG+Na]+** **XIC 909.7154 NL: 1.48E5**

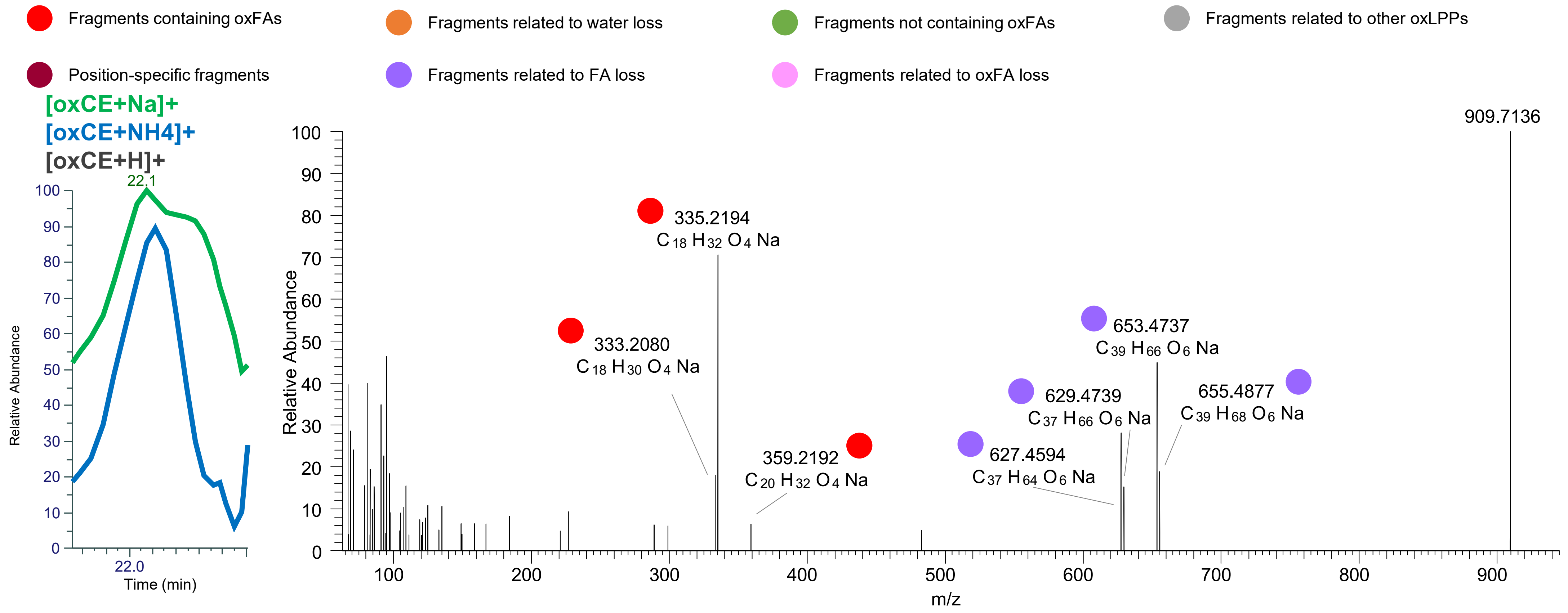

TG(16:0_18:2_18:2<OOH{13}>)
TG(16:1_18:1_18:2<OOH{13}>)
TG(16:0_18:1_18:3<2O>)
RT 23.0

[oxTG+Na]+ XIC 909.7154 NL: 2.29E6

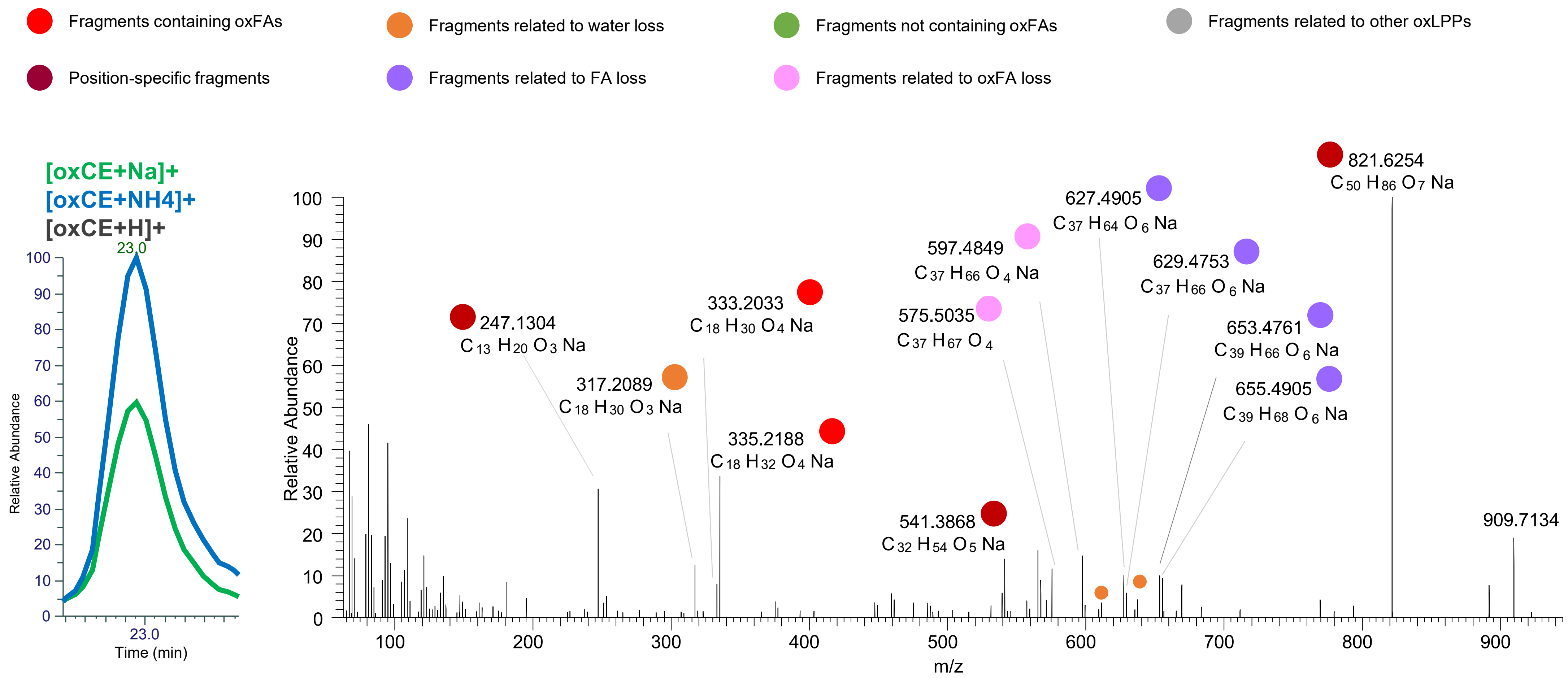

### TG(16:0_18:1_18:3<OOH{13}>)
### TG(16:0_18:1_18:3<OOH{16}>)
# RT 23.3

**[oxTG+Na]+** **XIC 909.7154 NL: 2.29E6**

- Fragments containing oxFAs
- Fragments related to water loss
- Fragments not containing oxFAs
- Fragments related to other oxLPPs
- Position-specific fragments
- Fragments related to FA loss
- Fragments related to oxFA loss

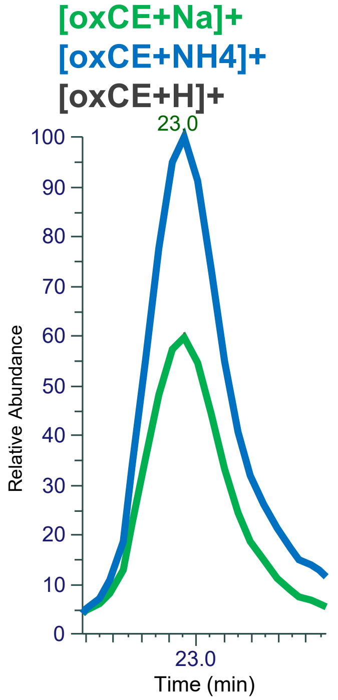


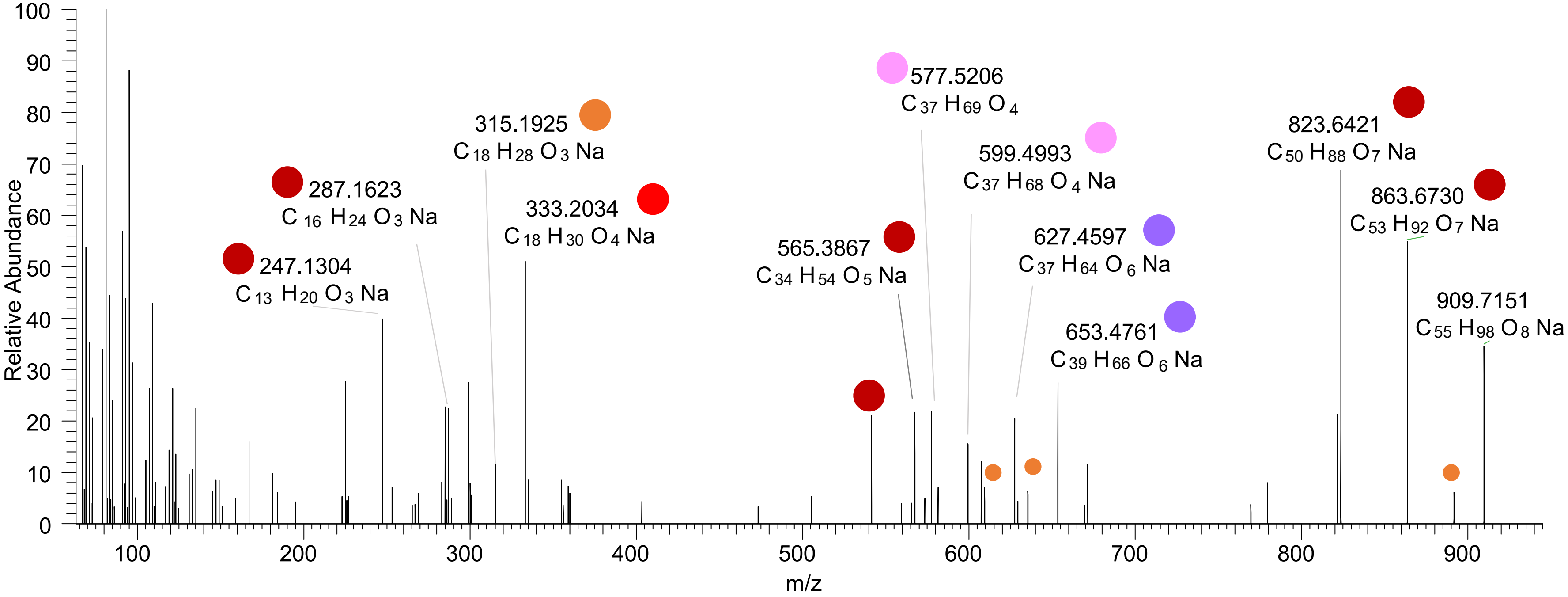

# TG(16:0_18:1_18:1<OH,ep>)
# RT 22.7

**[oxTG+Na]+ XIC 911.7310 NL: 1.52E5**

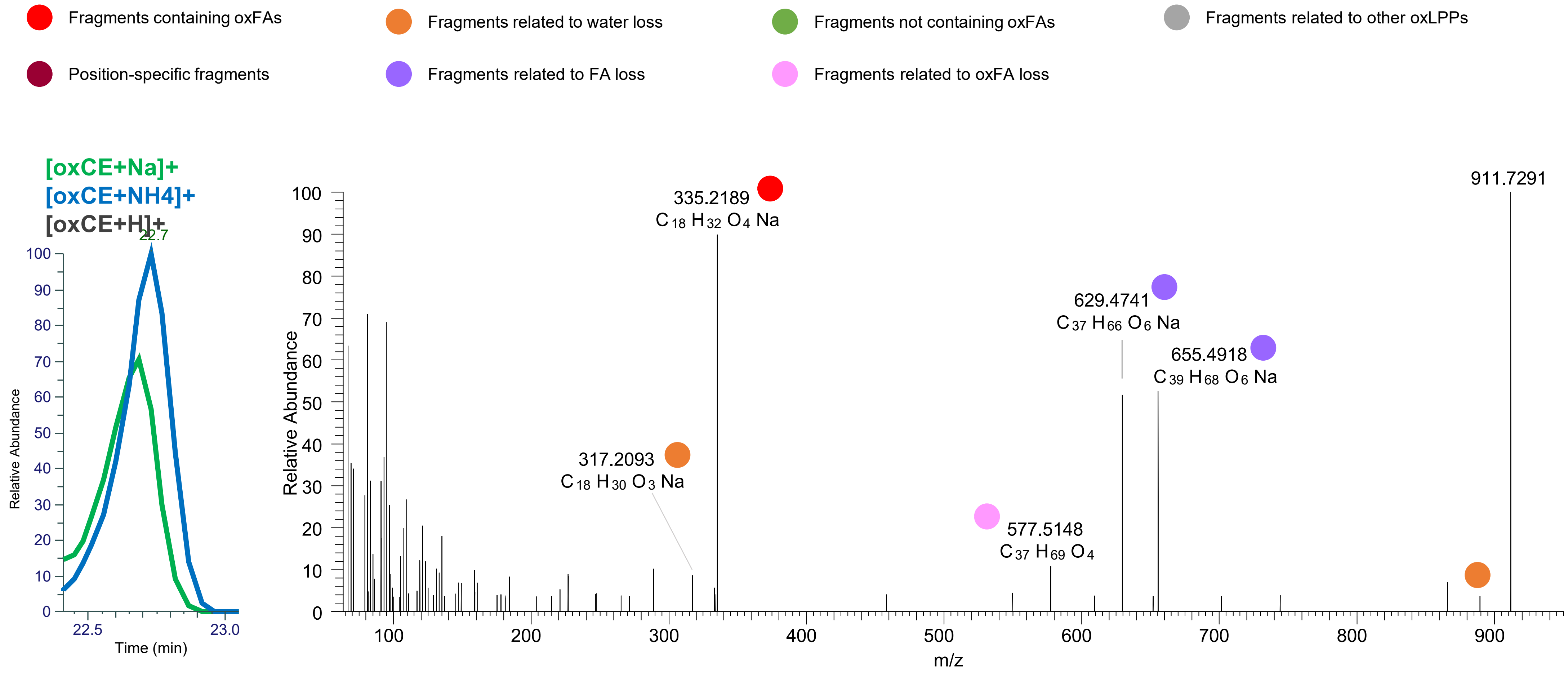

TG(16:0_18:1_18:2<OOH{11}>)
TG(16:0_18:1_18:2<OOH{13}>)
RT 23.4

[oxTG+Na]+ XIC 911.7310 NL: 9.73E5

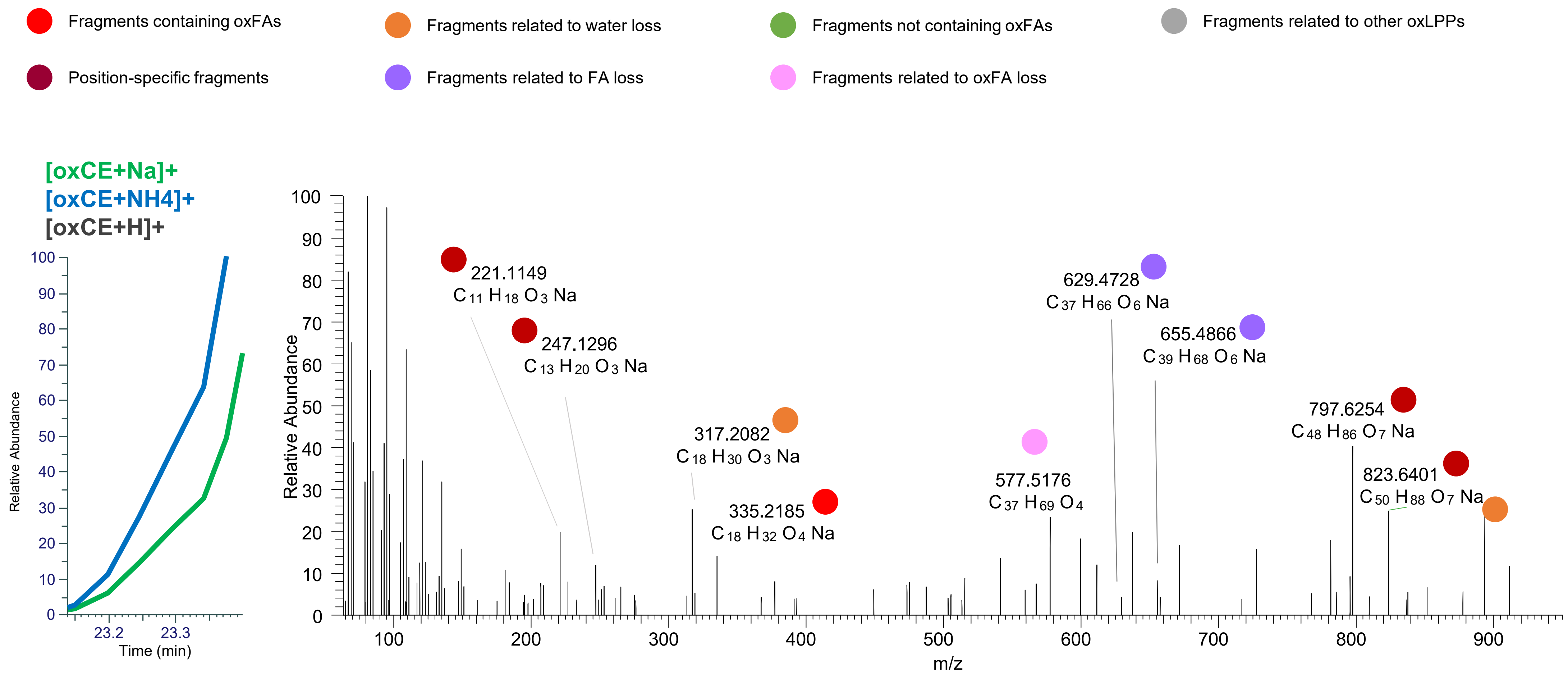

### TG(16:0_18:1_18:2<OOH{13}>)
# RT 23.5

**[oxTG+Na]+ XIC 911.7310 NL: 4.68E6**

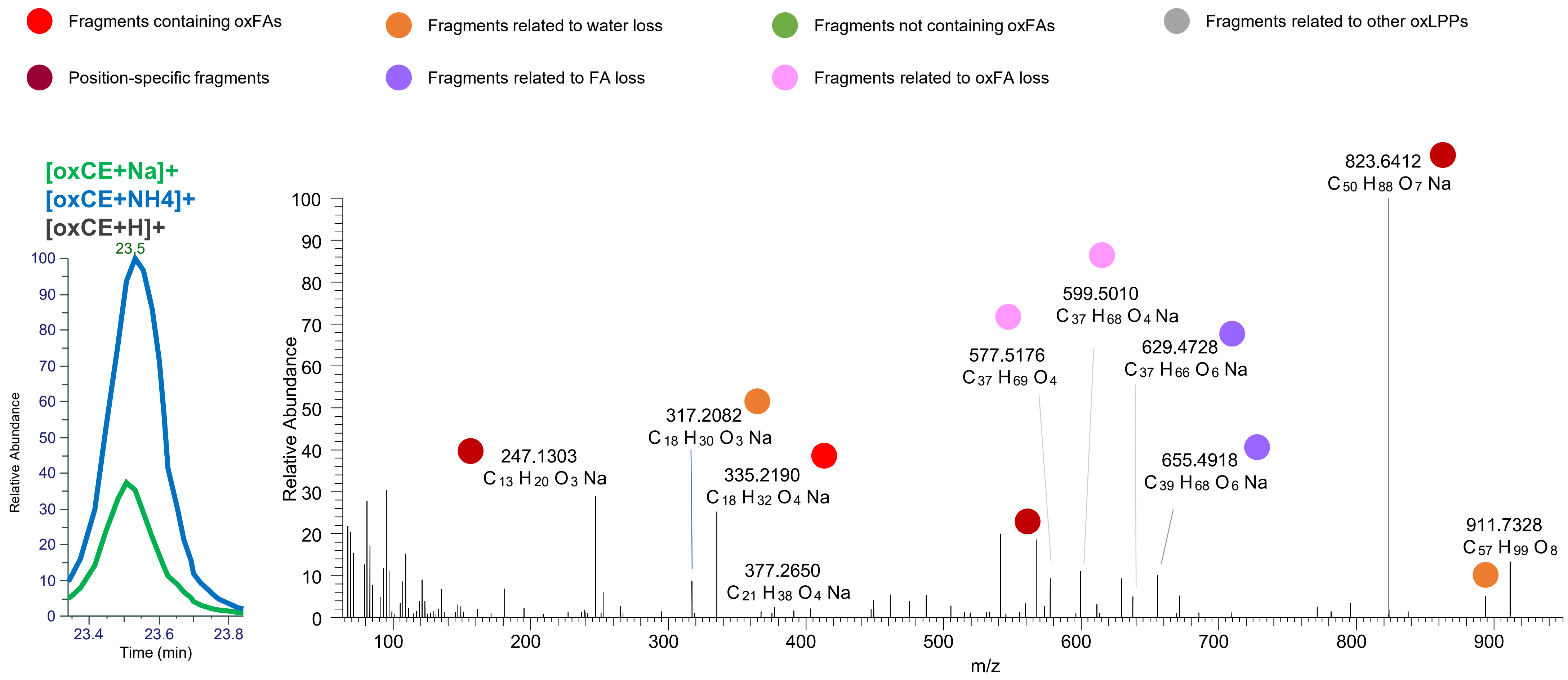

### TG(16:0_18:1_18:2<OOH{10}>)
### TG(16:0_18:1_18:2<OOH{13}>)
# RT 23.8

**[oxTG+Na]+** **XIC 911.7310 NL: 4.68E6**

- Fragments containing oxFAs
- Fragments related to water loss
- Fragments not containing oxFAs
- Fragments related to other oxLPPs
- Position-specific fragments
- Fragments related to FA loss
- Fragments related to oxFA loss

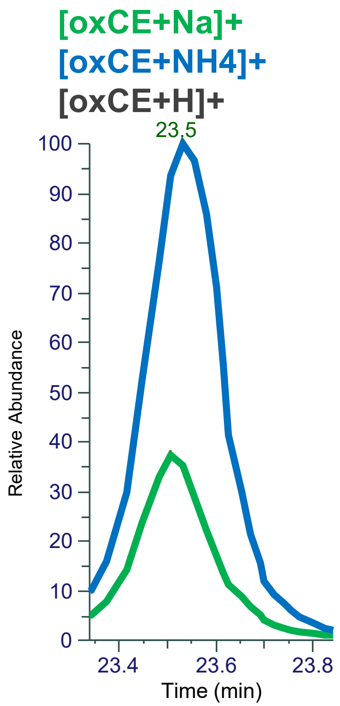


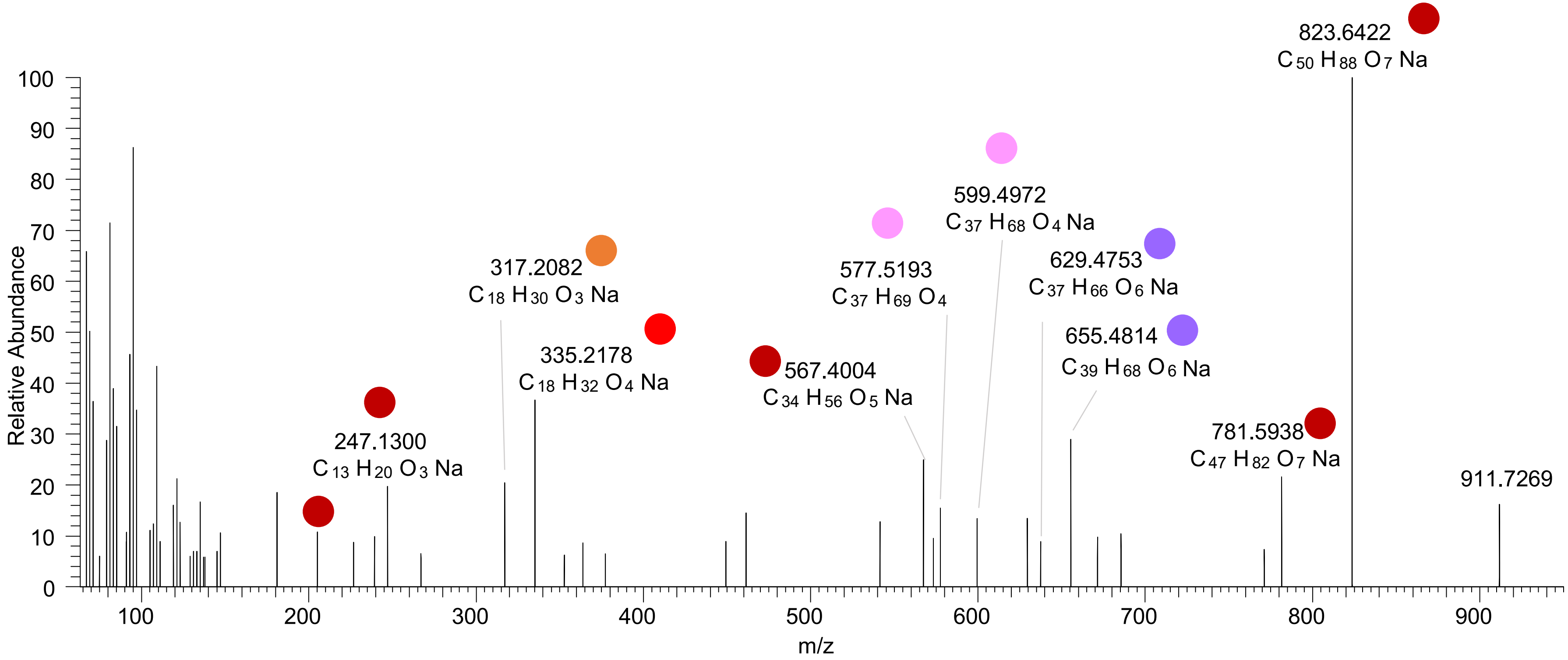

TG(18:2_18:2_18:2<OH>)
RT 22.5

[oxTG+Na]+ XIC 917.7204 NL: 5.85E5

Fragments containing oxFAs
Fragments related to water loss
Fragments not containing oxFAs
Fragments related to other oxLPPs
Position-specific fragments
Fragments related to FA loss
Fragments related to oxFA loss

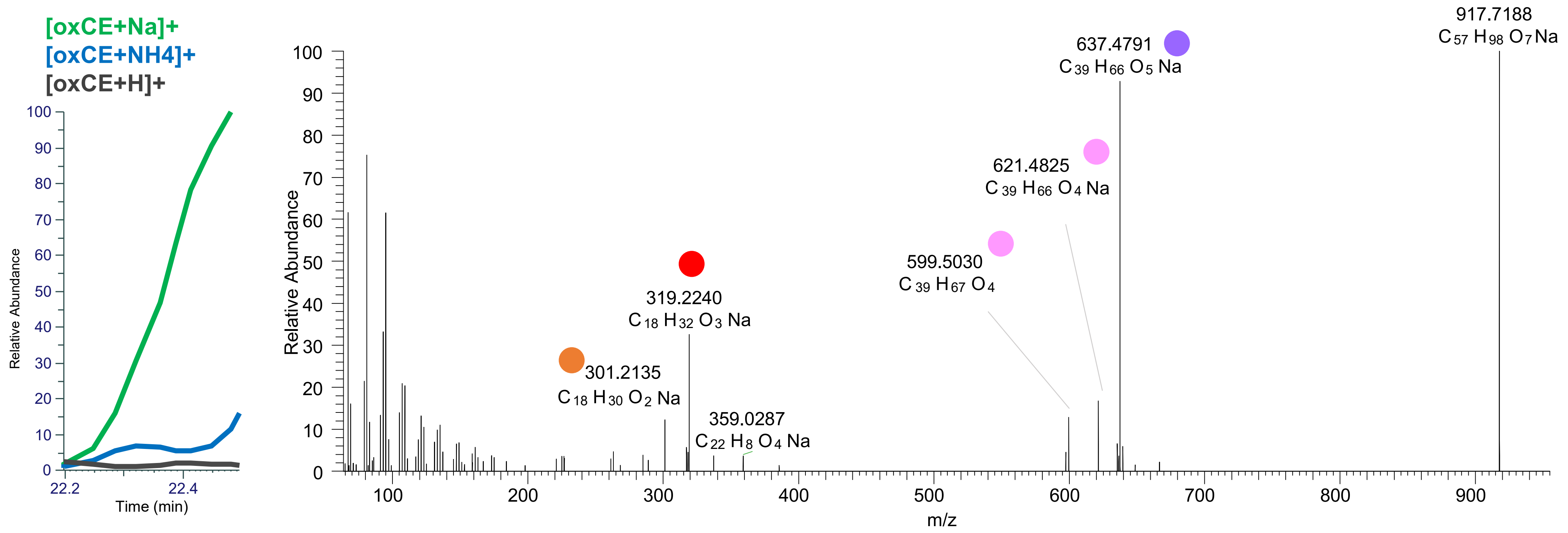

TG(18:2_18:2_18:2<OH>)
TG(18:1_18:2_18:3<OH>)
RT 22.7

[oxTG+Na]+ XIC 917.7204 NL: 7.14E5

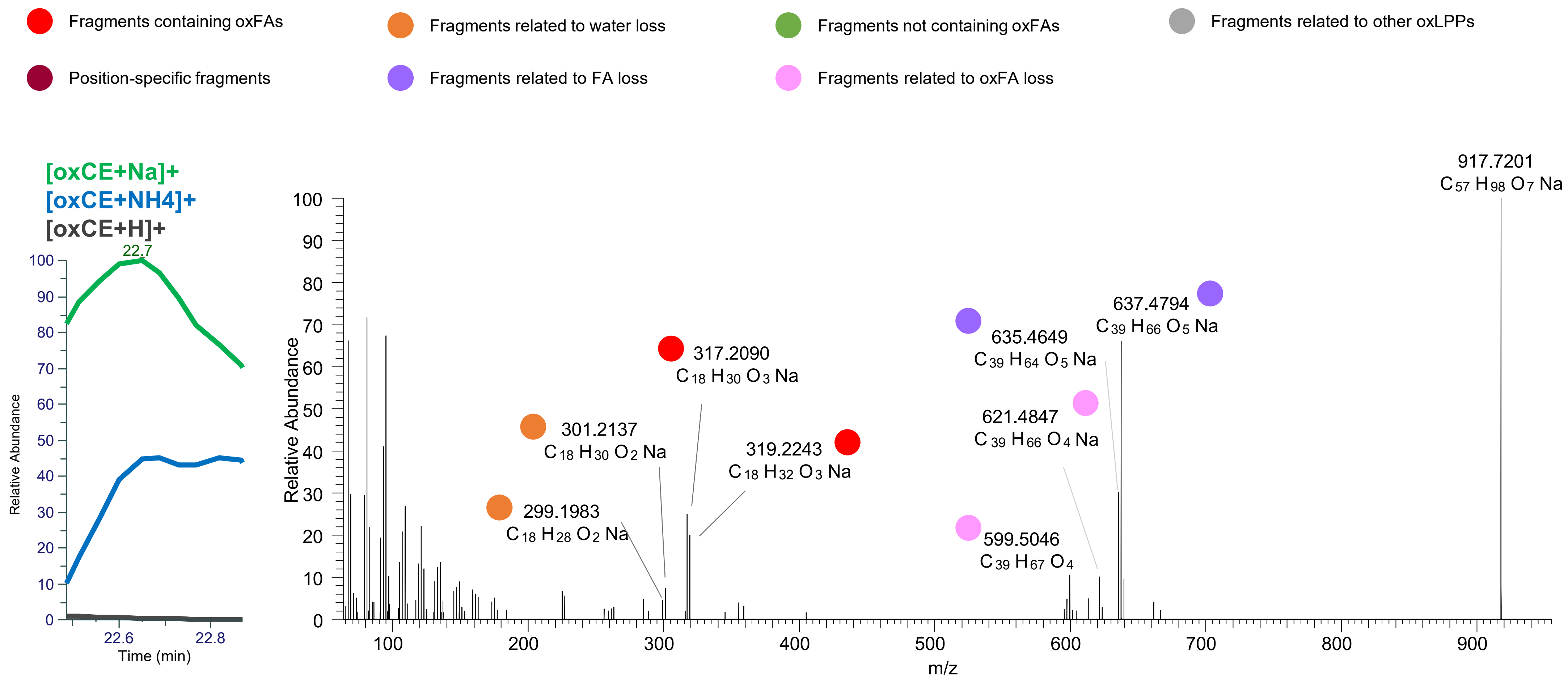

TG(18:1_18:2_18:3<OH>)
TG(18:2_18:2_18:2<OH>)
TG(16:0_18:2_20:4<O>)
RT 22.9

[oxTG+Na]+ XIC 917.7204 NL: 7.14E5

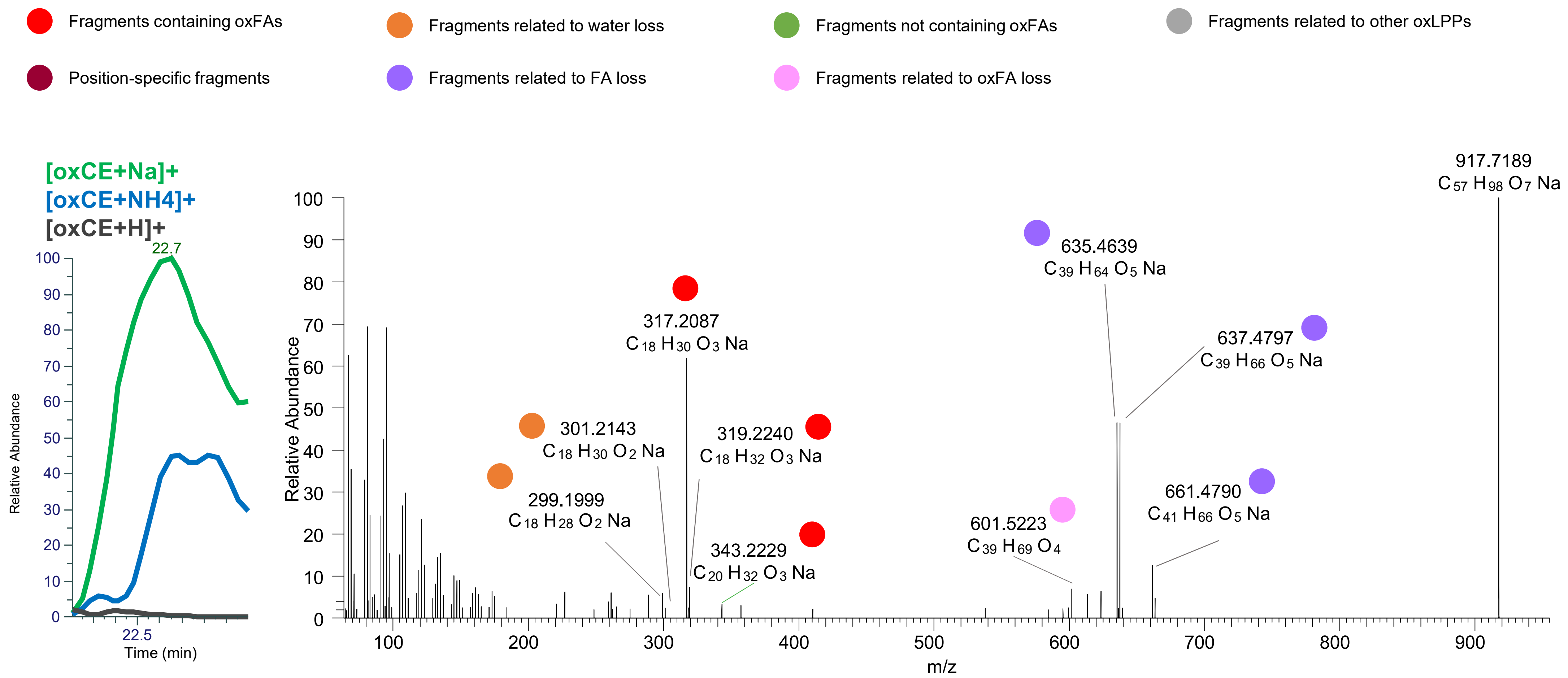

# TG(18:1_18:2_18:2<OH>)
# RT 23.2

**[oxTG+Na]+** **XIC 919.7361 NL: 1.35E6**

- Fragments containing oxFAs
- Fragments related to water loss
- Fragments not containing oxFAs
- Fragments related to other oxLPPs
- Position-specific fragments
- Fragments related to FA loss
- Fragments related to oxFA loss

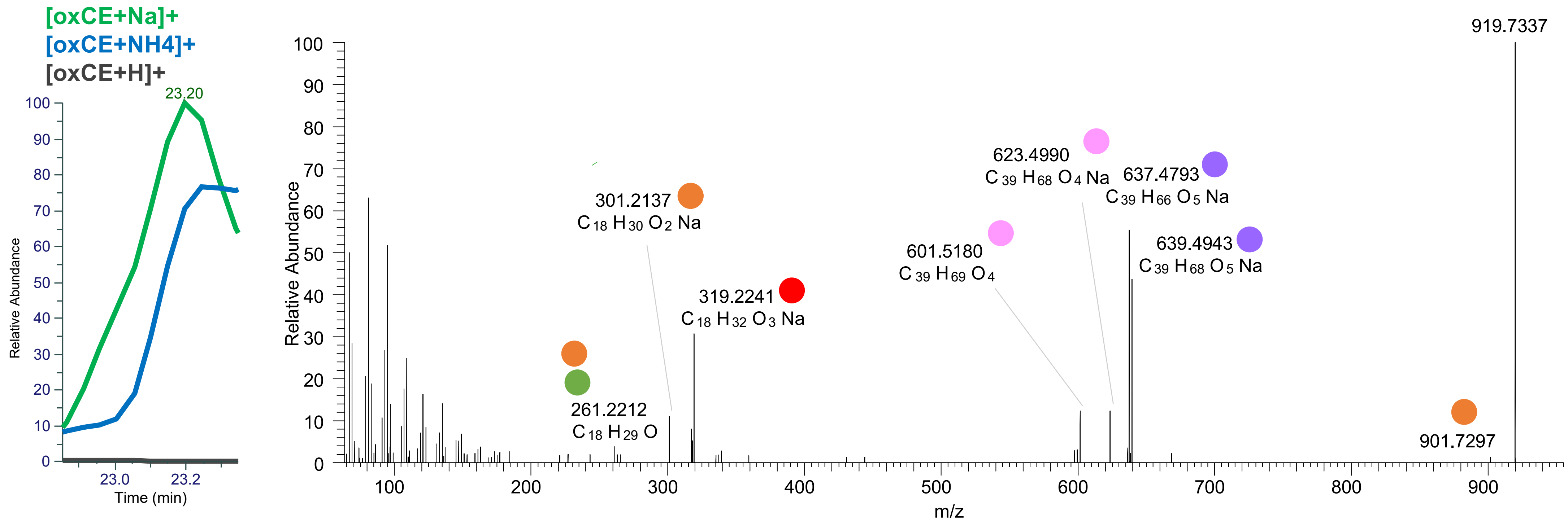

# TG(18:1_18:2_18:2<OH>)
# TG(18:1_18:1_18:3<OH>)
# TG(16:0_18:2_20:3<O>)
# RT 23.3

**[oxTG+Na]+** **XIC 919.7361 NL: 1.35E6**

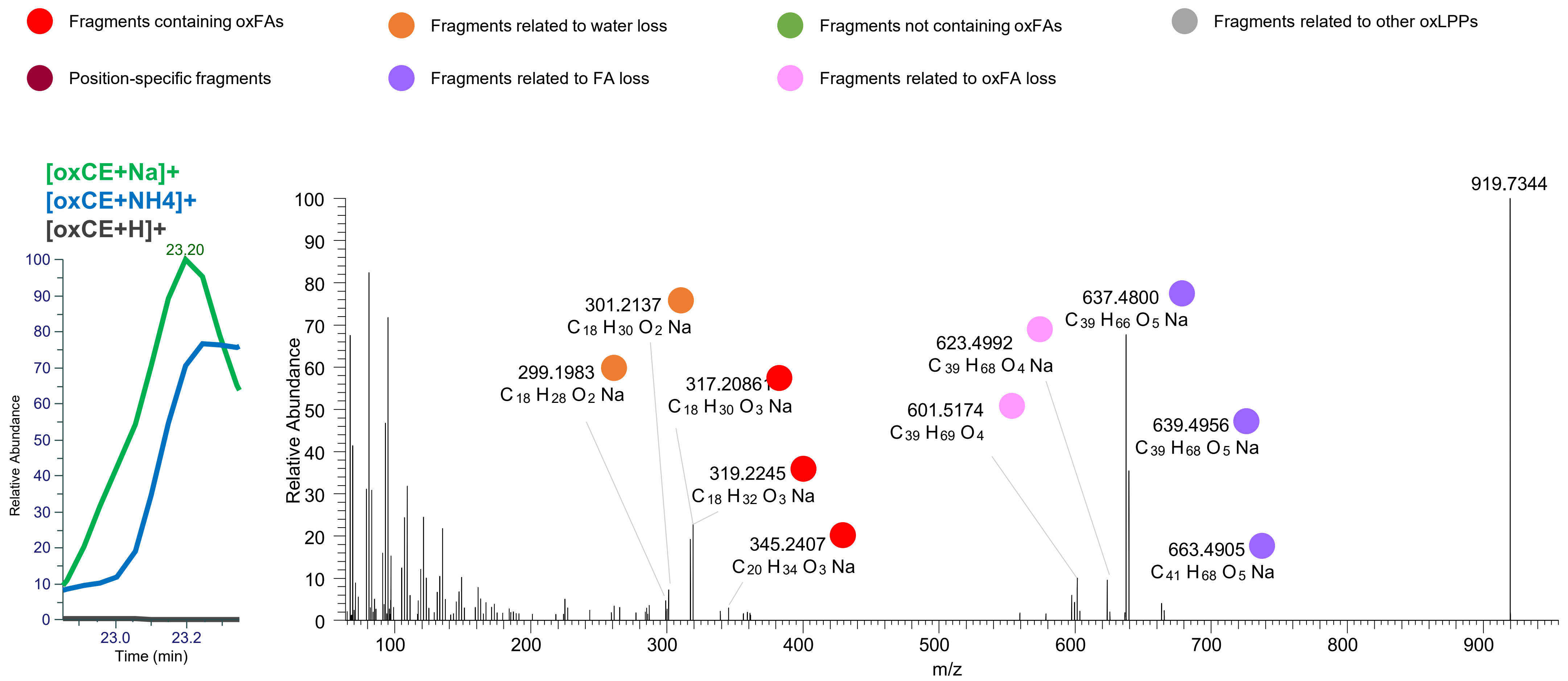

# TG(18:1_18:1_18:3<OH>)
### TG(16:0_18:1_20:3<oxo>)
# RT 23.6

**[oxTG+Na]+** **XIC 919.7361 NL: 5.74E5**

Fragments containing oxFAs
Fragments related to water loss
Fragments not containing oxFAs
Fragments related to other oxLPPs
Position-specific fragments
Fragments related to FA loss
Fragments related to oxFA loss

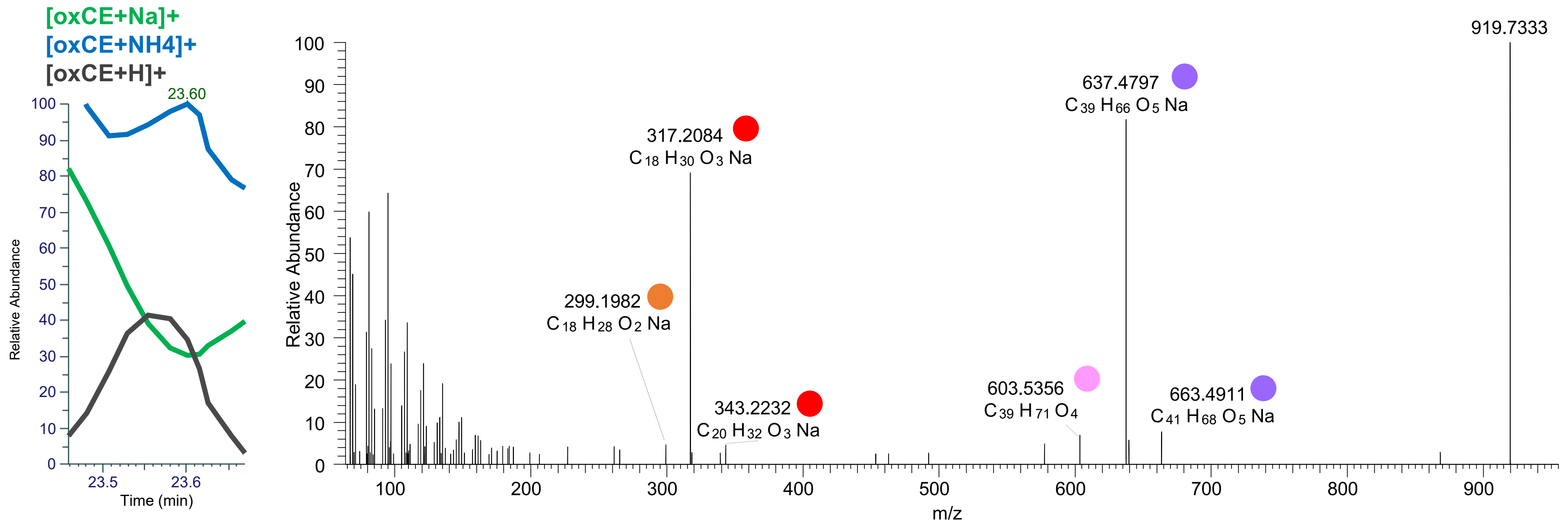

# TG(18:1_18:1_18:3<OH>)
# TG(16:0_18:1_20:4<OH>)
# RT 24.0

**[oxTG+Na]+** **XIC 919.7361 NL: 8.25E5**

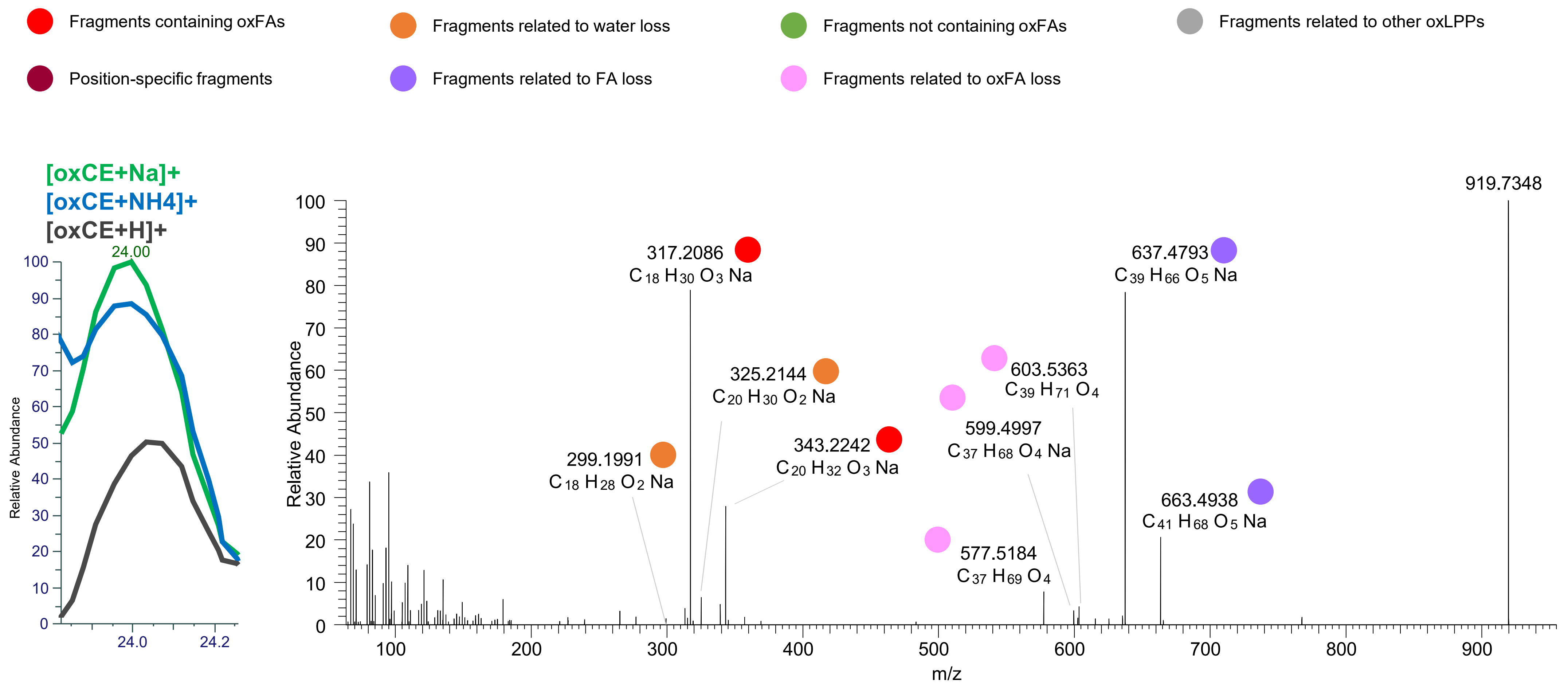

# TG(18:1_18:1_18:2<ep>)
# RT 25.4

**[oxTG+Na]+** **XIC 919.7361 NL: 1.17E5**

Fragments containing oxFAs
Fragments related to water loss
Fragments not containing oxFAs
Fragments related to other oxLPPs
Position-specific fragments
Fragments related to FA loss
Fragments related to oxFA loss

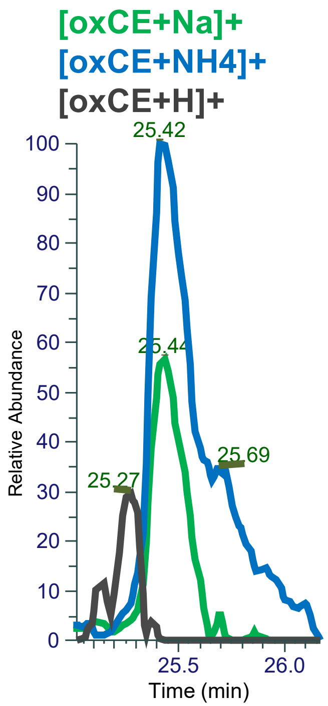


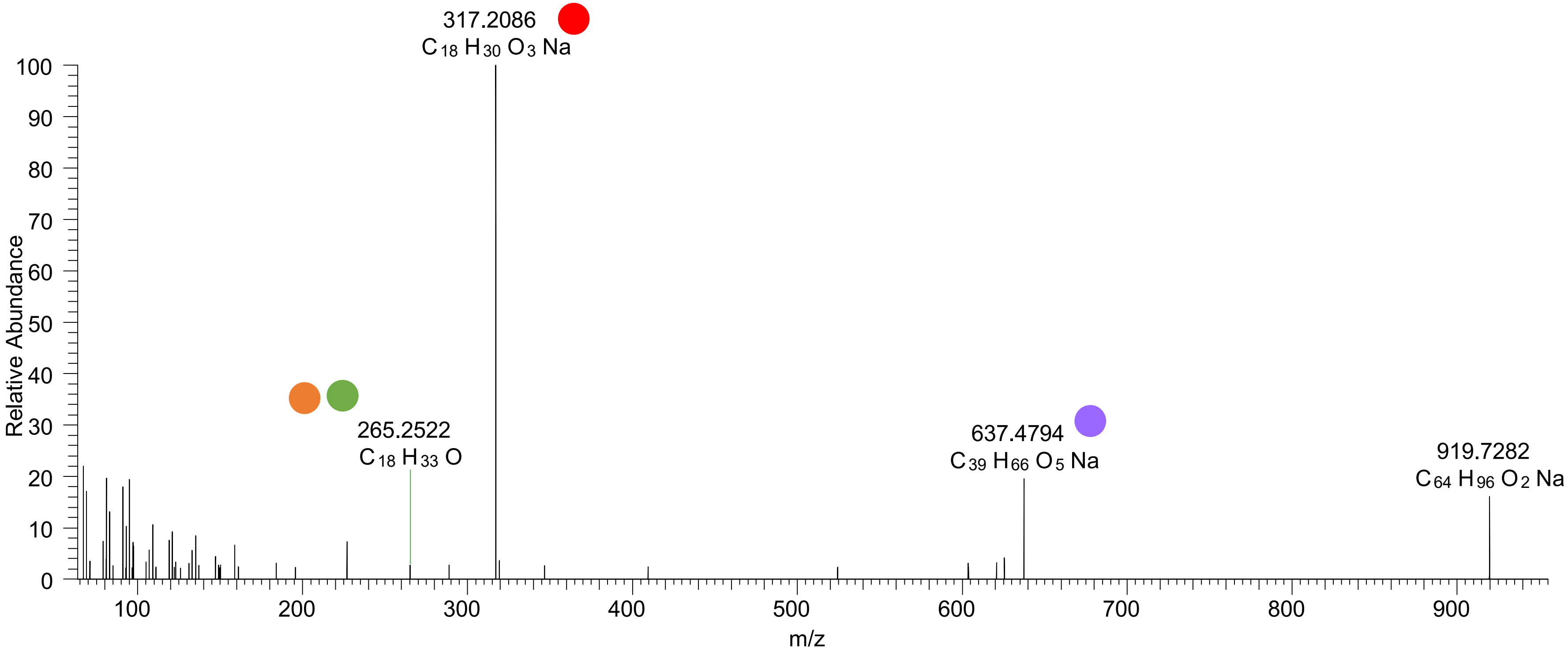

# TG(18:1_18:1_18:2<OH>)
# TG(18:0_18:1_18:3<OH>)
# RT 23.4

**[oxTG+Na]+** **XIC 921.7517 NL: 4.46E5**

- Fragments containing oxFAs
- Fragments related to water loss
- Fragments not containing oxFAs
- Fragments related to other oxLPPs
- Position-specific fragments
- Fragments related to FA loss
- Fragments related to oxFA loss

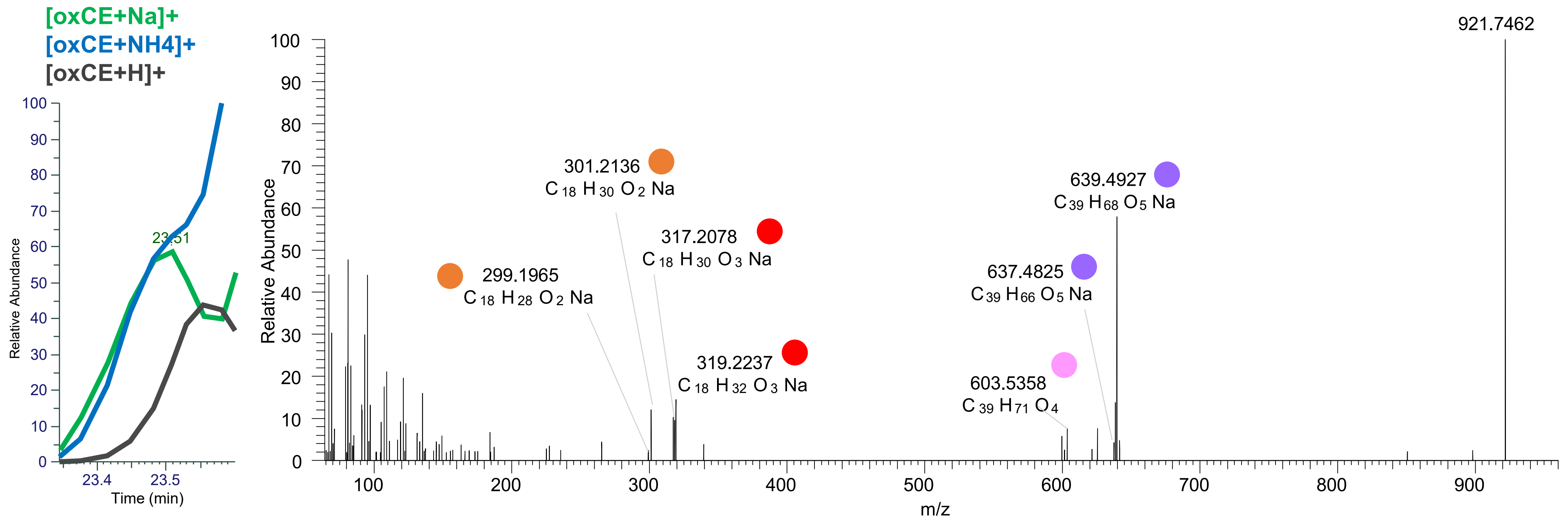

# TG(18:1_18:1_18:2<OH>)
# RT 23.7

**[oxTG+Na]+** **XIC 921.7517 NL: 2.21E6**

- Fragments containing oxFAs
- Fragments related to water loss
- Fragments not containing oxFAs
- Fragments related to other oxLPPs
- Position-specific fragments
- Fragments related to FA loss
- Fragments related to oxFA loss

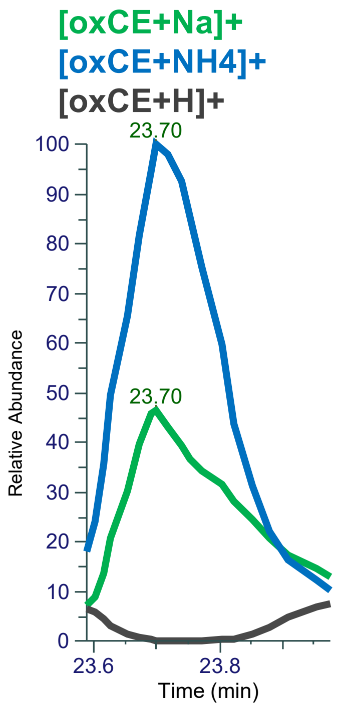


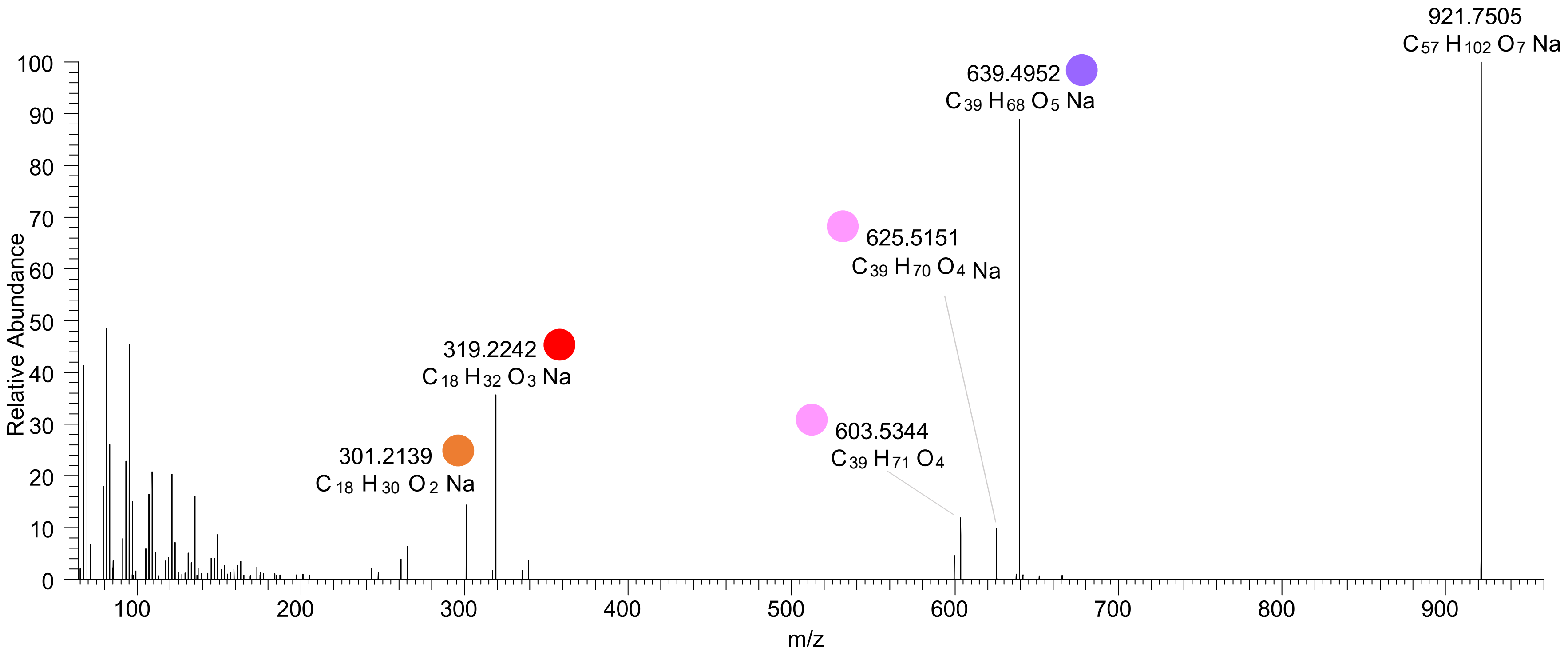

# TG(18:0_18:2_18:2<OH>)
# TG(16:0_18:1_20:3<OH>)
# RT 23.7

**[oxTG+Na]+** **XIC 921.7517 NL: 2.21E6**

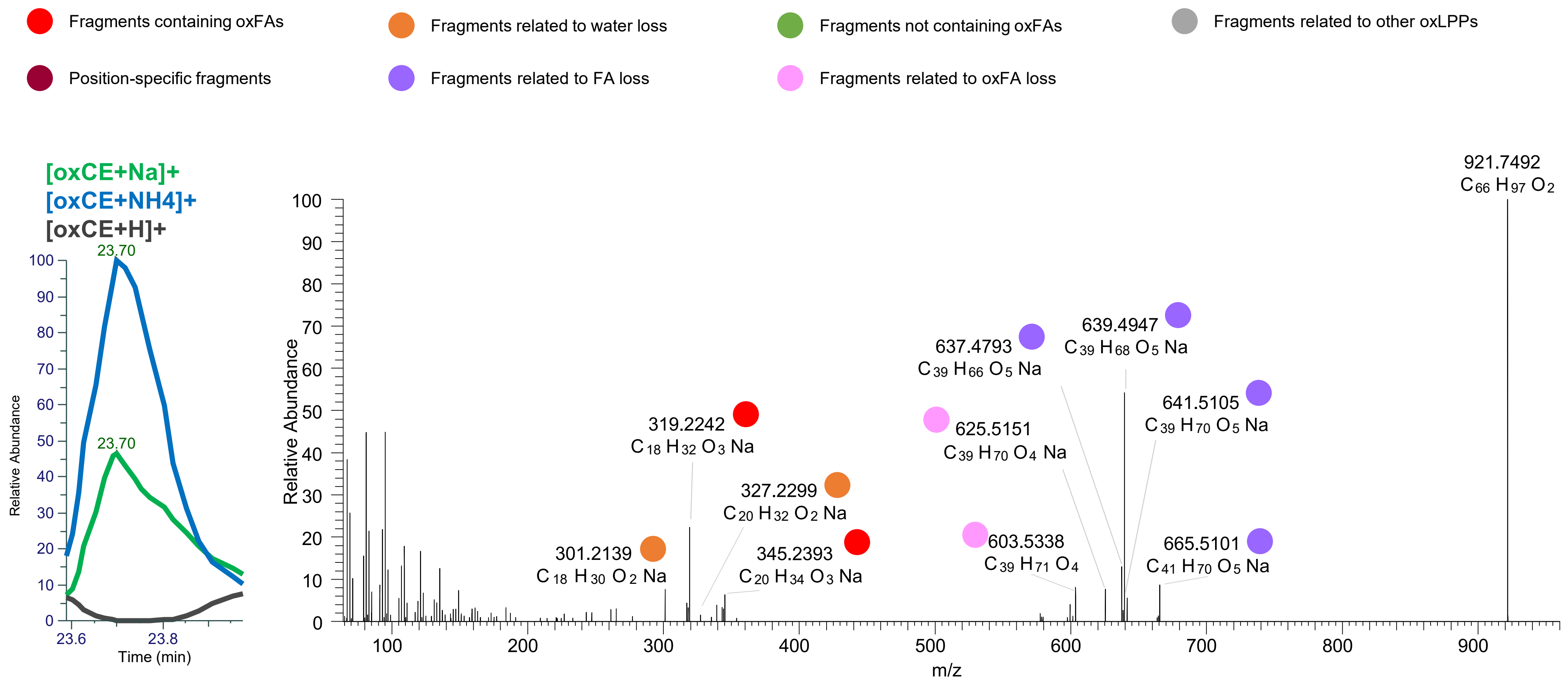

# TG(18:1_18:1_18:2<O>)
# TG(18:0_18:1_18:3<O>)
# RT 24.9

**[oxTG+Na]+** **XIC 921.7517 NL: 1.38E5**

- Fragments containing oxFAs
- Fragments related to water loss
- Fragments not containing oxFAs
- Fragments related to other oxLPPs
- Position-specific fragments
- Fragments related to FA loss
- Fragments related to oxFA loss

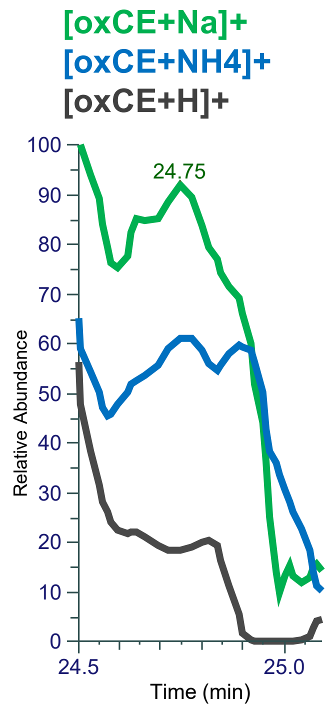


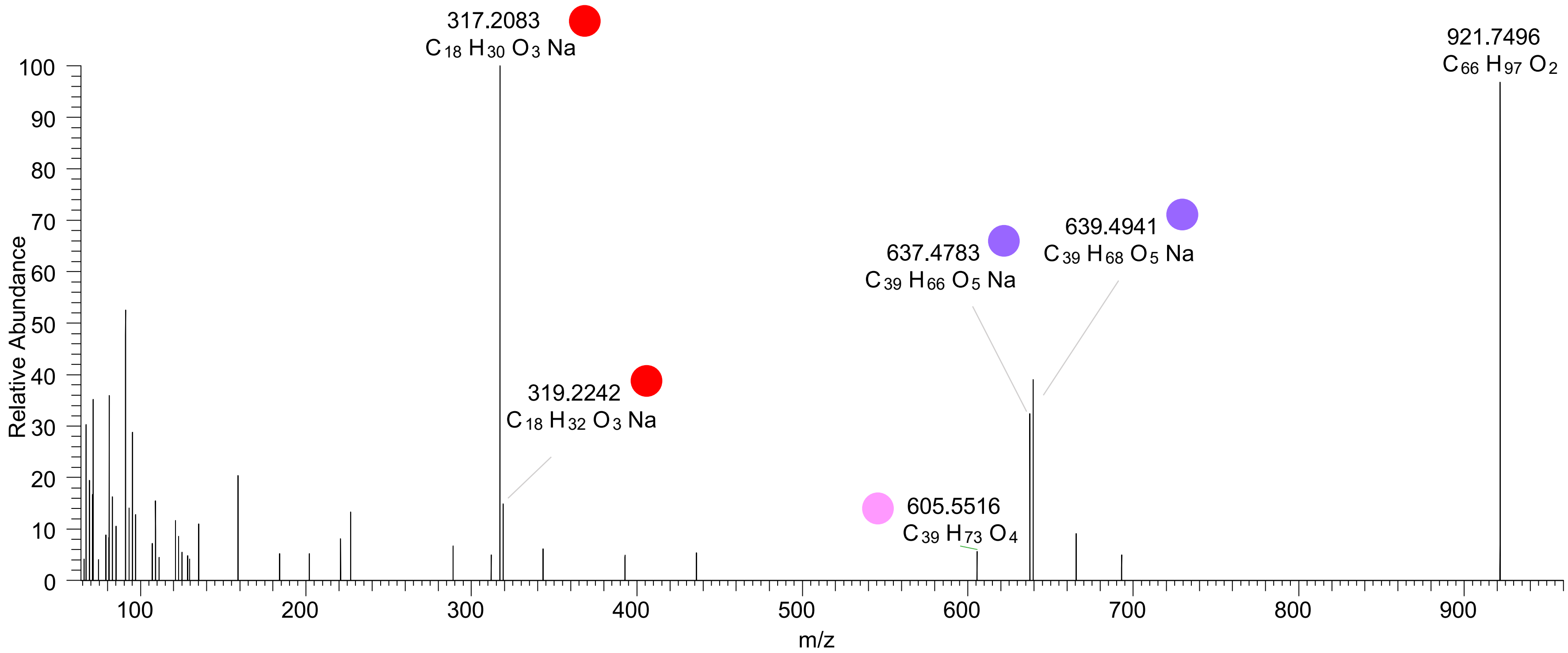

# TG(16:0_18:1_18:3<3O>)
# RT 21.8

**oxTG+Na]$^+$** **XIC 925.7103 NL: 4.70E4**

- Fragments containing oxFAs
- Fragments related to water loss
- Fragments not containing oxFAs
- Fragments related to other oxLPPs
- Position-specific fragments
- Fragments related to FA loss
- Fragments related to oxFA loss

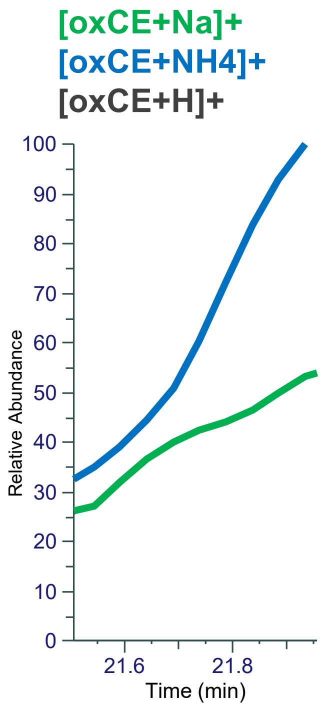


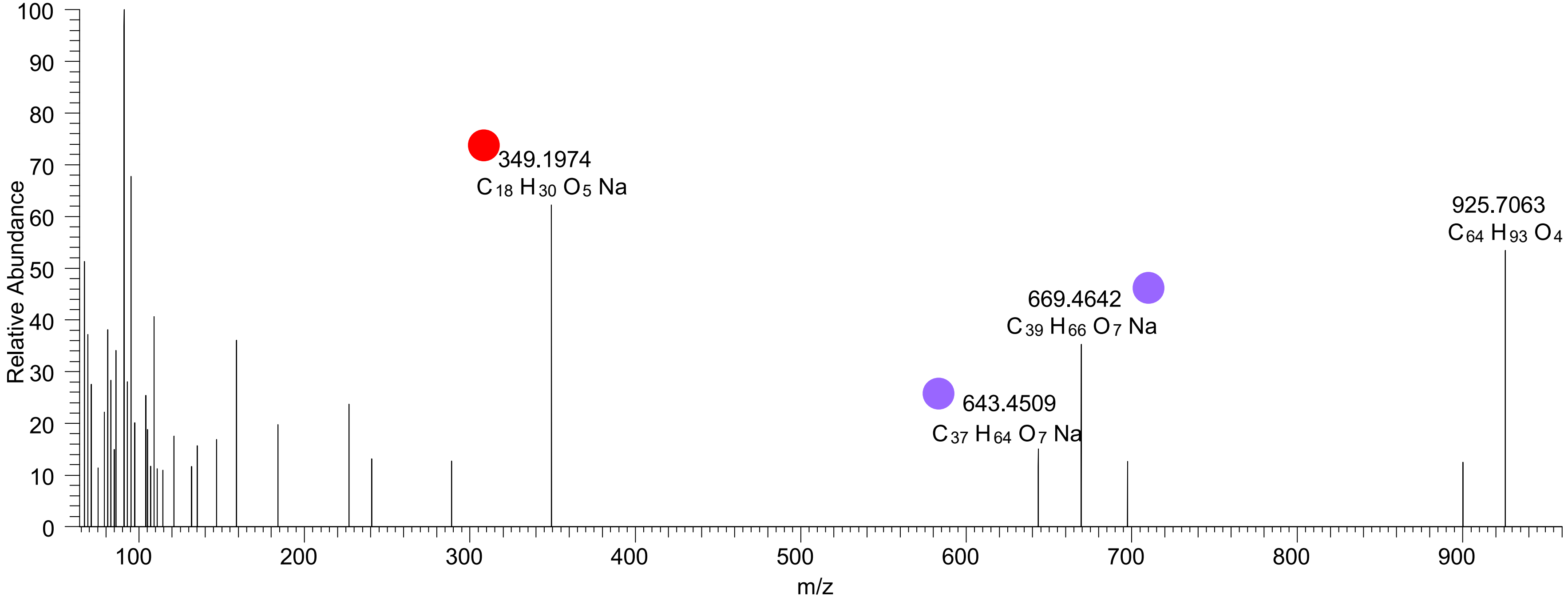

# TG(16:0_18:1_18:3<3O>)
# RT 23.4

**oxTG+Na]$^+$**

**XIC 925.7103 NL: 4.89E4**

- Fragments containing oxFAs
- Fragments related to water loss
- Fragments not containing oxFAs
- Fragments related to other oxLPPs
- Position-specific fragments
- Fragments related to FA loss
- Fragments related to oxFA loss

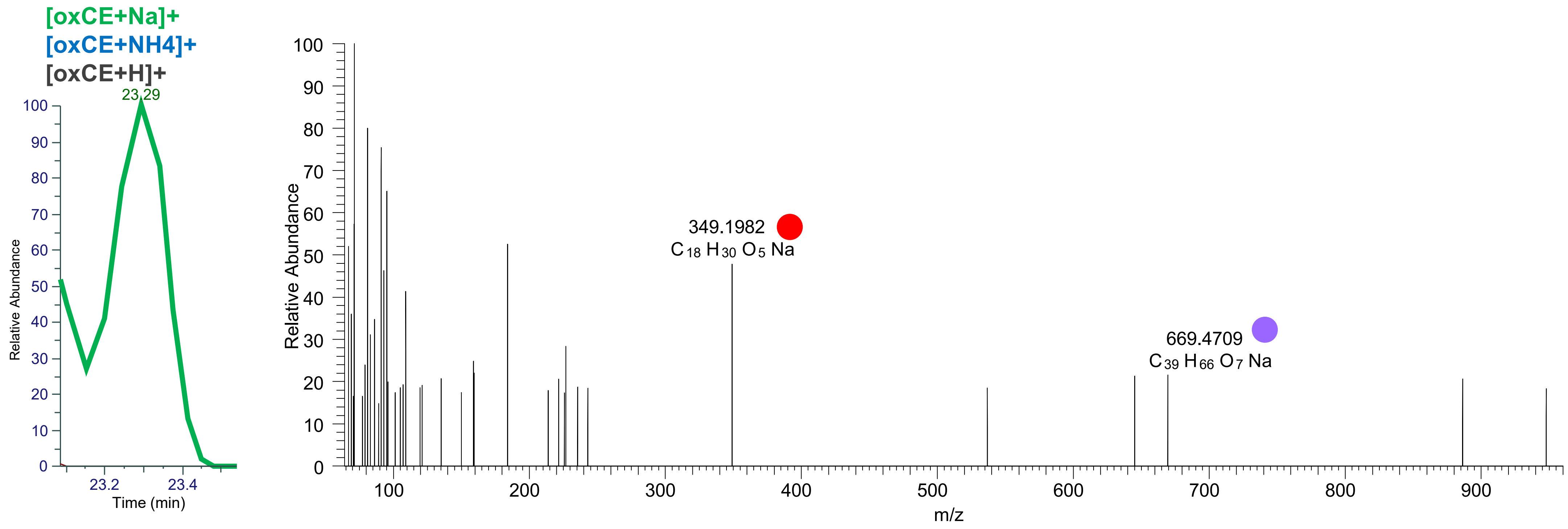

# TG(18:1_18:2<O>_18:3<O>)
# RT 20.6

**[oxTG+Na]+** **XIC 933.7154 NL: 1.13E5**

Fragments containing oxFAs
Fragments related to water loss
Fragments not containing oxFAs
Fragments related to other oxLPPs
Position-specific fragments
Fragments related to FA loss
Fragments related to oxFA loss

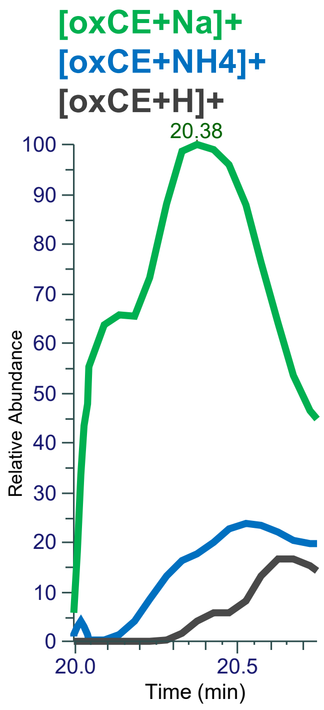


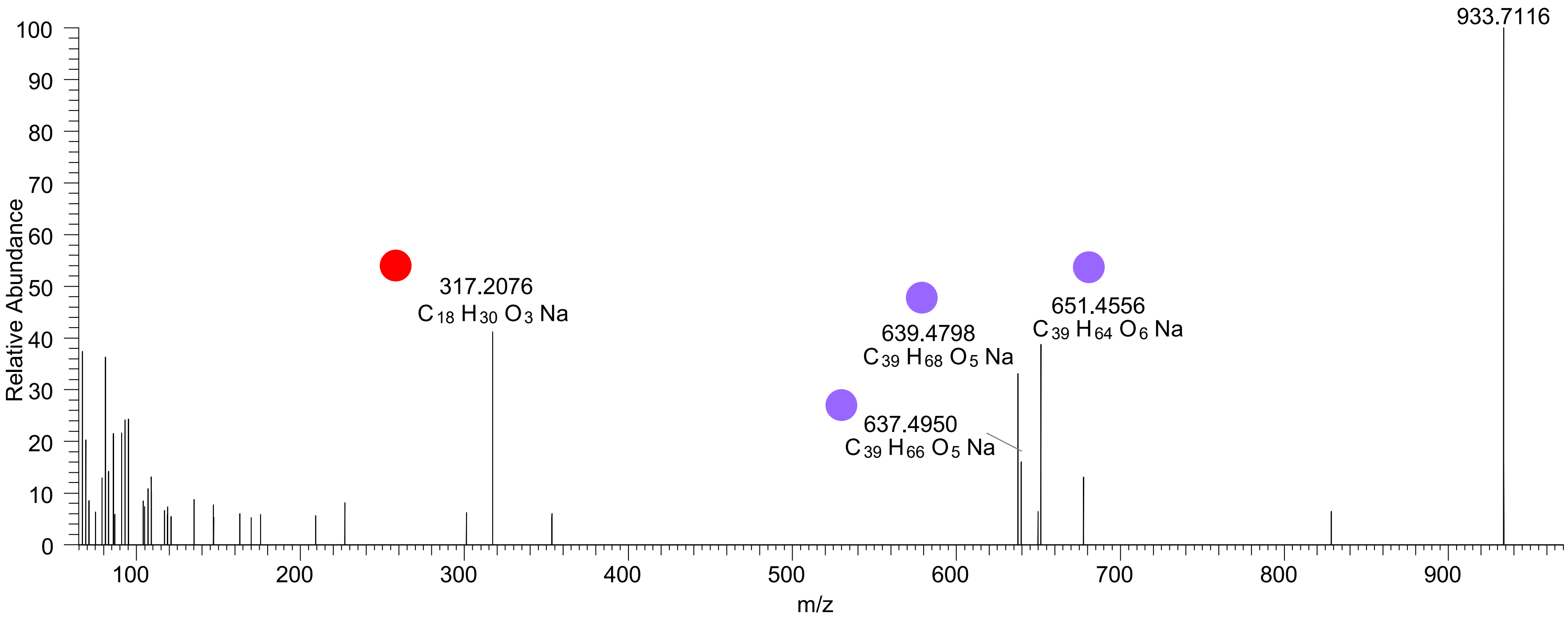

# TG(16:1_18:1_20:4<OH,O>)
# TG(18:1_18:2_18:3<OH,O>)
# RT 21.0

**[oxTG+Na]+** **XIC 933.7154 NL: 1.69E5**

- Fragments containing oxFAs
- Fragments related to water loss
- Fragments not containing oxFAs
- Fragments related to other oxLPPs
- Position-specific fragments
- Fragments related to FA loss
- Fragments related to oxFA loss

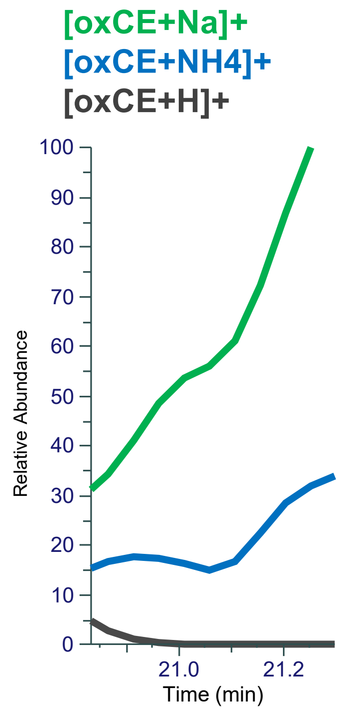


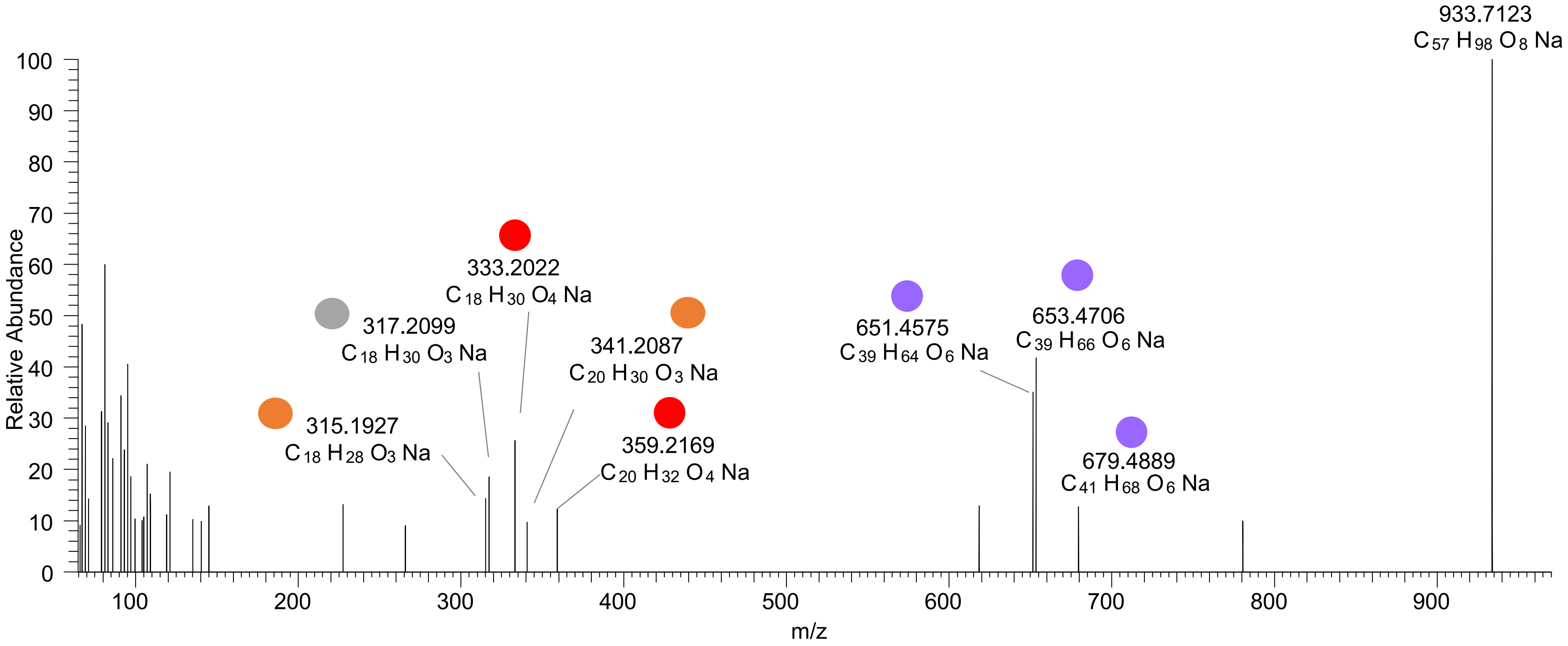

TG(16:1_18:1_20:4<OH,O>)
TG(18:2_18:2_18:2<2O>)
TG(18:1_18:2_18:3<OH,O>)
TG(18:1_18:1_18:4<2O>)
TG(16:0_16:1_22:5<2O>) RT 21.2

[oxTG+Na]+ XIC 933.7154 NL: 1.69E5

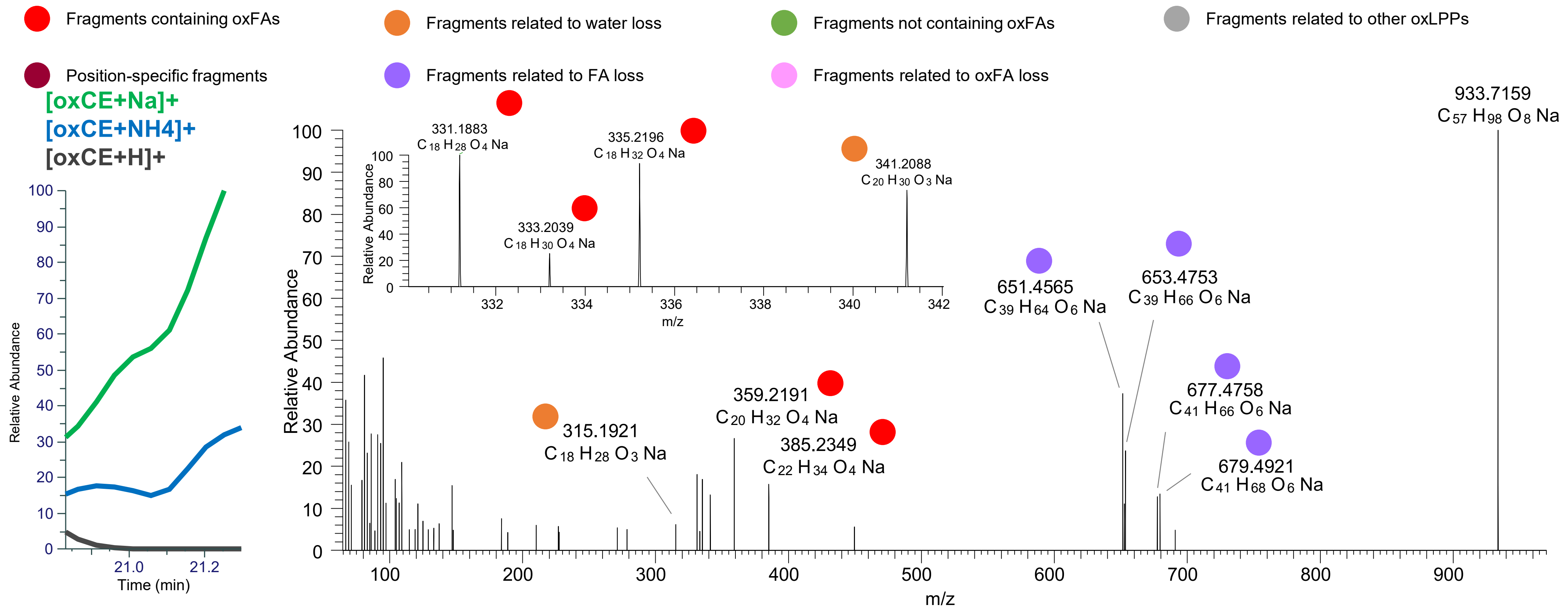

TG(16:0_18:1_20:5<OH,O>)
TG(16:1_18:1_20:4<OH,O>)
TG(16:0_18:2_20:4<OH,O>)
TG(18:1_18:1_18:4<2O>)
TG(18:1_18:2_18:3<2O>)
TG(18:1_18:3_18:2<OH,O>) RT 21.7

**[oxTG+Na]+ XIC 933.7154 NL: 2.41E5**

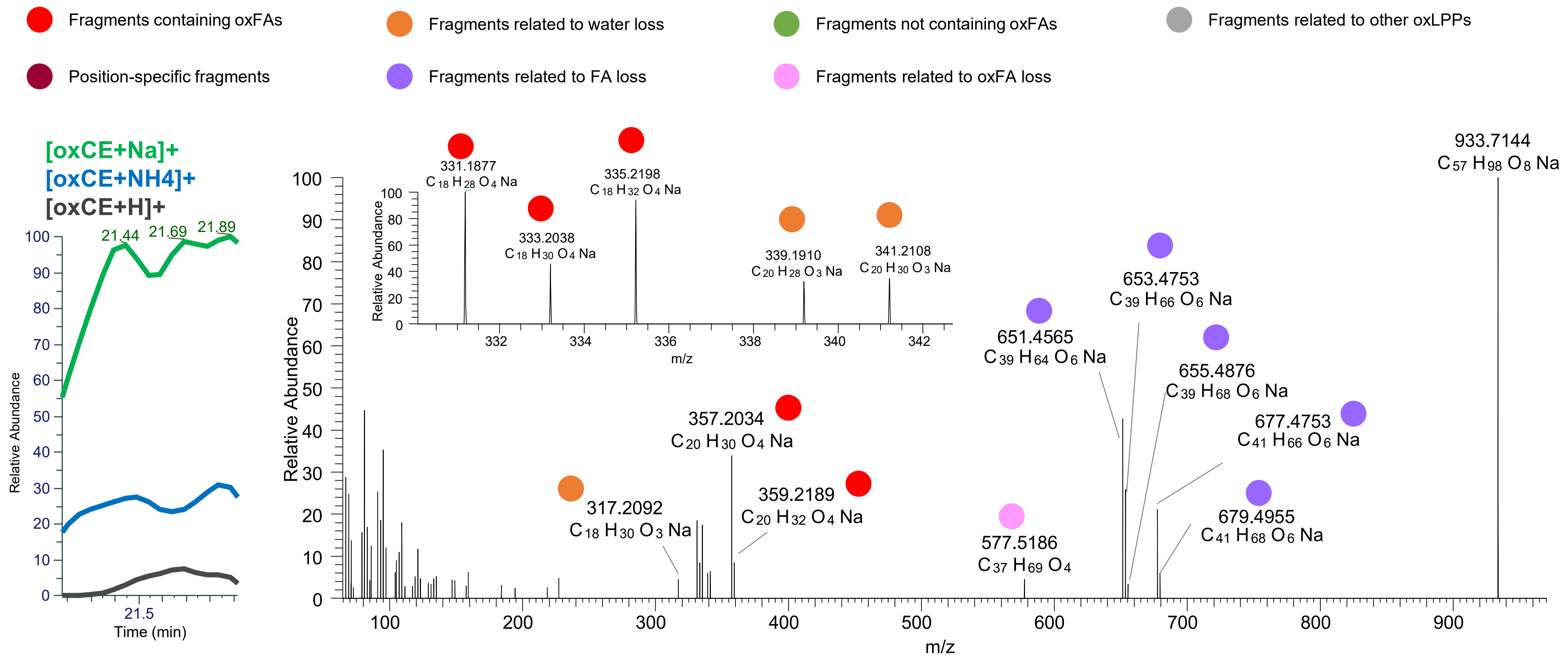

### TG(18:2_18:2_18:2<OOH{13}>)
# RT 22.5

**[oxTG+Na]+** **XIC 933.7154 NL: 5.39E5**

- Fragments containing oxFAs
- Fragments related to water loss
- Fragments not containing oxFAs
- Fragments related to other oxLPPs
- Position-specific fragments
- Fragments related to FA loss
- Fragments related to oxFA loss

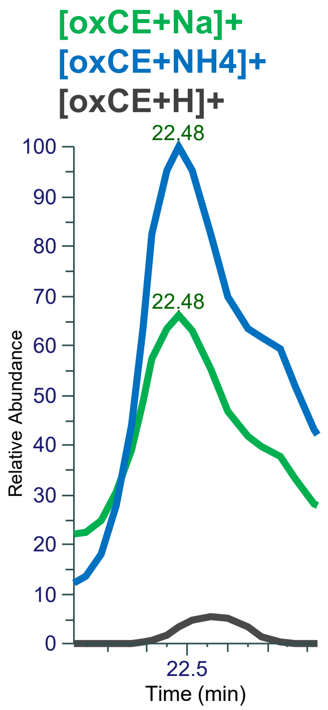


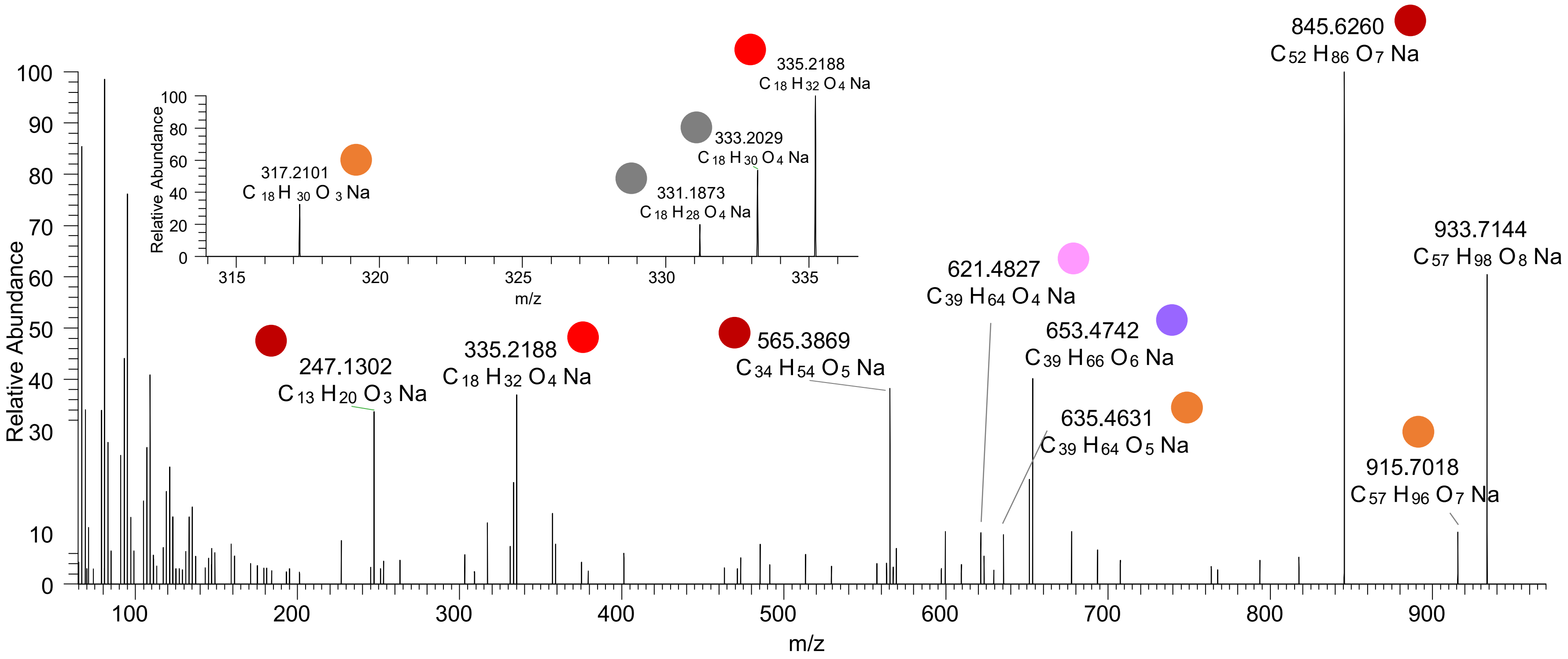

TG(18:2_18:2_18:2<OOH{13}>)
TG(18:1_18:2_18:3<OOH{16}>)
TG(18:1_18:1_18:4<OH,O>)
RT 22.6

[oxTG+Na]+ XIC 933.7154 NL: 5.39E5

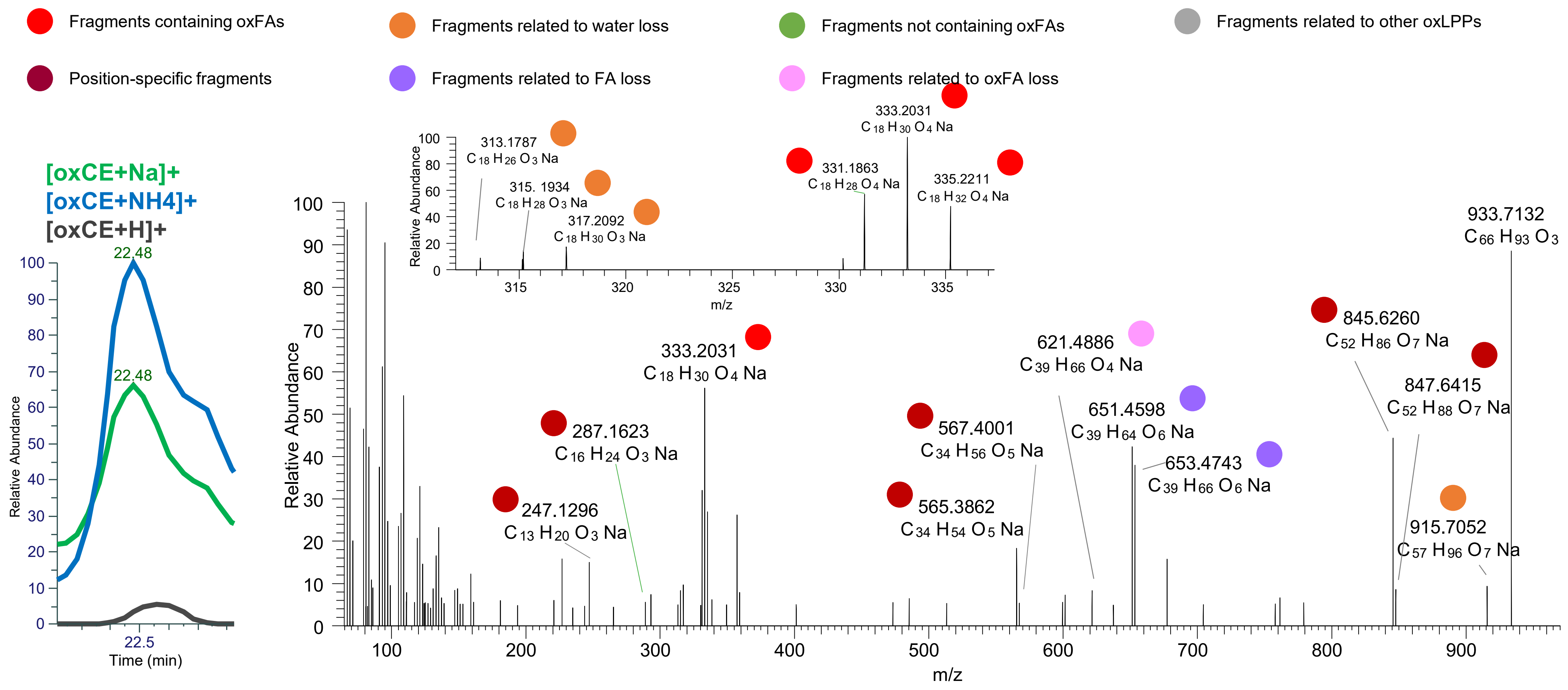

# TG(18:1_18:2<OH>_18:2<OH>)
# RT 20.2

**[oxTG+Na]+** **XIC 935.7310 NL: 3.20E5**

- Fragments containing oxFAs
- Fragments related to water loss
- Fragments not containing oxFAs
- Fragments related to other oxLPPs
- Position-specific fragments
- Fragments related to FA loss
- Fragments related to oxFA loss

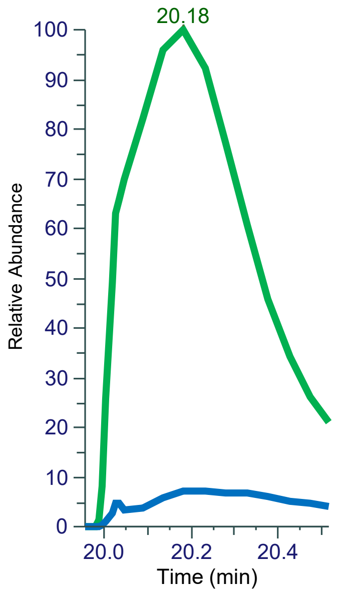


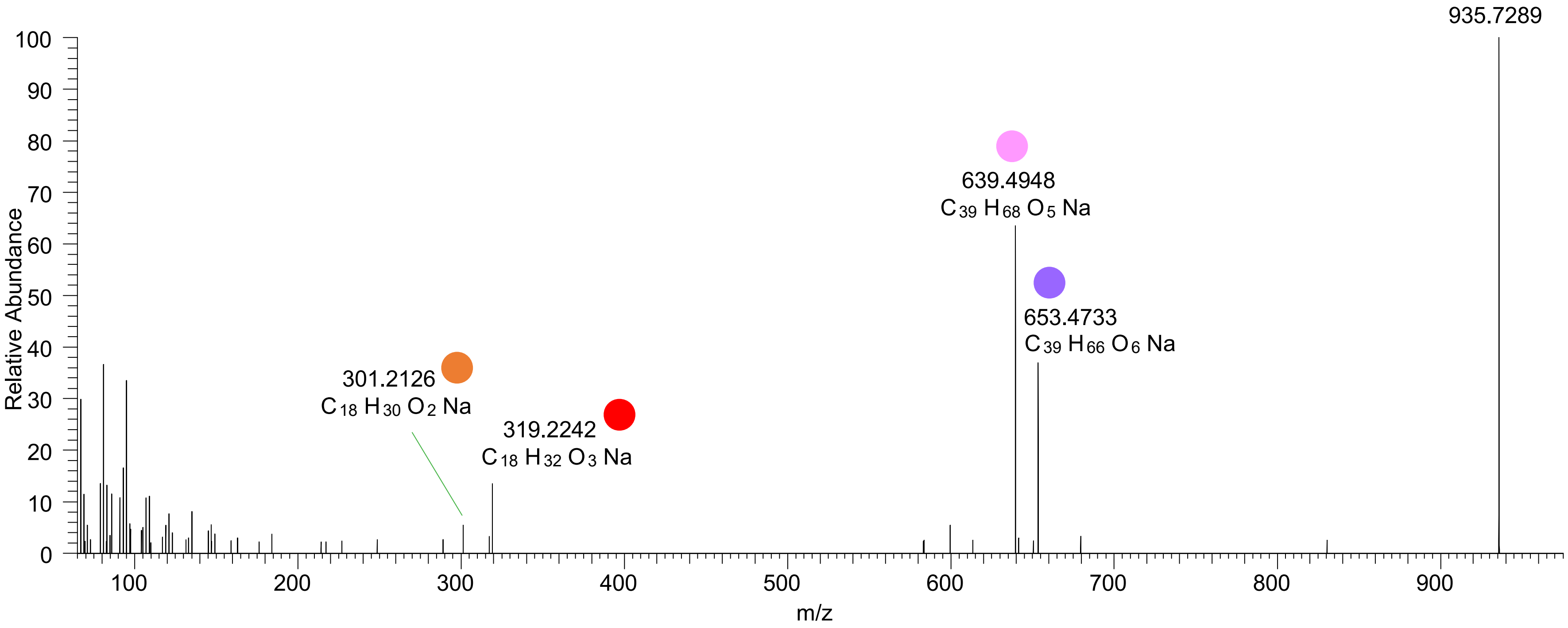

TG(16:0_18:1_20:4<OH,O>)
TG(18:1_18:2_18:2<2O>)
TG(18:1_18:1_18:3<OH,O>)
RT 22.2

**[oxTG+Na]+** **XIC 935.7310 NL: 2.72E6**

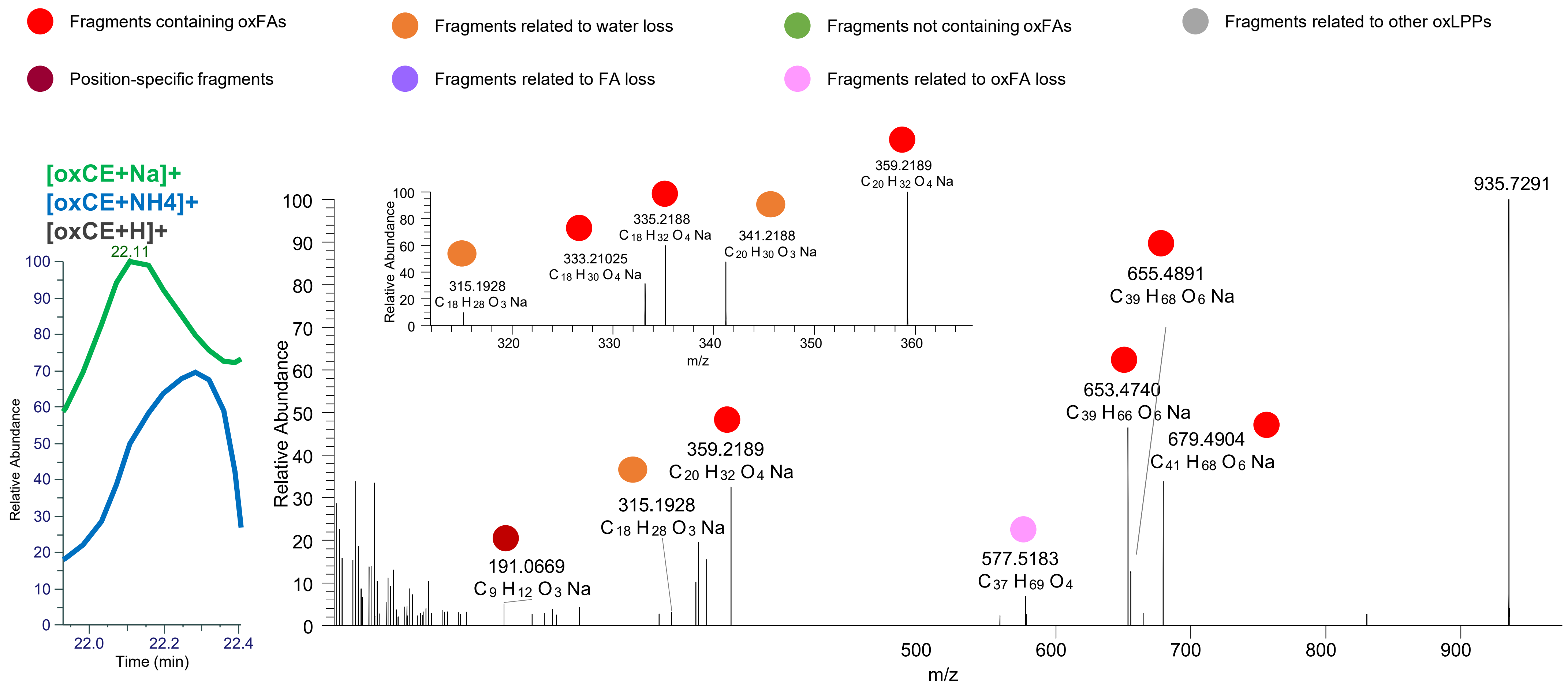

### TG(18:1_18:2_18:2<OOH{13})
# TG(18:1_18:1_18:3<2O>)
# RT 23.1

**[oxTG+Na]+** **XIC 935.7310 NL: 1.45E6**

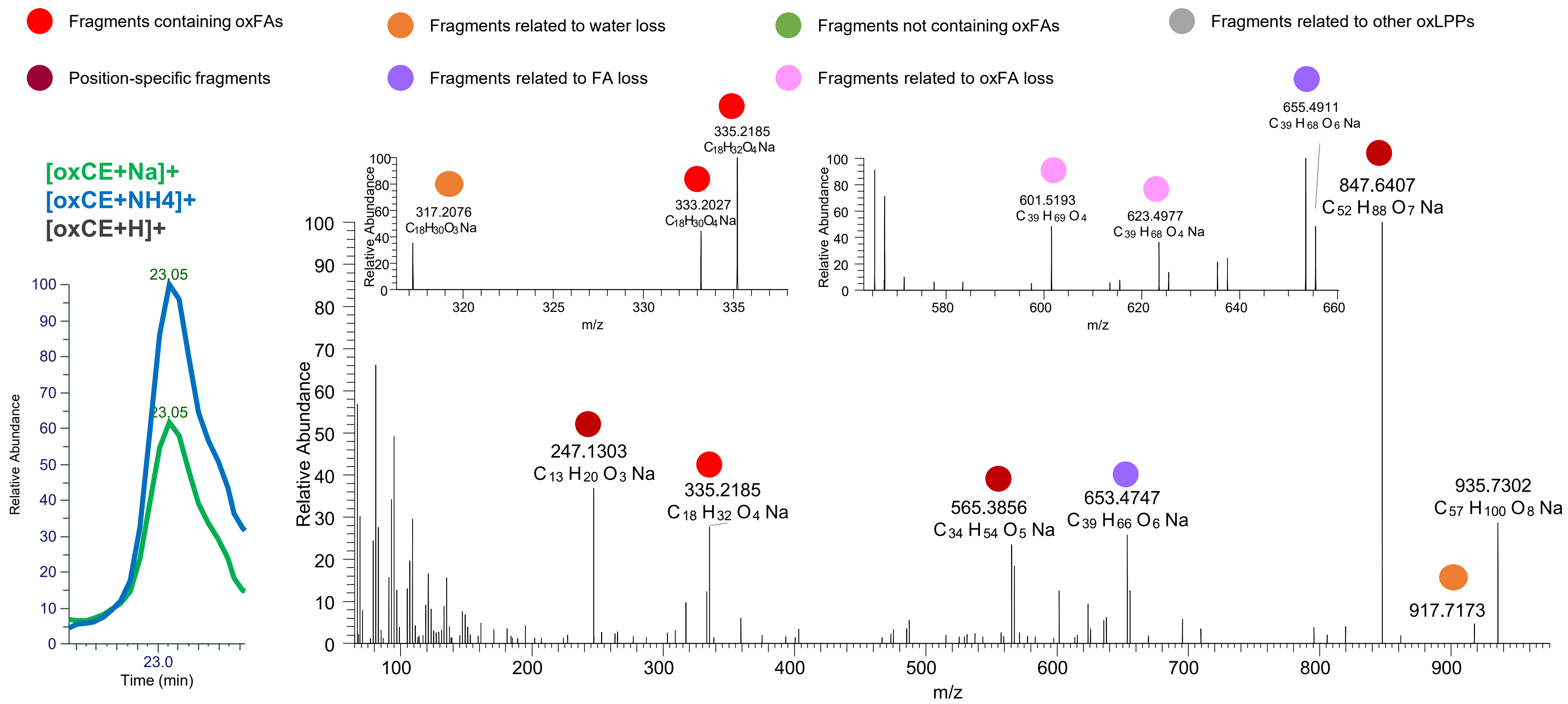

# TG(18:1_18:1_18:2<O,OH>)
# RT 22.8

**[oxTG+Na]+** **XIC 937.7467 NL: 1.35E5**

- Fragments containing oxFAs
- Fragments related to water loss
- Fragments not containing oxFAs
- Fragments related to other oxLPPs
- Position-specific fragments
- Fragments related to FA loss
- Fragments related to oxFA loss

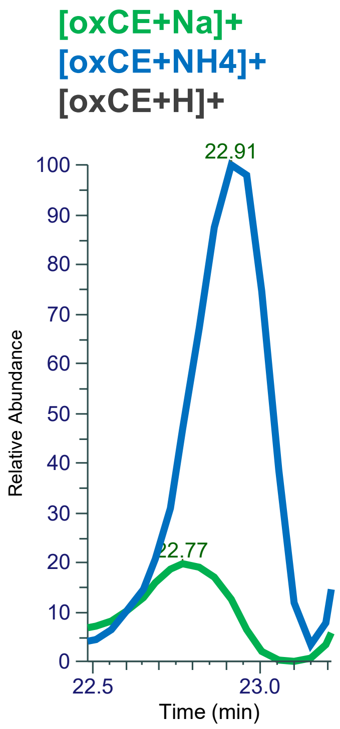


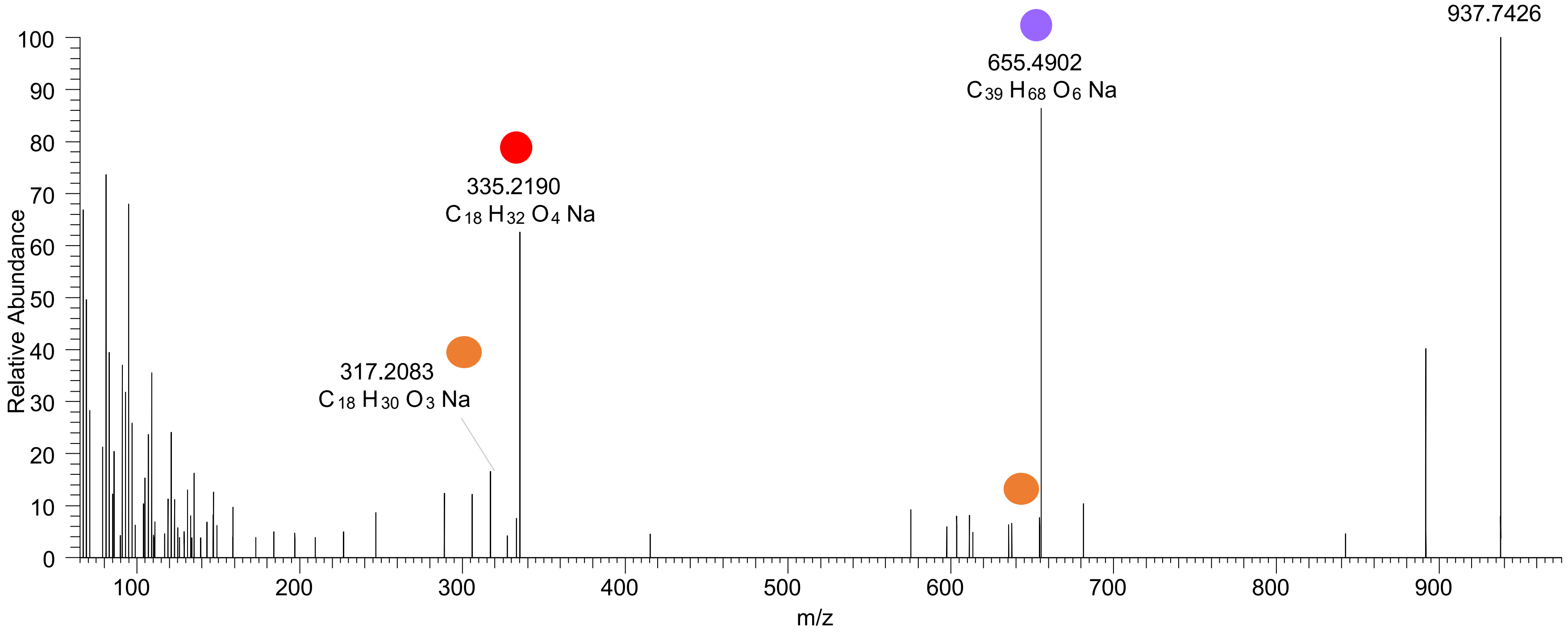

### TG(18:1_18:1_18:2<OOH{13}>)
# RT 23.6

[oxTG+Na]+ XIC 937.7467 NL: 3.40E6

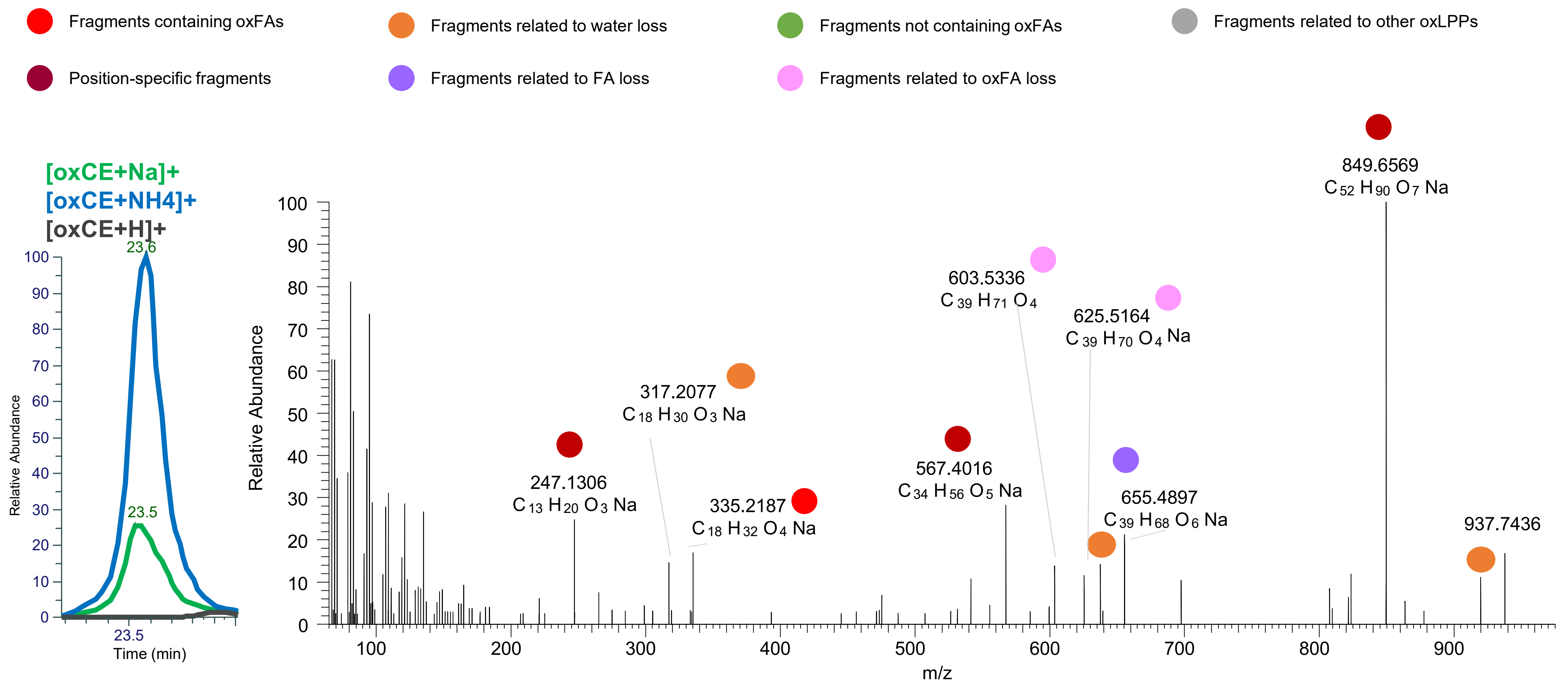

# TG(18:1_18:3<O>_18:2<2O>)
# RT 20.3

**[oxTG+Na]+** **XIC 949.7103 NL: 6.92E4**

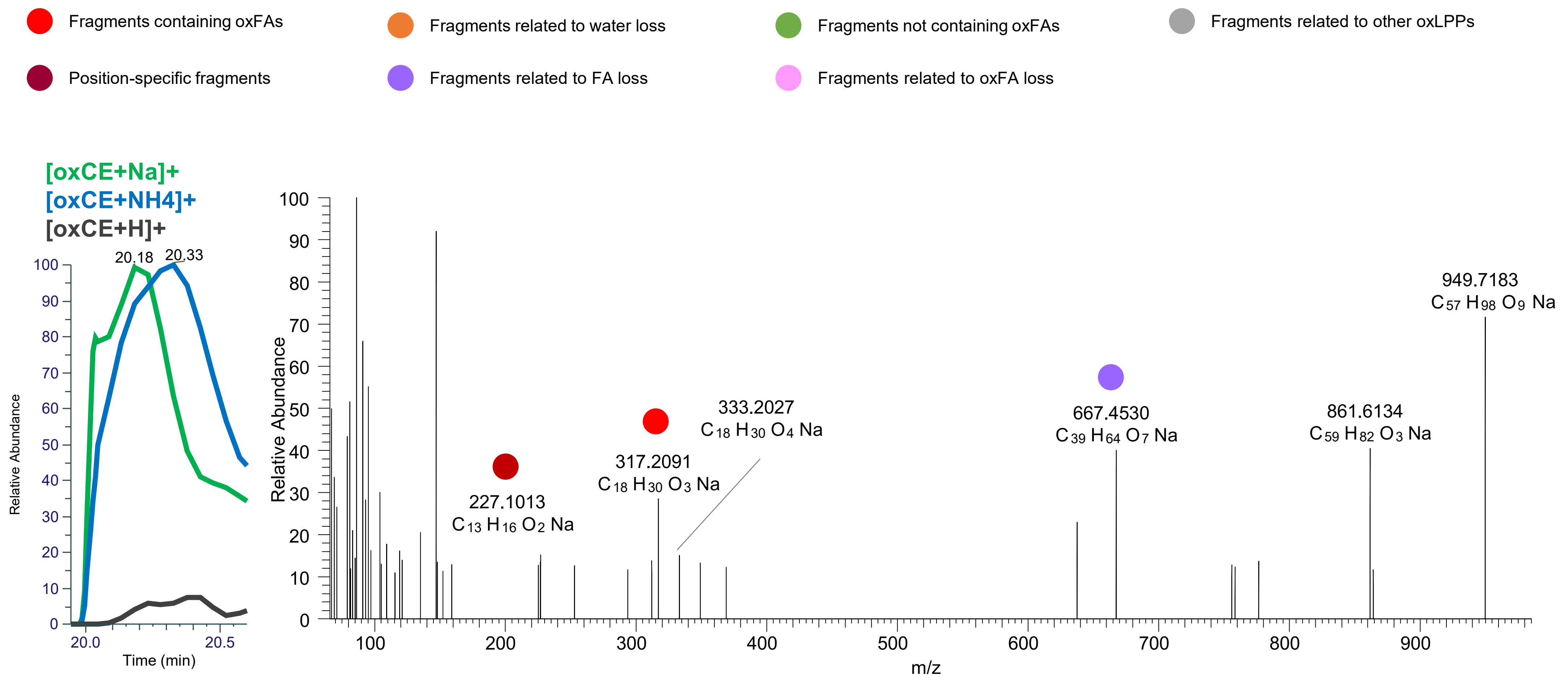

# TG(18:1_18:1_18:4<3O>)
# TG(16:0_18:1_20:5<3O>)
# RT 21.3

**[oxTG+Na]+** **XIC 949.7103 NL: 5.96E4**

- Fragments containing oxFAs
- Fragments related to water loss
- Fragments not containing oxFAs
- Fragments related to other oxLPPs
- Position-specific fragments
- Fragments related to FA loss
- Fragments related to oxFA loss

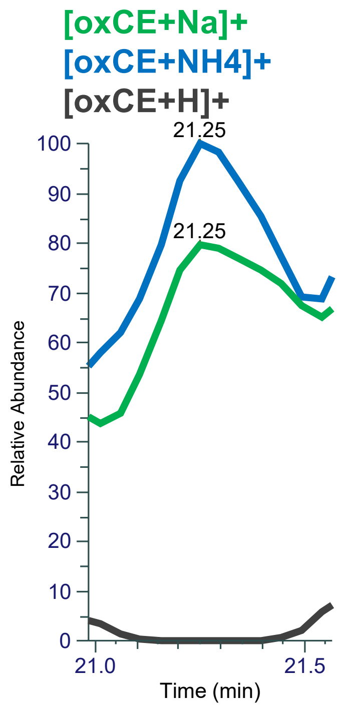


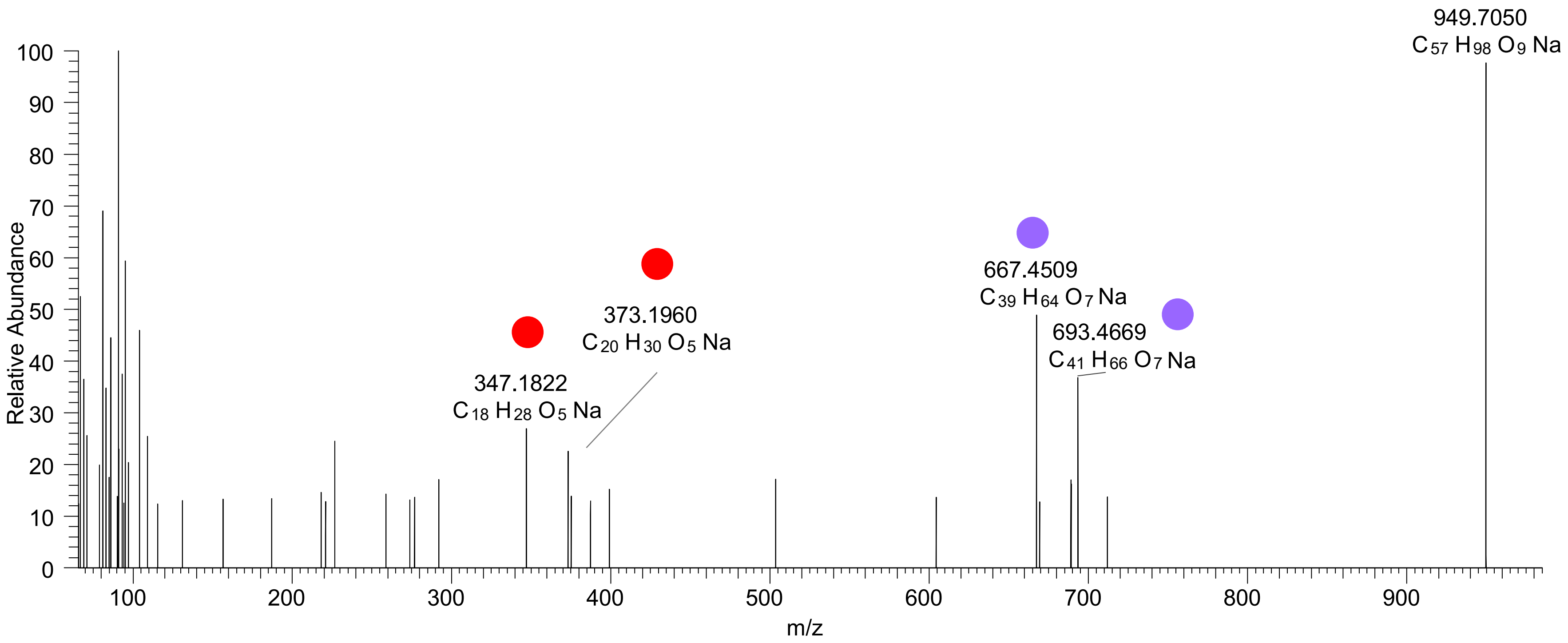

# TG(16:0_18:1_20:5<3O>)
# RT 21.8

**[oxTG+Na]+** **XIC 949.7103 NL: 5.53E4**

- Fragments containing oxFAs
- Fragments related to water loss
- Fragments not containing oxFAs
- Fragments related to other oxLPPs
- Position-specific fragments
- Fragments related to FA loss
- Fragments related to oxFA loss

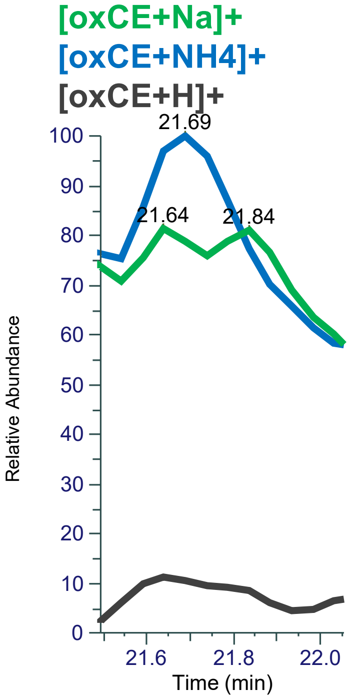


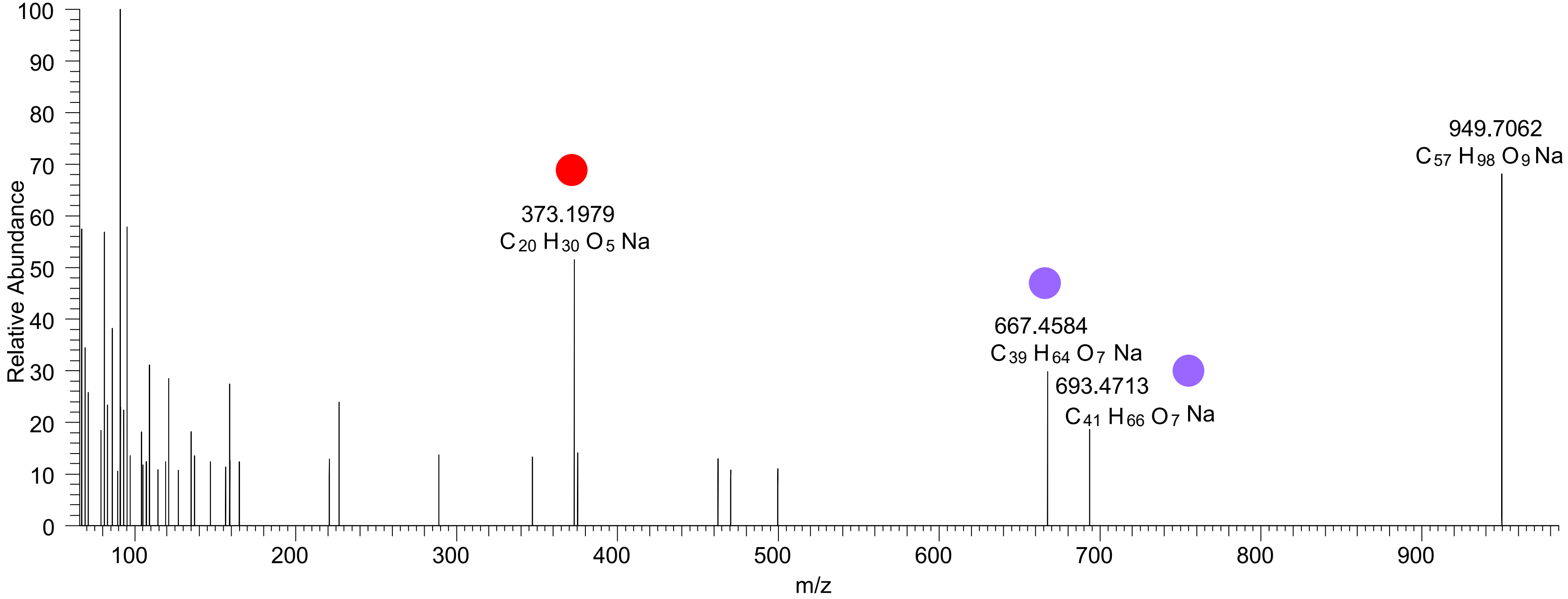

### TG(18:1_18:2<O>_18:2<OOH{13}>)
# RT 20.1

**[oxTG+Na]+** **XIC 951.7259 NL: 3.90E5**

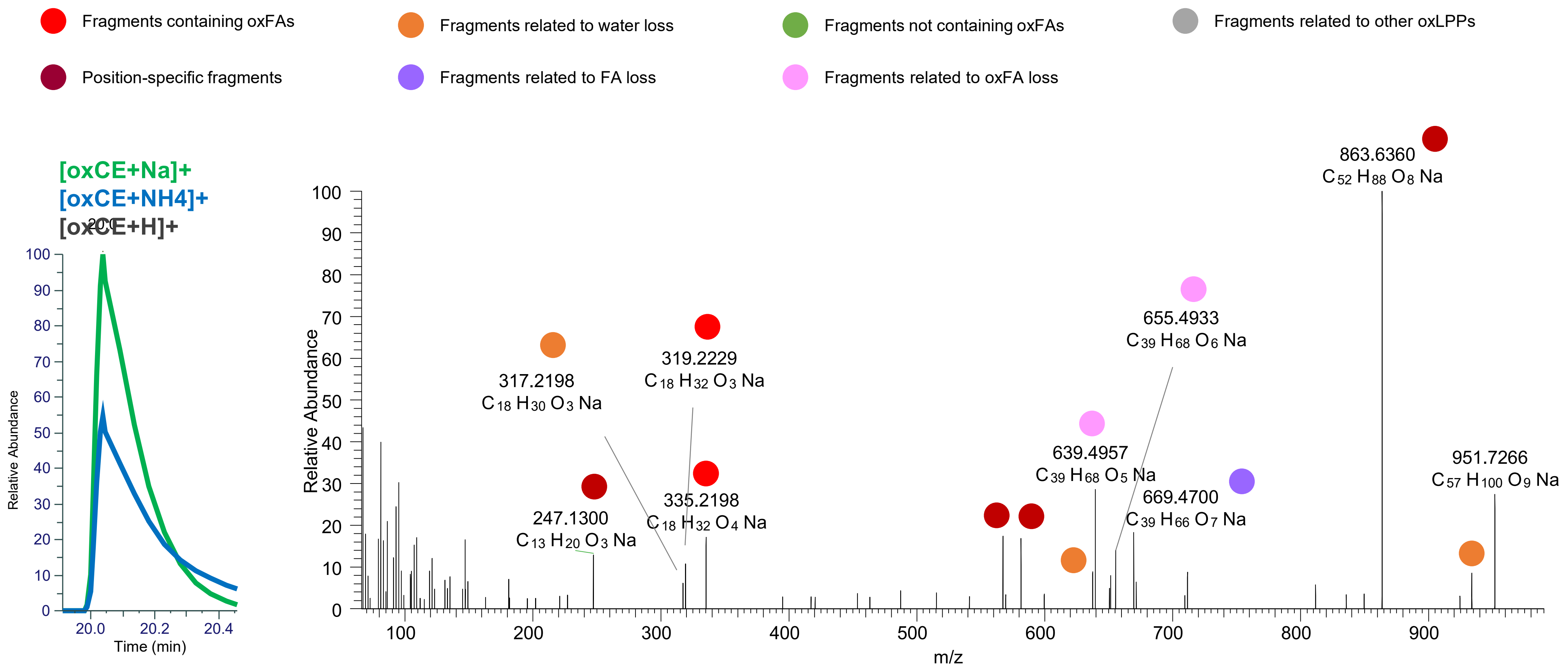

# TG(18:1_18:1_18:3<3O>)
# TG(16:0_18:1_20:4<3O>)
# RT 21.2

**[oxTG+Na]+** **XIC 951.7259 NL: 3.47E4**

- Fragments containing oxFAs
- Fragments related to water loss
- Fragments not containing oxFAs
- Fragments related to other oxLPPs
- Position-specific fragments
- Fragments related to FA loss
- Fragments related to oxFA loss

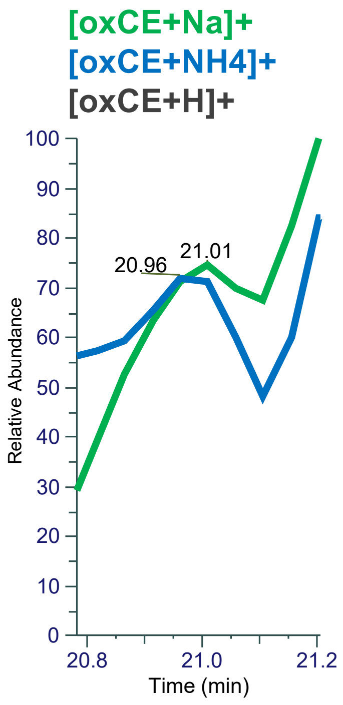


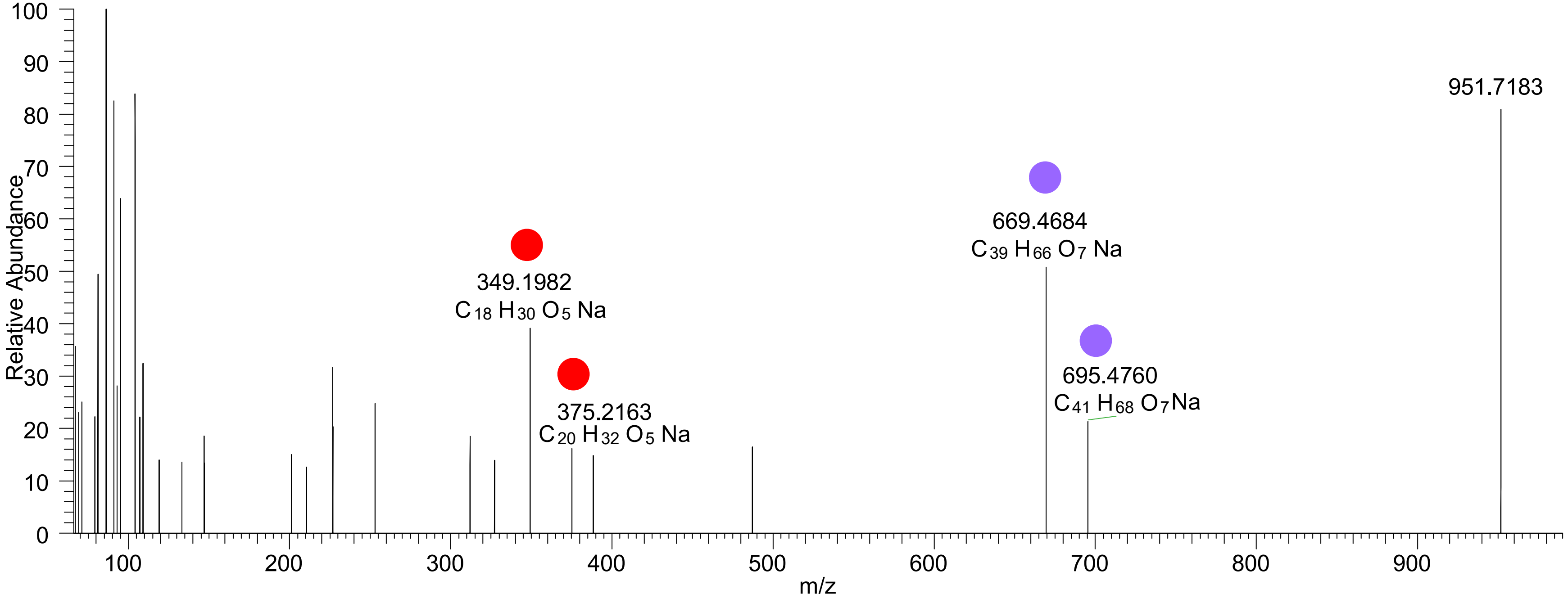

### TG(16:0_18:1_20:3<ep,OOH{15}>)
RT 22.1

**[oxTG+Na]+** **XIC 951.7259 NL: 9.20E4**

Fragments containing oxFAs
Fragments related to water loss
Fragments not containing oxFAs
Fragments related to other oxLPPs
Position-specific fragments
Fragments related to FA loss
Fragments related to oxFA loss

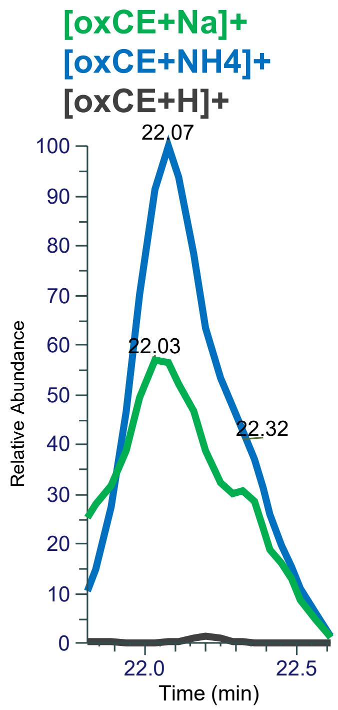


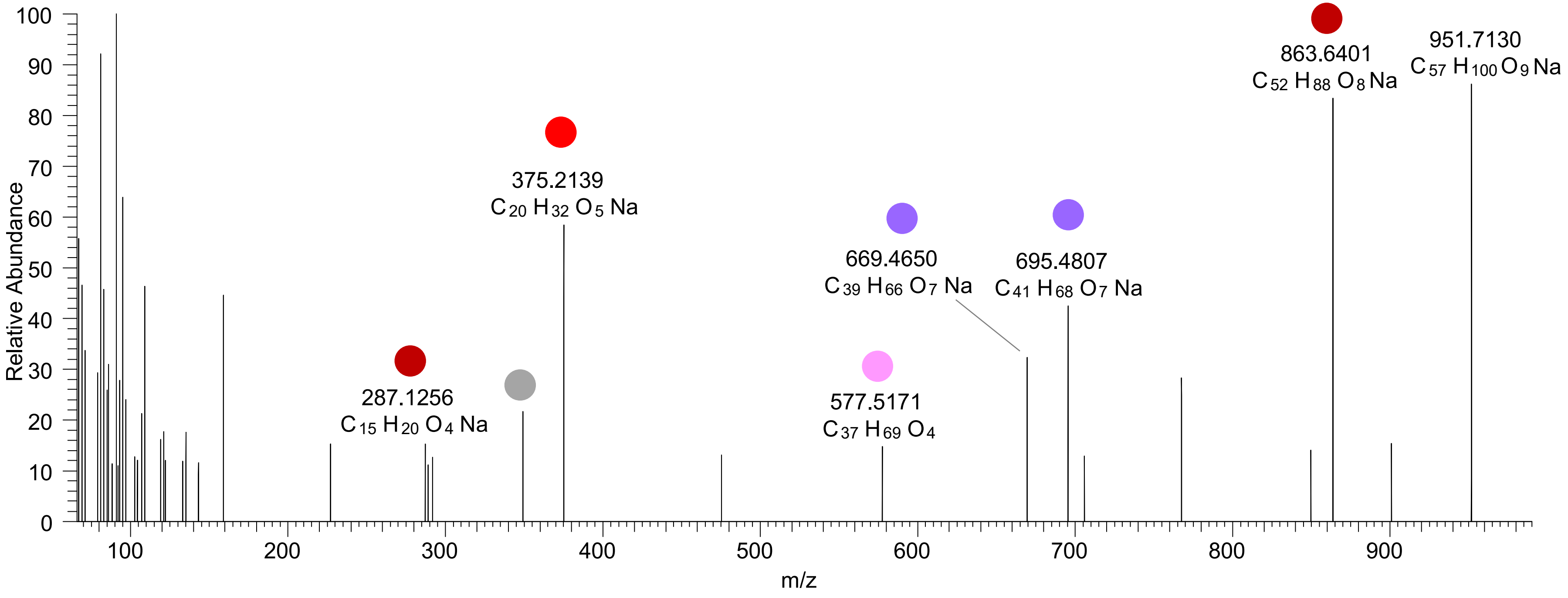

### OT2D_pool

# TG(16:1_18:1_16:0<OH>)
# RT 23.4

**[oxTG+Na]+** **XIC 869.7205 NL: 2.52E5**

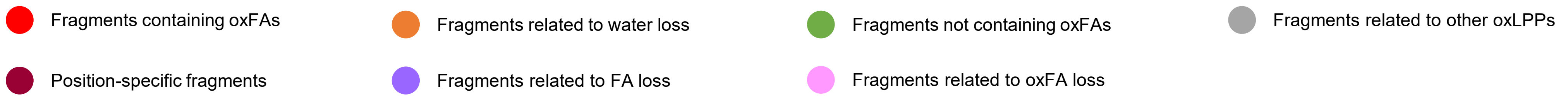


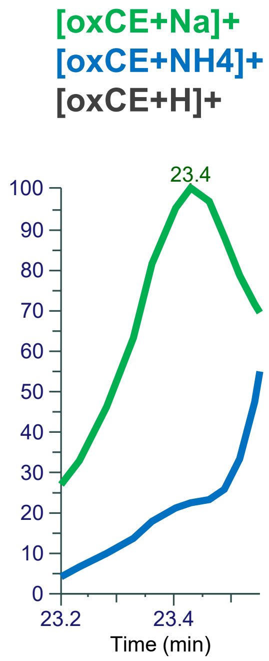


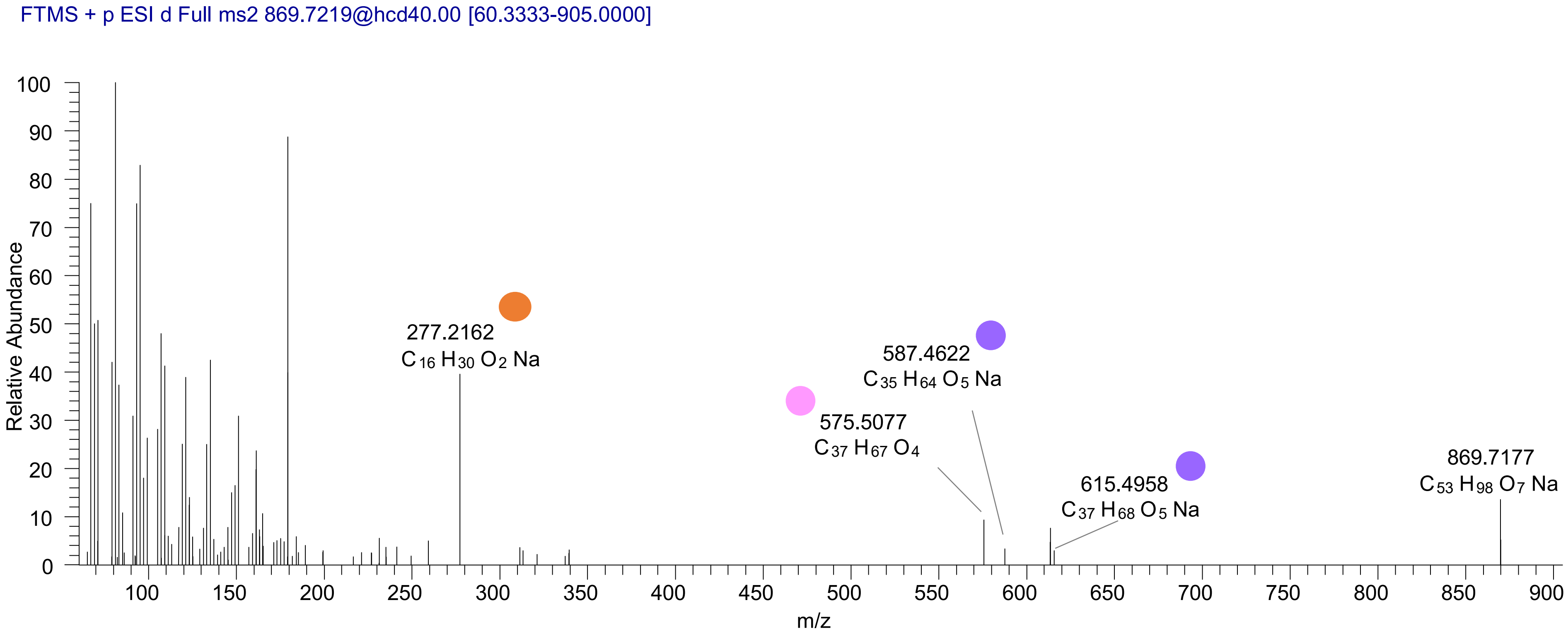

# TG(16:0_16:0_18:2<OH>)
# RT 23.7

[oxTG+Na]+ XIC 869.7205 NL: 2.70E5

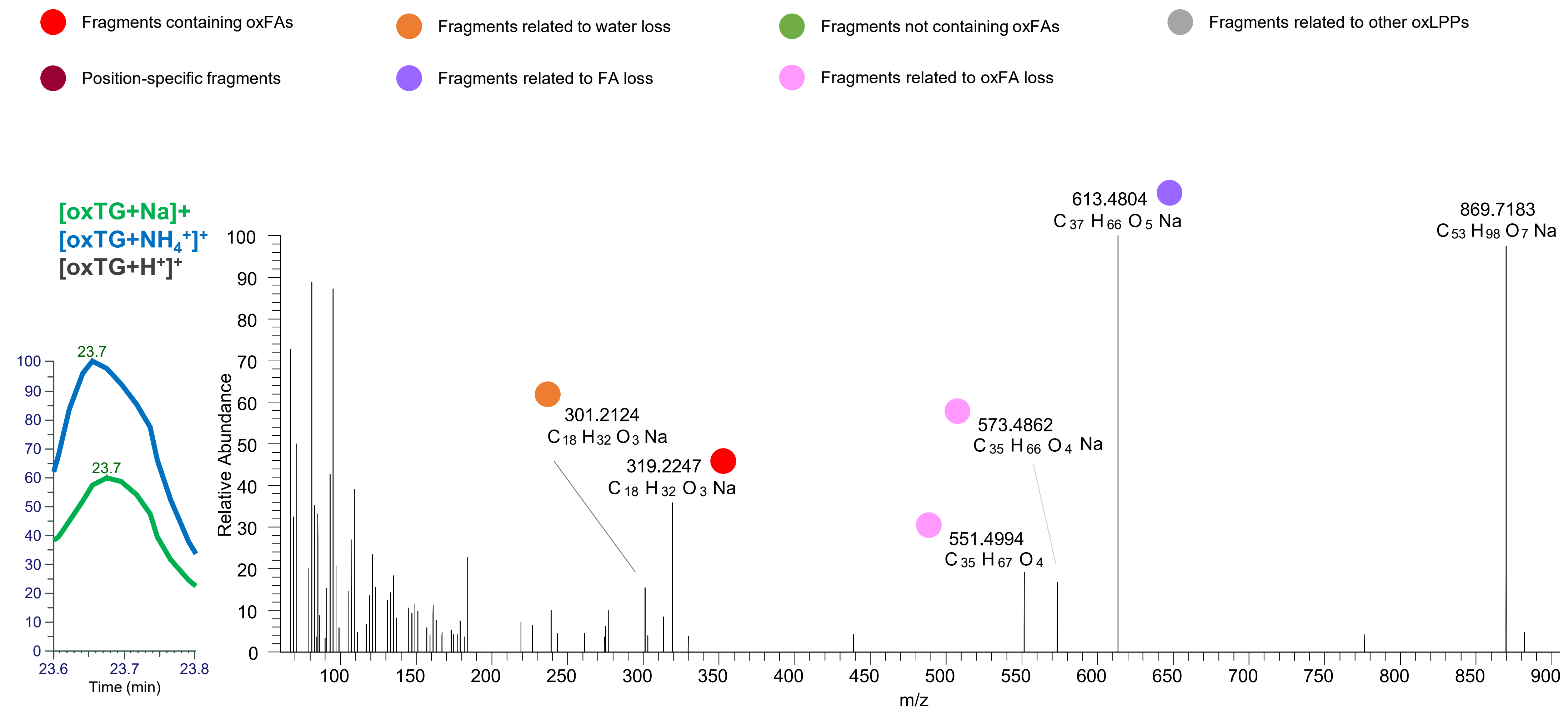

# TG(16:0_16:0_18:1<ep>)
# RT 24.3

**[oxTG+Na]+** **XIC 869.7205 NL: 3.51E4**

- Fragments containing oxFAs
- Fragments related to water loss
- Fragments not containing oxFAs
- Fragments related to other oxLPPs
- Position-specific fragments
- Fragments related to FA loss
- Fragments related to oxFA loss

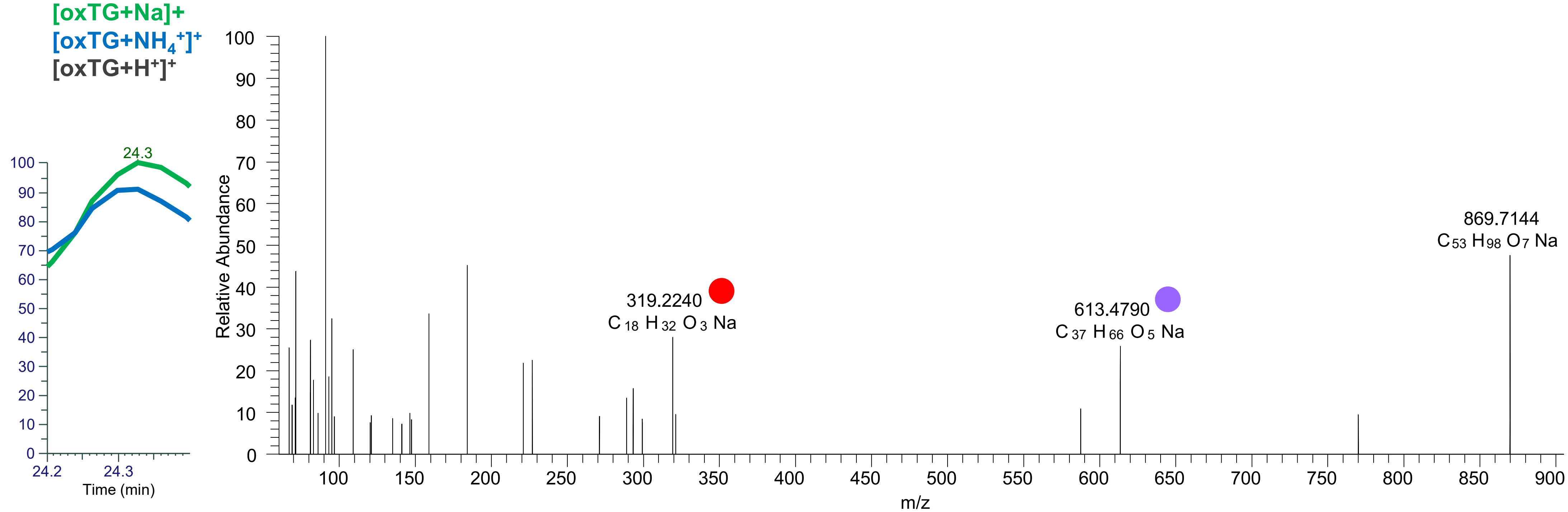

# TG(16:0_18:2_18:2<OH>)
# TG(16:1_18:1_18:2<OH>)
# RT 23.2

[oxTG+Na]+ XIC 893.7204 NL: 2.24E6

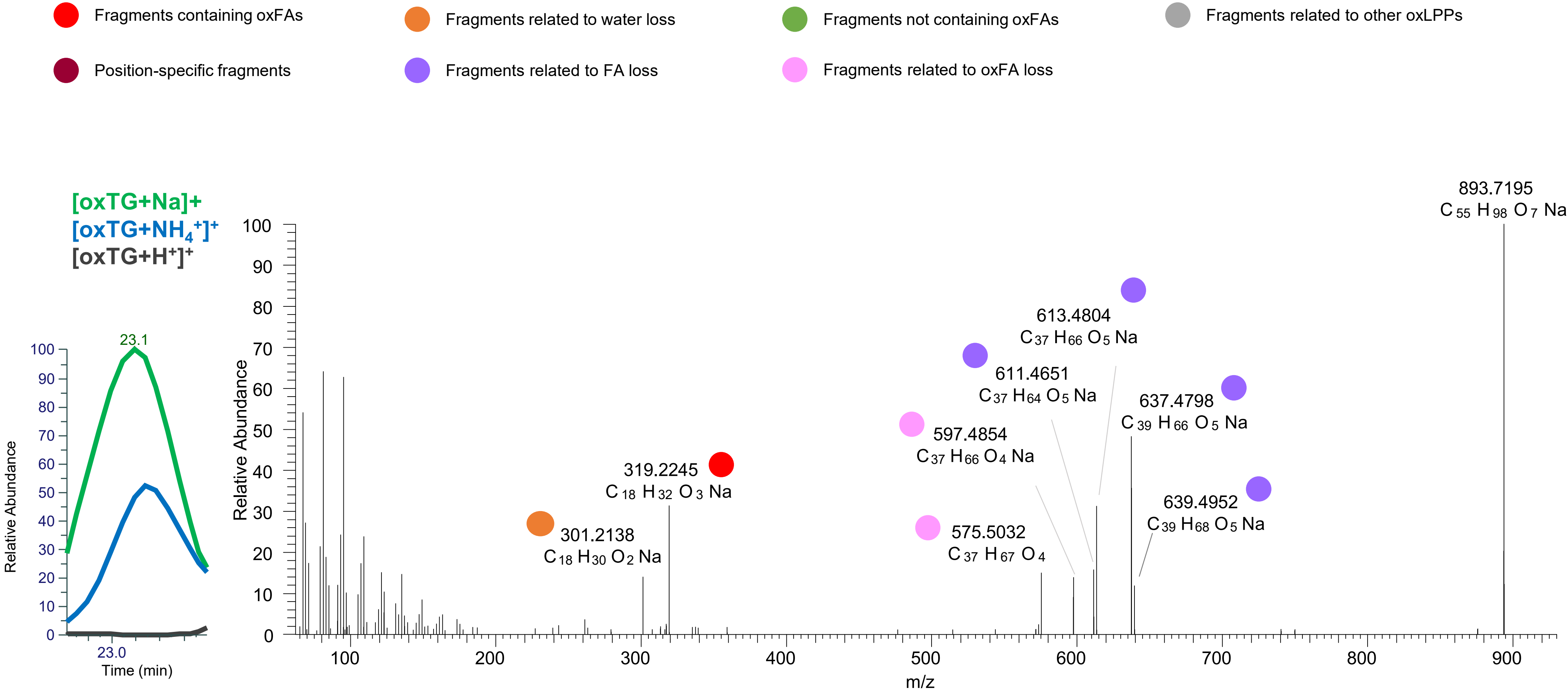

# TG(16:1_18:1_18:2<OH>)
# TG(16:0_18:1_18:3<OH>)
# RT 23.3

**[oxTG+Na]+** **XIC 893.7204 NL: 2.24E6**

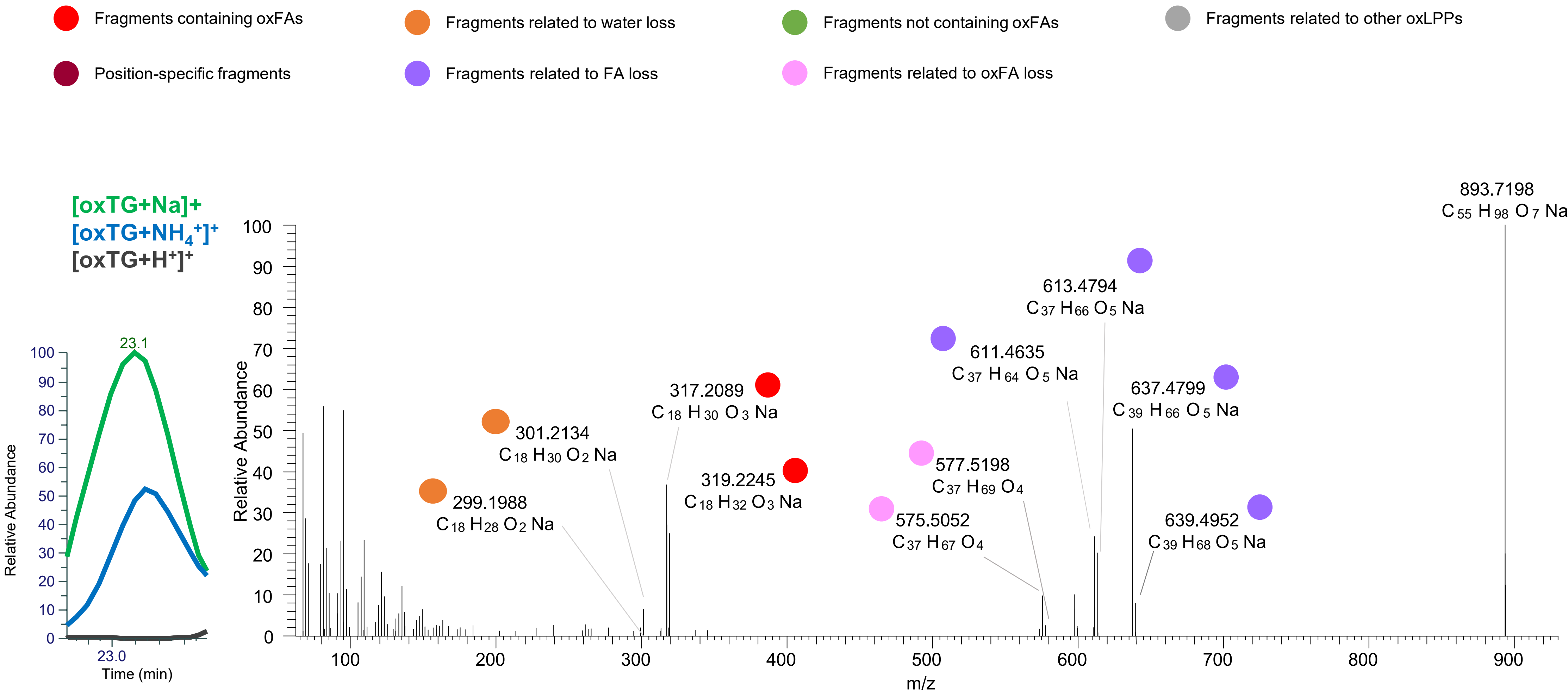

### TG(16:0_18:1_18:2<oxo>)
# RT 24.0

**[oxTG+Na]+** **XIC 893.7204 NL: 5.85E5**

- Fragments containing oxFAs
- Fragments related to water loss
- Fragments not containing oxFAs
- Fragments related to other oxLPPs
- Position-specific fragments
- Fragments related to FA loss
- Fragments related to oxFA loss

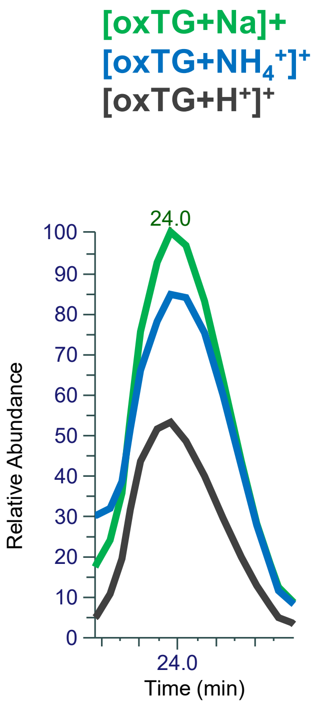


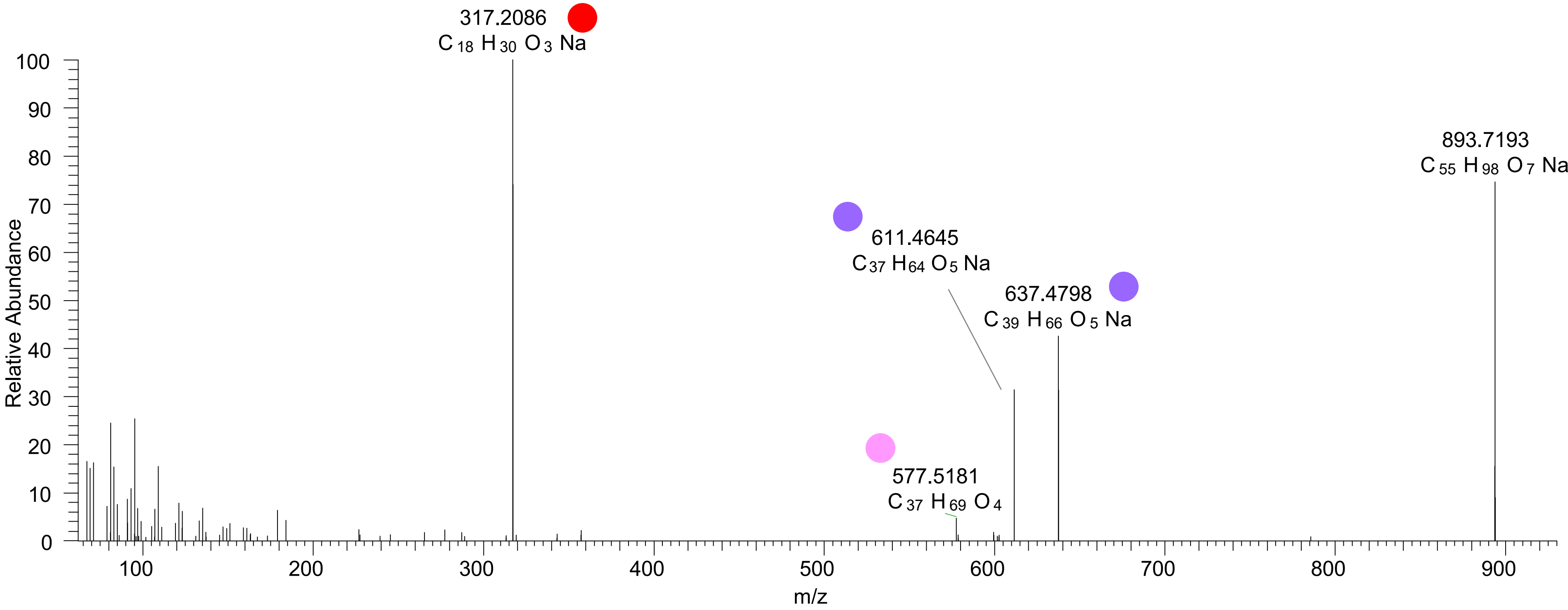

# TG(16:0_18:1_18:3<O>)
# RT 24.6

**[oxTG+Na]+** **XIC 893.7204 NL: 7.14E4**

- Fragments containing oxFAs
- Fragments related to water loss
- Fragments not containing oxFAs
- Fragments related to other oxLPPs
- Position-specific fragments
- Fragments related to FA loss
- Fragments related to oxFA loss

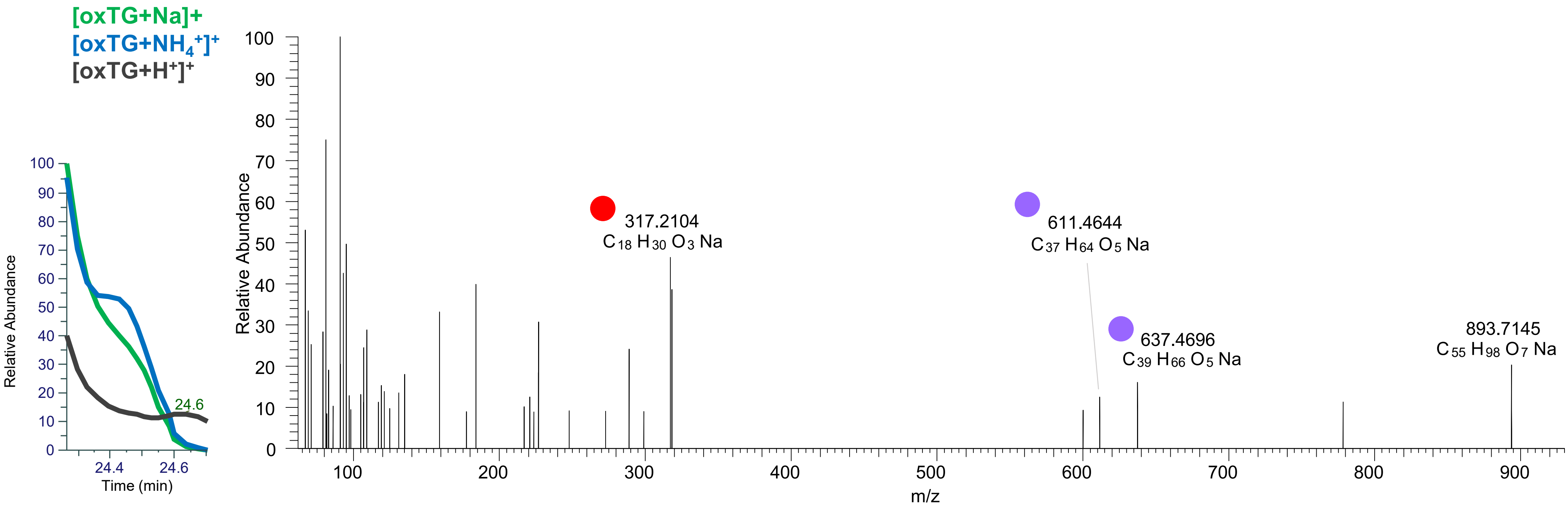

# TG(16:0_18:1_18:2<ep>)
# RT 25.4

**[oxTG+Na]+** **XIC 893.7204 NL: 3.85E5**

- Fragments containing oxFAs
- Fragments related to water loss
- Fragments not containing oxFAs
- Fragments related to other oxLPPs
- Position-specific fragments
- Fragments related to FA loss
- Fragments related to oxFA loss

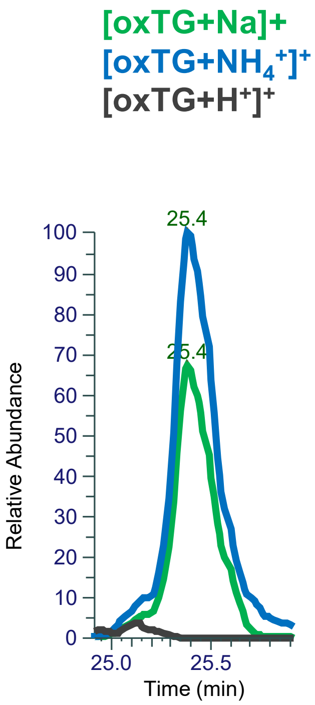


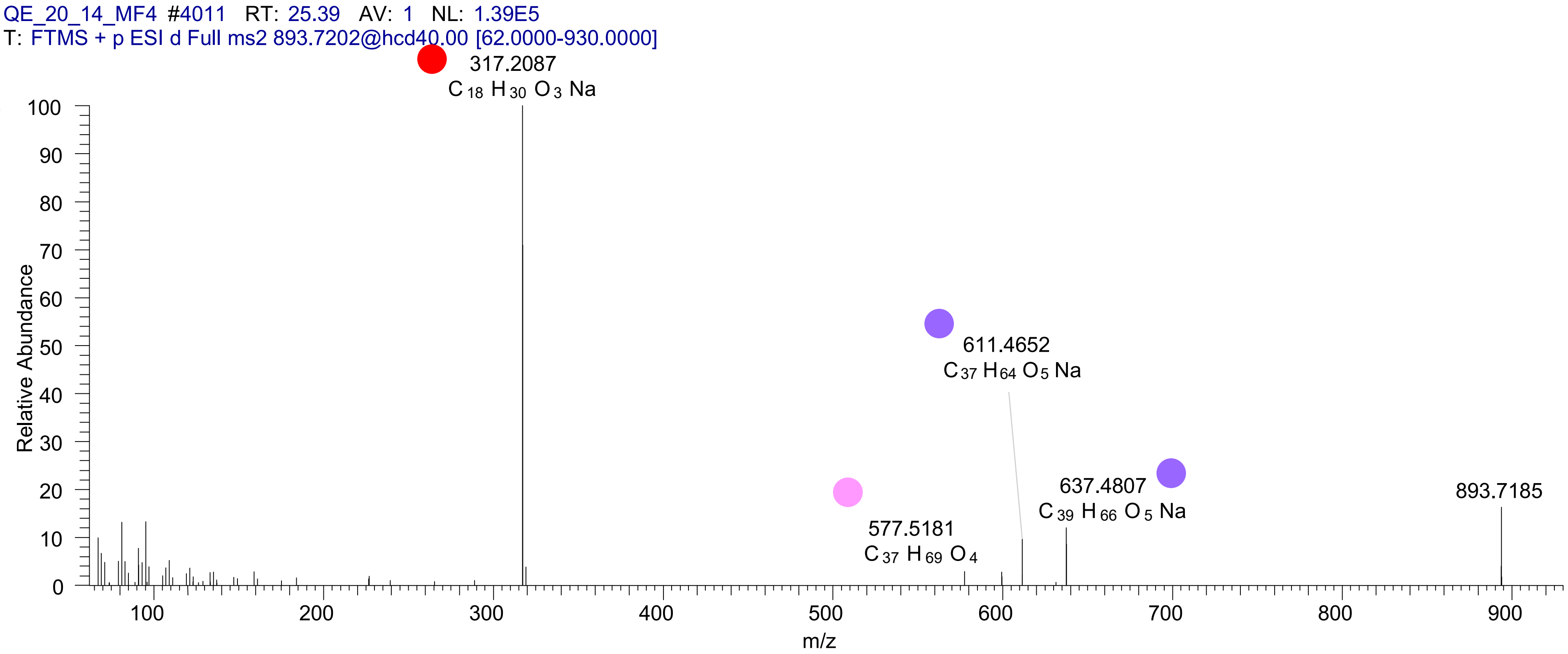

# TG(16:0_18:1_18:2<OH>)
# RT 23.5

**[oxTG+Na]+** **XIC 895.7361 NL: 6.83E5**

- Fragments containing oxFAs
- Fragments related to water loss
- Fragments not containing oxFAs
- Fragments related to other oxLPPs
- Position-specific fragments
- Fragments related to FA loss
- Fragments related to oxFA loss

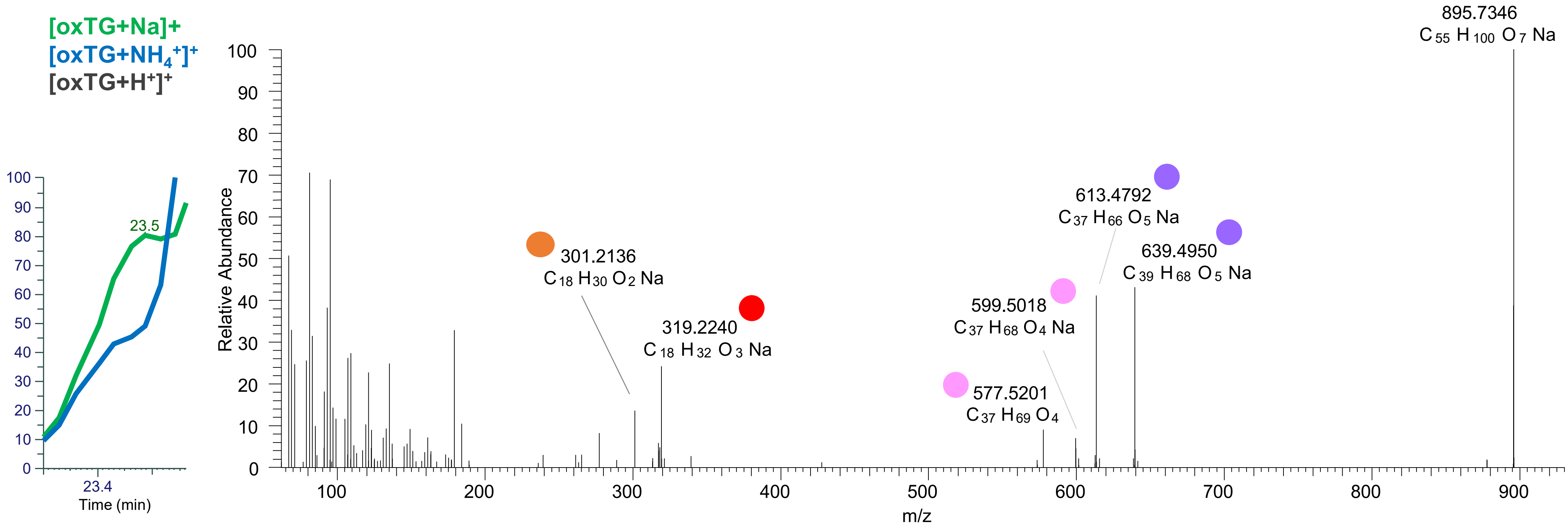

TG(16:0_18:1_18:2<O>)
TG(16:0_18:0_18:3<O>)
RT 24.7

**[oxTG+Na]+** **XIC 895.7361 NL: 8.14E4**

- Fragments containing oxFAs
- Fragments related to water loss
- Fragments not containing oxFAs
- Fragments related to other oxLPPs
- Position-specific fragments
- Fragments related to FA loss

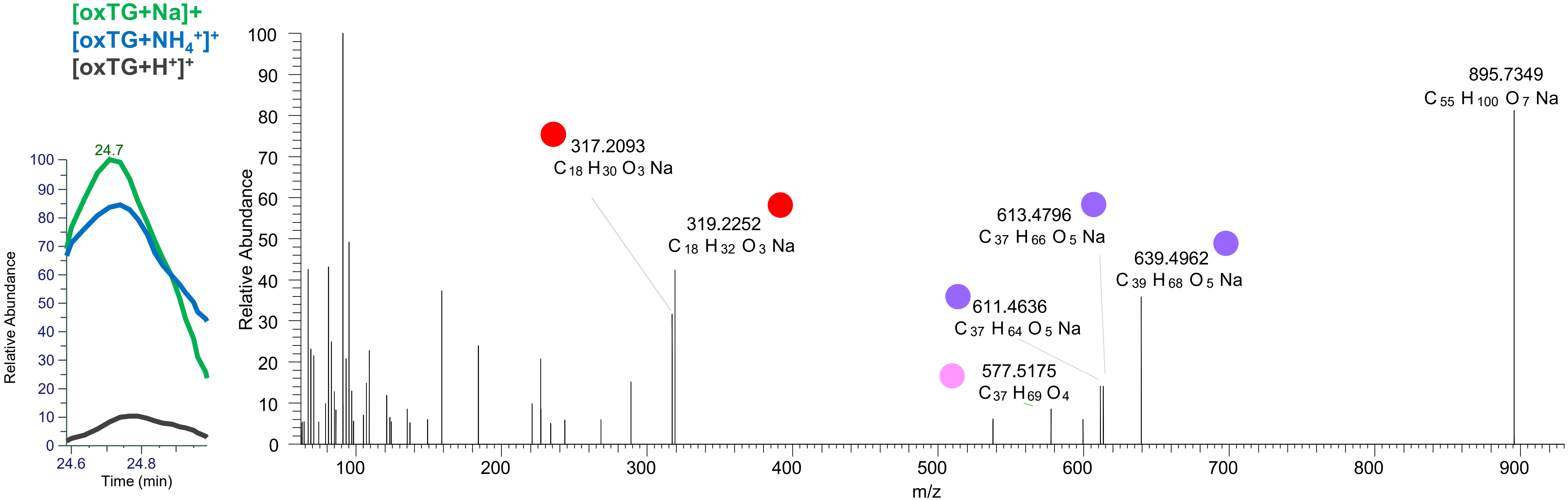

# TG(16:0_18:1_18:1<OH>)
# RT 24.1

**[oxTG+Na]+** **XIC 897.7517 NL: 3.43E5**

- Fragments containing oxFAs
- Fragments related to water loss
- Fragments not containing oxFAs
- Fragments related to other oxLPPs
- Position-specific fragments
- Fragments related to FA loss
- Fragments related to oxFA loss

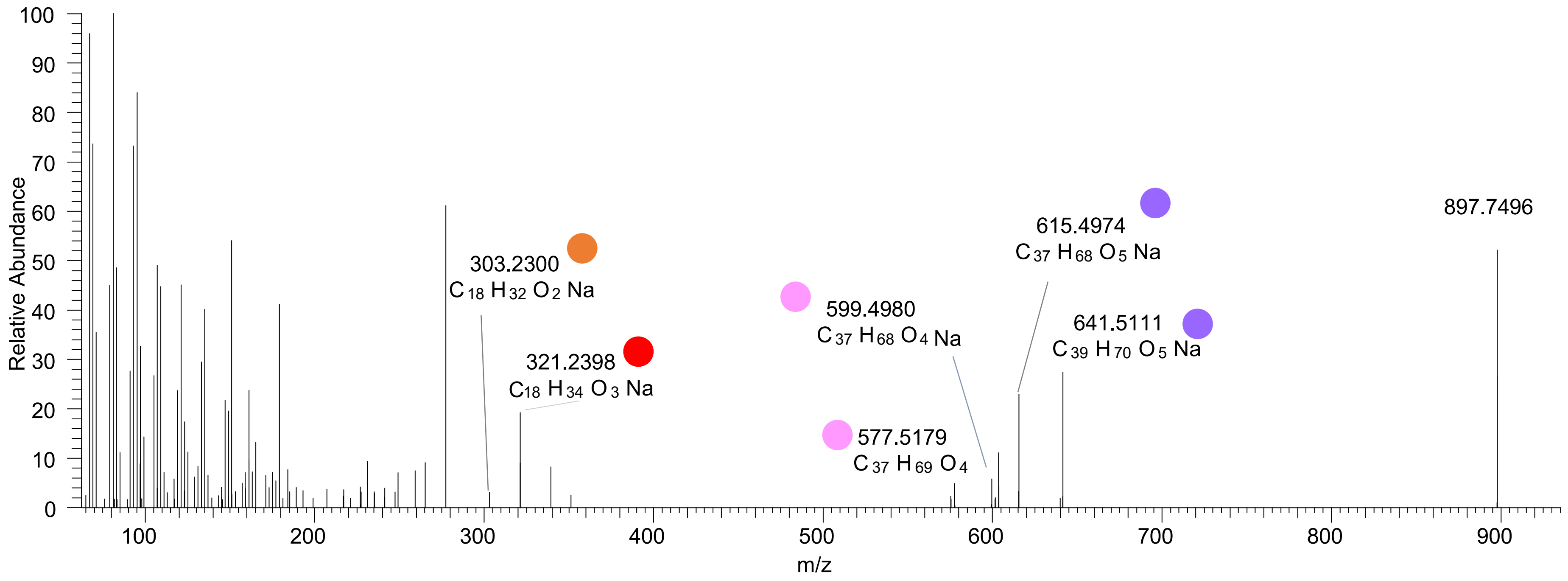

# TG(16:0_18:0_18:2<OH>)
# RT 24.3

**[oxTG+Na]$^{+}$**    **XIC 897.7517 NL: 3.43E5**

- Fragments containing oxFAs
- Fragments related to water loss
- Fragments not containing oxFAs
- Fragments related to other oxLPPs
- Position-specific fragments
- Fragments related to FA loss
- Fragments related to oxFA loss

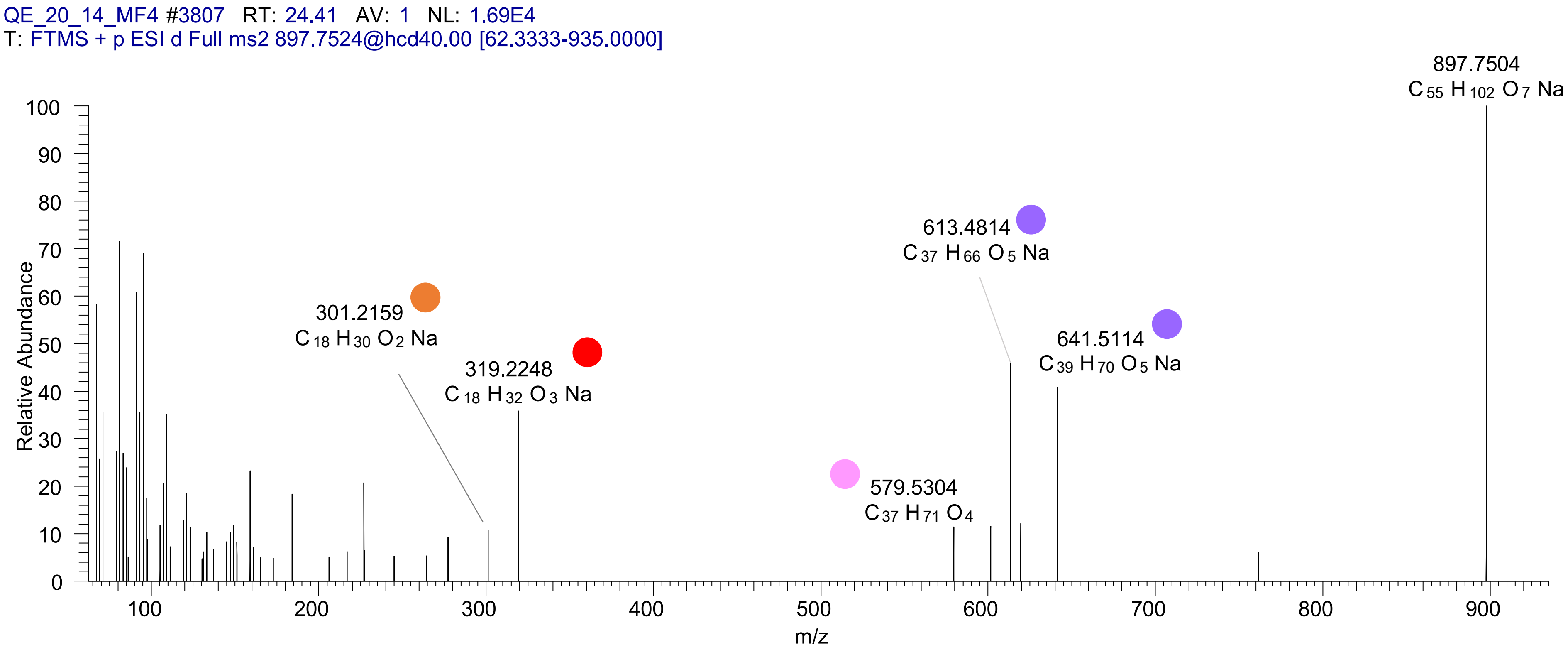

# TG(16:0_18:0_18:1<ep>)
# RT 24.6

**[oxTG+Na]$^+$** **XIC 897.7517 NL: 3.43E5**

- Fragments containing oxFAs
- Fragments related to water loss
- Fragments not containing oxFAs
- Fragments related to other oxLPPs
- Position-specific fragments
- Fragments related to FA loss
- Fragments related to oxFA loss

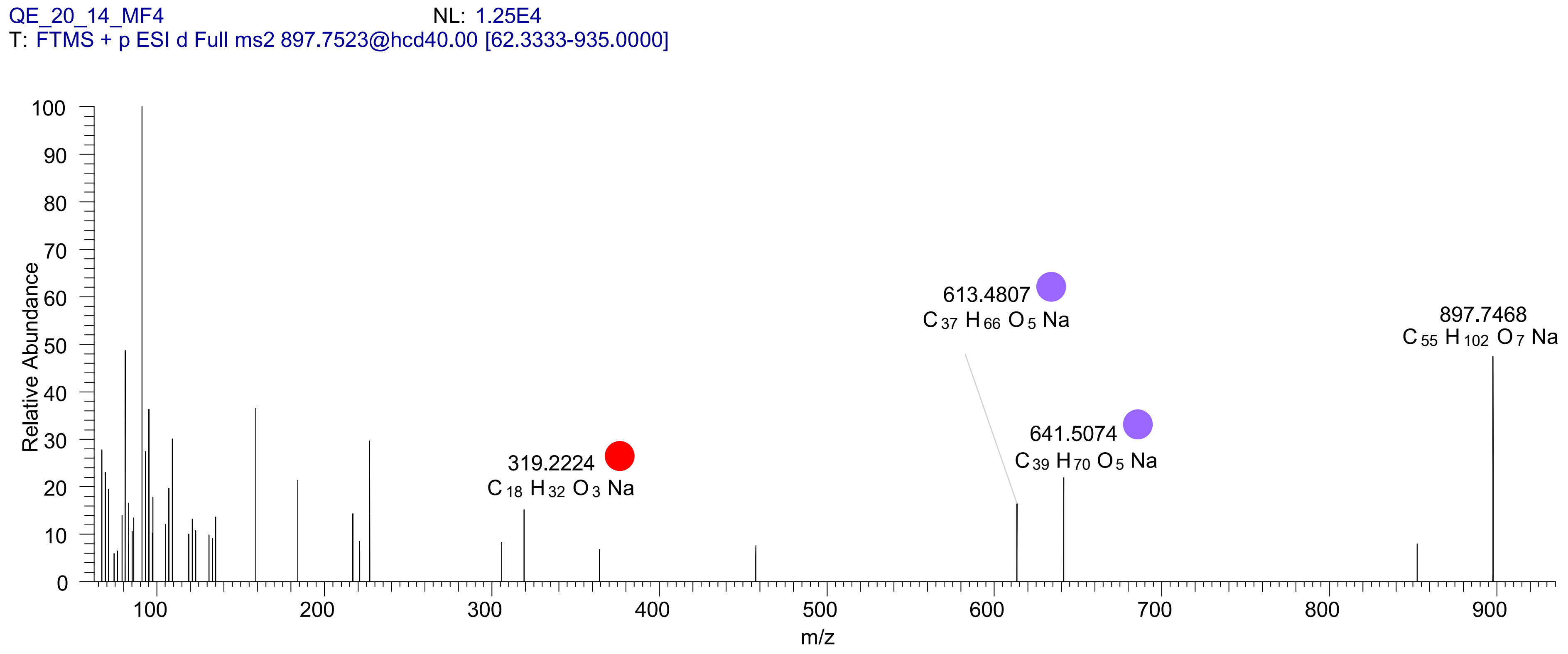

# TG(16:0_18:3_18:1<2OH>)
# RT 20.1

**[oxTG+Na]+** **XIC 909.7154 NL: 2.99E5**

- Fragments containing oxFAs
- Fragments related to water loss
- Fragments not containing oxFAs
- Fragments related to other oxLPPs
- Position-specific fragments
- Fragments related to FA loss
- Fragments related to oxFA loss

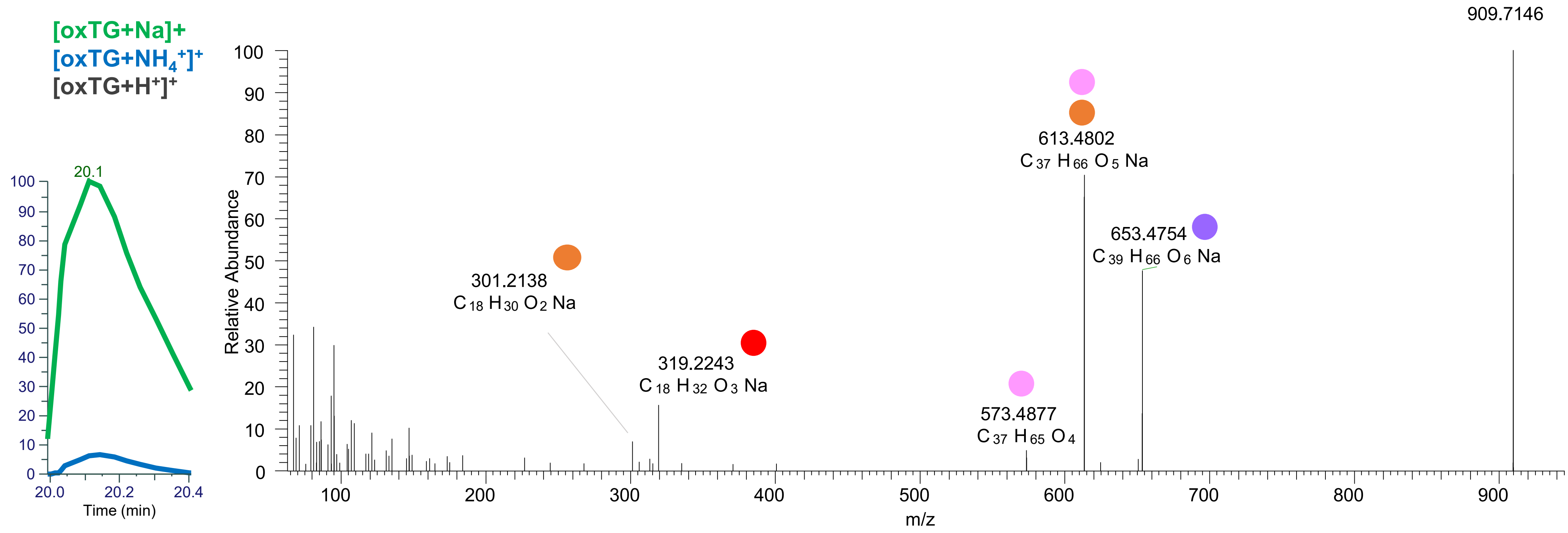

# TG(16:0_18:1_18:3<OH,O>)
# RT 21.6

**[oxTG+Na]+** **XIC 909.7154 NL: 8.98E4**

- Fragments containing oxFAs
- Fragments related to water loss
- Fragments not containing oxFAs
- Fragments related to other oxLPPs
- Position-specific fragments
- Fragments related to FA loss
- Fragments related to oxFA loss

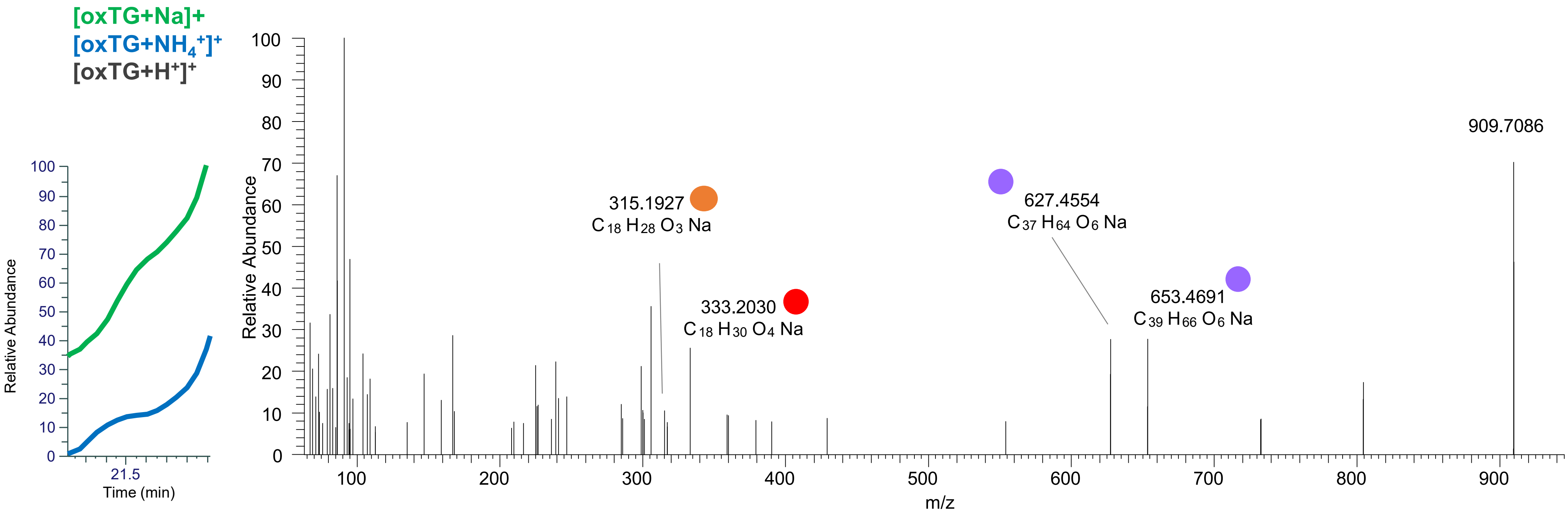

TG(16:0_18:1_18:3<2O>)
TG(16:1_18:1_18:2<2O>)
RT 21.8

[oxTG+Na]+ XIC 909.7154 NL: 8.98E4

Fragments containing oxFAs
Fragments related to water loss
Fragments not containing oxFAs
Fragments related to other oxLPPs
Position-specific fragments
Fragments related to FA loss
Fragments related to oxFA loss

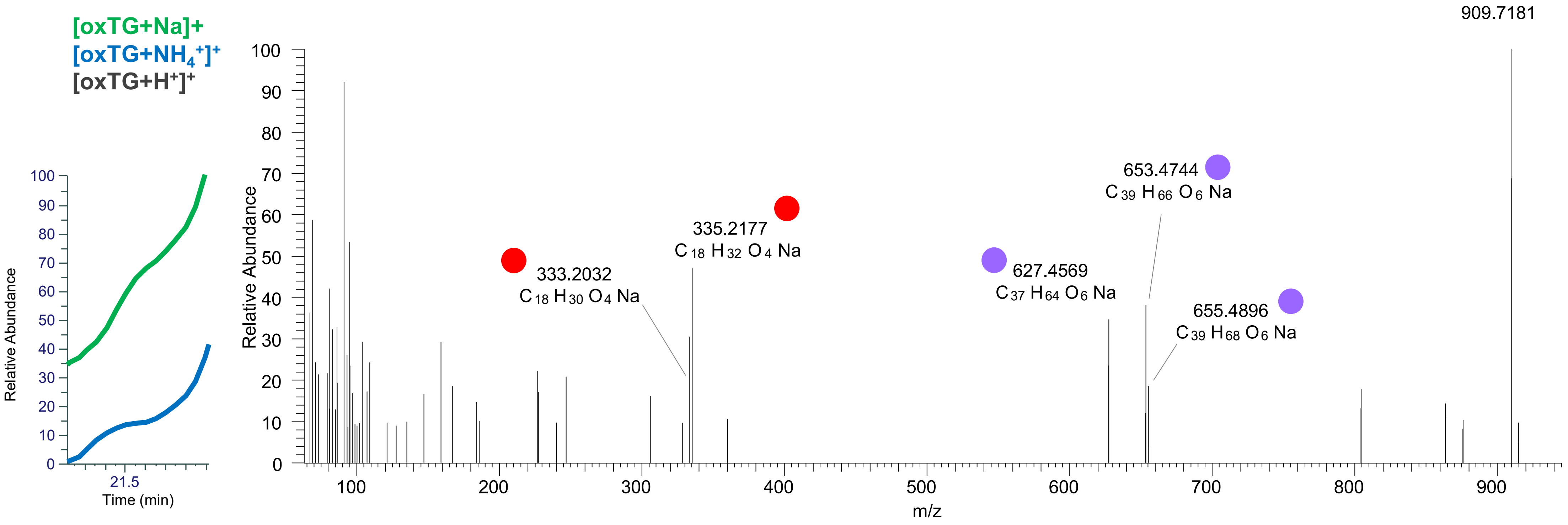

TG(16:1_18:1_18:2<2O>)
TG(16:0_18:1_18:3<2O>)
TG(16:0_18:2_18:2<2O>)
TG(16:0_16:0_20:4<2O>)
RT 22.1

**[oxTG+Na]+** **XIC 909.7154 NL: 1.18E5**

- Fragments containing oxFAs
- Fragments related to water loss
- Fragments not containing oxFAs
- Fragments related to other oxLPPs
- Position-specific fragments
- Fragments related to FA loss
- Fragments related to oxFA loss

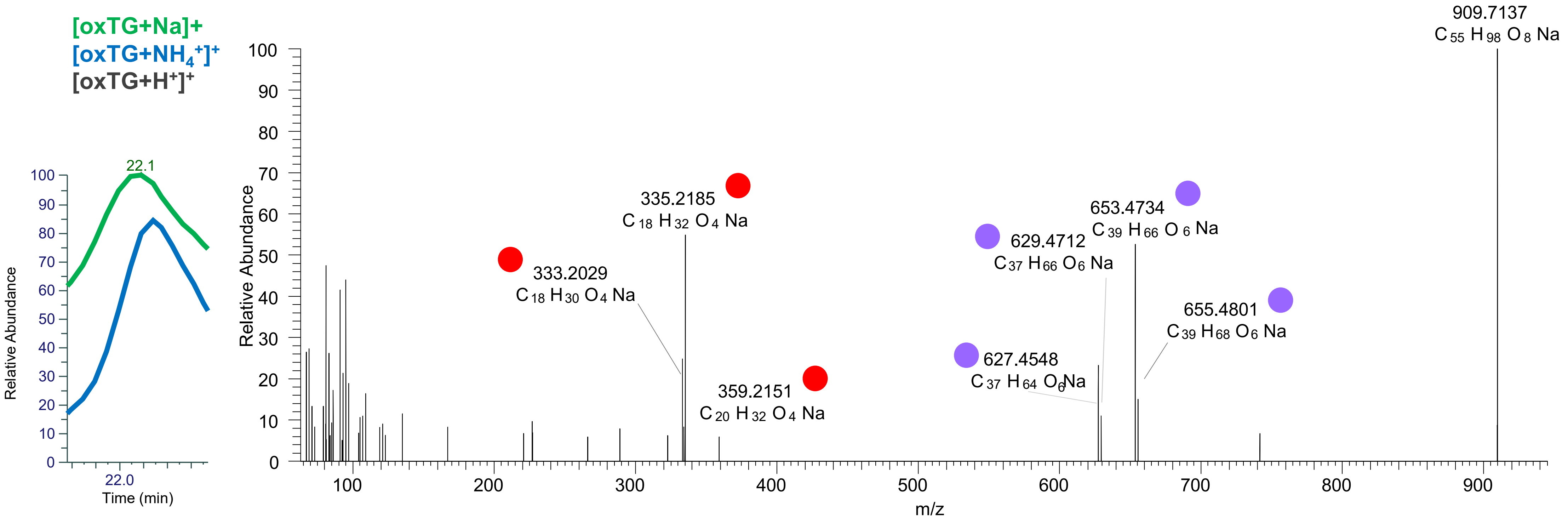

# TG(16:0_18:1_18:2<OH,ep>)
# TG(16:0_18:2_18:1<OH,ep>)
# RT 22.7

**[oxTG+Na]+** **XIC 909.7154 NL: 1.85E6**

- Fragments containing oxFAs
- Fragments related to water loss
- Fragments not containing oxFAs
- Fragments related to other oxLPPs
- Position-specific fragments
- Fragments related to FA loss
- Fragments related to oxFA loss

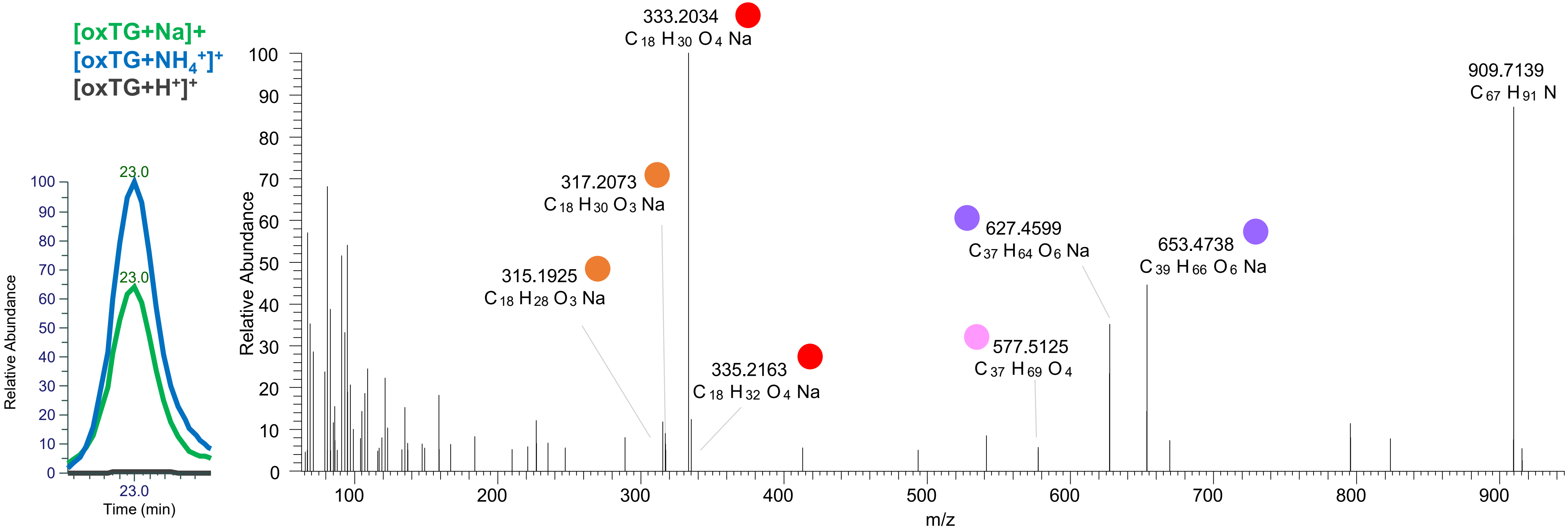

### TG(16:0_18:2_18:2<OOH{13}>)
### TG(16:0_18:1_18:3<OOH{13}>)
### TG(16:1_18:1_18:2<OOH{13}>)
# RT 23.0

**[oxTG+Na]+** **XIC 909.7154 NL: 1.85E6**

- Fragments containing oxFAs
- Fragments related to water loss
- Fragments not containing oxFAs
- Fragments related to other oxLPPs
- Position-specific fragments
- Fragments related to FA loss
- Fragments related to oxFA loss

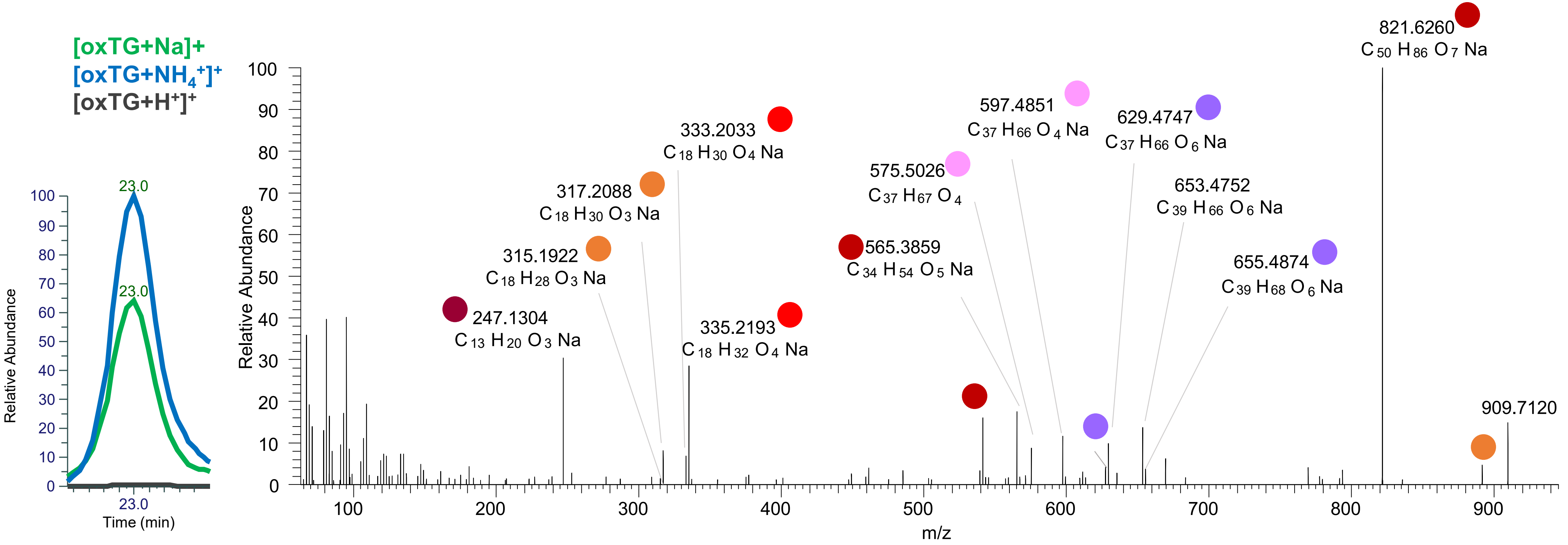

# TG(16:0_18:1_18:1<OH,ep>)
# RT 22.7

[oxTG+Na]+ XIC 911.7310 NL: 1.44E5

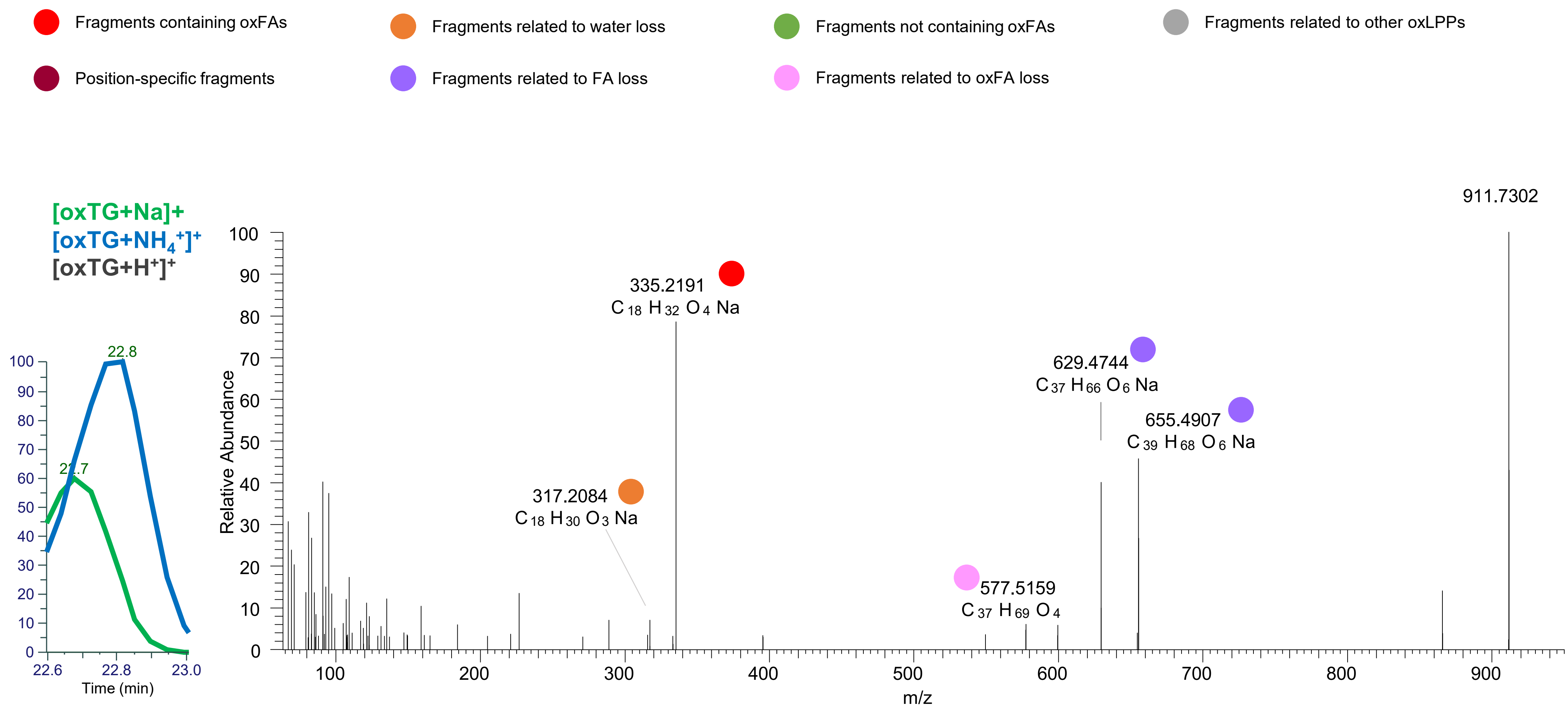

### TG(16:0_18:1_18:2<OOH{11}>)
### TG(16:0_18:1_18:2<OOH{13}>)
# RT 23.3

[oxTG+Na]+ XIC 911.7310 NL: 4.23E6

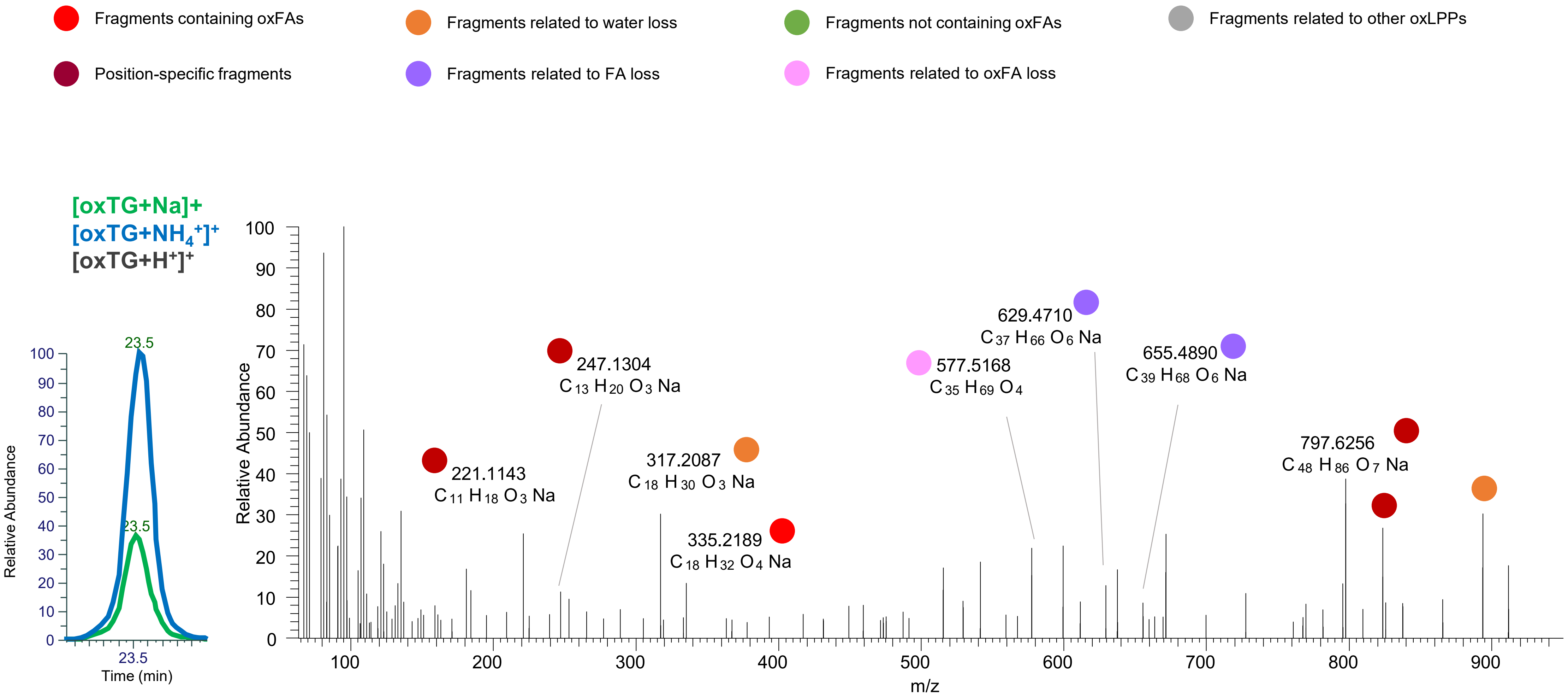

### TG(16:0_18:1_18:2<OOH{13}>)
# RT 23.5

**[oxTG+Na]+** **XIC 911.7310 NL: 4.23E6**

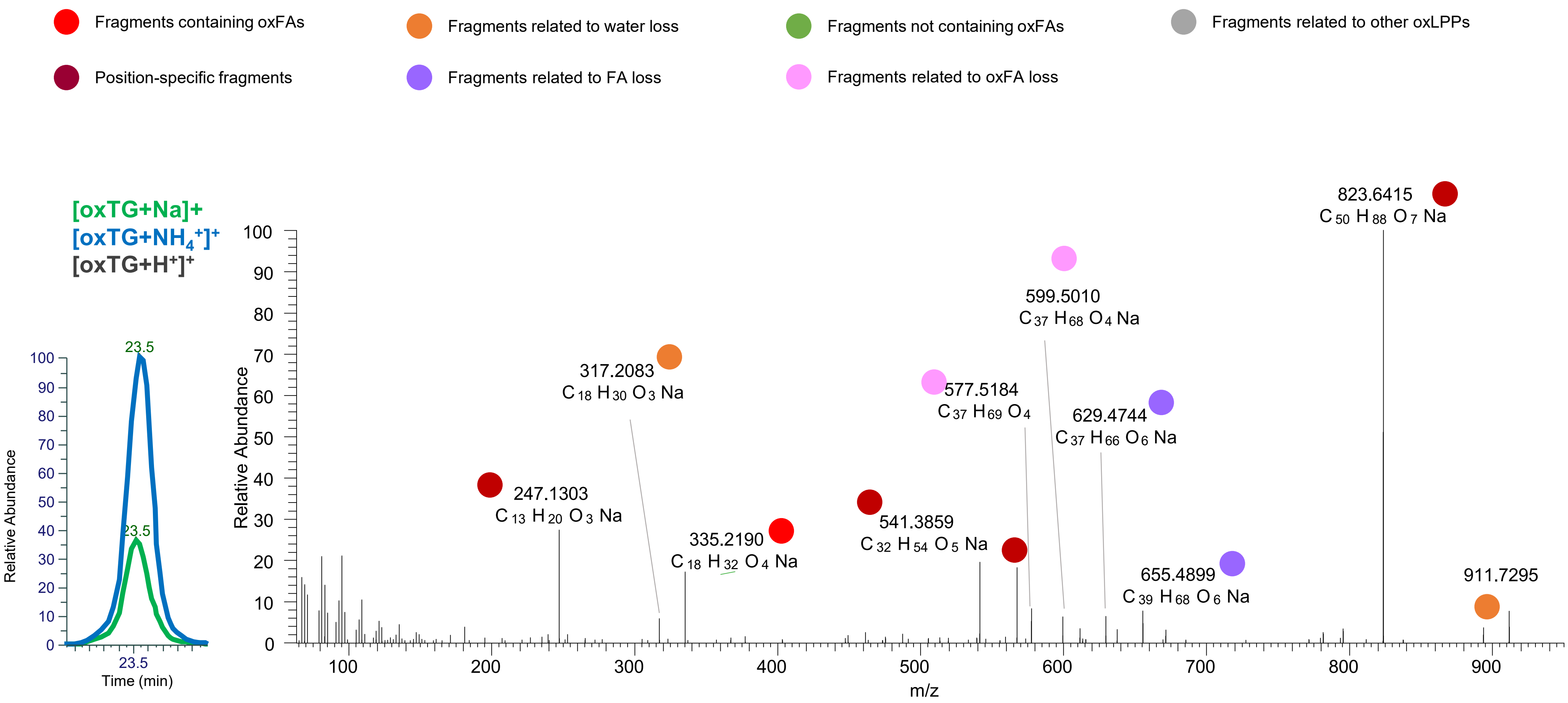

### TG(16:0_18:1_18:2<OOH{9}>)
### TG(16:0_18:1_18:2<OOH{13}>)
# RT 23.7

**[oxTG+Na]+** **XIC 911.7310 NL: 4.23E6**

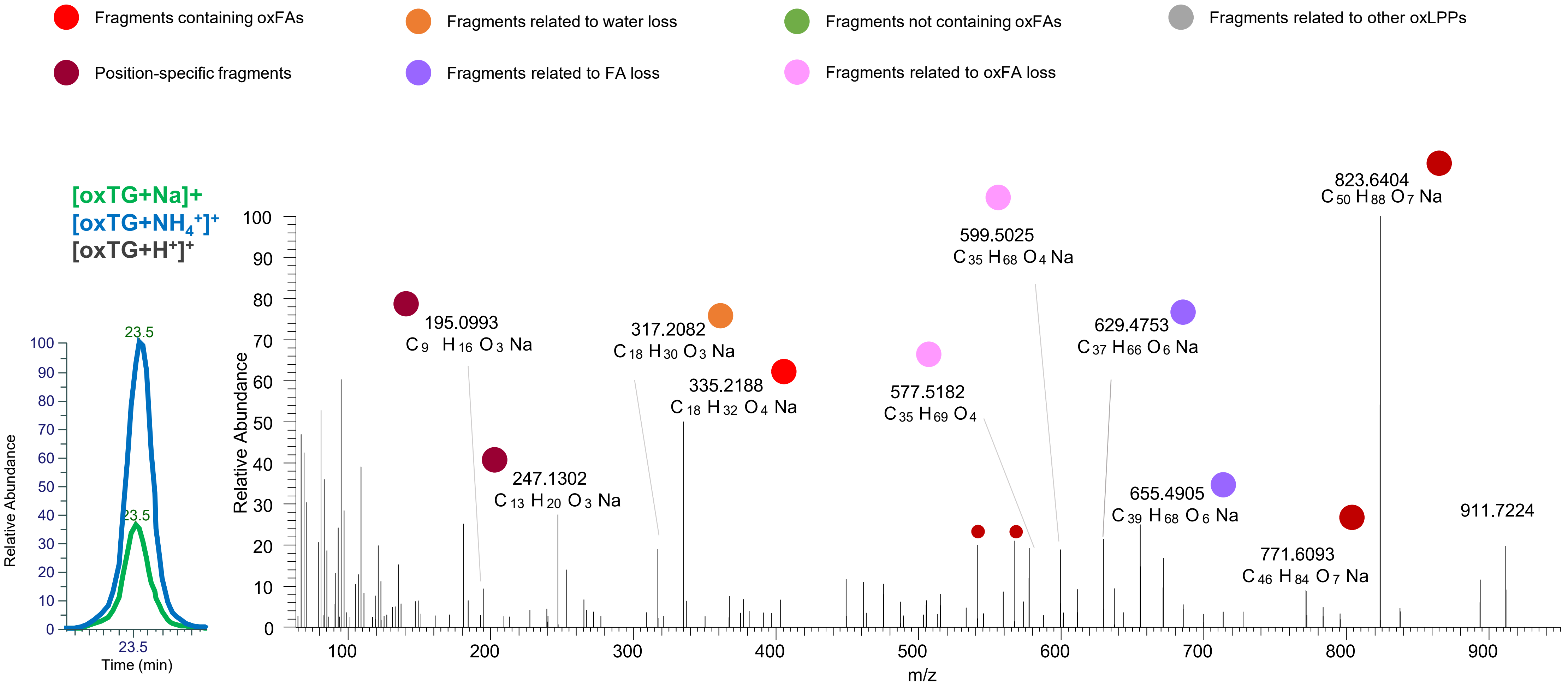

### TG(16:0_18:0_18:2<OOH{13}>)
# RT 24.3

**[oxTG+Na]+** **XIC 913.7467 NL: 1.32E5**

- Fragments containing oxFAs
- Fragments related to water loss
- Fragments not containing oxFAs
- Fragments related to other oxLPPs
- Position-specific fragments
- Fragments related to FA loss
- Fragments related to oxFA loss

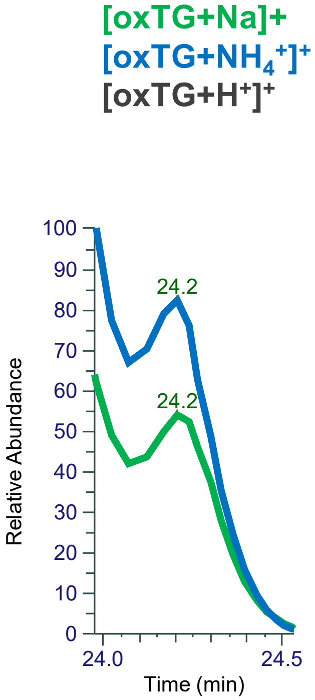


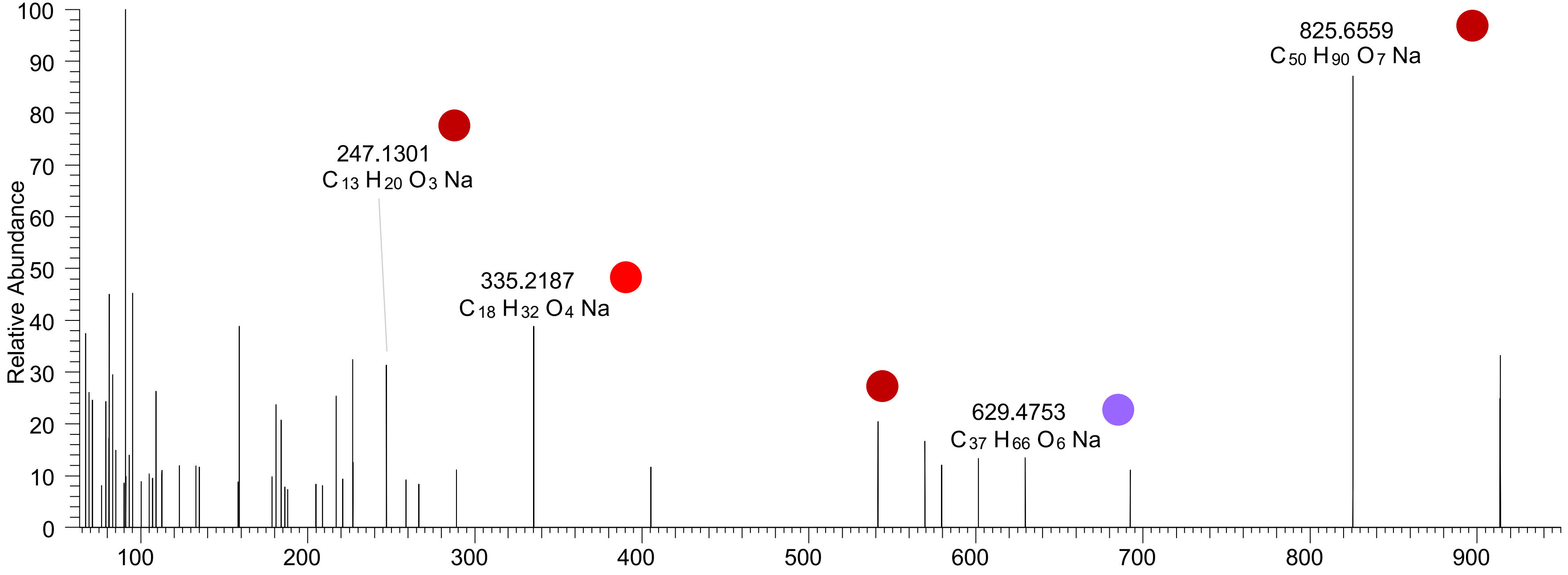

# TG(18:2_18:2_18:2<OH>)
# RT 22.6

**[oxTG+Na]+** **XIC 917.7204 NL: 3.82E5**

- Fragments containing oxFAs
- Fragments related to water loss
- Fragments not containing oxFAs
- Fragments related to other oxLPPs
- Position-specific fragments
- Fragments related to FA loss
- Fragments related to oxFA loss

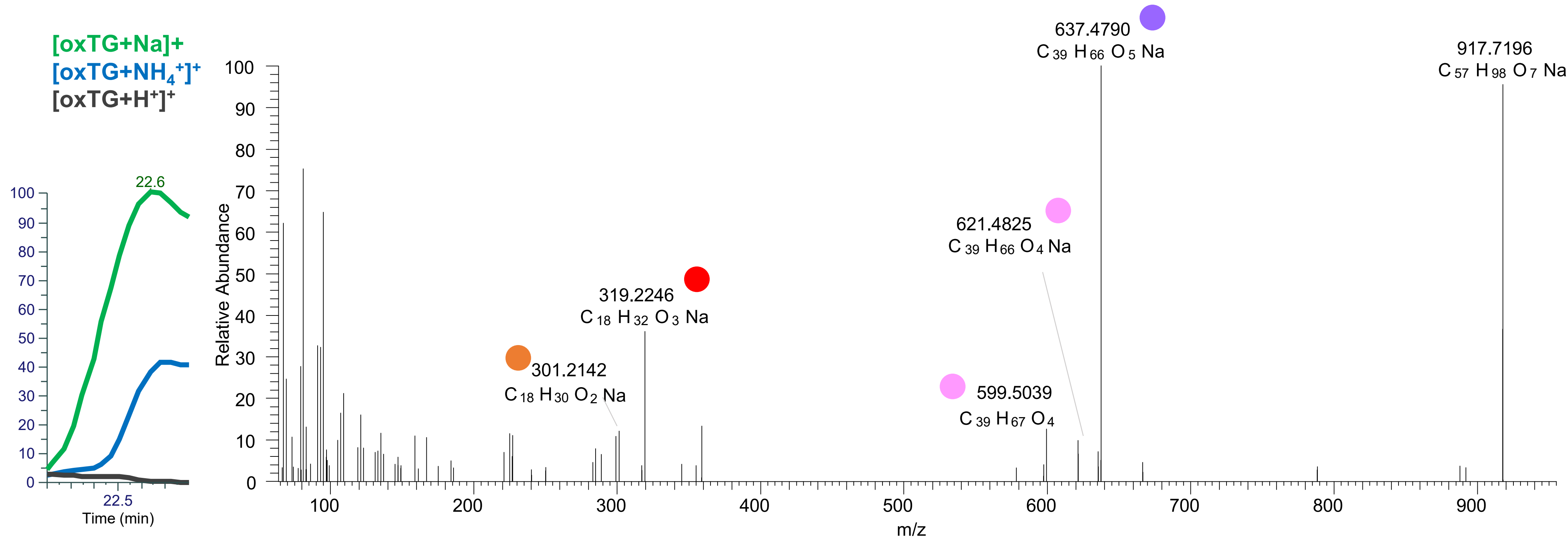

# TG(18:2_18:2_18:2<O>)
# TG(18:1_18:2_18:3<OH>)
# RT 23.0

**[oxTG+Na]+** **XIC 917.7204 NL: 3.53E5**

- Fragments containing oxFAs
- Fragments related to water loss
- Fragments not containing oxFAs
- Fragments related to other oxLPPs
- Position-specific fragments
- Fragments related to FA loss
- Fragments related to oxFA loss

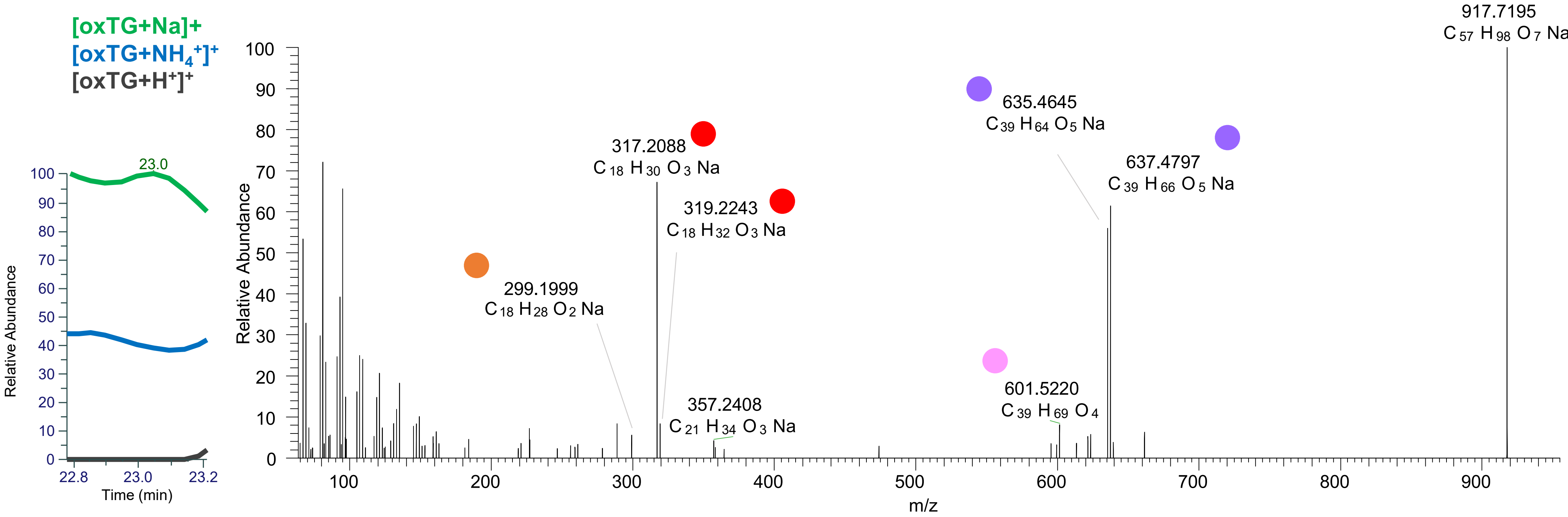

# TG(18:1_18:2_18:2<ep>)
### TG(18:1_18:2_18:2<oxo>)
# RT 24.7

**[oxTG+Na]+** **XIC 917.7204 NL: 1.08E5**

- Fragments containing oxFAs
- Fragments related to water loss
- Fragments not containing oxFAs
- Fragments related to other oxLPPs
- Position-specific fragments
- Fragments related to FA loss
- Fragments related to oxFA loss

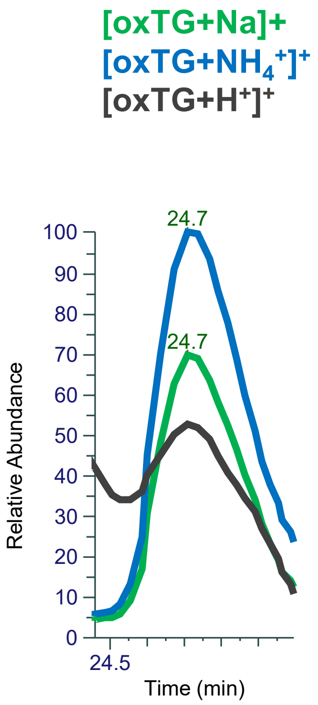


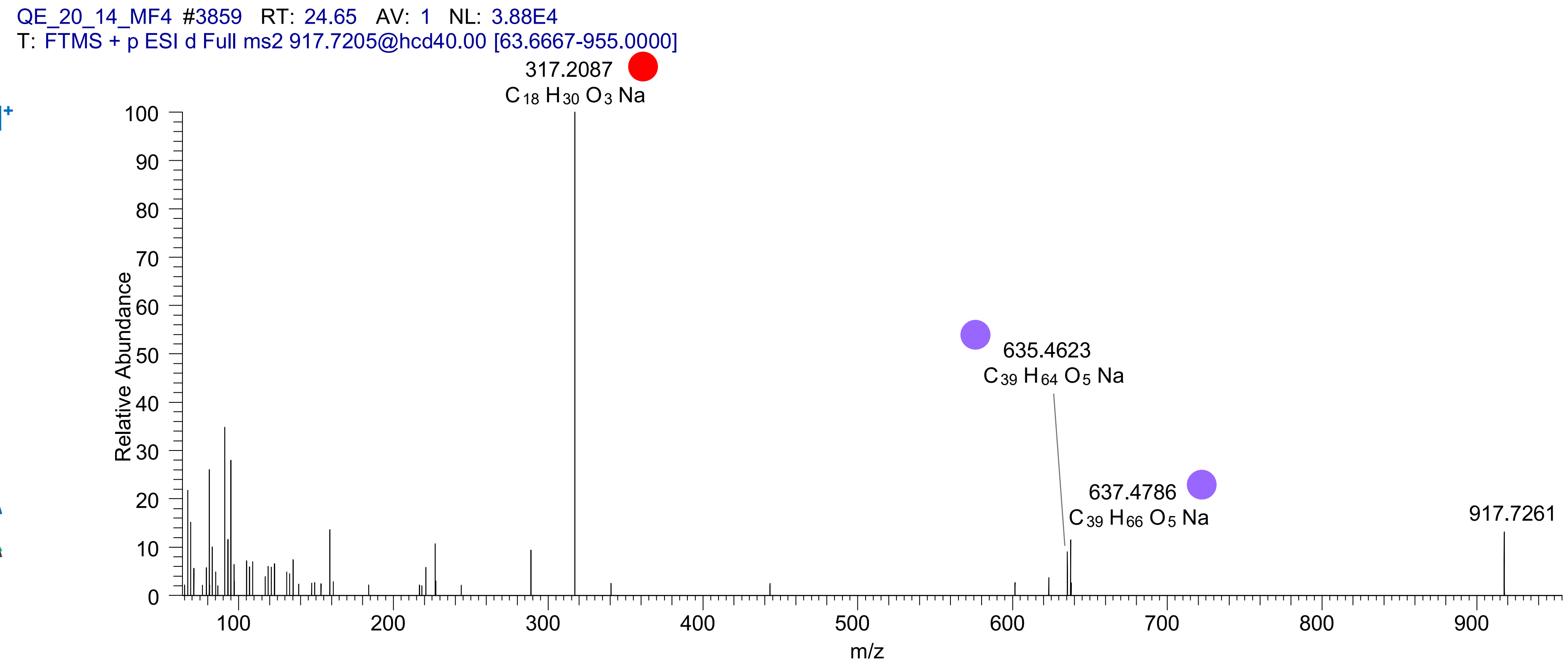

# TG(18:1_18:2_18:2<OH>)
# RT 23.2

**[oxTG+Na]+** **XIC 919.7361 NL: 1.09E6**

- Fragments containing oxFAs
- Fragments related to water loss
- Fragments not containing oxFAs
- Fragments related to other oxLPPs
- Position-specific fragments
- Fragments related to FA loss
- Fragments related to oxFA loss

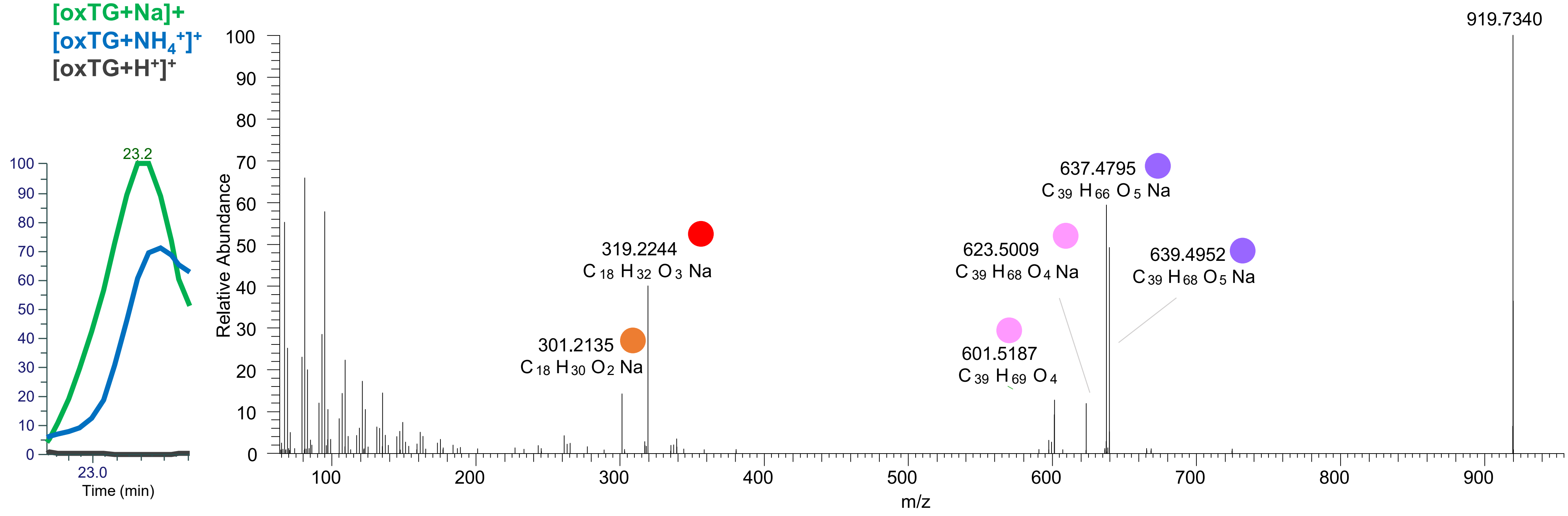

TG(18:1_18:2_18:2<O>)
TG(18:1_18:1_18:3<O>)
TG(16:0_18:2_20:3<O>)
RT 23.3

[oxTG+Na]+ XIC 919.7361 NL: 1.09E6

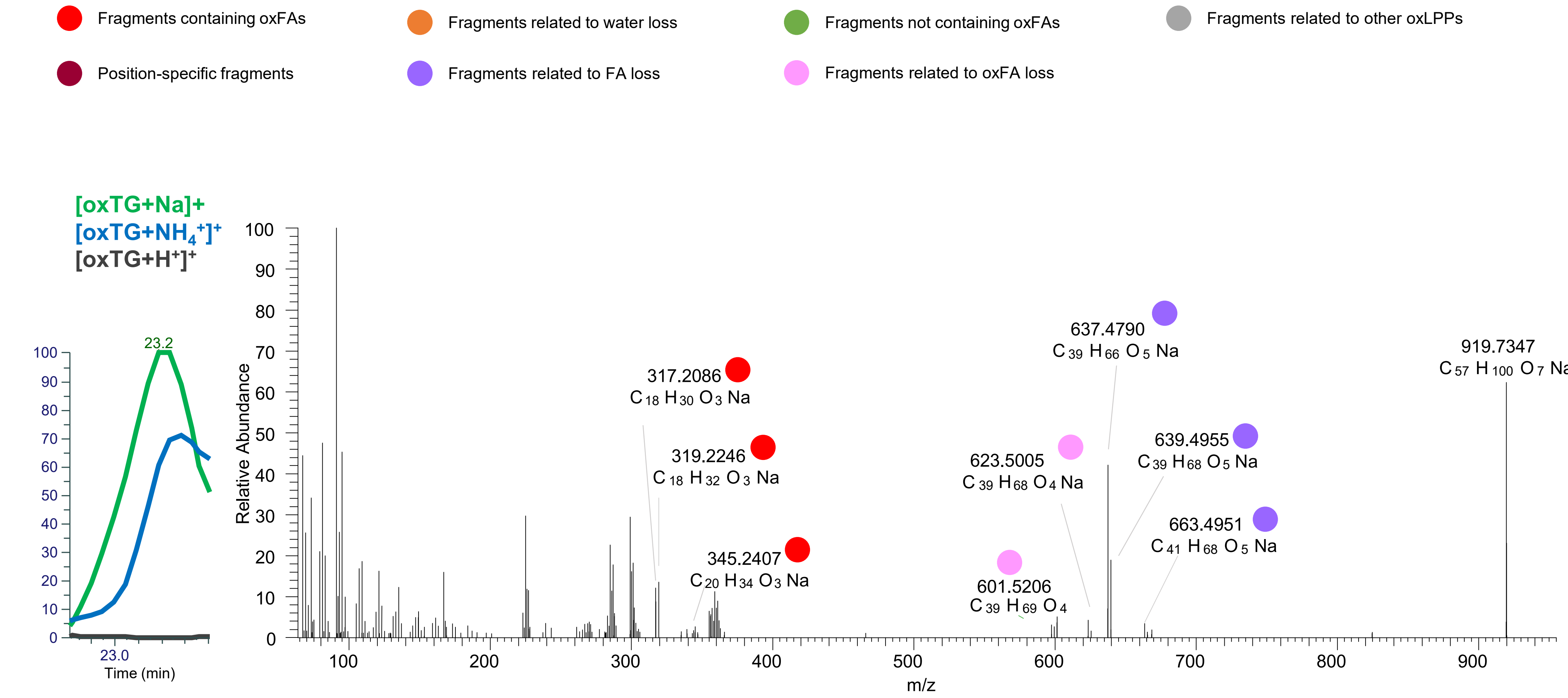

TG(18:1_18:1_18:3<OH>)
TG(18:1_18:2_18:2<O>)
TG(16:0_18:2_20:3<O>)
TG(16:0_18:1_20:4<O>)
RT 23.5

**[oxTG+Na]+** **XIC 919.7361 NL: 9.68E5**

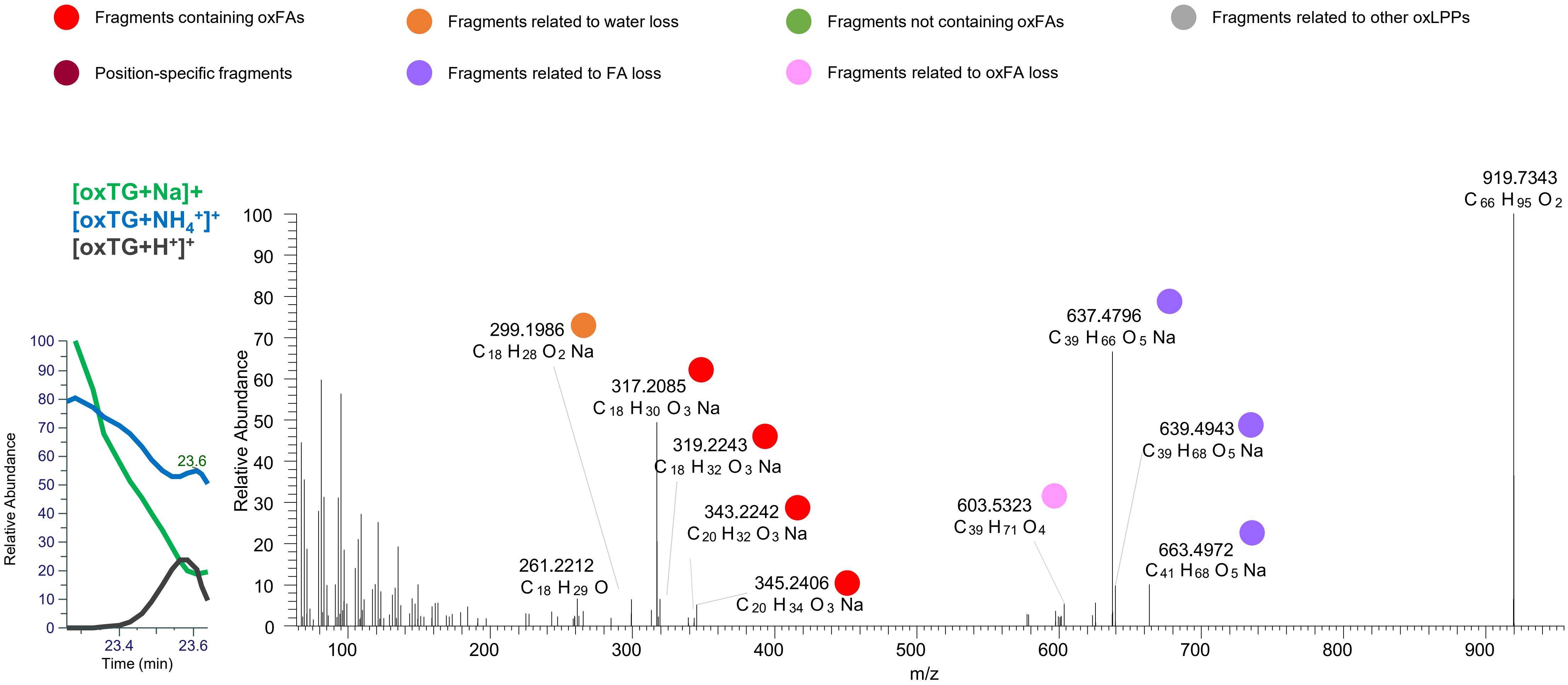

TG(18:1_18:1_18:3<OH>)
TG(16:0_18:2_20:2<oxo>)
TG(16:0_18:1_20:4<OH>)
RT 24.0

[oxTG+Na]+ XIC 919.7361 NL: 6.47E5

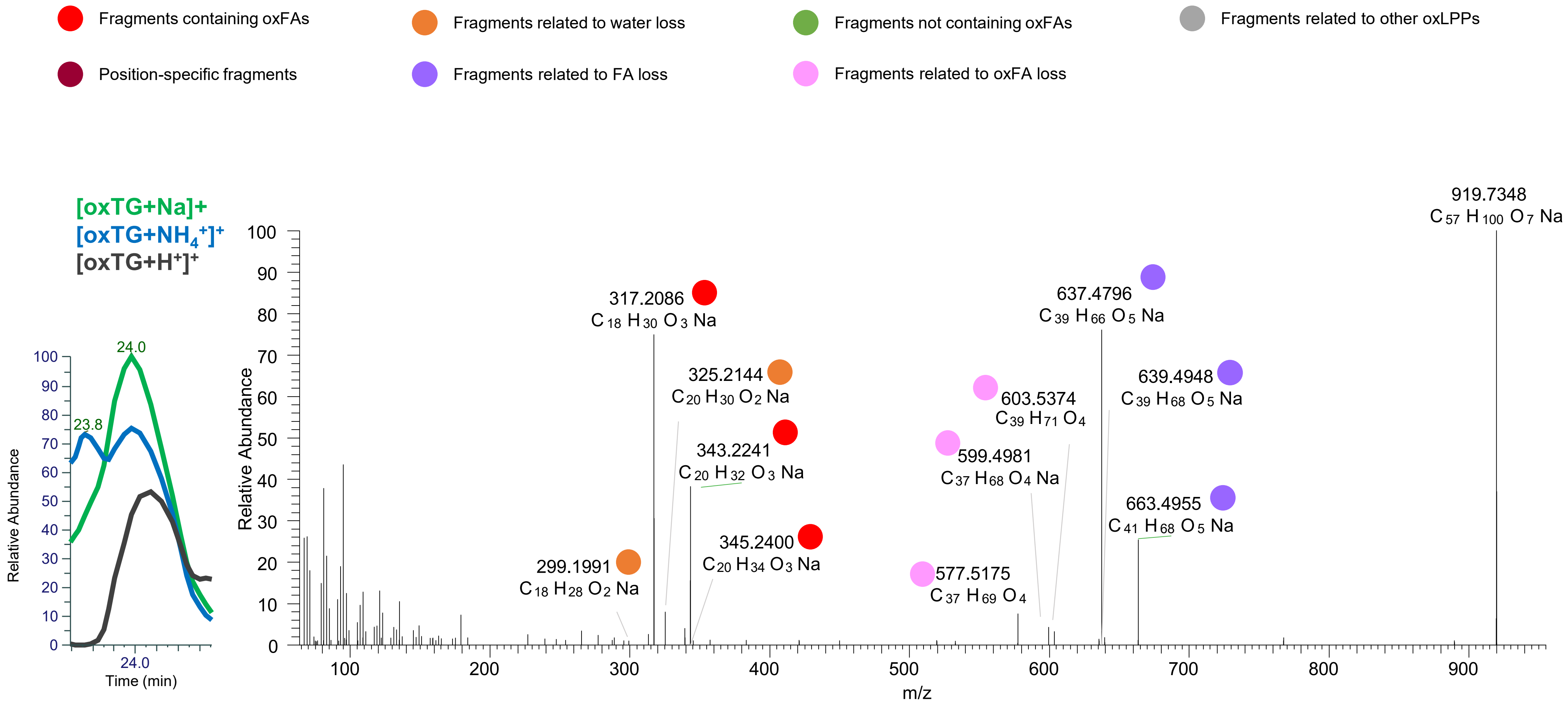

### TG(18:1_18:1_18:2<oxo>) RT 25.3 TG(18:1_18:1_18:2<ep>) RT 25.4

**[oxTG+Na]+** **XIC 919.7361 NL: 2.20E5**

- Fragments containing oxFAs
- Fragments related to water loss
- Fragments not containing oxFAs
- Fragments related to other oxLPPs
- Position-specific fragments
- Fragments related to FA loss
- Fragments related to oxFA loss

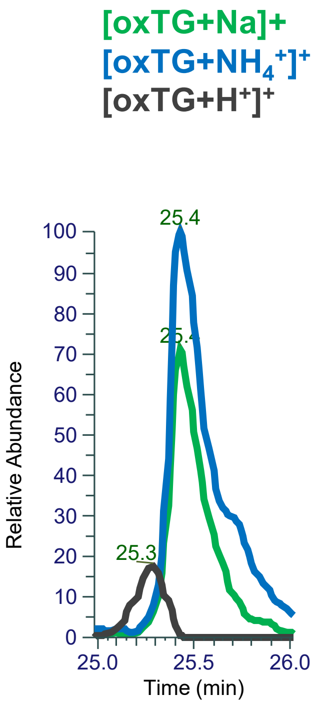


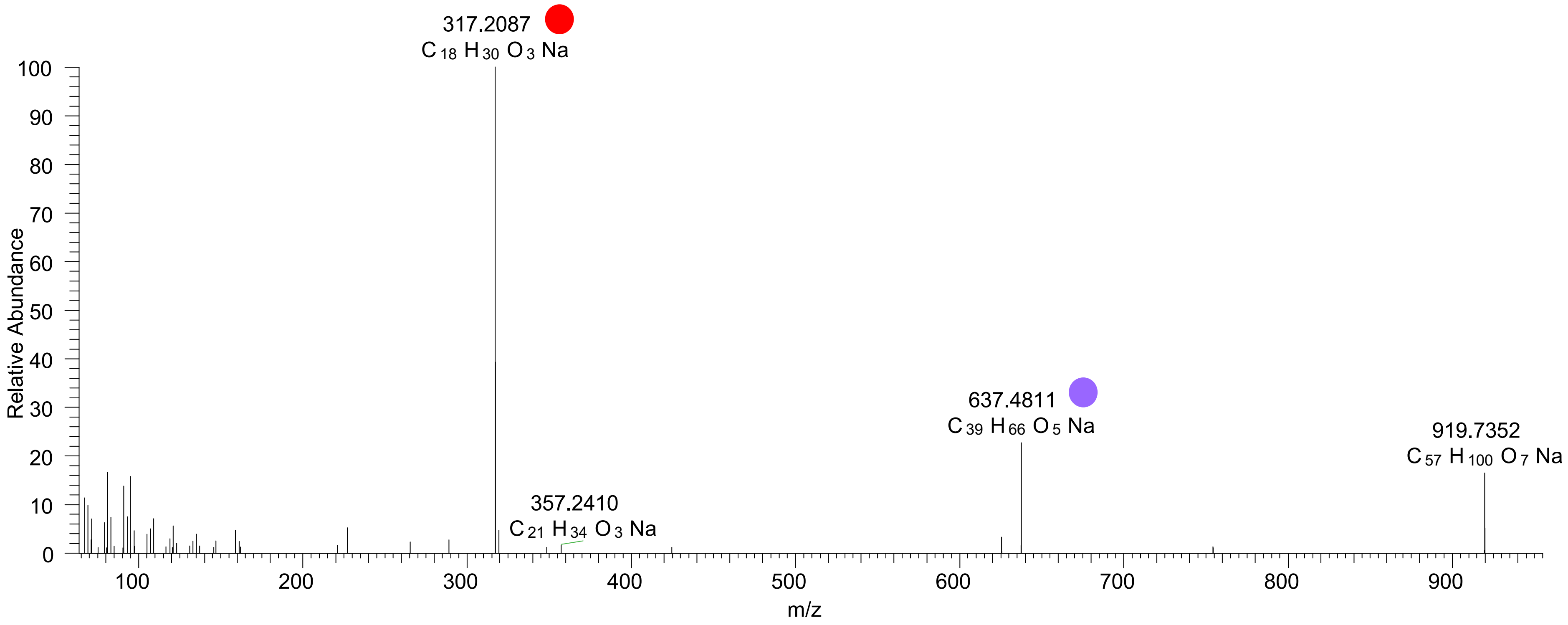

# TG(18:0_18:2_18:2<OH>)
# TG(18:0_18:1_18:2<ep>)
# RT 23.4

**[oxTG+Na]+** **XIC 921.7517 NL: 3.21E5**

- Fragments containing oxFAs
- Fragments related to water loss
- Fragments not containing oxFAs
- Fragments related to other oxLPPs
- Position-specific fragments
- Fragments related to FA loss
- Fragments related to oxFA loss

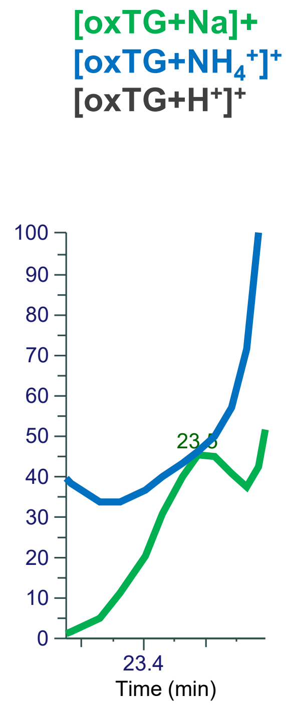


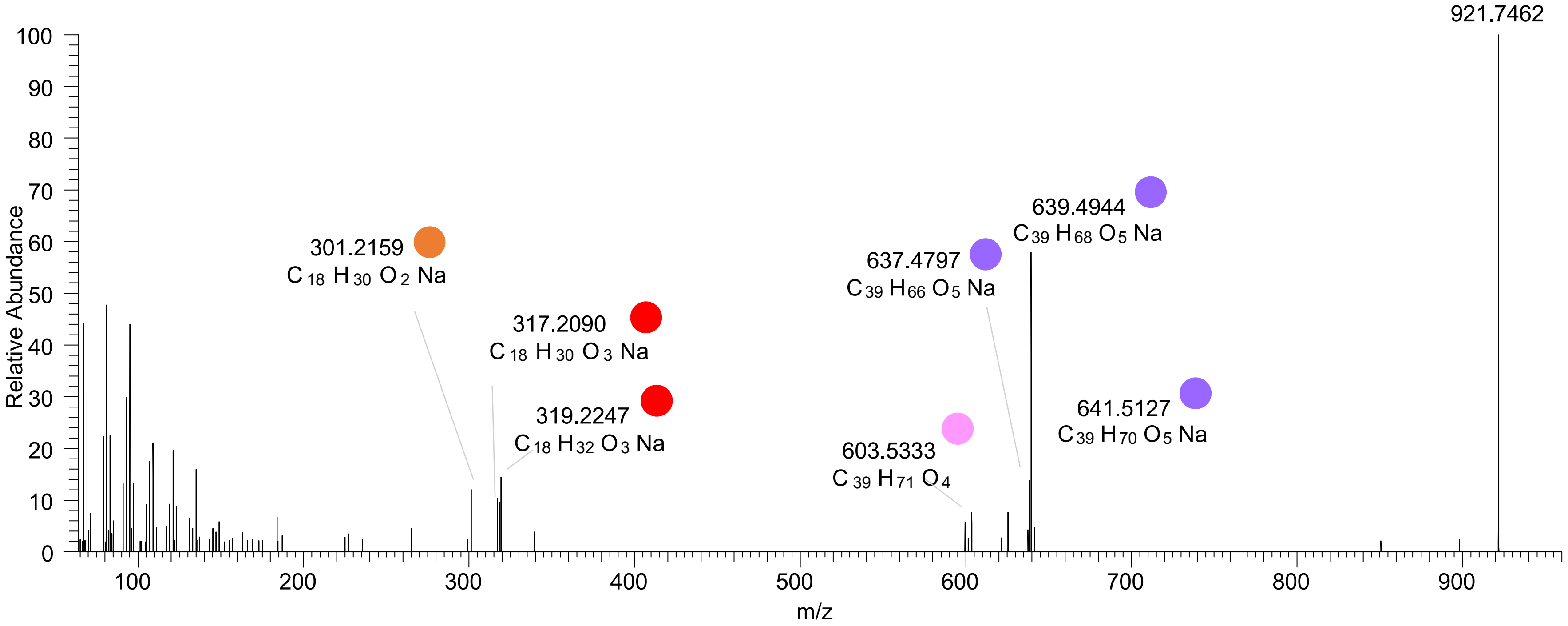

# TG(18:1_18:1_18:2<OH>)
# RT 23.7

**[oxTG+Na]+** **XIC 921.7517 NL: 3.21E5**

- Fragments containing oxFAs
- Fragments related to water loss
- Fragments not containing oxFAs
- Fragments related to other oxLPPs
- Position-specific fragments
- Fragments related to FA loss
- Fragments related to oxFA loss

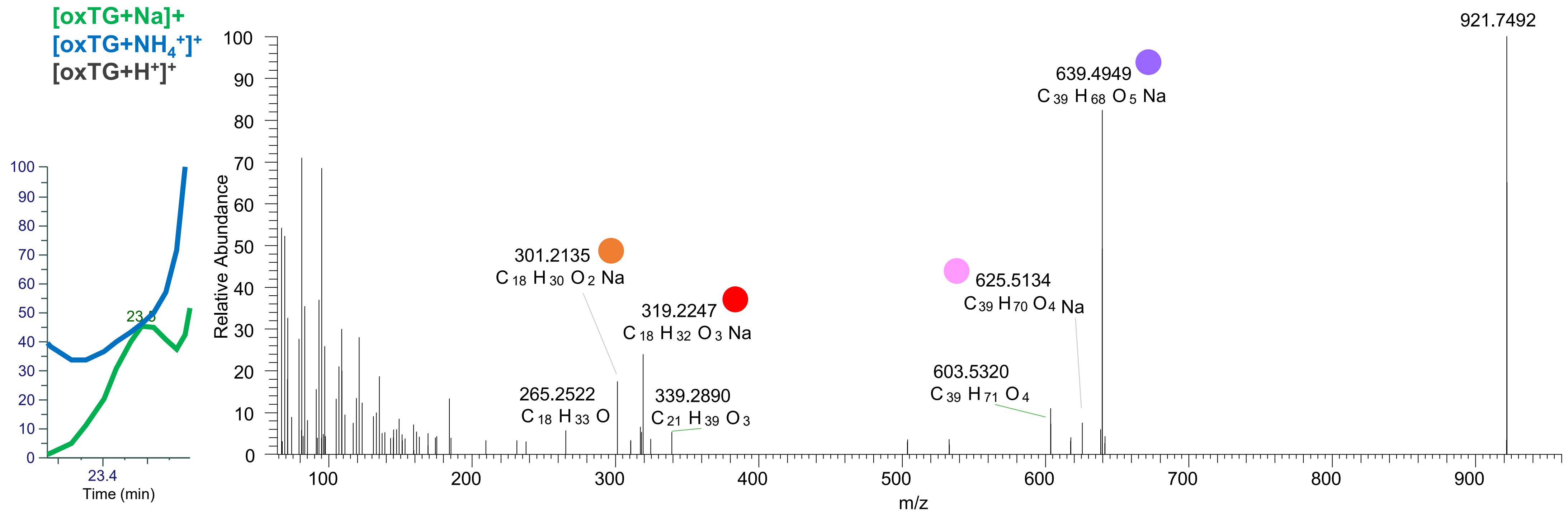

TG(18:1_18:1_18:2<OH>)
TG(16:0_18:1_20:3<OH>)
23.9

[oxTG+Na]+ XIC 921.7517 NL: 1.80E6

- Fragments containing oxFAs
- Fragments related to water loss
- Fragments not containing oxFAs
- Fragments related to other oxLPPs
- Position-specific fragments
- Fragments related to FA loss
- Fragments related to oxFA loss

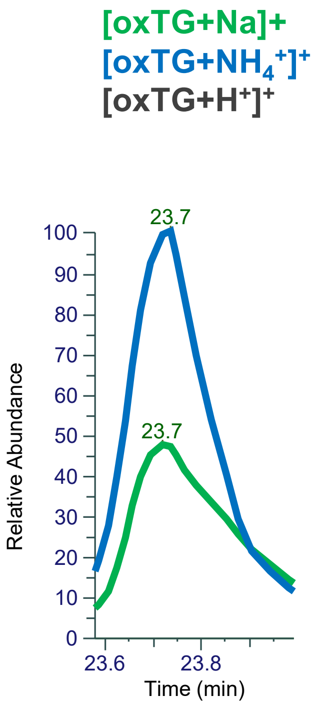


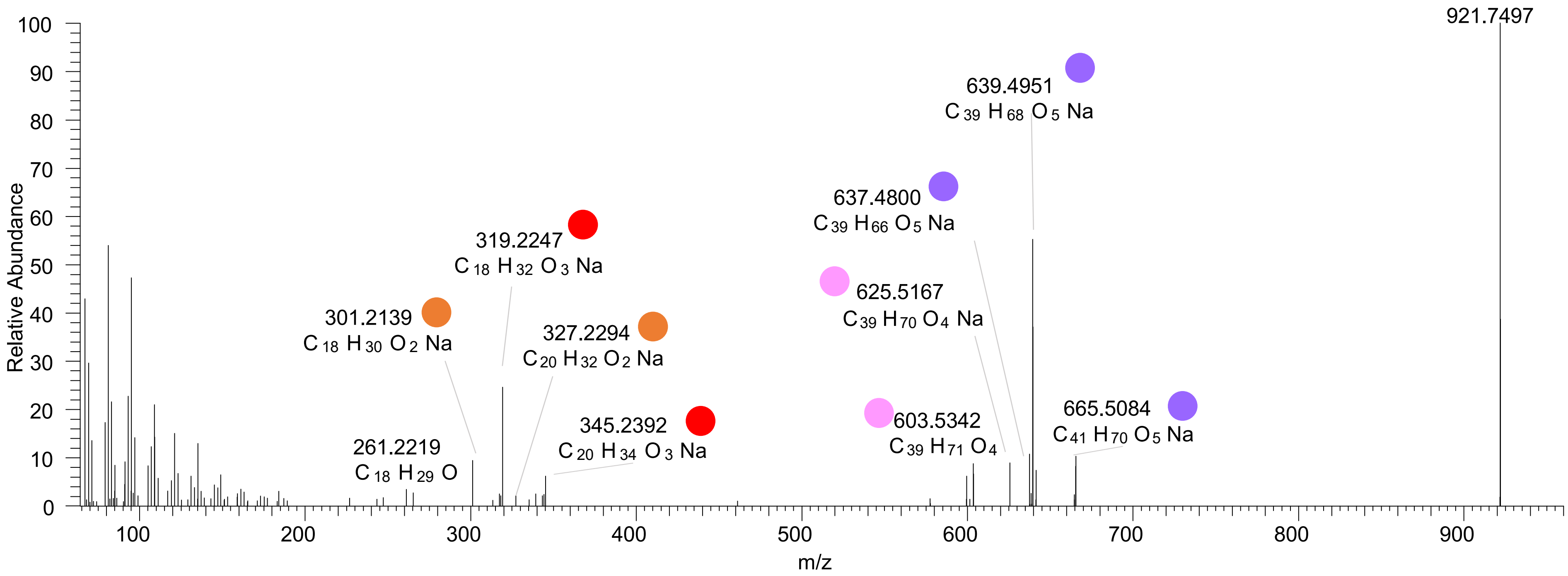

# TG(18:0_18:1_18:3<O>)
# RT 24.9

**[oxTG+Na]+** **XIC 921.7517 NL: 1.01E5**

- Fragments containing oxFAs
- Fragments related to water loss
- Fragments not containing oxFAs
- Fragments related to other oxLPPs
- Position-specific fragments
- Fragments related to FA loss
- Fragments related to oxFA loss

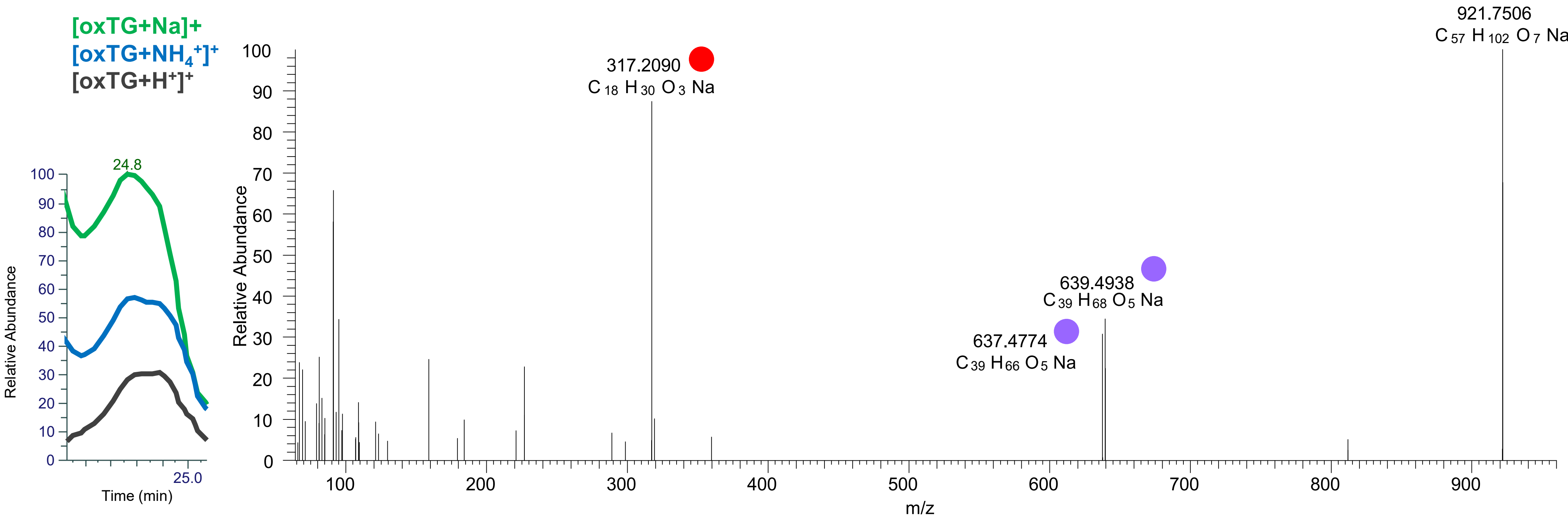

# TG(16:0_18:1_18:3<3O>)
# RT 22.0

**[oxTG+Na]+** **XIC 925.7103 NL: 3.65E4**

Fragments containing oxFAs
Fragments related to water loss
Fragments not containing oxFAs
Fragments related to other oxLPPs
Position-specific fragments
Fragments related to FA loss
Fragments related to oxFA loss

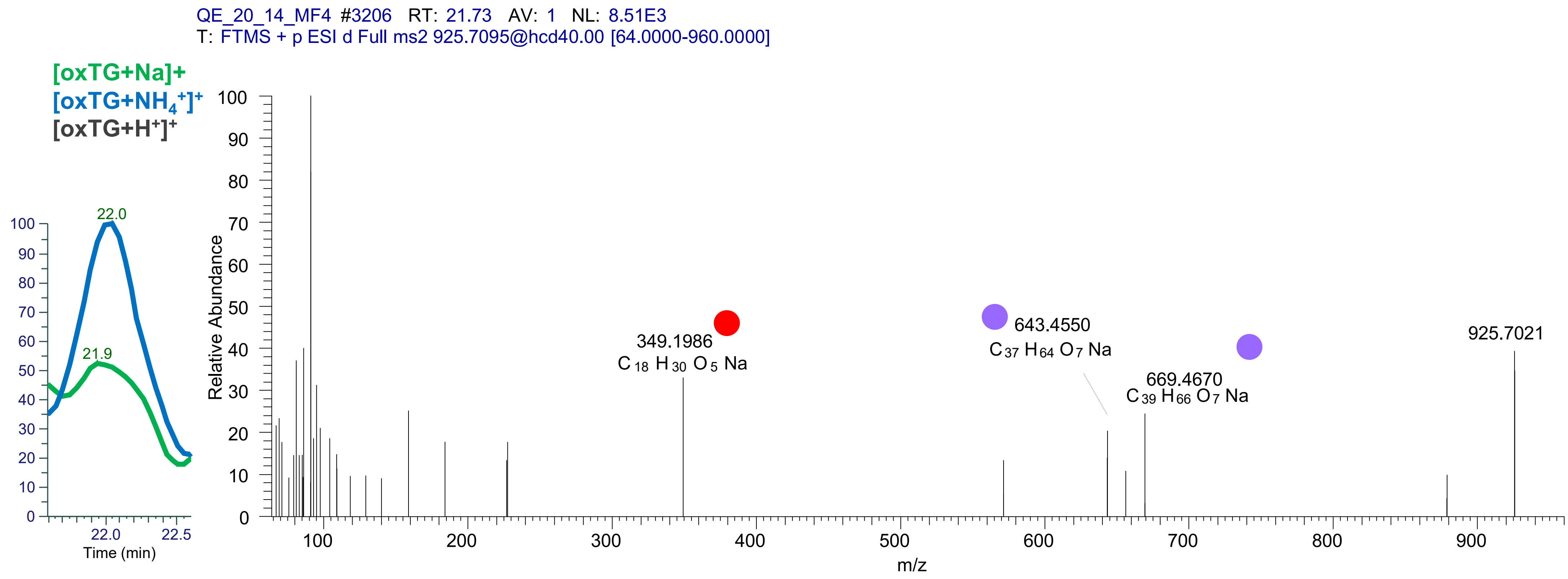

TG(16:1_18:1_20:4<2O>)
TG(16:0_18:2_20:4<2O>)
TG(18:1_18:2_18:3<2O>)
RT 21.1

[oxTG+Na]+ XIC 933.7154 NL: 9.25E4

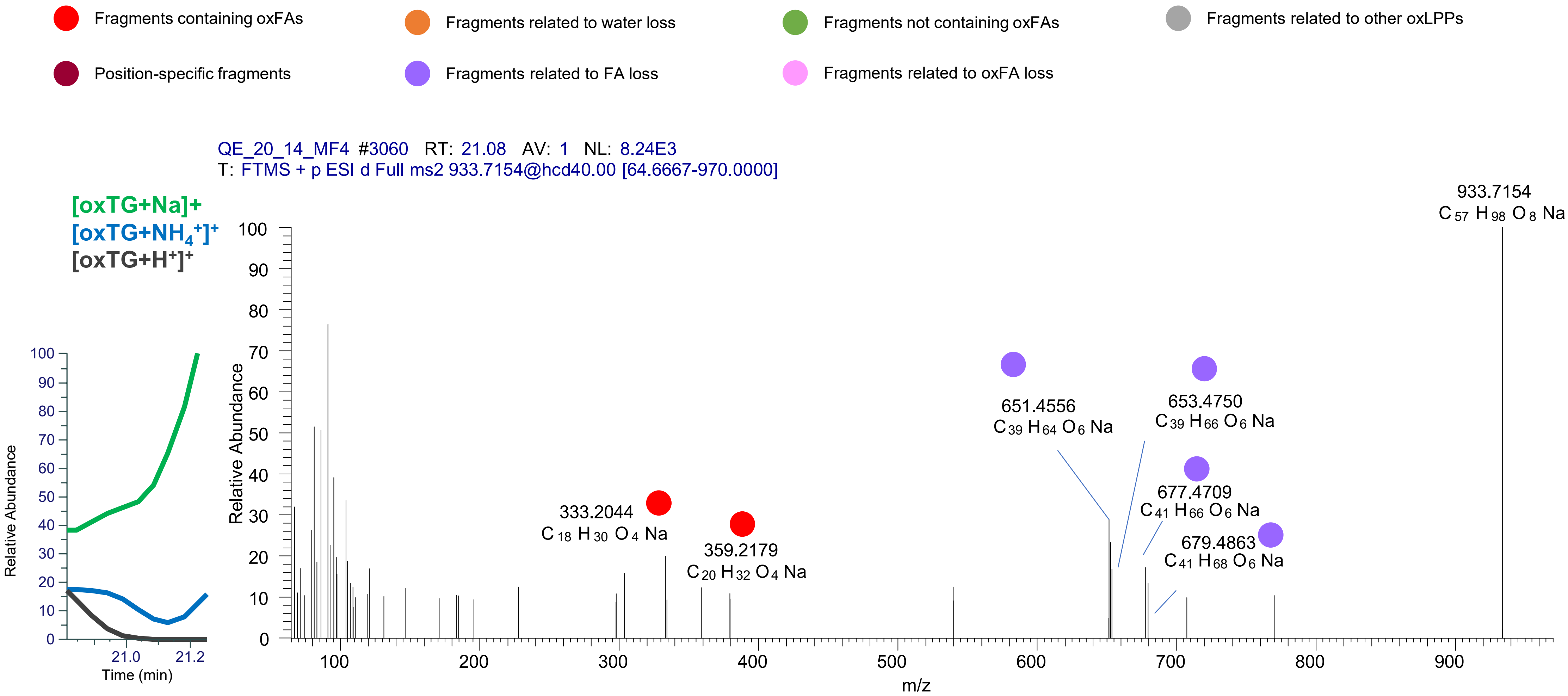

TG(16:0_18:2_20:4<O,OH>)
TG(18:1_18:1_18:4<2O>)
TG(16:0_16:1_22:5<2O>)
RT 21.5

[oxTG+Na]+ XIC 933.7154 NL: 1.40E5

- Fragments containing oxFAs
- Fragments related to water loss
- Fragments not containing oxFAs
- Fragments related to other oxLPPs
- Position-specific fragments
- Fragments related to FA loss
- Fragments related to oxFA loss

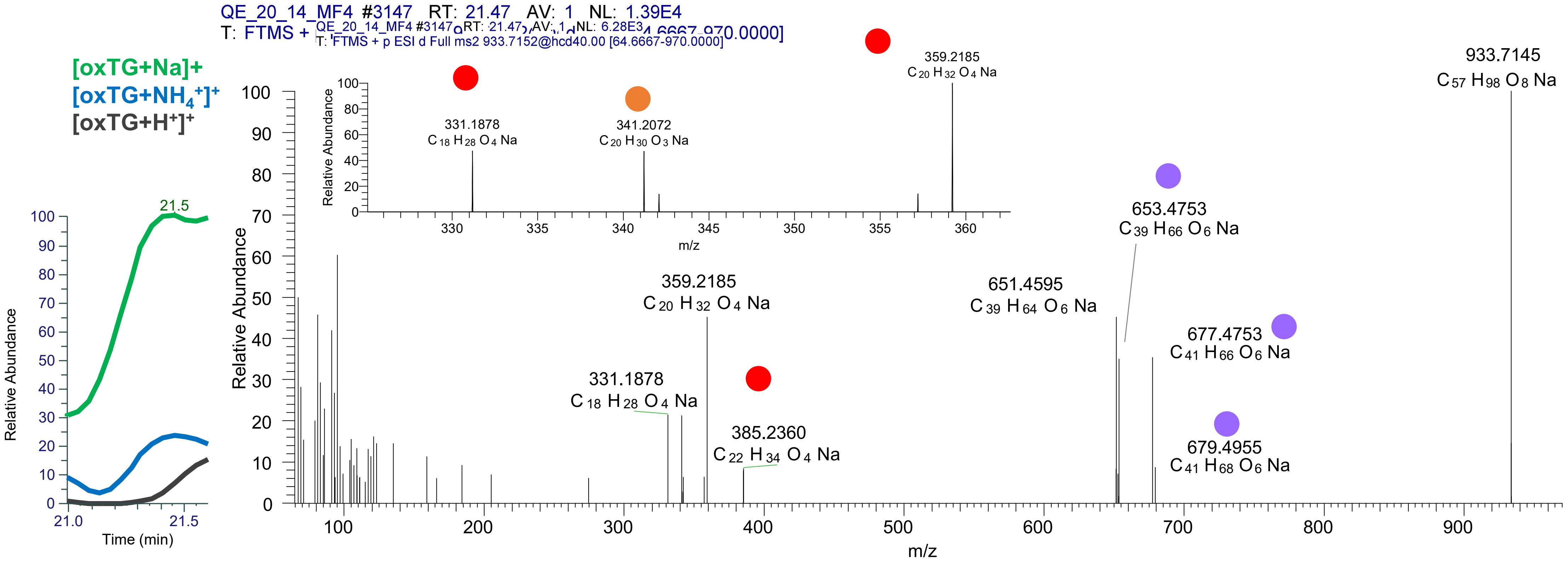

# TG(16:0_18:1_20:5<OH,O>)
# TG(16:0_18:2_20:4<OH,O>)
### TG(18:1_18:1_18:4<2O>) - ISF
# RT 21.8

[oxTG+Na]+ XIC 933.7154 NL: 1.57E5

Fragments containing oxFAs
Fragments related to water loss
Fragments not containing oxFAs
Fragments related to other oxLPPs
Position-specific fragments
Fragments related to FA loss
Fragments related to oxFA loss

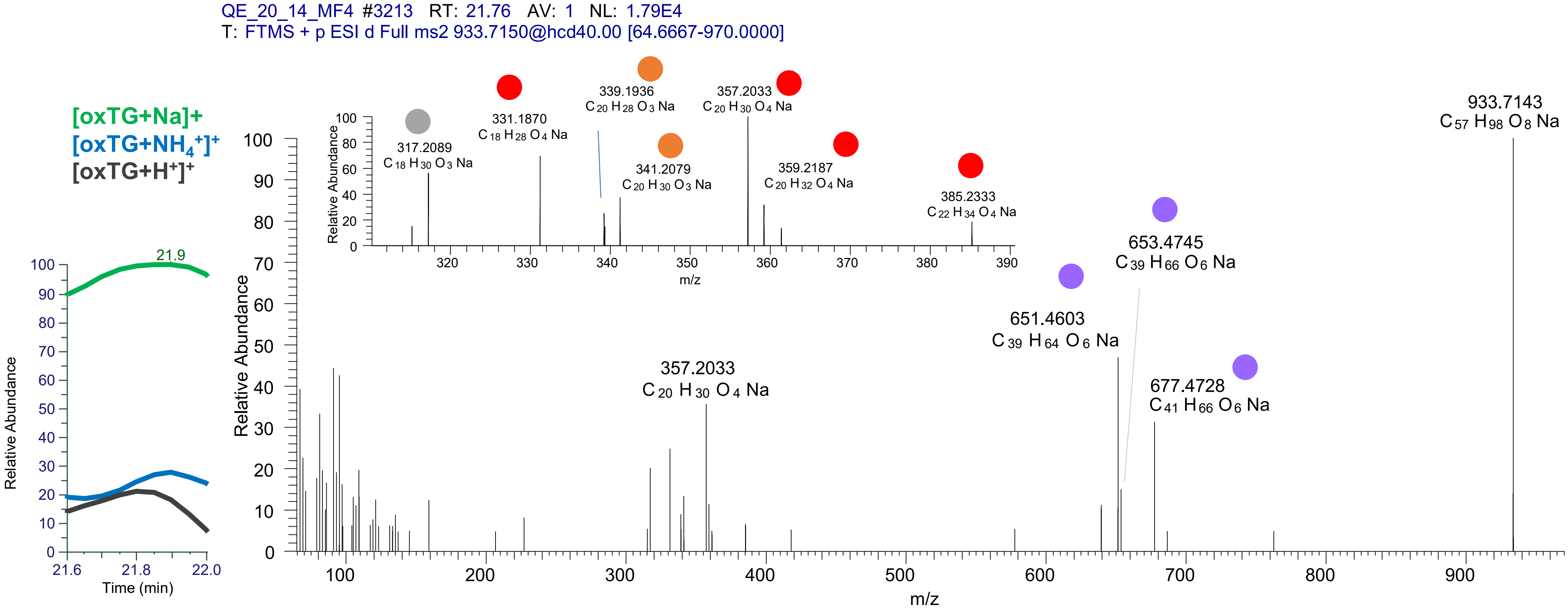

TG(18:1_18:2_18:3<OOH>)
TG(16:0_18:2_20:4<2O>)
TG(16:0_18:1_20:5<OOH>)
RT 22.3

[oxTG+Na]+ XIC 933.7154 NL: 2.25E5

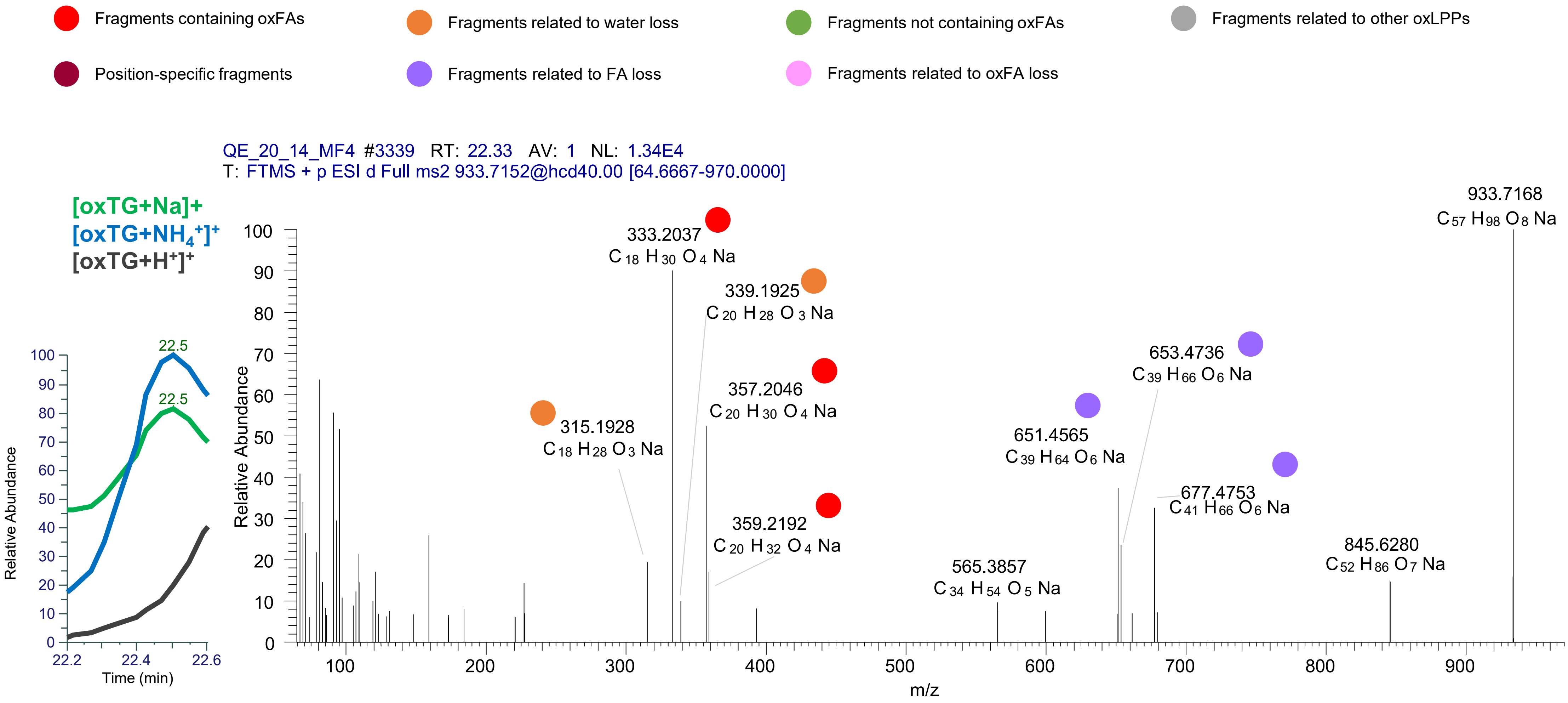

TG(18:1_18:1_18:4<2O>)
TG(18:1_18:2_18:3<2O>)
TG(18:2_18:2_18:2<2O>)
TG(16:0_18:1_20:5<2O>)
RT 22.5-22.7

[oxTG+Na]+ XIC 933.7154 NL: 2.25E5

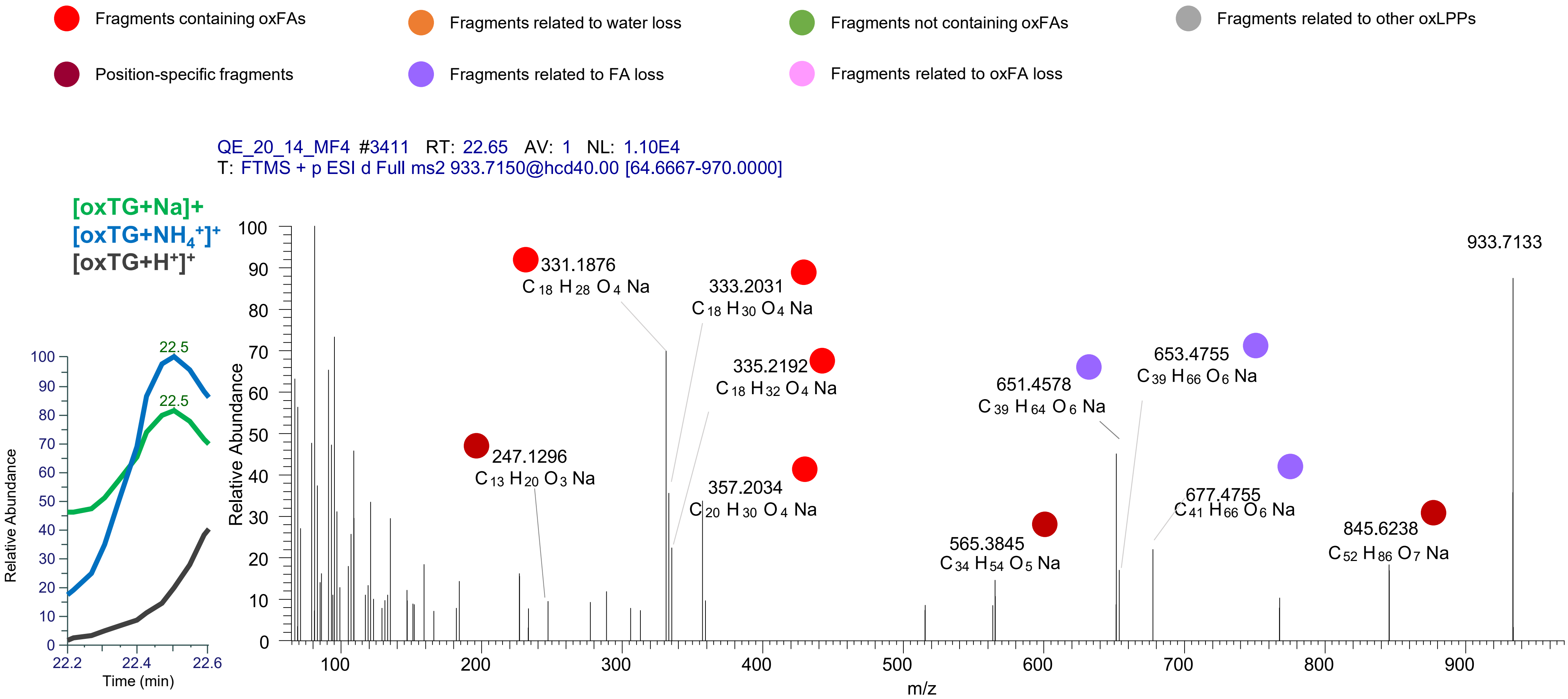

# TG(18:1_18:2<OH>_18:2<OH>)
# RT 20.2

**[oxTG+Na]+** **XIC 935.7310 NL: 3.13E5**

- Fragments containing oxFAs
- Fragments related to water loss
- Fragments not containing oxFAs
- Fragments related to other oxLPPs
- Position-specific fragments
- Fragments related to FA loss
- Fragments related to oxFA loss

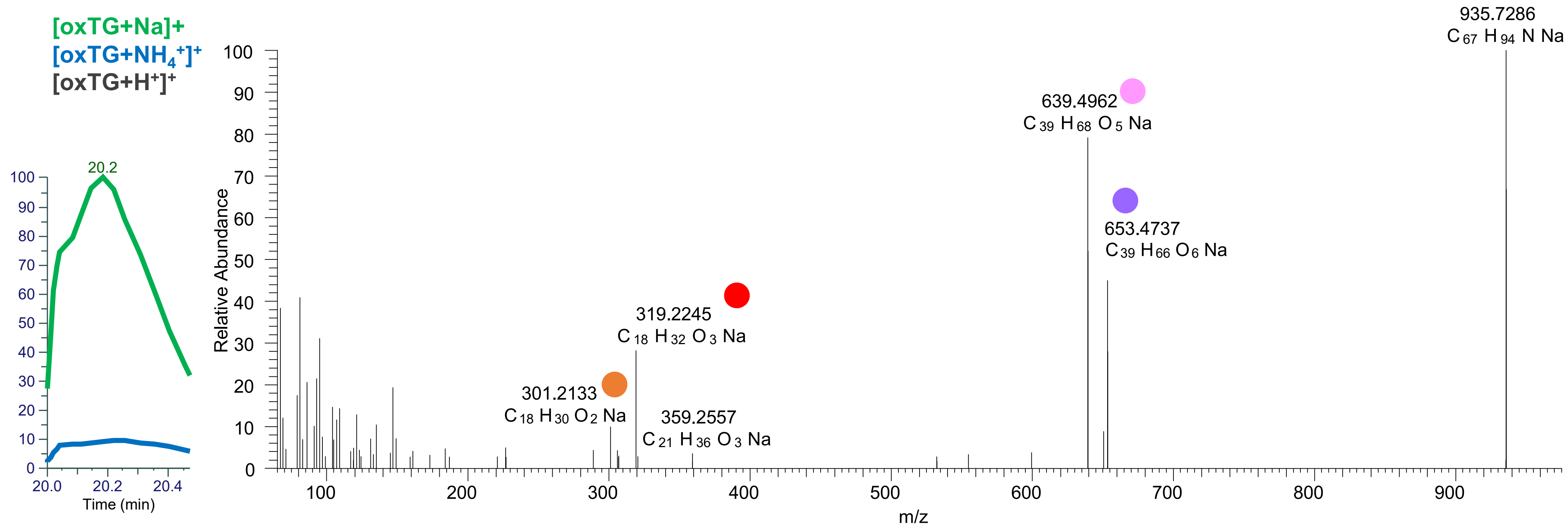

# TG(18:1_18:2_18:2<2O>)
# TG(18:1_18:1_18:3<OH,O>)
# RT 22.1

**[oxTG+Na]+** **XIC 935.7310 NL: 2.00E5**

- Fragments containing oxFAs
- Fragments related to water loss
- Fragments not containing oxFAs
- Fragments related to other oxLPPs
- Position-specific fragments
- Fragments related to FA loss
- Fragments related to oxFA loss

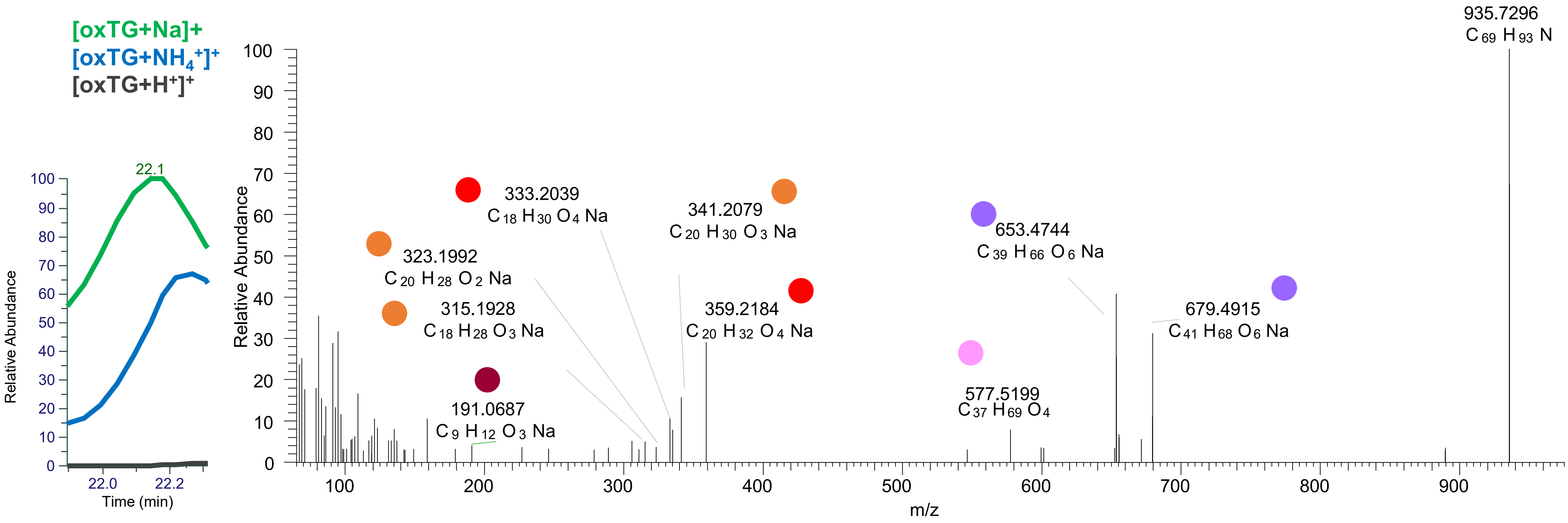

# TG(18:1_18:1_18:3<OH,O>)
# TG(16:0_18:1_20:4<OH,O>)
# RT 22.8

**[oxTG+Na]+** **XIC 935.7310 NL: 8.64E5**

- Fragments containing oxFAs
- Fragments related to water loss
- Fragments not containing oxFAs
- Fragments related to other oxLPPs
- Position-specific fragments
- Fragments related to FA loss
- Fragments related to oxFA loss

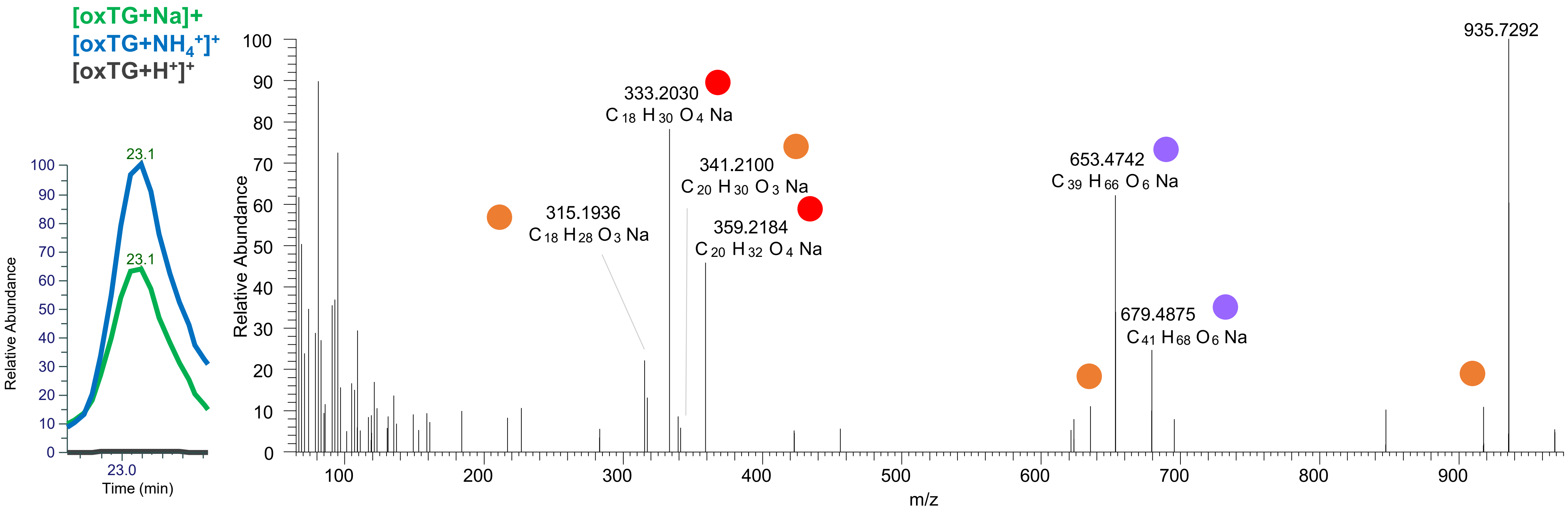

TG(18:1_18:1_18:3<OOH{13}>)
TG(18:1_18:2_18:2<OOH{13}>)
TG(16:0_18:1_20:4<OH,O>)
RT 23.1

[oxTG+Na]+ XIC 935.7310 NL: 8.64E5

- Fragments containing oxFAs
- Fragments related to water loss
- Fragments not containing oxFAs
- Fragments related to other oxLPPs
- Position-specific fragments
- Fragments related to FA loss
- Fragments related to oxFA loss

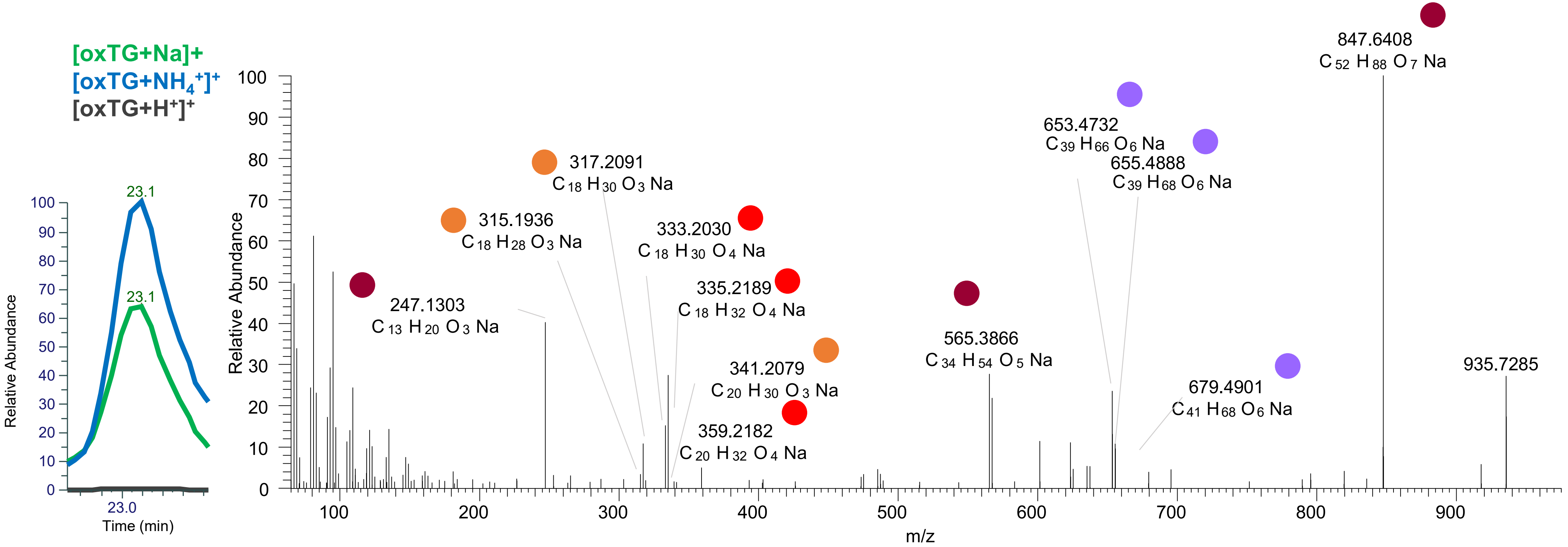

# TG(18:0_18:2<O>_18:2<O>)
# RT 21.0

**[oxTG+Na]+** **XIC 937.7467 NL: 7.18E4**

Fragments containing oxFAs
Fragments related to water loss
Fragments not containing oxFAs
Fragments related to other oxLPPs
Position-specific fragments
Fragments related to FA loss
Fragments related to oxFA loss

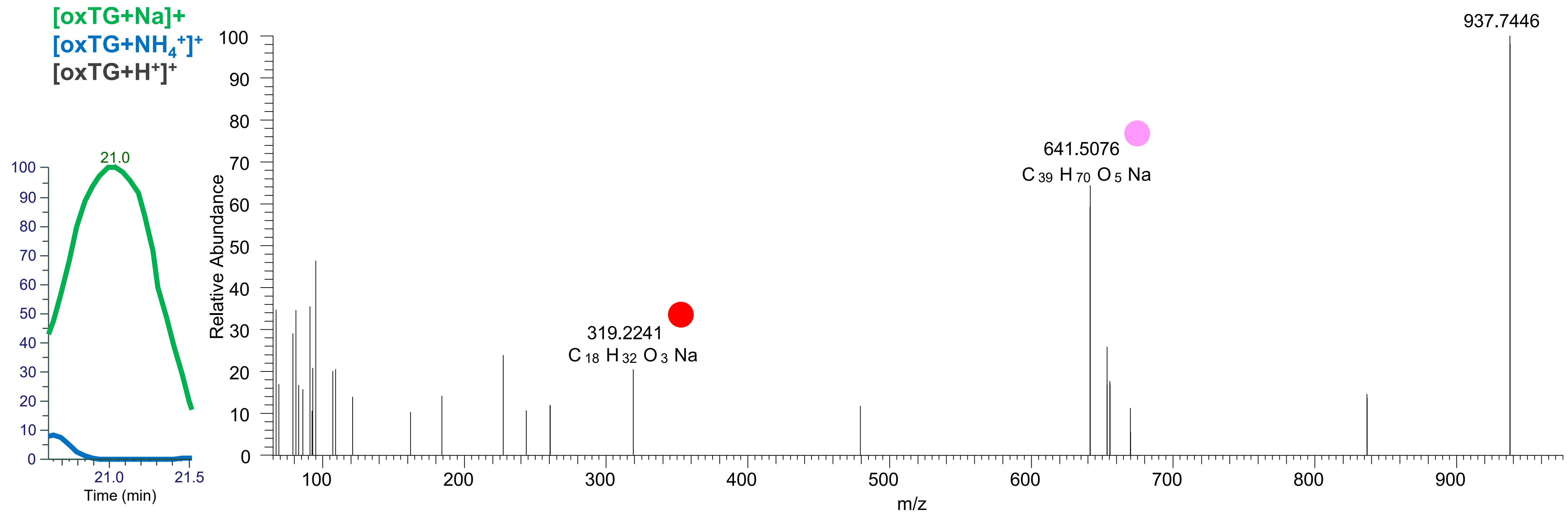

# TG(16:0_18:1_20:3<2O>)
# TG(18:1_18:1_18:2<OH,O>)
# RT 22.8

**[oxTG+Na]+** **XIC 937.7467 NL: 8.25E4**

- Fragments containing oxFAs
- Fragments related to water loss
- Fragments not containing oxFAs
- Fragments related to other oxLPPs
- Position-specific fragments
- Fragments related to FA loss
- Fragments related to oxFA loss

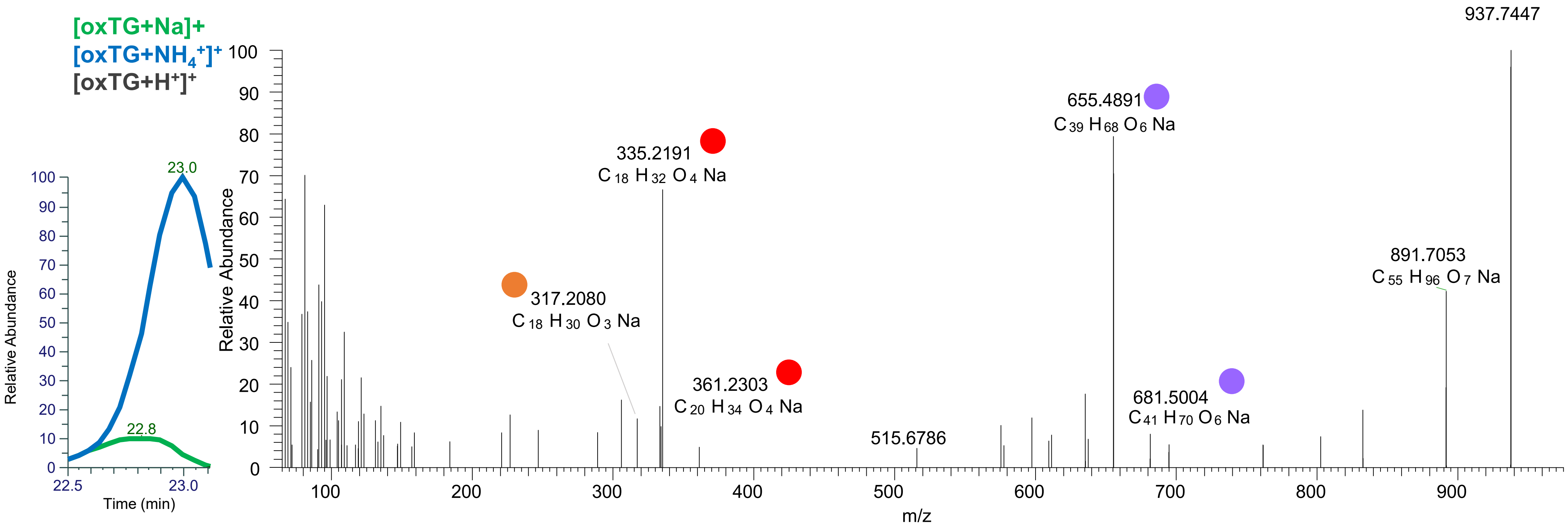

### TG(18:1_18:1_18:2<OOH{13}>)
# RT 23.6

**[oxTG+Na]+** **XIC 937.7467 NL: 2.48E6**

Fragments containing oxFAs
Fragments related to water loss
Fragments not containing oxFAs
Fragments related to other oxLPPs
Position-specific fragments
Fragments related to FA loss
Fragments related to oxFA loss

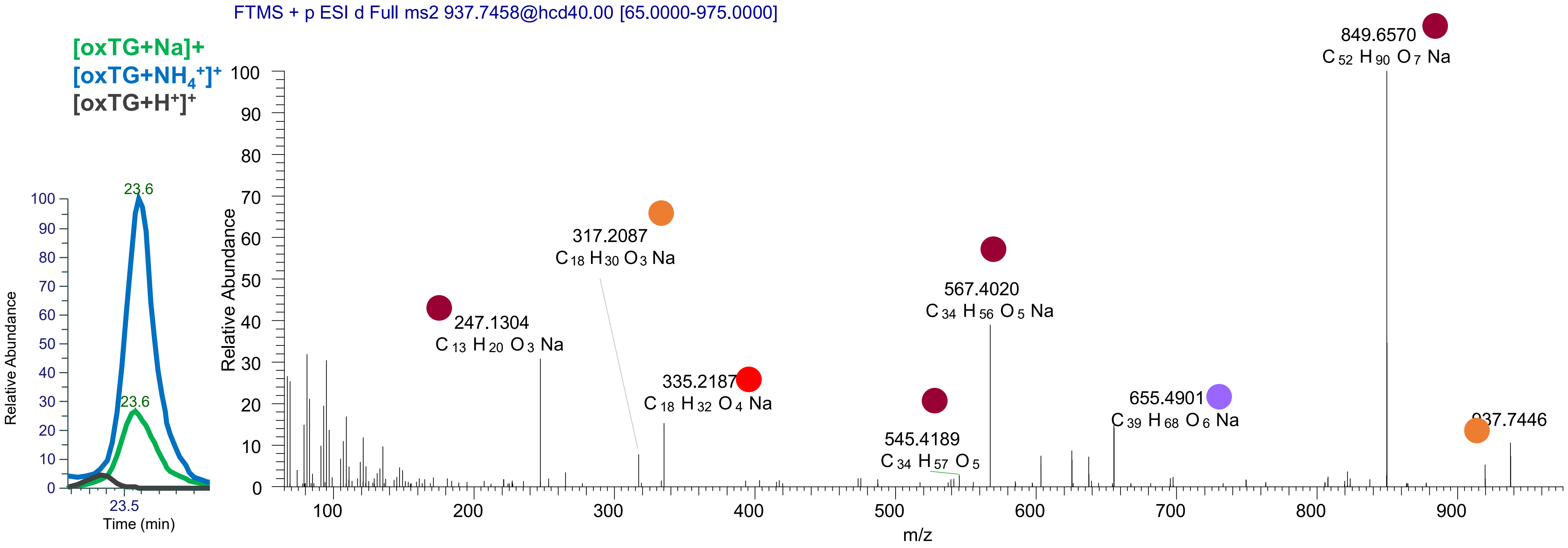

# TG(16:0_18:2_20:4<3O>)
# TG(16:0_18:1_20:5<3O>)
# TG(18:1_18:1_18:4<3O>)
# RT 21.3

**[oxTG+Na]+** **XIC 949.7103 NL: 4.15E4**

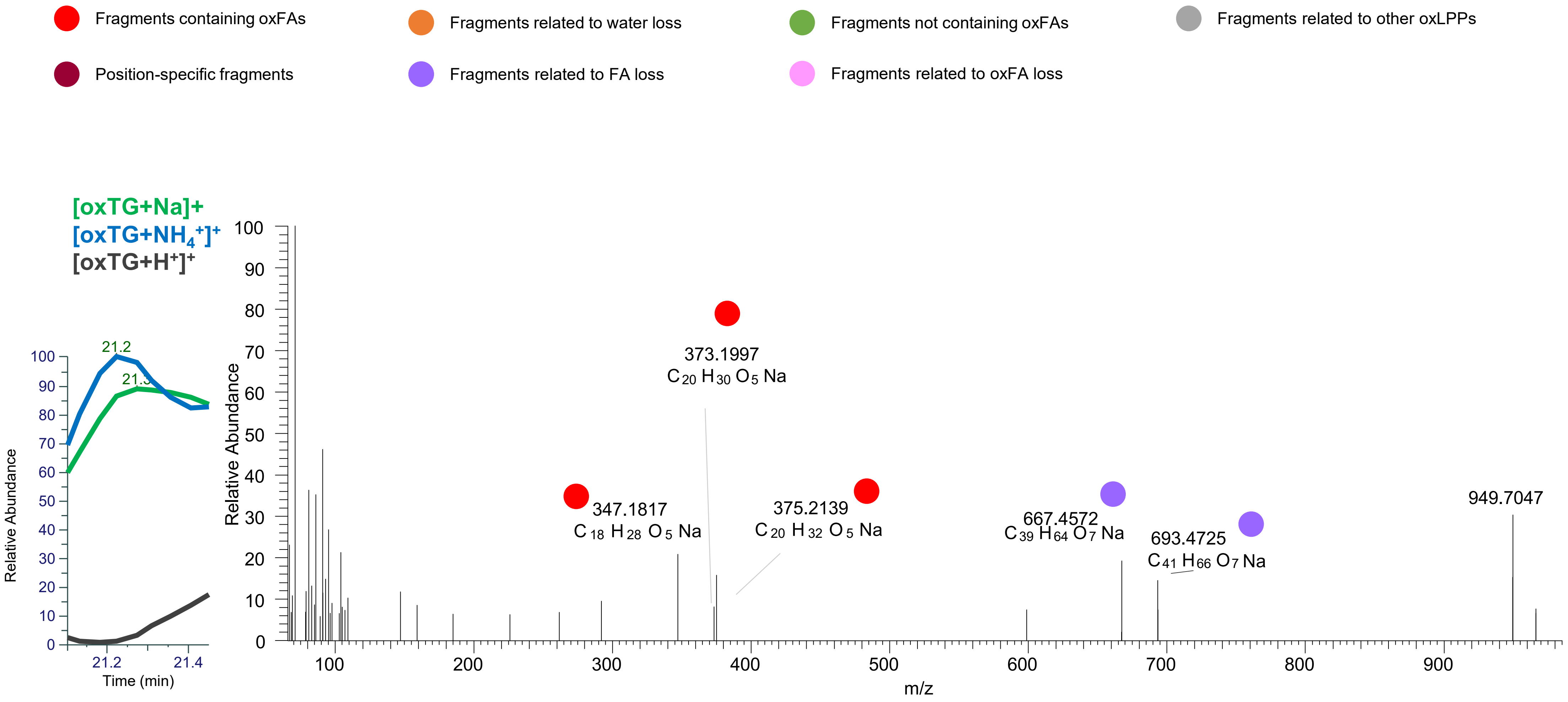

# TG(16:0_18:1_20:5<3O>)
# RT 21.8

**[oxTG+Na]+** **XIC 949.7103 NL: 4.45E4**

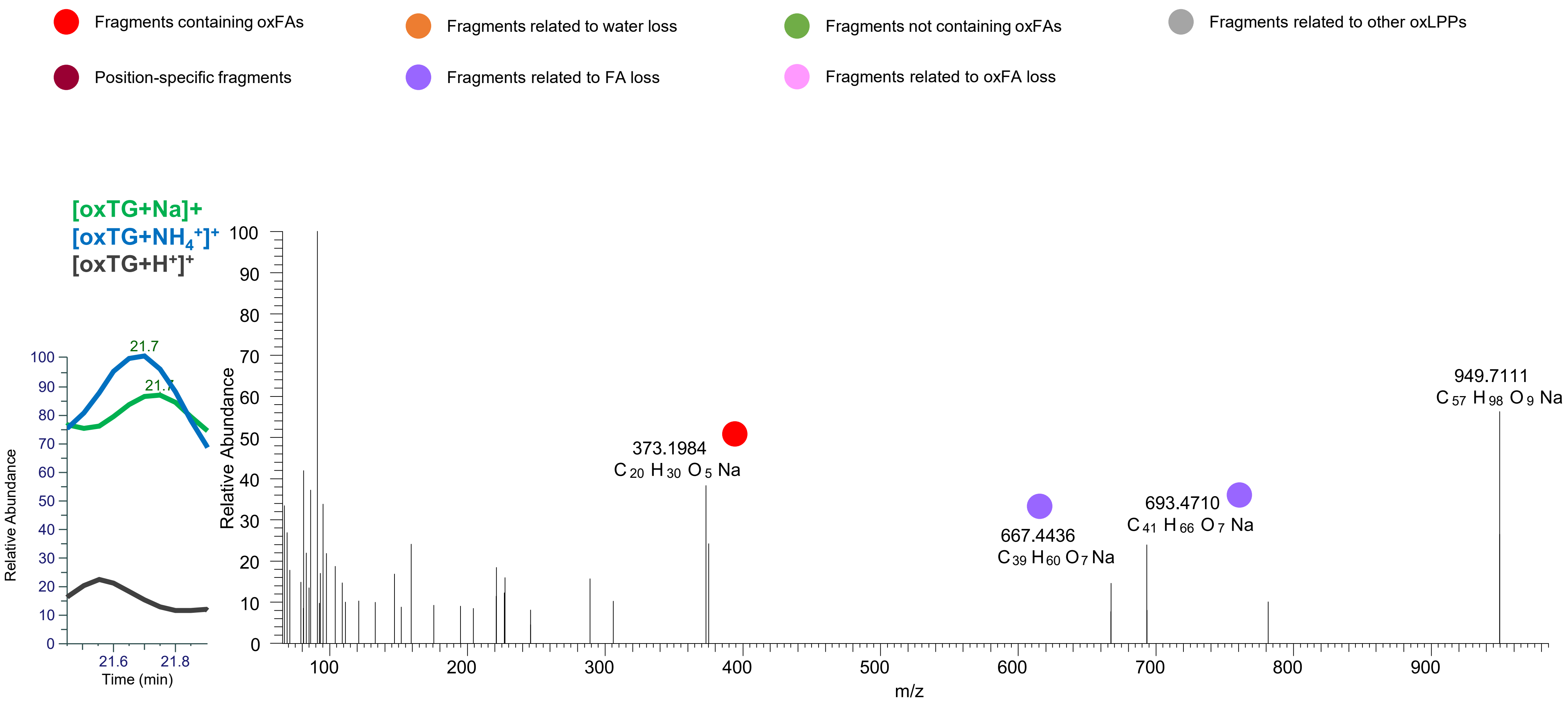

### TG(18:1_18:2<OH>_18:2<OOH{13}>)
# RT 20.1

**[oxTG+Na]+** **XIC 951.7259 NL: 3.66E5**

- Fragments containing oxFAs
- Fragments related to water loss
- Fragments not containing oxFAs
- Fragments related to other oxLPPs
- Position-specific fragments
- Fragments related to FA loss
- Fragments related to oxFA loss

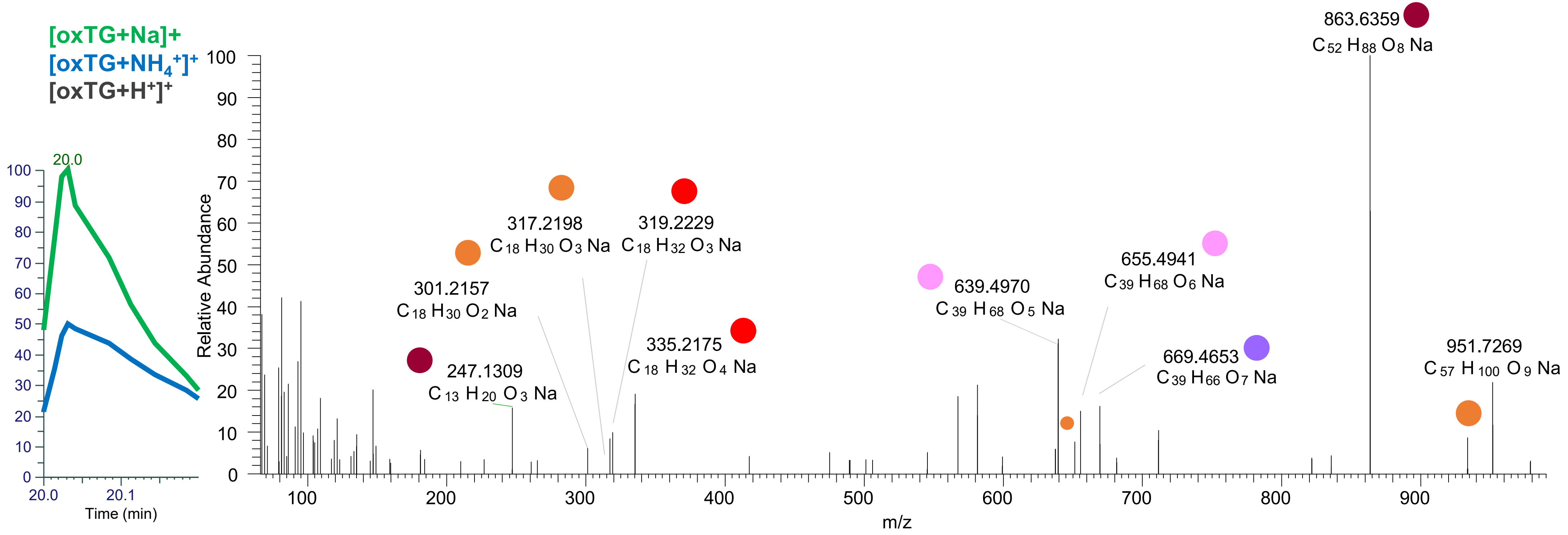

# TG(18:1_18:1_18:3<3O>)
# RT 21.1

[oxTG+Na]+ XIC 951.7259 NL: 3.39E4

- Fragments containing oxFAs
- Fragments related to water loss
- Fragments not containing oxFAs
- Fragments related to other oxLPPs
- Position-specific fragments
- Fragments related to FA loss
- Fragments related to oxFA loss

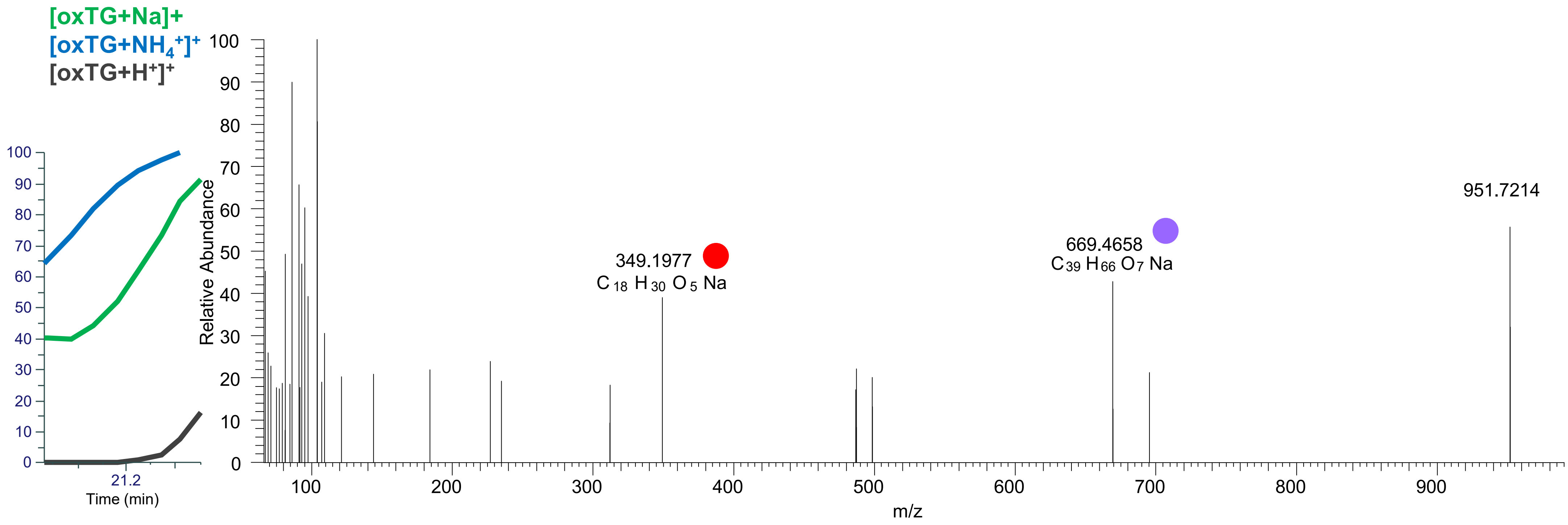

# TG(18:1_18:1_18:3<3O>)
# TG(16:0_18:1_20:4<OH,2O>)
# RT 21.7

**[oxTG+Na]+** **XIC 951.7259 NL: 4.45E4**

- Fragments containing oxFAs
- Fragments related to water loss
- Fragments not containing oxFAs
- Fragments related to other oxLPPs
- Position-specific fragments
- Fragments related to FA loss
- Fragments related to oxFA loss

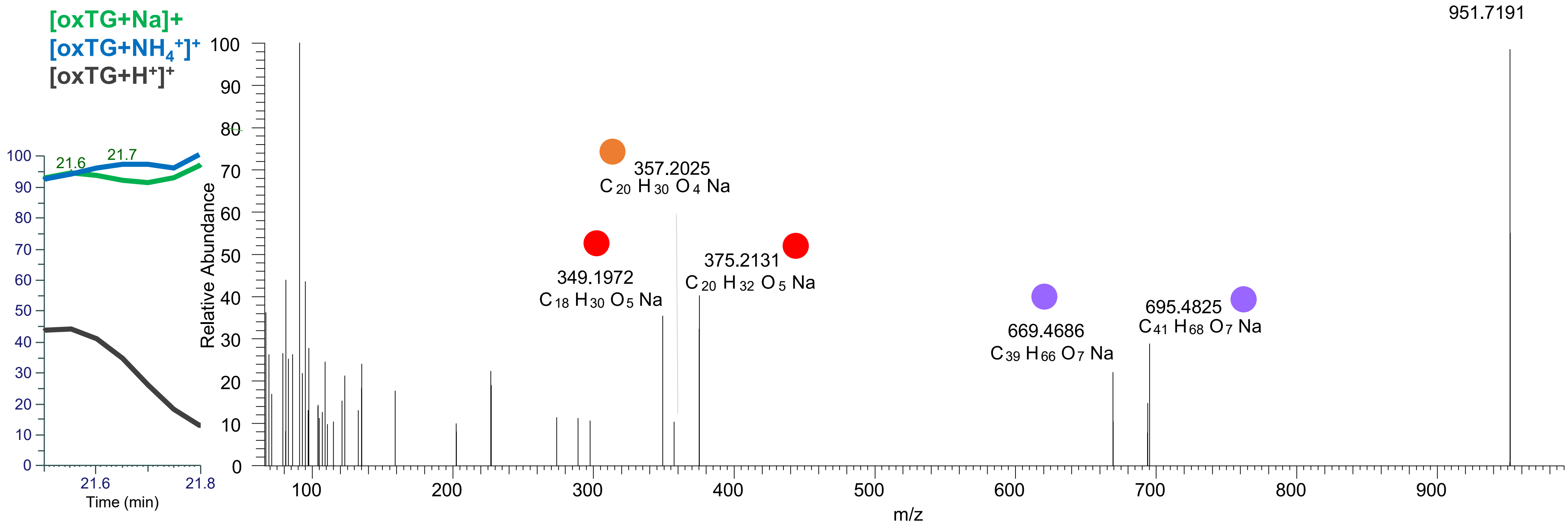

### TG(16:0_18:1_20:4<O,OOH{15}>)
# RT 22.0

**[oxTG+Na]+** **XIC 951.7259 NL: 7.27E4**

- Fragments containing oxFAs
- Fragments related to water loss
- Fragments not containing oxFAs
- Fragments related to other oxLPPs
- Position-specific fragments
- Fragments related to FA loss
- Fragments related to oxFA loss

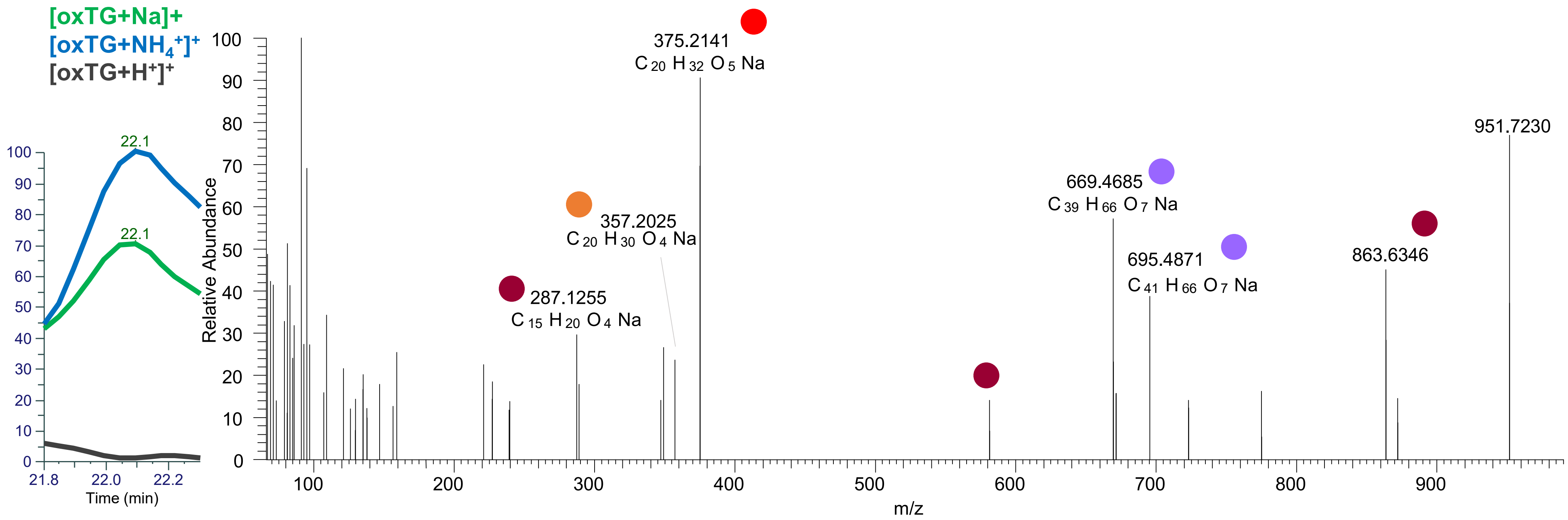

# TG(16:0_18:1_20:4<3O>)
# RT 22.2

**[oxTG+Na]+** **XIC 951.7259 NL: 7.27E4**

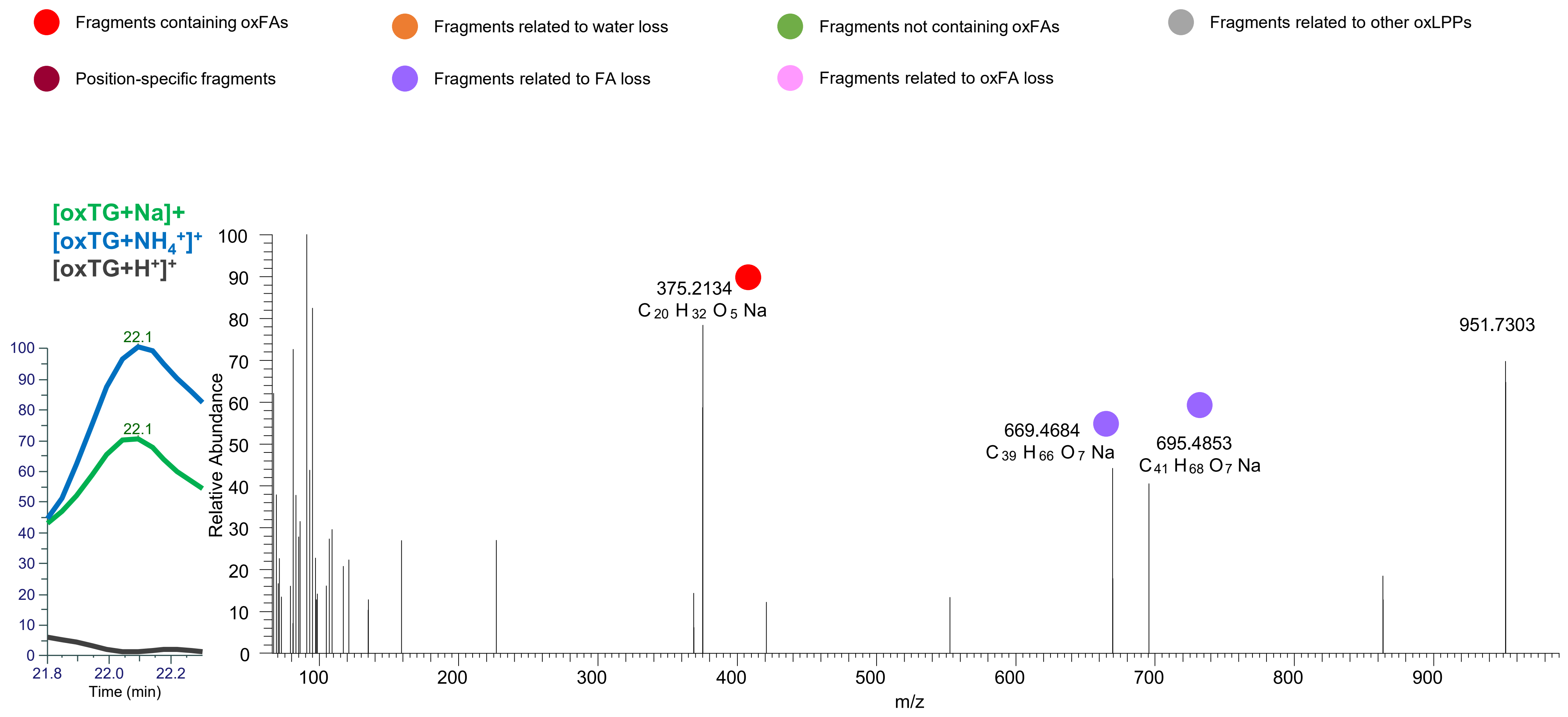

### TG(18:1_18:1_18:3<O,OOH{13}>)
# RT 22.8

[oxTG+Na]+ XIC 951.7259 NL: 3.52E4

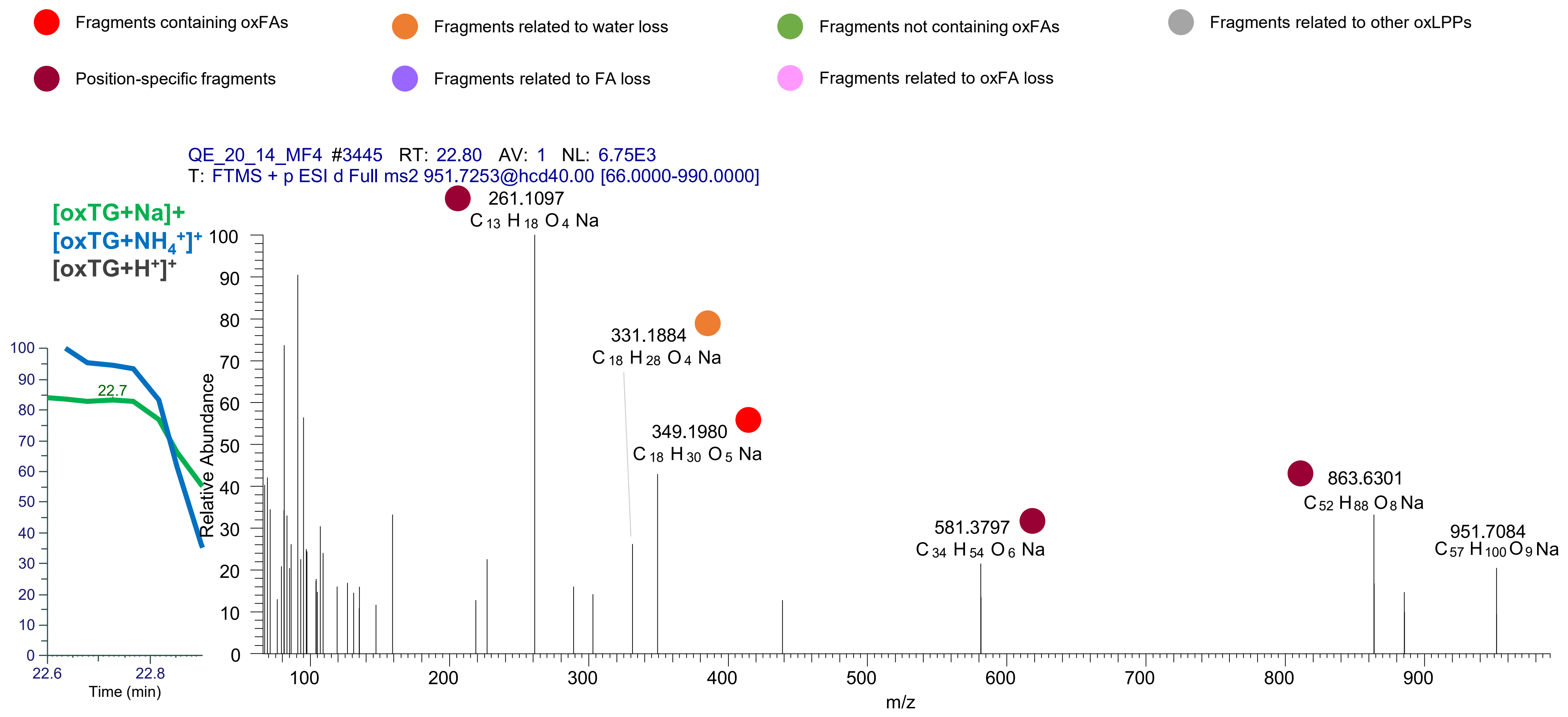
